# *GSDMA* drives the most replicated association with asthma in naïve CD4^+^ T cells

**DOI:** 10.1101/774760

**Authors:** Anne-Marie Madore, Lucile Pain, Anne-Marie Boucher-Lafleur, Jolyane Meloche, Andréanne Morin, Marie-Michelle Simon, Bing Ge, Tony Kwan, Warren A. Cheung, Tomi Pastinen, Catherine Laprise

**Author notes:** **Corresponding author** Catherine Laprise.

## Abstract

**Background:** The 17q12-21 locus is the most replicated association with asthma. However, no study had described the genetic mechanisms underlying this association considering all genes of the locus in immune cell samples isolated from asthmatic and non-asthmatic individuals.

**Objective:** This study takes benefit of samples from naïve CD4^+^ T cells and eosinophils isolated from the same 200 individuals to describe specific interactions between genetic variants, gene expression and DNA methylation levels for the 17q12-21 asthma locus.

**Methods and Results:** After isolation of naïve CD4^+^ T cells and eosinophils from blood samples, next generation sequencing was used to measure DNA methylation levels and gene expression counts. Genetic interactions were then evaluated considering genetic variants from imputed genotype data. In naïve CD4^+^ T cells but not eosinophils, 20 SNPs in the fourth and fifth haplotype blocks modulated both *GSDMA* expression and methylation levels, showing an opposite pattern of allele frequencies and expression counts in asthmatics compared to controls. Moreover, negative correlations have been measured between methylation levels of CpG sites located within the 1.5 kb region from the transcription start site of *GSDMA* and its expression counts.

**Conclusion:** Availability of sequencing data from two key cell types isolated from asthmatic and non-asthmatic individuals allowed identifying a new gene in naïve CD4^+^ T cells that drives the association with the 17q12-21 locus, leading to a better understanding of the genetic mechanisms taking place in it.

## Introduction

The 17q12-21 locus harboring ORMDL sphingolipid biosynthesis regulator 3 (*ORMDL3*) and Gasdermin B (*GSDMB*) genes exhibits one of the most significant and replicated associations with asthma [MIM: 600807].^1–12^ Even though this locus was associated with this disease without considering any specific age of onset as it was done using the Saguenay‒Lac-Saint-Jean (SLSJ) asthma familial cohort before,^11^ it is principally with the condition of childhood.^13, 14^ Several studies explored the complex interactions between SNPs, gene expressions and epigenetic modifications at this locus, primarily focusing on the levels of expressed *ORMDL3* and *GSDMB.*^12, 15–25^ Given that DNA methylation and gene expression signals are cell-type specific,^26, 27^ isolated materials such as lymphocytes from blood^17–21^ and, to a lesser extent, bronchial epithelial cells from biopsies of bronchi or bronchoalveolar lavages^15, 16^ have been used in a number of studies to investigate such interactions.

The study published by Schmiedel and colleagues described *ORMDL3* and *GSDMB* expression in the largest diversity of isolates, including eight different lymphocyte types, monocytes and dendritic cells.^20^ Theirs described genetic variants and epigenetic marks (H3K27ac, CTCF binding and DNase I hypersensitivity sites [DHS]), which regulated *ORMDL3* expression in various immune cell types and pinpointed T lymphocytes to be key effectors in this locus. However, this and further studies did not look at the underlying genetic mechanisms and the interactions among all the other genes in the locus, and they did not include individuals with asthma to decipher the disease-specific processes. Moreover, there is a lack of data from isolated granulocytes, even though eosinophils are well known key cells in asthma physiopathology.^28^

Furthermore, although DNA methylation profiles are key epigenetic mechanisms to understanding the association between 17q12-21 locus and asthma, they have been less studied than expression levels so far, and they were assessed with methods targeting specific methylation sites (pyrosequencing and microarrays).^29, 30^ Recently, a methylome bisulfite sequencing capture panel was designed to target regulatory regions specific to immune cells with the consideration on DHS or active chromatin sites, hypomethylated footprints, and autoimmune SNPs from genome-wide association study (GWAS) catalog,^31^ giving a better coverage of the 17q12-21 locus compared to widely used DNA methylation arrays (e.g. Illumina methylation beadchips).

This publication addresses the aforementioned missing components of earlier studies by using an integrative multi-omic approach to characterize the functional genetic mechanisms at this locus in eosinophils and naïve CD4^+^ T cells. Both were selected because of their important role regarding the immune response in the asthma pathogenesis, and according to previous results of methylation analyses with the SLSJ cohort; a French-Canadian one located in the northeastern part of the Quebec province.^20, 28, 32, 33^ Naïve CD4^+^ T cells and eosinophils were obtained from asthmatic and non-asthmatic individuals who were participants in this cohort.^34^ This study combines genomic (asthma-associated SNPs), epigenomic, and transcriptomic (DNA methylation levels and gene expression counts from next generation sequencing) data to better decipher the genetic mechanisms underlying the association between the 17q12-21 locus and this disease.

## Subjects and Methods

Figure 1 shows the different types of data used in this study regarding the sampling and sizing (number of samples, SNPs, CpGs methylation levels or gene expression counts) accessible for each analysis performed considering quality control filtering steps and covariates availability.

**Figure 1.**
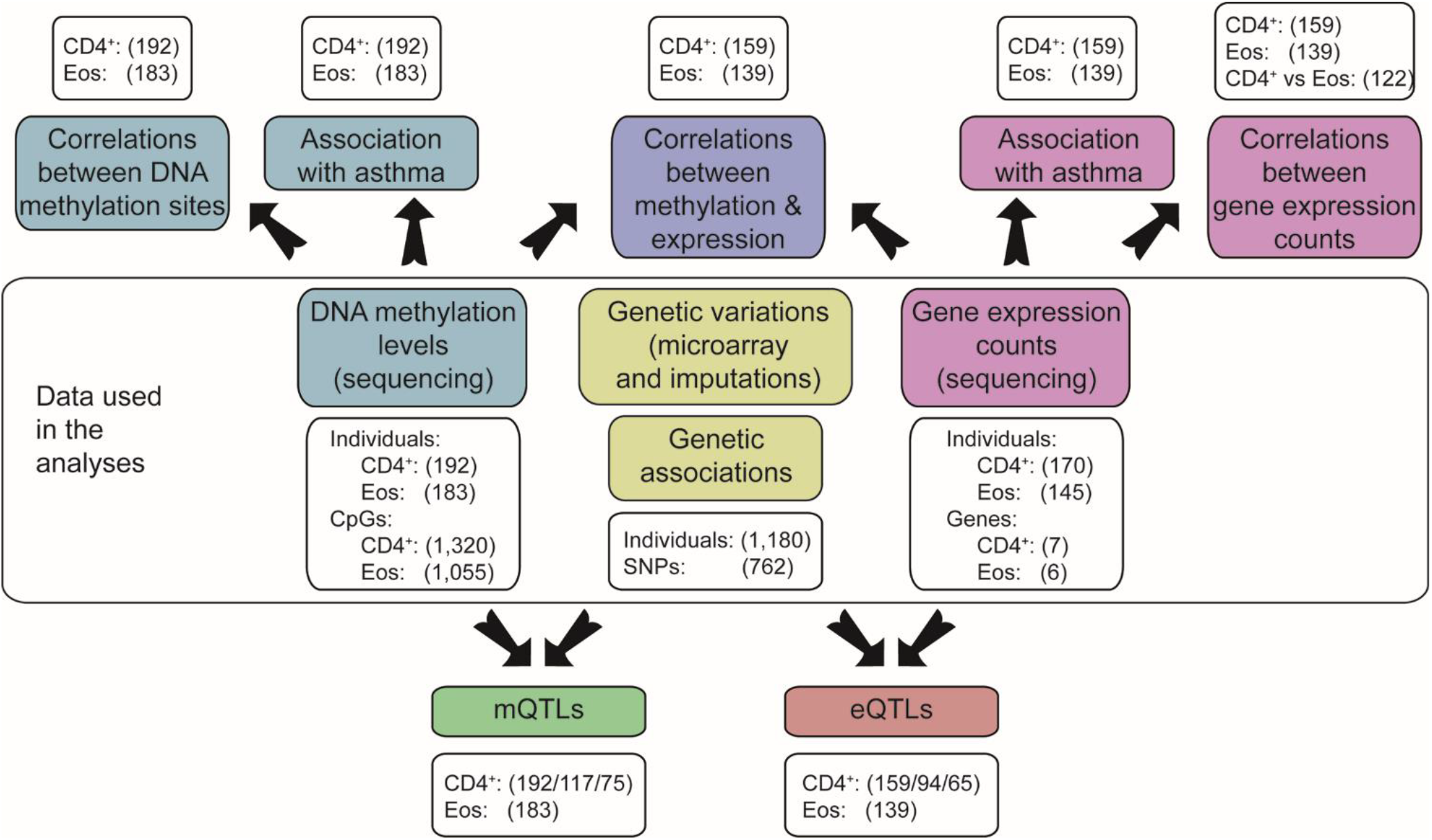
Schematic view of the data used and analyses performed in this study. Data used for the analyses included genetic variations from microarrays with imputations, as well as methylation levels and expression counts from next generation sequencing. The top part of the figure shows the analyses performed with methylation and expression data. The bottom part presents the integration of these data with genetic variations. Each white box indicates the number of samples and the one of genetic data (SNPs, CpGs or genes) included in the analyses, considering quality control filtering and the presence of all covariates in the models used. When three numbers are listed within parentheses, the first corresponds to the number of samples for the principal analysis (with all considered), the second where only individuals with asthma were analyzed and the third where controls were only. Abbreviations: CD4^+^ = naïve CD4^+^ T cells, Eos = eosinophils.

### SLSJ asthma cohort

The complete description of recruitment and clinical evaluation carried out with the SLSJ cohort can be found in Laprise et al.^34^ Briefly, families were included in the study through probands ascertained and the general and respiratory health of all individuals were evaluated using a standardized questionnaire and pulmonary function tests according to the American Thoracic Society guidelines.^35^

The SLSJ asthma cohort includes 1,394 individuals distributed in 271 families from which 1,214 individuals (representing 254 families) have genotypic information and 1,200 of them have imputed data available. See Table 1 for the phenotypic characteristics of the 1,200 individuals included in the association analysis with asthma. From these individuals, 215 were recruited for isolating naïve CD4^+^ T cells and eosinophils to measure methylation and expression levels by next generation sequencing. After being isolated and the quality control step (see below the isolation section), 173 naïve CD4^+^ T cell and 146 eosinophil samples were available for analyses. Shown in Table 1, their phenotypic characteristics. The *Centre intégré universitaire de santé et de services sociaux du Saguenay‒Lac-Saint-Jean* ethics committee approved the study, and all subjects gave informed consent.

**Table 1:**
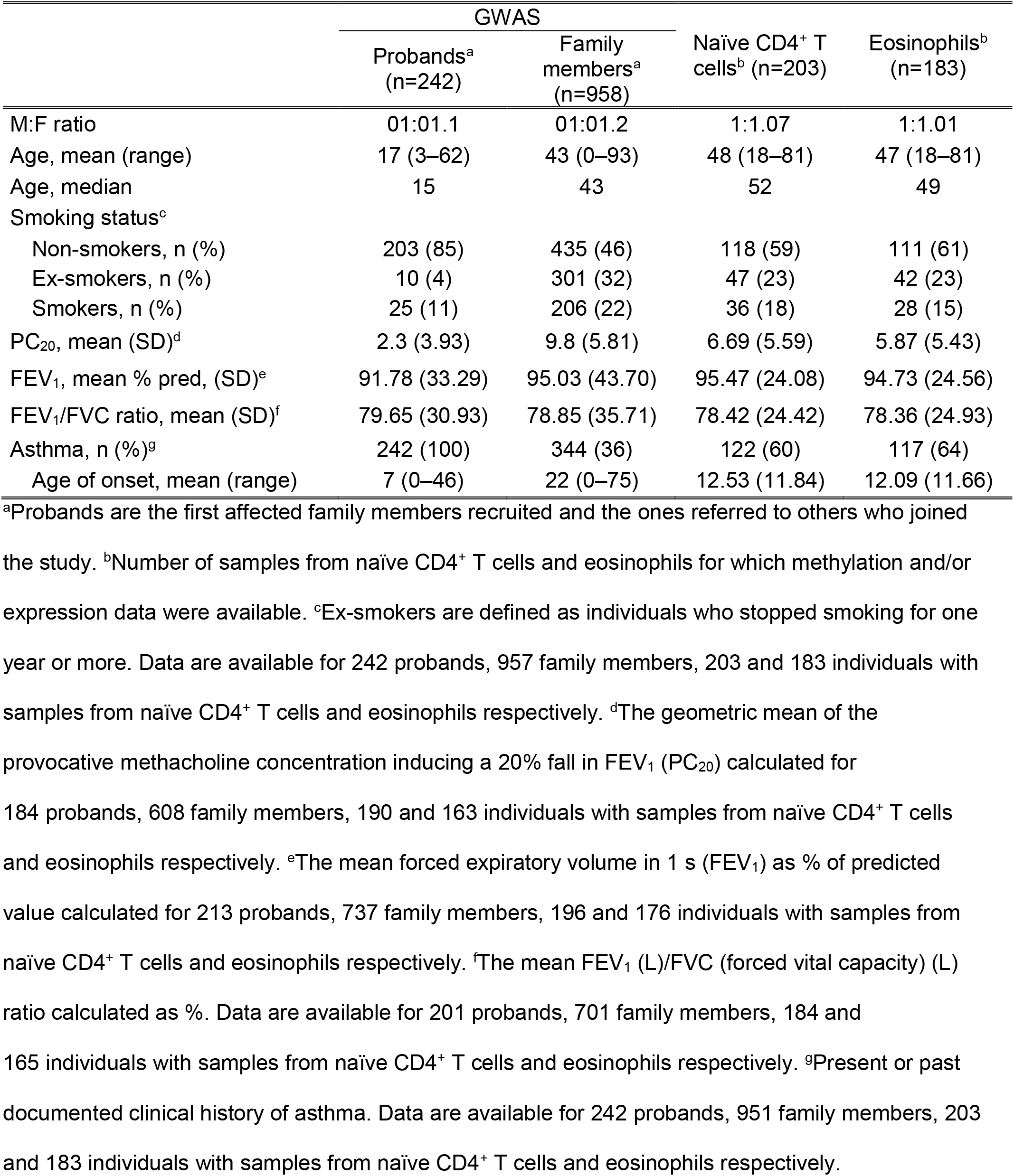
Phenotypic characteristics of the individuals genotyped and those included in the analyses performed on naïve CD4^+^ T cells and eosinophils

|  | GWAS |  | Naïve CD4<sup>+</sup> T cells<sup>b</sup> (n=203) | Eosinophils<sup>b</sup> (n=183) |
| --- | --- | --- | --- | --- |
|  | Probands<sup>a</sup> (n=242) | Family members<sup>a</sup> (n=958) |  |  |
| M:F ratio | 01:01.1 | 01:01.2 | 1:1.07 | 1:1.01 |
| Age, mean (range) | 17 (3–62) | 43 (0–93) | 48 (18–81) | 47 (18–81) |
| Age, median | 15 | 43 | 52 | 49 |
| Smoking status<sup>c</sup> |  |  |  |  |
| Non-smokers, n (%) | 203 (85) | 435 (46) | 118 (59) | 111 (61) |
| Ex-smokers, n (%) | 10 (4) | 301 (32) | 47 (23) | 42 (23) |
| Smokers, n (%) | 25 (11) | 206 (22) | 36 (18) | 28 (15) |
| PC<sub>20</sub>, mean (SD)<sup>d</sup> | 2.3 (3.93) | 9.8 (5.81) | 6.69 (5.59) | 5.87 (5.43) |
| FEV<sub>1</sub>, mean % pred, (SD)<sup>e</sup> | 91.78 (33.29) | 95.03 (43.70) | 95.47 (24.08) | 94.73 (24.56) |
| FEV<sub>1</sub>/FVC ratio, mean (SD)<sup>f</sup> | 79.65 (30.93) | 78.85 (35.71) | 78.42 (24.42) | 78.36 (24.93) |
| Asthma, n (%)<sup>g</sup> | 242 (100) | 344 (36) | 122 (60) | 117 (64) |
| Age of onset, mean (range) | 7 (0–46) | 22 (0–75) | 12.53 (11.84) | 12.09 (11.66) |

### Genetic data

The genotypes for the SLSJ asthma cohort were obtained from Illumina Human610-Quad BeadChip. The pre-phasing step for the imputation process was performed with the Shapeit2 Software using the duoHMM method that combines estimated haplotypes with pedigree information to take advantage of known structure related to the latter.^36^ Impute2 Software was used for imputation^37^ with the 1,000 Genome Project database (phase 3)^38^ and the UK10K one as reference samples. See Supplemental Data for details about quality control steps. Based on the review published by Stein et al.,^12^ the 17q12-21 locus was defined as chr17:37,826,875-38,134,519 for our analyses. This region includes 11 sequences coding for microRNAs or genes (Post-GPI attachment to proteins 3 [*PGAP3*], Erb-B2 receptor tyrosine kinase 2 [*ERBB2*], microRNA [miR]-4728, Migration and invasion enhancer 1 [*MIEN1*], Growth factor receptor-bound protein 7 [*GRB7*], IKAROS Family Zinc Finger 3 [*IKZF3*], Zona Pellucida-Binding Protein 2 [*ZPBP2*], Gasdermin A [*GSDMA*] and B [*GSDMB*]*, ORMDL3*, as well as Leucine-Rich Repeat-Containing 3C [*LRRC3C*]). *ORMDL3*, *GSDMB* and *ZPBP2* were the most replicated asthma associated genes. After quality filtering, association with asthma was evaluated for 762 SNPs for this specific region in 1,180 DNA samples.

### Isolation of naïve CD4^+^ T cells and eosinophils

Naïve CD4^+^ T cells, isolated in this project, were defined as units with CD3^+^, CD4^+^, CD45RA^+^, CD45RO^−^ markers. These and eosinophils were from 200 ml blood samples from 215 individuals, a subset of the SLSJ asthma cohort, using the EasySep Human Naïve CD4+ T Cell Isolation Kit (STEMCELL Technologies, Vancouver, BC, Canada) and anti-CD16 MicroBeads respectively according to protocols described (Miltenyi Biotec, Aubern, CA, USA)^39, 40^. A description of the different steps can be found in the Supplemental Data.

### Targeted bisulfite sequencing (DNA methylation levels)

Bisulfite sequencing was done for a custom methyl capture panel of identified functional immune genetic regions as previously described.^31, 41^ These covered a total of 4,609,564 CpGs for a sum of 822,884 regions and 119,089,296 bp sequenced.^31^ The MCC-Seq methylation was developed and optimized in collaboration with R&D at Roche NimbleGen. Details of the protocols used can be found in the Supplemental Data. After quality control filtering, methylation data were available for samples from 192 naïve CD4^+^ T cells and 183 eosinophils and 1,320 CpGs for analyses of the former type as well as 1,055 CpGs of the latter one.

### RNA sequencing (gene expression counts)

RNA sequencing was performed at the McGill University and Génome Québec Innovation Centre with the Illumina TruSeq Stranded Total RNA Sample Prep Kit (Illumina, CA, USA). A detailed description of the method used can be found in the Supplemental Data. After quality control filtering and given that the availability of all covariates included in statistical models, transcriptomic data were accessible for 159 naïve CD4^+^ T cell and 139 eosinophil samples. Using as a cut-off ten counts in totality while considering all samples together, a total of seven genes were expressed by naïve CD4^+^ T cells and six by eosinophils.

### Statistical methods

#### Genetic association and definition of haplotype blocks

The significance threshold was calculated with Nyholt’s method^42^ according to the number of independent SNPs in the 17q12-21 asthma locus and was set to 0.001. In this work, we carried out the MFQLS test implemented in the Workbench for Integrated Superfast Association study with Related Data (WISARD) toolkit, a quasi-likelihood method of estimation extended from the Cochran-Armitage test, suited for familial cohorts considering kinship coefficients, and can be used to associate multiple phenotypes with covariates.^43, 44^ The analyses were made with the asthma phenotype using sex, age and smoking status (a never or an ever smoker) as covariates.

The Haploview Software was used to delimit the haplotype blocks of the 17q12-21 asthma locus with an algorithm taken from Gabriel et al.^45, 46^ Methylation blocks for any cell type were also established using r^2^ values between each adjacent CpGs^47^ in a similar way haplotype blocks were defined with linkage disequilibrium ones (D’ and r^2^). For methylation, they were identified as two or more CpGs for which all pairwise r^2^ value is equal or above 0.5 as described by Guo and colleagues.^47^

#### Estimation of cell proportion and surrogate variables

The cell-type contribution approach was used to estimate the probable variation between the purity of samples from naïve CD4^+^ T cells and eosinophils and the values calculated, and the latter was employed as covariates in our analysis models. The method developed by Houseman^48^ and implemented in the RnBeads R package was applied using the whole genome methylation sequencing data. The reference samples used for hematocytes included ten types: CD14^+^, CD19^+^, CD4^+^, CD8^+^, eosinophils, granulocytes, neutrophils, PBMCs and white blood cells.^27^ Moreover, in order to take into account hidden confounders and relatedness between samples, surrogate variables were calculated using the R package sva.^49^ They were by considering all covariates included in models analyzed.

#### Analyses of gene expression counts and DNA methylation levels

Gene expression counts were normalized on the library size and log_2_ transformed while DNA methylation levels had undergone a logit transformation using the R packages DESeq2 and car respectively.^50^ Correlations between gene expression counts from the two cell types, or from within a unique one, were performed using Pearson’s r. Correlations between gene expression counts and DNA methylation levels were assessed with this and also with a negative binomial model (using normalized-only values for the former) considering age at sampling, sex, smoking status (a never or an ever smoker), proportion of naïve CD4^+^ T cells or eosinophils, according to the origin of data analyzed and surrogate variables as covariates. The differences, in gene expression counts (normalized-only data) and DNA methylation levels (non-transformed percentages) in accordance with the asthma status, were assessed using a negative binomial and a binomial model respectively, both considering the same covariates as for the previous analyses. False discovery rate (FDR) method was applied to take into account multiple testing.

QTL analyses were performed using the package MatrixEQTL in R^51^ for the gene expression counts (expression quantitative trait loci: eQTLs) and for the DNA methylation levels (methylation quantitative trait loci: mQTLs) with same covariates as previously. In mQTL analyses, CpGs were considered to be linked specifically if they were located within the gene body or the region within 1.5 kb from the transcription start site (TSS). As information from these focused on genetic variation in haplotype blocks interacting with the expression or methylation, significance threshold was calculated using Bonferroni correction considering the number of these structures included and the one of genes expressed by each cell type. The significance threshold was set to 0.001 for naïve CD4^+^ T cells and eosinophils (0.05/[7*7] and 0.05/[7*6] respectively).

Several analyses were performed on stratified samples according to asthma status or on affected and unaffected mother-child duos. In accordance with the small number of samples included, 1,000 analyses were performed after permutations between the asthmatic and non-asthmatic individuals to estimate the chance each result has to be a false positive.

Analyses using the mediation package in R were performed to describe more accurately the link between DNA methylation levels measured within the 1.5 kb from the *GSDMA* TSS region, asthma-associated SNPs in the fourth and fifth haplotype blocks identified in the SLSJ cohort and its gene expression counts. In the models used, methylation was considered as the mediator of the SNP-expression association of *GSDMA*. The same covariates as usual were applied to them, and 1,000 bootstraps were run to estimate the confidence intervals.

## Results

This project used genomic, epigenomic and transcriptomic data to better understand the genetic interactions underlying the association between a specific region of the 17q12-21 locus (chr17:37,826,875-38,134,519) and asthma in two immune cell types (naïve CD4^+^ T cells and eosinophils). SNPs, gene expression and methylation data were first analyzed separately to establish and compare their profiles in naïve CD4^+^ T cells (CD3^+^, CD4^+^, CD45RA^+^, CD45RO^−^) and eosinophils. Then, genetic interactions were assessed using eQTL and mQTL analyses and correlations between DNA methylation levels and gene expression counts (Figure 1). This figure also indicates the number of samples as well as the genetic data used for each analysis after quality control filtering.

### Haplotype blocks in the 17q12-21 locus are associated with asthma in the SLSJ cohort

The analysis of linkage disequilibrium between 762 SNPs in the 17q12-21 locus identified seven haplotype blocks in the SLSJ asthma cohort compared to the two from 17 SNPs in the European population^12^ (Figures S1-S2). Of the 248 SNPs associated with asthma with p <0.05, 20 remained of statistical significance after correction for multiple testing (p <0.001) and all were located in the fourth as well as the fifth haplotype blocks (Table S1). SNP rs869402 in the fourth block and rs9303281 in the fifth one exhibited the most statistically significant associations (p = 5.35e-04 and 2.60e-04, respectively). The analysis including only individuals with age of onset <17 years old as asthmatic individuals gave very similar results (p = 1.73e-04 and p = 2.01e-04 for the same two SNPs; see Table S1).

### Gene expression and DNA methylation profiles harbored by naïve CD4^+^ T cells and eosinophils in the 17q12-21 asthma locus are differents

#### Naïve CD4^+^ T cells and eosinophils express six genes in common at the 17q12-21 asthma locus

Gene expression analyses demonstrated that of the eleven ones located in the 17q12-21 locus, seven were detectable in naïve CD4^+^ T cells (*PGAP3, ERBB2, MIEN1, IKZF3, GSDMB, ORMDL3* and *GSDMA*) and six in eosinophils (*PGAP3, ERBB2, MIEN1, IKZF3, GSDMB* and *ORMDL3*). Detailed results for correlations between gene expression counts between the two cell types or within each one were available in Tables S2-S4. In both cell types, the most significant pairwise correlation was between *GSDMB* and *ORMDL3* (naïve CD4^+^ T cells: r = 0.668 [FDR <1.0e-20]; eosinophils: r = 0.888 [FDR <1.0e-20]) (Figure S3). No genes were differentially expressed in analyses that compare cells isolated from individuals with and without asthma (Table S5).

#### Methylation profiles in the 17q12-21 asthma locus are different in naïve CD4^+^ T cells and eosinophils

DNA methylation levels were measured in naïve CD4^+^ T cells and eosinophils in order to compare their profiles and determine if levels of CpGs in this region are associated with asthma. Using a custom sequencing panel, a total of 1,320 and 1,055 CpGs have been measured after quality control filtering for naïve CD4^+^ T cells and eosinophils respectively compared to the 166 CpGs evaluated by the Infinium Human Methylation 450k BeadChip in this same locus. Of these 166 CpGs, 156 and 115 were among the 1,320 and 1,055 CpGs included in this study on naïve CD4^+^ T cells and eosinophils, respectively. In the same way that haplotype blocks were evaluated for SNPs of the 17q12-21 asthma locus, r^2^ were computed for each neighboring pairs of CpGs in naïve CD4^+^ T cells and eosinophils separately to establish cell-type-specific methylation patterns in this region as previously described by Guo and colleagues^47^ (Figure 2). Some differences were observed between the two cell types particularly within or near 1.5 kb regions from the gene TSS. The most obvious one was in *IKZF3* as correlation values were between 0.20 and 0.74 (mean r^2^ = 0.57) in naïve CD4^+^ T cells compared to 0.02 and 0.55 (mean r^2^ = 0.19) in eosinophils. Moreover, the *IKZF3* promoter region included three 68–90 bp long methylation blocks encompassing four, five and six CpGs respectively in naïve CD4^+^ T cells compared to only one block of two CpGs in eosinophils (Figure 2). The methylation levels were compared between asthmatics and non-asthmatics individuals in naïve CD4^+^ T cells and eosinophils (Table S6). Eighty-six CpGs were associated with asthma in naïve CD4^+^ T cells (FDR 5%) of which, six were located within the 1.5 kb region from four gene TSSs (*ERBB2*, *PGAP3*, *IKZF3* and *ORMDL3*. Sixty-three CpGs were differentially methylated in eosinophils of which, 11 CpGs were located within five TSS regions (*ERBB2*, *PGAP3*, *IKZF3*, *GSDMB* and *ORMDL3*).

**Figure 2.**
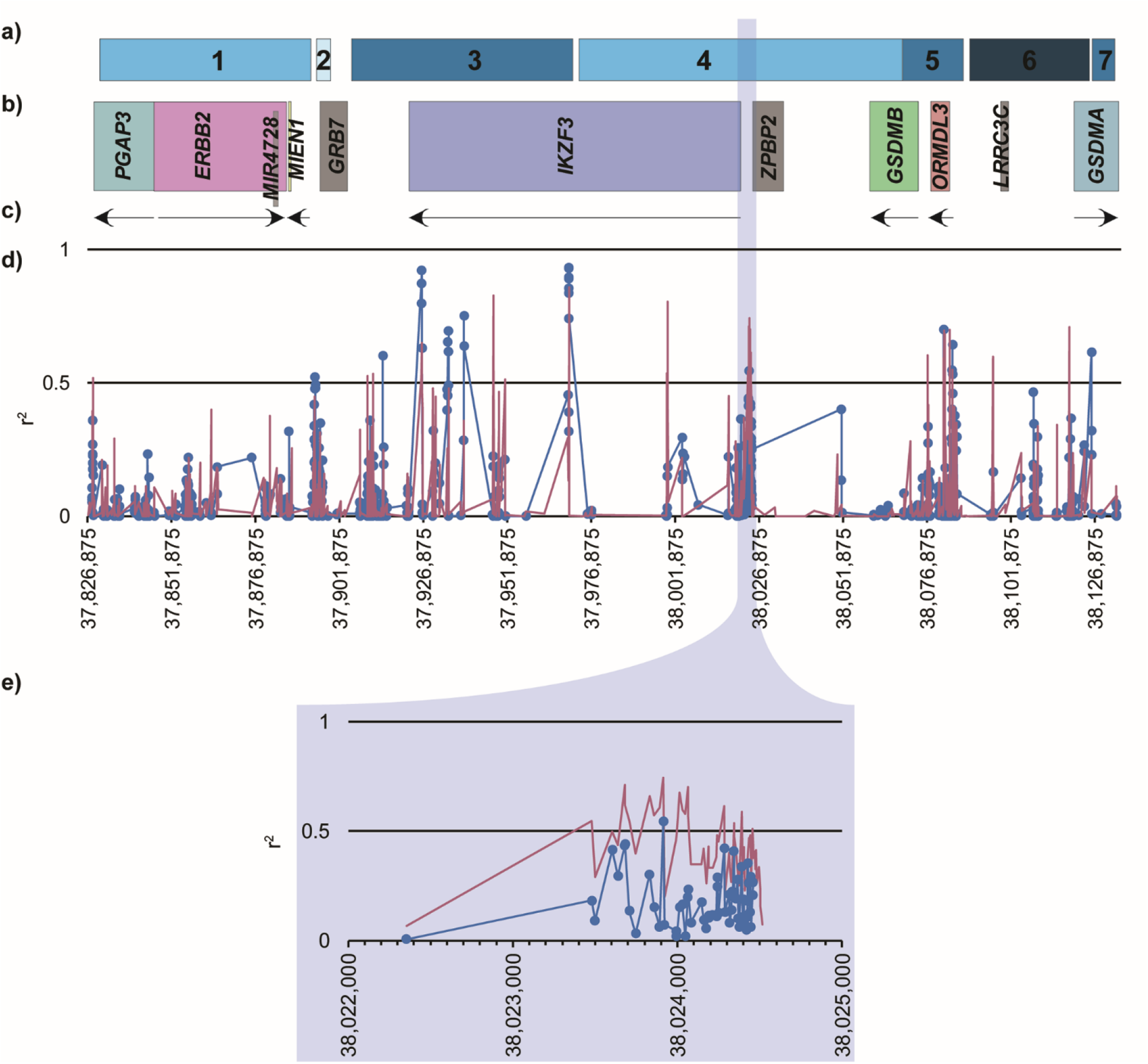
Correlation between methylation sites in naïve CD4^+^ T cells and eosinophils. Sections (a) and (b) depict the seven haplotype blocks identified in the Saguenay‒Lac-Saint-Jean asthma cohort and the eleven genes located in this locus. Genes colored in grey are those not expressed in this study by naïve CD4^+^ T cells. The section (c) indicates the strand of each gene expressed by one of the two cell types studied, and the section (d) depicts r^2^ that correlates between methylation sites in samples recovered from naïve CD4^+^ T cells (pink line) and eosinophils (blue line with round markers) across the region of interest. Section (e) zooms in the promoter region of *IKZF3* that includes several methylation blocks in naïve CD4^+^ T cells but not eosinophils (they are defined by two or more CpGs with r^2^ values >0.5).

### Associations between genetic, epigenetic and transcriptomic data at the 17q12-21 locus are specific to naïve CD4^+^ T cells and eosinophils

Several genetic interactions were analyzed to better understand the mechanisms underlying the association between the 17q12-21 locus and asthma. CpGs located within the 1.5 kb region from TSS and gene body were investigated separately given that they usually have opposite impacts on expression.^52^

#### SNPs of the 17q12-21 locus show different associations with gene expression counts (eQTLs) or DNA methylation levels (mQTLs) in naïve CD4^+^ T cells and eosinophils

In order to measure genetic interactions specific to asthma, disease-associated SNPs in the 17q12-21 locus were tested for association with gene expression and methylation levels at CpGs located within 1.5 kb region from TSSs. In naïve CD4^+^ T cell samples, 78 eQTLs in *IKZF3*, *GSDMB*, *ORMDL3* and *GSDMA* and 58 mQTLs in *MIEN1* and *GSDMA* were observed (Figure 3a). In eosinophil ones, 21 eQTLs in *GSDMB* and *ORMDL3* and no mQTL were detected (Figure 3b).

**Figure 3.**
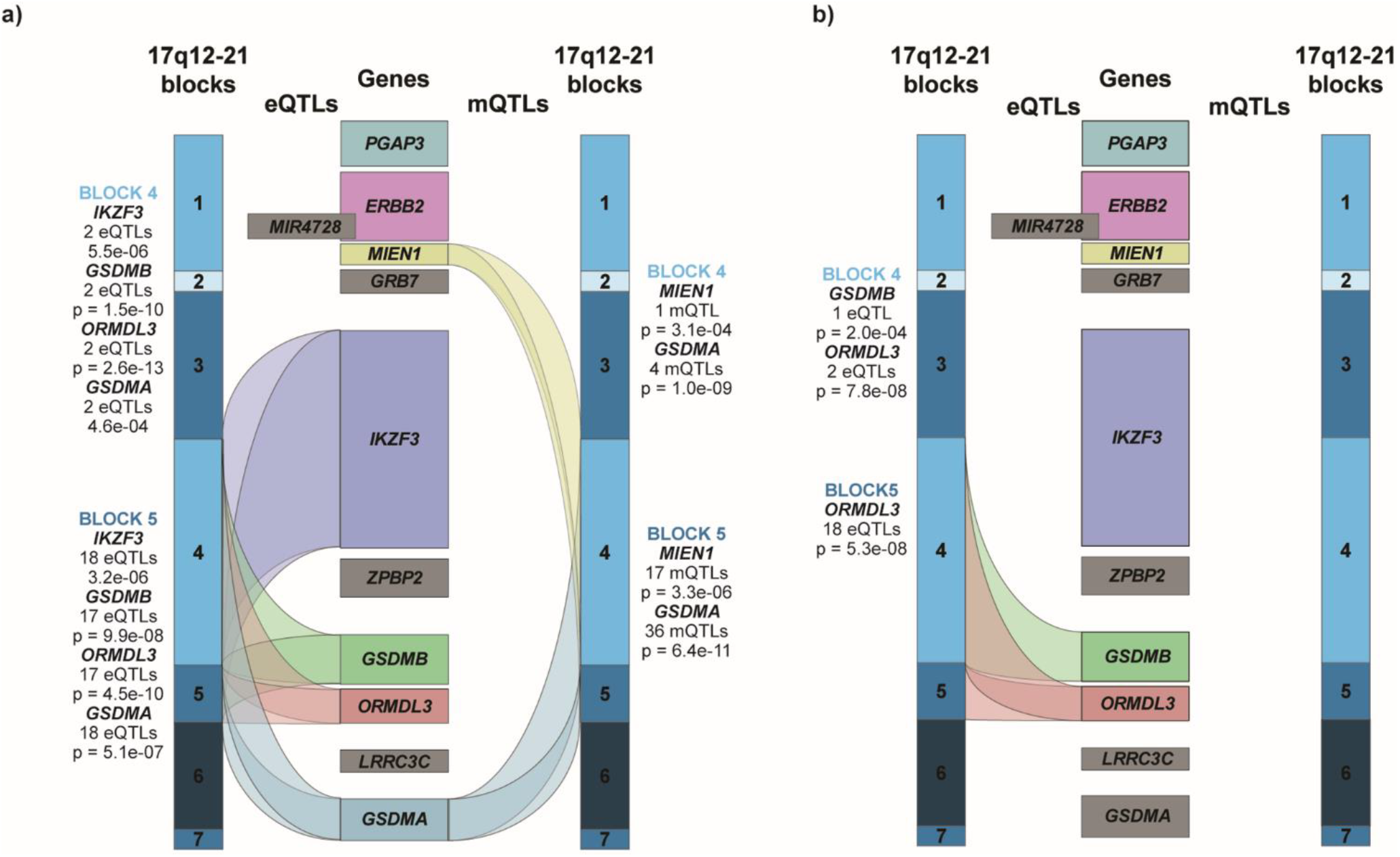
eQTLs and mQTLs of the 17q12-21 locus by cell types. This figure illustrates the eQTLs and mQTLs considering SNPs associated with asthma identified in naïve CD4^+^ T cells (a) and eosinophils (b). The seven haplotype blocks detected in the Saguenay‒ Lac-Saint-Jean asthma cohort for the chr17:37,826,875-38,134,519 region are at the outer edges in different shades of blue and the middle are genes located in the same area. Genes colored in grey are those not expressed by each cell type in this study. The number of eQTLs (left part of any figure) and mQTLs (right part) linked to each gene for each haplotype block are shown as indicated. P values for the most statistically significant QTLs are listed for each gene in each block.

Genetic interactions observed for asthma-associated SNPs were compared to mQTLs for methylation sites located within gene bodies or intergenic regions and to results considering all SNPs in the 17q12-21 locus for comprehensive and complementary purposes. The genes and the numbers of QTLs varied depending on the location of the CpGs (1.5 kb region from TSS or gene body) and the type of SNPs (asthma-associated or not) investigated. See Tables S7 to S10 and Figures S4 and S5 for all the significant results.

#### Different DNA methylation-expression associations are observed in naïve CD4+ T cells and eosinophils at the 17q12-21 locus

Regression analyses detected correlations between DNA methylation levels and gene expression counts in naïve CD4^+^ T cells (Table 2). Gene expression counts of *GSDMA* was negatively correlated with methylation levels of two CpGs located within its 1.5 kb region from TSS (r = −0.238 [FDR = 0.039] and −0.255 [FDR = 0.028]). Furthermore, those of *ORMDL3* had a positive correlation with one CpG within its 1.5 kb region from TSS (r = 0.308 [FDR = 7.55e-05]). The expression of *ERBB2, IKZF3*, *ORMDL3* and *GSDMA* correlated with methylation levels of CpGs within their gene body. All significant correlations also including intergenic methylation sites can be seen in Table S11. In eosinophils, *s*ignificant correlations were also uncovered for CpGs located in the gene body of *ERBB2* and *IKZF3* (Tables 3 and S12) but none with CpGs within the 1.5 kb from TSSs.

**Table 2.**
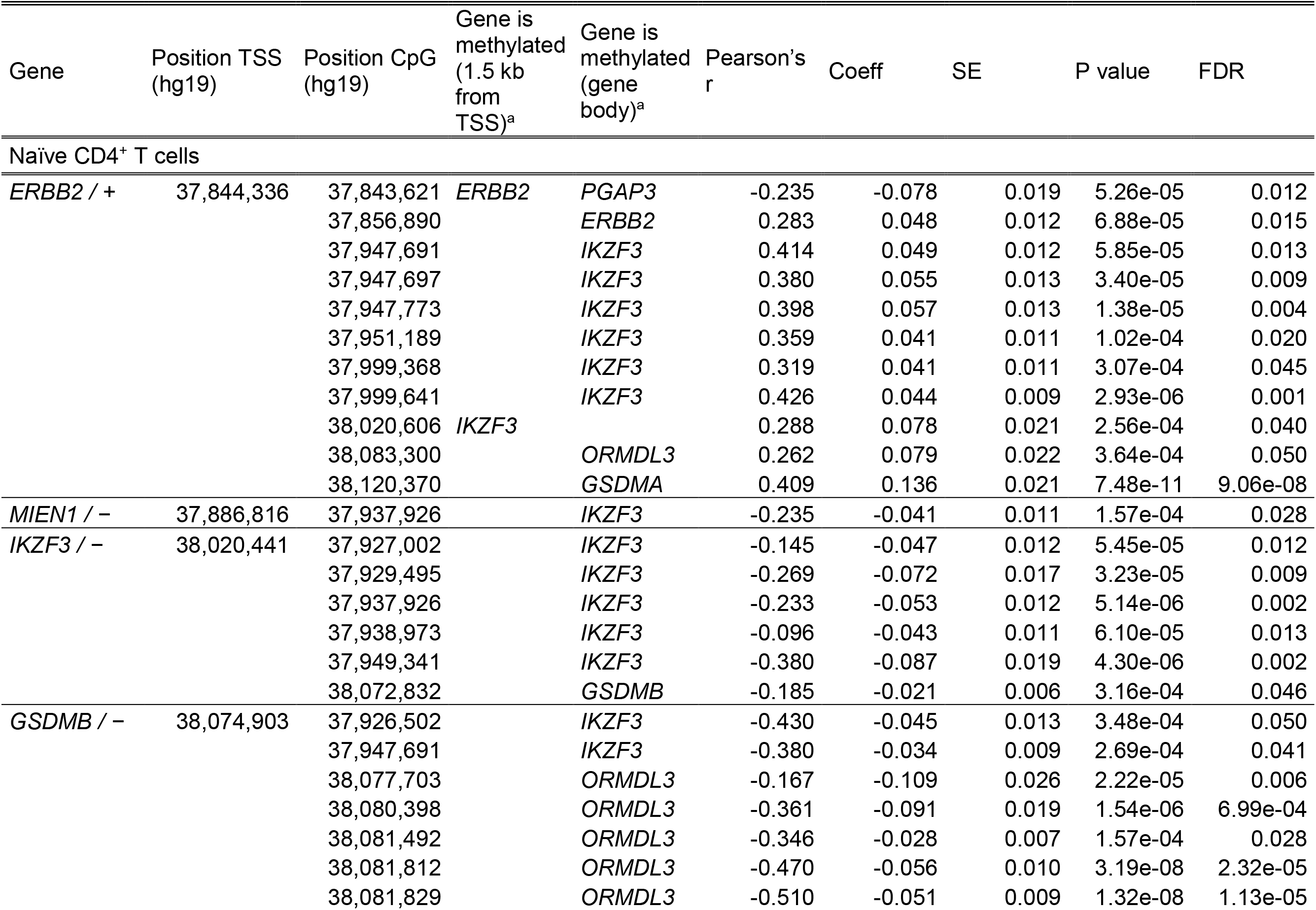
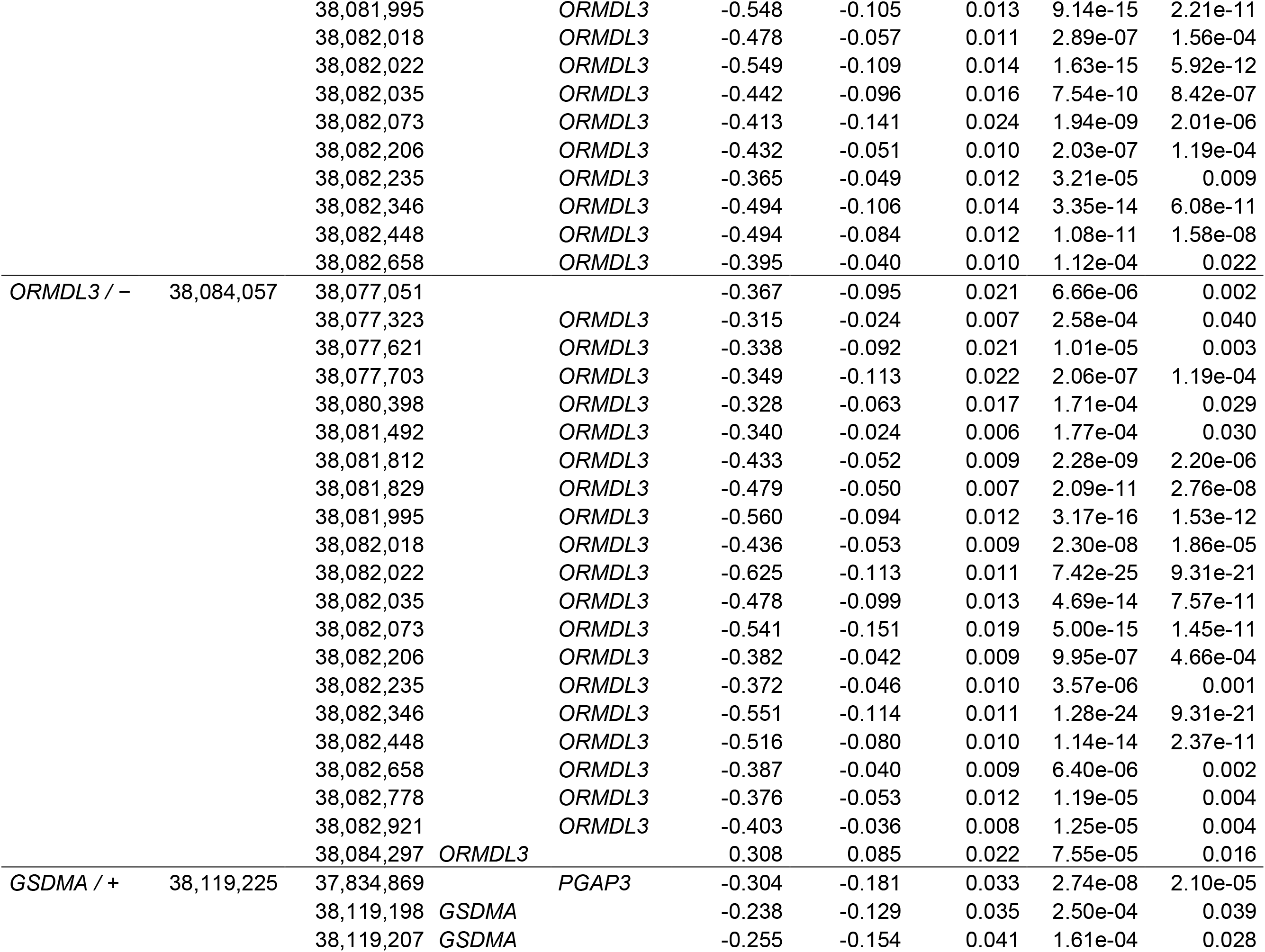
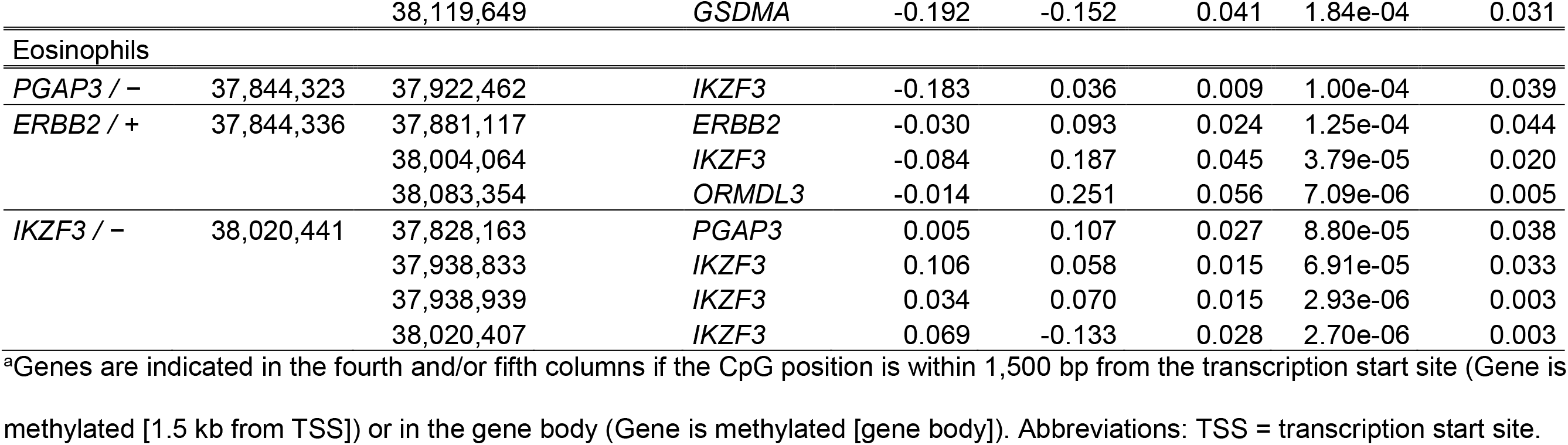
Significant associations between gene expression counts and DNA methylation levels of CpGs located within 1.5 kb from a TSS or a gene body in naïve CD4^+^ Tcells or eosinophils

### *GSDMA* expression is associated with genetic variations and methylation levels of the 17q12-21 locus in naïve CD4^+^ T cells

Different analyses in this study showed that genetic, epigenetic and expression of *GSDMA* were interacting. First, all asthma-associated SNPs (two in the fourth haplotype block and 18 in the fifth one) modulated both the gene expression counts and the DNA methylation levels within the 1.5 kb region from TSS of *GSDMA* (Figures 3a and 4). All these SNPs formed mQTLs with the same two methylation sites located 18 bp (chr17:38,119,207) and 27 bp (chr17:38,119,198) upstream of *GSDMA* TSS (Figure 4d and 4e). The two CpGs were positively correlated (r^2^ = 0.710 and FDR = 1.23e-47), and each was negatively with *GSDMA* expression (r = −0.238 with FDR = 0.039 and r = −0.255 with FDR = 0.028; Figure 4f). mQTLs were also observed for CpGs located in *GSDMA* gene body and one of these three DNA methylation sites has its level correlated with its expression (r = −0.192 with FDR = 0.031).

**Figure 4.**
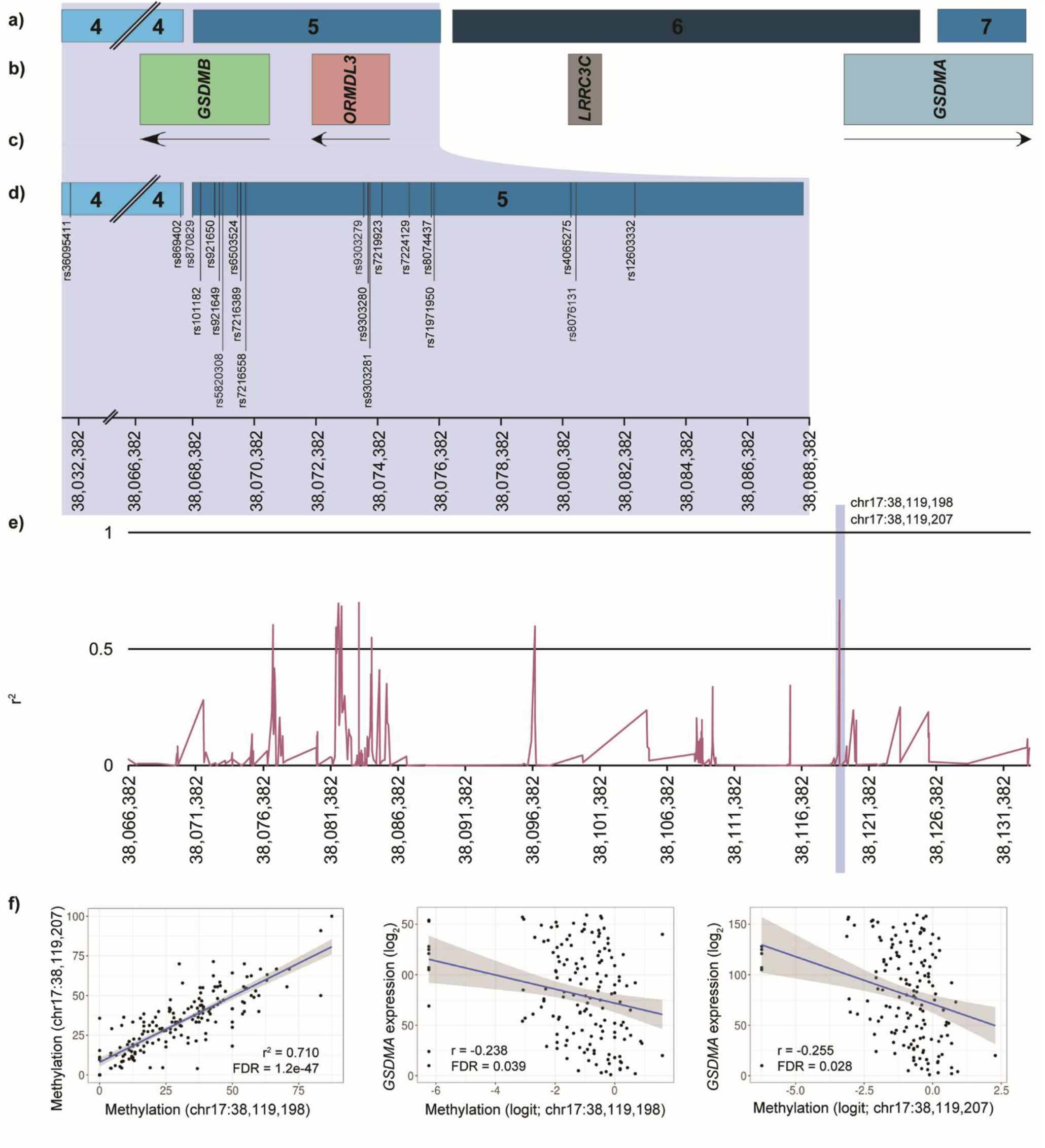
Methylation and expression profiles of *GSDMA* in samples from naïve CD4^+^ T cells. This figure shows: (a) the 17q12-21 region from the fourth to the seventh haplotype blocks; (b) the genes located in this region, and (c) the strand of these. Genes in grey are those not expressed by naïve CD4^+^ T cells. The section (d) zooms in the fourth and fifth haplotype blocks to show all SNPs in eQTLs and mQTLs that were identified in naïve CD4^+^ T cells for *GSDMA*. All SNPs included in eQTLs and are in mQTLs with the two correlated CpGs indicated in (e), where r^2^ is depicted for correlations between adjacent methylation sites located in the region including the end of the fourth block to the seventh one. The section (f) presents correlation between the two methylation sites included in mQTLs and the one between each of them and *GSDMA* expression.

Since the association between genetic variants and the disease occurring at the 17q12-21 locus is mainly driven by individuals with early-onset asthma,^13, 14^ correlation analyses were performed with DNA methylation levels from different mother-child duos for the two CpGs included in the significant mQTLs at *GSDMA* TSS. Analyses were between unaffected mother and affected child duos, all asthma status mother and affected child duos and affected mother and affected child duos to demonstrate the possible heritability of these DNA methylation levels. Also, they showed an increase of the correlation values for the two CpGs with Pearson’s r of 0.068 to 0.412 for the 17:38,119,198 CpG and of 0.112 to 0.238 for the 17:38,119,207 CpG in the case of the duos with unaffected mother and affected child and those with affected mother and affected child respectively (Figure S6).

Mediation analyses were performed to find out the proportion of the association between methylation levels near *GSDMA* TSS and its expression signal modulated by the 20 SNPs included in both eQTLs and mQTLs. These demonstrated no significant model (data not shown).

Finally, analyses of genetic interactions (correlation between methylation and expression levels of *GSDMA*, eQTLs and mQTLs) were conducted on individuals with and without asthma separately, in order to decipher if the latter is driven by subjects suffering from this disease. Neither of the correlations between methylation and expression, nor mQTL analysis provided significant results after permutations. However, eQTL analysis including SNPs located in the fourth and fifth haplotype blocks revealed 15 eQTLs to be significantly associated with *GSDMA* expression in individuals with asthma and 19 eQTLs in controls (See Table S13 for eQTL results regarding these subjects). Interestingly, minor allele frequencies of the asthma-associated SNPs were lower in asthmatic individuals for all SNPs compared to controls and expression level was higher in subjects with the disease in comparison with the others, showing an opposite pattern in the two groups (Figure 5).

**Figure 5.**
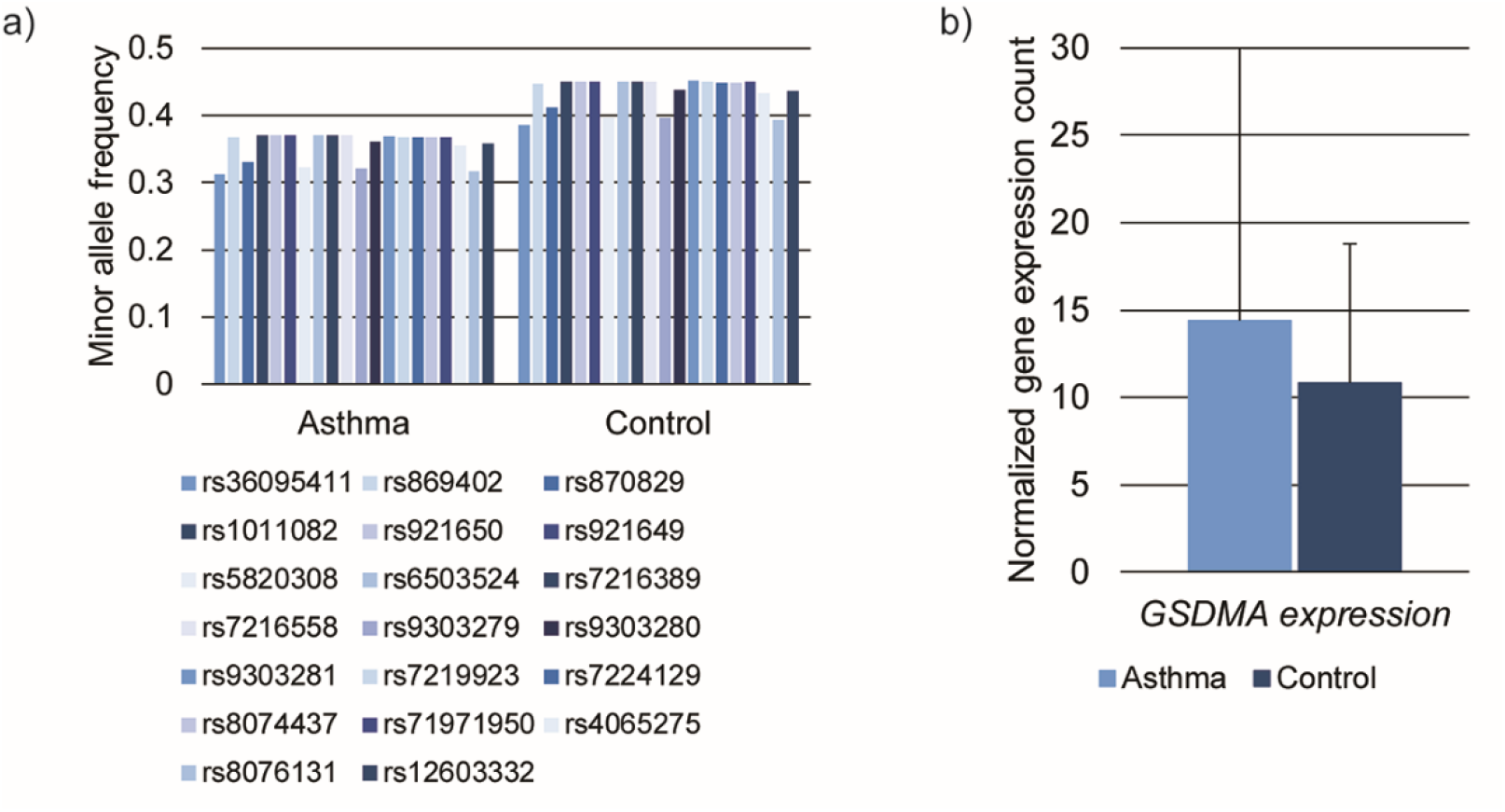
Minor allele frequencies of asthma-associated SNPs and *GSDMA* expression counts by the disease status. This figure shows: (a) the minor allele frequency of all 20 asthma-associated SNPs in the fourth and fifth haplotype blocks of individuals living with asthma and controls from the Saguenay‒Lac-Saint-Jean cohort and (b) the *GSDMA* expression counts in the same groups. Standard deviations are presented for mean values.

## Discussion

In this study, we described genetic, epigenetic and transcriptomic profiles as well as their interactions in a specific region (chr17:37,826,875-38,134,519) of the 17q12-21 asthma locus in naïve CD4^+^ T cells and eosinophils. The limit of this region was selected according to the review by Stein et al.^12^ to include genes from *PGAP3* to *GSDMA*. The feasibility to measure and analyze various data types of cells obtained from the same individuals increase the chance of detecting valid genetic interactions. Furthermore, using data from imputed SNPs combined with next generation sequencing for individuals suffering from asthma or not allowed for a more complete description regarding the genetic interactions specific to this disease phenotype in these two cell types, an approach that was never used for this locus.

Genetic analysis performed in this study identified seven haplotype blocks in the 17q12-21 locus in the SLSJ cohort, in which 20 SNPs located in the fourth and fifth ones were significantly associated with asthma after correction for multiple testing. Among them, eight were already in the literature.^6, 11, 53–55^ The haplotype blocks observed in the SLSJ cohort were similar to those described previously for the European population^12^ except that the second European one was represented by the third to the fifth in the SLSJ cohort. Given that imputed data were used in the SLSJ asthma cohort, the larger number of SNPs analyzed (762 compared to 17) improved the precision in linkage disequilibrium and led to the definition of more blocks and to a better accuracy in interpreting eQTL and mQTL results.

For cell-type-specific expression at this locus, seven genes (*PGAP3*, *ERBB2*, *MIEN1*, *IKZF3*, *GSDMB*, *ORMDL3* and *GSDMA*) were in naïve CD4^+^ T cells as previously also observed in tissue-resident memory ones or lymphoblastoid cell lines (LCLs).^12, 19, 20, 29^ In eosinophils, only one study has explored the gene expression.^17^ Thus, our findings show that six of the seven genes expressed by naïve CD4^+^ T cells were as well in eosinophils (excluding *GSDMA*), which significantly contributes to the research field. Our results also confirm the strong correlation observed between *GSDMB* and *ORMDL3* expression,^20^ independent of the asthma status. Regarding the methylation profiles, the difference at 3.2 kb upstream of the *IKZF3* TSS with blocks specific to naïve CD4^+^ T cells may be of interest since this gene is known to regulate the lymphocyte development^56^. Interestingly, none of the CpGs correlated in this region are included in the Infinium Human Methylation 450k BeadChip, and thus these results were obtain thanks to the custom capture panel used in this study.

To better understand the genetic interactions between SNPs, gene expression counts and DNA methylation levels, eQTL and mQTL analyses for each cell type were carried out. SNPs associated with the asthma phenotype in the SLSJ cohort were considered in order to identify QTLs, which are more likely to be involved in the disease. The two cell types exhibited different eQTL and mQTL patterns. Interestingly, both the expression and methylation levels of *GSDMA* in naïve CD4^+^ T cells were modulated by all asthma-associated SNPs (fourth and fifth haplotype blocks) and all measured mQTLs considering the 1.5 kb region from the TSS converged to two CpGs. The methylation profile at these two sites correlated with one another and both had a negative correlation with *GSDMA* expression counts; similar findings have been previously reported in LCLs.^24^ This gene is involved in pyroptosis, a process that induces cell death and the subsequent massive release of cellular contents and triggers strong inflammation.^57, 58^ Thus, the approach used in this study that combined various omic data on asthma phenotype pinpointed *GSDMA* as a new gene of interest in this locus in naïve CD4^+^ T cells.

The association between asthma and the 17q12-21 locus was first established in childhood asthma^6^ and was then confirmed to be mainly driven by individuals with the early-onset form of the disease.^13, 14^ Using a familial cohort with a mean age of onset of seven and 22 years old for probands and other family members respectively can be considered a caveat for the study of this locus, principally since gene expression counts and DNA methylation levels are changing over time. However, the genetic association was replicated in a previous study performed with the SLSJ asthma cohort^11^ and analyses using the imputed data in this study show similar results when taking account of all samples or only those with age of onset under 17 years old. Moreover, it was demonstrated that methylation marks can be transmitted to other generations.^59^ Analyses done on DNA methylation and mother-child duos in this study showed a marked increase of the correlation values between these levels in affected mother and affected child duos compared to the ones in the unaffected mother and affected child duos. Although the sample size was small in these stratified analyses, the above-mentioned increases strengthened the hypothesis that these two CpGs may be inherited and thus, the observation made in this study on *GSDMA* expression counts and methylation levels are of interest, even considering the age-specific association of this locus.

In this study, several observations previously described were reported; eQTLs between rs7216389, rs2290400 and rs4795405 as well as *ORMDL3* expression,^18, 19, 21, 60, 61^ mQTLs between rs7216389 and rs4065275 as well as methylation within the 1.5 kb region from *ORMDL3* TSS.^29^ Most interestingly, using the sequencing data for methylation and expression measures combined with the imputed genetic one provided new insights for the interactions in the 17q12-21 locus. This study design enabled the investigation of almost all potential genetic interactions with DNA methylation levels or gene expression counts in this locus for the two cell types. Only few studies have looked at all the genes in this locus for eQTL and even fewer ones for mQTL interactions.^12, 19^ The analyses considering all SNPs in the region also highlighted the important difference in the number of eQTLs and mQTLs observed in naïve CD4^+^ T cells compared to eosinophils, suggesting more complex interactions in the former type.

The investigation of asthma-associated SNPs in the eQTL and mQTL analyses allowed the detection of genetic interactions specific to this disease. Logically, its next step was to look for correlations, eQTLs and mQTLs in asthmatic and non-asthmatic individuals separately as an attempt to assess for a specific contribution of subjects suffering from the disease to the genetic interactions observed, focusing on the *GSDMA* gene. Unfortunately, it was not possible to conclude from the mQTLs data that the asthma status contributed to these interactions as p values resulting from permutations were not significant. However, it is interesting to note that the data in this study showed that eQTLs between SNPs located in the fourth and fifth haplotype blocks and *GSDMA* expression counts were linked to the asthma status of the individuals. In fact, 15 and 19 significant eQTLs were found in asthmatic individuals and controls respectively, showing a strong link between disease-associated SNPs and *GSDMA* expression counts in the two groups. More importantly, the pattern was opposite in both, the minor allele frequency is being lower in affected individuals for all SNPs and the expression being higher. This emphasizes the potential interest of this gene in the study of asthma.

To conclude, it will clearly be of interest to perform this approach on other immune cell types involved in asthma physiopathology to obtain a more complete picture. Nevertheless, by using sequencing data to measure DNA methylation level and gene expression counts on naïve CD4^+^ T cells and eosinophils from the same individuals (asthmatic and non-asthmatic), this study pinpointed *GSDMA* as a key new actor in the 17q12-21 locus and further confirmed the role of naïve CD4^+^ T cells in the pathogenesis of this disease.

## Supplemental Data

Supplemental Data include 6 figures and 13 tables.

## Acknowledgments

We wish to thank the participants recruited in the SLSJ asthma cohort for their valuable participation in this study. We also thank Dominique Fournier, scientific language expert, for the revision of this manuscript (http://www.serviceslinguistiquesdf.com/home). Anne-Marie Boucher-Lafleur, Andréanne Morin and Jolyane Meloche were supported by a *Fonds de recherche du Québec – Santé* (FRQS) Master, Doctoral or Postdoctoral training award respectively. Lucile Pain and Anne-Marie Boucher-Lafleur were supported by a Master or a Doctoral training award respectively from the Quebec Respiratory Health Network (RHN). Lucile Pain was also supported by the Natural Sciences and Engineering Graduate Scholarship from the Université du Québec à Chicoutimi Foundation (FUQAC). This project was supported by operating grants from the Canadian Institute of Health Research (CIHR; Laprise & Pastinen). Catherine Laprise is part of the Quebec Respiratory Health Network (RHN; https://rsr-qc.ca/en/), the investigator of CHILD Study, the director of the *Centre intersectoriel en santé durable de l’UQAC* and the chairholder of the Canada Research Chair in the Environment and Genetics of Respiratory Disorders and Allergies (http://www.chairs.gc.ca). The GWAS data were made available by the European Commission as part of GABRIEL (A multidisciplinary study to identify the genetic and environmental causes of asthma in the European Community) contract number 018996 under the Integrated Program LSH-2004-1.2.5-1 Post genomic approaches to understand the molecular basis of asthma aiming at a preventive or therapeutic control. This study makes use of data generated by the UK10K Consortium. A full list of the investigators who contributed to the generation of the data is available from www.UK10K.org. Funding for UK10K was provided by the Wellcome Trust under award WT091310.

## Web resources

OMIM, http://www.omim.org/

PLINK, https://www.cog-genomics.org/plink/1.9/

WISARD, http://statgen.snu.ac.kr/wisard/?act=aa_fam

