## Supplemental Methods and Figures for "*GSDMA* drives the most replicated association with asthma in naïve CD4^+^ T cells"

### Supplemental Data

### Subjects and Methods

#### Genetic data

GWAS data for the SLSJ asthma cohort were obtained from Illumina Human610-Quad BeadChip. After quality control assessment, data that followed these criteria were used for the imputation: minor allele frequency ≥1%, p value for Hardy-Weinberg equilibrium ≥1e-05, genotype and individual call rates ≥95%. The pre-phasing step was performed with the Shapeit2 software using the duoHMM method that combines estimated haplotypes with pedigree information to take advantage of known structure related to the latter.[1] Impute2 software was used for imputation[2] with the 1,000 Genome Project database (phase 3)[3] and the UK10K one as reference samples. Genotypes with a probability of 90% or more were kept after the imputation. Only imputed data that fulfilled the same criteria as the ones for GWAS were included in analyses (minor allele frequency ≥1%, p value for Hardy-Weinberg equilibrium ≥1e-05, genotype and individual call rates ≥95%). After quality filtering, association with asthma were evaluated for 762 SNPs of the chr17:37,826,875-38,134,519 region and for 1,180 DNA samples.

#### Isolation of naïve $CD4^+$ T cells and eosinophils

Naïve $CD4^+$ T cells and eosinophils were isolated from 200 ml blood samples from 215 individuals, a subset of the SLSJ asthma cohort, according to protocols described.[4, 5] The first step was to get rid of the platelet-rich plasma by centrifugation and to use dextran in order to remove the erythrocytes by sedimentation. A lymphocyte separation medium was chosen to withdraw mononuclear cells by density gradient. A positive selection was made on these using anti-CD45 and magnetic beads to retrieve naïve $CD4^+$ T cells (STEMCELL Technologies, Vancouver, BC, Canada). Counts were done with the Orflo Moxi Z Mini Automated Cell Counter (Thermo Fisher Scientific, Ottawa, ON, Canada). A proportion of cells (2e-06) was snap-frozen until DNA extraction and as much was suspended in Trizol for RNA's.

After separation of mononuclear cells and granulocytes, hypotonic lysis with water was performed to remove remaining erythrocytes from the latter type and negative selection with anti-CD16 MicroBeads and magnetic cell sorter allowed to isolate eosinophils (Miltenyi Biotec, Auburn, CA, USA). After monitoring for possible monocyte contamination using microscopy, all samples showing ≥ 90 % contamination were treated with another isolation step using anti-CD3 and anti-CD19 MicroBeads (Miltenyi Biotec, Auburn, CA, USA). The mean eosinophil purity was estimated to 95.5 %. Counts were done with the Orflo Moxi Z Mini Automated Cell Counter (Fisher Scientific, Ottawa, ON, Canada). A proportion of cells (1e-06) was snap-frozen until DNA extraction and the remaining (1e-06) was suspended in Trizol for RNA's.

DNA and RNA from both naïve $CD4^+$ T cells and eosinophils were extracted using the DNeasy Blood & Tissue Kit and the RNeasy Mini Kit respectively following the company's instructions (Qiagen, Toronto, ON, Canada).

**Targeted bisulfite sequencing (DNA methylation levels)**

Bisulfite sequencing was done for a custom methyl capture panel of identified functional immune genetic regions as previously described.[6, 7] These covered a total of 4,609,564 CpGs for a sum of 822,884 regions and 119,089,296 bp sequenced.[7] The MCC-Seq methylation was developed and optimized in collaboration with R&D at Roche NimbleGen. Libraries were prepared using the KAPA High Throughput Library Preparation Kit (Roche/KAPA Biosystems, Wilmington, MA, USA). Briefly,

1 ug of DNA was spiked with 0.1% (w/w) unmethylated λ and pUC19 DNA (Promega, Madison, WI, USA). DNA was sonicated (Covaris, Woburn, MA, USA) and fragments of 300–400 bp were controlled on a Bioanalyzer DNA 1000 Chip (Agilent, Santa Clara, CA, USA). Following fragmentation, end repair of double-stranded DNA breaks, 3'-end adenylation, adaptor ligation and clean-up steps were conducted according to KAPA Biosystems' protocols. The sample was then bisulfite converted using the EpiTect Fast Bisulfte Kit (Qiagen, Toronto, ON, Canada) following the manufacturer's instructions. The resulting bisulfite DNA was quantified with OliGreen (Life Technologies, Waltham, MA, USA) and amplified with 9–12 PCR cycles using the KAPA HiFi HotStart Uracil+ DNA Polymerase (Roche/KAPA Biosystems, Wilmington, MA, USA) according to suggested protocols. The final library was purified with AMPure beads that were validated on Bioanalyzer High Sensitivity DNA Chips (Agilent, Santa Clara, CA, USA) and quantified by PicoGreen (Thermo Fisher, Waltham, MA, USA). Then, the MCC-Seq protocol, developed and optimized by Roche NimbleGen, was applied to the following preparations for all individual samples (as described above). Briefly, the SeqCap Epi Enrichment System protocol (Roche NimbleGen, Wilmington, MA, USA) was used to capture the regions of interest. Equal quantities of multiplexed libraries (84 ng of each; 12 samples per capture) were combined to obtain 1 ug of total input and was hybridized to the capture panel at 47ºC for 72 h. Washing, recovery, PCR amplification of the captured libraries as well as final purification steps were conducted as recommended by the manufacturer. The DNA Chips were used to determine quality, concentration and size distribution of the final captured libraries. Six of these were sequenced over per lane of Illumina HiSeq 4000 System using 100 bp paired end sequencing.

Targeted bisulfite sequencing reads were aligned using the Epigenetics Pipeline available from the DRAGEN Bio-IT Platform (Edico Genomics/Illumina, San Diego, CA, USA). The latter provides highly optimized software, and hardware accelerated algorithms for secondary analysis of next generation sequencing. Data were processed on compute infrastructure available at the Center for Pediatric Genomic Medicine designed to support the needs of high throughput sequencing. Sequence reads were demultiplexed into FASTQ files with bcl2Fastq2 Conversion Software v2.19.1 from Illumina. They were then trimmed for quality (phred33 ≥20) as well as this manufacturer's adapters using Trimgalore v.0.4.2, a wrapper tool around Cutadapt[8] and FastQC. Reads were aligned to a bisulfite-converted reference genome with DRAGEN EP v2.6.3 or later in paired-end mode using the directional (Lister) methylation protocol presets; alignments were calculated for both Watson and Crick strands and the highest quality unique one is retained. A genome-wide methylation report was generated by DRAGEN to record counts of methylated and unmethylated cytosines where each is positioned in the genome. CpGs that were found to be overlapping with SNPs (dbSNP 137), the DAC Blacklisted Regions or Duke Excluded Regions (generated by the ENCODE project) were removed, CpGs having fewer than five reads were as well. Samples with more than 1.5 M CpGs and CpGs with data that concern more than 120 of these were kept for analyses. Targeted bisulfite sequencing data were available for 192 naïve $CD4^+$ T cell and 183 eosinophil samples. The methylation levels were calculated as total (forward and reverse) non-converted C reads over total (forward and reverse) ones.

**RNA sequencing (gene expression counts)**

RNA sequencing was performed at the McGill University and Génome Québec Innovation Centre. Total RNA was isolated from eosinophils and naïve $CD4^+$ T cells. As input, 500 ng RNA (of integrity number >7) was used for library preparations with the Illumina TruSeq Stranded Total RNA Sample Prep Kit according to the manufacturer's protocol (Illumina, San Diego, CA, USA). Final libraries were quality controlled on a Bioanalyzer and underwent 100 bp paired-end sequencing on the Illumina HiSeq2000 System. Generated raw reads were filtered for quality (phred33 ≥30) and length (n ≥32) as well as adapter sequences were removed using Trimmomatic v.0.32. Reads passing filters were then aligned to the human reference (hg19) with TopHat v.2.0.10 and Bowtie v.2.1.0. UCSC gene counts were obtained using HTSeq-count v.0.6.1 (http://htseq.readthedocs.io/). Transcriptomic data were available for 171 naïve $CD4^+$ T cell and 146 eosinophil samples.

### Figures

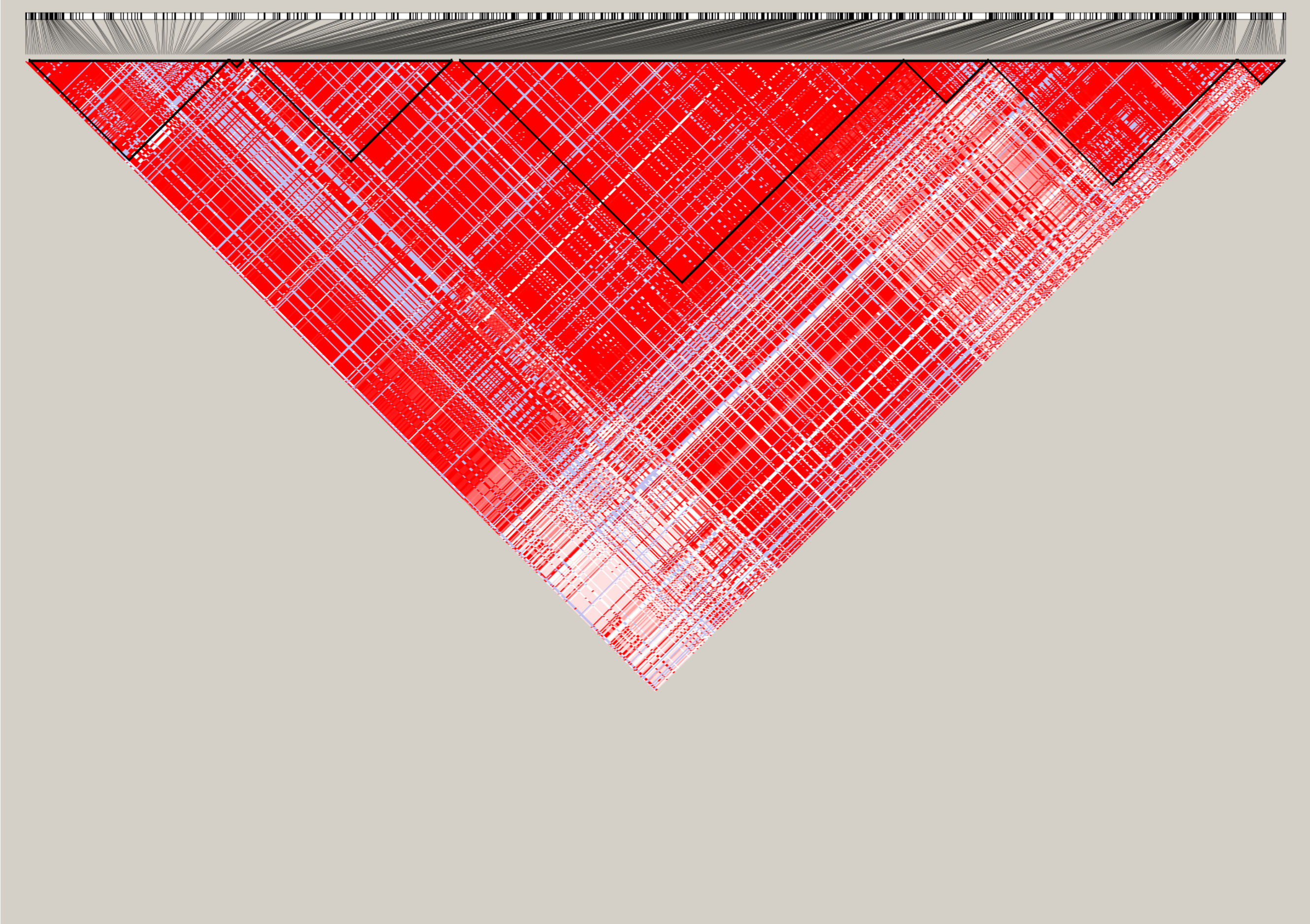

**Figure S1: Haplotype blocks identified in the Saguenay–Lac-Saint-Jean asthma cohort**

This figure shows the seven haplotype blocks identified in the Saguenay–Lac-Saint-Jean asthma cohort using the Haploview software.

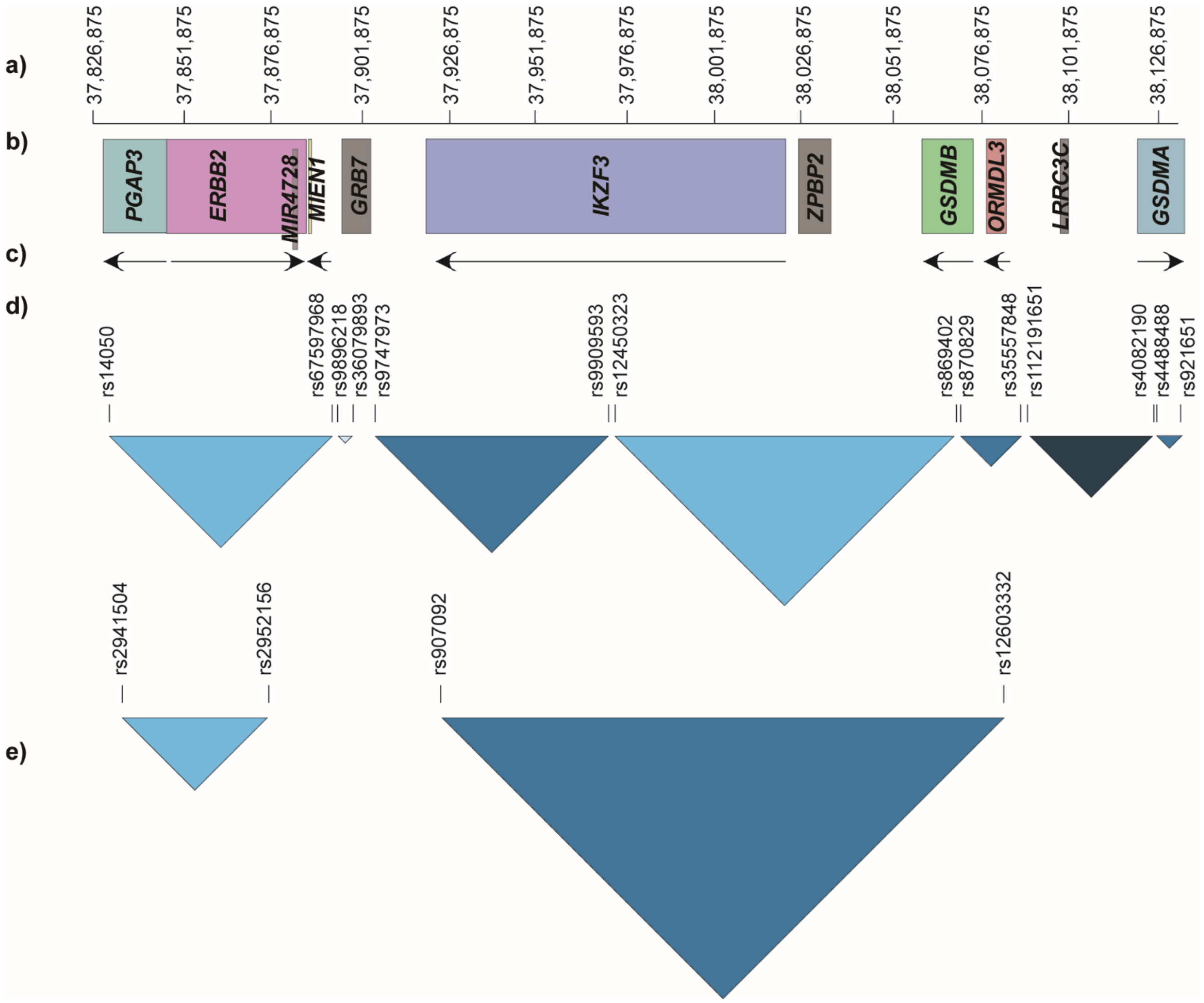


**Figure S2: Comparisons of the haplotype blocks identified in the Saguenay–Lac-Saint-Jean asthma cohort and those in the European population**

This figure shows: (a) the genetic position on the chromosome 17 in base pairs according to the human genome version 19; (b) the genes located in the chr17:37,826,875-38,134,519 region, and (c) the strand of these. The section (d) presents the haplotype blocks identified in the Saguenay–Lac-Saint-Jean asthma cohort with the first and last SNPs of each and in (e) the ones as published by Stein et al.[9] with also the first and last SNPs.

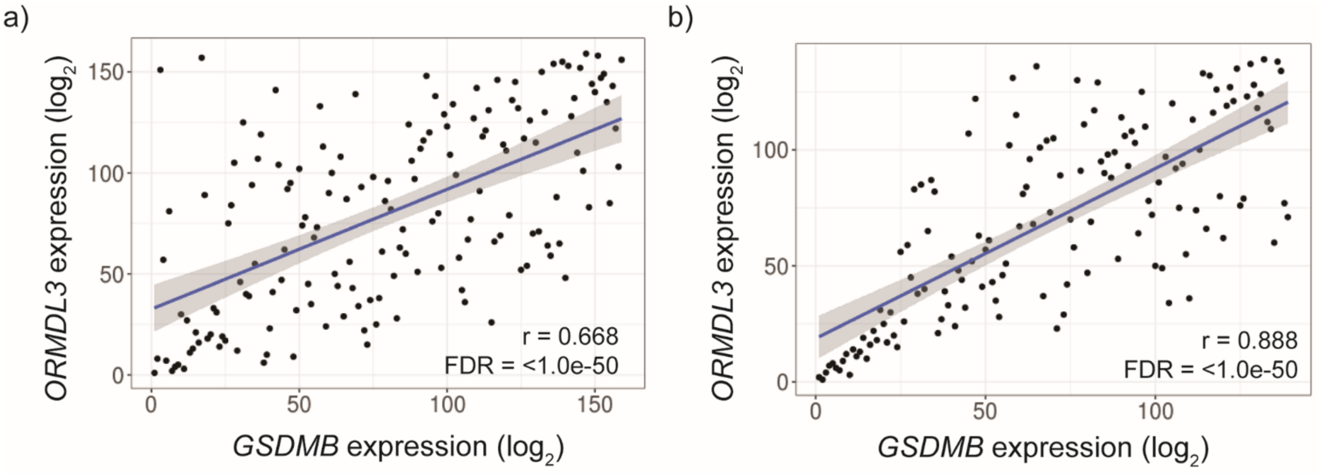


**Figure S3: Positive correlation between *GSDMB* and *ORMDL3* gene expression counts**

A positive correlation was observed between the gene expression of *ORMDL3* and *GSDMB* in (a) naïve $CD4^+$ T cells and (b) eosinophils. These data were normalized for library size and $log_2$-transformed.

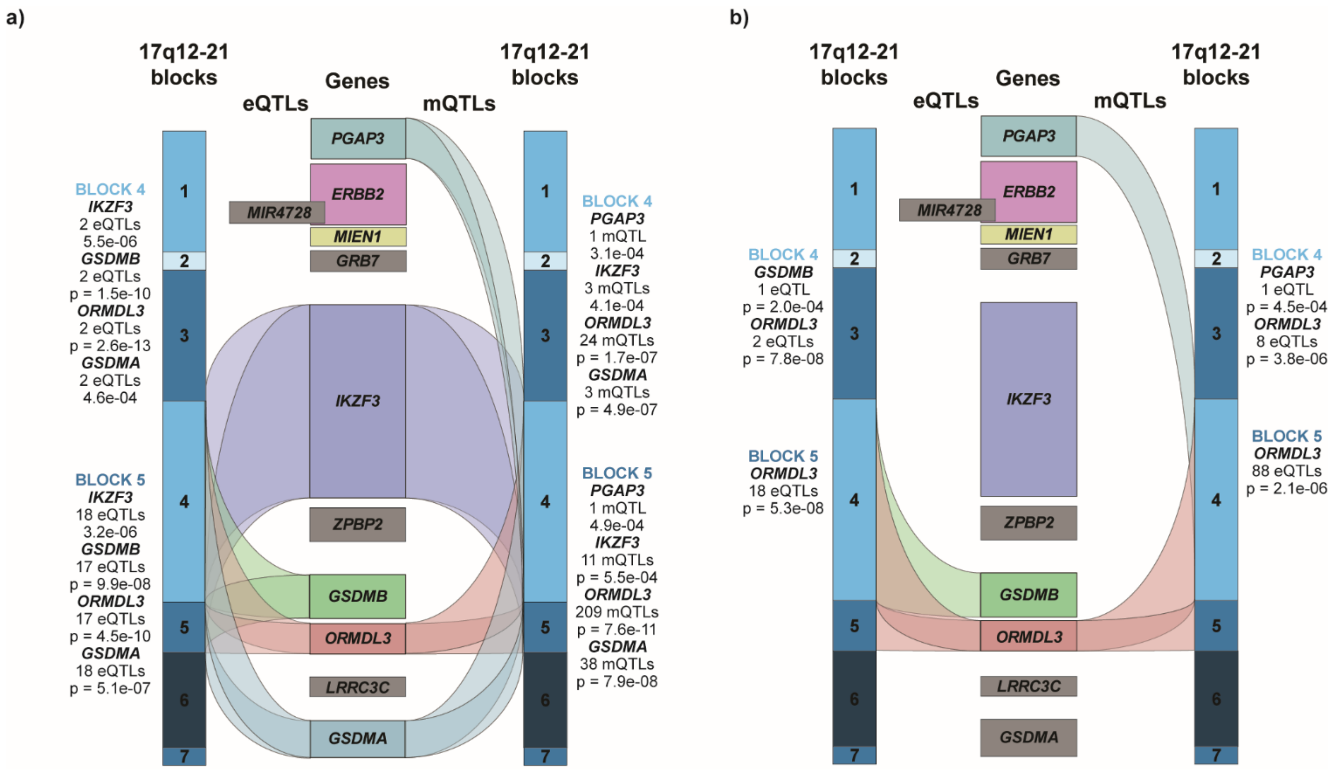


**Figure S4: eQTLs and mQTLs at the 17q12-21 locus for DNA methylation sites located in gene bodies and presented by cell type**

This figure illustrates the eQTLs and mQTLs considering SNPs associated with asthma identified in naïve CD4$^{+}$ T cells (a) and eosinophils (b) for methylation sites located in gene bodies. At the outer edges in different shades of blue are the seven haplotype blocks detected in the Saguenay–Lac-Saint-Jean asthma cohort for the chr17:37,826,875-38,134,519 region, and in the middle all the genes located in the same area. Genes colored in grey are those not expressed by each of the two cell types in this study. The number of eQTLs (left part of any figure) and mQTLs (right part) linked to each gene for each haplotype block are shown as indicated. P values for the most statistically significant QTLs are listed for each gene in each block.

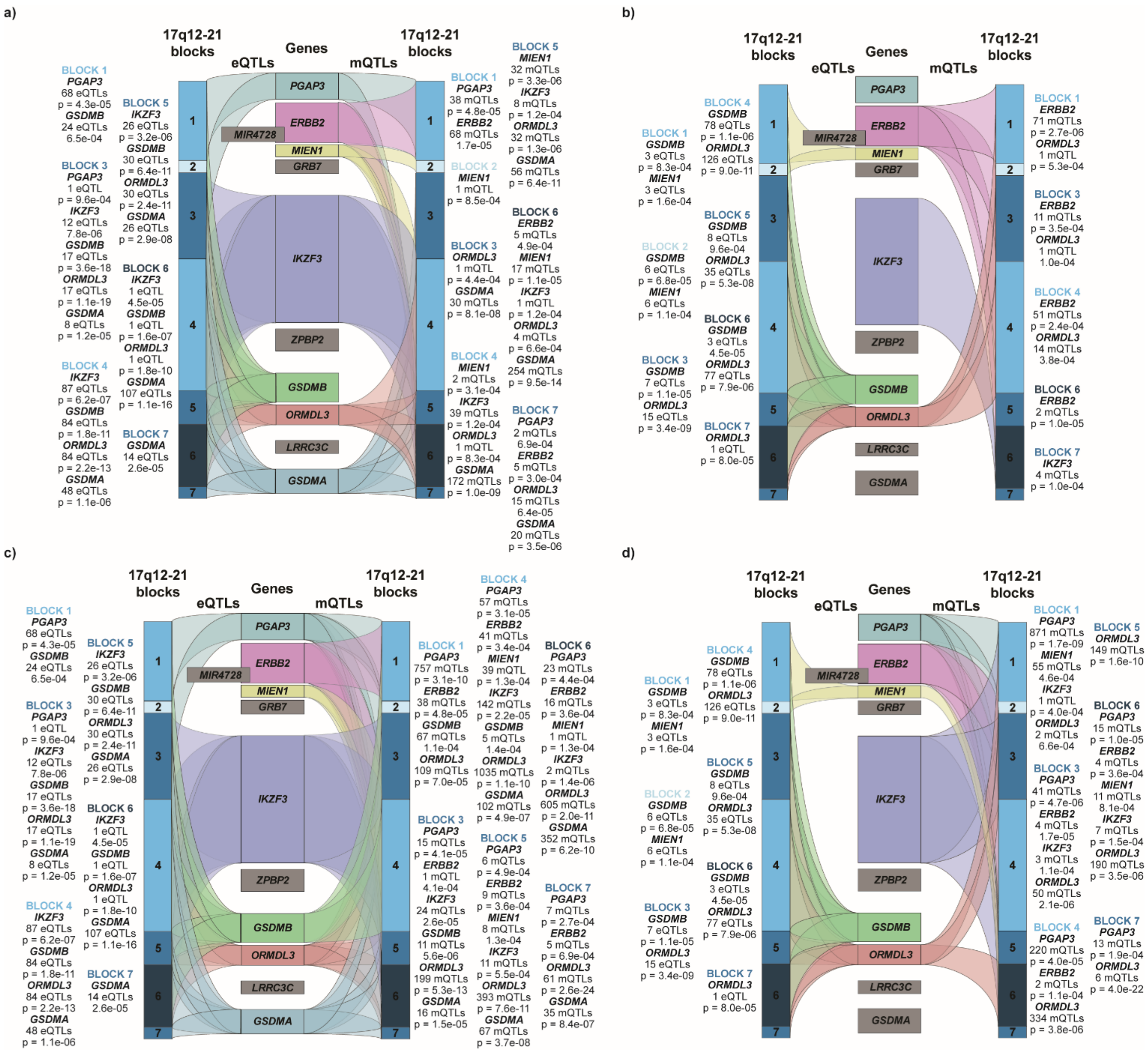


**Figure S5: eQTLs and mQTLs at the 17q12-21 locus including all SNPs analyzed and presented by cell types and location of methylation sites**

This figure illustrates the eQTLs and mQTLs identified considering all analyzed SNPs and methylation sites within 1.5 kb from TSS for (a) naïve CD4$^+$ T cells and (b) eosinophils and considering these located in gene bodies for (c) naïve CD4$^+$ T cells and (d) eosinophils. At the outer edges in different shades of blue are the seven haplotype blocks detected in the Saguenay–Lac-Saint-Jean asthma cohort for the chr17:37,826,875-38,134,519 region and in the middle all the genes located in the same area. Genes colored in grey are those not expressed by each of the two cell types in this study. The number of eQTLs (left part of any figure) and mQTLs (right part) linked to each gene for each haplotype block are shown as indicated. P values for the most statistically significant QTLs are listed for each gene in each block.

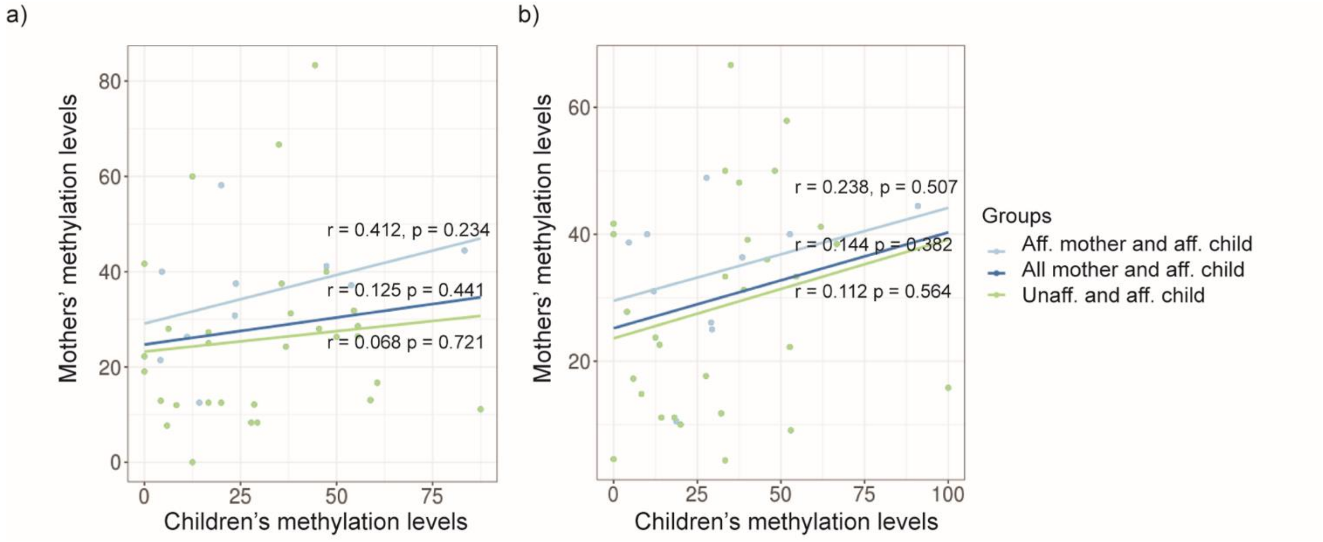


**Figure S6. Correlations between methylation sites in the 1.5 kb region from *GSDMA* TSS in mother-child duos**

This figure shows correlation values for the CpGs at (a) 17:38,119,198 and (b) 17:38,119,207 in the case of three mother-child duos. The green dots and line refer to the unaffected mother and affected child duos, the light blue dots and line refer to the affected mother and affected child duos and the dark blue line refers to all mother-child duos. Pearson's r correlation values and p values are indicated for each. Abbreviations: aff. = affected.
