## Supplemental Table 1 for "*GSDMA* drives the most replicated association with asthma in naïve CD4^+^ T cells"

**Table S1:** Significant associations with asthma

| Block | SNP id[a] | HGVS names | $r^{2}$ [b] | Chromosome | Minor allele | Asthma | | Asthma with age of onset <17 years old | | Previous associations[c] |
|---|---|---|---|---|---|---|---|---|---|---|
| | | | | | | Stat | P value | Stat | P value | |
| Fourth block | rs36095411 | g.38031865.G>T | 0.776 | 17 | G | 11.488 | 7.00E-04 | 11.085 | 8.71E-04 | |
| | rs869402 | g.38068043.T>C | 1.000 | 17 | T | 11.989 | 5.35E-04 | 14.101 | 1.73E-04 | |
| Fifth block | rs870829 | g.38068382.T>G | 0.821 | 17 | A | 11.428 | 7.23E-04 | 13.413 | 2.50E-04 | |
| | rs1011082 | g.38068514.A>G | 1.000 | 17 | T | 11.323 | 7.66E-04 | 12.808 | 3.45E-04 | |
| | rs921650 | g.38069076.C>T | 1.000 | 17 | G | 11.994 | 5.34E-04 | 13.551 | 2.32E-04 | |
| | rs921649 | g.38069274.G>A | 1.000 | 17 | C | 11.994 | 5.34E-04 | 13.551 | 2.32E-04 | |
| | rs5820308 | g.38069364_38069369del | 0.830 | 17 | TCAAAA | 11.212 | 8.13E-04 | 11.441 | 7.19E-04 | |
| | rs6503524 | g.38069809.C>T | 1.000 | 17 | C | 11.994 | 5.34E-04 | 13.551 | 2.32E-04 | |
| | rs7216389 | g.38069949.C>T | 1.000 | 17 | C | 11.994 | 5.34E-04 | 13.551 | 2.32E-04 | 1, 2 |
| | rs7216558 | g.38070071.C>T | 1.000 | 17 | C | 11.994 | 5.34E-04 | 13.551 | 2.32E-04 | 3 |
| | rs9303279 | g.38073968.G>C | 0.830 | 17 | G | 11.736 | 6.13E-04 | 11.079 | 8.73E-04 | |
| | rs9303280 | g.38074031.T>C | 0.958 | 17 | T | 11.130 | 8.49E-04 | 10.624 | NS | |
| | rs9303281 | g.38074046.G>A | 1.000 | 17 | G | 13.335 | 2.60E-04 | 13.821 | 2.01E-04 | 2 |
| | rs7219923 | g.38074518.C>T | 1.000 | 17 | C | 13.236 | 2.75E-04 | 13.761 | 2.08E-04 | 2 |
| | rs7224129 | g.38075426.G>A | 1.000 | 17 | G | 12.799 | 3.47E-04 | 12.973 | 3.16E-04 | 4 |
| | rs8074437 | g.38076137.T>G | 1.000 | 17 | T | 12.799 | 3.47E-04 | 12.973 | 3.16E-04 | |
| | rs71971950 | g.38076198_38076201del | 1.000 | 17 | TATA | 12.643 | 3.77E-04 | 12.635 | 3.79E-04 | |
| | rs4065275 | g.38080865.A>G | 0.946 | 17 | A | 11.197 | 8.19E-04 | 11.513 | 6.91E-04 | 5 |
| | rs8076131 | g.38080912.G>A | 0.822 | 17 | G | 12.067 | 5.13E-04 | 11.196 | 8.20E-04 | 2, 5 |
| | rs12603332 | g.38082807.T>C | 0.946 | 17 | T | 12.427 | 4.23E-04 | 12.552 | 3.96E-04 | 5 |

In grey: Best association for each haplotype block.

[a]SNPs shown are those with significant results (p <0.001) in the whole sample. [b]$r^2$ values between each SNP and the most associated SNP of the same haplotype block. [c]References for first association with asthma and/or association in the previous analysis performed in the SLSJ cohort:

1: Moffatt MF, Gut IG, Demenais F, Strachan DP, Bouzigon E, Heath S, et al. A large-scale, consortium-based genomewide association study of asthma. N Engl J Med 2010;363:1211–21. [PubMed: 20860503]

2: Madore AM, Tremblay K, Hudson TJ, Laprise C. Replication of an association between 17q21 SNPs and asthma in a French-Canadian familial collection. Hum Genet 2008;123:93–95. [PubMed: 17992541]

3: Shahid M, Sabar MF, Bano I, Rahman Z, Iqbal Z, Fatim Ali SS, Ghani MU, Iqbal M, Husnain T. Sequence variants on 17q21 are associated with the susceptibility of asthma in the population of Lahore, Pakistan. J Asthma. 2015;52:777-84. [PubMed: 26203825]

4: Tomita K, Sakashita M, Hirota T, Tanaka S, Masuyama K, Yamada T, Fujieda S, Miyatake A, Hizawa N, Kubo M, Nakamura Y, Tamari M. Variants in the 17q21 asthma susceptibility locus are associated with allergic rhinitis in the Japanese population. Allergy. 2013;68:92-100. [PubMed: 23157251]

5: Galanter J, Choudhry S, Eng C, Nazario S, Rodríguez-Santana JR, Casal J, Torres-Palacios A, Salas J, Chapela R, Watson HG, Meade K, LeNoir M, Rodríguez-Cintrón W, Avila PC, Burchard EG. ORMDL3 gene is associated with asthma in three ethnically diverse populations. Am J Respir Crit Care Med. 2008;177:1194-200. [PubMed: 18310477]
