## Supplemental Table 2 for "*GSDMA* drives the most replicated association with asthma in naïve CD4^+^ T cells"

**Table S2:** Difference of gene expression between samples from naïve CD4$^+$ T cells and eosinophils

| Gene | Mean naïve CD4$^+$ T cells | SD naïve CD4$^+$ T cells | Mean eosinophils | SD eosinophils | Fold change[a] | Pearson's r | zStat | P value | FDR |
|---|---|---|---|---|---|---|---|---|---|
| *PGAP3* | 543.098 | 101.092 | 134.286 | 35.015 | 4.044 | 0.124 | 1.372 | 0.173 | 0.432 |
| *ERBB2* | 113.571 | 67.750 | 54.958 | 51.890 | 2.067 | 0.001 | 0.011 | 0.991 | 0.991 |
| *MIEN1* | 212.650 | 65.444 | 209.044 | 58.535 | 1.017 | 0.221 | 2.477 | 0.015 | 0.073 |
| *IKZF3* | 1168.354 | 281.245 | 254.232 | 108.001 | 4.596 | 0.222 | 2.491 | 0.014 | 0.073 |
| *GSDMB* | 602.887 | 160.442 | 623.941 | 181.840 | 0.966 | -0.008 | -0.082 | 0.935 | 0.991 |
| *ORMDL3* | 2235.256 | 429.382 | 1637.832 | 404.431 | 1.365 | 0.023 | 0.256 | 0.799 | 0.991 |

[a]A positive fold change refers to an expression that is increased in naïve CD4$^+$ T cells compared to eosinophils and a negative fold change refers to an expression rather increased in eosinophils.
