## Supplemental Table 3 for "*GSDMA* drives the most replicated association with asthma in naïve CD4^+^ T cells"

**Table S3:** Correlations between gene expression counts in naïve CD4$^+$ T cells

| | *PGAP3* | *ERBB2* | *MIEN1* | *IKZF3* | *GSDMB* | *ORMDL3* | *GSDMA* |
|---|---|---|---|---|---|---|---|
| *PGAP3* | | 0.082 | 0.206 | 0.165 | *0.007* | 0.526 | 0.916 |
| *ERBB2* | -0.173 | | *0.003* | *0.030* | *0.007* | 0.200 | 0.082 |
| *MIEN1* | 0.128 | *0.278* | | *0.013* | *2.82E-12* | 0.515 | 0.547 |
| *IKZF3* | 0.140 | *0.212* | *-0.236* | | *6.88E-05* | 0.515 | 0.165 |
| *GSDMB* | *0.253* | *-0.260* | *-0.530* | *0.346* | | *< 1.00E-25* | 0.874 |
| *ORMDL3* | 0.065 | -0.130 | 0.074 | 0.074 | *0.668* | | 0.218 |
| *GSDMA* | 0.011 | 0.176 | 0.060 | 0.140 | -0.020 | 0.124 | |

Cells in grey include FDR values and the other ones the corresponding correlation r values. In italic, they are significant (FDR <0.05).
