## Supplemental Table 4 for "*GSDMA* drives the most replicated association with asthma in naïve CD4^+^ T cells"

**Table S4:** Correlations between gene expression counts in eosinophils

| | *PGAP3* | *ERBB2* | *MIEN1* | *IKZF3* | *GSDMB* | *ORMDL3* |
|---|---|---|---|---|---|---|
| *PGAP3* | | *0.001* | *0.044* | *1.23E-09* | *2.47E-05* | *2.16E-08* |
| *ERBB2* | *0.306* | | 0.642 | *0.021* | 0.365 | 0.642 |
| *MIEN1* | *-0.214* | -0.060 | | *0.002* | 0.056 | 0.208 |
| *IKZF3* | *0.506* | *0.237* | *-0.296* | | 0.642 | 0.726 |
| *GSDMB* | *0.375* | -0.104 | -0.204 | 0.058 | | *< 1.00E-25* |
| *ORMDL3* | *0.473* | -0.063 | -0.140 | 0.048 | *0.888* | |

Cells in grey include FDR values and the other ones the corresponding correlation r values. In italic, they are significant (FDR <0.05).
