## Supplemental Table 5 for "*GSDMA* drives the most replicated association with asthma in naïve CD4^+^ T cells"

**Table S5:** Associations between gene expression levels and asthma in naïve $CD4^+$ T cells and eosinophils

| Gene | Mean Aff | SD Aff | Mean Unaff | SD Unaff | Fold change[a] | Coeff | SE | P value | FDR |
|---|---|---|---|---|---|---|---|---|---|
| Naïve CD4+ T cells | | | | | | | | | |
| *PGAP3* | 534.045 | 90.892 | 553.330 | 94.166 | -1.036 | -0.027 | 0.020 | 0.188 | 0.988 |
| *ERBB2* | 112.654 | 47.032 | 110.610 | 84.345 | 1.018 | 0.108 | 0.055 | 0.051 | 0.561 |
| *MIEN1* | 211.220 | 71.878 | 212.302 | 44.352 | -1.005 | 0.007 | 0.031 | 0.809 | 0.988 |
| *IKZF3* | 1150.322 | 251.383 | 1179.409 | 260.505 | -1.025 | 0.021 | 0.030 | 0.497 | 0.988 |
| *GSDMB* | 598.076 | 147.693 | 607.352 | 159.215 | -1.016 | 0.001 | 0.038 | 0.970 | 0.988 |
| *ORMDL3* | 2216.841 | 384.674 | 2240.564 | 398.983 | -1.011 | -0.001 | 0.032 | 0.984 | 0.988 |
| *GSDMA* | 10.843 | 7.914 | 14.392 | 15.559 | -1.327 | 0.086 | 0.125 | 0.493 | 0.988 |
| Eosinophils | | | | | | | | | |
| *PGAP3* | 135.209 | 33.764 | 133.318 | 34.418 | 1.014 | 0.010 | 0.044 | 0.819 | 0.829 |
| *ERBB2* | 53.478 | 51.652 | 59.841 | 54.897 | -1.119 | -0.076 | 0.118 | 0.518 | 0.699 |
| *MIEN1* | 213.033 | 56.832 | 200.906 | 55.188 | 1.060 | 0.056 | 0.049 | 0.250 | 0.652 |
| *IKZF3* | 247.970 | 102.278 | 268.637 | 115.629 | -1.083 | -0.056 | 0.074 | 0.453 | 0.699 |
| *GSDMB* | 636.540 | 172.374 | 600.112 | 185.125 | 1.061 | 0.039 | 0.039 | 0.326 | 0.652 |
| *ORMDL3* | 1679.690 | 397.278 | 1560.377 | 365.119 | 1.076 | 0.053 | 0.031 | 0.087 | 0.526 |

[a]Positive fold change indicate an increase of gene expression in affected individuals compared to controls and negative fold change rather indicates a decrease in gene expression in affected individuals.

Abbreviations: Aff = samples from individuals affected with asthma, SD = standard deviation, SE = standard error, Unaff = samples from unaffected individuals.
