## Supplemental Table 6 for "*GSDMA* drives the most replicated association with asthma in naïve CD4^+^ T cells"

**Table S6:** Significant associations between CpG methylation levels and asthma in naïve $CD4^+$ T cells and eosinophils

| Position CpG (hg19) | Gene is methylated (1.5 kb from TSS)[a] | Gene is methylated (gene body)[a] | Mean Aff | SD Aff | Mean Unaff | SD Unaff | % difference[b] | Coeff | SE | P value | FDR |
|---|---|---|---|---|---|---|---|---|---|---|---|
| Naïve $CD4^+$ T cells | | | | | | | | | | | |
| 37,828,273 | | *PGAP3* | 38.8 | 13.6 | 42.0 | 13.9 | -3.2 | -0.174 | 0.056 | 0.002 | 0.010 |
| 37,828,296 | | *PGAP3* | 56.3 | 13.5 | 58.2 | 16.3 | -1.9 | -0.181 | 0.051 | 3.69E-04 | 0.002 |
| 37,831,276 | | *PGAP3* | 90.1 | 10.5 | 83.0 | 17.9 | 7.1 | 0.397 | 0.143 | 0.006 | 0.028 |
| 37,834,687 | | *PGAP3* | 95.4 | 4.5 | 96.4 | 5.1 | -1.0 | -0.488 | 0.169 | 0.004 | 0.020 |
| 37,836,263 | | *PGAP3* | 25.9 | 15.2 | 23.9 | 11.8 | 2.0 | 0.252 | 0.086 | 0.004 | 0.019 |
| 37,843,621 | *ERBB2* | *PGAP3* | 79.4 | 11.6 | 83.3 | 10.8 | -3.9 | -0.354 | 0.081 | 1.17E-05 | 7.82E-05 |
| 37,845,218 | *PGAP3* | *ERBB2* | 81.7 | 13.8 | 84.4 | 13.2 | -2.7 | -0.404 | 0.145 | 0.005 | 0.027 |
| 37,856,217 | | *ERBB2* | 19.4 | 12.4 | 26.0 | 13.5 | -6.6 | -0.255 | 0.085 | 0.003 | 0.015 |
| 37,856,364 | | *ERBB2* | 2.9 | 4.6 | 4.6 | 7.5 | -1.7 | -0.691 | 0.232 | 0.003 | 0.015 |
| 37,856,396 | | *ERBB2* | 3.5 | 6.9 | 4.5 | 7.2 | -1.0 | -0.969 | 0.297 | 0.001 | 0.006 |
| 37,856,663 | | *ERBB2* | 5.1 | 6.2 | 7.1 | 9.5 | -2.0 | -0.419 | 0.156 | 0.007 | 0.035 |
| 37,856,694 | | *ERBB2* | 4.6 | 4.9 | 7.1 | 7.0 | -2.5 | -0.404 | 0.129 | 0.002 | 0.009 |
| 37,856,884 | | *ERBB2* | 7.2 | 8.6 | 8.9 | 8.6 | -1.7 | -0.491 | 0.142 | 0.001 | 0.003 |
| 37,857,834 | | *ERBB2* | 79.0 | 10.6 | 77.6 | 9.8 | 1.4 | 0.169 | 0.064 | 0.008 | 0.040 |
| 37,860,411 | | *ERBB2* | 97.4 | 4.8 | 97.3 | 5.3 | 0.1 | -1.240 | 0.435 | 0.004 | 0.023 |
| 37,862,101 | | *ERBB2* | 94.2 | 6.1 | 95.6 | 7.0 | -1.4 | -0.523 | 0.167 | 0.002 | 0.010 |
| 37,862,292 | | *ERBB2* | 92.0 | 11.5 | 93.6 | 9.6 | -1.6 | -0.842 | 0.237 | 3.83E-04 | 0.002 |
| 37,893,978 | | | 11.5 | 15.6 | 5.9 | 9.2 | 5.6 | 0.942 | 0.310 | 0.002 | 0.013 |
| 37,894,826 | | | 5.9 | 9.6 | 7.4 | 9.8 | -1.5 | -1.109 | 0.328 | 0.001 | 0.004 |
| 37,895,157 | | | 13.2 | 13.2 | 11.7 | 11.7 | 1.5 | 0.488 | 0.148 | 0.001 | 0.006 |
| 37,895,529 | | | 74.3 | 16.2 | 74.9 | 17.0 | -0.6 | 0.300 | 0.103 | 0.004 | 0.019 |
| 37,895,642 | | | 42.5 | 16.9 | 43.3 | 16.8 | -0.8 | 0.204 | 0.073 | 0.005 | 0.027 |
| 37,896,063 | | | 62.3 | 17.1 | 67.6 | 15.4 | -5.3 | -0.261 | 0.082 | 0.001 | 0.009 |
| 37,896,165 | | | 47.5 | 18.5 | 44.9 | 14.3 | 2.6 | 0.252 | 0.084 | 0.003 | 0.014 |
| 37,896,203 | | | 71.5 | 18.8 | 70.8 | 17.7 | 0.7 | 0.366 | 0.118 | 0.002 | 0.011 |
| 37,896,363 | | | 83.0 | 14.9 | 86.9 | 12.7 | -3.9 | -0.497 | 0.155 | 0.001 | 0.008 |

| | | | | | | | | | | | |
|---|---|---|---|---|---|---|---|---|---|---|---|
| 37,907,944 | | | 87.4 | 10.0 | 89.6 | 8.8 | -2.2 | -0.262 | 0.101 | 0.009 | 0.043 |
| 37,910,316 | | | 0.2 | 1.0 | 0.5 | 1.6 | -0.3 | -2.148 | 0.709 | 0.002 | 0.013 |
| 37,910,688 | | | 0.3 | 1.1 | 1.1 | 3.0 | -0.8 | -1.118 | 0.416 | 0.007 | 0.035 |
| 37,911,929 | | | 23.4 | 13.0 | 25.8 | 13.6 | -2.4 | -0.200 | 0.074 | 0.007 | 0.033 |
| 37,913,515 | | | 76.2 | 15.6 | 81.8 | 11.7 | -5.6 | -0.515 | 0.113 | 4.96E-06 | 3.32E-05 |
| 37,914,977 | | | 26.2 | 12.0 | 25.0 | 9.6 | 1.2 | 0.227 | 0.060 | 1.65E-04 | 0.001 |
| 37,922,412 | | *IKZF3* | 95.2 | 7.5 | 93.4 | 6.6 | 1.8 | 0.558 | 0.148 | 1.61E-04 | 0.001 |
| 37,929,753 | | *IKZF3* | 7.2 | 11.1 | 6.6 | 9.9 | 0.6 | -0.508 | 0.182 | 0.005 | 0.027 |
| 37,939,157 | | *IKZF3* | 89.6 | 14.2 | 84.7 | 19.3 | 4.9 | 1.168 | 0.328 | 3.66E-04 | 0.002 |
| 37,949,286 | | *IKZF3* | 16.1 | 12.1 | 16.2 | 9.9 | -0.1 | -0.311 | 0.076 | 4.55E-05 | 3.00E-04 |
| 37,949,345 | | *IKZF3* | 25.6 | 13.0 | 25.6 | 12.3 | 0.0 | -0.240 | 0.069 | 4.71E-04 | 0.003 |
| 37,974,812 | | *IKZF3* | 64.8 | 21.4 | 64.6 | 23.7 | 0.2 | -0.954 | 0.256 | 1.88E-04 | 0.001 |
| 37,975,322 | | *IKZF3* | 92.1 | 10.6 | 95.4 | 7.9 | -3.3 | -0.604 | 0.228 | 0.008 | 0.039 |
| 37,999,368 | | *IKZF3* | 13.0 | 13.1 | 11.4 | 13.0 | 1.6 | 0.498 | 0.129 | 1.20E-04 | 0.001 |
| 37,999,393 | | *IKZF3* | 14.2 | 13.8 | 12.8 | 13.2 | 1.4 | 0.373 | 0.122 | 0.002 | 0.012 |
| 38,004,518 | | *IKZF3* | 49.4 | 16.0 | 53.6 | 17.9 | -4.2 | -0.235 | 0.091 | 0.010 | 0.046 |
| 38,020,407 | | *IKZF3* | 0.4 | 1.3 | 0.9 | 2.2 | -0.5 | -1.066 | 0.342 | 0.002 | 0.010 |
| 38,020,442 | *IKZF3* | | 0.5 | 1.2 | 0.7 | 3.7 | -0.2 | 1.391 | 0.420 | 0.001 | 0.005 |
| 38,020,544 | *IKZF3* | | 0.3 | 1.4 | 0.5 | 1.2 | -0.2 | -1.559 | 0.529 | 0.003 | 0.017 |
| 38,022,345 | | | 64.4 | 18.9 | 66.3 | 17.5 | -1.9 | -0.399 | 0.116 | 0.001 | 0.003 |
| 38,023,679 | | | 67.6 | 13.9 | 66.4 | 14.7 | 1.2 | -0.165 | 0.049 | 0.001 | 0.004 |
| 38,024,174 | | | 47.8 | 19.9 | 42.4 | 23.4 | 5.4 | 0.305 | 0.103 | 0.003 | 0.016 |
| 38,024,285 | | | 45.6 | 19.8 | 39.6 | 16.5 | 6.0 | 0.270 | 0.081 | 0.001 | 0.005 |
| 38,024,290 | | | 45.2 | 21.0 | 38.1 | 15.8 | 7.1 | 0.237 | 0.082 | 0.004 | 0.019 |
| 38,024,317 | | | 20.8 | 16.3 | 15.2 | 12.0 | 5.6 | 0.295 | 0.104 | 0.004 | 0.023 |
| 38,024,331 | | | 33.7 | 18.7 | 28.4 | 15.7 | 5.3 | 0.249 | 0.085 | 0.003 | 0.017 |
| 38,024,375 | | | 22.7 | 16.1 | 19.2 | 13.9 | 3.5 | 0.389 | 0.098 | 7.74E-05 | 0.001 |
| 38,024,390 | | | 16.8 | 14.0 | 14.8 | 14.3 | 2.0 | 0.331 | 0.119 | 0.005 | 0.027 |
| 38,024,394 | | | 21.7 | 15.8 | 17.5 | 15.2 | 4.2 | 0.425 | 0.106 | 5.71E-05 | 3.75E-04 |
| 38,024,456 | | | 33.3 | 20.0 | 31.0 | 18.6 | 2.3 | 0.374 | 0.103 | 3.03E-04 | 0.002 |
| 38,024,458 | | | 32.2 | 17.8 | 30.3 | 19.2 | 1.9 | 0.341 | 0.105 | 0.001 | 0.007 |
| 38,024,462 | | | 41.5 | 21.2 | 36.9 | 17.5 | 4.6 | 0.480 | 0.100 | 1.65E-06 | 1.11E-05 |
| 38,024,468 | | | 28.5 | 18.9 | 27.6 | 17.1 | 0.9 | 0.316 | 0.113 | 0.005 | 0.026 |
| 38,024,471 | | | 25.3 | 17.7 | 22.3 | 18.4 | 3.0 | 0.345 | 0.121 | 0.004 | 0.022 |

| | | | | | | | | | | | |
|---|---|---|---|---|---|---|---|---|---|---|---|
| 38,024,479 | | | 33.7 | 21.3 | 31.1 | 17.8 | 2.6 | 0.407 | 0.114 | 3.44E-04 | 0.002 |
| 38,024,496 | | | 31.8 | 22.5 | 28.1 | 20.5 | 3.7 | 0.592 | 0.121 | 9.22E-07 | 6.24E-06 |
| 38,024,502 | | | 35.3 | 20.5 | 33.6 | 18.0 | 1.7 | 0.360 | 0.107 | 0.001 | 0.005 |
| 38,024,515 | | | 63.2 | 20.9 | 62.5 | 19.8 | 0.7 | 0.296 | 0.114 | 0.010 | 0.046 |
| 38,074,066 | | *GSDMB* | 94.5 | 6.9 | 96.3 | 5.2 | -1.8 | -0.504 | 0.187 | 0.007 | 0.034 |
| 38,077,227 | | | 7.2 | 8.2 | 9.4 | 10.9 | -2.2 | -0.452 | 0.150 | 0.003 | 0.014 |
| 38,081,186 | | *ORMDL3* | 67.0 | 15.8 | 67.6 | 15.7 | -0.6 | -0.308 | 0.074 | 3.03E-05 | 2.01E-04 |
| 38,082,018 | | *ORMDL3* | 19.3 | 16.6 | 22.5 | 18.0 | -3.2 | -0.262 | 0.067 | 9.84E-05 | 0.001 |
| 38,082,206 | | *ORMDL3* | 15.4 | 13.9 | 19.0 | 16.7 | -3.6 | -0.289 | 0.085 | 0.001 | 0.004 |
| 38,082,346 | | *ORMDL3* | 30.6 | 17.1 | 28.3 | 16.5 | 2.3 | -0.207 | 0.067 | 0.002 | 0.011 |
| 38,084,037 | *ORMDL3* | | 0.4 | 1.4 | 0.9 | 2.2 | -0.5 | -1.848 | 0.550 | 0.001 | 0.005 |
| 38,085,045 | *ORMDL3* | | 84.2 | 12.4 | 83.1 | 12.7 | 1.1 | 0.270 | 0.097 | 0.006 | 0.028 |
| 38,085,835 | | | 85.5 | 10.0 | 79.8 | 15.3 | 5.7 | 0.339 | 0.092 | 2.23E-04 | 0.001 |
| 38,095,691 | | | 90.8 | 8.5 | 87.0 | 11.7 | 3.8 | 0.419 | 0.154 | 0.007 | 0.033 |
| 38,104,883 | | | 24.3 | 16.4 | 32.6 | 20.4 | -8.3 | -0.337 | 0.104 | 0.001 | 0.007 |
| 38,108,710 | | | 45.5 | 17.0 | 43.0 | 16.9 | 2.5 | 0.187 | 0.071 | 0.008 | 0.039 |
| 38,108,805 | | | 37.2 | 13.9 | 39.5 | 12.8 | -2.3 | -0.185 | 0.069 | 0.007 | 0.036 |
| 38,108,868 | | | 22.1 | 11.4 | 24.6 | 11.8 | -2.5 | -0.214 | 0.071 | 0.003 | 0.014 |
| 38,109,003 | | | 9.7 | 7.7 | 13.2 | 11.6 | -3.5 | -0.356 | 0.098 | 2.67E-04 | 0.002 |
| 38,109,870 | | | 73.4 | 12.0 | 76.2 | 11.2 | -2.8 | -0.199 | 0.075 | 0.008 | 0.038 |
| 38,115,399 | | | 0.6 | 2.0 | 0.3 | 1.4 | 0.3 | 1.749 | 0.555 | 0.002 | 0.009 |
| 38,119,649 | | *GSDMA* | 27.4 | 17.6 | 31.0 | 16.4 | -3.6 | -0.274 | 0.076 | 3.21E-04 | 0.002 |
| 38,119,760 | | *GSDMA* | 56.6 | 14.9 | 61.2 | 14.5 | -4.6 | -0.197 | 0.064 | 0.002 | 0.011 |
| 38,123,724 | | *GSDMA* | 78.8 | 10.9 | 74.7 | 13.0 | 4.1 | 0.281 | 0.086 | 0.001 | 0.006 |
| 38,133,185 | | *GSDMA* | 88.1 | 11.4 | 84.4 | 12.2 | 3.7 | 0.351 | 0.132 | 0.008 | 0.038 |
| 38,133,214 | | *GSDMA* | 90.4 | 11.2 | 91.6 | 8.7 | -1.2 | -0.426 | 0.160 | 0.008 | 0.038 |
| Eosinophils | | | | | | | | | | | |
| 37,828,205 | | *PGAP3* | 32.6 | 17.5 | 33.9 | 16.3 | -1.3 | -0.270 | 0.087 | 0.002 | 0.014 |
| 37,831,276 | | *PGAP3* | 75.8 | 14.6 | 73.3 | 15.4 | 2.5 | 0.324 | 0.116 | 0.005 | 0.033 |
| 37,831,660 | | *PGAP3* | 83.2 | 11.4 | 87.0 | 10.5 | -3.8 | -0.304 | 0.115 | 0.008 | 0.046 |
| 37,835,787 | | *PGAP3* | 13.5 | 7.6 | 15.8 | 10.3 | -2.3 | -0.264 | 0.086 | 0.002 | 0.014 |
| 37,844,210 | *ERBB2* | *PGAP3* | 0.4 | 1.2 | 0.2 | 0.7 | 0.2 | 2.468 | 0.837 | 0.003 | 0.022 |
| 37,844,223 | *ERBB2* | *PGAP3* | 0.7 | 1.7 | 0.3 | 1.0 | 0.4 | 1.678 | 0.537 | 0.002 | 0.013 |
| 37,844,950 | *PGAP3* | *ERBB2* | 4.3 | 5.3 | 4.2 | 5.1 | 0.1 | 0.564 | 0.154 | 2.59E-04 | 0.002 |

| | | | | | | | | | | | |
|---|---|---|---|---|---|---|---|---|---|---|---|
| 37,845,059 | *PGAP3* | *ERBB2* | 82.4 | 10.7 | 85.4 | 9.0 | -3.0 | -0.230 | 0.084 | 0.006 | 0.039 |
| 37,846,655 | | *ERBB2* | 94.2 | 8.1 | 96.8 | 6.3 | -2.6 | -0.926 | 0.352 | 0.009 | 0.049 |
| 37,851,022 | | *ERBB2* | 91.0 | 10.3 | 89.7 | 9.6 | 1.3 | 0.384 | 0.142 | 0.007 | 0.042 |
| 37,856,149 | | *ERBB2* | 16.9 | 10.4 | 16.3 | 11.1 | 0.6 | 0.265 | 0.094 | 0.005 | 0.031 |
| 37,856,430 | | *ERBB2* | 1.8 | 5.9 | 2.0 | 3.6 | -0.2 | -1.453 | 0.420 | 0.001 | 0.004 |
| 37,856,513 | | *ERBB2* | 1.9 | 4.7 | 4.0 | 6.8 | -2.1 | -0.940 | 0.331 | 0.005 | 0.029 |
| 37,857,767 | | *ERBB2* | 72.7 | 12.7 | 67.2 | 12.6 | 5.5 | 0.195 | 0.067 | 0.003 | 0.023 |
| 37,857,817 | | *ERBB2* | 35.8 | 11.6 | 32.3 | 12.0 | 3.5 | 0.224 | 0.058 | 1.12E-04 | 0.001 |
| 37,879,814 | | *ERBB2* | 94.8 | 7.6 | 95.1 | 6.6 | -0.3 | 0.575 | 0.215 | 0.007 | 0.043 |
| 37,881,102 | | *ERBB2* | 95.3 | 6.0 | 96.2 | 4.5 | -0.9 | -0.602 | 0.202 | 0.003 | 0.020 |
| 37,884,285 | | *ERBB2* | 93.1 | 8.5 | 95.7 | 6.3 | -2.6 | -0.614 | 0.193 | 0.001 | 0.011 |
| 37,886,523 | | *MIEN1* | 0.4 | 1.4 | 0.5 | 1.7 | -0.1 | -1.549 | 0.576 | 0.007 | 0.042 |
| 37,893,744 | | | 50.6 | 14.7 | 55.1 | 14.1 | -4.5 | -0.196 | 0.072 | 0.006 | 0.039 |
| 37,894,702 | | | 2.7 | 3.8 | 5.4 | 7.6 | -2.7 | -0.747 | 0.263 | 0.005 | 0.029 |
| 37,895,143 | | | 7.2 | 11.2 | 10.5 | 10.7 | -3.3 | -0.977 | 0.180 | 5.70E-08 | 4.85E-07 |
| 37,895,775 | | | 3.7 | 6.4 | 5.5 | 9.0 | -1.8 | -0.600 | 0.221 | 0.007 | 0.040 |
| 37,896,165 | | | 13.2 | 11.5 | 10.6 | 9.6 | 2.6 | 0.411 | 0.133 | 0.002 | 0.014 |
| 37,910,184 | | | 0.5 | 2.3 | 0.9 | 3.2 | -0.4 | -2.408 | 0.737 | 0.001 | 0.008 |
| 37,910,294 | | | 0.3 | 1.1 | 0.5 | 1.3 | -0.2 | -2.630 | 1.001 | 0.009 | 0.049 |
| 37,910,569 | | | 0.4 | 1.1 | 0.2 | 0.8 | 0.2 | 2.426 | 0.890 | 0.006 | 0.039 |
| 37,913,899 | | | 76.5 | 14.0 | 80.5 | 13.6 | -4.0 | -0.292 | 0.101 | 0.004 | 0.025 |
| 37,933,933 | | *IKZF3* | 6.0 | 9.7 | 8.5 | 11.0 | -2.5 | -0.463 | 0.166 | 0.005 | 0.032 |
| 37,937,868 | | *IKZF3* | 82.4 | 15.2 | 79.9 | 15.7 | 2.5 | 0.383 | 0.137 | 0.005 | 0.032 |
| 37,970,196 | | *IKZF3* | 10.0 | 15.3 | 12.7 | 16.1 | -2.7 | -0.492 | 0.115 | 1.73E-05 | 1.44E-04 |
| 37,970,198 | | *IKZF3* | 13.0 | 17.5 | 14.3 | 17.6 | -1.3 | -0.292 | 0.104 | 0.005 | 0.032 |
| 37,999,093 | | *IKZF3* | 90.0 | 12.2 | 93.5 | 8.8 | -3.5 | -0.654 | 0.206 | 0.002 | 0.011 |
| 37,999,368 | | *IKZF3* | 88.3 | 11.0 | 86.0 | 12.0 | 2.3 | 0.317 | 0.121 | 0.009 | 0.049 |
| 38,003,876 | | *IKZF3* | 13.1 | 10.4 | 12.4 | 9.3 | 0.7 | 0.356 | 0.107 | 0.001 | 0.007 |
| 38,020,344 | | *IKZF3* | 4.3 | 5.4 | 3.4 | 4.3 | 0.9 | 0.398 | 0.141 | 0.005 | 0.030 |
| 38,021,072 | *IKZF3* | | 6.5 | 7.4 | 8.6 | 10.7 | -2.1 | -0.644 | 0.202 | 0.001 | 0.011 |
| 38,024,315 | | | 18.7 | 15.1 | 21.8 | 17.5 | -3.1 | -0.412 | 0.109 | 1.59E-04 | 0.001 |
| 38,024,317 | | | 19.4 | 15.5 | 24.6 | 17.8 | -5.2 | -0.474 | 0.107 | 1.01E-05 | 8.47E-05 |
| 38,024,331 | | | 40.5 | 18.7 | 43.9 | 21.0 | -3.4 | -0.297 | 0.084 | 3.88E-04 | 0.003 |
| 38,024,335 | | | 14.8 | 11.0 | 17.7 | 12.8 | -2.9 | -0.371 | 0.112 | 0.001 | 0.007 |

| | | | | | | | | | | | |
|---|---|---|---|---|---|---|---|---|---|---|---|
| 38,024,342 | | | 20.6 | 14.8 | 22.6 | 15.0 | -2.0 | -0.391 | 0.101 | 1.11E-04 | 0.001 |
| 38,024,367 | | | 20.8 | 14.7 | 23.7 | 16.6 | -2.9 | -0.316 | 0.101 | 0.002 | 0.013 |
| 38,024,456 | | | 22.9 | 15.2 | 27.4 | 15.8 | -4.5 | -0.441 | 0.115 | 0.000 | 0.001 |
| 38,024,458 | | | 23.5 | 17.5 | 28.3 | 17.8 | -4.8 | -0.314 | 0.118 | 0.008 | 0.044 |
| 38,075,408 | *GSDMB* | | 92.9 | 7.5 | 94.8 | 5.9 | -1.9 | -0.415 | 0.154 | 0.007 | 0.041 |
| 38,077,098 | | | 54.3 | 15.1 | 53.5 | 15.6 | 0.8 | 0.200 | 0.068 | 0.003 | 0.022 |
| 38,077,104 | | | 62.4 | 13.9 | 59.2 | 14.9 | 3.2 | 0.296 | 0.069 | 1.82E-05 | 1.50E-04 |
| 38,077,107 | | | 58.1 | 15.9 | 53.3 | 16.2 | 4.8 | 0.333 | 0.068 | 1.02E-06 | 8.59E-06 |
| 38,077,703 | | *ORMDL3* | 3.5 | 3.8 | 6.8 | 8.8 | -3.3 | -0.388 | 0.124 | 0.002 | 0.013 |
| 38,077,798 | | *ORMDL3* | 0.5 | 1.4 | 1.0 | 1.9 | -0.5 | -1.039 | 0.360 | 0.004 | 0.026 |
| 38,077,826 | | *ORMDL3* | 1.9 | 3.2 | 3.2 | 6.3 | -1.3 | -0.796 | 0.241 | 0.001 | 0.007 |
| 38,081,710 | | *ORMDL3* | 64.4 | 14.9 | 67.5 | 13.0 | -3.1 | -0.216 | 0.079 | 0.006 | 0.038 |
| 38,084,256 | *ORMDL3* | | 8.1 | 11.5 | 7.3 | 6.0 | 0.8 | -0.308 | 0.107 | 0.004 | 0.027 |
| 38,084,445 | *ORMDL3* | | 41.4 | 15.8 | 43.7 | 18.5 | -2.3 | -0.211 | 0.069 | 0.002 | 0.016 |
| 38,084,450 | *ORMDL3* | | 35.4 | 17.7 | 37.2 | 19.0 | -1.8 | -0.220 | 0.071 | 0.002 | 0.013 |
| 38,084,474 | *ORMDL3* | | 46.5 | 18.1 | 46.3 | 16.6 | 0.2 | -0.191 | 0.067 | 0.004 | 0.027 |
| 38,084,816 | *ORMDL3* | | 84.2 | 11.9 | 80.2 | 15.7 | 4.0 | 0.387 | 0.132 | 0.003 | 0.023 |
| 38,085,536 | | | 42.5 | 20.3 | 40.0 | 22.2 | 2.5 | 0.196 | 0.072 | 0.006 | 0.039 |
| 38,108,509 | | | 93.3 | 5.4 | 91.8 | 6.7 | 1.5 | 0.505 | 0.126 | 6.43E-05 | 0.001 |
| 38,108,523 | | | 94.9 | 5.5 | 92.3 | 6.3 | 2.6 | 0.464 | 0.132 | 4.22E-04 | 0.003 |
| 38,108,630 | | | 77.4 | 12.7 | 72.4 | 13.0 | 5.0 | 0.199 | 0.075 | 0.008 | 0.044 |
| 38,133,359 | | *GSDMA* | 67.3 | 14.2 | 71.1 | 16.1 | -3.8 | -0.272 | 0.091 | 0.003 | 0.019 |

[a]Genes are indicated in the second and/or third columns if the CpG position is within 1,500 bp from the transcription start site (Gene is methylated [1.5 kb from TSS]) or in the gene body (Gene is methylated [gene body]). [b]Positive difference refers to the percentage of methylation levels increased in individuals affected by asthma compared to controls, and a negative difference refers to methylation levels decreased in individuals affected by asthma.

Abbreviations: Aff = samples from individuals affected with asthma, SD = standard deviation, SE = standard error, TSS = transcription start site, Unaff = samples from unaffected individuals.
