## Supplemental Table 7 for "*GSDMA* drives the most replicated association with asthma in naïve CD4^+^ T cells"

**Table S7:** Significant associations between gene expression and methylation levels in naïve CD4$^{+}$ T cells

| Gene | Position TSS (hg19) | Strand | Position CpG (hg19) | Gene is methylated (1.5 kb from TSS)[a] | Gene is methylated (gene body)[a] | Pearson's r | Coeff | SE | P value | FDR |
|---|---|---|---|---|---|---|---|---|---|---|
| *PGAP3* | 37,844,323 | - | 38,023,747 | | | 0.307 | 0.038 | 0.009 | 5.14E-05 | 0.012 |
| | | | 38,085,469 | | | 0.223 | 0.027 | 0.005 | 4.90E-07 | 2.54E-04 |
| *ERBB2* | 37,844,336 | + | 37,843,621 | *ERBB2* | *PGAP3* | -0.235 | -0.078 | 0.019 | 5.26E-05 | 0.012 |
| | | | 37,856,890 | | *ERBB2* | 0.283 | 0.048 | 0.012 | 6.88E-05 | 0.014 |
| | | | 37,907,944 | | | -0.200 | -0.047 | 0.013 | 3.77E-04 | 0.050 |
| | | | 37,914,933 | | | -0.258 | -0.099 | 0.027 | 2.59E-04 | 0.040 |
| | | | 37,914,953 | | | -0.220 | -0.134 | 0.038 | 3.69E-04 | 0.050 |
| | | | 37,947,691 | | *IKZF3* | 0.414 | 0.049 | 0.012 | 5.85E-05 | 0.013 |
| | | | 37,947,697 | | *IKZF3* | 0.380 | 0.055 | 0.013 | 3.40E-05 | 0.009 |
| | | | 37,947,773 | | *IKZF3* | 0.398 | 0.057 | 0.013 | 1.38E-05 | 0.004 |
| | | | 37,951,189 | | *IKZF3* | 0.359 | 0.041 | 0.011 | 1.02E-04 | 0.020 |
| | | | 37,999,368 | | *IKZF3* | 0.319 | 0.041 | 0.011 | 3.07E-04 | 0.045 |
| | | | 37,999,641 | | *IKZF3* | 0.426 | 0.044 | 0.009 | 2.93E-06 | 0.001 |
| | | | 38,020,606 | *IKZF3* | | 0.288 | 0.078 | 0.021 | 2.56E-04 | 0.040 |
| | | | 38,083,300 | | *ORMDL3* | 0.262 | 0.079 | 0.022 | 3.64E-04 | 0.050 |
| | | | 38,109,595 | | | 0.328 | 0.090 | 0.023 | 1.16E-04 | 0.022 |
| | | | 38,120,370 | | *GSDMA* | 0.409 | 0.136 | 0.021 | 7.48E-11 | 9.06E-08 |
| *MIEN1* | 37,886,816 | - | 37,914,933 | | | -0.034 | -0.055 | 0.015 | 2.10E-04 | 0.034 |
| | | | 37,937,926 | | *IKZF3* | -0.235 | -0.041 | 0.011 | 1.57E-04 | 0.028 |
| *IKZF3* | 38,020,441 | - | 37,894,120 | | | 0.229 | 0.042 | 0.012 | 3.76E-04 | 0.050 |
| | | | 37,894,659 | | | 0.265 | 0.032 | 0.008 | 1.27E-04 | 0.024 |
| | | | 37,914,933 | | | -0.217 | -0.058 | 0.016 | 3.16E-04 | 0.046 |
| | | | 37,914,953 | | | -0.251 | -0.110 | 0.022 | 5.16E-07 | 2.58E-04 |
| | | | 37,927,002 | | *IKZF3* | -0.145 | -0.047 | 0.012 | 5.45E-05 | 0.012 |
| | | | 37,929,495 | | *IKZF3* | -0.269 | -0.072 | 0.017 | 3.23E-05 | 0.009 |
| | | | 37,937,926 | | *IKZF3* | -0.233 | -0.053 | 0.012 | 5.14E-06 | 0.002 |
| | | | 37,938,973 | | *IKZF3* | -0.096 | -0.043 | 0.011 | 6.10E-05 | 0.013 |

| | | | | | | | | |
|---|---|---|---|---|---|---|---|---|
| | | 37,949,341 | *IKZF3* | -0.380 | -0.087 | 0.019 | 4.30E-06 | 0.002 |
| | | 38,023,480 | | -0.409 | -0.098 | 0.021 | 4.07E-06 | 0.002 |
| | | 38,023,500 | | -0.355 | -0.078 | 0.018 | 1.16E-05 | 0.004 |
| | | 38,023,639 | | -0.397 | -0.094 | 0.024 | 8.53E-05 | 0.017 |
| | | 38,023,679 | | -0.327 | -0.082 | 0.020 | 5.77E-05 | 0.013 |
| | | 38,023,681 | | -0.309 | -0.074 | 0.013 | 3.82E-09 | 3.47E-06 |
| | | 38,023,707 | | -0.392 | -0.096 | 0.018 | 1.59E-07 | 1.00E-04 |
| | | 38,023,747 | | -0.324 | -0.071 | 0.013 | 1.08E-07 | 7.15E-05 |
| | | 38,023,830 | | -0.398 | -0.108 | 0.020 | 4.15E-08 | 2.87E-05 |
| | | 38,023,858 | | -0.345 | -0.080 | 0.017 | 2.16E-06 | 0.001 |
| | | 38,023,891 | | -0.398 | -0.114 | 0.022 | 2.18E-07 | 1.21E-04 |
| | | 38,023,914 | | -0.419 | -0.074 | 0.016 | 2.13E-06 | 0.001 |
| | | 38,023,920 | | -0.337 | -0.062 | 0.017 | 2.07E-04 | 0.034 |
| | | 38,023,993 | | -0.219 | -0.070 | 0.020 | 3.65E-04 | 0.050 |
| | | 38,024,048 | | -0.307 | -0.077 | 0.019 | 7.92E-05 | 0.016 |
| | | 38,024,063 | | -0.257 | -0.048 | 0.014 | 3.48E-04 | 0.050 |
| | | 38,024,159 | | -0.196 | -0.040 | 0.009 | 6.55E-06 | 0.002 |
| | | 38,051,451 | | -0.047 | -0.041 | 0.011 | 3.04E-04 | 0.045 |
| | | 38,072,832 | *GSDMB* | -0.185 | -0.021 | 0.006 | 3.16E-04 | 0.046 |
| *GSDMB* | 38,074,903 - | 37,926,502 | *IKZF3* | -0.430 | -0.045 | 0.013 | 3.48E-04 | 0.050 |
| | | 37,947,691 | *IKZF3* | -0.380 | -0.034 | 0.009 | 2.69E-04 | 0.041 |
| | | 38,077,051 | | -0.383 | -0.106 | 0.024 | 1.49E-05 | 0.005 |
| | | 38,077,104 | | -0.320 | -0.080 | 0.019 | 4.07E-05 | 0.010 |
| | | 38,077,107 | | -0.294 | -0.058 | 0.015 | 1.53E-04 | 0.027 |
| | | 38,077,176 | | -0.337 | -0.061 | 0.015 | 4.25E-05 | 0.010 |
| | | 38,077,234 | | -0.315 | -0.036 | 0.009 | 5.33E-05 | 0.012 |
| | | 38,077,703 | *ORMDL3* | -0.167 | -0.109 | 0.026 | 2.22E-05 | 0.006 |
| | | 38,080,398 | *ORMDL3* | -0.361 | -0.091 | 0.019 | 1.54E-06 | 0.001 |
| | | 38,081,492 | *ORMDL3* | -0.346 | -0.028 | 0.007 | 1.57E-04 | 0.028 |
| | | 38,081,812 | *ORMDL3* | -0.470 | -0.056 | 0.010 | 3.19E-08 | 2.32E-05 |
| | | 38,081,829 | *ORMDL3* | -0.510 | -0.051 | 0.009 | 1.32E-08 | 1.13E-05 |
| | | 38,081,995 | *ORMDL3* | -0.548 | -0.105 | 0.013 | 9.14E-15 | 2.21E-11 |
| | | 38,082,018 | *ORMDL3* | -0.478 | -0.057 | 0.011 | 2.89E-07 | 1.56E-04 |
| | | 38,082,022 | *ORMDL3* | -0.549 | -0.109 | 0.014 | 1.63E-15 | 5.92E-12 |

| | | | | | | | | |
|---|---|---|---|---|---|---|---|---|
| | | 38,082,035 | *ORMDL3* | -0.442 | -0.096 | 0.016 | 7.54E-10 | 8.42E-07 |
| | | 38,082,073 | *ORMDL3* | -0.413 | -0.141 | 0.024 | 1.94E-09 | 2.01E-06 |
| | | 38,082,206 | *ORMDL3* | -0.432 | -0.051 | 0.010 | 2.03E-07 | 1.19E-04 |
| | | 38,082,235 | *ORMDL3* | -0.365 | -0.049 | 0.012 | 3.21E-05 | 0.009 |
| | | 38,082,346 | *ORMDL3* | -0.494 | -0.106 | 0.014 | 3.35E-14 | 6.08E-11 |
| | | 38,082,448 | *ORMDL3* | -0.494 | -0.084 | 0.012 | 1.08E-11 | 1.58E-08 |
| | | 38,082,658 | *ORMDL3* | -0.395 | -0.040 | 0.010 | 1.12E-04 | 0.022 |
| *ORMDL3* | 38,084,057 - | 38,077,051 | | -0.367 | -0.095 | 0.021 | 6.66E-06 | 0.002 |
| | | 38,077,104 | | -0.334 | -0.070 | 0.017 | 2.86E-05 | 0.008 |
| | | 38,077,107 | | -0.312 | -0.049 | 0.013 | 1.98E-04 | 0.033 |
| | | 38,077,176 | | -0.331 | -0.053 | 0.013 | 3.67E-05 | 0.009 |
| | | 38,077,191 | | -0.359 | -0.029 | 0.007 | 4.00E-05 | 0.010 |
| | | 38,077,234 | | -0.360 | -0.033 | 0.008 | 1.60E-05 | 0.005 |
| | | 38,077,277 | | -0.268 | -0.037 | 0.010 | 1.49E-04 | 0.027 |
| | | 38,077,323 | *ORMDL3* | -0.315 | -0.024 | 0.007 | 2.58E-04 | 0.040 |
| | | 38,077,621 | *ORMDL3* | -0.338 | -0.092 | 0.021 | 1.01E-05 | 0.003 |
| | | 38,077,703 | *ORMDL3* | -0.349 | -0.113 | 0.022 | 2.06E-07 | 1.19E-04 |
| | | 38,080,398 | *ORMDL3* | -0.328 | -0.063 | 0.017 | 1.71E-04 | 0.029 |
| | | 38,081,492 | *ORMDL3* | -0.340 | -0.024 | 0.006 | 1.77E-04 | 0.030 |
| | | 38,081,812 | *ORMDL3* | -0.433 | -0.052 | 0.009 | 2.28E-09 | 2.20E-06 |
| | | 38,081,829 | *ORMDL3* | -0.479 | -0.050 | 0.007 | 2.09E-11 | 2.76E-08 |
| | | 38,081,995 | *ORMDL3* | -0.560 | -0.094 | 0.012 | 3.17E-16 | 1.53E-12 |
| | | 38,082,018 | *ORMDL3* | -0.436 | -0.053 | 0.009 | 2.30E-08 | 1.86E-05 |
| | | 38,082,022 | *ORMDL3* | -0.625 | -0.113 | 0.011 | 7.42E-25 | 9.31E-21 |
| | | 38,082,035 | *ORMDL3* | -0.478 | -0.099 | 0.013 | 4.69E-14 | 7.57E-11 |
| | | 38,082,073 | *ORMDL3* | -0.541 | -0.151 | 0.019 | 5.00E-15 | 1.45E-11 |
| | | 38,082,206 | *ORMDL3* | -0.382 | -0.042 | 0.009 | 9.95E-07 | 4.66E-04 |
| | | 38,082,235 | *ORMDL3* | -0.372 | -0.046 | 0.010 | 3.57E-06 | 0.001 |
| | | 38,082,346 | *ORMDL3* | -0.551 | -0.114 | 0.011 | 1.28E-24 | 9.31E-21 |
| | | 38,082,448 | *ORMDL3* | -0.516 | -0.080 | 0.010 | 1.14E-14 | 2.37E-11 |
| | | 38,082,658 | *ORMDL3* | -0.387 | -0.040 | 0.009 | 6.40E-06 | 0.002 |
| | | 38,082,778 | *ORMDL3* | -0.376 | -0.053 | 0.012 | 1.19E-05 | 0.004 |
| | | 38,082,921 | *ORMDL3* | -0.403 | -0.036 | 0.008 | 1.25E-05 | 0.004 |
| | | 38,084,297 | *ORMDL3* | 0.308 | 0.085 | 0.022 | 7.55E-05 | 0.016 |

| | | | | | | | | | | |
|---|---|---|---|---|---|---|---|---|---|---|
| *GSDMA* | 38,119,225 | + | 37,834,869 | | *PGAP3* | -0.304 | -0.181 | 0.033 | 2.74E-08 | 2.10E-05 |
| | | | 37,894,258 | | | 0.310 | 0.229 | 0.056 | 4.67E-05 | 0.011 |
| | | | 38,023,747 | | | -0.350 | -0.259 | 0.062 | 3.45E-05 | 0.009 |
| | | | 38,109,870 | | | -0.224 | -0.220 | 0.062 | 3.76E-04 | 0.050 |
| | | | 38,119,198 | *GSDMA* | | -0.238 | -0.129 | 0.035 | 2.50E-04 | 0.039 |
| | | | 38,119,207 | *GSDMA* | | -0.255 | -0.154 | 0.041 | 1.61E-04 | 0.028 |
| | | | 38,119,649 | | *GSDMA* | -0.192 | -0.152 | 0.041 | 1.84E-04 | 0.031 |

[a]Genes are indicated in the fourth and/or fifth columns if the CpG position is within 1,500 bp from the transcription start site (Gene is methylated [1.5 kb from TSS]) or in the gene body (Gene is methylated [gene body]).

Abbreviations: TSS = transcription start site.
