## Supplemental Table 8 for "*GSDMA* drives the most replicated association with asthma in naïve CD4^+^ T cells"

**Table S8:** Significant associations between gene expression and methylation levels in eosinophils

| Gene | Position TSS (hg19) | Strand | Position CpG (hg19) | Gene is methylated (1.5 kb from TSS)[a] | Gene is methylated (Gene body)[a] | Pearson's r | Coeff | SE | P value | FDR |
|---|---|---|---|---|---|---|---|---|---|---|
| *PGAP3* | 37,844,323 | - | 37,896,750 | | | 0.015 | -0.060 | 0.013 | 1.89E-06 | 0.002 |
| | | | 37,911,851 | | | -0.161 | -0.087 | 0.021 | 3.87E-05 | 0.020 |
| | | | 37,922,462 | | *IKZF3* | -0.183 | 0.036 | 0.009 | 1.00E-04 | 0.039 |
| | | | 38,024,174 | | | -0.037 | -0.070 | 0.014 | 7.47E-07 | 0.002 |
| *ERBB2* | 37,844,336 | + | 37,881,117 | | *ERBB2* | -0.030 | 0.093 | 0.024 | 1.25E-04 | 0.044 |
| | | | 37,895,384 | | | 0.016 | 0.133 | 0.030 | 8.07E-06 | 0.005 |
| | | | 37,898,348 | | | -0.130 | 0.088 | 0.023 | 1.01E-04 | 0.039 |
| | | | 38,004,064 | | *IKZF3* | -0.084 | 0.187 | 0.045 | 3.79E-05 | 0.020 |
| | | | 38,083,354 | | *ORMDL3* | -0.014 | 0.251 | 0.056 | 7.09E-06 | 0.005 |
| | | | 38,108,459 | | | -0.067 | 0.098 | 0.026 | 1.47E-04 | 0.048 |
| | | | 38,128,699 | | | -0.109 | 0.114 | 0.023 | 1.29E-06 | 0.002 |
| *IKZF3* | 38,020,441 | - | 37,828,163 | | *PGAP3* | 0.005 | 0.107 | 0.027 | 8.80E-05 | 0.038 |
| | | | 37,894,403 | | | 0.215 | 0.068 | 0.015 | 7.44E-06 | 0.005 |
| | | | 37,894,413 | | | 0.213 | 0.072 | 0.015 | 1.97E-06 | 0.002 |
| | | | 37,894,590 | | | 0.191 | 0.056 | 0.014 | 1.09E-04 | 0.041 |
| | | | 37,894,608 | | | 0.098 | 0.064 | 0.016 | 6.90E-05 | 0.033 |
| | | | 37,896,063 | | | 0.113 | 0.060 | 0.016 | 1.57E-04 | 0.048 |
| | | | 37,914,977 | | | -0.129 | -0.207 | 0.053 | 9.24E-05 | 0.039 |
| | | | 37,938,833 | | *IKZF3* | 0.106 | 0.058 | 0.015 | 6.91E-05 | 0.033 |
| | | | 37,938,939 | | *IKZF3* | 0.034 | 0.070 | 0.015 | 2.93E-06 | 0.003 |
| | | | 38,020,407 | | *IKZF3* | 0.069 | -0.133 | 0.028 | 2.70E-06 | 0.003 |
| | | | 38,023,914 | | | -0.419 | -0.092 | 0.020 | 2.47E-06 | 0.003 |
| | | | 38,023,920 | | | -0.337 | -0.097 | 0.020 | 1.65E-06 | 0.002 |
| | | | 38,024,174 | | | -0.186 | -0.109 | 0.024 | 6.58E-06 | 0.005 |
| | | | 38,024,285 | | | -0.090 | -0.172 | 0.035 | 1.22E-06 | 0.002 |
| | | | 38,024,290 | | | -0.065 | -0.116 | 0.028 | 2.97E-05 | 0.017 |

[a]Genes are indicated in the fourth and/or fifth columns if the CpG position is within 1,500 bp from the transcription start site (Gene is methylated [1.5 kb from TSS]) or int he gene body (Gene is methylated [gene body]).
Abbreviations: TSS = transcription start site.
