## Supplemental Table 9 for "*GSDMA* drives the most replicated association with asthma in naïve CD4^+^ T cells"

**Table S9:** Significant eQTLs in naïve $CD4^+$ T cells

| Haplotype block | SNP id | HGVS names (hg19) | Gene | Statistic | P eQTL | P asthma[a] |
|---|---|---|---|---|---|---|
| First block | rs14050 | g.37828072.G>A | *PGAP3* | -3.95 | 1.24E-04 | |
| | rs2952151 | g.37828496.T>C | *PGAP3* | -3.95 | 1.24E-04 | |
| | rs907087 | g.37828787.G>A | *PGAP3* | -3.95 | 1.24E-04 | |
| | rs2247862 | g.37829129.T>C | *PGAP3* | -3.95 | 1.24E-04 | |
| | rs903503 | g.37829571.T>G | *PGAP3* | -3.71 | 3.00E-04 | |
| | rs903502 | g.37829604.A>G | *PGAP3* | -3.84 | 1.86E-04 | |
| | rs2934956 | g.37830447.T>A | *PGAP3* | -3.84 | 1.86E-04 | |
| | rs1565920 | g.37831613.T>C | *PGAP3* | -3.84 | 1.86E-04 | |
| | rs1495102 | g.37832093.A>G | *PGAP3* | -3.95 | 1.24E-04 | |
| | rs1495101 | g.37832103.G>T | *PGAP3* | -3.95 | 1.24E-04 | |
| | rs1495100 | g.37832279.A>G | *PGAP3* | -3.84 | 1.86E-04 | |
| | rs2934953 | g.37832315.A>T | *PGAP3* | -3.95 | 1.24E-04 | |
| | rs2934952 | g.37832366.C>T | *PGAP3* | -3.95 | 1.24E-04 | |
| | rs2941505 | g.37832704.A>G | *PGAP3* | -3.84 | 1.86E-04 | |
| | rs2952152 | g.37832735.T>C | *PGAP3* | -3.84 | 1.86E-04 | |
| | rs2941506 | g.37833035.A>G | *PGAP3* | -3.84 | 1.86E-04 | |
| | rs2934951 | g.37833328.T>C | *PGAP3* | -3.84 | 1.86E-04 | |
| | rs907088 | g.37833567.G>C | *PGAP3* | -4.04 | 8.62E-05 | |
| | rs907089 | g.37833600.G>A | *PGAP3* | -4.04 | 8.62E-05 | |
| | rs907090 | g.37833632.T>C | *PGAP3* | -4.04 | 8.62E-05 | |
| | rs9675194 | g.37833805.T>C | *PGAP3* | -4.04 | 8.62E-05 | |
| | rs2313171 | g.37833842.T>C | *PGAP3* | -4.04 | 8.62E-05 | |
| | rs732083 | g.37834367.T>C | *PGAP3* | -4.04 | 8.62E-05 | |
| | rs12150603 | g.37834715.A>G | *PGAP3* | -4.04 | 8.62E-05 | |
| | rs8077172 | g.37834977.G>T | *PGAP3* | -4.04 | 8.62E-05 | |
| | rs8078228 | g.37834998.C>T | *PGAP3* | -4.04 | 8.62E-05 | |
| | rs1018246 | g.37835240.C>T | *PGAP3* | -4.04 | 8.62E-05 | |
| | rs11078919 | g.37835755.T>C | *PGAP3* | -4.04 | 8.62E-05 | |
| | rs1476278 | g.37836243.A>G | *PGAP3* | -4.04 | 8.62E-05 | |
| | rs9303274 | g.37836353.C>T | *PGAP3* | -4.04 | 8.62E-05 | |

| | | | | |
|---|---|---|---|---|
| rs12940986 | g.37836581.A>G | *PGAP3* | -4.04 | 8.62E-05 |
| rs2517952 | g.37838301.A>G | *PGAP3* | -3.95 | 1.24E-04 |
| rs2517957 | g.37838716.A>G | *PGAP3* | -4.04 | 8.62E-05 |
| rs2517958 | g.37838751.A>G | *PGAP3* | -4.04 | 8.62E-05 |
| rs903501 | g.37839493.A>G | *PGAP3* | -4.15 | 5.68E-05 |
| rs2517953 | g.37841211.C>G | *GSDMB* | 3.40 | 8.74E-04 |
| | | *PGAP3* | -3.93 | 1.31E-04 |
| rs2517954 | g.37843550.T>C | *GSDMB* | 3.40 | 8.78E-04 |
| | | *PGAP3* | -4.04 | 8.56E-05 |
| rs2517955 | g.37843681.T>C | *GSDMB* | 3.44 | 7.66E-04 |
| | | *PGAP3* | -3.93 | 1.29E-04 |
| rs2517956 | g.37843859.G>A | *GSDMB* | 3.40 | 8.78E-04 |
| | | *PGAP3* | -4.04 | 8.56E-05 |
| rs2517959 | g.37846512.A>T | *GSDMB* | 3.40 | 8.78E-04 |
| | | *PGAP3* | -4.04 | 8.56E-05 |
| rs2517960 | g.37846521.C>T | *GSDMB* | 3.44 | 7.66E-04 |
| | | *PGAP3* | -3.93 | 1.29E-04 |
| rs557885708 | g.37847799_37847800insAAA | *GSDMB* | 3.40 | 8.78E-04 |
| | | *PGAP3* | -4.04 | 8.56E-05 |
| rs2643194 | g.37853048.C>T | *GSDMB* | 3.39 | 9.03E-04 |
| | | *PGAP3* | -3.84 | 1.80E-04 |
| rs2517951 | g.37853097.T>C | *PGAP3* | -3.62 | 4.13E-04 |
| rs2643195 | g.37853118.A>G | *PGAP3* | -3.73 | 2.75E-04 |
| rs2934971 | g.37854507.G>T | *GSDMB* | 3.39 | 9.03E-04 |
| | | *PGAP3* | -3.84 | 1.80E-04 |
| rs1565923 | g.37858678.A>G | *GSDMB* | 3.39 | 9.03E-04 |
| | | *PGAP3* | -3.84 | 1.80E-04 |
| rs2934967 | g.37870378.G>A | *GSDMB* | 3.39 | 9.03E-04 |
| | | *PGAP3* | -3.84 | 1.80E-04 |
| rs2952156 | g.37876835.A>G | *GSDMB* | 3.39 | 8.91E-04 |
| | | *PGAP3* | -3.64 | 3.85E-04 |
| rs2952157 | g.37877412.G>A | *GSDMB* | 3.39 | 8.91E-04 |
| | | *PGAP3* | -3.64 | 3.85E-04 |

| | | | | | |
|---|---|---|---|---|---|
| | rs11653998 | g.37877447.G>C | *GSDMB* | 3.43 | 7.90E-04 |
| | | | *PGAP3* | -3.53 | 5.63E-04 |
| | rs2088126 | g.37879030.A>G | *GSDMB* | 3.39 | 8.91E-04 |
| | | | *PGAP3* | -3.64 | 3.85E-04 |
| | rs903506 | g.37879762.G>A | *GSDMB* | 3.43 | 7.90E-04 |
| | | | *PGAP3* | -3.53 | 5.63E-04 |
| | rs547454526 | g.37880525_37880526insTTTTT | *GSDMB* | 3.44 | 7.70E-04 |
| | | | *PGAP3* | -3.53 | 5.61E-04 |
| | rs1058808 | g.37884037.C>G (Pro.1155.Ala) | *GSDMB* | 3.49 | 6.51E-04 |
| | | | *PGAP3* | -3.63 | 3.98E-04 |
| | rs10558975 | g.37831304.G>C | *PGAP3* | -3.90 | 1.46E-04 |
| | rs12150298 | g.37834541.T>C | *PGAP3* | -4.04 | 8.62E-05 |
| | rs1565922 | g.37831035.T>C | *PGAP3* | -3.95 | 1.24E-04 |
| | rs1810132 | g.37866005.C>T | *GSDMB* | 3.39 | 9.03E-04 |
| | | | *PGAP3* | -3.84 | 1.80E-04 |
| | rs2904765 | g.37848675.T>G | *GSDMB* | 3.36 | 9.96E-04 |
| | | | *PGAP3* | -4.22 | 4.29E-05 |
| | rs2904766 | g.37848677.A>G | *GSDMB* | 3.40 | 8.78E-04 |
| | | | *PGAP3* | -4.04 | 8.56E-05 |
| | rs2904768 | g.37850571.C>T | *GSDMB* | 3.39 | 9.03E-04 |
| | | | *PGAP3* | -3.84 | 1.80E-04 |
| | rs2941503 | g.37828745.A>G | *PGAP3* | -3.95 | 1.24E-04 |
| | rs2941504 | g.37830900.A>G (Val.155.Val) | *PGAP3* | -3.95 | 1.24E-04 |
| | rs4252627 | g.37868715.C>T | *GSDMB* | 3.43 | 7.98E-04 |
| | | | *PGAP3* | -3.74 | 2.69E-04 |
| | rs55717377 | g.37850569.C>T | *GSDMB* | 3.39 | 9.03E-04 |
| | | | *PGAP3* | -3.84 | 1.80E-04 |
| | rs732084 | g.37834357.T>G | *PGAP3* | -4.04 | 8.62E-05 |
| | rs903504 | g.37829570.C>G | *PGAP3* | -3.84 | 1.86E-04 |
| Third block | rs9747973 | g.37905107.C>T | *GSDMB* | -5.97 | 1.70E-08 |
| | | | *IKZF3* | 4.40 | 2.05E-05 |
| | | | *ORMDL3* | -6.00 | 1.49E-08 |

| | | | | |
|---|---|---|---|---|
| rs11078921 | g.37908867.A>C | *GSDMA* | -4.54 | 1.17E-05 |
| rs2941522 | g.37910368.G>A | *GSDMB* | -5.97 | 1.70E-08 |
| | | *IKZF3* | 4.40 | 2.05E-05 |
| | | *ORMDL3* | -6.00 | 1.49E-08 |
| rs12946510 | g.37912377.T>C | *GSDMA* | -3.47 | 6.85E-04 |
| | | *GSDMB* | -6.05 | 1.15E-08 |
| | | *ORMDL3* | -6.75 | 3.37E-10 |
| rs35833706 | g.37916390_37916390insT | *GSDMB* | -6.10 | 9.01E-09 |
| | | *IKZF3* | 4.26 | 3.73E-05 |
| | | *ORMDL3* | -6.15 | 7.02E-09 |
| rs3764354 | g.37916823.A>G | *GSDMB* | -9.98 | 3.58E-18 |
| | | *ORMDL3* | -10.56 | 1.10E-19 |
| rs907091 | g.37921742.C>T | *GSDMB* | -6.10 | 9.01E-09 |
| | | *IKZF3* | 4.26 | 3.73E-05 |
| | | *ORMDL3* | -6.15 | 7.02E-09 |
| rs2952140 | g.37928059.G>A | *GSDMB* | -6.86 | 1.89E-10 |
| | | *IKZF3* | 4.63 | 7.93E-06 |
| | | *ORMDL3* | -6.86 | 1.87E-10 |
| rs2313430 | g.37929816.A>G | *GSDMB* | -6.90 | 1.46E-10 |
| | | *IKZF3* | 4.64 | 7.76E-06 |
| | | *ORMDL3* | -6.88 | 1.71E-10 |
| rs10445308 | g.37938047.T>C | *GSDMA* | -4.04 | 8.63E-05 |
| | | *GSDMB* | -6.80 | 2.49E-10 |
| | | *IKZF3* | 3.70 | 3.06E-04 |
| | | *ORMDL3* | -7.54 | 4.64E-12 |
| rs12942330 | g.37939839.T>C | *GSDMA* | -4.04 | 8.63E-05 |
| | | *GSDMB* | -6.80 | 2.49E-10 |
| | | *IKZF3* | 3.70 | 3.06E-04 |
| | | *ORMDL3* | -7.54 | 4.64E-12 |
| rs11658993 | g.37940808.T>C | *GSDMA* | -4.04 | 8.63E-05 |
| | | *GSDMB* | -6.80 | 2.49E-10 |
| | | *IKZF3* | 3.70 | 3.06E-04 |
| | | *ORMDL3* | -7.54 | 4.64E-12 |
| rs35395438 | g.37952891.G>T | *GSDMB* | -8.57 | 1.37E-14 |

| | | | | | |
|---|---|---|---|---|---|
| | | | *ORMDL3* | -9.53 | 5.26E-17 |
| | rs112350333 | g.37952989.T>C | *GSDMB* | -9.23 | 3.02E-16 |
| | | | *ORMDL3* | -10.34 | 4.09E-19 |
| | rs2952144 | g.37960017.C>T | *GSDMB* | -6.90 | 1.46E-10 |
| | | | *IKZF3* | 4.64 | 7.76E-06 |
| | | | *ORMDL3* | -6.88 | 1.71E-10 |
| | rs4795395 | g.37962987.A>T | *GSDMA* | -4.04 | 8.63E-05 |
| | | | *GSDMB* | -6.80 | 2.49E-10 |
| | | | *IKZF3* | 3.70 | 3.06E-04 |
| | | | *ORMDL3* | -7.54 | 4.64E-12 |
| | rs9909593 | g.37970149.G>A | *GSDMA* | -4.04 | 8.63E-05 |
| | | | *GSDMB* | -6.80 | 2.49E-10 |
| | | | *IKZF3* | 3.70 | 3.06E-04 |
| | | | *ORMDL3* | -7.54 | 4.64E-12 |
| | rs17676191 | g.37949924.G>A | *PGAP3* | -3.37 | 9.54E-04 |
| | rs907092 | g.37922259.A>G (Ser.438.Ser) | *GSDMA* | -3.50 | 6.23E-04 |
| | | | *GSDMB* | -5.95 | 1.93E-08 |
| | | | *ORMDL3* | -6.77 | 2.90E-10 |
| Fourth block | rs113897057 | g.37975214_37975214insTTCTA | *GSDMB* | -6.83 | 2.18E-10 |
| | | | *IKZF3* | 4.60 | 9.06E-06 |
| | | | *ORMDL3* | -6.74 | 3.52E-10 |
| | rs12944882 | g.37983492.C>T | *GSDMB* | -7.11 | 4.96E-11 |
| | | | *IKZF3* | 4.50 | 1.37E-05 |
| | | | *ORMDL3* | -7.43 | 8.75E-12 |
| | rs3816470 | g.37985801.C>T | *GSDMB* | -7.11 | 4.96E-11 |
| | | | *IKZF3* | 4.50 | 1.37E-05 |
| | | | *ORMDL3* | -7.43 | 8.75E-12 |
| | rs62066988 | g.37992281.T>C | *GSDMA* | -5.08 | 1.12E-06 |
| | | | *IKZF3* | 3.76 | 2.43E-04 |
| | rs34233420 | g.38004929_38004933del | *GSDMA* | -3.89 | 1.50E-04 |
| | | | *GSDMB* | -6.79 | 2.67E-10 |
| | | | *IKZF3* | 4.03 | 8.84E-05 |

| | | | | |
|---|---|---|---|---|
| | | *ORMDL3* | -7.63 | 2.94E-12 |
| rs9916765 | g.38005595.C>T | *GSDMB* | -7.15 | 3.99E-11 |
| | | *IKZF3* | 4.85 | 3.14E-06 |
| | | *ORMDL3* | -7.10 | 5.04E-11 |
| rs35564481 | g.38020058_38020058insC | *GSDMB* | -6.79 | 2.71E-10 |
| | | *IKZF3* | 4.97 | 1.86E-06 |
| | | *ORMDL3* | -6.85 | 1.92E-10 |
| rs1453559 | g.38020419.G>A | *GSDMB* | -7.15 | 3.99E-11 |
| | | *IKZF3* | 4.85 | 3.14E-06 |
| | | *ORMDL3* | -7.10 | 5.04E-11 |
| rs4795397 | g.38023745.G>A | *GSDMA* | -4.00 | 9.90E-05 |
| | | *GSDMB* | -7.05 | 6.69E-11 |
| | | *IKZF3* | 3.94 | 1.27E-04 |
| | | *ORMDL3* | -7.82 | 9.97E-13 |
| rs11557466 | g.38024626.T>C (Leu.7.Leu) | *GSDMA* | -3.67 | 3.35E-04 |
| | | *GSDMB* | -6.98 | 9.54E-11 |
| | | *IKZF3* | 4.15 | 5.60E-05 |
| | | *ORMDL3* | -7.99 | 3.76E-13 |
| rs11078925 | g.38025208.C>T | *GSDMA* | -3.67 | 3.35E-04 |
| | | *GSDMB* | -6.98 | 9.54E-11 |
| | | *IKZF3* | 4.15 | 5.60E-05 |
| | | *ORMDL3* | -7.99 | 3.76E-13 |
| rs34120102 | g.38026035.A>G | *GSDMA* | -3.67 | 3.35E-04 |
| | | *GSDMB* | -6.98 | 9.54E-11 |
| | | *IKZF3* | 4.15 | 5.60E-05 |
| | | *ORMDL3* | -7.99 | 3.76E-13 |
| rs11655198 | g.38026169.T>C | *GSDMB* | -7.05 | 6.78E-11 |
| | | *IKZF3* | 5.07 | 1.20E-06 |
| | | *ORMDL3* | -7.25 | 2.27E-11 |
| rs11650661 | g.38026286.T>A | *GSDMB* | -7.05 | 6.78E-11 |
| | | *IKZF3* | 5.07 | 1.20E-06 |
| | | *ORMDL3* | -7.25 | 2.27E-11 |
| rs11655292 | g.38026361.G>C | *GSDMB* | -7.05 | 6.78E-11 |
| | | *IKZF3* | 5.07 | 1.20E-06 |

| | | | | |
|---|---|---|---|---|
| | | *ORMDL3* | -7.25 | 2.27E-11 |
| rs12709365 | g.38027400.G>A | *GSDMA* | -3.67 | 3.35E-04 |
| | | *GSDMB* | -6.98 | 9.54E-11 |
| | | *IKZF3* | 4.15 | 5.60E-05 |
| | | *ORMDL3* | -7.99 | 3.76E-13 |
| rs13380815 | g.38027583.G>A | *GSDMA* | -3.67 | 3.35E-04 |
| | | *GSDMB* | -6.98 | 9.54E-11 |
| | | *IKZF3* | 4.15 | 5.60E-05 |
| | | *ORMDL3* | -7.99 | 3.76E-13 |
| rs12936231 | g.38029120.G>C | *GSDMB* | -7.05 | 6.78E-11 |
| | | *IKZF3* | 5.07 | 1.20E-06 |
| | | *ORMDL3* | -7.25 | 2.27E-11 |
| rs11870965 | g.38030205.A>T | *GSDMA* | -3.62 | 4.02E-04 |
| | | *GSDMB* | -6.91 | 1.39E-10 |
| | | *IKZF3* | 4.10 | 6.74E-05 |
| | | *ORMDL3* | -8.05 | 2.80E-13 |
| rs9903250 | g.38031030.A>G | *GSDMB* | -7.05 | 6.78E-11 |
| | | *IKZF3* | 5.07 | 1.20E-06 |
| | | *ORMDL3* | -7.25 | 2.27E-11 |
| rs10852935 | g.38031674.T>C (Cys.292.Cys) | *GSDMA* | -3.67 | 3.35E-04 |
| | | *GSDMB* | -6.98 | 9.54E-11 |
| | | *IKZF3* | 4.15 | 5.60E-05 |
| | | *ORMDL3* | -7.99 | 3.76E-13 |
| rs10852936 | g.38031714.T>C | *GSDMA* | -3.75 | 2.52E-04 |
| | | *GSDMB* | -6.97 | 1.01E-10 |
| | | *IKZF3* | 4.26 | 3.72E-05 |
| | | *ORMDL3* | -8.02 | 3.29E-13 |
| rs9891174 | g.38031802.A>T | *GSDMA* | -3.67 | 3.35E-04 |
| | | *GSDMB* | -6.98 | 9.54E-11 |
| | | *IKZF3* | 4.15 | 5.60E-05 |
| | | *ORMDL3* | -7.99 | 3.76E-13 |
| rs59716545 | g.38031857.G>T | *GSDMB* | -6.42 | 1.83E-09 |
| | | *IKZF3* | 3.87 | 1.66E-04 |

| | | | | |
|---|---|---|---|---|
| rs34189114 | g.38032460.T>C | *ORMDL3* | -6.94 | 1.22E-10 |
| | | *GSDMA* | -3.67 | 3.35E-04 |
| | | *GSDMB* | -6.98 | 9.54E-11 |
| | | *IKZF3* | 4.15 | 5.60E-05 |
| rs35736272 | g.38032680.C>T | *ORMDL3* | -7.99 | 3.76E-13 |
| | | *GSDMA* | -3.67 | 3.35E-04 |
| | | *GSDMB* | -6.98 | 9.54E-11 |
| | | *IKZF3* | 4.15 | 5.60E-05 |
| rs1054609 | g.38033277.C>A | *ORMDL3* | -7.99 | 3.76E-13 |
| | | *GSDMA* | -3.67 | 3.35E-04 |
| | | *GSDMB* | -6.98 | 9.54E-11 |
| | | *IKZF3* | 4.15 | 5.60E-05 |
| rs9907088 | g.38035116.A>G | *ORMDL3* | -7.99 | 3.76E-13 |
| | | *GSDMA* | -3.67 | 3.35E-04 |
| | | *GSDMB* | -6.98 | 9.54E-11 |
| | | *IKZF3* | 4.15 | 5.60E-05 |
| rs36038753 | g.38035370.T>G | *ORMDL3* | -7.99 | 3.76E-13 |
| | | *GSDMA* | -3.67 | 3.35E-04 |
| | | *GSDMB* | -6.98 | 9.54E-11 |
| | | *IKZF3* | 4.15 | 5.60E-05 |
| rs35569035 | g.38035624.T>C | *ORMDL3* | -7.99 | 3.76E-13 |
| | | *GSDMB* | -7.09 | 5.45E-11 |
| | | *IKZF3* | 4.41 | 2.00E-05 |
| rs9910826 | g.38035648.G>A | *ORMDL3* | -7.78 | 1.24E-12 |
| | | *GSDMB* | -7.09 | 5.45E-11 |
| | | *IKZF3* | 4.41 | 2.00E-05 |
| rs34074973 | g.38035766_38035769del | *ORMDL3* | -7.78 | 1.24E-12 |
| | | *GSDMA* | -3.67 | 3.35E-04 |
| | | *GSDMB* | -6.98 | 9.54E-11 |
| | | *IKZF3* | 4.15 | 5.60E-05 |
| rs9904624 | g.38036586.G>A | *ORMDL3* | -7.99 | 3.76E-13 |
| | | *GSDMA* | -3.67 | 3.35E-04 |
| | | *GSDMB* | -6.98 | 9.54E-11 |
| | | *IKZF3* | 4.15 | 5.60E-05 |

| | | | | |
|---|---|---|---|---|
| rs4795398 | g.38038179.T>C | *ORMDL3* | -7.99 | 3.76E-13 |
| | | *GSDMA* | -3.67 | 3.35E-04 |
| | | *GSDMB* | -6.98 | 9.54E-11 |
| | | *IKZF3* | 4.15 | 5.60E-05 |
| rs148094956 | g.38039561_38039565del | *ORMDL3* | -7.99 | 3.76E-13 |
| | | *GSDMB* | -7.05 | 6.78E-11 |
| | | *IKZF3* | 5.07 | 1.20E-06 |
| rs12232497 | g.38040119.C>T | *ORMDL3* | -7.25 | 2.27E-11 |
| | | *GSDMA* | -3.67 | 3.35E-04 |
| | | *GSDMB* | -6.98 | 9.54E-11 |
| | | *IKZF3* | 4.15 | 5.60E-05 |
| rs12232498 | g.38040363.C>T | *ORMDL3* | -7.99 | 3.76E-13 |
| | | *GSDMA* | -3.66 | 3.50E-04 |
| | | *GSDMB* | -7.01 | 8.13E-11 |
| | | *IKZF3* | 4.18 | 4.99E-05 |
| rs12941333 | g.38040534.T>C | *ORMDL3* | -7.92 | 5.68E-13 |
| | | *GSDMA* | -3.67 | 3.35E-04 |
| | | *GSDMB* | -6.98 | 9.54E-11 |
| | | *IKZF3* | 4.15 | 5.60E-05 |
| rs9908132 | g.38042777.A>T | *ORMDL3* | -7.99 | 3.76E-13 |
| | | *GSDMB* | -7.05 | 6.78E-11 |
| | | *IKZF3* | 5.07 | 1.20E-06 |
| rs9901146 | g.38043343.A>G | *ORMDL3* | -7.25 | 2.27E-11 |
| | | *GSDMB* | -7.05 | 6.78E-11 |
| | | *IKZF3* | 5.07 | 1.20E-06 |
| rs12936409 | g.38043649.T>C | *ORMDL3* | -7.25 | 2.27E-11 |
| | | *GSDMA* | -3.67 | 3.35E-04 |
| | | *GSDMB* | -6.98 | 9.54E-11 |
| | | *IKZF3* | 4.15 | 5.60E-05 |
| rs12103884 | g.38045725.T>C | *ORMDL3* | -7.99 | 3.76E-13 |
| | | *GSDMB* | -7.05 | 6.78E-11 |
| | | *IKZF3* | 5.07 | 1.20E-06 |
| rs9906951 | g.38048244.C>T | *ORMDL3* | -7.25 | 2.27E-11 |
| | | *GSDMB* | -7.05 | 6.78E-11 |

| | | | | |
|---|---|---|---|---|
| | | *IKZF3* | 5.07 | 1.20E-06 |
| | | *ORMDL3* | -7.25 | 2.27E-11 |
| rs12950209 | g.38049102.C>T | *GSDMB* | -7.05 | 6.78E-11 |
| | | *IKZF3* | 5.07 | 1.20E-06 |
| | | *ORMDL3* | -7.25 | 2.27E-11 |
| rs12950743 | g.38049233.C>T | *GSDMB* | -7.05 | 6.78E-11 |
| | | *IKZF3* | 5.07 | 1.20E-06 |
| | | *ORMDL3* | -7.25 | 2.27E-11 |
| rs7359623 | g.38049589.T>C | *GSDMB* | -7.08 | 5.80E-11 |
| | | *IKZF3* | 4.97 | 1.87E-06 |
| | | *ORMDL3* | -7.34 | 1.43E-11 |
| rs12453507 | g.38053207.G>C | *GSDMB* | -6.87 | 1.73E-10 |
| | | *IKZF3* | 4.88 | 2.73E-06 |
| | | *ORMDL3* | -7.37 | 1.17E-11 |
| rs11651596 | g.38056116.C>T | *GSDMA* | -3.78 | 2.26E-04 |
| | | *GSDMB* | -6.91 | 1.45E-10 |
| | | *IKZF3* | 4.17 | 5.17E-05 |
| | | *ORMDL3* | -8.09 | 2.16E-13 |
| rs12949100 | g.38057189.A>G | *GSDMA* | -3.70 | 3.06E-04 |
| | | *GSDMB* | -6.91 | 1.46E-10 |
| | | *IKZF3* | 3.98 | 1.09E-04 |
| | | *ORMDL3* | -8.05 | 2.77E-13 |
| rs8069176 | g.38057197.A>G | *GSDMA* | -3.70 | 3.06E-04 |
| | | *GSDMB* | -6.91 | 1.46E-10 |
| | | *IKZF3* | 3.98 | 1.09E-04 |
| | | *ORMDL3* | -8.05 | 2.77E-13 |
| rs11657449 | g.38057841.C>G | *GSDMA* | -5.01 | 1.56E-06 |
| | | *IKZF3* | 4.05 | 8.40E-05 |
| rs4795399 | g.38061439.C>T | *GSDMA* | -3.78 | 2.26E-04 |
| | | *GSDMB* | -6.91 | 1.45E-10 |
| | | *IKZF3* | 4.17 | 5.17E-05 |
| | | *ORMDL3* | -8.09 | 2.16E-13 |
| rs2305479 | g.38062217.A>G (Gly.304.Arg) | *GSDMB* | -6.96 | 1.10E-10 |

| | | | | |
|---|---|---|---|---|
| | | *IKZF3* | 5.09 | 1.09E-06 |
| | | *ORMDL3* | -7.31 | 1.66E-11 |
| rs883770 | g.38063381.T>C | *GSDMB* | -6.96 | 1.10E-10 |
| | | *IKZF3* | 5.09 | 1.09E-06 |
| | | *ORMDL3* | -7.31 | 1.66E-11 |
| rs62067034 | g.38063738.T>C | *GSDMB* | -6.96 | 1.10E-10 |
| | | *IKZF3* | 5.09 | 1.09E-06 |
| | | *ORMDL3* | -7.31 | 1.66E-11 |
| rs36000226 | g.38063929.C>T | *GSDMB* | -6.96 | 1.10E-10 |
| | | *IKZF3* | 5.09 | 1.09E-06 |
| | | *ORMDL3* | -7.31 | 1.66E-11 |
| rs36084703 | g.38063980_38063981del | *GSDMB* | -6.96 | 1.10E-10 |
| | | *IKZF3* | 5.09 | 1.09E-06 |
| | | *ORMDL3* | -7.31 | 1.66E-11 |
| rs11078927 | g.38064405.T>C | *GSDMA* | -3.78 | 2.26E-04 |
| | | *GSDMB* | -6.91 | 1.45E-10 |
| | | *IKZF3* | 4.17 | 5.17E-05 |
| | | *ORMDL3* | -8.09 | 2.16E-13 |
| rs11078928 | g.38064469.C>T (Leu.224.Leu) | *GSDMA* | -3.78 | 2.26E-04 |
| | | *GSDMB* | -6.91 | 1.45E-10 |
| | | *IKZF3* | 4.17 | 5.17E-05 |
| | | *ORMDL3* | -8.09 | 2.16E-13 |
| rs1008723 | g.38066267.T>G | *GSDMB* | -6.96 | 1.10E-10 |
| | | *IKZF3* | 5.09 | 1.09E-06 |
| | | *ORMDL3* | -7.31 | 1.66E-11 |
| rs56380902 | g.38066372.C>T | *GSDMB* | -6.96 | 1.10E-10 |
| | | *IKZF3* | 5.09 | 1.09E-06 |
| | | *ORMDL3* | -7.31 | 1.66E-11 |
| rs4795400 | g.38067020.T>C | *GSDMA* | -3.78 | 2.26E-04 |
| | | *GSDMB* | -6.91 | 1.45E-10 |
| | | *IKZF3* | 4.17 | 5.17E-05 |
| | | *ORMDL3* | -8.09 | 2.16E-13 |
| rs4795401 | g.38067533.G>A | *GSDMB* | -6.96 | 1.10E-10 |

| | | | | | |
|---|---|---|---|---|---|
| | | *IKZF3* | 5.09 | 1.09E-06 | |
| | | *ORMDL3* | -7.31 | 1.66E-11 | |
| rs869402 | g.38068043.T>C | *GSDMA* | -3.49 | 6.35E-04 | 5.35E-04 |
| | | *GSDMB* | -5.59 | 1.11E-07 | 5.35E-04 |
| | | *IKZF3* | 4.72 | 5.47E-06 | 5.35E-04 |
| | | *ORMDL3* | -6.21 | 5.20E-09 | 5.35E-04 |
| rs11078926 | g.38062976.A>G | *GSDMA* | -3.78 | 2.26E-04 | |
| | | *GSDMB* | -6.91 | 1.45E-10 | |
| | | *IKZF3* | 4.17 | 5.17E-05 | |
| | | *ORMDL3* | -8.09 | 2.16E-13 | |
| rs11557467 | g.38028634.T>G (Ser.173.Ile) | *GSDMB* | -7.05 | 6.78E-11 | |
| | | *IKZF3* | 5.07 | 1.20E-06 | |
| | | *ORMDL3* | -7.25 | 2.27E-11 | |
| rs11658278 | g.38031164.C>T | *GSDMB* | -7.05 | 6.78E-11 | |
| | | *IKZF3* | 5.07 | 1.20E-06 | |
| | | *ORMDL3* | -7.25 | 2.27E-11 | |
| rs12150079 | g.38025417.A>G | *GSDMA* | -4.87 | 2.93E-06 | |
| | | *IKZF3* | 4.02 | 9.20E-05 | |
| rs12939457 | g.38032188.C>T | *GSDMA* | -3.74 | 2.67E-04 | |
| | | *GSDMB* | -7.00 | 8.66E-11 | |
| | | *IKZF3* | 4.21 | 4.53E-05 | |
| | | *ORMDL3* | -8.01 | 3.52E-13 | |
| rs12939565 | g.38038389.T>A | *GSDMB* | -7.05 | 6.78E-11 | |
| | | *IKZF3* | 5.07 | 1.20E-06 | |
| | | *ORMDL3* | -7.25 | 2.27E-11 | |
| rs12939566 | g.38038390.T>A | *GSDMA* | -3.67 | 3.35E-04 | |
| | | *GSDMB* | -6.98 | 9.54E-11 | |
| | | *IKZF3* | 4.15 | 5.60E-05 | |
| | | *ORMDL3* | -7.99 | 3.76E-13 | |
| rs200216139 | g.38032132_38032132insC | *GSDMB* | -7.01 | 8.23E-11 | |
| | | *IKZF3* | 5.22 | 6.18E-07 | |
| | | *ORMDL3* | -7.23 | 2.49E-11 | |
| rs202126107 | g.38006767_38006776del | *GSDMA* | -4.00 | 9.90E-05 | |

| | | | | | |
|---|---|---|---|---|---|
| | | *GSDMB* | -7.05 | 6.69E-11 | |
| | | *IKZF3* | 3.94 | 1.27E-04 | |
| | | *ORMDL3* | -7.82 | 9.97E-13 | |
| rs2060941 | g.37982883.T>G | *GSDMB* | -7.29 | 1.82E-11 | |
| | | *ORMDL3* | -7.66 | 2.39E-12 | |
| rs2290400 | g.38066240.G>A | *GSDMB* | -6.96 | 1.10E-10 | |
| | | *IKZF3* | 5.09 | 1.09E-06 | |
| | | *ORMDL3* | -7.31 | 1.66E-11 | |
| rs2305480 | g.38062196.T>C (Pro.311.Ser) | *GSDMA* | -3.78 | 2.26E-04 | |
| | | *GSDMB* | -6.91 | 1.45E-10 | |
| | | *IKZF3* | 4.17 | 5.17E-05 | |
| | | *ORMDL3* | -8.09 | 2.16E-13 | |
| rs2872507 | g.38040763.A>G | *GSDMA* | -3.67 | 3.35E-04 | |
| | | *GSDMB* | -6.98 | 9.54E-11 | |
| | | *IKZF3* | 4.15 | 5.60E-05 | |
| | | *ORMDL3* | -7.99 | 3.76E-13 | |
| rs33938760 | | *GSDMB* | -7.17 | 3.61E-11 | |
| | | *IKZF3* | 4.92 | 2.31E-06 | |
| | | *ORMDL3* | -7.13 | 4.48E-11 | |
| rs34170568 | g.38055921_38055922ins | *GSDMA* | -4.32 | 2.91E-05 | |
| | | *GSDMB* | -5.05 | 1.28E-06 | |
| | | *IKZF3* | 4.38 | 2.23E-05 | |
| | | *ORMDL3* | -5.89 | 2.53E-08 | |
| rs35196450 | g.38062942_38062942insC | *GSDMA* | -3.78 | 2.26E-04 | |
| | | *GSDMB* | -6.91 | 1.45E-10 | |
| | | *IKZF3* | 4.17 | 5.17E-05 | |
| | | *ORMDL3* | -8.09 | 2.16E-13 | |
| rs35503505 | g.38057780_38057783del | *GSDMA* | -4.99 | 1.68E-06 | |
| | | *IKZF3* | 3.95 | 1.20E-04 | |
| rs36095411 | g.38031865.G>T | *GSDMA* | -3.58 | 4.63E-04 | 7.00E-04 |
| | | *GSDMB* | -6.90 | 1.48E-10 | 7.00E-04 |
| | | *IKZF3* | 4.21 | 4.54E-05 | 7.00E-04 |
| | | *ORMDL3* | -8.06 | 2.57E-13 | 7.00E-04 |

| | | | | | | |
|---|---|---|---|---|---|---|
| | rs367998020 | g.38032200_38032217del | *GSDMA* | -3.74 | 2.67E-04 | |
| | | | *GSDMB* | -7.00 | 8.66E-11 | |
| | | | *IKZF3* | 4.21 | 4.53E-05 | |
| | | | *ORMDL3* | -8.01 | 3.52E-13 | |
| | rs56750287 | g.38062944.C>A | *GSDMA* | -3.78 | 2.26E-04 | |
| | | | *GSDMB* | -6.91 | 1.45E-10 | |
| | | | *IKZF3* | 4.17 | 5.17E-05 | |
| | | | *ORMDL3* | -8.09 | 2.16E-13 | |
| | rs68122720 | | *GSDMA* | -3.67 | 3.35E-04 | |
| | | | *GSDMB* | -6.98 | 9.54E-11 | |
| | | | *IKZF3* | 4.15 | 5.60E-05 | |
| | | | *ORMDL3* | -7.99 | 3.76E-13 | |
| | rs8067378 | g.38051348.G>A | *GSDMB* | -7.05 | 6.78E-11 | |
| | | | *IKZF3* | 5.07 | 1.20E-06 | |
| | | | *ORMDL3* | -7.25 | 2.27E-11 | |
| | rs9303277 | g.37976469.T>C | *GSDMB* | -6.90 | 1.46E-10 | |
| | | | *IKZF3* | 4.64 | 7.76E-06 | |
| | | | *ORMDL3* | -6.88 | 1.71E-10 | |
| | rs9905959 | g.38031138.G>A | *GSDMA* | -3.67 | 3.35E-04 | |
| | | | *GSDMB* | -6.98 | 9.54E-11 | |
| | | | *IKZF3* | 4.15 | 5.60E-05 | |
| | | | *ORMDL3* | -7.99 | 3.76E-13 | |
| Fifth block | rs870829 | g.38068382.T>G | *GSDMA* | -3.96 | 1.16E-04 | 7.23E-04 |
| | | | *IKZF3* | 3.93 | 1.34E-04 | 7.23E-04 |
| | rs1011082 | g.38068514.A>G | *GSDMA* | -3.46 | 7.16E-04 | 7.66E-04 |
| | | | *GSDMB* | -5.48 | 1.80E-07 | 7.66E-04 |
| | | | *IKZF3* | 4.78 | 4.24E-06 | 7.66E-04 |
| | | | *ORMDL3* | -6.15 | 7.29E-09 | 7.66E-04 |
| | rs921650 | g.38069076.C>T | *GSDMA* | -3.49 | 6.35E-04 | 5.34E-04 |
| | | | *GSDMB* | -5.59 | 1.11E-07 | 5.34E-04 |
| | | | *IKZF3* | 4.72 | 5.47E-06 | 5.34E-04 |
| | | | *ORMDL3* | -6.21 | 5.20E-09 | 5.34E-04 |
| | rs921649 | g.38069274.G>A | *GSDMA* | -3.49 | 6.35E-04 | 5.34E-04 |
| | | | *GSDMB* | -5.59 | 1.11E-07 | 5.34E-04 |

| | | | | | |
|---|---|---|---|---|---|
| | | *IKZF3* | 4.72 | 5.47E-06 | 5.34E-04 |
| | | *ORMDL3* | -6.21 | 5.20E-09 | 5.34E-04 |
| rs5820308 | g.38069364_38069369del | *GSDMA* | -4.93 | 2.21E-06 | 8.13E-04 |
| | | *GSDMB* | -5.37 | 3.08E-07 | 8.13E-04 |
| | | *IKZF3* | 3.76 | 2.44E-04 | 8.13E-04 |
| | | *ORMDL3* | -6.69 | 4.49E-10 | 8.13E-04 |
| rs6503524 | g.38069809.C>T | *GSDMA* | -3.49 | 6.35E-04 | 5.34E-04 |
| | | *GSDMB* | -5.59 | 1.11E-07 | 5.34E-04 |
| | | *IKZF3* | 4.72 | 5.47E-06 | 5.34E-04 |
| | | *ORMDL3* | -6.21 | 5.20E-09 | 5.34E-04 |
| rs7216558 | g.38070071.C>T | *GSDMA* | -3.49 | 6.35E-04 | 5.34E-04 |
| | | *GSDMB* | -5.59 | 1.11E-07 | 5.34E-04 |
| | | *IKZF3* | 4.72 | 5.47E-06 | 5.34E-04 |
| | | *ORMDL3* | -6.21 | 5.20E-09 | 5.34E-04 |
| rs7221605 | g.38070789.C>T | *GSDMA* | -3.55 | 5.16E-04 | |
| | | *GSDMB* | -5.53 | 1.47E-07 | |
| | | *IKZF3* | 4.82 | 3.56E-06 | |
| | | *ORMDL3* | -6.08 | 1.04E-08 | |
| rs1031458 | g.38072173.G>T | *GSDMA* | -3.55 | 5.16E-04 | |
| | | *GSDMB* | -5.53 | 1.47E-07 | |
| | | *IKZF3* | 4.82 | 3.56E-06 | |
| | | *ORMDL3* | -6.08 | 1.04E-08 | |
| rs1031459 | g.38072245.C>G | *GSDMA* | -5.86 | 2.94E-08 | |
| | | *IKZF3* | 3.58 | 4.73E-04 | |
| rs1031460 | g.38072247.T>G | *GSDMA* | -3.55 | 5.16E-04 | |
| | | *GSDMB* | -5.53 | 1.47E-07 | |
| | | *IKZF3* | 4.82 | 3.56E-06 | |
| | | *ORMDL3* | -6.08 | 1.04E-08 | |
| rs8065777 | g.38072402.C>T | *GSDMA* | -3.55 | 5.16E-04 | |
| | | *GSDMB* | -5.53 | 1.47E-07 | |
| | | *IKZF3* | 4.82 | 3.56E-06 | |
| | | *ORMDL3* | -6.08 | 1.04E-08 | |
| rs2872516 | g.38072727.C>T | *GSDMA* | -5.01 | 1.57E-06 | |
| | | *GSDMB* | -5.32 | 3.90E-07 | |

| | | | | | |
|---|---|---|---|---|---|
| | | *IKZF3* | 3.87 | 1.65E-04 | |
| | | *ORMDL3* | -6.56 | 8.94E-10 | |
| rs9303279 | g.38073968.G>C | *GSDMA* | -4.93 | 2.21E-06 | 6.13E-04 |
| | | *GSDMB* | -5.37 | 3.08E-07 | 6.13E-04 |
| | | *IKZF3* | 3.76 | 2.44E-04 | 6.13E-04 |
| | | *ORMDL3* | -6.69 | 4.49E-10 | 6.13E-04 |
| rs9303280 | g.38074031.T>C | *GSDMA* | -3.55 | 5.16E-04 | 8.49E-04 |
| | | *GSDMB* | -5.53 | 1.47E-07 | 8.49E-04 |
| | | *IKZF3* | 4.82 | 3.56E-06 | 8.49E-04 |
| | | *ORMDL3* | -6.08 | 1.04E-08 | 8.49E-04 |
| rs9303281 | g.38074046.G>A | *GSDMA* | -3.49 | 6.35E-04 | 2.60E-04 |
| | | *GSDMB* | -5.59 | 1.11E-07 | 2.60E-04 |
| | | *IKZF3* | 4.72 | 5.47E-06 | 2.60E-04 |
| | | *ORMDL3* | -6.21 | 5.20E-09 | 2.60E-04 |
| rs7219923 | g.38074518.C>T | *GSDMA* | -3.49 | 6.35E-04 | 2.75E-04 |
| | | *GSDMB* | -5.59 | 1.11E-07 | 2.75E-04 |
| | | *IKZF3* | 4.72 | 5.47E-06 | 2.75E-04 |
| | | *ORMDL3* | -6.21 | 5.20E-09 | 2.75E-04 |
| rs7224129 | g.38075426.G>A | *GSDMA* | -3.44 | 7.48E-04 | 3.47E-04 |
| | | *GSDMB* | -5.51 | 1.59E-07 | 3.47E-04 |
| | | *IKZF3* | 4.65 | 7.55E-06 | 3.47E-04 |
| | | *ORMDL3* | -6.08 | 1.00E-08 | 3.47E-04 |
| rs8074437 | g.38076137.T>G | *GSDMA* | -3.44 | 7.48E-04 | 3.47E-04 |
| | | *GSDMB* | -5.51 | 1.59E-07 | 3.47E-04 |
| | | *IKZF3* | 4.65 | 7.55E-06 | 3.47E-04 |
| | | *ORMDL3* | -6.08 | 1.00E-08 | 3.47E-04 |
| rs71971950 | g.38076198_38076201del | *GSDMA* | -3.66 | 3.58E-04 | 3.77E-04 |
| | | *GSDMB* | -5.52 | 1.50E-07 | 3.77E-04 |
| | | *IKZF3* | 4.45 | 1.68E-05 | 3.77E-04 |
| | | *ORMDL3* | -6.13 | 7.85E-09 | 3.77E-04 |
| rs4065275 | g.38080865.A>G | *GSDMA* | -3.84 | 1.84E-04 | 8.19E-04 |
| | | *GSDMB* | -5.61 | 1.00E-07 | 8.19E-04 |
| | | *IKZF3* | 4.85 | 3.17E-06 | 8.19E-04 |
| | | *ORMDL3* | -6.17 | 6.43E-09 | 8.19E-04 |

| | | | | | | |
|---|---|---|---|---|---|---|
| | rs8076131 | g.38080912.G>A | *GSDMA* | -5.26 | 5.06E-07 | 5.13E-04 |
| | | | *GSDMB* | -5.40 | 2.69E-07 | 5.13E-04 |
| | | | *IKZF3* | 3.75 | 2.57E-04 | 5.13E-04 |
| | | | *ORMDL3* | -6.66 | 5.17E-10 | 5.13E-04 |
| | rs12603332 | g.38082807.T>C | *GSDMA* | -3.82 | 1.98E-04 | 4.23E-04 |
| | | | *GSDMB* | -5.61 | 9.86E-08 | 4.23E-04 |
| | | | *IKZF3* | 4.83 | 3.42E-06 | 4.23E-04 |
| | | | *ORMDL3* | -6.17 | 6.52E-09 | 4.23E-04 |
| | rs4795403 | g.38085722.T>C | *GSDMB* | -6.95 | 1.13E-10 | |
| | | | *ORMDL3* | -7.21 | 2.85E-11 | |
| | rs4795404 | g.38085791.A>C | *GSDMB* | -6.95 | 1.13E-10 | |
| | | | *ORMDL3* | -7.21 | 2.85E-11 | |
| | rs7224908 | g.38086854.A>G | *GSDMB* | -7.06 | 6.38E-11 | |
| | | | *ORMDL3* | -7.24 | 2.41E-11 | |
| | rs12946393 | g.38087429.T>G | *GSDMB* | -7.06 | 6.38E-11 | |
| | | | *ORMDL3* | -7.24 | 2.41E-11 | |
| | rs143385463 | g.38071855_38071858del | *GSDMA* | -3.55 | 5.16E-04 | |
| | | | *GSDMB* | -5.53 | 1.47E-07 | |
| | | | *IKZF3* | 4.82 | 3.56E-06 | |
| | | | *ORMDL3* | -6.08 | 1.04E-08 | |
| | rs150597688 | g.38071086_38071086insTT | *GSDMA* | -3.55 | 5.16E-04 | |
| | | | *GSDMB* | -5.53 | 1.47E-07 | |
| | | | *IKZF3* | 4.82 | 3.56E-06 | |
| | | | *ORMDL3* | -6.08 | 1.04E-08 | |
| | rs3744246 | g.38084350.T>C | *GSDMB* | -6.95 | 1.13E-10 | |
| | | | *ORMDL3* | -7.21 | 2.85E-11 | |
| | rs4795402 | g.38085385.A>C | *GSDMB* | -6.74 | 3.50E-10 | |
| | | | *ORMDL3* | -6.24 | 4.50E-09 | |
| | rs7216389 | g.38069949.C>T | *GSDMA* | -3.49 | 6.35E-04 | 5.34E-04 |
| | | | *GSDMB* | -5.59 | 1.11E-07 | 5.34E-04 |
| | | | *IKZF3* | 4.72 | 5.47E-06 | 5.34E-04 |
| | | | *ORMDL3* | -6.21 | 5.20E-09 | 5.34E-04 |
| Sixth block | rs56199421 | g.38090808.C>T | *GSDMA* | -8.35 | 4.86E-14 | |
| | rs8065244 | g.38091228.G>C | *GSDMA* | 5.40 | 2.61E-07 | |

| | | | | |
|---|---|---|---|---|
| rs72832957 | g.38093085.G>A | *GSDMA* | 5.32 | 3.85E-07 |
| rs12601749 | g.38093315.A>G | *GSDMA* | -8.97 | 1.40E-15 |
| rs12603481 | g.38093339.A>G | *GSDMA* | -4.92 | 2.37E-06 |
| rs72832958 | g.38093881.A>C | *GSDMA* | 5.32 | 3.85E-07 |
| rs113778191 | g.38094183.T>G | *GSDMA* | 5.32 | 3.85E-07 |
| rs6503525 | g.38095174.C>G | *GSDMA* | -4.92 | 2.37E-06 |
| rs72832962 | g.38096501.C>A | *GSDMA* | 5.32 | 3.85E-07 |
| rs72832964 | g.38097008.C>T | *GSDMA* | 5.32 | 3.85E-07 |
| rs72832965 | g.38097143.C>T | *GSDMA* | 5.32 | 3.85E-07 |
| rs7216564 | g.38097172.A>T | *GSDMA* | -4.92 | 2.37E-06 |
| rs79785426 | g.38098027.A>G | *GSDMA* | 5.32 | 3.85E-07 |
| rs144007425 | g.38098781_38098781insG | *GSDMA* | -4.92 | 2.37E-06 |
| rs8065126 | g.38099035.T>C | *GSDMA* | -7.69 | 2.00E-12 |
| rs72832966 | g.38100129.A>G | *GSDMA* | 5.32 | 3.85E-07 |
| rs4795406 | g.38100134.C>G | *GSDMA* | -4.92 | 2.37E-06 |
| rs74717022 | g.38100633.T>C (Ser.158.Ser) | *GSDMA* | 5.32 | 3.85E-07 |
| rs3848395 | g.38101468.T>C | *GSDMA* | -7.69 | 2.00E-12 |
| rs4065985 | g.38101932.G>C | *GSDMA* | -8.97 | 1.40E-15 |
| rs3893044 | g.38103016.A>G | *GSDMA* | -8.97 | 1.40E-15 |
| rs62068170 | g.38103210.G>A | *GSDMA* | -8.97 | 1.40E-15 |
| rs62068171 | g.38103242.G>A | *GSDMA* | -8.97 | 1.40E-15 |
| rs8080734 | g.38103285.G>A | *GSDMA* | -9.16 | 4.67E-16 |
| rs8071050 | g.38106599.A>G | *GSDMA* | -4.14 | 5.96E-05 |
| rs4795408 | g.38107627.A>G | *GSDMA* | -4.92 | 2.37E-06 |
| rs10589831 | g.38108023_38108025del | *GSDMA* | -8.19 | 1.25E-13 |
| rs9889716 | g.38108298.A>G | *GSDMA* | -7.96 | 4.47E-13 |
| rs9895948 | g.38108363.T>C | *GSDMA* | -8.19 | 1.25E-13 |
| rs7223318 | g.38108553.G>C | *GSDMA* | -8.19 | 1.25E-13 |
| rs72832971 | g.38108619.T>C | *GSDMA* | 5.39 | 2.79E-07 |
| rs7209742 | g.38108708.A>G | *GSDMA* | -8.19 | 1.25E-13 |
| rs28618095 | g.38109075.T>C | *GSDMA* | -7.93 | 5.32E-13 |
| rs56301252 | g.38109155.T>G | *GSDMA* | -8.19 | 1.25E-13 |
| rs76137456 | g.38109590.G>C | *GSDMA* | 5.60 | 1.03E-07 |

| | | | | |
|---|---|---|---|---|
| rs60667221 | g.38110390.T>A | *GSDMA* | -8.35 | 5.07E-14 |
| rs17609240 | g.38110689.T>G | *GSDMA* | -8.35 | 5.07E-14 |
| rs8076474 | g.38111234.G>C | *GSDMA* | -8.35 | 5.07E-14 |
| rs1007655 | g.38111419.G>A | *GSDMA* | -8.35 | 5.07E-14 |
| rs1563103 | g.38111740.G>A | *GSDMA* | -8.35 | 5.07E-14 |
| rs2313640 | g.38111845.G>A | *GSDMA* | -8.35 | 5.07E-14 |
| rs8068522 | g.38112076.T>C | *GSDMA* | -8.32 | 5.95E-14 |
| rs8081437 | g.38112114.G>A | *GSDMA* | -8.35 | 5.07E-14 |
| rs8081462 | g.38112190.C>G | *GSDMA* | -4.92 | 2.37E-06 |
| rs62068174 | g.38112255.G>C | *GSDMA* | -8.35 | 5.07E-14 |
| rs62068175 | g.38112300.A>C | *GSDMA* | -8.35 | 5.07E-14 |
| rs62068176 | g.38112438.A>G | *GSDMA* | -8.41 | 3.45E-14 |
| rs62068177 | g.38112601.G>A | *GSDMA* | -6.49 | 1.30E-09 |
| rs62068178 | g.38112608.C>T | *GSDMA* | -6.49 | 1.30E-09 |
| rs62068179 | g.38112617.G>A | *GSDMA* | -6.49 | 1.30E-09 |
| rs67480438 | g.38112774.A>G | *GSDMA* | -8.33 | 5.44E-14 |
| rs62068182 | g.38112864.G>A | *GSDMA* | -8.33 | 5.44E-14 |
| rs62068184 | g.38113054.C>T | *GSDMA* | -8.35 | 5.07E-14 |
| rs7218742 | g.38114361.A>G | *GSDMA* | -8.35 | 5.07E-14 |
| rs7218321 | g.38114469.C>T | *GSDMA* | -8.35 | 5.07E-14 |
| rs6503526 | g.38114598.T>C | *GSDMA* | -4.71 | 5.82E-06 |
| rs6503527 | g.38114719.G>A | *GSDMA* | -8.35 | 5.07E-14 |
| rs3931960 | g.38114977.G>C | *GSDMA* | -4.14 | 5.96E-05 |
| rs7223717 | g.38115333.C>A | *GSDMA* | -4.14 | 5.96E-05 |
| rs10693935 | g.38115429_38115429insTG | *GSDMA* | -4.14 | 5.96E-05 |
| rs112979959 | g.38115873.G>T | *GSDMA* | 5.60 | 1.03E-07 |
| rs112570995 | g.38116212.A>C | *GSDMA* | 5.60 | 1.03E-07 |
| rs111522131 | g.38116225.G>A | *GSDMA* | 5.60 | 1.03E-07 |
| rs11424020 | g.38116339_38116339insT | *GSDMA* | -7.83 | 9.21E-13 |
| rs11869855 | g.38117653.A>G | *GSDMA* | -7.82 | 9.82E-13 |
| rs113510790 | g.38118010.T>C | *GSDMA* | 5.65 | 8.28E-08 |
| rs564455927 | g.38118235_38118235insA | *GSDMA* | -4.91 | 2.41E-06 |
| rs72832987 | g.38118468.C>A | *GSDMA* | 5.60 | 1.03E-07 |
| rs72832988 | g.38118541.A>G | *GSDMA* | 5.60 | 1.03E-07 |

| | | | | |
|---|---|---|---|---|
| rs72832989 | g.38118752.T>C | *GSDMA* | 5.60 | 1.03E-07 |
| rs72832991 | g.38119448.T>C | *GSDMA* | 5.60 | 1.03E-07 |
| rs3902024 | g.38119548.T>C | *GSDMA* | -7.82 | 9.82E-13 |
| rs55739615 | g.38119638.C>T | *GSDMA* | -9.41 | 1.05E-16 |
| rs78295195 | g.38119663.G>C | *GSDMA* | 5.60 | 1.03E-07 |
| rs56396280 | g.38119708.A>C | *GSDMA* | -7.82 | 9.82E-13 |
| rs12451084 | g.38119757.T>G | *GSDMA* | -4.92 | 2.37E-06 |
| rs12451100 | g.38119831.T>G | *GSDMA* | -4.92 | 2.37E-06 |
| rs111690387 | g.38119882.A>G | *GSDMA* | 5.60 | 1.03E-07 |
| rs60701125 | g.38120179.T>C | *GSDMA* | 5.60 | 1.03E-07 |
| rs2001476 | g.38120604.T>C | *GSDMA* | -7.82 | 9.82E-13 |
| rs4795409 | g.38120736.T>C | *GSDMA* | -4.92 | 2.37E-06 |
| rs142218801 | g.38120827_38120829del | *GSDMA* | 5.60 | 1.03E-07 |
| rs4458030 | g.38121706.C>T | *GSDMA* | -7.60 | 3.44E-12 |
| rs8069202 | g.38122200.A>G | *GSDMA* | -4.66 | 7.24E-06 |
| rs1007654 | g.38111354.A>G | *GSDMA* | -8.35 | 5.07E-14 |
| rs11399056 | g.38114046_38114046insA | *GSDMA* | -7.35 | 1.32E-11 |
| rs35123741 | g.38092930.G>A | *GSDMA* | -4.92 | 2.37E-06 |
| rs3894194 | g.38121993.T>C (Arg.18.Gln) | *GSDMA* | -4.66 | 7.24E-06 |
| rs3902025 | g.38119254.C>A | *GSDMA* | -9.41 | 1.05E-16 |
| rs4065986 | g.38102641.A>G | *GSDMA* | -4.92 | 2.37E-06 |
| rs4134417 | g.38113278.G>A | *GSDMA* | -8.32 | 6.07E-14 |
| rs4134498 | g.38113282.G>A | *GSDMA* | -8.32 | 6.07E-14 |
| rs4134499 | g.38113274.G>A | *GSDMA* | -8.32 | 6.07E-14 |
| rs4134500 | g.38113270.G>A | *GSDMA* | -8.32 | 6.07E-14 |
| rs4134501 | g.38113290.G>A | *GSDMA* | -8.32 | 6.07E-14 |
| rs4134502 | g.38113286.G>A | *GSDMA* | -8.32 | 6.07E-14 |
| rs4795405 | g.38088417.T>C | *GSDMA* | -5.14 | 8.74E-07 |
| | | *GSDMB* | -5.51 | 1.57E-07 |
| | | *IKZF3* | 4.21 | 4.48E-05 |
| | | *ORMDL3* | -6.68 | 4.78E-10 |
| rs55927420 | g.38109251.G>T | *GSDMA* | -8.19 | 1.25E-13 |
| rs56340811 | g.38115430.A>C | *GSDMA* | -4.14 | 5.96E-05 |

| | | | | | |
|---|---|---|---|---|---|
| | rs56410675 | g.38109254.A>G | *GSDMA* | -8.19 | 1.25E-13 |
| | rs62068180 | g.38112825.T>C | *GSDMA* | -8.33 | 5.44E-14 |
| | rs62068181 | g.38112832.T>C | *GSDMA* | -8.33 | 5.44E-14 |
| | rs62068183 | g.38112918.C>T | *GSDMA* | -8.35 | 5.07E-14 |
| | rs7219080 | g.38114516.A>C | *GSDMA* | -8.35 | 5.07E-14 |
| | rs140941705 | g.38115333_38115333insA | *GSDMA* | 5.60 | 1.03E-07 |
| | rs75541765 | g.38096881.C>T | *GSDMA* | 5.32 | 3.85E-07 |
| | rs8079416 | g.38092713.C>T | *GSDMA* | -4.92 | 2.37E-06 |
| Seventh block | rs4239225 | g.38127112.T>C | *GSDMA* | -4.35 | 2.60E-05 |
| | rs74727658 | g.38128181_38128182del | *GSDMA* | -3.92 | 1.37E-04 |
| | rs3859191 | g.38128714.A>G | *GSDMA* | -4.35 | 2.59E-05 |
| | rs8077456 | g.38128765.C>G | *GSDMA* | -3.92 | 1.37E-04 |
| | rs4065876 | g.38129506.A>G | *GSDMA* | -4.23 | 4.15E-05 |
| | rs748908389 | g.38129525_38129525ins | *GSDMA* | -4.17 | 5.31E-05 |
| | rs11870683 | g.38129841.A>T | *GSDMA* | -3.66 | 3.47E-04 |
| | rs60137005 | g.38129996.T>A | *GSDMA* | -4.23 | 4.15E-05 |
| | rs56326707 | g.38130139.T>C | *GSDMA* | -4.23 | 4.15E-05 |
| | rs56030650 | g.38131187.A>C (Thr.314.Asn) | *GSDMA* | -4.23 | 4.15E-05 |
| | rs60134943 | g.38133792.T>G | *GSDMA* | -4.23 | 4.15E-05 |
| | rs139141843 | g.38133914_38133914insA | *GSDMA* | -3.67 | 3.39E-04 |
| | rs556061928 | g.38126911_38126919del | *GSDMA* | -3.92 | 1.37E-04 |
| | rs3859192 | g.38128648.T>C | *GSDMA* | -4.35 | 2.59E-05 |
| NA | rs407307 | g.37827163.G>A | *PGAP3* | -3.89 | 1.52E-04 |
| | rs118036166 | g.37860994.G>A | *PGAP3* | -3.64 | 3.81E-04 |
| | rs79026872 | g.37965932.C>T | *PGAP3* | -3.62 | 4.04E-04 |
| | rs17608925 | g.38082831.C>T | *GSDMB* | -13.28 | 7.86E-27 |
| | | | *ORMDL3* | -14.63 | 2.39E-30 |
| | rs373861118 | g.38118929_38118930insCACACACACACA | *GSDMA* | -6.24 | 4.60E-09 |
| | rs149274005 | g.38129546.G>A | *GSDMB* | -4.31 | 2.98E-05 |
| | | | *ORMDL3* | -5.18 | 7.35E-07 |
| | rs12946335 | g.38133764.T>C | *GSDMB* | -4.31 | 2.98E-05 |
| | | | *ORMDL3* | -5.18 | 7.35E-07 |

| | | | | |
|---|---|---|---|---|
| rs59132767 | g.38134300_38134301del | *GSDMA* | -4.23 | 4.15E-05 |

[a] P values for the association between the SNPs included in the eQTL and asthma. Ps are only shown if significant (P <0.001).
