## Supplemental Table 10 for "*GSDMA* drives the most replicated association with asthma in naïve CD4^+^ T cells"

**Table S10:** Significant mQTLs in naïve CD4$^+$ T cells

| Haplotype block | SNP id | HGVS names (hg19) | Position CpG (hg19) | Gene is methylated (1.5 kb from TSS)[a] | Gene is methylated (gene body)[a] | Statistic | P mQTL | $P_{SNP}$ asthma[b] | $FDR_{CpG}$ asthma[c] |
|---|---|---|---|---|---|---|---|---|---|
| First block | rs14050 | g.37828072.G>A | 37,828,195 | | *PGAP3* | -5.016 | 1.70E-06 | | |
| | | | 37,828,197 | | *PGAP3* | -4.142 | 6.15E-05 | | |
| | | | 37,828,220 | | *PGAP3* | -4.815 | 4.03E-06 | | |
| | | | 37,828,273 | | *PGAP3* | -4.350 | 2.72E-05 | | 0.010 |
| | | | 37,828,283 | | *PGAP3* | -3.941 | 1.32E-04 | | 0.010 |
| | | | 37,828,296 | | *PGAP3* | -5.849 | 3.79E-08 | | 0.002 |
| | | | 37,828,338 | | *PGAP3* | -5.116 | 1.09E-06 | | 0.002 |
| | | | 37,828,377 | | *PGAP3* | -6.790 | 3.60E-10 | | 0.002 |
| | | | 37,828,384 | | *PGAP3* | -6.144 | 9.12E-09 | | 0.002 |
| | | | 37,828,560 | | *PGAP3* | -4.411 | 2.14E-05 | | 0.002 |
| | | | 37,843,658 | *ERBB2* | *PGAP3* | -3.662 | 3.62E-04 | | 7.82E-05 |
| | | | 38,072,514 | | *GSDMB* | -3.691 | 3.27E-04 | | 0.046 |
| | | | 38,077,039 | | | -3.551 | 5.36E-04 | | 0.034 |
| | | | 38,082,022 | | *ORMDL3* | -3.901 | 1.53E-04 | | 6.40E-04 |
| | rs2952151 | g.37828496.T>C | 37,828,195 | | *PGAP3* | -5.016 | 1.70E-06 | | |
| | | | 37,828,197 | | *PGAP3* | -4.142 | 6.15E-05 | | |
| | | | 37,828,220 | | *PGAP3* | -4.815 | 4.03E-06 | | |
| | | | 37,828,273 | | *PGAP3* | -4.350 | 2.72E-05 | | 0.010 |
| | | | 37,828,283 | | *PGAP3* | -3.941 | 1.32E-04 | | 0.010 |
| | | | 37,828,296 | | *PGAP3* | -5.849 | 3.79E-08 | | 0.002 |
| | | | 37,828,338 | | *PGAP3* | -5.116 | 1.09E-06 | | 0.002 |
| | | | 37,828,377 | | *PGAP3* | -6.790 | 3.60E-10 | | 0.002 |
| | | | 37,828,384 | | *PGAP3* | -6.144 | 9.12E-09 | | 0.002 |
| | | | 37,828,560 | | *PGAP3* | -4.411 | 2.14E-05 | | 0.002 |
| | | | 37,843,658 | *ERBB2* | *PGAP3* | -3.662 | 3.62E-04 | | 7.82E-05 |
| | | | 38,072,514 | | *GSDMB* | -3.691 | 3.27E-04 | | 0.046 |
| | | | 38,077,039 | | | -3.551 | 5.36E-04 | | 0.034 |
| | | | 38,082,022 | | *ORMDL3* | -3.901 | 1.53E-04 | | 6.40E-04 |
| | rs2941503 | g.37828745.A>G | 37,828,195 | | *PGAP3* | -5.016 | 1.70E-06 | | |
| | | | 37,828,197 | | *PGAP3* | -4.142 | 6.15E-05 | | |
| | | | 37,828,220 | | *PGAP3* | -4.815 | 4.03E-06 | | |
| | | | 37,828,273 | | *PGAP3* | -4.350 | 2.72E-05 | | 0.010 |
| | | | 37,828,283 | | *PGAP3* | -3.941 | 1.32E-04 | | 0.010 |
| | | | 37,828,296 | | *PGAP3* | -5.849 | 3.79E-08 | | 0.002 |
| | | | 37,828,338 | | *PGAP3* | -5.116 | 1.09E-06 | | 0.002 |
| | | | 37,828,377 | | *PGAP3* | -6.790 | 3.60E-10 | | 0.002 |

| | | | | | | | |
|---|---|---|---|---|---|---|---|
| | | 37,828,384 | | *PGAP3* | -6.144 | 9.12E-09 | 0.002 |
| | | 37,828,560 | | *PGAP3* | -4.411 | 2.14E-05 | 0.002 |
| | | 37,843,658 | *ERBB2* | *PGAP3* | -3.662 | 3.62E-04 | 7.82E-05 |
| | | 38,072,514 | | *GSDMB* | -3.691 | 3.27E-04 | 0.046 |
| | | 38,077,039 | | | -3.551 | 5.36E-04 | 0.034 |
| | | 38,082,022 | | *ORMDL3* | -3.901 | 1.53E-04 | 6.40E-04 |
| rs907087 | g.37828787.G>A | 37,828,195 | | *PGAP3* | -5.016 | 1.70E-06 | |
| | | 37,828,197 | | *PGAP3* | -4.142 | 6.15E-05 | |
| | | 37,828,220 | | *PGAP3* | -4.815 | 4.03E-06 | |
| | | 37,828,273 | | *PGAP3* | -4.350 | 2.72E-05 | 0.010 |
| | | 37,828,283 | | *PGAP3* | -3.941 | 1.32E-04 | 0.010 |
| | | 37,828,296 | | *PGAP3* | -5.849 | 3.79E-08 | 0.002 |
| | | 37,828,338 | | *PGAP3* | -5.116 | 1.09E-06 | 0.002 |
| | | 37,828,377 | | *PGAP3* | -6.790 | 3.60E-10 | 0.002 |
| | | 37,828,384 | | *PGAP3* | -6.144 | 9.12E-09 | 0.002 |
| | | 37,828,560 | | *PGAP3* | -4.411 | 2.14E-05 | 0.002 |
| | | 37,843,658 | *ERBB2* | *PGAP3* | -3.662 | 3.62E-04 | 7.82E-05 |
| | | 38,072,514 | | *GSDMB* | -3.691 | 3.27E-04 | 0.046 |
| | | 38,077,039 | | | -3.551 | 5.36E-04 | 0.034 |
| | | 38,082,022 | | *ORMDL3* | -3.901 | 1.53E-04 | 6.40E-04 |
| rs2247862 | g.37829129.T>C | 37,828,195 | | *PGAP3* | -5.016 | 1.70E-06 | |
| | | 37,828,197 | | *PGAP3* | -4.142 | 6.15E-05 | |
| | | 37,828,220 | | *PGAP3* | -4.815 | 4.03E-06 | |
| | | 37,828,273 | | *PGAP3* | -4.350 | 2.72E-05 | 0.010 |
| | | 37,828,283 | | *PGAP3* | -3.941 | 1.32E-04 | 0.010 |
| | | 37,828,296 | | *PGAP3* | -5.849 | 3.79E-08 | 0.002 |
| | | 37,828,338 | | *PGAP3* | -5.116 | 1.09E-06 | 0.002 |
| | | 37,828,377 | | *PGAP3* | -6.790 | 3.60E-10 | 0.002 |
| | | 37,828,384 | | *PGAP3* | -6.144 | 9.12E-09 | 0.002 |
| | | 37,828,560 | | *PGAP3* | -4.411 | 2.14E-05 | 0.002 |
| | | 37,843,658 | *ERBB2* | *PGAP3* | -3.662 | 3.62E-04 | 7.82E-05 |
| | | 38,072,514 | | *GSDMB* | -3.691 | 3.27E-04 | 0.046 |
| | | 38,077,039 | | | -3.551 | 5.36E-04 | 0.034 |
| | | 38,082,022 | | *ORMDL3* | -3.901 | 1.53E-04 | 6.40E-04 |
| rs903504 | g.37829570.C>G | 37,828,195 | | *PGAP3* | -5.000 | 1.82E-06 | |
| | | 37,828,197 | | *PGAP3* | -4.400 | 2.23E-05 | |
| | | 37,828,220 | | *PGAP3* | -4.897 | 2.84E-06 | |
| | | 37,828,273 | | *PGAP3* | -4.687 | 6.89E-06 | 0.010 |
| | | 37,828,283 | | *PGAP3* | -4.018 | 9.88E-05 | 0.010 |
| | | 37,828,296 | | *PGAP3* | -5.643 | 1.00E-07 | 0.002 |
| | | 37,828,338 | | *PGAP3* | -5.249 | 6.07E-07 | 0.002 |

| | | | | | | | |
|---|---|---|---|---|---|---|---|
| | | 37,828,377 | | *PGAP3* | -6.797 | 3.48E-10 | 0.002 |
| | | 37,828,384 | | *PGAP3* | -6.071 | 1.30E-08 | 0.002 |
| | | 37,828,560 | | *PGAP3* | -4.337 | 2.87E-05 | 0.002 |
| | | 37,843,658 | *ERBB2* | *PGAP3* | -4.429 | 1.99E-05 | 7.82E-05 |
| | | 37,844,480 | *PGAP3* | *ERBB2* | 3.999 | 1.06E-04 | 7.82E-05 |
| | | 38,072,514 | | *GSDMB* | -3.537 | 5.61E-04 | 0.046 |
| | | 38,082,022 | | *ORMDL3* | -3.993 | 1.09E-04 | 6.40E-04 |
| | | 38,082,035 | | *ORMDL3* | -3.905 | 1.51E-04 | 6.40E-04 |
| rs903503 | g.37829571.T>G | 37,828,195 | | *PGAP3* | -4.898 | 2.83E-06 | |
| | | 37,828,197 | | *PGAP3* | -4.524 | 1.35E-05 | |
| | | 37,828,220 | | *PGAP3* | -4.809 | 4.12E-06 | |
| | | 37,828,273 | | *PGAP3* | -4.661 | 7.70E-06 | 0.010 |
| | | 37,828,283 | | *PGAP3* | -3.889 | 1.60E-04 | 0.010 |
| | | 37,828,296 | | *PGAP3* | -5.678 | 8.49E-08 | 0.002 |
| | | 37,828,338 | | *PGAP3* | -5.143 | 9.71E-07 | 0.002 |
| | | 37,828,377 | | *PGAP3* | -6.695 | 5.86E-10 | 0.002 |
| | | 37,828,384 | | *PGAP3* | -5.948 | 2.36E-08 | 0.002 |
| | | 37,828,560 | | *PGAP3* | -4.278 | 3.62E-05 | 0.002 |
| | | 37,843,658 | *ERBB2* | *PGAP3* | -4.339 | 2.85E-05 | 7.82E-05 |
| | | 37,844,480 | *PGAP3* | *ERBB2* | 4.207 | 4.80E-05 | 7.82E-05 |
| | | 38,072,514 | | *GSDMB* | -3.654 | 3.73E-04 | 0.046 |
| | | 38,082,022 | | *ORMDL3* | -3.993 | 1.09E-04 | 6.40E-04 |
| | | 38,082,035 | | *ORMDL3* | -3.778 | 2.40E-04 | 6.40E-04 |
| rs903502 | g.37829604.A>G | 37,828,195 | | *PGAP3* | -5.000 | 1.82E-06 | |
| | | 37,828,197 | | *PGAP3* | -4.400 | 2.23E-05 | |
| | | 37,828,220 | | *PGAP3* | -4.897 | 2.84E-06 | |
| | | 37,828,273 | | *PGAP3* | -4.687 | 6.89E-06 | 0.010 |
| | | 37,828,283 | | *PGAP3* | -4.018 | 9.88E-05 | 0.010 |
| | | 37,828,296 | | *PGAP3* | -5.643 | 1.00E-07 | 0.002 |
| | | 37,828,338 | | *PGAP3* | -5.249 | 6.07E-07 | 0.002 |
| | | 37,828,377 | | *PGAP3* | -6.797 | 3.48E-10 | 0.002 |
| | | 37,828,384 | | *PGAP3* | -6.071 | 1.30E-08 | 0.002 |
| | | 37,828,560 | | *PGAP3* | -4.337 | 2.87E-05 | 0.002 |
| | | 37,843,658 | *ERBB2* | *PGAP3* | -4.429 | 1.99E-05 | 7.82E-05 |
| | | 37,844,480 | *PGAP3* | *ERBB2* | 3.999 | 1.06E-04 | 7.82E-05 |
| | | 38,072,514 | | *GSDMB* | -3.537 | 5.61E-04 | 0.046 |
| | | 38,082,022 | | *ORMDL3* | -3.993 | 1.09E-04 | 6.40E-04 |
| | | 38,082,035 | | *ORMDL3* | -3.905 | 1.51E-04 | 6.40E-04 |
| rs2934956 | g.37830447.T>A | 37,828,195 | | *PGAP3* | -5.000 | 1.82E-06 | |
| | | 37,828,197 | | *PGAP3* | -4.400 | 2.23E-05 | |
| | | 37,828,220 | | *PGAP3* | -4.897 | 2.84E-06 | |

| | | | | | | | |
|---|---|---|---|---|---|---|---|
| | | 37,828,273 | | *PGAP3* | -4.687 | 6.89E-06 | 0.010 |
| | | 37,828,283 | | *PGAP3* | -4.018 | 9.88E-05 | 0.010 |
| | | 37,828,296 | | *PGAP3* | -5.643 | 1.00E-07 | 0.002 |
| | | 37,828,338 | | *PGAP3* | -5.249 | 6.07E-07 | 0.002 |
| | | 37,828,377 | | *PGAP3* | -6.797 | 3.48E-10 | 0.002 |
| | | 37,828,384 | | *PGAP3* | -6.071 | 1.30E-08 | 0.002 |
| | | 37,828,560 | | *PGAP3* | -4.337 | 2.87E-05 | 0.002 |
| | | 37,843,658 | *ERBB2* | *PGAP3* | -4.429 | 1.99E-05 | 7.82E-05 |
| | | 37,844,480 | *PGAP3* | *ERBB2* | 3.999 | 1.06E-04 | 7.82E-05 |
| | | 38,072,514 | | *GSDMB* | -3.537 | 5.61E-04 | 0.046 |
| | | 38,082,022 | | *ORMDL3* | -3.993 | 1.09E-04 | 6.40E-04 |
| | | 38,082,035 | | *ORMDL3* | -3.905 | 1.51E-04 | 6.40E-04 |
| rs2941504 | g.37830900.A>G (Val.155.Val) | 37,828,195 | | *PGAP3* | -5.016 | 1.70E-06 | |
| | | 37,828,197 | | *PGAP3* | -4.142 | 6.15E-05 | |
| | | 37,828,220 | | *PGAP3* | -4.815 | 4.03E-06 | |
| | | 37,828,273 | | *PGAP3* | -4.350 | 2.72E-05 | 0.010 |
| | | 37,828,283 | | *PGAP3* | -3.941 | 1.32E-04 | 0.010 |
| | | 37,828,296 | | *PGAP3* | -5.849 | 3.79E-08 | 0.002 |
| | | 37,828,338 | | *PGAP3* | -5.116 | 1.09E-06 | 0.002 |
| | | 37,828,377 | | *PGAP3* | -6.790 | 3.60E-10 | 0.002 |
| | | 37,828,384 | | *PGAP3* | -6.144 | 9.12E-09 | 0.002 |
| | | 37,828,560 | | *PGAP3* | -4.411 | 2.14E-05 | 0.002 |
| | | 37,843,658 | *ERBB2* | *PGAP3* | -3.662 | 3.62E-04 | 7.82E-05 |
| | | 38,072,514 | | *GSDMB* | -3.691 | 3.27E-04 | 0.046 |
| | | 38,077,039 | | | -3.551 | 5.36E-04 | 0.034 |
| | | 38,082,022 | | *ORMDL3* | -3.901 | 1.53E-04 | 6.40E-04 |
| rs1565922 | g.37831035.T>C | 37,828,195 | | *PGAP3* | -5.016 | 1.70E-06 | |
| | | 37,828,197 | | *PGAP3* | -4.142 | 6.15E-05 | |
| | | 37,828,220 | | *PGAP3* | -4.815 | 4.03E-06 | |
| | | 37,828,273 | | *PGAP3* | -4.350 | 2.72E-05 | 0.010 |
| | | 37,828,283 | | *PGAP3* | -3.941 | 1.32E-04 | 0.010 |
| | | 37,828,296 | | *PGAP3* | -5.849 | 3.79E-08 | 0.002 |
| | | 37,828,338 | | *PGAP3* | -5.116 | 1.09E-06 | 0.002 |
| | | 37,828,377 | | *PGAP3* | -6.790 | 3.60E-10 | 0.002 |
| | | 37,828,384 | | *PGAP3* | -6.144 | 9.12E-09 | 0.002 |
| | | 37,828,560 | | *PGAP3* | -4.411 | 2.14E-05 | 0.002 |
| | | 37,843,658 | *ERBB2* | *PGAP3* | -3.662 | 3.62E-04 | 7.82E-05 |
| | | 38,072,514 | | *GSDMB* | -3.691 | 3.27E-04 | 0.046 |
| | | 38,077,039 | | | -3.551 | 5.36E-04 | 0.034 |
| | | 38,082,022 | | *ORMDL3* | -3.901 | 1.53E-04 | 6.40E-04 |

| | | | | | | | |
|---|---|---|---|---|---|---|---|
| rs10558975 | g.37831304.G>C | 37,828,195 | | *PGAP3* | -4.987 | 1.92E-06 | |
| | | 37,828,197 | | *PGAP3* | -4.418 | 2.07E-05 | |
| | | 37,828,220 | | *PGAP3* | -4.900 | 2.81E-06 | |
| | | 37,828,273 | | *PGAP3* | -4.695 | 6.66E-06 | 0.010 |
| | | 37,828,283 | | *PGAP3* | -4.042 | 9.02E-05 | 0.010 |
| | | 37,828,296 | | *PGAP3* | -5.679 | 8.43E-08 | 0.002 |
| | | 37,828,338 | | *PGAP3* | -5.255 | 5.90E-07 | 0.002 |
| | | 37,828,377 | | *PGAP3* | -6.822 | 3.06E-10 | 0.002 |
| | | 37,828,384 | | *PGAP3* | -6.080 | 1.25E-08 | 0.002 |
| | | 37,828,560 | | *PGAP3* | -4.352 | 2.71E-05 | 0.002 |
| | | 37,843,658 | *ERBB2* | *PGAP3* | -4.440 | 1.90E-05 | 7.82E-05 |
| | | 37,844,480 | *PGAP3* | *ERBB2* | 3.976 | 1.16E-04 | 7.82E-05 |
| | | 38,072,514 | | *GSDMB* | -3.543 | 5.49E-04 | 0.046 |
| | | 38,082,022 | | *ORMDL3* | -4.005 | 1.04E-04 | 6.40E-04 |
| | | 38,082,035 | | *ORMDL3* | -3.902 | 1.52E-04 | 6.40E-04 |
| rs1565920 | g.37831613.T>C | 37,828,195 | | *PGAP3* | -5.000 | 1.82E-06 | |
| | | 37,828,197 | | *PGAP3* | -4.400 | 2.23E-05 | |
| | | 37,828,220 | | *PGAP3* | -4.897 | 2.84E-06 | |
| | | 37,828,273 | | *PGAP3* | -4.687 | 6.89E-06 | 0.010 |
| | | 37,828,283 | | *PGAP3* | -4.018 | 9.88E-05 | 0.010 |
| | | 37,828,296 | | *PGAP3* | -5.643 | 1.00E-07 | 0.002 |
| | | 37,828,338 | | *PGAP3* | -5.249 | 6.07E-07 | 0.002 |
| | | 37,828,377 | | *PGAP3* | -6.797 | 3.48E-10 | 0.002 |
| | | 37,828,384 | | *PGAP3* | -6.071 | 1.30E-08 | 0.002 |
| | | 37,828,560 | | *PGAP3* | -4.337 | 2.87E-05 | 0.002 |
| | | 37,843,658 | *ERBB2* | *PGAP3* | -4.429 | 1.99E-05 | 7.82E-05 |
| | | 37,844,480 | *PGAP3* | *ERBB2* | 3.999 | 1.06E-04 | 7.82E-05 |
| | | 38,072,514 | | *GSDMB* | -3.537 | 5.61E-04 | 0.046 |
| | | 38,082,022 | | *ORMDL3* | -3.993 | 1.09E-04 | 6.40E-04 |
| | | 38,082,035 | | *ORMDL3* | -3.905 | 1.51E-04 | 6.40E-04 |
| rs1495102 | g.37832093.A>G | 37,828,195 | | *PGAP3* | -5.016 | 1.70E-06 | |
| | | 37,828,197 | | *PGAP3* | -4.142 | 6.15E-05 | |
| | | 37,828,220 | | *PGAP3* | -4.815 | 4.03E-06 | |
| | | 37,828,273 | | *PGAP3* | -4.350 | 2.72E-05 | 0.010 |
| | | 37,828,283 | | *PGAP3* | -3.941 | 1.32E-04 | 0.010 |
| | | 37,828,296 | | *PGAP3* | -5.849 | 3.79E-08 | 0.002 |
| | | 37,828,338 | | *PGAP3* | -5.116 | 1.09E-06 | 0.002 |
| | | 37,828,377 | | *PGAP3* | -6.790 | 3.60E-10 | 0.002 |
| | | 37,828,384 | | *PGAP3* | -6.144 | 9.12E-09 | 0.002 |
| | | 37,828,560 | | *PGAP3* | -4.411 | 2.14E-05 | 0.002 |
| | | 37,843,658 | *ERBB2* | *PGAP3* | -3.662 | 3.62E-04 | 7.82E-05 |

| | | | | | | | |
|---|---|---|---|---|---|---|---|
| | | 38,072,514 | | *GSDMB* | -3.691 | 3.27E-04 | 0.046 |
| | | 38,077,039 | | | -3.551 | 5.36E-04 | 0.034 |
| | | 38,082,022 | | *ORMDL3* | -3.901 | 1.53E-04 | 6.40E-04 |
| rs1495101 | g.37832103.G>T | 37,828,195 | | *PGAP3* | -5.016 | 1.70E-06 | |
| | | 37,828,197 | | *PGAP3* | -4.142 | 6.15E-05 | |
| | | 37,828,220 | | *PGAP3* | -4.815 | 4.03E-06 | |
| | | 37,828,273 | | *PGAP3* | -4.350 | 2.72E-05 | 0.010 |
| | | 37,828,283 | | *PGAP3* | -3.941 | 1.32E-04 | 0.010 |
| | | 37,828,296 | | *PGAP3* | -5.849 | 3.79E-08 | 0.002 |
| | | 37,828,338 | | *PGAP3* | -5.116 | 1.09E-06 | 0.002 |
| | | 37,828,377 | | *PGAP3* | -6.790 | 3.60E-10 | 0.002 |
| | | 37,828,384 | | *PGAP3* | -6.144 | 9.12E-09 | 0.002 |
| | | 37,828,560 | | *PGAP3* | -4.411 | 2.14E-05 | 0.002 |
| | | 37,843,658 | *ERBB2* | *PGAP3* | -3.662 | 3.62E-04 | 7.82E-05 |
| | | 38,072,514 | | *GSDMB* | -3.691 | 3.27E-04 | 0.046 |
| | | 38,077,039 | | | -3.551 | 5.36E-04 | 0.034 |
| | | 38,082,022 | | *ORMDL3* | -3.901 | 1.53E-04 | 6.40E-04 |
| rs1495100 | g.37832279.A>G | 37,828,195 | | *PGAP3* | -5.000 | 1.82E-06 | |
| | | 37,828,197 | | *PGAP3* | -4.400 | 2.23E-05 | |
| | | 37,828,220 | | *PGAP3* | -4.897 | 2.84E-06 | |
| | | 37,828,273 | | *PGAP3* | -4.687 | 6.89E-06 | 0.010 |
| | | 37,828,283 | | *PGAP3* | -4.018 | 9.88E-05 | 0.010 |
| | | 37,828,296 | | *PGAP3* | -5.643 | 1.00E-07 | 0.002 |
| | | 37,828,338 | | *PGAP3* | -5.249 | 6.07E-07 | 0.002 |
| | | 37,828,377 | | *PGAP3* | -6.797 | 3.48E-10 | 0.002 |
| | | 37,828,384 | | *PGAP3* | -6.071 | 1.30E-08 | 0.002 |
| | | 37,828,560 | | *PGAP3* | -4.337 | 2.87E-05 | 0.002 |
| | | 37,843,658 | *ERBB2* | *PGAP3* | -4.429 | 1.99E-05 | 7.82E-05 |
| | | 37,844,480 | *PGAP3* | *ERBB2* | 3.999 | 1.06E-04 | 7.82E-05 |
| | | 38,072,514 | | *GSDMB* | -3.537 | 5.61E-04 | 0.046 |
| | | 38,082,022 | | *ORMDL3* | -3.993 | 1.09E-04 | 6.40E-04 |
| | | 38,082,035 | | *ORMDL3* | -3.905 | 1.51E-04 | 6.40E-04 |
| rs2934953 | g.37832315.A>T | 37,828,195 | | *PGAP3* | -5.016 | 1.70E-06 | |
| | | 37,828,197 | | *PGAP3* | -4.142 | 6.15E-05 | |
| | | 37,828,220 | | *PGAP3* | -4.815 | 4.03E-06 | |
| | | 37,828,273 | | *PGAP3* | -4.350 | 2.72E-05 | 0.010 |
| | | 37,828,283 | | *PGAP3* | -3.941 | 1.32E-04 | 0.010 |
| | | 37,828,296 | | *PGAP3* | -5.849 | 3.79E-08 | 0.002 |
| | | 37,828,338 | | *PGAP3* | -5.116 | 1.09E-06 | 0.002 |
| | | 37,828,377 | | *PGAP3* | -6.790 | 3.60E-10 | 0.002 |
| | | 37,828,384 | | *PGAP3* | -6.144 | 9.12E-09 | 0.002 |

| | | | | | | | |
|---|---|---|---|---|---|---|---|
| | | 37,828,560 | | *PGAP3* | -4.411 | 2.14E-05 | 0.002 |
| | | 37,843,658 | *ERBB2* | *PGAP3* | -3.662 | 3.62E-04 | 7.82E-05 |
| | | 38,072,514 | | *GSDMB* | -3.691 | 3.27E-04 | 0.046 |
| | | 38,077,039 | | | -3.551 | 5.36E-04 | 0.034 |
| | | 38,082,022 | | *ORMDL3* | -3.901 | 1.53E-04 | 6.40E-04 |
| rs2934952 | g.37832366.C>T | 37,828,195 | | *PGAP3* | -5.016 | 1.70E-06 | |
| | | 37,828,197 | | *PGAP3* | -4.142 | 6.15E-05 | |
| | | 37,828,220 | | *PGAP3* | -4.815 | 4.03E-06 | |
| | | 37,828,273 | | *PGAP3* | -4.350 | 2.72E-05 | 0.010 |
| | | 37,828,283 | | *PGAP3* | -3.941 | 1.32E-04 | 0.010 |
| | | 37,828,296 | | *PGAP3* | -5.849 | 3.79E-08 | 0.002 |
| | | 37,828,338 | | *PGAP3* | -5.116 | 1.09E-06 | 0.002 |
| | | 37,828,377 | | *PGAP3* | -6.790 | 3.60E-10 | 0.002 |
| | | 37,828,384 | | *PGAP3* | -6.144 | 9.12E-09 | 0.002 |
| | | 37,828,560 | | *PGAP3* | -4.411 | 2.14E-05 | 0.002 |
| | | 37,843,658 | *ERBB2* | *PGAP3* | -3.662 | 3.62E-04 | 7.82E-05 |
| | | 38,072,514 | | *GSDMB* | -3.691 | 3.27E-04 | 0.046 |
| | | 38,077,039 | | | -3.551 | 5.36E-04 | 0.034 |
| | | 38,082,022 | | *ORMDL3* | -3.901 | 1.53E-04 | 6.40E-04 |
| rs2941505 | g.37832704.A>G | 37,828,195 | | *PGAP3* | -5.000 | 1.82E-06 | |
| | | 37,828,197 | | *PGAP3* | -4.400 | 2.23E-05 | |
| | | 37,828,220 | | *PGAP3* | -4.897 | 2.84E-06 | |
| | | 37,828,273 | | *PGAP3* | -4.687 | 6.89E-06 | 0.010 |
| | | 37,828,283 | | *PGAP3* | -4.018 | 9.88E-05 | 0.010 |
| | | 37,828,296 | | *PGAP3* | -5.643 | 1.00E-07 | 0.002 |
| | | 37,828,338 | | *PGAP3* | -5.249 | 6.07E-07 | 0.002 |
| | | 37,828,377 | | *PGAP3* | -6.797 | 3.48E-10 | 0.002 |
| | | 37,828,384 | | *PGAP3* | -6.071 | 1.30E-08 | 0.002 |
| | | 37,828,560 | | *PGAP3* | -4.337 | 2.87E-05 | 0.002 |
| | | 37,843,658 | *ERBB2* | *PGAP3* | -4.429 | 1.99E-05 | 7.82E-05 |
| | | 37,844,480 | *PGAP3* | *ERBB2* | 3.999 | 1.06E-04 | 7.82E-05 |
| | | 38,072,514 | | *GSDMB* | -3.537 | 5.61E-04 | 0.046 |
| | | 38,082,022 | | *ORMDL3* | -3.993 | 1.09E-04 | 6.40E-04 |
| | | 38,082,035 | | *ORMDL3* | -3.905 | 1.51E-04 | 6.40E-04 |
| rs2952152 | g.37832735.T>C | 37,828,195 | | *PGAP3* | -5.000 | 1.82E-06 | |
| | | 37,828,197 | | *PGAP3* | -4.400 | 2.23E-05 | |
| | | 37,828,220 | | *PGAP3* | -4.897 | 2.84E-06 | |
| | | 37,828,273 | | *PGAP3* | -4.687 | 6.89E-06 | 0.010 |
| | | 37,828,283 | | *PGAP3* | -4.018 | 9.88E-05 | 0.010 |
| | | 37,828,296 | | *PGAP3* | -5.643 | 1.00E-07 | 0.002 |
| | | 37,828,338 | | *PGAP3* | -5.249 | 6.07E-07 | 0.002 |

| | | | | | | | |
|---|---|---|---|---|---|---|---|
| | | 37,828,377 | | *PGAP3* | -6.797 | 3.48E-10 | 0.002 |
| | | 37,828,384 | | *PGAP3* | -6.071 | 1.30E-08 | 0.002 |
| | | 37,828,560 | | *PGAP3* | -4.337 | 2.87E-05 | 0.002 |
| | | 37,843,658 | *ERBB2* | *PGAP3* | -4.429 | 1.99E-05 | 7.82E-05 |
| | | 37,844,480 | *PGAP3* | *ERBB2* | 3.999 | 1.06E-04 | 7.82E-05 |
| | | 38,072,514 | | *GSDMB* | -3.537 | 5.61E-04 | 0.046 |
| | | 38,082,022 | | *ORMDL3* | -3.993 | 1.09E-04 | 6.40E-04 |
| | | 38,082,035 | | *ORMDL3* | -3.905 | 1.51E-04 | 6.40E-04 |
| rs2941506 | g.37833035.A>G | 37,828,195 | | *PGAP3* | -5.000 | 1.82E-06 | |
| | | 37,828,197 | | *PGAP3* | -4.400 | 2.23E-05 | |
| | | 37,828,220 | | *PGAP3* | -4.897 | 2.84E-06 | |
| | | 37,828,273 | | *PGAP3* | -4.687 | 6.89E-06 | 0.010 |
| | | 37,828,283 | | *PGAP3* | -4.018 | 9.88E-05 | 0.010 |
| | | 37,828,296 | | *PGAP3* | -5.643 | 1.00E-07 | 0.002 |
| | | 37,828,338 | | *PGAP3* | -5.249 | 6.07E-07 | 0.002 |
| | | 37,828,377 | | *PGAP3* | -6.797 | 3.48E-10 | 0.002 |
| | | 37,828,384 | | *PGAP3* | -6.071 | 1.30E-08 | 0.002 |
| | | 37,828,560 | | *PGAP3* | -4.337 | 2.87E-05 | 0.002 |
| | | 37,843,658 | *ERBB2* | *PGAP3* | -4.429 | 1.99E-05 | 7.82E-05 |
| | | 37,844,480 | *PGAP3* | *ERBB2* | 3.999 | 1.06E-04 | 7.82E-05 |
| | | 38,072,514 | | *GSDMB* | -3.537 | 5.61E-04 | 0.046 |
| | | 38,082,022 | | *ORMDL3* | -3.993 | 1.09E-04 | 6.40E-04 |
| | | 38,082,035 | | *ORMDL3* | -3.905 | 1.51E-04 | 6.40E-04 |
| rs2934951 | g.37833328.T>C | 37,828,195 | | *PGAP3* | -5.000 | 1.82E-06 | |
| | | 37,828,197 | | *PGAP3* | -4.400 | 2.23E-05 | |
| | | 37,828,220 | | *PGAP3* | -4.897 | 2.84E-06 | |
| | | 37,828,273 | | *PGAP3* | -4.687 | 6.89E-06 | 0.010 |
| | | 37,828,283 | | *PGAP3* | -4.018 | 9.88E-05 | 0.010 |
| | | 37,828,296 | | *PGAP3* | -5.643 | 1.00E-07 | 0.002 |
| | | 37,828,338 | | *PGAP3* | -5.249 | 6.07E-07 | 0.002 |
| | | 37,828,377 | | *PGAP3* | -6.797 | 3.48E-10 | 0.002 |
| | | 37,828,384 | | *PGAP3* | -6.071 | 1.30E-08 | 0.002 |
| | | 37,828,560 | | *PGAP3* | -4.337 | 2.87E-05 | 0.002 |
| | | 37,843,658 | *ERBB2* | *PGAP3* | -4.429 | 1.99E-05 | 7.82E-05 |
| | | 37,844,480 | *PGAP3* | *ERBB2* | 3.999 | 1.06E-04 | 7.82E-05 |
| | | 38,072,514 | | *GSDMB* | -3.537 | 5.61E-04 | 0.046 |
| | | 38,082,022 | | *ORMDL3* | -3.993 | 1.09E-04 | 6.40E-04 |
| | | 38,082,035 | | *ORMDL3* | -3.905 | 1.51E-04 | 6.40E-04 |
| rs907088 | g.37833567.G>C | 37,828,195 | | *PGAP3* | -5.004 | 1.79E-06 | |
| | | 37,828,197 | | *PGAP3* | -4.400 | 2.23E-05 | |
| | | 37,828,220 | | *PGAP3* | -4.879 | 3.06E-06 | |

| | | | | | | | |
|---|---|---|---|---|---|---|---|
| | | 37,828,273 | | *PGAP3* | -4.644 | 8.25E-06 | 0.010 |
| | | 37,828,283 | | *PGAP3* | -3.997 | 1.07E-04 | 0.010 |
| | | 37,828,296 | | *PGAP3* | -5.610 | 1.16E-07 | 0.002 |
| | | 37,828,338 | | *PGAP3* | -5.217 | 6.98E-07 | 0.002 |
| | | 37,828,377 | | *PGAP3* | -6.760 | 4.21E-10 | 0.002 |
| | | 37,828,384 | | *PGAP3* | -6.036 | 1.55E-08 | 0.002 |
| | | 37,828,560 | | *PGAP3* | -4.336 | 2.88E-05 | 0.002 |
| | | 37,843,658 | *ERBB2* | *PGAP3* | -4.414 | 2.11E-05 | 7.82E-05 |
| | | 37,844,480 | *PGAP3* | *ERBB2* | 4.015 | 9.98E-05 | 7.82E-05 |
| | | 38,072,514 | | *GSDMB* | -3.544 | 5.48E-04 | 0.046 |
| | | 38,077,039 | | | -3.374 | 9.77E-04 | 0.034 |
| | | 38,082,022 | | *ORMDL3* | -4.007 | 1.03E-04 | 6.40E-04 |
| | | 38,082,035 | | *ORMDL3* | -3.929 | 1.38E-04 | 6.40E-04 |
| rs907089 | g.37833600.G>A | 37,828,195 | | *PGAP3* | -5.004 | 1.79E-06 | |
| | | 37,828,197 | | *PGAP3* | -4.400 | 2.23E-05 | |
| | | 37,828,220 | | *PGAP3* | -4.879 | 3.06E-06 | |
| | | 37,828,273 | | *PGAP3* | -4.644 | 8.25E-06 | 0.010 |
| | | 37,828,283 | | *PGAP3* | -3.997 | 1.07E-04 | 0.010 |
| | | 37,828,296 | | *PGAP3* | -5.610 | 1.16E-07 | 0.002 |
| | | 37,828,338 | | *PGAP3* | -5.217 | 6.98E-07 | 0.002 |
| | | 37,828,377 | | *PGAP3* | -6.760 | 4.21E-10 | 0.002 |
| | | 37,828,384 | | *PGAP3* | -6.036 | 1.55E-08 | 0.002 |
| | | 37,828,560 | | *PGAP3* | -4.336 | 2.88E-05 | 0.002 |
| | | 37,843,658 | *ERBB2* | *PGAP3* | -4.414 | 2.11E-05 | 7.82E-05 |
| | | 37,844,480 | *PGAP3* | *ERBB2* | 4.015 | 9.98E-05 | 7.82E-05 |
| | | 38,072,514 | | *GSDMB* | -3.544 | 5.48E-04 | 0.046 |
| | | 38,077,039 | | | -3.374 | 9.77E-04 | 0.034 |
| | | 38,082,022 | | *ORMDL3* | -4.007 | 1.03E-04 | 6.40E-04 |
| | | 38,082,035 | | *ORMDL3* | -3.929 | 1.38E-04 | 6.40E-04 |
| rs907090 | g.37833632.T>C | 37,828,195 | | *PGAP3* | -5.004 | 1.79E-06 | |
| | | 37,828,197 | | *PGAP3* | -4.400 | 2.23E-05 | |
| | | 37,828,220 | | *PGAP3* | -4.879 | 3.06E-06 | |
| | | 37,828,273 | | *PGAP3* | -4.644 | 8.25E-06 | 0.010 |
| | | 37,828,283 | | *PGAP3* | -3.997 | 1.07E-04 | 0.010 |
| | | 37,828,296 | | *PGAP3* | -5.610 | 1.16E-07 | 0.002 |
| | | 37,828,338 | | *PGAP3* | -5.217 | 6.98E-07 | 0.002 |
| | | 37,828,377 | | *PGAP3* | -6.760 | 4.21E-10 | 0.002 |
| | | 37,828,384 | | *PGAP3* | -6.036 | 1.55E-08 | 0.002 |
| | | 37,828,560 | | *PGAP3* | -4.336 | 2.88E-05 | 0.002 |
| | | 37,843,658 | *ERBB2* | *PGAP3* | -4.414 | 2.11E-05 | 7.82E-05 |
| | | 37,844,480 | *PGAP3* | *ERBB2* | 4.015 | 9.98E-05 | 7.82E-05 |

| | | | | | | | |
|---|---|---|---|---|---|---|---|
| | | 38,072,514 | | *GSDMB* | -3.544 | 5.48E-04 | 0.046 |
| | | 38,077,039 | | | -3.374 | 9.77E-04 | 0.034 |
| | | 38,082,022 | | *ORMDL3* | -4.007 | 1.03E-04 | 6.40E-04 |
| | | 38,082,035 | | *ORMDL3* | -3.929 | 1.38E-04 | 6.40E-04 |
| rs9675194 | g.37833805.T>C | 37,828,195 | | *PGAP3* | -4.992 | 1.88E-06 | |
| | | 37,828,197 | | *PGAP3* | -4.382 | 2.40E-05 | |
| | | 37,828,220 | | *PGAP3* | -4.835 | 3.70E-06 | |
| | | 37,828,273 | | *PGAP3* | -4.648 | 8.13E-06 | 0.010 |
| | | 37,828,283 | | *PGAP3* | -4.007 | 1.03E-04 | 0.010 |
| | | 37,828,296 | | *PGAP3* | -5.608 | 1.18E-07 | 0.002 |
| | | 37,828,338 | | *PGAP3* | -5.246 | 6.13E-07 | 0.002 |
| | | 37,828,377 | | *PGAP3* | -6.773 | 3.94E-10 | 0.002 |
| | | 37,828,384 | | *PGAP3* | -6.060 | 1.38E-08 | 0.002 |
| | | 37,828,560 | | *PGAP3* | -4.385 | 2.37E-05 | 0.002 |
| | | 37,843,658 | *ERBB2* | *PGAP3* | -4.373 | 2.49E-05 | 7.82E-05 |
| | | 37,844,480 | *PGAP3* | *ERBB2* | 3.915 | 1.45E-04 | 7.82E-05 |
| | | 38,072,514 | | *GSDMB* | -3.484 | 6.73E-04 | 0.046 |
| | | 38,077,039 | | | -3.370 | 9.90E-04 | 0.034 |
| | | 38,082,022 | | *ORMDL3* | -4.004 | 1.04E-04 | 6.40E-04 |
| | | 38,082,035 | | *ORMDL3* | -3.919 | 1.43E-04 | 6.40E-04 |
| rs2313171 | g.37833842.T>C | 37,828,195 | | *PGAP3* | -4.992 | 1.88E-06 | |
| | | 37,828,197 | | *PGAP3* | -4.382 | 2.40E-05 | |
| | | 37,828,220 | | *PGAP3* | -4.835 | 3.70E-06 | |
| | | 37,828,273 | | *PGAP3* | -4.648 | 8.13E-06 | 0.010 |
| | | 37,828,283 | | *PGAP3* | -4.007 | 1.03E-04 | 0.010 |
| | | 37,828,296 | | *PGAP3* | -5.608 | 1.18E-07 | 0.002 |
| | | 37,828,338 | | *PGAP3* | -5.246 | 6.13E-07 | 0.002 |
| | | 37,828,377 | | *PGAP3* | -6.773 | 3.94E-10 | 0.002 |
| | | 37,828,384 | | *PGAP3* | -6.060 | 1.38E-08 | 0.002 |
| | | 37,828,560 | | *PGAP3* | -4.385 | 2.37E-05 | 0.002 |
| | | 37,843,658 | *ERBB2* | *PGAP3* | -4.373 | 2.49E-05 | 7.82E-05 |
| | | 37,844,480 | *PGAP3* | *ERBB2* | 3.915 | 1.45E-04 | 7.82E-05 |
| | | 38,072,514 | | *GSDMB* | -3.484 | 6.73E-04 | 0.046 |
| | | 38,077,039 | | | -3.370 | 9.90E-04 | 0.034 |
| | | 38,082,022 | | *ORMDL3* | -4.004 | 1.04E-04 | 6.40E-04 |
| | | 38,082,035 | | *ORMDL3* | -3.919 | 1.43E-04 | 6.40E-04 |
| rs732084 | g.37834357.T>G | 37,828,195 | | *PGAP3* | -5.004 | 1.79E-06 | |
| | | 37,828,197 | | *PGAP3* | -4.400 | 2.23E-05 | |
| | | 37,828,220 | | *PGAP3* | -4.879 | 3.06E-06 | |
| | | 37,828,273 | | *PGAP3* | -4.644 | 8.25E-06 | 0.010 |
| | | 37,828,283 | | *PGAP3* | -3.997 | 1.07E-04 | 0.010 |

| | | | | | | | |
|---|---|---|---|---|---|---|---|
| | | 37,828,296 | | *PGAP3* | -5.610 | 1.16E-07 | 0.002 |
| | | 37,828,338 | | *PGAP3* | -5.217 | 6.98E-07 | 0.002 |
| | | 37,828,377 | | *PGAP3* | -6.760 | 4.21E-10 | 0.002 |
| | | 37,828,384 | | *PGAP3* | -6.036 | 1.55E-08 | 0.002 |
| | | 37,828,560 | | *PGAP3* | -4.336 | 2.88E-05 | 0.002 |
| | | 37,843,658 | *ERBB2* | *PGAP3* | -4.414 | 2.11E-05 | 7.82E-05 |
| | | 37,844,480 | *PGAP3* | *ERBB2* | 4.015 | 9.98E-05 | 7.82E-05 |
| | | 38,072,514 | | *GSDMB* | -3.544 | 5.48E-04 | 0.046 |
| | | 38,077,039 | | | -3.374 | 9.77E-04 | 0.034 |
| | | 38,082,022 | | *ORMDL3* | -4.007 | 1.03E-04 | 6.40E-04 |
| | | 38,082,035 | | *ORMDL3* | -3.929 | 1.38E-04 | 6.40E-04 |
| rs732083 | g.37834367.T>C | 37,828,195 | | *PGAP3* | -5.004 | 1.79E-06 | |
| | | 37,828,197 | | *PGAP3* | -4.400 | 2.23E-05 | |
| | | 37,828,220 | | *PGAP3* | -4.879 | 3.06E-06 | |
| | | 37,828,273 | | *PGAP3* | -4.644 | 8.25E-06 | 0.010 |
| | | 37,828,283 | | *PGAP3* | -3.997 | 1.07E-04 | 0.010 |
| | | 37,828,296 | | *PGAP3* | -5.610 | 1.16E-07 | 0.002 |
| | | 37,828,338 | | *PGAP3* | -5.217 | 6.98E-07 | 0.002 |
| | | 37,828,377 | | *PGAP3* | -6.760 | 4.21E-10 | 0.002 |
| | | 37,828,384 | | *PGAP3* | -6.036 | 1.55E-08 | 0.002 |
| | | 37,828,560 | | *PGAP3* | -4.336 | 2.88E-05 | 0.002 |
| | | 37,843,658 | *ERBB2* | *PGAP3* | -4.414 | 2.11E-05 | 7.82E-05 |
| | | 37,844,480 | *PGAP3* | *ERBB2* | 4.015 | 9.98E-05 | 7.82E-05 |
| | | 38,072,514 | | *GSDMB* | -3.544 | 5.48E-04 | 0.046 |
| | | 38,077,039 | | | -3.374 | 9.77E-04 | 0.034 |
| | | 38,082,022 | | *ORMDL3* | -4.007 | 1.03E-04 | 6.40E-04 |
| | | 38,082,035 | | *ORMDL3* | -3.929 | 1.38E-04 | 6.40E-04 |
| rs12150298 | g.37834541.T>C | 37,828,195 | | *PGAP3* | -5.004 | 1.79E-06 | |
| | | 37,828,197 | | *PGAP3* | -4.400 | 2.23E-05 | |
| | | 37,828,220 | | *PGAP3* | -4.879 | 3.06E-06 | |
| | | 37,828,273 | | *PGAP3* | -4.644 | 8.25E-06 | 0.010 |
| | | 37,828,283 | | *PGAP3* | -3.997 | 1.07E-04 | 0.010 |
| | | 37,828,296 | | *PGAP3* | -5.610 | 1.16E-07 | 0.002 |
| | | 37,828,338 | | *PGAP3* | -5.217 | 6.98E-07 | 0.002 |
| | | 37,828,377 | | *PGAP3* | -6.760 | 4.21E-10 | 0.002 |
| | | 37,828,384 | | *PGAP3* | -6.036 | 1.55E-08 | 0.002 |
| | | 37,828,560 | | *PGAP3* | -4.336 | 2.88E-05 | 0.002 |
| | | 37,843,658 | *ERBB2* | *PGAP3* | -4.414 | 2.11E-05 | 7.82E-05 |
| | | 37,844,480 | *PGAP3* | *ERBB2* | 4.015 | 9.98E-05 | 7.82E-05 |
| | | 38,072,514 | | *GSDMB* | -3.544 | 5.48E-04 | 0.046 |
| | | 38,077,039 | | | -3.374 | 9.77E-04 | 0.034 |

| | | | | | | | |
|---|---|---|---|---|---|---|---|
| | | 38,082,022 | | *ORMDL3* | -4.007 | 1.03E-04 | 6.40E-04 |
| | | 38,082,035 | | *ORMDL3* | -3.929 | 1.38E-04 | 6.40E-04 |
| rs12150603 | g.37834715.A>G | 37,828,195 | | *PGAP3* | -5.004 | 1.79E-06 | |
| | | 37,828,197 | | *PGAP3* | -4.400 | 2.23E-05 | |
| | | 37,828,220 | | *PGAP3* | -4.879 | 3.06E-06 | |
| | | 37,828,273 | | *PGAP3* | -4.644 | 8.25E-06 | 0.010 |
| | | 37,828,283 | | *PGAP3* | -3.997 | 1.07E-04 | 0.010 |
| | | 37,828,296 | | *PGAP3* | -5.610 | 1.16E-07 | 0.002 |
| | | 37,828,338 | | *PGAP3* | -5.217 | 6.98E-07 | 0.002 |
| | | 37,828,377 | | *PGAP3* | -6.760 | 4.21E-10 | 0.002 |
| | | 37,828,384 | | *PGAP3* | -6.036 | 1.55E-08 | 0.002 |
| | | 37,828,560 | | *PGAP3* | -4.336 | 2.88E-05 | 0.002 |
| | | 37,843,658 | *ERBB2* | *PGAP3* | -4.414 | 2.11E-05 | 7.82E-05 |
| | | 37,844,480 | *PGAP3* | *ERBB2* | 4.015 | 9.98E-05 | 7.82E-05 |
| | | 38,072,514 | | *GSDMB* | -3.544 | 5.48E-04 | 0.046 |
| | | 38,077,039 | | | -3.374 | 9.77E-04 | 0.034 |
| | | 38,082,022 | | *ORMDL3* | -4.007 | 1.03E-04 | 6.40E-04 |
| | | 38,082,035 | | *ORMDL3* | -3.929 | 1.38E-04 | 6.40E-04 |
| rs8077172 | g.37834977.G>T | 37,828,195 | | *PGAP3* | -5.004 | 1.79E-06 | |
| | | 37,828,197 | | *PGAP3* | -4.400 | 2.23E-05 | |
| | | 37,828,220 | | *PGAP3* | -4.879 | 3.06E-06 | |
| | | 37,828,273 | | *PGAP3* | -4.644 | 8.25E-06 | 0.010 |
| | | 37,828,283 | | *PGAP3* | -3.997 | 1.07E-04 | 0.010 |
| | | 37,828,296 | | *PGAP3* | -5.610 | 1.16E-07 | 0.002 |
| | | 37,828,338 | | *PGAP3* | -5.217 | 6.98E-07 | 0.002 |
| | | 37,828,377 | | *PGAP3* | -6.760 | 4.21E-10 | 0.002 |
| | | 37,828,384 | | *PGAP3* | -6.036 | 1.55E-08 | 0.002 |
| | | 37,828,560 | | *PGAP3* | -4.336 | 2.88E-05 | 0.002 |
| | | 37,843,658 | *ERBB2* | *PGAP3* | -4.414 | 2.11E-05 | 7.82E-05 |
| | | 37,844,480 | *PGAP3* | *ERBB2* | 4.015 | 9.98E-05 | 7.82E-05 |
| | | 38,072,514 | | *GSDMB* | -3.544 | 5.48E-04 | 0.046 |
| | | 38,077,039 | | | -3.374 | 9.77E-04 | 0.034 |
| | | 38,082,022 | | *ORMDL3* | -4.007 | 1.03E-04 | 6.40E-04 |
| | | 38,082,035 | | *ORMDL3* | -3.929 | 1.38E-04 | 6.40E-04 |
| rs8078228 | g.37834998.C>T | 37,828,195 | | *PGAP3* | -5.004 | 1.79E-06 | |
| | | 37,828,197 | | *PGAP3* | -4.400 | 2.23E-05 | |
| | | 37,828,220 | | *PGAP3* | -4.879 | 3.06E-06 | |
| | | 37,828,273 | | *PGAP3* | -4.644 | 8.25E-06 | 0.010 |
| | | 37,828,283 | | *PGAP3* | -3.997 | 1.07E-04 | 0.010 |
| | | 37,828,296 | | *PGAP3* | -5.610 | 1.16E-07 | 0.002 |
| | | 37,828,338 | | *PGAP3* | -5.217 | 6.98E-07 | 0.002 |

| | | | | | | | |
|---|---|---|---|---|---|---|---|
| | | 37,828,377 | | *PGAP3* | -6.760 | 4.21E-10 | 0.002 |
| | | 37,828,384 | | *PGAP3* | -6.036 | 1.55E-08 | 0.002 |
| | | 37,828,560 | | *PGAP3* | -4.336 | 2.88E-05 | 0.002 |
| | | 37,843,658 | *ERBB2* | *PGAP3* | -4.414 | 2.11E-05 | 7.82E-05 |
| | | 37,844,480 | *PGAP3* | *ERBB2* | 4.015 | 9.98E-05 | 7.82E-05 |
| | | 38,072,514 | | *GSDMB* | -3.544 | 5.48E-04 | 0.046 |
| | | 38,077,039 | | | -3.374 | 9.77E-04 | 0.034 |
| | | 38,082,022 | | *ORMDL3* | -4.007 | 1.03E-04 | 6.40E-04 |
| | | 38,082,035 | | *ORMDL3* | -3.929 | 1.38E-04 | 6.40E-04 |
| rs1018246 | g.37835240.C>T | 37,828,195 | | *PGAP3* | -5.004 | 1.79E-06 | |
| | | 37,828,197 | | *PGAP3* | -4.400 | 2.23E-05 | |
| | | 37,828,220 | | *PGAP3* | -4.879 | 3.06E-06 | |
| | | 37,828,273 | | *PGAP3* | -4.644 | 8.25E-06 | 0.010 |
| | | 37,828,283 | | *PGAP3* | -3.997 | 1.07E-04 | 0.010 |
| | | 37,828,296 | | *PGAP3* | -5.610 | 1.16E-07 | 0.002 |
| | | 37,828,338 | | *PGAP3* | -5.217 | 6.98E-07 | 0.002 |
| | | 37,828,377 | | *PGAP3* | -6.760 | 4.21E-10 | 0.002 |
| | | 37,828,384 | | *PGAP3* | -6.036 | 1.55E-08 | 0.002 |
| | | 37,828,560 | | *PGAP3* | -4.336 | 2.88E-05 | 0.002 |
| | | 37,843,658 | *ERBB2* | *PGAP3* | -4.414 | 2.11E-05 | 7.82E-05 |
| | | 37,844,480 | *PGAP3* | *ERBB2* | 4.015 | 9.98E-05 | 7.82E-05 |
| | | 38,072,514 | | *GSDMB* | -3.544 | 5.48E-04 | 0.046 |
| | | 38,077,039 | | | -3.374 | 9.77E-04 | 0.034 |
| | | 38,082,022 | | *ORMDL3* | -4.007 | 1.03E-04 | 6.40E-04 |
| | | 38,082,035 | | *ORMDL3* | -3.929 | 1.38E-04 | 6.40E-04 |
| rs11078919 | g.37835755.T>C | 37,828,195 | | *PGAP3* | -5.004 | 1.79E-06 | |
| | | 37,828,197 | | *PGAP3* | -4.400 | 2.23E-05 | |
| | | 37,828,220 | | *PGAP3* | -4.879 | 3.06E-06 | |
| | | 37,828,273 | | *PGAP3* | -4.644 | 8.25E-06 | 0.010 |
| | | 37,828,283 | | *PGAP3* | -3.997 | 1.07E-04 | 0.010 |
| | | 37,828,296 | | *PGAP3* | -5.610 | 1.16E-07 | 0.002 |
| | | 37,828,338 | | *PGAP3* | -5.217 | 6.98E-07 | 0.002 |
| | | 37,828,377 | | *PGAP3* | -6.760 | 4.21E-10 | 0.002 |
| | | 37,828,384 | | *PGAP3* | -6.036 | 1.55E-08 | 0.002 |
| | | 37,828,560 | | *PGAP3* | -4.336 | 2.88E-05 | 0.002 |
| | | 37,843,658 | *ERBB2* | *PGAP3* | -4.414 | 2.11E-05 | 7.82E-05 |
| | | 37,844,480 | *PGAP3* | *ERBB2* | 4.015 | 9.98E-05 | 7.82E-05 |
| | | 38,072,514 | | *GSDMB* | -3.544 | 5.48E-04 | 0.046 |
| | | 38,077,039 | | | -3.374 | 9.77E-04 | 0.034 |
| | | 38,082,022 | | *ORMDL3* | -4.007 | 1.03E-04 | 6.40E-04 |
| | | 38,082,035 | | *ORMDL3* | -3.929 | 1.38E-04 | 6.40E-04 |

| | | | | | | | |
|---|---|---|---|---|---|---|---|
| rs1476278 | g.37836243.A>G | 37,828,195 | | *PGAP3* | -5.004 | 1.79E-06 | |
| | | 37,828,197 | | *PGAP3* | -4.400 | 2.23E-05 | |
| | | 37,828,220 | | *PGAP3* | -4.879 | 3.06E-06 | |
| | | 37,828,273 | | *PGAP3* | -4.644 | 8.25E-06 | 0.010 |
| | | 37,828,283 | | *PGAP3* | -3.997 | 1.07E-04 | 0.010 |
| | | 37,828,296 | | *PGAP3* | -5.610 | 1.16E-07 | 0.002 |
| | | 37,828,338 | | *PGAP3* | -5.217 | 6.98E-07 | 0.002 |
| | | 37,828,377 | | *PGAP3* | -6.760 | 4.21E-10 | 0.002 |
| | | 37,828,384 | | *PGAP3* | -6.036 | 1.55E-08 | 0.002 |
| | | 37,828,560 | | *PGAP3* | -4.336 | 2.88E-05 | 0.002 |
| | | 37,843,658 | *ERBB2* | *PGAP3* | -4.414 | 2.11E-05 | 7.82E-05 |
| | | 37,844,480 | *PGAP3* | *ERBB2* | 4.015 | 9.98E-05 | 7.82E-05 |
| | | 38,072,514 | | *GSDMB* | -3.544 | 5.48E-04 | 0.046 |
| | | 38,077,039 | | | -3.374 | 9.77E-04 | 0.034 |
| | | 38,082,022 | | *ORMDL3* | -4.007 | 1.03E-04 | 6.40E-04 |
| | | 38,082,035 | | *ORMDL3* | -3.929 | 1.38E-04 | 6.40E-04 |
| rs9303274 | g.37836353.C>T | 37,828,195 | | *PGAP3* | -5.004 | 1.79E-06 | |
| | | 37,828,197 | | *PGAP3* | -4.400 | 2.23E-05 | |
| | | 37,828,220 | | *PGAP3* | -4.879 | 3.06E-06 | |
| | | 37,828,273 | | *PGAP3* | -4.644 | 8.25E-06 | 0.010 |
| | | 37,828,283 | | *PGAP3* | -3.997 | 1.07E-04 | 0.010 |
| | | 37,828,296 | | *PGAP3* | -5.610 | 1.16E-07 | 0.002 |
| | | 37,828,338 | | *PGAP3* | -5.217 | 6.98E-07 | 0.002 |
| | | 37,828,377 | | *PGAP3* | -6.760 | 4.21E-10 | 0.002 |
| | | 37,828,384 | | *PGAP3* | -6.036 | 1.55E-08 | 0.002 |
| | | 37,828,560 | | *PGAP3* | -4.336 | 2.88E-05 | 0.002 |
| | | 37,843,658 | *ERBB2* | *PGAP3* | -4.414 | 2.11E-05 | 7.82E-05 |
| | | 37,844,480 | *PGAP3* | *ERBB2* | 4.015 | 9.98E-05 | 7.82E-05 |
| | | 38,072,514 | | *GSDMB* | -3.544 | 5.48E-04 | 0.046 |
| | | 38,077,039 | | | -3.374 | 9.77E-04 | 0.034 |
| | | 38,082,022 | | *ORMDL3* | -4.007 | 1.03E-04 | 6.40E-04 |
| | | 38,082,035 | | *ORMDL3* | -3.929 | 1.38E-04 | 6.40E-04 |
| rs12940986 | g.37836581.A>G | 37,828,195 | | *PGAP3* | -5.004 | 1.79E-06 | |
| | | 37,828,197 | | *PGAP3* | -4.400 | 2.23E-05 | |
| | | 37,828,220 | | *PGAP3* | -4.879 | 3.06E-06 | |
| | | 37,828,273 | | *PGAP3* | -4.644 | 8.25E-06 | 0.010 |
| | | 37,828,283 | | *PGAP3* | -3.997 | 1.07E-04 | 0.010 |
| | | 37,828,296 | | *PGAP3* | -5.610 | 1.16E-07 | 0.002 |
| | | 37,828,338 | | *PGAP3* | -5.217 | 6.98E-07 | 0.002 |
| | | 37,828,377 | | *PGAP3* | -6.760 | 4.21E-10 | 0.002 |
| | | 37,828,384 | | *PGAP3* | -6.036 | 1.55E-08 | 0.002 |

| | | | | | | | |
|---|---|---|---|---|---|---|---|
| | | 37,828,560 | | *PGAP3* | -4.336 | 2.88E-05 | 0.002 |
| | | 37,843,658 | *ERBB2* | *PGAP3* | -4.414 | 2.11E-05 | 7.82E-05 |
| | | 37,844,480 | *PGAP3* | *ERBB2* | 4.015 | 9.98E-05 | 7.82E-05 |
| | | 38,072,514 | | *GSDMB* | -3.544 | 5.48E-04 | 0.046 |
| | | 38,077,039 | | | -3.374 | 9.77E-04 | 0.034 |
| | | 38,082,022 | | *ORMDL3* | -4.007 | 1.03E-04 | 6.40E-04 |
| | | 38,082,035 | | *ORMDL3* | -3.929 | 1.38E-04 | 6.40E-04 |
| rs2517952 | g.37838301.A>G | 37,828,195 | | *PGAP3* | -5.016 | 1.70E-06 | |
| | | 37,828,197 | | *PGAP3* | -4.142 | 6.15E-05 | |
| | | 37,828,220 | | *PGAP3* | -4.815 | 4.03E-06 | |
| | | 37,828,273 | | *PGAP3* | -4.350 | 2.72E-05 | 0.010 |
| | | 37,828,283 | | *PGAP3* | -3.941 | 1.32E-04 | 0.010 |
| | | 37,828,296 | | *PGAP3* | -5.849 | 3.79E-08 | 0.002 |
| | | 37,828,338 | | *PGAP3* | -5.116 | 1.09E-06 | 0.002 |
| | | 37,828,377 | | *PGAP3* | -6.790 | 3.60E-10 | 0.002 |
| | | 37,828,384 | | *PGAP3* | -6.144 | 9.12E-09 | 0.002 |
| | | 37,828,560 | | *PGAP3* | -4.411 | 2.14E-05 | 0.002 |
| | | 37,843,658 | *ERBB2* | *PGAP3* | -3.662 | 3.62E-04 | 7.82E-05 |
| | | 38,072,514 | | *GSDMB* | -3.691 | 3.27E-04 | 0.046 |
| | | 38,077,039 | | | -3.551 | 5.36E-04 | 0.034 |
| | | 38,082,022 | | *ORMDL3* | -3.901 | 1.53E-04 | 6.40E-04 |
| rs2517957 | g.37838716.A>G | 37,828,195 | | *PGAP3* | -5.004 | 1.79E-06 | |
| | | 37,828,197 | | *PGAP3* | -4.400 | 2.23E-05 | |
| | | 37,828,220 | | *PGAP3* | -4.879 | 3.06E-06 | |
| | | 37,828,273 | | *PGAP3* | -4.644 | 8.25E-06 | 0.010 |
| | | 37,828,283 | | *PGAP3* | -3.997 | 1.07E-04 | 0.010 |
| | | 37,828,296 | | *PGAP3* | -5.610 | 1.16E-07 | 0.002 |
| | | 37,828,338 | | *PGAP3* | -5.217 | 6.98E-07 | 0.002 |
| | | 37,828,377 | | *PGAP3* | -6.760 | 4.21E-10 | 0.002 |
| | | 37,828,384 | | *PGAP3* | -6.036 | 1.55E-08 | 0.002 |
| | | 37,828,560 | | *PGAP3* | -4.336 | 2.88E-05 | 0.002 |
| | | 37,843,658 | *ERBB2* | *PGAP3* | -4.414 | 2.11E-05 | 7.82E-05 |
| | | 37,844,480 | *PGAP3* | *ERBB2* | 4.015 | 9.98E-05 | 7.82E-05 |
| | | 38,072,514 | | *GSDMB* | -3.544 | 5.48E-04 | 0.046 |
| | | 38,077,039 | | | -3.374 | 9.77E-04 | 0.034 |
| | | 38,082,022 | | *ORMDL3* | -4.007 | 1.03E-04 | 6.40E-04 |
| | | 38,082,035 | | *ORMDL3* | -3.929 | 1.38E-04 | 6.40E-04 |
| rs2517958 | g.37838751.A>G | 37,828,195 | | *PGAP3* | -5.004 | 1.79E-06 | |
| | | 37,828,197 | | *PGAP3* | -4.400 | 2.23E-05 | |
| | | 37,828,220 | | *PGAP3* | -4.879 | 3.06E-06 | |
| | | 37,828,273 | | *PGAP3* | -4.644 | 8.25E-06 | 0.010 |

| | | | | | | | |
|---|---|---|---|---|---|---|---|
| | | 37,828,283 | | *PGAP3* | -3.997 | 1.07E-04 | 0.010 |
| | | 37,828,296 | | *PGAP3* | -5.610 | 1.16E-07 | 0.002 |
| | | 37,828,338 | | *PGAP3* | -5.217 | 6.98E-07 | 0.002 |
| | | 37,828,377 | | *PGAP3* | -6.760 | 4.21E-10 | 0.002 |
| | | 37,828,384 | | *PGAP3* | -6.036 | 1.55E-08 | 0.002 |
| | | 37,828,560 | | *PGAP3* | -4.336 | 2.88E-05 | 0.002 |
| | | 37,843,658 | *ERBB2* | *PGAP3* | -4.414 | 2.11E-05 | 7.82E-05 |
| | | 37,844,480 | *PGAP3* | *ERBB2* | 4.015 | 9.98E-05 | 7.82E-05 |
| | | 38,072,514 | | *GSDMB* | -3.544 | 5.48E-04 | 0.046 |
| | | 38,077,039 | | | -3.374 | 9.77E-04 | 0.034 |
| | | 38,082,022 | | *ORMDL3* | -4.007 | 1.03E-04 | 6.40E-04 |
| | | 38,082,035 | | *ORMDL3* | -3.929 | 1.38E-04 | 6.40E-04 |
| rs903501 | g.37839493.A>G | 37,828,195 | | *PGAP3* | -5.022 | 1.65E-06 | |
| | | 37,828,197 | | *PGAP3* | -4.144 | 6.11E-05 | |
| | | 37,828,220 | | *PGAP3* | -4.799 | 4.30E-06 | |
| | | 37,828,273 | | *PGAP3* | -4.310 | 3.20E-05 | 0.010 |
| | | 37,828,283 | | *PGAP3* | -3.922 | 1.42E-04 | 0.010 |
| | | 37,828,296 | | *PGAP3* | -5.818 | 4.39E-08 | 0.002 |
| | | 37,828,338 | | *PGAP3* | -5.087 | 1.24E-06 | 0.002 |
| | | 37,828,377 | | *PGAP3* | -6.756 | 4.29E-10 | 0.002 |
| | | 37,828,384 | | *PGAP3* | -6.111 | 1.07E-08 | 0.002 |
| | | 37,828,560 | | *PGAP3* | -4.412 | 2.13E-05 | 0.002 |
| | | 37,843,658 | *ERBB2* | *PGAP3* | -3.649 | 3.79E-04 | 7.82E-05 |
| | | 38,072,514 | | *GSDMB* | -3.699 | 3.18E-04 | 0.046 |
| | | 38,077,039 | | | -3.572 | 4.97E-04 | 0.034 |
| | | 38,082,022 | | *ORMDL3* | -3.917 | 1.44E-04 | 6.40E-04 |
| rs2517953 | g.37841211.C>G | 37,828,195 | | *PGAP3* | -4.956 | 2.20E-06 | |
| | | 37,828,197 | | *PGAP3* | -4.648 | 8.11E-06 | |
| | | 37,828,220 | | *PGAP3* | -4.817 | 3.99E-06 | |
| | | 37,828,273 | | *PGAP3* | -4.760 | 5.07E-06 | 0.010 |
| | | 37,828,283 | | *PGAP3* | -3.937 | 1.34E-04 | 0.010 |
| | | 37,828,296 | | *PGAP3* | -5.864 | 3.52E-08 | 0.002 |
| | | 37,828,338 | | *PGAP3* | -5.243 | 6.22E-07 | 0.002 |
| | | 37,828,377 | | *PGAP3* | -6.800 | 3.43E-10 | 0.002 |
| | | 37,828,384 | | *PGAP3* | -6.073 | 1.29E-08 | 0.002 |
| | | 37,828,560 | | *PGAP3* | -4.452 | 1.82E-05 | 0.002 |
| | | 37,843,658 | *ERBB2* | *PGAP3* | -4.466 | 1.71E-05 | 7.82E-05 |
| | | 37,844,480 | *PGAP3* | *ERBB2* | 4.008 | 1.03E-04 | 7.82E-05 |
| | | 38,072,514 | | *GSDMB* | -3.630 | 4.06E-04 | 0.046 |
| | | 38,082,022 | | *ORMDL3* | -4.108 | 7.02E-05 | 6.40E-04 |
| | | 38,082,035 | | *ORMDL3* | -3.741 | 2.74E-04 | 6.40E-04 |

| | | | | | | | |
|---|---|---|---|---|---|---|---|
| rs2517954 | g.37843550.T>C | 37,828,195 | | *PGAP3* | -5.015 | 1.70E-06 | |
| | | 37,828,197 | | *PGAP3* | -4.367 | 2.55E-05 | |
| | | 37,828,220 | | *PGAP3* | -4.726 | 5.86E-06 | |
| | | 37,828,273 | | *PGAP3* | -4.268 | 3.77E-05 | 0.010 |
| | | 37,828,283 | | *PGAP3* | -3.745 | 2.70E-04 | 0.010 |
| | | 37,828,296 | | *PGAP3* | -5.780 | 5.24E-08 | 0.002 |
| | | 37,828,338 | | *PGAP3* | -4.945 | 2.31E-06 | 0.002 |
| | | 37,828,377 | | *PGAP3* | -6.574 | 1.08E-09 | 0.002 |
| | | 37,828,384 | | *PGAP3* | -6.005 | 1.79E-08 | 0.002 |
| | | 37,828,560 | | *PGAP3* | -4.424 | 2.03E-05 | 0.002 |
| | | 37,843,658 | *ERBB2* | *PGAP3* | -3.601 | 4.50E-04 | 7.82E-05 |
| | | 38,072,514 | | *GSDMB* | -3.779 | 2.38E-04 | 0.046 |
| | | 38,077,039 | | | -3.504 | 6.29E-04 | 0.034 |
| | | 38,082,022 | | *ORMDL3* | -3.994 | 1.08E-04 | 6.40E-04 |
| rs2517955 | g.37843681.T>C | 37,828,195 | | *PGAP3* | -4.998 | 1.83E-06 | |
| | | 37,828,197 | | *PGAP3* | -4.627 | 8.87E-06 | |
| | | 37,828,220 | | *PGAP3* | -4.807 | 4.17E-06 | |
| | | 37,828,273 | | *PGAP3* | -4.602 | 9.81E-06 | 0.010 |
| | | 37,828,283 | | *PGAP3* | -3.821 | 2.05E-04 | 0.010 |
| | | 37,828,296 | | *PGAP3* | -5.575 | 1.37E-07 | 0.002 |
| | | 37,828,338 | | *PGAP3* | -5.075 | 1.31E-06 | 0.002 |
| | | 37,828,377 | | *PGAP3* | -6.581 | 1.04E-09 | 0.002 |
| | | 37,828,384 | | *PGAP3* | -5.933 | 2.54E-08 | 0.002 |
| | | 37,828,560 | | *PGAP3* | -4.349 | 2.74E-05 | 0.002 |
| | | 37,843,658 | *ERBB2* | *PGAP3* | -4.363 | 2.59E-05 | 7.82E-05 |
| | | 37,844,480 | *PGAP3* | *ERBB2* | 3.942 | 1.31E-04 | 7.82E-05 |
| | | 38,072,514 | | *GSDMB* | -3.623 | 4.16E-04 | 0.046 |
| | | 38,082,022 | | *ORMDL3* | -4.085 | 7.67E-05 | 6.40E-04 |
| | | 38,082,035 | | *ORMDL3* | -3.770 | 2.47E-04 | 6.40E-04 |
| rs2517956 | g.37843859.G>A | 37,828,195 | | *PGAP3* | -5.015 | 1.70E-06 | |
| | | 37,828,197 | | *PGAP3* | -4.367 | 2.55E-05 | |
| | | 37,828,220 | | *PGAP3* | -4.726 | 5.86E-06 | |
| | | 37,828,273 | | *PGAP3* | -4.268 | 3.77E-05 | 0.010 |
| | | 37,828,283 | | *PGAP3* | -3.745 | 2.70E-04 | 0.010 |
| | | 37,828,296 | | *PGAP3* | -5.780 | 5.24E-08 | 0.002 |
| | | 37,828,338 | | *PGAP3* | -4.945 | 2.31E-06 | 0.002 |
| | | 37,828,377 | | *PGAP3* | -6.574 | 1.08E-09 | 0.002 |
| | | 37,828,384 | | *PGAP3* | -6.005 | 1.79E-08 | 0.002 |
| | | 37,828,560 | | *PGAP3* | -4.424 | 2.03E-05 | 0.002 |
| | | 37,843,658 | *ERBB2* | *PGAP3* | -3.601 | 4.50E-04 | 7.82E-05 |
| | | 38,072,514 | | *GSDMB* | -3.779 | 2.38E-04 | 0.046 |

| | | | | | | | |
|---|---|---|---|---|---|---|---|
| | | 38,077,039 | | | -3.504 | 6.29E-04 | 0.034 |
| | | 38,082,022 | | *ORMDL3* | -3.994 | 1.08E-04 | 6.40E-04 |
| rs2517959 | g.37846512.A>T | 37,828,195 | | *PGAP3* | -5.015 | 1.70E-06 | |
| | | 37,828,197 | | *PGAP3* | -4.367 | 2.55E-05 | |
| | | 37,828,220 | | *PGAP3* | -4.726 | 5.86E-06 | |
| | | 37,828,273 | | *PGAP3* | -4.268 | 3.77E-05 | 0.010 |
| | | 37,828,283 | | *PGAP3* | -3.745 | 2.70E-04 | 0.010 |
| | | 37,828,296 | | *PGAP3* | -5.780 | 5.24E-08 | 0.002 |
| | | 37,828,338 | | *PGAP3* | -4.945 | 2.31E-06 | 0.002 |
| | | 37,828,377 | | *PGAP3* | -6.574 | 1.08E-09 | 0.002 |
| | | 37,828,384 | | *PGAP3* | -6.005 | 1.79E-08 | 0.002 |
| | | 37,828,560 | | *PGAP3* | -4.424 | 2.03E-05 | 0.002 |
| | | 37,843,658 | *ERBB2* | *PGAP3* | -3.601 | 4.50E-04 | 7.82E-05 |
| | | 38,072,514 | | *GSDMB* | -3.779 | 2.38E-04 | 0.046 |
| | | 38,077,039 | | | -3.504 | 6.29E-04 | 0.034 |
| | | 38,082,022 | | *ORMDL3* | -3.994 | 1.08E-04 | 6.40E-04 |
| rs2517960 | g.37846521.C>T | 37,828,195 | | *PGAP3* | -4.998 | 1.83E-06 | |
| | | 37,828,197 | | *PGAP3* | -4.627 | 8.87E-06 | |
| | | 37,828,220 | | *PGAP3* | -4.807 | 4.17E-06 | |
| | | 37,828,273 | | *PGAP3* | -4.602 | 9.81E-06 | 0.010 |
| | | 37,828,283 | | *PGAP3* | -3.821 | 2.05E-04 | 0.010 |
| | | 37,828,296 | | *PGAP3* | -5.575 | 1.37E-07 | 0.002 |
| | | 37,828,338 | | *PGAP3* | -5.075 | 1.31E-06 | 0.002 |
| | | 37,828,377 | | *PGAP3* | -6.581 | 1.04E-09 | 0.002 |
| | | 37,828,384 | | *PGAP3* | -5.933 | 2.54E-08 | 0.002 |
| | | 37,828,560 | | *PGAP3* | -4.349 | 2.74E-05 | 0.002 |
| | | 37,843,658 | *ERBB2* | *PGAP3* | -4.363 | 2.59E-05 | 7.82E-05 |
| | | 37,844,480 | *PGAP3* | *ERBB2* | 3.942 | 1.31E-04 | 7.82E-05 |
| | | 38,072,514 | | *GSDMB* | -3.623 | 4.16E-04 | 0.046 |
| | | 38,082,022 | | *ORMDL3* | -4.085 | 7.67E-05 | 6.40E-04 |
| | | 38,082,035 | | *ORMDL3* | -3.770 | 2.47E-04 | 6.40E-04 |
| rs2517961 | g.37847457.A>C | 37,911,152 | | | 3.786 | 2.33E-04 | 0.035 |
| rs557885708 | g.37847799_37847800ins AAA | 37,828,195 | | *PGAP3* | -5.015 | 1.70E-06 | |
| | | 37,828,197 | | *PGAP3* | -4.367 | 2.55E-05 | |
| | | 37,828,220 | | *PGAP3* | -4.726 | 5.86E-06 | |
| | | 37,828,273 | | *PGAP3* | -4.268 | 3.77E-05 | 0.010 |
| | | 37,828,283 | | *PGAP3* | -3.745 | 2.70E-04 | 0.010 |
| | | 37,828,296 | | *PGAP3* | -5.780 | 5.24E-08 | 0.002 |
| | | 37,828,338 | | *PGAP3* | -4.945 | 2.31E-06 | 0.002 |
| | | 37,828,377 | | *PGAP3* | -6.574 | 1.08E-09 | 0.002 |

| | | | | | | | |
|---|---|---|---|---|---|---|---|
| | | 37,828,384 | | *PGAP3* | -6.005 | 1.79E-08 | 0.002 |
| | | 37,828,560 | | *PGAP3* | -4.424 | 2.03E-05 | 0.002 |
| | | 37,843,658 | *ERBB2* | *PGAP3* | -3.601 | 4.50E-04 | 7.82E-05 |
| | | 38,072,514 | | *GSDMB* | -3.779 | 2.38E-04 | 0.046 |
| | | 38,077,039 | | | -3.504 | 6.29E-04 | 0.034 |
| | | 38,082,022 | | *ORMDL3* | -3.994 | 1.08E-04 | 6.40E-04 |
| rs2904765 | g.37848675.T>G | 37,828,195 | | *PGAP3* | -4.970 | 2.07E-06 | |
| | | 37,828,197 | | *PGAP3* | -4.264 | 3.83E-05 | |
| | | 37,828,220 | | *PGAP3* | -4.677 | 7.19E-06 | |
| | | 37,828,273 | | *PGAP3* | -4.335 | 2.89E-05 | 0.010 |
| | | 37,828,283 | | *PGAP3* | -3.708 | 3.08E-04 | 0.010 |
| | | 37,828,296 | | *PGAP3* | -5.692 | 7.93E-08 | 0.002 |
| | | 37,828,338 | | *PGAP3* | -4.803 | 4.23E-06 | 0.002 |
| | | 37,828,377 | | *PGAP3* | -6.477 | 1.76E-09 | 0.002 |
| | | 37,828,384 | | *PGAP3* | -5.933 | 2.53E-08 | 0.002 |
| | | 37,828,560 | | *PGAP3* | -4.382 | 2.40E-05 | 0.002 |
| | | 37,843,658 | *ERBB2* | *PGAP3* | -3.549 | 5.39E-04 | 7.82E-05 |
| | | 38,072,514 | | *GSDMB* | -3.558 | 5.22E-04 | 0.046 |
| | | 38,077,039 | | | -3.392 | 9.18E-04 | 0.034 |
| | | 38,082,022 | | *ORMDL3* | -4.007 | 1.03E-04 | 6.40E-04 |
| rs2904766 | g.37848677.A>G | 37,828,195 | | *PGAP3* | -5.015 | 1.70E-06 | |
| | | 37,828,197 | | *PGAP3* | -4.367 | 2.55E-05 | |
| | | 37,828,220 | | *PGAP3* | -4.726 | 5.86E-06 | |
| | | 37,828,273 | | *PGAP3* | -4.268 | 3.77E-05 | 0.010 |
| | | 37,828,283 | | *PGAP3* | -3.745 | 2.70E-04 | 0.010 |
| | | 37,828,296 | | *PGAP3* | -5.780 | 5.24E-08 | 0.002 |
| | | 37,828,338 | | *PGAP3* | -4.945 | 2.31E-06 | 0.002 |
| | | 37,828,377 | | *PGAP3* | -6.574 | 1.08E-09 | 0.002 |
| | | 37,828,384 | | *PGAP3* | -6.005 | 1.79E-08 | 0.002 |
| | | 37,828,560 | | *PGAP3* | -4.424 | 2.03E-05 | 0.002 |
| | | 37,843,658 | *ERBB2* | *PGAP3* | -3.601 | 4.50E-04 | 7.82E-05 |
| | | 38,072,514 | | *GSDMB* | -3.779 | 2.38E-04 | 0.046 |
| | | 38,077,039 | | | -3.504 | 6.29E-04 | 0.034 |
| | | 38,082,022 | | *ORMDL3* | -3.994 | 1.08E-04 | 6.40E-04 |
| rs55717377 | g.37850569.C>T | 37,828,195 | | *PGAP3* | -5.009 | 1.74E-06 | |
| | | 37,828,197 | | *PGAP3* | -4.365 | 2.57E-05 | |
| | | 37,828,220 | | *PGAP3* | -4.741 | 5.50E-06 | |
| | | 37,828,273 | | *PGAP3* | -4.309 | 3.21E-05 | 0.010 |
| | | 37,828,283 | | *PGAP3* | -3.764 | 2.52E-04 | 0.010 |
| | | 37,828,296 | | *PGAP3* | -5.811 | 4.53E-08 | 0.002 |
| | | 37,828,338 | | *PGAP3* | -4.974 | 2.04E-06 | 0.002 |

| | | | | | | | |
|---|---|---|---|---|---|---|---|
| | | 37,828,377 | | *PGAP3* | -6.608 | 9.11E-10 | 0.002 |
| | | 37,828,384 | | *PGAP3* | -6.038 | 1.53E-08 | 0.002 |
| | | 37,828,560 | | *PGAP3* | -4.422 | 2.04E-05 | 0.002 |
| | | 37,843,658 | *ERBB2* | *PGAP3* | -3.614 | 4.30E-04 | 7.82E-05 |
| | | 38,072,514 | | *GSDMB* | -3.771 | 2.46E-04 | 0.046 |
| | | 38,077,039 | | | -3.483 | 6.77E-04 | 0.034 |
| | | 38,082,022 | | *ORMDL3* | -3.978 | 1.15E-04 | 6.40E-04 |
| rs2904768 | g.37850571.C>T | 37,828,195 | | *PGAP3* | -5.009 | 1.74E-06 | |
| | | 37,828,197 | | *PGAP3* | -4.365 | 2.57E-05 | |
| | | 37,828,220 | | *PGAP3* | -4.741 | 5.50E-06 | |
| | | 37,828,273 | | *PGAP3* | -4.309 | 3.21E-05 | 0.010 |
| | | 37,828,283 | | *PGAP3* | -3.764 | 2.52E-04 | 0.010 |
| | | 37,828,296 | | *PGAP3* | -5.811 | 4.53E-08 | 0.002 |
| | | 37,828,338 | | *PGAP3* | -4.974 | 2.04E-06 | 0.002 |
| | | 37,828,377 | | *PGAP3* | -6.608 | 9.11E-10 | 0.002 |
| | | 37,828,384 | | *PGAP3* | -6.038 | 1.53E-08 | 0.002 |
| | | 37,828,560 | | *PGAP3* | -4.422 | 2.04E-05 | 0.002 |
| | | 37,843,658 | *ERBB2* | *PGAP3* | -3.614 | 4.30E-04 | 7.82E-05 |
| | | 38,072,514 | | *GSDMB* | -3.771 | 2.46E-04 | 0.046 |
| | | 38,077,039 | | | -3.483 | 6.77E-04 | 0.034 |
| | | 38,082,022 | | *ORMDL3* | -3.978 | 1.15E-04 | 6.40E-04 |
| rs2643194 | g.37853048.C>T | 37,828,195 | | *PGAP3* | -5.009 | 1.74E-06 | |
| | | 37,828,197 | | *PGAP3* | -4.365 | 2.57E-05 | |
| | | 37,828,220 | | *PGAP3* | -4.741 | 5.50E-06 | |
| | | 37,828,273 | | *PGAP3* | -4.309 | 3.21E-05 | 0.010 |
| | | 37,828,283 | | *PGAP3* | -3.764 | 2.52E-04 | 0.010 |
| | | 37,828,296 | | *PGAP3* | -5.811 | 4.53E-08 | 0.002 |
| | | 37,828,338 | | *PGAP3* | -4.974 | 2.04E-06 | 0.002 |
| | | 37,828,377 | | *PGAP3* | -6.608 | 9.11E-10 | 0.002 |
| | | 37,828,384 | | *PGAP3* | -6.038 | 1.53E-08 | 0.002 |
| | | 37,828,560 | | *PGAP3* | -4.422 | 2.04E-05 | 0.002 |
| | | 37,843,658 | *ERBB2* | *PGAP3* | -3.614 | 4.30E-04 | 7.82E-05 |
| | | 38,072,514 | | *GSDMB* | -3.771 | 2.46E-04 | 0.046 |
| | | 38,077,039 | | | -3.483 | 6.77E-04 | 0.034 |
| | | 38,082,022 | | *ORMDL3* | -3.978 | 1.15E-04 | 6.40E-04 |
| rs2517951 | g.37853097.T>C | 37,828,195 | | *PGAP3* | -4.897 | 2.84E-06 | |
| | | 37,828,197 | | *PGAP3* | -4.454 | 1.79E-05 | |
| | | 37,828,220 | | *PGAP3* | -4.791 | 4.45E-06 | |
| | | 37,828,273 | | *PGAP3* | -4.531 | 1.31E-05 | 0.010 |
| | | 37,828,283 | | *PGAP3* | -3.907 | 1.50E-04 | 0.010 |
| | | 37,828,296 | | *PGAP3* | -5.454 | 2.40E-07 | 0.002 |

| | | | | | | | |
|---|---|---|---|---|---|---|---|
| | | 37,828,338 | | *PGAP3* | -5.023 | 1.64E-06 | 0.002 |
| | | 37,828,377 | | *PGAP3* | -6.720 | 5.15E-10 | 0.002 |
| | | 37,828,384 | | *PGAP3* | -6.100 | 1.13E-08 | 0.002 |
| | | 37,828,560 | | *PGAP3* | -4.360 | 2.62E-05 | 0.002 |
| | | 37,843,658 | *ERBB2* | *PGAP3* | -4.317 | 3.11E-05 | 7.82E-05 |
| | | 37,844,480 | *PGAP3* | *ERBB2* | 3.895 | 1.56E-04 | 7.82E-05 |
| | | 38,072,514 | | *GSDMB* | -3.839 | 1.92E-04 | 0.046 |
| | | 38,082,022 | | *ORMDL3* | -4.033 | 9.32E-05 | 6.40E-04 |
| | | 38,082,035 | | *ORMDL3* | -3.724 | 2.91E-04 | 6.40E-04 |
| rs2643195 | g.37853118.A>G | 37,828,195 | | *PGAP3* | -4.911 | 2.67E-06 | |
| | | 37,828,197 | | *PGAP3* | -4.198 | 4.96E-05 | |
| | | 37,828,220 | | *PGAP3* | -4.712 | 6.22E-06 | |
| | | 37,828,273 | | *PGAP3* | -4.190 | 5.11E-05 | 0.010 |
| | | 37,828,283 | | *PGAP3* | -3.827 | 2.01E-04 | 0.010 |
| | | 37,828,296 | | *PGAP3* | -5.651 | 9.61E-08 | 0.002 |
| | | 37,828,338 | | *PGAP3* | -4.888 | 2.95E-06 | 0.002 |
| | | 37,828,377 | | *PGAP3* | -6.700 | 5.72E-10 | 0.002 |
| | | 37,828,384 | | *PGAP3* | -6.162 | 8.39E-09 | 0.002 |
| | | 37,828,560 | | *PGAP3* | -4.432 | 1.96E-05 | 0.002 |
| | | 37,843,658 | *ERBB2* | *PGAP3* | -3.551 | 5.35E-04 | 7.82E-05 |
| | | 38,072,514 | | *GSDMB* | -4.000 | 1.06E-04 | 0.046 |
| | | 38,077,039 | | | -3.472 | 7.03E-04 | 0.034 |
| | | 38,082,022 | | *ORMDL3* | -3.945 | 1.30E-04 | 6.40E-04 |
| rs2934971 | g.37854507.G>T | 37,828,195 | | *PGAP3* | -5.009 | 1.74E-06 | |
| | | 37,828,197 | | *PGAP3* | -4.365 | 2.57E-05 | |
| | | 37,828,220 | | *PGAP3* | -4.741 | 5.50E-06 | |
| | | 37,828,273 | | *PGAP3* | -4.309 | 3.21E-05 | 0.010 |
| | | 37,828,283 | | *PGAP3* | -3.764 | 2.52E-04 | 0.010 |
| | | 37,828,296 | | *PGAP3* | -5.811 | 4.53E-08 | 0.002 |
| | | 37,828,338 | | *PGAP3* | -4.974 | 2.04E-06 | 0.002 |
| | | 37,828,377 | | *PGAP3* | -6.608 | 9.11E-10 | 0.002 |
| | | 37,828,384 | | *PGAP3* | -6.038 | 1.53E-08 | 0.002 |
| | | 37,828,560 | | *PGAP3* | -4.422 | 2.04E-05 | 0.002 |
| | | 37,843,658 | *ERBB2* | *PGAP3* | -3.614 | 4.30E-04 | 7.82E-05 |
| | | 38,072,514 | | *GSDMB* | -3.771 | 2.46E-04 | 0.046 |
| | | 38,077,039 | | | -3.483 | 6.77E-04 | 0.034 |
| | | 38,082,022 | | *ORMDL3* | -3.978 | 1.15E-04 | 6.40E-04 |
| rs1565923 | g.37858678.A>G | 37,828,195 | | *PGAP3* | -5.009 | 1.74E-06 | |
| | | 37,828,197 | | *PGAP3* | -4.365 | 2.57E-05 | |
| | | 37,828,220 | | *PGAP3* | -4.741 | 5.50E-06 | |
| | | 37,828,273 | | *PGAP3* | -4.309 | 3.21E-05 | 0.010 |

| | | | | | | | |
|---|---|---|---|---|---|---|---|
| | | 37,828,283 | | *PGAP3* | -3.764 | 2.52E-04 | 0.010 |
| | | 37,828,296 | | *PGAP3* | -5.811 | 4.53E-08 | 0.002 |
| | | 37,828,338 | | *PGAP3* | -4.974 | 2.04E-06 | 0.002 |
| | | 37,828,377 | | *PGAP3* | -6.608 | 9.11E-10 | 0.002 |
| | | 37,828,384 | | *PGAP3* | -6.038 | 1.53E-08 | 0.002 |
| | | 37,828,560 | | *PGAP3* | -4.422 | 2.04E-05 | 0.002 |
| | | 37,843,658 | *ERBB2* | *PGAP3* | -3.614 | 4.30E-04 | 7.82E-05 |
| | | 38,072,514 | | *GSDMB* | -3.771 | 2.46E-04 | 0.046 |
| | | 38,077,039 | | | -3.483 | 6.77E-04 | 0.034 |
| | | 38,082,022 | | *ORMDL3* | -3.978 | 1.15E-04 | 6.40E-04 |
| rs2952155 | g.37861718.T>C | 37,828,195 | | *PGAP3* | -3.720 | 2.96E-04 | |
| | | 37,828,220 | | *PGAP3* | -4.110 | 6.95E-05 | |
| | | 37,828,273 | | *PGAP3* | -3.367 | 9.98E-04 | 0.010 |
| | | 37,828,283 | | *PGAP3* | -3.640 | 3.92E-04 | 0.010 |
| | | 37,828,296 | | *PGAP3* | -5.036 | 1.55E-06 | 0.002 |
| | | 37,828,338 | | *PGAP3* | -4.650 | 8.03E-06 | 0.002 |
| | | 37,828,377 | | *PGAP3* | -5.806 | 4.63E-08 | 0.002 |
| | | 37,828,384 | | *PGAP3* | -5.355 | 3.75E-07 | 0.002 |
| | | 37,828,560 | | *PGAP3* | -4.837 | 3.67E-06 | 0.002 |
| | | 37,902,268 | | | -3.620 | 4.20E-04 | 0.008 |
| | | 38,109,098 | | | 3.504 | 6.28E-04 | 0.002 |
| rs1810132 | g.37866005.C>T | 37,828,195 | | *PGAP3* | -5.009 | 1.74E-06 | |
| | | 37,828,197 | | *PGAP3* | -4.365 | 2.57E-05 | |
| | | 37,828,220 | | *PGAP3* | -4.741 | 5.50E-06 | |
| | | 37,828,273 | | *PGAP3* | -4.309 | 3.21E-05 | 0.010 |
| | | 37,828,283 | | *PGAP3* | -3.764 | 2.52E-04 | 0.010 |
| | | 37,828,296 | | *PGAP3* | -5.811 | 4.53E-08 | 0.002 |
| | | 37,828,338 | | *PGAP3* | -4.974 | 2.04E-06 | 0.002 |
| | | 37,828,377 | | *PGAP3* | -6.608 | 9.11E-10 | 0.002 |
| | | 37,828,384 | | *PGAP3* | -6.038 | 1.53E-08 | 0.002 |
| | | 37,828,560 | | *PGAP3* | -4.422 | 2.04E-05 | 0.002 |
| | | 37,843,658 | *ERBB2* | *PGAP3* | -3.614 | 4.30E-04 | 7.82E-05 |
| | | 38,072,514 | | *GSDMB* | -3.771 | 2.46E-04 | 0.046 |
| | | 38,077,039 | | | -3.483 | 6.77E-04 | 0.034 |
| | | 38,082,022 | | *ORMDL3* | -3.978 | 1.15E-04 | 6.40E-04 |
| rs4252627 | g.37868715.C>T | 37,828,195 | | *PGAP3* | -4.994 | 1.86E-06 | |
| | | 37,828,197 | | *PGAP3* | -4.627 | 8.87E-06 | |
| | | 37,828,220 | | *PGAP3* | -4.824 | 3.88E-06 | |
| | | 37,828,273 | | *PGAP3* | -4.646 | 8.20E-06 | 0.010 |
| | | 37,828,283 | | *PGAP3* | -3.841 | 1.90E-04 | 0.010 |
| | | 37,828,296 | | *PGAP3* | -5.607 | 1.18E-07 | 0.002 |

| | | | | | | | |
|---|---|---|---|---|---|---|---|
| | | 37,828,338 | | *PGAP3* | -5.106 | 1.14E-06 | 0.002 |
| | | 37,828,377 | | *PGAP3* | -6.617 | 8.69E-10 | 0.002 |
| | | 37,828,384 | | *PGAP3* | -5.968 | 2.15E-08 | 0.002 |
| | | 37,828,560 | | *PGAP3* | -4.349 | 2.74E-05 | 0.002 |
| | | 37,843,658 | *ERBB2* | *PGAP3* | -4.378 | 2.44E-05 | 7.82E-05 |
| | | 37,844,480 | *PGAP3* | *ERBB2* | 3.926 | 1.39E-04 | 7.82E-05 |
| | | 38,072,514 | | *GSDMB* | -3.616 | 4.26E-04 | 0.046 |
| | | 38,082,022 | | *ORMDL3* | -4.070 | 8.10E-05 | 6.40E-04 |
| | | 38,082,035 | | *ORMDL3* | -3.745 | 2.69E-04 | 6.40E-04 |
| rs2934967 | g.37870378.G>A | 37,828,195 | | *PGAP3* | -5.009 | 1.74E-06 | |
| | | 37,828,197 | | *PGAP3* | -4.365 | 2.57E-05 | |
| | | 37,828,220 | | *PGAP3* | -4.741 | 5.50E-06 | |
| | | 37,828,273 | | *PGAP3* | -4.309 | 3.21E-05 | 0.010 |
| | | 37,828,283 | | *PGAP3* | -3.764 | 2.52E-04 | 0.010 |
| | | 37,828,296 | | *PGAP3* | -5.811 | 4.53E-08 | 0.002 |
| | | 37,828,338 | | *PGAP3* | -4.974 | 2.04E-06 | 0.002 |
| | | 37,828,377 | | *PGAP3* | -6.608 | 9.11E-10 | 0.002 |
| | | 37,828,384 | | *PGAP3* | -6.038 | 1.53E-08 | 0.002 |
| | | 37,828,560 | | *PGAP3* | -4.422 | 2.04E-05 | 0.002 |
| | | 37,843,658 | *ERBB2* | *PGAP3* | -3.614 | 4.30E-04 | 7.82E-05 |
| | | 38,072,514 | | *GSDMB* | -3.771 | 2.46E-04 | 0.046 |
| | | 38,077,039 | | | -3.483 | 6.77E-04 | 0.034 |
| | | 38,082,022 | | *ORMDL3* | -3.978 | 1.15E-04 | 6.40E-04 |
| rs2952156 | g.37876835.A>G | 37,828,195 | | *PGAP3* | -4.732 | 5.72E-06 | |
| | | 37,828,197 | | *PGAP3* | -4.312 | 3.17E-05 | |
| | | 37,828,220 | | *PGAP3* | -4.735 | 5.65E-06 | |
| | | 37,828,273 | | *PGAP3* | -4.042 | 9.02E-05 | 0.010 |
| | | 37,828,283 | | *PGAP3* | -3.690 | 3.28E-04 | 0.010 |
| | | 37,828,296 | | *PGAP3* | -5.886 | 3.17E-08 | 0.002 |
| | | 37,828,338 | | *PGAP3* | -4.706 | 6.36E-06 | 0.002 |
| | | 37,828,377 | | *PGAP3* | -6.519 | 1.43E-09 | 0.002 |
| | | 37,828,384 | | *PGAP3* | -5.833 | 4.09E-08 | 0.002 |
| | | 37,828,560 | | *PGAP3* | -4.282 | 3.57E-05 | 0.002 |
| | | 37,843,658 | *ERBB2* | *PGAP3* | -3.449 | 7.59E-04 | 7.82E-05 |
| | | 38,072,514 | | *GSDMB* | -3.795 | 2.25E-04 | 0.046 |
| | | 38,077,039 | | | -3.377 | 9.67E-04 | 0.034 |
| | | 38,082,022 | | *ORMDL3* | -4.001 | 1.05E-04 | 6.40E-04 |
| | | 38,082,073 | | *ORMDL3* | -3.638 | 3.95E-04 | 6.40E-04 |
| rs2952157 | g.37877412.G>A | 37,828,195 | | *PGAP3* | -4.732 | 5.72E-06 | |
| | | 37,828,197 | | *PGAP3* | -4.312 | 3.17E-05 | |
| | | 37,828,220 | | *PGAP3* | -4.735 | 5.65E-06 | |

| | | | | | | | |
|---|---|---|---|---|---|---|---|
| | | 37,828,273 | | *PGAP3* | -4.042 | 9.02E-05 | 0.010 |
| | | 37,828,283 | | *PGAP3* | -3.690 | 3.28E-04 | 0.010 |
| | | 37,828,296 | | *PGAP3* | -5.886 | 3.17E-08 | 0.002 |
| | | 37,828,338 | | *PGAP3* | -4.706 | 6.36E-06 | 0.002 |
| | | 37,828,377 | | *PGAP3* | -6.519 | 1.43E-09 | 0.002 |
| | | 37,828,384 | | *PGAP3* | -5.833 | 4.09E-08 | 0.002 |
| | | 37,828,560 | | *PGAP3* | -4.282 | 3.57E-05 | 0.002 |
| | | 37,843,658 | *ERBB2* | *PGAP3* | -3.449 | 7.59E-04 | 7.82E-05 |
| | | 38,072,514 | | *GSDMB* | -3.795 | 2.25E-04 | 0.046 |
| | | 38,077,039 | | | -3.377 | 9.67E-04 | 0.034 |
| | | 38,082,022 | | *ORMDL3* | -4.001 | 1.05E-04 | 6.40E-04 |
| | | 38,082,073 | | *ORMDL3* | -3.638 | 3.95E-04 | 6.40E-04 |
| rs11653998 | g.37877447.G>C | 37,828,195 | | *PGAP3* | -4.722 | 5.97E-06 | |
| | | 37,828,197 | | *PGAP3* | -4.573 | 1.11E-05 | |
| | | 37,828,220 | | *PGAP3* | -4.819 | 3.95E-06 | |
| | | 37,828,273 | | *PGAP3* | -4.372 | 2.50E-05 | 0.010 |
| | | 37,828,283 | | *PGAP3* | -3.769 | 2.48E-04 | 0.010 |
| | | 37,828,296 | | *PGAP3* | -5.685 | 8.20E-08 | 0.002 |
| | | 37,828,338 | | *PGAP3* | -4.839 | 3.64E-06 | 0.002 |
| | | 37,828,377 | | *PGAP3* | -6.533 | 1.33E-09 | 0.002 |
| | | 37,828,384 | | *PGAP3* | -5.769 | 5.53E-08 | 0.002 |
| | | 37,828,560 | | *PGAP3* | -4.213 | 4.67E-05 | 0.002 |
| | | 37,843,658 | *ERBB2* | *PGAP3* | -4.199 | 4.94E-05 | 7.82E-05 |
| | | 37,844,480 | *PGAP3* | *ERBB2* | 4.045 | 8.91E-05 | 7.82E-05 |
| | | 38,072,514 | | *GSDMB* | -3.643 | 3.87E-04 | 0.046 |
| | | 38,082,022 | | *ORMDL3* | -4.094 | 7.39E-05 | 6.40E-04 |
| | | 38,082,035 | | *ORMDL3* | -3.819 | 2.07E-04 | 6.40E-04 |
| rs2088126 | g.37879030.A>G | 37,828,195 | | *PGAP3* | -4.732 | 5.72E-06 | |
| | | 37,828,197 | | *PGAP3* | -4.312 | 3.17E-05 | |
| | | 37,828,220 | | *PGAP3* | -4.735 | 5.65E-06 | |
| | | 37,828,273 | | *PGAP3* | -4.042 | 9.02E-05 | 0.010 |
| | | 37,828,283 | | *PGAP3* | -3.690 | 3.28E-04 | 0.010 |
| | | 37,828,296 | | *PGAP3* | -5.886 | 3.17E-08 | 0.002 |
| | | 37,828,338 | | *PGAP3* | -4.706 | 6.36E-06 | 0.002 |
| | | 37,828,377 | | *PGAP3* | -6.519 | 1.43E-09 | 0.002 |
| | | 37,828,384 | | *PGAP3* | -5.833 | 4.09E-08 | 0.002 |
| | | 37,828,560 | | *PGAP3* | -4.282 | 3.57E-05 | 0.002 |
| | | 37,843,658 | *ERBB2* | *PGAP3* | -3.449 | 7.59E-04 | 7.82E-05 |
| | | 38,072,514 | | *GSDMB* | -3.795 | 2.25E-04 | 0.046 |
| | | 38,077,039 | | | -3.377 | 9.67E-04 | 0.034 |
| | | 38,082,022 | | *ORMDL3* | -4.001 | 1.05E-04 | 6.40E-04 |

| | | | | | | | |
|---|---|---|---|---|---|---|---|
| | | 38,082,073 | | *ORMDL3* | -3.638 | 3.95E-04 | 6.40E-04 |
| rs903506 | g.37879762.G>A | 37,828,195 | | *PGAP3* | -4.722 | 5.97E-06 | |
| | | 37,828,197 | | *PGAP3* | -4.573 | 1.11E-05 | |
| | | 37,828,220 | | *PGAP3* | -4.819 | 3.95E-06 | |
| | | 37,828,273 | | *PGAP3* | -4.372 | 2.50E-05 | 0.010 |
| | | 37,828,283 | | *PGAP3* | -3.769 | 2.48E-04 | 0.010 |
| | | 37,828,296 | | *PGAP3* | -5.685 | 8.20E-08 | 0.002 |
| | | 37,828,338 | | *PGAP3* | -4.839 | 3.64E-06 | 0.002 |
| | | 37,828,377 | | *PGAP3* | -6.533 | 1.33E-09 | 0.002 |
| | | 37,828,384 | | *PGAP3* | -5.769 | 5.53E-08 | 0.002 |
| | | 37,828,560 | | *PGAP3* | -4.213 | 4.67E-05 | 0.002 |
| | | 37,843,658 | *ERBB2* | *PGAP3* | -4.199 | 4.94E-05 | 7.82E-05 |
| | | 37,844,480 | *PGAP3* | *ERBB2* | 4.045 | 8.91E-05 | 7.82E-05 |
| | | 38,072,514 | | *GSDMB* | -3.643 | 3.87E-04 | 0.046 |
| | | 38,082,022 | | *ORMDL3* | -4.094 | 7.39E-05 | 6.40E-04 |
| | | 38,082,035 | | *ORMDL3* | -3.819 | 2.07E-04 | 6.40E-04 |
| rs547454526 | g.37880525_37880526ins TTTTT | 37,828,195 | | *PGAP3* | -4.823 | 3.89E-06 | |
| | | 37,828,197 | | *PGAP3* | -4.198 | 4.95E-05 | |
| | | 37,828,220 | | *PGAP3* | -4.571 | 1.12E-05 | |
| | | 37,828,273 | | *PGAP3* | -4.412 | 2.13E-05 | 0.010 |
| | | 37,828,283 | | *PGAP3* | -3.708 | 3.09E-04 | 0.010 |
| | | 37,828,296 | | *PGAP3* | -5.711 | 7.26E-08 | 0.002 |
| | | 37,828,338 | | *PGAP3* | -4.696 | 6.65E-06 | 0.002 |
| | | 37,828,377 | | *PGAP3* | -6.245 | 5.57E-09 | 0.002 |
| | | 37,828,384 | | *PGAP3* | -5.746 | 6.16E-08 | 0.002 |
| | | 37,828,560 | | *PGAP3* | -4.147 | 6.03E-05 | 0.002 |
| | | 37,843,658 | *ERBB2* | *PGAP3* | -4.163 | 5.68E-05 | 7.82E-05 |
| | | 37,844,480 | *PGAP3* | *ERBB2* | 3.970 | 1.18E-04 | 7.82E-05 |
| | | 38,082,022 | | *ORMDL3* | -3.945 | 1.30E-04 | 6.40E-04 |
| | | 38,082,035 | | *ORMDL3* | -3.626 | 4.12E-04 | 6.40E-04 |
| rs1058808 | g.37884037.C>G (Pro.1155.Ala) | 37,828,195 | | *PGAP3* | -4.724 | 5.91E-06 | |
| | | 37,828,197 | | *PGAP3* | -4.564 | 1.15E-05 | |
| | | 37,828,220 | | *PGAP3* | -4.814 | 4.05E-06 | |
| | | 37,828,273 | | *PGAP3* | -4.357 | 2.65E-05 | 0.010 |
| | | 37,828,283 | | *PGAP3* | -3.766 | 2.50E-04 | 0.010 |
| | | 37,828,296 | | *PGAP3* | -5.680 | 8.41E-08 | 0.002 |
| | | 37,828,338 | | *PGAP3* | -4.845 | 3.54E-06 | 0.002 |
| | | 37,828,377 | | *PGAP3* | -6.517 | 1.44E-09 | 0.002 |
| | | 37,828,384 | | *PGAP3* | -5.763 | 5.68E-08 | 0.002 |

| | | | | | | | | |
|---|---|---|---|---|---|---|---|---|
| | | | 37,828,560 | | *PGAP3* | -4.210 | 4.72E-05 | 0.002 |
| | | | 37,843,658 | *ERBB2* | *PGAP3* | -4.192 | 5.08E-05 | 7.82E-05 |
| | | | 37,844,480 | *PGAP3* | *ERBB2* | 4.049 | 8.77E-05 | 7.82E-05 |
| | | | 38,072,514 | | *GSDMB* | -3.645 | 3.85E-04 | 0.046 |
| | | | 38,082,022 | | *ORMDL3* | -4.102 | 7.19E-05 | 6.40E-04 |
| | | | 38,082,035 | | *ORMDL3* | -3.820 | 2.06E-04 | 6.40E-04 |
| | rs34503675 | g.37885640_37885640insG | 37,895,335 | | | 3.865 | 1.75E-04 | 0.006 |
| | | | 38,096,333 | | | -4.602 | 9.82E-06 | 0.033 |
| | | | 38,096,571 | | | -6.738 | 4.70E-10 | 0.033 |
| | | | 38,096,578 | | | -7.472 | 1.03E-11 | 0.033 |
| | | | 38,096,648 | | | -4.245 | 4.12E-05 | 0.033 |
| | rs66459548 | g.37892697.A>T | 37,895,335 | | | 3.857 | 1.80E-04 | 0.006 |
| | | | 38,096,333 | | | -4.691 | 6.79E-06 | 0.033 |
| | | | 38,096,571 | | | -6.631 | 8.11E-10 | 0.033 |
| | | | 38,096,578 | | | -7.428 | 1.30E-11 | 0.033 |
| | | | 38,096,648 | | | -3.865 | 1.75E-04 | 0.033 |
| | rs67597968 | g.37893458.G>A | 37,895,335 | | | 3.857 | 1.80E-04 | 0.006 |
| | | | 38,096,333 | | | -4.691 | 6.79E-06 | 0.033 |
| | | | 38,096,571 | | | -6.631 | 8.11E-10 | 0.033 |
| | | | 38,096,578 | | | -7.428 | 1.30E-11 | 0.033 |
| | | | 38,096,648 | | | -3.865 | 1.75E-04 | 0.033 |
| Second block | rs9896218 | g.37894463.C>A | 37,895,335 | | | 3.857 | 1.80E-04 | 0.006 |
| | | | 38,096,333 | | | -4.691 | 6.79E-06 | 0.033 |
| | | | 38,096,571 | | | -6.631 | 8.11E-10 | 0.033 |
| | | | 38,096,578 | | | -7.428 | 1.30E-11 | 0.033 |
| | | | 38,096,648 | | | -3.865 | 1.75E-04 | 0.033 |
| | rs71355409 | g.37895367_37895371del | 37,886,749 | | *MIEN1* | 3.966 | 1.20E-04 | 0.002 |
| | | | 37,886,788 | *MIEN1* | *MIEN1* | 3.415 | 8.51E-04 | 0.002 |
| | | | 37,894,352 | | | -3.403 | 8.87E-04 | 0.013 |
| | | | 38,096,571 | | | -5.813 | 4.50E-08 | 0.033 |
| | | | 38,096,578 | | | -5.492 | 2.01E-07 | 0.033 |
| | rs59667301 | g.37895379.C>T | 37,895,335 | | | 3.857 | 1.80E-04 | 0.006 |
| | | | 38,096,333 | | | -4.691 | 6.79E-06 | 0.033 |
| | | | 38,096,571 | | | -6.631 | 8.11E-10 | 0.033 |
| | | | 38,096,578 | | | -7.428 | 1.30E-11 | 0.033 |
| | | | 38,096,648 | | | -3.865 | 1.75E-04 | 0.033 |
| | rs35585925 | g.37898210.C>T | 37,895,335 | | | 3.857 | 1.80E-04 | 0.006 |
| | | | 38,096,333 | | | -4.691 | 6.79E-06 | 0.033 |
| | | | 38,096,571 | | | -6.631 | 8.11E-10 | 0.033 |

| | | | | | | | | |
|---|---|---|---|---|---|---|---|---|
| | | | 38,096,578 | | | -7.428 | 1.30E-11 | 0.033 |
| | | | 38,096,648 | | | -3.865 | 1.75E-04 | 0.033 |
| | rs34093201 | g.37898249.A>G | 37,895,335 | | | 3.818 | 2.07E-04 | 0.006 |
| | | | 37,896,063 | | | 3.432 | 8.04E-04 | 0.009 |
| | | | 38,096,333 | | | -4.665 | 7.58E-06 | 0.033 |
| | | | 38,096,571 | | | -6.760 | 4.21E-10 | 0.033 |
| | | | 38,096,578 | | | -7.488 | 9.41E-12 | 0.033 |
| | | | 38,096,648 | | | -3.941 | 1.32E-04 | 0.033 |
| | rs36079893 | g.37898909.C>A (Pro.105.Pro) | 37,895,335 | | | 3.857 | 1.80E-04 | 0.006 |
| | | | 38,096,333 | | | -4.691 | 6.79E-06 | 0.033 |
| | | | 38,096,571 | | | -6.631 | 8.11E-10 | 0.033 |
| | | | 38,096,578 | | | -7.428 | 1.30E-11 | 0.033 |
| | | | 38,096,648 | | | -3.865 | 1.75E-04 | 0.033 |
| Third block | rs9747973 | g.37905107.C>T | 37,828,296 | | *PGAP3* | 4.124 | 6.59E-05 | 0.002 |
| | | | 37,916,005 | | | 3.522 | 5.92E-04 | 0.001 |
| | | | 37,927,002 | | *IKZF3* | -3.430 | 8.08E-04 | 0.001 |
| | | | 38,018,804 | | *IKZF3* | 3.618 | 4.23E-04 | 0.046 |
| | | | 38,077,191 | | | 4.058 | 8.49E-05 | 0.034 |
| | | | 38,080,829 | | *ORMDL3* | 4.895 | 2.86E-06 | 0.014 |
| | | | 38,081,492 | | *ORMDL3* | 3.598 | 4.55E-04 | 2.01E-04 |
| | | | 38,081,995 | | *ORMDL3* | 4.067 | 8.21E-05 | 2.01E-04 |
| | | | 38,082,022 | | *ORMDL3* | 4.600 | 9.91E-06 | 6.40E-04 |
| | | | 38,082,035 | | *ORMDL3* | 5.355 | 3.75E-07 | 6.40E-04 |
| | | | 38,082,073 | | *ORMDL3* | 4.569 | 1.13E-05 | 6.40E-04 |
| | | | 38,082,235 | | *ORMDL3* | 3.434 | 7.99E-04 | 0.004 |
| | | | 38,082,346 | | *ORMDL3* | 3.895 | 1.56E-04 | 0.011 |
| | | | 38,082,448 | | *ORMDL3* | 3.717 | 2.98E-04 | 0.011 |
| | | | 38,082,658 | | *ORMDL3* | 3.442 | 7.78E-04 | 0.011 |
| | | | 38,082,778 | | *ORMDL3* | 3.637 | 3.96E-04 | 0.011 |
| | | | 38,119,198 | *GSDMA* | | 3.388 | 9.31E-04 | 0.009 |
| | | | 38,119,207 | *GSDMA* | | 4.733 | 5.69E-06 | 0.009 |
| | | | 38,119,649 | | *GSDMA* | 3.572 | 4.98E-04 | 0.002 |
| | rs11078921 | g.37908867.A>C | 37,896,050 | | | 3.582 | 4.80E-04 | 0.027 |
| | | | 38,024,331 | | | -4.274 | 3.68E-05 | 0.017 |
| | | | 38,024,422 | | | -3.440 | 7.83E-04 | 3.75E-04 |
| | | | 38,072,514 | | *GSDMB* | 4.184 | 5.23E-05 | 0.046 |
| | | | 38,082,073 | | *ORMDL3* | 3.695 | 3.23E-04 | 6.40E-04 |
| | | | 38,119,198 | *GSDMA* | | 3.550 | 5.37E-04 | 0.009 |
| | | | 38,119,207 | *GSDMA* | | 3.757 | 2.59E-04 | 0.009 |
| | | | 38,119,649 | | *GSDMA* | 3.670 | 3.53E-04 | 0.002 |

| | | | | | | | |
|---|---|---|---|---|---|---|---|
| | | 38,121,965 | | *GSDMA* | -3.433 | 8.00E-04 | 0.011 |
| rs2941522 | g.37910368.G>A | 37,828,296 | | *PGAP3* | 4.124 | 6.59E-05 | 0.002 |
| | | 37,916,005 | | | 3.522 | 5.92E-04 | 0.001 |
| | | 37,927,002 | | *IKZF3* | -3.430 | 8.08E-04 | 0.001 |
| | | 38,018,804 | | *IKZF3* | 3.618 | 4.23E-04 | 0.046 |
| | | 38,077,191 | | | 4.058 | 8.49E-05 | 0.034 |
| | | 38,080,829 | | *ORMDL3* | 4.895 | 2.86E-06 | 0.014 |
| | | 38,081,492 | | *ORMDL3* | 3.598 | 4.55E-04 | 2.01E-04 |
| | | 38,081,995 | | *ORMDL3* | 4.067 | 8.21E-05 | 2.01E-04 |
| | | 38,082,022 | | *ORMDL3* | 4.600 | 9.91E-06 | 6.40E-04 |
| | | 38,082,035 | | *ORMDL3* | 5.355 | 3.75E-07 | 6.40E-04 |
| | | 38,082,073 | | *ORMDL3* | 4.569 | 1.13E-05 | 6.40E-04 |
| | | 38,082,235 | | *ORMDL3* | 3.434 | 7.99E-04 | 0.004 |
| | | 38,082,346 | | *ORMDL3* | 3.895 | 1.56E-04 | 0.011 |
| | | 38,082,448 | | *ORMDL3* | 3.717 | 2.98E-04 | 0.011 |
| | | 38,082,658 | | *ORMDL3* | 3.442 | 7.78E-04 | 0.011 |
| | | 38,082,778 | | *ORMDL3* | 3.637 | 3.96E-04 | 0.011 |
| | | 38,119,198 | *GSDMA* | | 3.388 | 9.31E-04 | 0.009 |
| | | 38,119,207 | *GSDMA* | | 4.733 | 5.69E-06 | 0.009 |
| | | 38,119,649 | | *GSDMA* | 3.572 | 4.98E-04 | 0.002 |
| rs12946510 | g.37912377.T>C | 37,828,296 | | *PGAP3* | 3.947 | 1.29E-04 | 0.002 |
| | | 37,927,002 | | *IKZF3* | -3.492 | 6.55E-04 | 0.001 |
| | | 38,072,514 | | *GSDMB* | 3.849 | 1.85E-04 | 0.046 |
| | | 38,077,191 | | | 4.118 | 6.76E-05 | 0.034 |
| | | 38,081,995 | | *ORMDL3* | 5.621 | 1.11E-07 | 2.01E-04 |
| | | 38,082,018 | | *ORMDL3* | 3.957 | 1.24E-04 | 6.40E-04 |
| | | 38,082,022 | | *ORMDL3* | 6.211 | 6.58E-09 | 6.40E-04 |
| | | 38,082,035 | | *ORMDL3* | 6.144 | 9.12E-09 | 6.40E-04 |
| | | 38,082,073 | | *ORMDL3* | 6.311 | 4.02E-09 | 6.40E-04 |
| | | 38,082,206 | | *ORMDL3* | 3.997 | 1.07E-04 | 0.004 |
| | | 38,082,235 | | *ORMDL3* | 3.937 | 1.34E-04 | 0.004 |
| | | 38,082,346 | | *ORMDL3* | 3.783 | 2.35E-04 | 0.011 |
| | | 38,082,448 | | *ORMDL3* | 3.486 | 6.69E-04 | 0.011 |
| | | 38,082,658 | | *ORMDL3* | 3.793 | 2.27E-04 | 0.011 |
| | | 38,082,778 | | *ORMDL3* | 4.118 | 6.74E-05 | 0.011 |
| | | 38,108,710 | | | -4.046 | 8.89E-05 | 0.039 |
| | | 38,119,198 | *GSDMA* | | 3.742 | 2.73E-04 | 0.009 |
| | | 38,119,207 | *GSDMA* | | 4.735 | 5.65E-06 | 0.009 |
| | | 38,119,649 | | *GSDMA* | 3.913 | 1.46E-04 | 0.002 |
| rs35833706 | g.37916390_37916390insT | 37,828,296 | | *PGAP3* | 3.791 | 2.29E-04 | 0.002 |

| | | | | | | | |
|---|---|---|---|---|---|---|---|
| | | 37,916,005 | | | 3.621 | 4.19E-04 | 0.001 |
| | | 37,927,002 | | *IKZF3* | -3.395 | 9.12E-04 | 0.001 |
| | | 38,018,804 | | *IKZF3* | 3.982 | 1.13E-04 | 0.046 |
| | | 38,049,272 | | | 3.454 | 7.47E-04 | 0.046 |
| | | 38,077,191 | | | 3.949 | 1.28E-04 | 0.034 |
| | | 38,080,829 | | *ORMDL3* | 5.188 | 7.94E-07 | 0.014 |
| | | 38,081,995 | | *ORMDL3* | 4.569 | 1.13E-05 | 2.01E-04 |
| | | 38,082,022 | | *ORMDL3* | 4.670 | 7.41E-06 | 6.40E-04 |
| | | 38,082,035 | | *ORMDL3* | 5.559 | 1.48E-07 | 6.40E-04 |
| | | 38,082,073 | | *ORMDL3* | 5.344 | 3.95E-07 | 6.40E-04 |
| | | 38,082,346 | | *ORMDL3* | 4.011 | 1.01E-04 | 0.011 |
| | | 38,082,448 | | *ORMDL3* | 3.937 | 1.34E-04 | 0.011 |
| | | 38,082,658 | | *ORMDL3* | 3.981 | 1.14E-04 | 0.011 |
| | | 38,082,778 | | *ORMDL3* | 3.591 | 4.65E-04 | 0.011 |
| | | 38,108,710 | | | -3.751 | 2.64E-04 | 0.039 |
| | | 38,119,198 | *GSDMA* | | 3.857 | 1.80E-04 | 0.009 |
| | | 38,119,207 | *GSDMA* | | 5.435 | 2.62E-07 | 0.009 |
| | | 38,119,649 | | *GSDMA* | 3.891 | 1.59E-04 | 0.002 |
| rs3764354 | g.37916823.A>G | 37,893,689 | | | 3.496 | 6.46E-04 | 0.002 |
| | | 37,926,608 | | *IKZF3* | -3.571 | 4.99E-04 | 0.001 |
| | | 38,020,208 | | *IKZF3* | 3.385 | 9.41E-04 | 0.046 |
| | | 38,025,241 | | | -3.741 | 2.74E-04 | 0.046 |
| | | 38,065,264 | | *GSDMB* | 4.319 | 3.09E-05 | 0.046 |
| | | 38,077,191 | | | 3.424 | 8.25E-04 | 0.034 |
| | | 38,081,812 | | *ORMDL3* | 3.935 | 1.35E-04 | 2.01E-04 |
| | | 38,081,829 | | *ORMDL3* | 5.046 | 1.49E-06 | 2.01E-04 |
| | | 38,081,995 | | *ORMDL3* | 8.024 | 5.25E-13 | 2.01E-04 |
| | | 38,082,018 | | *ORMDL3* | 5.178 | 8.31E-07 | 6.40E-04 |
| | | 38,082,022 | | *ORMDL3* | 5.750 | 6.06E-08 | 6.40E-04 |
| | | 38,082,035 | | *ORMDL3* | 5.054 | 1.44E-06 | 6.40E-04 |
| | | 38,082,206 | | *ORMDL3* | 4.463 | 1.73E-05 | 0.004 |
| | | 38,082,235 | | *ORMDL3* | 4.806 | 4.18E-06 | 0.004 |
| | | 38,082,346 | | *ORMDL3* | 5.646 | 9.85E-08 | 0.011 |
| | | 38,082,448 | | *ORMDL3* | 5.015 | 1.70E-06 | 0.011 |
| | | 38,082,658 | | *ORMDL3* | 4.641 | 8.36E-06 | 0.011 |
| | | 38,082,921 | | *ORMDL3* | 3.473 | 7.00E-04 | 0.011 |
| rs907091 | g.37921742.C>T | 37,828,296 | | *PGAP3* | 3.803 | 2.19E-04 | 0.002 |
| | | 37,916,005 | | | 3.601 | 4.50E-04 | 0.001 |
| | | 37,927,002 | | *IKZF3* | -3.436 | 7.92E-04 | 0.001 |
| | | 38,018,804 | | *IKZF3* | 3.992 | 1.09E-04 | 0.046 |
| | | 38,049,272 | | | 3.476 | 6.93E-04 | 0.046 |

| | | | | | | | |
|---|---|---|---|---|---|---|---|
| | | 38,077,191 | | | 3.984 | 1.12E-04 | 0.034 |
| | | 38,080,829 | | *ORMDL3* | 5.174 | 8.46E-07 | 0.014 |
| | | 38,081,995 | | *ORMDL3* | 4.572 | 1.11E-05 | 2.01E-04 |
| | | 38,082,022 | | *ORMDL3* | 4.714 | 6.16E-06 | 6.40E-04 |
| | | 38,082,035 | | *ORMDL3* | 5.590 | 1.28E-07 | 6.40E-04 |
| | | 38,082,073 | | *ORMDL3* | 5.398 | 3.08E-07 | 6.40E-04 |
| | | 38,082,346 | | *ORMDL3* | 4.036 | 9.23E-05 | 0.011 |
| | | 38,082,448 | | *ORMDL3* | 3.945 | 1.30E-04 | 0.011 |
| | | 38,082,658 | | *ORMDL3* | 3.982 | 1.13E-04 | 0.011 |
| | | 38,082,778 | | *ORMDL3* | 3.596 | 4.57E-04 | 0.011 |
| | | 38,108,710 | | | -3.733 | 2.82E-04 | 0.039 |
| | | 38,119,198 | *GSDMA* | | 3.852 | 1.83E-04 | 0.009 |
| | | 38,119,207 | *GSDMA* | | 5.413 | 2.89E-07 | 0.009 |
| | | 38,119,649 | | *GSDMA* | 3.900 | 1.54E-04 | 0.002 |
| rs907092 | g.37922259.A>G (Ser.438.Ser) | 37,828,296 | | *PGAP3* | 4.245 | 4.12E-05 | 0.002 |
| | | 37,927,002 | | *IKZF3* | -3.408 | 8.72E-04 | 0.001 |
| | | 38,024,331 | | | -3.380 | 9.59E-04 | 0.017 |
| | | 38,072,514 | | *GSDMB* | 3.674 | 3.48E-04 | 0.046 |
| | | 38,077,191 | | | 4.240 | 4.21E-05 | 0.034 |
| | | 38,080,829 | | *ORMDL3* | 3.389 | 9.29E-04 | 0.014 |
| | | 38,081,829 | | *ORMDL3* | 3.410 | 8.67E-04 | 2.01E-04 |
| | | 38,081,995 | | *ORMDL3* | 5.953 | 2.30E-08 | 2.01E-04 |
| | | 38,082,018 | | *ORMDL3* | 3.785 | 2.34E-04 | 6.40E-04 |
| | | 38,082,022 | | *ORMDL3* | 6.359 | 3.16E-09 | 6.40E-04 |
| | | 38,082,035 | | *ORMDL3* | 6.335 | 3.56E-09 | 6.40E-04 |
| | | 38,082,073 | | *ORMDL3* | 6.301 | 4.22E-09 | 6.40E-04 |
| | | 38,082,206 | | *ORMDL3* | 3.805 | 2.18E-04 | 0.004 |
| | | 38,082,235 | | *ORMDL3* | 3.678 | 3.43E-04 | 0.004 |
| | | 38,082,346 | | *ORMDL3* | 4.166 | 5.61E-05 | 0.011 |
| | | 38,082,448 | | *ORMDL3* | 3.502 | 6.33E-04 | 0.011 |
| | | 38,082,658 | | *ORMDL3* | 3.961 | 1.23E-04 | 0.011 |
| | | 38,082,778 | | *ORMDL3* | 4.055 | 8.57E-05 | 0.011 |
| | | 38,108,710 | | | -4.323 | 3.04E-05 | 0.039 |
| | | 38,119,198 | *GSDMA* | | 3.982 | 1.13E-04 | 0.009 |
| | | 38,119,207 | *GSDMA* | | 5.131 | 1.03E-06 | 0.009 |
| | | 38,119,649 | | *GSDMA* | 4.151 | 5.95E-05 | 0.002 |
| rs2952140 | g.37928059.G>A | 37,828,296 | | *PGAP3* | 3.778 | 2.40E-04 | 0.002 |
| | | 37,927,002 | | *IKZF3* | -3.460 | 7.31E-04 | 0.001 |
| | | 38,018,804 | | *IKZF3* | 4.320 | 3.07E-05 | 0.046 |
| | | 38,049,272 | | | 3.402 | 8.89E-04 | 0.046 |

| | | | | | | | |
|---|---|---|---|---|---|---|---|
| | | 38,077,191 | | | 4.230 | 4.38E-05 | 0.034 |
| | | 38,080,829 | | *ORMDL3* | 5.263 | 5.69E-07 | 0.014 |
| | | 38,081,492 | | *ORMDL3* | 3.374 | 9.75E-04 | 2.01E-04 |
| | | 38,081,812 | | *ORMDL3* | 3.527 | 5.81E-04 | 2.01E-04 |
| | | 38,081,995 | | *ORMDL3* | 4.777 | 4.73E-06 | 2.01E-04 |
| | | 38,082,022 | | *ORMDL3* | 5.131 | 1.02E-06 | 6.40E-04 |
| | | 38,082,035 | | *ORMDL3* | 6.518 | 1.43E-09 | 6.40E-04 |
| | | 38,082,073 | | *ORMDL3* | 5.615 | 1.14E-07 | 6.40E-04 |
| | | 38,082,235 | | *ORMDL3* | 3.719 | 2.96E-04 | 0.004 |
| | | 38,082,346 | | *ORMDL3* | 4.935 | 2.40E-06 | 0.011 |
| | | 38,082,448 | | *ORMDL3* | 4.302 | 3.30E-05 | 0.011 |
| | | 38,082,658 | | *ORMDL3* | 4.456 | 1.78E-05 | 0.011 |
| | | 38,082,778 | | *ORMDL3* | 3.680 | 3.41E-04 | 0.011 |
| | | 38,108,710 | | | -3.858 | 1.79E-04 | 0.039 |
| | | 38,119,198 | *GSDMA* | | 4.235 | 4.30E-05 | 0.009 |
| | | 38,119,207 | *GSDMA* | | 5.681 | 8.39E-08 | 0.009 |
| | | 38,119,649 | | *GSDMA* | 4.187 | 5.18E-05 | 0.002 |
| rs2313430 | g.37929816.A>G | 37,828,296 | | *PGAP3* | 3.713 | 3.03E-04 | 0.002 |
| | | 37,927,002 | | *IKZF3* | -3.431 | 8.08E-04 | 0.001 |
| | | 38,018,804 | | *IKZF3* | 4.364 | 2.58E-05 | 0.046 |
| | | 38,049,272 | | | 3.372 | 9.82E-04 | 0.046 |
| | | 38,077,191 | | | 4.240 | 4.21E-05 | 0.034 |
| | | 38,080,829 | | *ORMDL3* | 5.181 | 8.21E-07 | 0.014 |
| | | 38,081,492 | | *ORMDL3* | 3.390 | 9.27E-04 | 2.01E-04 |
| | | 38,081,812 | | *ORMDL3* | 3.541 | 5.53E-04 | 2.01E-04 |
| | | 38,081,995 | | *ORMDL3* | 4.771 | 4.84E-06 | 2.01E-04 |
| | | 38,082,022 | | *ORMDL3* | 5.151 | 9.38E-07 | 6.40E-04 |
| | | 38,082,035 | | *ORMDL3* | 6.590 | 9.97E-10 | 6.40E-04 |
| | | 38,082,073 | | *ORMDL3* | 5.656 | 9.39E-08 | 6.40E-04 |
| | | 38,082,235 | | *ORMDL3* | 3.750 | 2.65E-04 | 0.004 |
| | | 38,082,346 | | *ORMDL3* | 4.945 | 2.31E-06 | 0.011 |
| | | 38,082,448 | | *ORMDL3* | 4.327 | 2.99E-05 | 0.011 |
| | | 38,082,658 | | *ORMDL3* | 4.490 | 1.55E-05 | 0.011 |
| | | 38,082,778 | | *ORMDL3* | 3.691 | 3.27E-04 | 0.011 |
| | | 38,108,710 | | | -3.906 | 1.50E-04 | 0.039 |
| | | 38,119,198 | *GSDMA* | | 4.245 | 4.12E-05 | 0.009 |
| | | 38,119,207 | *GSDMA* | | 5.687 | 8.13E-08 | 0.009 |
| | | 38,119,649 | | *GSDMA* | 4.237 | 4.25E-05 | 0.002 |
| rs10445308 | g.37938047.T>C | 37,828,296 | | *PGAP3* | 4.110 | 6.95E-05 | 0.002 |
| | | 38,018,804 | | *IKZF3* | 3.691 | 3.27E-04 | 0.046 |
| | | 38,072,514 | | *GSDMB* | 3.478 | 6.88E-04 | 0.046 |

| | | | | | | | |
|---|---|---|---|---|---|---|---|
| | | 38,077,191 | | | 4.469 | 1.69E-05 | 0.034 |
| | | 38,081,812 | | *ORMDL3* | 3.613 | 4.31E-04 | 2.01E-04 |
| | | 38,081,829 | | *ORMDL3* | 3.789 | 2.30E-04 | 2.01E-04 |
| | | 38,081,995 | | *ORMDL3* | 6.147 | 9.00E-09 | 2.01E-04 |
| | | 38,082,018 | | *ORMDL3* | 3.887 | 1.61E-04 | 6.40E-04 |
| | | 38,082,022 | | *ORMDL3* | 6.811 | 3.25E-10 | 6.40E-04 |
| | | 38,082,035 | | *ORMDL3* | 7.272 | 2.96E-11 | 6.40E-04 |
| | | 38,082,073 | | *ORMDL3* | 6.558 | 1.17E-09 | 6.40E-04 |
| | | 38,082,206 | | *ORMDL3* | 3.980 | 1.14E-04 | 0.004 |
| | | 38,082,235 | | *ORMDL3* | 4.048 | 8.83E-05 | 0.004 |
| | | 38,082,346 | | *ORMDL3* | 4.893 | 2.88E-06 | 0.011 |
| | | 38,082,448 | | *ORMDL3* | 3.824 | 2.03E-04 | 0.011 |
| | | 38,082,658 | | *ORMDL3* | 4.418 | 2.08E-05 | 0.011 |
| | | 38,082,778 | | *ORMDL3* | 4.155 | 5.85E-05 | 0.011 |
| | | 38,108,710 | | | -4.508 | 1.44E-05 | 0.039 |
| | | 38,119,198 | *GSDMA* | | 4.288 | 3.49E-05 | 0.009 |
| | | 38,119,207 | *GSDMA* | | 5.321 | 4.38E-07 | 0.009 |
| | | 38,119,649 | | *GSDMA* | 4.492 | 1.54E-05 | 0.002 |
| rs12942330 | g.37939839.T>C | 37,828,296 | | *PGAP3* | 4.110 | 6.95E-05 | 0.002 |
| | | 38,018,804 | | *IKZF3* | 3.691 | 3.27E-04 | 0.046 |
| | | 38,072,514 | | *GSDMB* | 3.478 | 6.88E-04 | 0.046 |
| | | 38,077,191 | | | 4.469 | 1.69E-05 | 0.034 |
| | | 38,081,812 | | *ORMDL3* | 3.613 | 4.31E-04 | 2.01E-04 |
| | | 38,081,829 | | *ORMDL3* | 3.789 | 2.30E-04 | 2.01E-04 |
| | | 38,081,995 | | *ORMDL3* | 6.147 | 9.00E-09 | 2.01E-04 |
| | | 38,082,018 | | *ORMDL3* | 3.887 | 1.61E-04 | 6.40E-04 |
| | | 38,082,022 | | *ORMDL3* | 6.811 | 3.25E-10 | 6.40E-04 |
| | | 38,082,035 | | *ORMDL3* | 7.272 | 2.96E-11 | 6.40E-04 |
| | | 38,082,073 | | *ORMDL3* | 6.558 | 1.17E-09 | 6.40E-04 |
| | | 38,082,206 | | *ORMDL3* | 3.980 | 1.14E-04 | 0.004 |
| | | 38,082,235 | | *ORMDL3* | 4.048 | 8.83E-05 | 0.004 |
| | | 38,082,346 | | *ORMDL3* | 4.893 | 2.88E-06 | 0.011 |
| | | 38,082,448 | | *ORMDL3* | 3.824 | 2.03E-04 | 0.011 |
| | | 38,082,658 | | *ORMDL3* | 4.418 | 2.08E-05 | 0.011 |
| | | 38,082,778 | | *ORMDL3* | 4.155 | 5.85E-05 | 0.011 |
| | | 38,108,710 | | | -4.508 | 1.44E-05 | 0.039 |
| | | 38,119,198 | *GSDMA* | | 4.288 | 3.49E-05 | 0.009 |
| | | 38,119,207 | *GSDMA* | | 5.321 | 4.38E-07 | 0.009 |
| | | 38,119,649 | | *GSDMA* | 4.492 | 1.54E-05 | 0.002 |
| rs11658993 | g.37940808.T>C | 37,828,296 | | *PGAP3* | 4.110 | 6.95E-05 | 0.002 |
| | | 38,018,804 | | *IKZF3* | 3.691 | 3.27E-04 | 0.046 |

| | | | | | | | |
|---|---|---|---|---|---|---|---|
| | | 38,072,514 | | *GSDMB* | 3.478 | 6.88E-04 | 0.046 |
| | | 38,077,191 | | | 4.469 | 1.69E-05 | 0.034 |
| | | 38,081,812 | | *ORMDL3* | 3.613 | 4.31E-04 | 2.01E-04 |
| | | 38,081,829 | | *ORMDL3* | 3.789 | 2.30E-04 | 2.01E-04 |
| | | 38,081,995 | | *ORMDL3* | 6.147 | 9.00E-09 | 2.01E-04 |
| | | 38,082,018 | | *ORMDL3* | 3.887 | 1.61E-04 | 6.40E-04 |
| | | 38,082,022 | | *ORMDL3* | 6.811 | 3.25E-10 | 6.40E-04 |
| | | 38,082,035 | | *ORMDL3* | 7.272 | 2.96E-11 | 6.40E-04 |
| | | 38,082,073 | | *ORMDL3* | 6.558 | 1.17E-09 | 6.40E-04 |
| | | 38,082,206 | | *ORMDL3* | 3.980 | 1.14E-04 | 0.004 |
| | | 38,082,235 | | *ORMDL3* | 4.048 | 8.83E-05 | 0.004 |
| | | 38,082,346 | | *ORMDL3* | 4.893 | 2.88E-06 | 0.011 |
| | | 38,082,448 | | *ORMDL3* | 3.824 | 2.03E-04 | 0.011 |
| | | 38,082,658 | | *ORMDL3* | 4.418 | 2.08E-05 | 0.011 |
| | | 38,082,778 | | *ORMDL3* | 4.155 | 5.85E-05 | 0.011 |
| | | 38,108,710 | | | -4.508 | 1.44E-05 | 0.039 |
| | | 38,119,198 | *GSDMA* | | 4.288 | 3.49E-05 | 0.009 |
| | | 38,119,207 | *GSDMA* | | 5.321 | 4.38E-07 | 0.009 |
| | | 38,119,649 | | *GSDMA* | 4.492 | 1.54E-05 | 0.002 |
| rs17676191 | g.37949924.G>A | 37,828,195 | | *PGAP3* | -4.004 | 1.04E-04 | |
| | | 37,895,953 | | | -3.395 | 9.12E-04 | 0.027 |
| | | 38,077,464 | | *ORMDL3* | 4.624 | 8.98E-06 | 0.014 |
| | | 38,084,012 | *ORMDL3* | | 3.610 | 4.35E-04 | 0.011 |
| rs35395438 | g.37952891.G>T | 37,863,595 | | *ERBB2* | -3.625 | 4.13E-04 | 0.002 |
| | | 37,893,689 | | | 4.186 | 5.20E-05 | 0.002 |
| | | 38,025,241 | | | -4.081 | 7.79E-05 | 0.046 |
| | | 38,065,264 | | *GSDMB* | 4.640 | 8.40E-06 | 0.046 |
| | | 38,081,829 | | *ORMDL3* | 3.724 | 2.91E-04 | 2.01E-04 |
| | | 38,081,995 | | *ORMDL3* | 6.045 | 1.48E-08 | 2.01E-04 |
| | | 38,082,018 | | *ORMDL3* | 3.649 | 3.80E-04 | 6.40E-04 |
| | | 38,082,022 | | *ORMDL3* | 4.585 | 1.05E-05 | 6.40E-04 |
| | | 38,082,035 | | *ORMDL3* | 4.229 | 4.39E-05 | 6.40E-04 |
| | | 38,082,235 | | *ORMDL3* | 3.461 | 7.30E-04 | 0.004 |
| | | 38,082,346 | | *ORMDL3* | 4.867 | 3.22E-06 | 0.011 |
| | | 38,082,448 | | *ORMDL3* | 3.542 | 5.53E-04 | 0.011 |
| | | 38,082,658 | | *ORMDL3* | 4.517 | 1.39E-05 | 0.011 |
| | | 38,109,763 | | | -3.701 | 3.16E-04 | 0.002 |
| rs112350333 | g.37952989.T>C | 37,893,689 | | | 3.635 | 3.99E-04 | 0.002 |
| | | 37,912,051 | | | 3.489 | 6.63E-04 | 0.033 |
| | | 38,020,208 | | *IKZF3* | 3.573 | 4.97E-04 | 0.046 |
| | | 38,025,241 | | | -4.068 | 8.18E-05 | 0.046 |

| | | | | | | | |
|---|---|---|---|---|---|---|---|
| | | 38,065,264 | | *GSDMB* | 4.738 | 5.57E-06 | 0.046 |
| | | 38,081,812 | | *ORMDL3* | 3.417 | 8.46E-04 | 2.01E-04 |
| | | 38,081,829 | | *ORMDL3* | 4.479 | 1.63E-05 | 2.01E-04 |
| | | 38,081,995 | | *ORMDL3* | 7.259 | 3.17E-11 | 2.01E-04 |
| | | 38,082,018 | | *ORMDL3* | 4.543 | 1.25E-05 | 6.40E-04 |
| | | 38,082,022 | | *ORMDL3* | 5.453 | 2.41E-07 | 6.40E-04 |
| | | 38,082,035 | | *ORMDL3* | 5.220 | 6.89E-07 | 6.40E-04 |
| | | 38,082,206 | | *ORMDL3* | 3.882 | 1.64E-04 | 0.004 |
| | | 38,082,235 | | *ORMDL3* | 4.264 | 3.83E-05 | 0.004 |
| | | 38,082,346 | | *ORMDL3* | 5.707 | 7.39E-08 | 0.011 |
| | | 38,082,448 | | *ORMDL3* | 4.498 | 1.50E-05 | 0.011 |
| | | 38,082,658 | | *ORMDL3* | 4.804 | 4.22E-06 | 0.011 |
| rs2952144 | g.37960017.C>T | 37,828,296 | | *PGAP3* | 3.713 | 3.03E-04 | 0.002 |
| | | 37,927,002 | | *IKZF3* | -3.431 | 8.08E-04 | 0.001 |
| | | 38,018,804 | | *IKZF3* | 4.364 | 2.58E-05 | 0.046 |
| | | 38,049,272 | | | 3.372 | 9.82E-04 | 0.046 |
| | | 38,077,191 | | | 4.240 | 4.21E-05 | 0.034 |
| | | 38,080,829 | | *ORMDL3* | 5.181 | 8.21E-07 | 0.014 |
| | | 38,081,492 | | *ORMDL3* | 3.390 | 9.27E-04 | 2.01E-04 |
| | | 38,081,812 | | *ORMDL3* | 3.541 | 5.53E-04 | 2.01E-04 |
| | | 38,081,995 | | *ORMDL3* | 4.771 | 4.84E-06 | 2.01E-04 |
| | | 38,082,022 | | *ORMDL3* | 5.151 | 9.38E-07 | 6.40E-04 |
| | | 38,082,035 | | *ORMDL3* | 6.590 | 9.97E-10 | 6.40E-04 |
| | | 38,082,073 | | *ORMDL3* | 5.656 | 9.39E-08 | 6.40E-04 |
| | | 38,082,235 | | *ORMDL3* | 3.750 | 2.65E-04 | 0.004 |
| | | 38,082,346 | | *ORMDL3* | 4.945 | 2.31E-06 | 0.011 |
| | | 38,082,448 | | *ORMDL3* | 4.327 | 2.99E-05 | 0.011 |
| | | 38,082,658 | | *ORMDL3* | 4.490 | 1.55E-05 | 0.011 |
| | | 38,082,778 | | *ORMDL3* | 3.691 | 3.27E-04 | 0.011 |
| | | 38,108,710 | | | -3.906 | 1.50E-04 | 0.039 |
| | | 38,119,198 | *GSDMA* | | 4.245 | 4.12E-05 | 0.009 |
| | | 38,119,207 | *GSDMA* | | 5.687 | 8.13E-08 | 0.009 |
| | | 38,119,649 | | *GSDMA* | 4.237 | 4.25E-05 | 0.002 |
| rs4795395 | g.37962987.A>T | 37,828,296 | | *PGAP3* | 4.110 | 6.95E-05 | 0.002 |
| | | 38,018,804 | | *IKZF3* | 3.691 | 3.27E-04 | 0.046 |
| | | 38,072,514 | | *GSDMB* | 3.478 | 6.88E-04 | 0.046 |
| | | 38,077,191 | | | 4.469 | 1.69E-05 | 0.034 |
| | | 38,081,812 | | *ORMDL3* | 3.613 | 4.31E-04 | 2.01E-04 |
| | | 38,081,829 | | *ORMDL3* | 3.789 | 2.30E-04 | 2.01E-04 |
| | | 38,081,995 | | *ORMDL3* | 6.147 | 9.00E-09 | 2.01E-04 |
| | | 38,082,018 | | *ORMDL3* | 3.887 | 1.61E-04 | 6.40E-04 |

| | | | | | | | | |
|---|---|---|---|---|---|---|---|---|
| | | | 38,082,022 | | *ORMDL3* | 6.811 | 3.25E-10 | 6.40E-04 |
| | | | 38,082,035 | | *ORMDL3* | 7.272 | 2.96E-11 | 6.40E-04 |
| | | | 38,082,073 | | *ORMDL3* | 6.558 | 1.17E-09 | 6.40E-04 |
| | | | 38,082,206 | | *ORMDL3* | 3.980 | 1.14E-04 | 0.004 |
| | | | 38,082,235 | | *ORMDL3* | 4.048 | 8.83E-05 | 0.004 |
| | | | 38,082,346 | | *ORMDL3* | 4.893 | 2.88E-06 | 0.011 |
| | | | 38,082,448 | | *ORMDL3* | 3.824 | 2.03E-04 | 0.011 |
| | | | 38,082,658 | | *ORMDL3* | 4.418 | 2.08E-05 | 0.011 |
| | | | 38,082,778 | | *ORMDL3* | 4.155 | 5.85E-05 | 0.011 |
| | | | 38,108,710 | | | -4.508 | 1.44E-05 | 0.039 |
| | | | 38,119,198 | *GSDMA* | | 4.288 | 3.49E-05 | 0.009 |
| | | | 38,119,207 | *GSDMA* | | 5.321 | 4.38E-07 | 0.009 |
| | | | 38,119,649 | | *GSDMA* | 4.492 | 1.54E-05 | 0.002 |
| | rs9909593 | g.37970149.G>A | 37,828,296 | | *PGAP3* | 4.110 | 6.95E-05 | 0.002 |
| | | | 38,018,804 | | *IKZF3* | 3.691 | 3.27E-04 | 0.046 |
| | | | 38,072,514 | | *GSDMB* | 3.478 | 6.88E-04 | 0.046 |
| | | | 38,077,191 | | | 4.469 | 1.69E-05 | 0.034 |
| | | | 38,081,812 | | *ORMDL3* | 3.613 | 4.31E-04 | 2.01E-04 |
| | | | 38,081,829 | | *ORMDL3* | 3.789 | 2.30E-04 | 2.01E-04 |
| | | | 38,081,995 | | *ORMDL3* | 6.147 | 9.00E-09 | 2.01E-04 |
| | | | 38,082,018 | | *ORMDL3* | 3.887 | 1.61E-04 | 6.40E-04 |
| | | | 38,082,022 | | *ORMDL3* | 6.811 | 3.25E-10 | 6.40E-04 |
| | | | 38,082,035 | | *ORMDL3* | 7.272 | 2.96E-11 | 6.40E-04 |
| | | | 38,082,073 | | *ORMDL3* | 6.558 | 1.17E-09 | 6.40E-04 |
| | | | 38,082,206 | | *ORMDL3* | 3.980 | 1.14E-04 | 0.004 |
| | | | 38,082,235 | | *ORMDL3* | 4.048 | 8.83E-05 | 0.004 |
| | | | 38,082,346 | | *ORMDL3* | 4.893 | 2.88E-06 | 0.011 |
| | | | 38,082,448 | | *ORMDL3* | 3.824 | 2.03E-04 | 0.011 |
| | | | 38,082,658 | | *ORMDL3* | 4.418 | 2.08E-05 | 0.011 |
| | | | 38,082,778 | | *ORMDL3* | 4.155 | 5.85E-05 | 0.011 |
| | | | 38,108,710 | | | -4.508 | 1.44E-05 | 0.039 |
| | | | 38,119,198 | *GSDMA* | | 4.288 | 3.49E-05 | 0.009 |
| | | | 38,119,207 | *GSDMA* | | 5.321 | 4.38E-07 | 0.009 |
| | | | 38,119,649 | | *GSDMA* | 4.492 | 1.54E-05 | 0.002 |
| Fourth block | rs12450323 | g.37972708.T>G | 37,828,296 | | *PGAP3* | -4.127 | 6.53E-05 | 0.002 |
| | | | 37,828,560 | | *PGAP3* | -4.049 | 8.77E-05 | 0.002 |
| | | | 37,856,551 | | *ERBB2* | 3.378 | 9.64E-04 | 0.006 |
| | | | 38,082,035 | | *ORMDL3* | -3.693 | 3.25E-04 | 6.40E-04 |
| | | | 38,082,073 | | *ORMDL3* | -4.654 | 7.93E-06 | 6.40E-04 |
| | rs113897057 | g.37975214_37975214ins TTCTA | 37,828,296 | | *PGAP3* | 3.648 | 3.81E-04 | 0.002 |

| | | | | | | | |
|---|---|---|---|---|---|---|---|
| | | 37,927,002 | | *IKZF3* | -3.592 | 4.65E-04 | 0.001 |
| | | 38,018,804 | | *IKZF3* | 4.401 | 2.23E-05 | 0.046 |
| | | 38,049,272 | | | 3.480 | 6.82E-04 | 0.046 |
| | | 38,077,191 | | | 4.095 | 7.37E-05 | 0.034 |
| | | 38,080,829 | | *ORMDL3* | 5.101 | 1.17E-06 | 0.014 |
| | | 38,081,492 | | *ORMDL3* | 3.611 | 4.35E-04 | 2.01E-04 |
| | | 38,081,995 | | *ORMDL3* | 4.772 | 4.84E-06 | 2.01E-04 |
| | | 38,082,022 | | *ORMDL3* | 5.205 | 7.37E-07 | 6.40E-04 |
| | | 38,082,035 | | *ORMDL3* | 6.863 | 2.48E-10 | 6.40E-04 |
| | | 38,082,073 | | *ORMDL3* | 5.646 | 9.86E-08 | 6.40E-04 |
| | | 38,082,235 | | *ORMDL3* | 3.663 | 3.62E-04 | 0.004 |
| | | 38,082,346 | | *ORMDL3* | 5.288 | 5.08E-07 | 0.011 |
| | | 38,082,448 | | *ORMDL3* | 4.305 | 3.26E-05 | 0.011 |
| | | 38,082,658 | | *ORMDL3* | 4.478 | 1.63E-05 | 0.011 |
| | | 38,082,778 | | *ORMDL3* | 3.651 | 3.78E-04 | 0.011 |
| | | 38,108,710 | | | -3.889 | 1.60E-04 | 0.039 |
| | | 38,119,198 | *GSDMA* | | 4.466 | 1.71E-05 | 0.009 |
| | | 38,119,207 | *GSDMA* | | 5.826 | 4.22E-08 | 0.009 |
| | | 38,119,649 | | *GSDMA* | 4.232 | 4.35E-05 | 0.002 |
| rs9303277 | g.37976469.T>C | 37,828,296 | | *PGAP3* | 3.713 | 3.03E-04 | 0.002 |
| | | 37,927,002 | | *IKZF3* | -3.431 | 8.08E-04 | 0.001 |
| | | 38,018,804 | | *IKZF3* | 4.364 | 2.58E-05 | 0.046 |
| | | 38,049,272 | | | 3.372 | 9.82E-04 | 0.046 |
| | | 38,077,191 | | | 4.240 | 4.21E-05 | 0.034 |
| | | 38,080,829 | | *ORMDL3* | 5.181 | 8.21E-07 | 0.014 |
| | | 38,081,492 | | *ORMDL3* | 3.390 | 9.27E-04 | 2.01E-04 |
| | | 38,081,812 | | *ORMDL3* | 3.541 | 5.53E-04 | 2.01E-04 |
| | | 38,081,995 | | *ORMDL3* | 4.771 | 4.84E-06 | 2.01E-04 |
| | | 38,082,022 | | *ORMDL3* | 5.151 | 9.38E-07 | 6.40E-04 |
| | | 38,082,035 | | *ORMDL3* | 6.590 | 9.97E-10 | 6.40E-04 |
| | | 38,082,073 | | *ORMDL3* | 5.656 | 9.39E-08 | 6.40E-04 |
| | | 38,082,235 | | *ORMDL3* | 3.750 | 2.65E-04 | 0.004 |
| | | 38,082,346 | | *ORMDL3* | 4.945 | 2.31E-06 | 0.011 |
| | | 38,082,448 | | *ORMDL3* | 4.327 | 2.99E-05 | 0.011 |
| | | 38,082,658 | | *ORMDL3* | 4.490 | 1.55E-05 | 0.011 |
| | | 38,082,778 | | *ORMDL3* | 3.691 | 3.27E-04 | 0.011 |
| | | 38,108,710 | | | -3.906 | 1.50E-04 | 0.039 |
| | | 38,119,198 | *GSDMA* | | 4.245 | 4.12E-05 | 0.009 |
| | | 38,119,207 | *GSDMA* | | 5.687 | 8.13E-08 | 0.009 |
| | | 38,119,649 | | *GSDMA* | 4.237 | 4.25E-05 | 0.002 |
| rs2060941 | g.37982883.T>G | 37,911,152 | | | 3.404 | 8.83E-04 | 0.035 |

| | | | | | | | |
|---|---|---|---|---|---|---|---|
| | | 38,024,328 | | | 3.810 | 2.13E-04 | 0.023 |
| | | 38,081,812 | | *ORMDL3* | 3.779 | 2.39E-04 | 2.01E-04 |
| | | 38,081,829 | | *ORMDL3* | 3.526 | 5.84E-04 | 2.01E-04 |
| | | 38,081,995 | | *ORMDL3* | 4.587 | 1.04E-05 | 2.01E-04 |
| | | 38,082,018 | | *ORMDL3* | 3.834 | 1.96E-04 | 6.40E-04 |
| | | 38,082,035 | | *ORMDL3* | 3.585 | 4.76E-04 | 6.40E-04 |
| | | 38,082,346 | | *ORMDL3* | 4.645 | 8.23E-06 | 0.011 |
| | | 38,083,537 | | *ORMDL3* | 3.912 | 1.47E-04 | 0.011 |
| rs12944882 | g.37983492.C>T | 37,828,296 | | *PGAP3* | 3.381 | 9.53E-04 | 0.002 |
| | | 37,927,002 | | *IKZF3* | -3.507 | 6.24E-04 | 0.001 |
| | | 38,018,804 | | *IKZF3* | 3.950 | 1.28E-04 | 0.046 |
| | | 38,077,191 | | | 4.307 | 3.24E-05 | 0.034 |
| | | 38,080,829 | | *ORMDL3* | 4.195 | 5.02E-05 | 0.014 |
| | | 38,081,812 | | *ORMDL3* | 3.968 | 1.19E-04 | 2.01E-04 |
| | | 38,081,995 | | *ORMDL3* | 4.443 | 1.88E-05 | 2.01E-04 |
| | | 38,082,022 | | *ORMDL3* | 4.750 | 5.30E-06 | 6.40E-04 |
| | | 38,082,035 | | *ORMDL3* | 6.058 | 1.39E-08 | 6.40E-04 |
| | | 38,082,073 | | *ORMDL3* | 6.009 | 1.76E-08 | 6.40E-04 |
| | | 38,082,235 | | *ORMDL3* | 3.735 | 2.80E-04 | 0.004 |
| | | 38,082,346 | | *ORMDL3* | 5.482 | 2.11E-07 | 0.011 |
| | | 38,082,448 | | *ORMDL3* | 3.997 | 1.07E-04 | 0.011 |
| | | 38,082,658 | | *ORMDL3* | 3.607 | 4.40E-04 | 0.011 |
| | | 38,082,778 | | *ORMDL3* | 3.529 | 5.78E-04 | 0.011 |
| | | 38,119,207 | *GSDMA* | | 4.644 | 8.26E-06 | 0.009 |
| | | 38,119,649 | | *GSDMA* | 4.299 | 3.34E-05 | 0.002 |
| rs3816470 | g.37985801.C>T | 37,828,296 | | *PGAP3* | 3.381 | 9.53E-04 | 0.002 |
| | | 37,927,002 | | *IKZF3* | -3.507 | 6.24E-04 | 0.001 |
| | | 38,018,804 | | *IKZF3* | 3.950 | 1.28E-04 | 0.046 |
| | | 38,077,191 | | | 4.307 | 3.24E-05 | 0.034 |
| | | 38,080,829 | | *ORMDL3* | 4.195 | 5.02E-05 | 0.014 |
| | | 38,081,812 | | *ORMDL3* | 3.968 | 1.19E-04 | 2.01E-04 |
| | | 38,081,995 | | *ORMDL3* | 4.443 | 1.88E-05 | 2.01E-04 |
| | | 38,082,022 | | *ORMDL3* | 4.750 | 5.30E-06 | 6.40E-04 |
| | | 38,082,035 | | *ORMDL3* | 6.058 | 1.39E-08 | 6.40E-04 |
| | | 38,082,073 | | *ORMDL3* | 6.009 | 1.76E-08 | 6.40E-04 |
| | | 38,082,235 | | *ORMDL3* | 3.735 | 2.80E-04 | 0.004 |
| | | 38,082,346 | | *ORMDL3* | 5.482 | 2.11E-07 | 0.011 |
| | | 38,082,448 | | *ORMDL3* | 3.997 | 1.07E-04 | 0.011 |
| | | 38,082,658 | | *ORMDL3* | 3.607 | 4.40E-04 | 0.011 |
| | | 38,082,778 | | *ORMDL3* | 3.529 | 5.78E-04 | 0.011 |
| | | 38,119,207 | *GSDMA* | | 4.644 | 8.26E-06 | 0.009 |

| | | | | | | | |
|---|---|---|---|---|---|---|---|
| | | 38,119,649 | | *GSDMA* | 4.299 | 3.34E-05 | 0.002 |
| rs62066988 | g.37992281.T>C | 38,018,804 | | *IKZF3* | 3.455 | 7.44E-04 | 0.046 |
| | | 38,024,331 | | | -3.954 | 1.26E-04 | 0.017 |
| | | 38,024,353 | | | -3.385 | 9.43E-04 | 0.017 |
| | | 38,072,514 | | *GSDMB* | 3.931 | 1.37E-04 | 0.046 |
| | | 38,082,035 | | *ORMDL3* | 3.670 | 3.52E-04 | 6.40E-04 |
| | | 38,082,073 | | *ORMDL3* | 4.454 | 1.80E-05 | 6.40E-04 |
| | | 38,108,710 | | | -3.868 | 1.73E-04 | 0.039 |
| | | 38,119,198 | *GSDMA* | | 4.130 | 6.44E-05 | 0.009 |
| | | 38,119,207 | *GSDMA* | | 4.557 | 1.18E-05 | 0.009 |
| | | 38,119,649 | | *GSDMA* | 4.330 | 2.95E-05 | 0.002 |
| rs34233420 | g.38004929_38004933del | 37,828,296 | | *PGAP3* | 4.004 | 1.04E-04 | 0.002 |
| | | 37,887,061 | *MIEN1* | | 3.395 | 9.09E-04 | 0.002 |
| | | 38,018,804 | | *IKZF3* | 3.393 | 9.17E-04 | 0.046 |
| | | 38,077,098 | | | 3.419 | 8.39E-04 | 0.034 |
| | | 38,077,191 | | | 4.160 | 5.75E-05 | 0.034 |
| | | 38,080,829 | | *ORMDL3* | 3.540 | 5.56E-04 | 0.014 |
| | | 38,081,812 | | *ORMDL3* | 3.845 | 1.88E-04 | 2.01E-04 |
| | | 38,081,829 | | *ORMDL3* | 3.797 | 2.23E-04 | 2.01E-04 |
| | | 38,081,995 | | *ORMDL3* | 6.022 | 1.65E-08 | 2.01E-04 |
| | | 38,082,022 | | *ORMDL3* | 6.106 | 1.10E-08 | 6.40E-04 |
| | | 38,082,035 | | *ORMDL3* | 6.810 | 3.26E-10 | 6.40E-04 |
| | | 38,082,073 | | *ORMDL3* | 6.701 | 5.68E-10 | 6.40E-04 |
| | | 38,082,206 | | *ORMDL3* | 3.828 | 2.00E-04 | 0.004 |
| | | 38,082,235 | | *ORMDL3* | 3.796 | 2.25E-04 | 0.004 |
| | | 38,082,346 | | *ORMDL3* | 4.413 | 2.12E-05 | 0.011 |
| | | 38,082,448 | | *ORMDL3* | 3.633 | 4.02E-04 | 0.011 |
| | | 38,082,658 | | *ORMDL3* | 4.452 | 1.81E-05 | 0.011 |
| | | 38,082,778 | | *ORMDL3* | 4.051 | 8.71E-05 | 0.011 |
| | | 38,108,710 | | | -4.415 | 2.10E-05 | 0.039 |
| | | 38,119,198 | *GSDMA* | | 4.797 | 4.34E-06 | 0.009 |
| | | 38,119,207 | *GSDMA* | | 5.722 | 6.91E-08 | 0.009 |
| | | 38,119,649 | | *GSDMA* | 4.784 | 4.60E-06 | 0.002 |
| | | 38,120,370 | | *GSDMA* | 3.626 | 4.13E-04 | 0.011 |
| rs9916765 | g.38005595.C>T | 37,828,296 | | *PGAP3* | 3.522 | 5.91E-04 | 0.002 |
| | | 37,927,002 | | *IKZF3* | -3.572 | 4.98E-04 | 0.001 |
| | | 38,018,804 | | *IKZF3* | 4.164 | 5.66E-05 | 0.046 |
| | | 38,077,191 | | | 4.055 | 8.59E-05 | 0.034 |
| | | 38,080,829 | | *ORMDL3* | 5.194 | 7.75E-07 | 0.014 |
| | | 38,081,812 | | *ORMDL3* | 3.646 | 3.84E-04 | 2.01E-04 |

| | | | | | | | |
|---|---|---|---|---|---|---|---|
| | | 38,081,995 | | *ORMDL3* | 4.541 | 1.26E-05 | 2.01E-04 |
| | | 38,082,022 | | *ORMDL3* | 5.042 | 1.51E-06 | 6.40E-04 |
| | | 38,082,035 | | *ORMDL3* | 6.398 | 2.61E-09 | 6.40E-04 |
| | | 38,082,073 | | *ORMDL3* | 5.731 | 6.63E-08 | 6.40E-04 |
| | | 38,082,235 | | *ORMDL3* | 3.673 | 3.49E-04 | 0.004 |
| | | 38,082,346 | | *ORMDL3* | 4.786 | 4.54E-06 | 0.011 |
| | | 38,082,448 | | *ORMDL3* | 4.304 | 3.28E-05 | 0.011 |
| | | 38,082,658 | | *ORMDL3* | 4.404 | 2.20E-05 | 0.011 |
| | | 38,082,778 | | *ORMDL3* | 3.634 | 4.00E-04 | 0.011 |
| | | 38,108,710 | | | -4.178 | 5.37E-05 | 0.039 |
| | | 38,119,198 | *GSDMA* | | 4.357 | 2.65E-05 | 0.009 |
| | | 38,119,207 | *GSDMA* | | 5.856 | 3.67E-08 | 0.009 |
| | | 38,119,649 | | *GSDMA* | 4.338 | 2.86E-05 | 0.002 |
| rs202126107 | g.38006767_38006776del | 37,828,296 | | *PGAP3* | 3.890 | 1.59E-04 | 0.002 |
| | | 37,927,002 | | *IKZF3* | -3.505 | 6.26E-04 | 0.001 |
| | | 38,018,804 | | *IKZF3* | 3.500 | 6.38E-04 | 0.046 |
| | | 38,077,191 | | | 4.279 | 3.61E-05 | 0.034 |
| | | 38,081,812 | | *ORMDL3* | 3.691 | 3.27E-04 | 2.01E-04 |
| | | 38,081,829 | | *ORMDL3* | 3.902 | 1.52E-04 | 2.01E-04 |
| | | 38,081,995 | | *ORMDL3* | 5.878 | 3.29E-08 | 2.01E-04 |
| | | 38,082,022 | | *ORMDL3* | 6.688 | 6.07E-10 | 6.40E-04 |
| | | 38,082,035 | | *ORMDL3* | 7.026 | 1.07E-10 | 6.40E-04 |
| | | 38,082,073 | | *ORMDL3* | 6.666 | 6.80E-10 | 6.40E-04 |
| | | 38,082,206 | | *ORMDL3* | 3.732 | 2.83E-04 | 0.004 |
| | | 38,082,235 | | *ORMDL3* | 3.999 | 1.06E-04 | 0.004 |
| | | 38,082,346 | | *ORMDL3* | 4.711 | 6.24E-06 | 0.011 |
| | | 38,082,448 | | *ORMDL3* | 3.776 | 2.41E-04 | 0.011 |
| | | 38,082,658 | | *ORMDL3* | 4.319 | 3.08E-05 | 0.011 |
| | | 38,082,778 | | *ORMDL3* | 4.101 | 7.20E-05 | 0.011 |
| | | 38,108,710 | | | -4.792 | 4.44E-06 | 0.039 |
| | | 38,119,198 | *GSDMA* | | 4.384 | 2.38E-05 | 0.009 |
| | | 38,119,207 | *GSDMA* | | 5.463 | 2.29E-07 | 0.009 |
| | | 38,119,649 | | *GSDMA* | 4.576 | 1.09E-05 | 0.002 |
| rs35564481 | g.38020058_38020058insC | 37,828,296 | | *PGAP3* | 3.538 | 5.59E-04 | 0.002 |
| | | 37,927,002 | | *IKZF3* | -3.751 | 2.64E-04 | 0.001 |
| | | 38,018,804 | | *IKZF3* | 4.189 | 5.14E-05 | 0.046 |
| | | 38,077,191 | | | 3.985 | 1.12E-04 | 0.034 |
| | | 38,080,829 | | *ORMDL3* | 4.942 | 2.34E-06 | 0.014 |
| | | 38,081,492 | | *ORMDL3* | 3.542 | 5.52E-04 | 2.01E-04 |

| | | | | | | | |
|---|---|---|---|---|---|---|---|
| | | 38,081,812 | | *ORMDL3* | 3.829 | 1.99E-04 | 2.01E-04 |
| | | 38,081,995 | | *ORMDL3* | 4.171 | 5.51E-05 | 2.01E-04 |
| | | 38,082,022 | | *ORMDL3* | 5.143 | 9.72E-07 | 6.40E-04 |
| | | 38,082,035 | | *ORMDL3* | 6.548 | 1.23E-09 | 6.40E-04 |
| | | 38,082,073 | | *ORMDL3* | 5.237 | 6.38E-07 | 6.40E-04 |
| | | 38,082,235 | | *ORMDL3* | 3.564 | 5.12E-04 | 0.004 |
| | | 38,082,346 | | *ORMDL3* | 4.650 | 8.03E-06 | 0.011 |
| | | 38,082,448 | | *ORMDL3* | 4.413 | 2.12E-05 | 0.011 |
| | | 38,082,658 | | *ORMDL3* | 4.624 | 8.97E-06 | 0.011 |
| | | 38,082,778 | | *ORMDL3* | 3.506 | 6.25E-04 | 0.011 |
| | | 38,108,710 | | | -3.999 | 1.06E-04 | 0.039 |
| | | 38,119,198 | *GSDMA* | | 4.226 | 4.44E-05 | 0.009 |
| | | 38,119,207 | *GSDMA* | | 5.417 | 2.83E-07 | 0.009 |
| | | 38,119,649 | | *GSDMA* | 3.666 | 3.57E-04 | 0.002 |
| rs9635726 | g.38020141.T>C | 37,828,296 | | *PGAP3* | -3.934 | 1.35E-04 | 0.002 |
| | | 37,828,560 | | *PGAP3* | -4.321 | 3.06E-05 | 0.002 |
| | | 37,856,551 | | *ERBB2* | 3.507 | 6.22E-04 | 0.006 |
| | | 38,082,035 | | *ORMDL3* | -3.381 | 9.54E-04 | 6.40E-04 |
| | | 38,082,073 | | *ORMDL3* | -4.555 | 1.19E-05 | 6.40E-04 |
| rs1453559 | g.38020419.G>A | 37,828,296 | | *PGAP3* | 3.522 | 5.91E-04 | 0.002 |
| | | 37,927,002 | | *IKZF3* | -3.572 | 4.98E-04 | 0.001 |
| | | 38,018,804 | | *IKZF3* | 4.164 | 5.66E-05 | 0.046 |
| | | 38,077,191 | | | 4.055 | 8.59E-05 | 0.034 |
| | | 38,080,829 | | *ORMDL3* | 5.194 | 7.75E-07 | 0.014 |
| | | 38,081,812 | | *ORMDL3* | 3.646 | 3.84E-04 | 2.01E-04 |
| | | 38,081,995 | | *ORMDL3* | 4.541 | 1.26E-05 | 2.01E-04 |
| | | 38,082,022 | | *ORMDL3* | 5.042 | 1.51E-06 | 6.40E-04 |
| | | 38,082,035 | | *ORMDL3* | 6.398 | 2.61E-09 | 6.40E-04 |
| | | 38,082,073 | | *ORMDL3* | 5.731 | 6.63E-08 | 6.40E-04 |
| | | 38,082,235 | | *ORMDL3* | 3.673 | 3.49E-04 | 0.004 |
| | | 38,082,346 | | *ORMDL3* | 4.786 | 4.54E-06 | 0.011 |
| | | 38,082,448 | | *ORMDL3* | 4.304 | 3.28E-05 | 0.011 |
| | | 38,082,658 | | *ORMDL3* | 4.404 | 2.20E-05 | 0.011 |
| | | 38,082,778 | | *ORMDL3* | 3.634 | 4.00E-04 | 0.011 |
| | | 38,108,710 | | | -4.178 | 5.37E-05 | 0.039 |
| | | 38,119,198 | *GSDMA* | | 4.357 | 2.65E-05 | 0.009 |
| | | 38,119,207 | *GSDMA* | | 5.856 | 3.67E-08 | 0.009 |
| | | 38,119,649 | | *GSDMA* | 4.338 | 2.86E-05 | 0.002 |
| rs4795397 | g.38023745.G>A | 37,828,296 | | *PGAP3* | 3.890 | 1.59E-04 | 0.002 |
| | | 37,927,002 | | *IKZF3* | -3.505 | 6.26E-04 | 0.001 |
| | | 38,018,804 | | *IKZF3* | 3.500 | 6.38E-04 | 0.046 |

| | | | | | | | |
|---|---|---|---|---|---|---|---|
| | | 38,077,191 | | | 4.279 | 3.61E-05 | 0.034 |
| | | 38,081,812 | | *ORMDL3* | 3.691 | 3.27E-04 | 2.01E-04 |
| | | 38,081,829 | | *ORMDL3* | 3.902 | 1.52E-04 | 2.01E-04 |
| | | 38,081,995 | | *ORMDL3* | 5.878 | 3.29E-08 | 2.01E-04 |
| | | 38,082,022 | | *ORMDL3* | 6.688 | 6.07E-10 | 6.40E-04 |
| | | 38,082,035 | | *ORMDL3* | 7.026 | 1.07E-10 | 6.40E-04 |
| | | 38,082,073 | | *ORMDL3* | 6.666 | 6.80E-10 | 6.40E-04 |
| | | 38,082,206 | | *ORMDL3* | 3.732 | 2.83E-04 | 0.004 |
| | | 38,082,235 | | *ORMDL3* | 3.999 | 1.06E-04 | 0.004 |
| | | 38,082,346 | | *ORMDL3* | 4.711 | 6.24E-06 | 0.011 |
| | | 38,082,448 | | *ORMDL3* | 3.776 | 2.41E-04 | 0.011 |
| | | 38,082,658 | | *ORMDL3* | 4.319 | 3.08E-05 | 0.011 |
| | | 38,082,778 | | *ORMDL3* | 4.101 | 7.20E-05 | 0.011 |
| | | 38,108,710 | | | -4.792 | 4.44E-06 | 0.039 |
| | | 38,119,198 | *GSDMA* | | 4.384 | 2.38E-05 | 0.009 |
| | | 38,119,207 | *GSDMA* | | 5.463 | 2.29E-07 | 0.009 |
| | | 38,119,649 | | *GSDMA* | 4.576 | 1.09E-05 | 0.002 |
| rs11557466 | g.38024626.T>C (Leu.7.Leu) | 37,828,296 | | *PGAP3* | 3.739 | 2.75E-04 | 0.002 |
| | | 37,927,002 | | *IKZF3* | -3.528 | 5.80E-04 | 0.001 |
| | | 38,018,804 | | *IKZF3* | 3.616 | 4.26E-04 | 0.046 |
| | | 38,077,191 | | | 4.332 | 2.93E-05 | 0.034 |
| | | 38,080,829 | | *ORMDL3* | 3.435 | 7.95E-04 | 0.014 |
| | | 38,081,812 | | *ORMDL3* | 3.459 | 7.34E-04 | 2.01E-04 |
| | | 38,081,829 | | *ORMDL3* | 3.730 | 2.85E-04 | 2.01E-04 |
| | | 38,081,995 | | *ORMDL3* | 5.842 | 3.92E-08 | 2.01E-04 |
| | | 38,082,022 | | *ORMDL3* | 6.694 | 5.90E-10 | 6.40E-04 |
| | | 38,082,035 | | *ORMDL3* | 6.772 | 3.96E-10 | 6.40E-04 |
| | | 38,082,073 | | *ORMDL3* | 6.832 | 2.91E-10 | 6.40E-04 |
| | | 38,082,206 | | *ORMDL3* | 3.847 | 1.86E-04 | 0.004 |
| | | 38,082,235 | | *ORMDL3* | 4.278 | 3.62E-05 | 0.004 |
| | | 38,082,346 | | *ORMDL3* | 4.590 | 1.03E-05 | 0.011 |
| | | 38,082,448 | | *ORMDL3* | 3.644 | 3.87E-04 | 0.011 |
| | | 38,082,658 | | *ORMDL3* | 4.266 | 3.79E-05 | 0.011 |
| | | 38,082,778 | | *ORMDL3* | 4.147 | 6.05E-05 | 0.011 |
| | | 38,108,710 | | | -4.669 | 7.42E-06 | 0.039 |
| | | 38,119,198 | *GSDMA* | | 4.246 | 4.11E-05 | 0.009 |
| | | 38,119,207 | *GSDMA* | | 5.331 | 4.19E-07 | 0.009 |
| | | 38,119,649 | | *GSDMA* | 4.404 | 2.19E-05 | 0.002 |
| rs11078925 | g.38025208.C>T | 37,828,296 | | *PGAP3* | 3.739 | 2.75E-04 | 0.002 |
| | | 37,927,002 | | *IKZF3* | -3.528 | 5.80E-04 | 0.001 |

| | | | | | | | |
|---|---|---|---|---|---|---|---|
| | | 38,018,804 | | *IKZF3* | 3.616 | 4.26E-04 | 0.046 |
| | | 38,077,191 | | | 4.332 | 2.93E-05 | 0.034 |
| | | 38,080,829 | | *ORMDL3* | 3.435 | 7.95E-04 | 0.014 |
| | | 38,081,812 | | *ORMDL3* | 3.459 | 7.34E-04 | 2.01E-04 |
| | | 38,081,829 | | *ORMDL3* | 3.730 | 2.85E-04 | 2.01E-04 |
| | | 38,081,995 | | *ORMDL3* | 5.842 | 3.92E-08 | 2.01E-04 |
| | | 38,082,022 | | *ORMDL3* | 6.694 | 5.90E-10 | 6.40E-04 |
| | | 38,082,035 | | *ORMDL3* | 6.772 | 3.96E-10 | 6.40E-04 |
| | | 38,082,073 | | *ORMDL3* | 6.832 | 2.91E-10 | 6.40E-04 |
| | | 38,082,206 | | *ORMDL3* | 3.847 | 1.86E-04 | 0.004 |
| | | 38,082,235 | | *ORMDL3* | 4.278 | 3.62E-05 | 0.004 |
| | | 38,082,346 | | *ORMDL3* | 4.590 | 1.03E-05 | 0.011 |
| | | 38,082,448 | | *ORMDL3* | 3.644 | 3.87E-04 | 0.011 |
| | | 38,082,658 | | *ORMDL3* | 4.266 | 3.79E-05 | 0.011 |
| | | 38,082,778 | | *ORMDL3* | 4.147 | 6.05E-05 | 0.011 |
| | | 38,108,710 | | | -4.669 | 7.42E-06 | 0.039 |
| | | 38,119,198 | *GSDMA* | | 4.246 | 4.11E-05 | 0.009 |
| | | 38,119,207 | *GSDMA* | | 5.331 | 4.19E-07 | 0.009 |
| | | 38,119,649 | | *GSDMA* | 4.404 | 2.19E-05 | 0.002 |
| rs12150079 | g.38025417.A>G | 38,024,331 | | | -3.759 | 2.57E-04 | 0.017 |
| | | 38,072,514 | | *GSDMB* | 3.730 | 2.85E-04 | 0.046 |
| | | 38,082,073 | | *ORMDL3* | 4.507 | 1.45E-05 | 6.40E-04 |
| | | 38,108,710 | | | -4.053 | 8.65E-05 | 0.039 |
| | | 38,119,198 | *GSDMA* | | 4.128 | 6.51E-05 | 0.009 |
| | | 38,119,207 | *GSDMA* | | 4.584 | 1.06E-05 | 0.009 |
| | | 38,119,649 | | *GSDMA* | 4.300 | 3.33E-05 | 0.002 |
| rs111940435 | g.38025904.T>C | 37,856,732 | | *ERBB2* | -3.668 | 3.56E-04 | 0.009 |
| | | 37,886,749 | | *MIEN1* | 3.945 | 1.30E-04 | 0.002 |
| | | 37,896,063 | | | 3.476 | 6.92E-04 | 0.009 |
| | | 38,020,485 | *IKZF3* | | 3.962 | 1.22E-04 | 0.005 |
| | | 38,080,829 | | *ORMDL3* | 3.995 | 1.08E-04 | 0.014 |
| | | 38,096,333 | | | -3.664 | 3.60E-04 | 0.033 |
| | | 38,096,571 | | | -4.780 | 4.67E-06 | 0.033 |
| | | 38,096,578 | | | -5.642 | 1.00E-07 | 0.033 |
| | | 38,096,648 | | | -3.887 | 1.61E-04 | 0.033 |
| rs34120102 | g.38026035.A>G | 37,828,296 | | *PGAP3* | 3.739 | 2.75E-04 | 0.002 |
| | | 37,927,002 | | *IKZF3* | -3.528 | 5.80E-04 | 0.001 |
| | | 38,018,804 | | *IKZF3* | 3.616 | 4.26E-04 | 0.046 |
| | | 38,077,191 | | | 4.332 | 2.93E-05 | 0.034 |
| | | 38,080,829 | | *ORMDL3* | 3.435 | 7.95E-04 | 0.014 |
| | | 38,081,812 | | *ORMDL3* | 3.459 | 7.34E-04 | 2.01E-04 |

| | | | | | | | |
|---|---|---|---|---|---|---|---|
| | | 38,081,829 | | *ORMDL3* | 3.730 | 2.85E-04 | 2.01E-04 |
| | | 38,081,995 | | *ORMDL3* | 5.842 | 3.92E-08 | 2.01E-04 |
| | | 38,082,022 | | *ORMDL3* | 6.694 | 5.90E-10 | 6.40E-04 |
| | | 38,082,035 | | *ORMDL3* | 6.772 | 3.96E-10 | 6.40E-04 |
| | | 38,082,073 | | *ORMDL3* | 6.832 | 2.91E-10 | 6.40E-04 |
| | | 38,082,206 | | *ORMDL3* | 3.847 | 1.86E-04 | 0.004 |
| | | 38,082,235 | | *ORMDL3* | 4.278 | 3.62E-05 | 0.004 |
| | | 38,082,346 | | *ORMDL3* | 4.590 | 1.03E-05 | 0.011 |
| | | 38,082,448 | | *ORMDL3* | 3.644 | 3.87E-04 | 0.011 |
| | | 38,082,658 | | *ORMDL3* | 4.266 | 3.79E-05 | 0.011 |
| | | 38,082,778 | | *ORMDL3* | 4.147 | 6.05E-05 | 0.011 |
| | | 38,108,710 | | | -4.669 | 7.42E-06 | 0.039 |
| | | 38,119,198 | *GSDMA* | | 4.246 | 4.11E-05 | 0.009 |
| | | 38,119,207 | *GSDMA* | | 5.331 | 4.19E-07 | 0.009 |
| | | 38,119,649 | | *GSDMA* | 4.404 | 2.19E-05 | 0.002 |
| rs11655198 | g.38026169.T>C | 37,927,002 | | *IKZF3* | -3.588 | 4.70E-04 | 0.001 |
| | | 38,018,804 | | *IKZF3* | 4.291 | 3.45E-05 | 0.046 |
| | | 38,049,272 | | | 3.531 | 5.74E-04 | 0.046 |
| | | 38,077,191 | | | 4.101 | 7.22E-05 | 0.034 |
| | | 38,080,829 | | *ORMDL3* | 5.326 | 4.28E-07 | 0.014 |
| | | 38,081,812 | | *ORMDL3* | 3.387 | 9.35E-04 | 2.01E-04 |
| | | 38,081,995 | | *ORMDL3* | 4.489 | 1.56E-05 | 2.01E-04 |
| | | 38,082,022 | | *ORMDL3* | 5.027 | 1.62E-06 | 6.40E-04 |
| | | 38,082,035 | | *ORMDL3* | 6.122 | 1.02E-08 | 6.40E-04 |
| | | 38,082,073 | | *ORMDL3* | 5.874 | 3.36E-08 | 6.40E-04 |
| | | 38,082,235 | | *ORMDL3* | 3.957 | 1.24E-04 | 0.004 |
| | | 38,082,346 | | *ORMDL3* | 4.643 | 8.28E-06 | 0.011 |
| | | 38,082,448 | | *ORMDL3* | 4.146 | 6.05E-05 | 0.011 |
| | | 38,082,658 | | *ORMDL3* | 4.337 | 2.87E-05 | 0.011 |
| | | 38,082,778 | | *ORMDL3* | 3.671 | 3.51E-04 | 0.011 |
| | | 38,108,710 | | | -4.040 | 9.10E-05 | 0.039 |
| | | 38,119,198 | *GSDMA* | | 4.199 | 4.94E-05 | 0.009 |
| | | 38,119,207 | *GSDMA* | | 5.693 | 7.91E-08 | 0.009 |
| | | 38,119,649 | | *GSDMA* | 4.145 | 6.08E-05 | 0.002 |
| rs11650661 | g.38026286.T>A | 37,927,002 | | *IKZF3* | -3.588 | 4.70E-04 | 0.001 |
| | | 38,018,804 | | *IKZF3* | 4.291 | 3.45E-05 | 0.046 |
| | | 38,049,272 | | | 3.531 | 5.74E-04 | 0.046 |
| | | 38,077,191 | | | 4.101 | 7.22E-05 | 0.034 |
| | | 38,080,829 | | *ORMDL3* | 5.326 | 4.28E-07 | 0.014 |
| | | 38,081,812 | | *ORMDL3* | 3.387 | 9.35E-04 | 2.01E-04 |
| | | 38,081,995 | | *ORMDL3* | 4.489 | 1.56E-05 | 2.01E-04 |

| | | | | | | | |
|---|---|---|---|---|---|---|---|
| | | 38,082,022 | | *ORMDL3* | 5.027 | 1.62E-06 | 6.40E-04 |
| | | 38,082,035 | | *ORMDL3* | 6.122 | 1.02E-08 | 6.40E-04 |
| | | 38,082,073 | | *ORMDL3* | 5.874 | 3.36E-08 | 6.40E-04 |
| | | 38,082,235 | | *ORMDL3* | 3.957 | 1.24E-04 | 0.004 |
| | | 38,082,346 | | *ORMDL3* | 4.643 | 8.28E-06 | 0.011 |
| | | 38,082,448 | | *ORMDL3* | 4.146 | 6.05E-05 | 0.011 |
| | | 38,082,658 | | *ORMDL3* | 4.337 | 2.87E-05 | 0.011 |
| | | 38,082,778 | | *ORMDL3* | 3.671 | 3.51E-04 | 0.011 |
| | | 38,108,710 | | | -4.040 | 9.10E-05 | 0.039 |
| | | 38,119,198 | *GSDMA* | | 4.199 | 4.94E-05 | 0.009 |
| | | 38,119,207 | *GSDMA* | | 5.693 | 7.91E-08 | 0.009 |
| | | 38,119,649 | | *GSDMA* | 4.145 | 6.08E-05 | 0.002 |
| rs11655292 | g.38026361.G>C | 37,927,002 | | *IKZF3* | -3.588 | 4.70E-04 | 0.001 |
| | | 38,018,804 | | *IKZF3* | 4.291 | 3.45E-05 | 0.046 |
| | | 38,049,272 | | | 3.531 | 5.74E-04 | 0.046 |
| | | 38,077,191 | | | 4.101 | 7.22E-05 | 0.034 |
| | | 38,080,829 | | *ORMDL3* | 5.326 | 4.28E-07 | 0.014 |
| | | 38,081,812 | | *ORMDL3* | 3.387 | 9.35E-04 | 2.01E-04 |
| | | 38,081,995 | | *ORMDL3* | 4.489 | 1.56E-05 | 2.01E-04 |
| | | 38,082,022 | | *ORMDL3* | 5.027 | 1.62E-06 | 6.40E-04 |
| | | 38,082,035 | | *ORMDL3* | 6.122 | 1.02E-08 | 6.40E-04 |
| | | 38,082,073 | | *ORMDL3* | 5.874 | 3.36E-08 | 6.40E-04 |
| | | 38,082,235 | | *ORMDL3* | 3.957 | 1.24E-04 | 0.004 |
| | | 38,082,346 | | *ORMDL3* | 4.643 | 8.28E-06 | 0.011 |
| | | 38,082,448 | | *ORMDL3* | 4.146 | 6.05E-05 | 0.011 |
| | | 38,082,658 | | *ORMDL3* | 4.337 | 2.87E-05 | 0.011 |
| | | 38,082,778 | | *ORMDL3* | 3.671 | 3.51E-04 | 0.011 |
| | | 38,108,710 | | | -4.040 | 9.10E-05 | 0.039 |
| | | 38,119,198 | *GSDMA* | | 4.199 | 4.94E-05 | 0.009 |
| | | 38,119,207 | *GSDMA* | | 5.693 | 7.91E-08 | 0.009 |
| | | 38,119,649 | | *GSDMA* | 4.145 | 6.08E-05 | 0.002 |
| rs12709365 | g.38027400.G>A | 37,828,296 | | *PGAP3* | 3.739 | 2.75E-04 | 0.002 |
| | | 37,927,002 | | *IKZF3* | -3.528 | 5.80E-04 | 0.001 |
| | | 38,018,804 | | *IKZF3* | 3.616 | 4.26E-04 | 0.046 |
| | | 38,077,191 | | | 4.332 | 2.93E-05 | 0.034 |
| | | 38,080,829 | | *ORMDL3* | 3.435 | 7.95E-04 | 0.014 |
| | | 38,081,812 | | *ORMDL3* | 3.459 | 7.34E-04 | 2.01E-04 |
| | | 38,081,829 | | *ORMDL3* | 3.730 | 2.85E-04 | 2.01E-04 |
| | | 38,081,995 | | *ORMDL3* | 5.842 | 3.92E-08 | 2.01E-04 |
| | | 38,082,022 | | *ORMDL3* | 6.694 | 5.90E-10 | 6.40E-04 |
| | | 38,082,035 | | *ORMDL3* | 6.772 | 3.96E-10 | 6.40E-04 |

| | | | | | | | |
|---|---|---|---|---|---|---|---|
| | | 38,082,073 | | *ORMDL3* | 6.832 | 2.91E-10 | 6.40E-04 |
| | | 38,082,206 | | *ORMDL3* | 3.847 | 1.86E-04 | 0.004 |
| | | 38,082,235 | | *ORMDL3* | 4.278 | 3.62E-05 | 0.004 |
| | | 38,082,346 | | *ORMDL3* | 4.590 | 1.03E-05 | 0.011 |
| | | 38,082,448 | | *ORMDL3* | 3.644 | 3.87E-04 | 0.011 |
| | | 38,082,658 | | *ORMDL3* | 4.266 | 3.79E-05 | 0.011 |
| | | 38,082,778 | | *ORMDL3* | 4.147 | 6.05E-05 | 0.011 |
| | | 38,108,710 | | | -4.669 | 7.42E-06 | 0.039 |
| | | 38,119,198 | *GSDMA* | | 4.246 | 4.11E-05 | 0.009 |
| | | 38,119,207 | *GSDMA* | | 5.331 | 4.19E-07 | 0.009 |
| | | 38,119,649 | | *GSDMA* | 4.404 | 2.19E-05 | 0.002 |
| rs13380815 | g.38027583.G>A | 37,828,296 | | *PGAP3* | 3.739 | 2.75E-04 | 0.002 |
| | | 37,927,002 | | *IKZF3* | -3.528 | 5.80E-04 | 0.001 |
| | | 38,018,804 | | *IKZF3* | 3.616 | 4.26E-04 | 0.046 |
| | | 38,077,191 | | | 4.332 | 2.93E-05 | 0.034 |
| | | 38,080,829 | | *ORMDL3* | 3.435 | 7.95E-04 | 0.014 |
| | | 38,081,812 | | *ORMDL3* | 3.459 | 7.34E-04 | 2.01E-04 |
| | | 38,081,829 | | *ORMDL3* | 3.730 | 2.85E-04 | 2.01E-04 |
| | | 38,081,995 | | *ORMDL3* | 5.842 | 3.92E-08 | 2.01E-04 |
| | | 38,082,022 | | *ORMDL3* | 6.694 | 5.90E-10 | 6.40E-04 |
| | | 38,082,035 | | *ORMDL3* | 6.772 | 3.96E-10 | 6.40E-04 |
| | | 38,082,073 | | *ORMDL3* | 6.832 | 2.91E-10 | 6.40E-04 |
| | | 38,082,206 | | *ORMDL3* | 3.847 | 1.86E-04 | 0.004 |
| | | 38,082,235 | | *ORMDL3* | 4.278 | 3.62E-05 | 0.004 |
| | | 38,082,346 | | *ORMDL3* | 4.590 | 1.03E-05 | 0.011 |
| | | 38,082,448 | | *ORMDL3* | 3.644 | 3.87E-04 | 0.011 |
| | | 38,082,658 | | *ORMDL3* | 4.266 | 3.79E-05 | 0.011 |
| | | 38,082,778 | | *ORMDL3* | 4.147 | 6.05E-05 | 0.011 |
| | | 38,108,710 | | | -4.669 | 7.42E-06 | 0.039 |
| | | 38,119,198 | *GSDMA* | | 4.246 | 4.11E-05 | 0.009 |
| | | 38,119,207 | *GSDMA* | | 5.331 | 4.19E-07 | 0.009 |
| | | 38,119,649 | | *GSDMA* | 4.404 | 2.19E-05 | 0.002 |
| rs113606921 | g.38028254.C>T | 37,856,732 | | *ERBB2* | -3.668 | 3.56E-04 | 0.009 |
| | | 37,886,749 | | *MIEN1* | 3.945 | 1.30E-04 | 0.002 |
| | | 37,896,063 | | | 3.476 | 6.92E-04 | 0.009 |
| | | 38,020,485 | *IKZF3* | | 3.962 | 1.22E-04 | 0.005 |
| | | 38,080,829 | | *ORMDL3* | 3.995 | 1.08E-04 | 0.014 |
| | | 38,096,333 | | | -3.664 | 3.60E-04 | 0.033 |
| | | 38,096,571 | | | -4.780 | 4.67E-06 | 0.033 |
| | | 38,096,578 | | | -5.642 | 1.00E-07 | 0.033 |
| | | 38,096,648 | | | -3.887 | 1.61E-04 | 0.033 |

| | | | | | | | |
|---|---|---|---|---|---|---|---|
| rs111845187 | g.38028497.G>C | 37,856,732 | | *ERBB2* | -3.668 | 3.56E-04 | 0.009 |
| | | 37,886,749 | | *MIEN1* | 3.945 | 1.30E-04 | 0.002 |
| | | 37,896,063 | | | 3.476 | 6.92E-04 | 0.009 |
| | | 38,020,485 | *IKZF3* | | 3.962 | 1.22E-04 | 0.005 |
| | | 38,080,829 | | *ORMDL3* | 3.995 | 1.08E-04 | 0.014 |
| | | 38,096,333 | | | -3.664 | 3.60E-04 | 0.033 |
| | | 38,096,571 | | | -4.780 | 4.67E-06 | 0.033 |
| | | 38,096,578 | | | -5.642 | 1.00E-07 | 0.033 |
| | | 38,096,648 | | | -3.887 | 1.61E-04 | 0.033 |
| rs11557467 | g.38028634.T>G (Ser.173.Ile) | 37,927,002 | | *IKZF3* | -3.588 | 4.70E-04 | 0.001 |
| | | 38,018,804 | | *IKZF3* | 4.291 | 3.45E-05 | 0.046 |
| | | 38,049,272 | | | 3.531 | 5.74E-04 | 0.046 |
| | | 38,077,191 | | | 4.101 | 7.22E-05 | 0.034 |
| | | 38,080,829 | | *ORMDL3* | 5.326 | 4.28E-07 | 0.014 |
| | | 38,081,812 | | *ORMDL3* | 3.387 | 9.35E-04 | 2.01E-04 |
| | | 38,081,995 | | *ORMDL3* | 4.489 | 1.56E-05 | 2.01E-04 |
| | | 38,082,022 | | *ORMDL3* | 5.027 | 1.62E-06 | 6.40E-04 |
| | | 38,082,035 | | *ORMDL3* | 6.122 | 1.02E-08 | 6.40E-04 |
| | | 38,082,073 | | *ORMDL3* | 5.874 | 3.36E-08 | 6.40E-04 |
| | | 38,082,235 | | *ORMDL3* | 3.957 | 1.24E-04 | 0.004 |
| | | 38,082,346 | | *ORMDL3* | 4.643 | 8.28E-06 | 0.011 |
| | | 38,082,448 | | *ORMDL3* | 4.146 | 6.05E-05 | 0.011 |
| | | 38,082,658 | | *ORMDL3* | 4.337 | 2.87E-05 | 0.011 |
| | | 38,082,778 | | *ORMDL3* | 3.671 | 3.51E-04 | 0.011 |
| | | 38,108,710 | | | -4.040 | 9.10E-05 | 0.039 |
| | | 38,119,198 | *GSDMA* | | 4.199 | 4.94E-05 | 0.009 |
| | | 38,119,207 | *GSDMA* | | 5.693 | 7.91E-08 | 0.009 |
| | | 38,119,649 | | *GSDMA* | 4.145 | 6.08E-05 | 0.002 |
| rs112561568 | g.38028857.A>G | 37,856,732 | | *ERBB2* | -3.668 | 3.56E-04 | 0.009 |
| | | 37,886,749 | | *MIEN1* | 3.945 | 1.30E-04 | 0.002 |
| | | 37,896,063 | | | 3.476 | 6.92E-04 | 0.009 |
| | | 38,020,485 | *IKZF3* | | 3.962 | 1.22E-04 | 0.005 |
| | | 38,080,829 | | *ORMDL3* | 3.995 | 1.08E-04 | 0.014 |
| | | 38,096,333 | | | -3.664 | 3.60E-04 | 0.033 |
| | | 38,096,571 | | | -4.780 | 4.67E-06 | 0.033 |
| | | 38,096,578 | | | -5.642 | 1.00E-07 | 0.033 |
| | | 38,096,648 | | | -3.887 | 1.61E-04 | 0.033 |
| rs12936231 | g.38029120.G>C | 37,927,002 | | *IKZF3* | -3.588 | 4.70E-04 | 0.001 |
| | | 38,018,804 | | *IKZF3* | 4.291 | 3.45E-05 | 0.046 |
| | | 38,049,272 | | | 3.531 | 5.74E-04 | 0.046 |

| | | | | | | | |
|---|---|---|---|---|---|---|---|
| | | 38,077,191 | | | 4.101 | 7.22E-05 | 0.034 |
| | | 38,080,829 | | *ORMDL3* | 5.326 | 4.28E-07 | 0.014 |
| | | 38,081,812 | | *ORMDL3* | 3.387 | 9.35E-04 | 2.01E-04 |
| | | 38,081,995 | | *ORMDL3* | 4.489 | 1.56E-05 | 2.01E-04 |
| | | 38,082,022 | | *ORMDL3* | 5.027 | 1.62E-06 | 6.40E-04 |
| | | 38,082,035 | | *ORMDL3* | 6.122 | 1.02E-08 | 6.40E-04 |
| | | 38,082,073 | | *ORMDL3* | 5.874 | 3.36E-08 | 6.40E-04 |
| | | 38,082,235 | | *ORMDL3* | 3.957 | 1.24E-04 | 0.004 |
| | | 38,082,346 | | *ORMDL3* | 4.643 | 8.28E-06 | 0.011 |
| | | 38,082,448 | | *ORMDL3* | 4.146 | 6.05E-05 | 0.011 |
| | | 38,082,658 | | *ORMDL3* | 4.337 | 2.87E-05 | 0.011 |
| | | 38,082,778 | | *ORMDL3* | 3.671 | 3.51E-04 | 0.011 |
| | | 38,108,710 | | | -4.040 | 9.10E-05 | 0.039 |
| | | 38,119,198 | *GSDMA* | | 4.199 | 4.94E-05 | 0.009 |
| | | 38,119,207 | *GSDMA* | | 5.693 | 7.91E-08 | 0.009 |
| | | 38,119,649 | | *GSDMA* | 4.145 | 6.08E-05 | 0.002 |
| rs147134135 | g.38029867.C>A | 37,856,732 | | *ERBB2* | -3.668 | 3.56E-04 | 0.009 |
| | | 37,886,749 | | *MIEN1* | 3.945 | 1.30E-04 | 0.002 |
| | | 37,896,063 | | | 3.476 | 6.92E-04 | 0.009 |
| | | 38,020,485 | *IKZF3* | | 3.962 | 1.22E-04 | 0.005 |
| | | 38,080,829 | | *ORMDL3* | 3.995 | 1.08E-04 | 0.014 |
| | | 38,096,333 | | | -3.664 | 3.60E-04 | 0.033 |
| | | 38,096,571 | | | -4.780 | 4.67E-06 | 0.033 |
| | | 38,096,578 | | | -5.642 | 1.00E-07 | 0.033 |
| | | 38,096,648 | | | -3.887 | 1.61E-04 | 0.033 |
| rs11870965 | g.38030205.A>T | 37,828,296 | | *PGAP3* | 3.784 | 2.34E-04 | 0.002 |
| | | 37,927,002 | | *IKZF3* | -3.552 | 5.33E-04 | 0.001 |
| | | 38,018,804 | | *IKZF3* | 3.727 | 2.88E-04 | 0.046 |
| | | 38,077,191 | | | 4.473 | 1.67E-05 | 0.034 |
| | | 38,081,812 | | *ORMDL3* | 3.501 | 6.36E-04 | 2.01E-04 |
| | | 38,081,829 | | *ORMDL3* | 3.794 | 2.26E-04 | 2.01E-04 |
| | | 38,081,995 | | *ORMDL3* | 5.870 | 3.43E-08 | 2.01E-04 |
| | | 38,082,022 | | *ORMDL3* | 6.643 | 7.62E-10 | 6.40E-04 |
| | | 38,082,035 | | *ORMDL3* | 6.734 | 4.80E-10 | 6.40E-04 |
| | | 38,082,073 | | *ORMDL3* | 6.834 | 2.88E-10 | 6.40E-04 |
| | | 38,082,206 | | *ORMDL3* | 3.794 | 2.27E-04 | 0.004 |
| | | 38,082,235 | | *ORMDL3* | 4.231 | 4.36E-05 | 0.004 |
| | | 38,082,346 | | *ORMDL3* | 4.572 | 1.11E-05 | 0.011 |
| | | 38,082,448 | | *ORMDL3* | 3.631 | 4.05E-04 | 0.011 |
| | | 38,082,658 | | *ORMDL3* | 4.371 | 2.51E-05 | 0.011 |
| | | 38,082,778 | | *ORMDL3* | 4.187 | 5.18E-05 | 0.011 |

| | | | | | | | |
|---|---|---|---|---|---|---|---|
| | | 38,108,710 | | | -4.589 | 1.03E-05 | 0.039 |
| | | 38,119,198 | *GSDMA* | | 4.235 | 4.30E-05 | 0.009 |
| | | 38,119,207 | *GSDMA* | | 5.357 | 3.72E-07 | 0.009 |
| | | 38,119,649 | | *GSDMA* | 4.398 | 2.26E-05 | 0.002 |
| rs9903250 | g.38031030.A>G | 37,927,002 | | *IKZF3* | -3.588 | 4.70E-04 | 0.001 |
| | | 38,018,804 | | *IKZF3* | 4.291 | 3.45E-05 | 0.046 |
| | | 38,049,272 | | | 3.531 | 5.74E-04 | 0.046 |
| | | 38,077,191 | | | 4.101 | 7.22E-05 | 0.034 |
| | | 38,080,829 | | *ORMDL3* | 5.326 | 4.28E-07 | 0.014 |
| | | 38,081,812 | | *ORMDL3* | 3.387 | 9.35E-04 | 2.01E-04 |
| | | 38,081,995 | | *ORMDL3* | 4.489 | 1.56E-05 | 2.01E-04 |
| | | 38,082,022 | | *ORMDL3* | 5.027 | 1.62E-06 | 6.40E-04 |
| | | 38,082,035 | | *ORMDL3* | 6.122 | 1.02E-08 | 6.40E-04 |
| | | 38,082,073 | | *ORMDL3* | 5.874 | 3.36E-08 | 6.40E-04 |
| | | 38,082,235 | | *ORMDL3* | 3.957 | 1.24E-04 | 0.004 |
| | | 38,082,346 | | *ORMDL3* | 4.643 | 8.28E-06 | 0.011 |
| | | 38,082,448 | | *ORMDL3* | 4.146 | 6.05E-05 | 0.011 |
| | | 38,082,658 | | *ORMDL3* | 4.337 | 2.87E-05 | 0.011 |
| | | 38,082,778 | | *ORMDL3* | 3.671 | 3.51E-04 | 0.011 |
| | | 38,108,710 | | | -4.040 | 9.10E-05 | 0.039 |
| | | 38,119,198 | *GSDMA* | | 4.199 | 4.94E-05 | 0.009 |
| | | 38,119,207 | *GSDMA* | | 5.693 | 7.91E-08 | 0.009 |
| | | 38,119,649 | | *GSDMA* | 4.145 | 6.08E-05 | 0.002 |
| rs112389508 | g.38031067.A>G | 37,856,732 | | *ERBB2* | -3.668 | 3.56E-04 | 0.009 |
| | | 37,886,749 | | *MIEN1* | 3.945 | 1.30E-04 | 0.002 |
| | | 37,896,063 | | | 3.476 | 6.92E-04 | 0.009 |
| | | 38,020,485 | *IKZF3* | | 3.962 | 1.22E-04 | 0.005 |
| | | 38,080,829 | | *ORMDL3* | 3.995 | 1.08E-04 | 0.014 |
| | | 38,096,333 | | | -3.664 | 3.60E-04 | 0.033 |
| | | 38,096,571 | | | -4.780 | 4.67E-06 | 0.033 |
| | | 38,096,578 | | | -5.642 | 1.00E-07 | 0.033 |
| | | 38,096,648 | | | -3.887 | 1.61E-04 | 0.033 |
| rs9905959 | g.38031138.G>A | 37,828,296 | | *PGAP3* | 3.739 | 2.75E-04 | 0.002 |
| | | 37,927,002 | | *IKZF3* | -3.528 | 5.80E-04 | 0.001 |
| | | 38,018,804 | | *IKZF3* | 3.616 | 4.26E-04 | 0.046 |
| | | 38,077,191 | | | 4.332 | 2.93E-05 | 0.034 |
| | | 38,080,829 | | *ORMDL3* | 3.435 | 7.95E-04 | 0.014 |
| | | 38,081,812 | | *ORMDL3* | 3.459 | 7.34E-04 | 2.01E-04 |
| | | 38,081,829 | | *ORMDL3* | 3.730 | 2.85E-04 | 2.01E-04 |
| | | 38,081,995 | | *ORMDL3* | 5.842 | 3.92E-08 | 2.01E-04 |
| | | 38,082,022 | | *ORMDL3* | 6.694 | 5.90E-10 | 6.40E-04 |

| | | | | | | | |
|---|---|---|---|---|---|---|---|
| | | 38,082,035 | | *ORMDL3* | 6.772 | 3.96E-10 | 6.40E-04 |
| | | 38,082,073 | | *ORMDL3* | 6.832 | 2.91E-10 | 6.40E-04 |
| | | 38,082,206 | | *ORMDL3* | 3.847 | 1.86E-04 | 0.004 |
| | | 38,082,235 | | *ORMDL3* | 4.278 | 3.62E-05 | 0.004 |
| | | 38,082,346 | | *ORMDL3* | 4.590 | 1.03E-05 | 0.011 |
| | | 38,082,448 | | *ORMDL3* | 3.644 | 3.87E-04 | 0.011 |
| | | 38,082,658 | | *ORMDL3* | 4.266 | 3.79E-05 | 0.011 |
| | | 38,082,778 | | *ORMDL3* | 4.147 | 6.05E-05 | 0.011 |
| | | 38,108,710 | | | -4.669 | 7.42E-06 | 0.039 |
| | | 38,119,198 | *GSDMA* | | 4.246 | 4.11E-05 | 0.009 |
| | | 38,119,207 | *GSDMA* | | 5.331 | 4.19E-07 | 0.009 |
| | | 38,119,649 | | *GSDMA* | 4.404 | 2.19E-05 | 0.002 |
| rs11658278 | g.38031164.C>T | 37,927,002 | | *IKZF3* | -3.588 | 4.70E-04 | 0.001 |
| | | 38,018,804 | | *IKZF3* | 4.291 | 3.45E-05 | 0.046 |
| | | 38,049,272 | | | 3.531 | 5.74E-04 | 0.046 |
| | | 38,077,191 | | | 4.101 | 7.22E-05 | 0.034 |
| | | 38,080,829 | | *ORMDL3* | 5.326 | 4.28E-07 | 0.014 |
| | | 38,081,812 | | *ORMDL3* | 3.387 | 9.35E-04 | 2.01E-04 |
| | | 38,081,995 | | *ORMDL3* | 4.489 | 1.56E-05 | 2.01E-04 |
| | | 38,082,022 | | *ORMDL3* | 5.027 | 1.62E-06 | 6.40E-04 |
| | | 38,082,035 | | *ORMDL3* | 6.122 | 1.02E-08 | 6.40E-04 |
| | | 38,082,073 | | *ORMDL3* | 5.874 | 3.36E-08 | 6.40E-04 |
| | | 38,082,235 | | *ORMDL3* | 3.957 | 1.24E-04 | 0.004 |
| | | 38,082,346 | | *ORMDL3* | 4.643 | 8.28E-06 | 0.011 |
| | | 38,082,448 | | *ORMDL3* | 4.146 | 6.05E-05 | 0.011 |
| | | 38,082,658 | | *ORMDL3* | 4.337 | 2.87E-05 | 0.011 |
| | | 38,082,778 | | *ORMDL3* | 3.671 | 3.51E-04 | 0.011 |
| | | 38,108,710 | | | -4.040 | 9.10E-05 | 0.039 |
| | | 38,119,198 | *GSDMA* | | 4.199 | 4.94E-05 | 0.009 |
| | | 38,119,207 | *GSDMA* | | 5.693 | 7.91E-08 | 0.009 |
| | | 38,119,649 | | *GSDMA* | 4.145 | 6.08E-05 | 0.002 |
| rs111396994 | g.38031325.G>A | 37,856,732 | | *ERBB2* | -3.668 | 3.56E-04 | 0.009 |
| | | 37,886,749 | | *MIEN1* | 3.945 | 1.30E-04 | 0.002 |
| | | 37,896,063 | | | 3.476 | 6.92E-04 | 0.009 |
| | | 38,020,485 | *IKZF3* | | 3.962 | 1.22E-04 | 0.005 |
| | | 38,080,829 | | *ORMDL3* | 3.995 | 1.08E-04 | 0.014 |
| | | 38,096,333 | | | -3.664 | 3.60E-04 | 0.033 |
| | | 38,096,571 | | | -4.780 | 4.67E-06 | 0.033 |
| | | 38,096,578 | | | -5.642 | 1.00E-07 | 0.033 |
| | | 38,096,648 | | | -3.887 | 1.61E-04 | 0.033 |

| | | | | | | | |
|---|---|---|---|---|---|---|---|
| rs10852935 | g.38031674.T>C (Cys.292.Cys) | 37,828,296 | | *PGAP3* | 3.739 | 2.75E-04 | 0.002 |
| | | 37,927,002 | | *IKZF3* | -3.528 | 5.80E-04 | 0.001 |
| | | 38,018,804 | | *IKZF3* | 3.616 | 4.26E-04 | 0.046 |
| | | 38,077,191 | | | 4.332 | 2.93E-05 | 0.034 |
| | | 38,080,829 | | *ORMDL3* | 3.435 | 7.95E-04 | 0.014 |
| | | 38,081,812 | | *ORMDL3* | 3.459 | 7.34E-04 | 2.01E-04 |
| | | 38,081,829 | | *ORMDL3* | 3.730 | 2.85E-04 | 2.01E-04 |
| | | 38,081,995 | | *ORMDL3* | 5.842 | 3.92E-08 | 2.01E-04 |
| | | 38,082,022 | | *ORMDL3* | 6.694 | 5.90E-10 | 6.40E-04 |
| | | 38,082,035 | | *ORMDL3* | 6.772 | 3.96E-10 | 6.40E-04 |
| | | 38,082,073 | | *ORMDL3* | 6.832 | 2.91E-10 | 6.40E-04 |
| | | 38,082,206 | | *ORMDL3* | 3.847 | 1.86E-04 | 0.004 |
| | | 38,082,235 | | *ORMDL3* | 4.278 | 3.62E-05 | 0.004 |
| | | 38,082,346 | | *ORMDL3* | 4.590 | 1.03E-05 | 0.011 |
| | | 38,082,448 | | *ORMDL3* | 3.644 | 3.87E-04 | 0.011 |
| | | 38,082,658 | | *ORMDL3* | 4.266 | 3.79E-05 | 0.011 |
| | | 38,082,778 | | *ORMDL3* | 4.147 | 6.05E-05 | 0.011 |
| | | 38,108,710 | | | -4.669 | 7.42E-06 | 0.039 |
| | | 38,119,198 | *GSDMA* | | 4.246 | 4.11E-05 | 0.009 |
| | | 38,119,207 | *GSDMA* | | 5.331 | 4.19E-07 | 0.009 |
| | | 38,119,649 | | *GSDMA* | 4.404 | 2.19E-05 | 0.002 |
| rs10852936 | g.38031714.T>C | 37,828,296 | | *PGAP3* | 3.807 | 2.16E-04 | 0.002 |
| | | 37,927,002 | | *IKZF3* | -3.510 | 6.17E-04 | 0.001 |
| | | 38,018,804 | | *IKZF3* | 3.610 | 4.36E-04 | 0.046 |
| | | 38,072,514 | | *GSDMB* | 3.397 | 9.05E-04 | 0.046 |
| | | 38,077,191 | | | 4.357 | 2.65E-05 | 0.034 |
| | | 38,081,812 | | *ORMDL3* | 3.430 | 8.11E-04 | 2.01E-04 |
| | | 38,081,829 | | *ORMDL3* | 3.718 | 2.98E-04 | 2.01E-04 |
| | | 38,081,995 | | *ORMDL3* | 5.849 | 3.78E-08 | 2.01E-04 |
| | | 38,082,022 | | *ORMDL3* | 6.803 | 3.37E-10 | 6.40E-04 |
| | | 38,082,035 | | *ORMDL3* | 6.894 | 2.12E-10 | 6.40E-04 |
| | | 38,082,073 | | *ORMDL3* | 6.816 | 3.16E-10 | 6.40E-04 |
| | | 38,082,206 | | *ORMDL3* | 3.907 | 1.50E-04 | 0.004 |
| | | 38,082,235 | | *ORMDL3* | 4.336 | 2.88E-05 | 0.004 |
| | | 38,082,346 | | *ORMDL3* | 4.603 | 9.80E-06 | 0.011 |
| | | 38,082,448 | | *ORMDL3* | 3.621 | 4.19E-04 | 0.011 |
| | | 38,082,658 | | *ORMDL3* | 4.298 | 3.35E-05 | 0.011 |
| | | 38,082,778 | | *ORMDL3* | 4.232 | 4.35E-05 | 0.011 |
| | | 38,108,710 | | | -4.538 | 1.28E-05 | 0.039 |
| | | 38,119,198 | *GSDMA* | | 4.169 | 5.55E-05 | 0.009 |

| | | | | | | | |
|---|---|---|---|---|---|---|---|
| | | 38,119,207 | *GSDMA* | | 5.247 | 6.11E-07 | 0.009 |
| | | 38,119,649 | | *GSDMA* | 4.644 | 8.24E-06 | 0.002 |
| rs9891174 | g.38031802.A>T | 37,828,296 | | *PGAP3* | 3.739 | 2.75E-04 | 0.002 |
| | | 37,927,002 | | *IKZF3* | -3.528 | 5.80E-04 | 0.001 |
| | | 38,018,804 | | *IKZF3* | 3.616 | 4.26E-04 | 0.046 |
| | | 38,077,191 | | | 4.332 | 2.93E-05 | 0.034 |
| | | 38,080,829 | | *ORMDL3* | 3.435 | 7.95E-04 | 0.014 |
| | | 38,081,812 | | *ORMDL3* | 3.459 | 7.34E-04 | 2.01E-04 |
| | | 38,081,829 | | *ORMDL3* | 3.730 | 2.85E-04 | 2.01E-04 |
| | | 38,081,995 | | *ORMDL3* | 5.842 | 3.92E-08 | 2.01E-04 |
| | | 38,082,022 | | *ORMDL3* | 6.694 | 5.90E-10 | 6.40E-04 |
| | | 38,082,035 | | *ORMDL3* | 6.772 | 3.96E-10 | 6.40E-04 |
| | | 38,082,073 | | *ORMDL3* | 6.832 | 2.91E-10 | 6.40E-04 |
| | | 38,082,206 | | *ORMDL3* | 3.847 | 1.86E-04 | 0.004 |
| | | 38,082,235 | | *ORMDL3* | 4.278 | 3.62E-05 | 0.004 |
| | | 38,082,346 | | *ORMDL3* | 4.590 | 1.03E-05 | 0.011 |
| | | 38,082,448 | | *ORMDL3* | 3.644 | 3.87E-04 | 0.011 |
| | | 38,082,658 | | *ORMDL3* | 4.266 | 3.79E-05 | 0.011 |
| | | 38,082,778 | | *ORMDL3* | 4.147 | 6.05E-05 | 0.011 |
| | | 38,108,710 | | | -4.669 | 7.42E-06 | 0.039 |
| | | 38,119,198 | *GSDMA* | | 4.246 | 4.11E-05 | 0.009 |
| | | 38,119,207 | *GSDMA* | | 5.331 | 4.19E-07 | 0.009 |
| | | 38,119,649 | | *GSDMA* | 4.404 | 2.19E-05 | 0.002 |
| rs781123329 | g.38031846_38031848del | 37,856,732 | | *ERBB2* | -3.668 | 3.56E-04 | 0.009 |
| | | 37,886,749 | | *MIEN1* | 3.945 | 1.30E-04 | 0.002 |
| | | 37,896,063 | | | 3.476 | 6.92E-04 | 0.009 |
| | | 38,020,485 | *IKZF3* | | 3.962 | 1.22E-04 | 0.005 |
| | | 38,080,829 | | *ORMDL3* | 3.995 | 1.08E-04 | 0.014 |
| | | 38,096,333 | | | -3.664 | 3.60E-04 | 0.033 |
| | | 38,096,571 | | | -4.780 | 4.67E-06 | 0.033 |
| | | 38,096,578 | | | -5.642 | 1.00E-07 | 0.033 |
| | | 38,096,648 | | | -3.887 | 1.61E-04 | 0.033 |
| rs59716545 | g.38031857.G>T | 37,828,296 | | *PGAP3* | 3.484 | 6.73E-04 | 0.002 |
| | | 38,072,514 | | *GSDMB* | 3.574 | 4.94E-04 | 0.046 |
| | | 38,077,191 | | | 4.278 | 3.63E-05 | 0.034 |
| | | 38,080,829 | | *ORMDL3* | 3.523 | 5.90E-04 | 0.014 |
| | | 38,081,812 | | *ORMDL3* | 3.547 | 5.43E-04 | 2.01E-04 |
| | | 38,081,829 | | *ORMDL3* | 3.484 | 6.73E-04 | 2.01E-04 |
| | | 38,081,995 | | *ORMDL3* | 5.522 | 1.75E-07 | 2.01E-04 |
| | | 38,082,022 | | *ORMDL3* | 6.461 | 1.90E-09 | 6.40E-04 |

| | | | | | | | | |
|---|---|---|---|---|---|---|---|---|
| | | 38,082,035 | | *ORMDL3* | 6.455 | 1.96E-09 | | 6.40E-04 |
| | | 38,082,073 | | *ORMDL3* | 6.478 | 1.75E-09 | | 6.40E-04 |
| | | 38,082,206 | | *ORMDL3* | 4.075 | 7.97E-05 | | 0.004 |
| | | 38,082,235 | | *ORMDL3* | 4.348 | 2.74E-05 | | 0.004 |
| | | 38,082,346 | | *ORMDL3* | 5.032 | 1.58E-06 | | 0.011 |
| | | 38,082,448 | | *ORMDL3* | 3.673 | 3.49E-04 | | 0.011 |
| | | 38,082,658 | | *ORMDL3* | 4.374 | 2.48E-05 | | 0.011 |
| | | 38,082,778 | | *ORMDL3* | 3.570 | 5.02E-04 | | 0.011 |
| | | 38,108,710 | | | -4.169 | 5.54E-05 | | 0.039 |
| | | 38,119,198 | *GSDMA* | | 4.022 | 9.75E-05 | | 0.009 |
| | | 38,119,207 | *GSDMA* | | 4.998 | 1.84E-06 | | 0.009 |
| | | 38,119,649 | | *GSDMA* | 4.099 | 7.27E-05 | | 0.002 |
| rs36095411 | g.38031865.G>T | 37,828,296 | | *PGAP3* | 3.703 | 3.14E-04 | 7.00E-04 | 0.002 |
| | | 37,927,002 | | *IKZF3* | -3.454 | 7.46E-04 | 7.00E-04 | 0.001 |
| | | 38,018,804 | | *IKZF3* | 3.629 | 4.08E-04 | 7.00E-04 | 0.046 |
| | | 38,077,191 | | | 4.035 | 9.25E-05 | 7.00E-04 | 0.034 |
| | | 38,080,829 | | *ORMDL3* | 4.116 | 6.80E-05 | 7.00E-04 | 0.014 |
| | | 38,081,812 | | *ORMDL3* | 3.398 | 9.02E-04 | 7.00E-04 | 2.01E-04 |
| | | 38,081,829 | | *ORMDL3* | 3.445 | 7.71E-04 | 7.00E-04 | 2.01E-04 |
| | | 38,081,995 | | *ORMDL3* | 5.085 | 1.25E-06 | 7.00E-04 | 2.01E-04 |
| | | 38,082,022 | | *ORMDL3* | 6.582 | 1.04E-09 | 7.00E-04 | 6.40E-04 |
| | | 38,082,035 | | *ORMDL3* | 6.989 | 1.30E-10 | 7.00E-04 | 6.40E-04 |
| | | 38,082,073 | | *ORMDL3* | 6.298 | 4.29E-09 | 7.00E-04 | 6.40E-04 |
| | | 38,082,206 | | *ORMDL3* | 3.490 | 6.59E-04 | 7.00E-04 | 0.004 |
| | | 38,082,235 | | *ORMDL3* | 4.110 | 6.95E-05 | 7.00E-04 | 0.004 |
| | | 38,082,346 | | *ORMDL3* | 4.880 | 3.05E-06 | 7.00E-04 | 0.011 |
| | | 38,082,448 | | *ORMDL3* | 3.694 | 3.23E-04 | 7.00E-04 | 0.011 |
| | | 38,082,658 | | *ORMDL3* | 4.740 | 5.52E-06 | 7.00E-04 | 0.011 |
| | | 38,082,778 | | *ORMDL3* | 3.900 | 1.53E-04 | 7.00E-04 | 0.011 |
| | | 38,108,710 | | | -4.313 | 3.16E-05 | 7.00E-04 | 0.039 |
| | | 38,119,198 | *GSDMA* | | 4.374 | 2.48E-05 | 7.00E-04 | 0.009 |
| | | 38,119,207 | *GSDMA* | | 5.423 | 2.76E-07 | 7.00E-04 | 0.009 |
| | | 38,119,649 | | *GSDMA* | 4.717 | 6.09E-06 | 7.00E-04 | 0.002 |
| rs200216139 | g.38032132_38032132ins C | 37,828,296 | | *PGAP3* | 3.694 | 3.24E-04 | | 0.002 |
| | | 37,927,002 | | *IKZF3* | -3.513 | 6.10E-04 | | 0.001 |
| | | 38,018,804 | | *IKZF3* | 4.225 | 4.46E-05 | | 0.046 |
| | | 38,049,272 | | | 3.664 | 3.60E-04 | | 0.046 |
| | | 38,077,191 | | | 3.953 | 1.26E-04 | | 0.034 |
| | | 38,080,829 | | *ORMDL3* | 4.847 | 3.51E-06 | | 0.014 |
| | | 38,081,812 | | *ORMDL3* | 3.419 | 8.40E-04 | | 2.01E-04 |

| | | | | | | | |
|---|---|---|---|---|---|---|---|
| | | 38,081,995 | | *ORMDL3* | 4.428 | 2.00E-05 | 2.01E-04 |
| | | 38,082,022 | | *ORMDL3* | 5.300 | 4.81E-07 | 6.40E-04 |
| | | 38,082,035 | | *ORMDL3* | 6.421 | 2.33E-09 | 6.40E-04 |
| | | 38,082,073 | | *ORMDL3* | 5.642 | 1.01E-07 | 6.40E-04 |
| | | 38,082,235 | | *ORMDL3* | 4.043 | 8.99E-05 | 0.004 |
| | | 38,082,346 | | *ORMDL3* | 4.648 | 8.13E-06 | 0.011 |
| | | 38,082,448 | | *ORMDL3* | 3.984 | 1.12E-04 | 0.011 |
| | | 38,082,658 | | *ORMDL3* | 4.053 | 8.66E-05 | 0.011 |
| | | 38,082,778 | | *ORMDL3* | 3.785 | 2.34E-04 | 0.011 |
| | | 38,108,710 | | | -3.686 | 3.33E-04 | 0.039 |
| | | 38,119,198 | *GSDMA* | | 4.214 | 4.67E-05 | 0.009 |
| | | 38,119,207 | *GSDMA* | | 5.718 | 7.03E-08 | 0.009 |
| | | 38,119,649 | | *GSDMA* | 5.012 | 1.73E-06 | 0.002 |
| rs12939457 | g.38032188.C>T | 37,828,296 | | *PGAP3* | 3.758 | 2.57E-04 | 0.002 |
| | | 37,927,002 | | *IKZF3* | -3.565 | 5.10E-04 | 0.001 |
| | | 38,018,804 | | *IKZF3* | 3.591 | 4.65E-04 | 0.046 |
| | | 38,077,191 | | | 4.413 | 2.12E-05 | 0.034 |
| | | 38,080,829 | | *ORMDL3* | 3.542 | 5.52E-04 | 0.014 |
| | | 38,081,812 | | *ORMDL3* | 3.488 | 6.65E-04 | 2.01E-04 |
| | | 38,081,829 | | *ORMDL3* | 3.767 | 2.49E-04 | 2.01E-04 |
| | | 38,081,995 | | *ORMDL3* | 5.840 | 3.95E-08 | 2.01E-04 |
| | | 38,082,022 | | *ORMDL3* | 6.712 | 5.37E-10 | 6.40E-04 |
| | | 38,082,035 | | *ORMDL3* | 6.837 | 2.83E-10 | 6.40E-04 |
| | | 38,082,073 | | *ORMDL3* | 6.850 | 2.65E-10 | 6.40E-04 |
| | | 38,082,206 | | *ORMDL3* | 3.797 | 2.24E-04 | 0.004 |
| | | 38,082,235 | | *ORMDL3* | 4.241 | 4.19E-05 | 0.004 |
| | | 38,082,346 | | *ORMDL3* | 4.594 | 1.02E-05 | 0.011 |
| | | 38,082,448 | | *ORMDL3* | 3.685 | 3.34E-04 | 0.011 |
| | | 38,082,658 | | *ORMDL3* | 4.301 | 3.31E-05 | 0.011 |
| | | 38,082,778 | | *ORMDL3* | 4.178 | 5.36E-05 | 0.011 |
| | | 38,108,710 | | | -4.684 | 7.00E-06 | 0.039 |
| | | 38,119,198 | *GSDMA* | | 4.246 | 4.12E-05 | 0.009 |
| | | 38,119,207 | *GSDMA* | | 5.308 | 4.65E-07 | 0.009 |
| | | 38,119,649 | | *GSDMA* | 4.423 | 2.03E-05 | 0.002 |
| rs367998020 | g.38032200_38032217del | 37,828,296 | | *PGAP3* | 3.758 | 2.57E-04 | 0.002 |
| | | 37,927,002 | | *IKZF3* | -3.565 | 5.10E-04 | 0.001 |
| | | 38,018,804 | | *IKZF3* | 3.591 | 4.65E-04 | 0.046 |
| | | 38,077,191 | | | 4.413 | 2.12E-05 | 0.034 |
| | | 38,080,829 | | *ORMDL3* | 3.542 | 5.52E-04 | 0.014 |
| | | 38,081,812 | | *ORMDL3* | 3.488 | 6.65E-04 | 2.01E-04 |

| | | | | | | | |
|---|---|---|---|---|---|---|---|
| | | 38,081,829 | | *ORMDL3* | 3.767 | 2.49E-04 | 2.01E-04 |
| | | 38,081,995 | | *ORMDL3* | 5.840 | 3.95E-08 | 2.01E-04 |
| | | 38,082,022 | | *ORMDL3* | 6.712 | 5.37E-10 | 6.40E-04 |
| | | 38,082,035 | | *ORMDL3* | 6.837 | 2.83E-10 | 6.40E-04 |
| | | 38,082,073 | | *ORMDL3* | 6.850 | 2.65E-10 | 6.40E-04 |
| | | 38,082,206 | | *ORMDL3* | 3.797 | 2.24E-04 | 0.004 |
| | | 38,082,235 | | *ORMDL3* | 4.241 | 4.19E-05 | 0.004 |
| | | 38,082,346 | | *ORMDL3* | 4.594 | 1.02E-05 | 0.011 |
| | | 38,082,448 | | *ORMDL3* | 3.685 | 3.34E-04 | 0.011 |
| | | 38,082,658 | | *ORMDL3* | 4.301 | 3.31E-05 | 0.011 |
| | | 38,082,778 | | *ORMDL3* | 4.178 | 5.36E-05 | 0.011 |
| | | 38,108,710 | | | -4.684 | 7.00E-06 | 0.039 |
| | | 38,119,198 | *GSDMA* | | 4.246 | 4.12E-05 | 0.009 |
| | | 38,119,207 | *GSDMA* | | 5.308 | 4.65E-07 | 0.009 |
| | | 38,119,649 | | *GSDMA* | 4.423 | 2.03E-05 | 0.002 |
| rs34189114 | g.38032460.T>C | 37,828,296 | | *PGAP3* | 3.739 | 2.75E-04 | 0.002 |
| | | 37,927,002 | | *IKZF3* | -3.528 | 5.80E-04 | 0.001 |
| | | 38,018,804 | | *IKZF3* | 3.616 | 4.26E-04 | 0.046 |
| | | 38,077,191 | | | 4.332 | 2.93E-05 | 0.034 |
| | | 38,080,829 | | *ORMDL3* | 3.435 | 7.95E-04 | 0.014 |
| | | 38,081,812 | | *ORMDL3* | 3.459 | 7.34E-04 | 2.01E-04 |
| | | 38,081,829 | | *ORMDL3* | 3.730 | 2.85E-04 | 2.01E-04 |
| | | 38,081,995 | | *ORMDL3* | 5.842 | 3.92E-08 | 2.01E-04 |
| | | 38,082,022 | | *ORMDL3* | 6.694 | 5.90E-10 | 6.40E-04 |
| | | 38,082,035 | | *ORMDL3* | 6.772 | 3.96E-10 | 6.40E-04 |
| | | 38,082,073 | | *ORMDL3* | 6.832 | 2.91E-10 | 6.40E-04 |
| | | 38,082,206 | | *ORMDL3* | 3.847 | 1.86E-04 | 0.004 |
| | | 38,082,235 | | *ORMDL3* | 4.278 | 3.62E-05 | 0.004 |
| | | 38,082,346 | | *ORMDL3* | 4.590 | 1.03E-05 | 0.011 |
| | | 38,082,448 | | *ORMDL3* | 3.644 | 3.87E-04 | 0.011 |
| | | 38,082,658 | | *ORMDL3* | 4.266 | 3.79E-05 | 0.011 |
| | | 38,082,778 | | *ORMDL3* | 4.147 | 6.05E-05 | 0.011 |
| | | 38,108,710 | | | -4.669 | 7.42E-06 | 0.039 |
| | | 38,119,198 | *GSDMA* | | 4.246 | 4.11E-05 | 0.009 |
| | | 38,119,207 | *GSDMA* | | 5.331 | 4.19E-07 | 0.009 |
| | | 38,119,649 | | *GSDMA* | 4.404 | 2.19E-05 | 0.002 |
| rs35736272 | g.38032680.C>T | 37,828,296 | | *PGAP3* | 3.739 | 2.75E-04 | 0.002 |
| | | 37,927,002 | | *IKZF3* | -3.528 | 5.80E-04 | 0.001 |
| | | 38,018,804 | | *IKZF3* | 3.616 | 4.26E-04 | 0.046 |
| | | 38,077,191 | | | 4.332 | 2.93E-05 | 0.034 |
| | | 38,080,829 | | *ORMDL3* | 3.435 | 7.95E-04 | 0.014 |

| | | | | | | | |
|---|---|---|---|---|---|---|---|
| | | 38,081,812 | | *ORMDL3* | 3.459 | 7.34E-04 | 2.01E-04 |
| | | 38,081,829 | | *ORMDL3* | 3.730 | 2.85E-04 | 2.01E-04 |
| | | 38,081,995 | | *ORMDL3* | 5.842 | 3.92E-08 | 2.01E-04 |
| | | 38,082,022 | | *ORMDL3* | 6.694 | 5.90E-10 | 6.40E-04 |
| | | 38,082,035 | | *ORMDL3* | 6.772 | 3.96E-10 | 6.40E-04 |
| | | 38,082,073 | | *ORMDL3* | 6.832 | 2.91E-10 | 6.40E-04 |
| | | 38,082,206 | | *ORMDL3* | 3.847 | 1.86E-04 | 0.004 |
| | | 38,082,235 | | *ORMDL3* | 4.278 | 3.62E-05 | 0.004 |
| | | 38,082,346 | | *ORMDL3* | 4.590 | 1.03E-05 | 0.011 |
| | | 38,082,448 | | *ORMDL3* | 3.644 | 3.87E-04 | 0.011 |
| | | 38,082,658 | | *ORMDL3* | 4.266 | 3.79E-05 | 0.011 |
| | | 38,082,778 | | *ORMDL3* | 4.147 | 6.05E-05 | 0.011 |
| | | 38,108,710 | | | -4.669 | 7.42E-06 | 0.039 |
| | | 38,119,198 | *GSDMA* | | 4.246 | 4.11E-05 | 0.009 |
| | | 38,119,207 | *GSDMA* | | 5.331 | 4.19E-07 | 0.009 |
| | | 38,119,649 | | *GSDMA* | 4.404 | 2.19E-05 | 0.002 |
| rs1054609 | g.38033277.C>A | 37,828,296 | | *PGAP3* | 3.739 | 2.75E-04 | 0.002 |
| | | 37,927,002 | | *IKZF3* | -3.528 | 5.80E-04 | 0.001 |
| | | 38,018,804 | | *IKZF3* | 3.616 | 4.26E-04 | 0.046 |
| | | 38,077,191 | | | 4.332 | 2.93E-05 | 0.034 |
| | | 38,080,829 | | *ORMDL3* | 3.435 | 7.95E-04 | 0.014 |
| | | 38,081,812 | | *ORMDL3* | 3.459 | 7.34E-04 | 2.01E-04 |
| | | 38,081,829 | | *ORMDL3* | 3.730 | 2.85E-04 | 2.01E-04 |
| | | 38,081,995 | | *ORMDL3* | 5.842 | 3.92E-08 | 2.01E-04 |
| | | 38,082,022 | | *ORMDL3* | 6.694 | 5.90E-10 | 6.40E-04 |
| | | 38,082,035 | | *ORMDL3* | 6.772 | 3.96E-10 | 6.40E-04 |
| | | 38,082,073 | | *ORMDL3* | 6.832 | 2.91E-10 | 6.40E-04 |
| | | 38,082,206 | | *ORMDL3* | 3.847 | 1.86E-04 | 0.004 |
| | | 38,082,235 | | *ORMDL3* | 4.278 | 3.62E-05 | 0.004 |
| | | 38,082,346 | | *ORMDL3* | 4.590 | 1.03E-05 | 0.011 |
| | | 38,082,448 | | *ORMDL3* | 3.644 | 3.87E-04 | 0.011 |
| | | 38,082,658 | | *ORMDL3* | 4.266 | 3.79E-05 | 0.011 |
| | | 38,082,778 | | *ORMDL3* | 4.147 | 6.05E-05 | 0.011 |
| | | 38,108,710 | | | -4.669 | 7.42E-06 | 0.039 |
| | | 38,119,198 | *GSDMA* | | 4.246 | 4.11E-05 | 0.009 |
| | | 38,119,207 | *GSDMA* | | 5.331 | 4.19E-07 | 0.009 |
| | | 38,119,649 | | *GSDMA* | 4.404 | 2.19E-05 | 0.002 |
| rs117846478 | g.38033447.G>A | 37,856,732 | | *ERBB2* | -3.668 | 3.56E-04 | 0.009 |
| | | 37,886,749 | | *MIEN1* | 3.945 | 1.30E-04 | 0.002 |
| | | 37,896,063 | | | 3.476 | 6.92E-04 | 0.009 |
| | | 38,020,485 | *IKZF3* | | 3.962 | 1.22E-04 | 0.005 |

| | | | | | | | |
|---|---|---|---|---|---|---|---|
| | | 38,080,829 | | *ORMDL3* | 3.995 | 1.08E-04 | 0.014 |
| | | 38,096,333 | | | -3.664 | 3.60E-04 | 0.033 |
| | | 38,096,571 | | | -4.780 | 4.67E-06 | 0.033 |
| | | 38,096,578 | | | -5.642 | 1.00E-07 | 0.033 |
| | | 38,096,648 | | | -3.887 | 1.61E-04 | 0.033 |
| rs75972983 | g.38035050.T>C | 37,856,732 | | *ERBB2* | -3.668 | 3.56E-04 | 0.009 |
| | | 37,886,749 | | *MIEN1* | 3.945 | 1.30E-04 | 0.002 |
| | | 37,896,063 | | | 3.476 | 6.92E-04 | 0.009 |
| | | 38,020,485 | *IKZF3* | | 3.962 | 1.22E-04 | 0.005 |
| | | 38,080,829 | | *ORMDL3* | 3.995 | 1.08E-04 | 0.014 |
| | | 38,096,333 | | | -3.664 | 3.60E-04 | 0.033 |
| | | 38,096,571 | | | -4.780 | 4.67E-06 | 0.033 |
| | | 38,096,578 | | | -5.642 | 1.00E-07 | 0.033 |
| | | 38,096,648 | | | -3.887 | 1.61E-04 | 0.033 |
| rs9907088 | g.38035116.A>G | 37,828,296 | | *PGAP3* | 3.739 | 2.75E-04 | 0.002 |
| | | 37,927,002 | | *IKZF3* | -3.528 | 5.80E-04 | 0.001 |
| | | 38,018,804 | | *IKZF3* | 3.616 | 4.26E-04 | 0.046 |
| | | 38,077,191 | | | 4.332 | 2.93E-05 | 0.034 |
| | | 38,080,829 | | *ORMDL3* | 3.435 | 7.95E-04 | 0.014 |
| | | 38,081,812 | | *ORMDL3* | 3.459 | 7.34E-04 | 2.01E-04 |
| | | 38,081,829 | | *ORMDL3* | 3.730 | 2.85E-04 | 2.01E-04 |
| | | 38,081,995 | | *ORMDL3* | 5.842 | 3.92E-08 | 2.01E-04 |
| | | 38,082,022 | | *ORMDL3* | 6.694 | 5.90E-10 | 6.40E-04 |
| | | 38,082,035 | | *ORMDL3* | 6.772 | 3.96E-10 | 6.40E-04 |
| | | 38,082,073 | | *ORMDL3* | 6.832 | 2.91E-10 | 6.40E-04 |
| | | 38,082,206 | | *ORMDL3* | 3.847 | 1.86E-04 | 0.004 |
| | | 38,082,235 | | *ORMDL3* | 4.278 | 3.62E-05 | 0.004 |
| | | 38,082,346 | | *ORMDL3* | 4.590 | 1.03E-05 | 0.011 |
| | | 38,082,448 | | *ORMDL3* | 3.644 | 3.87E-04 | 0.011 |
| | | 38,082,658 | | *ORMDL3* | 4.266 | 3.79E-05 | 0.011 |
| | | 38,082,778 | | *ORMDL3* | 4.147 | 6.05E-05 | 0.011 |
| | | 38,108,710 | | | -4.669 | 7.42E-06 | 0.039 |
| | | 38,119,198 | *GSDMA* | | 4.246 | 4.11E-05 | 0.009 |
| | | 38,119,207 | *GSDMA* | | 5.331 | 4.19E-07 | 0.009 |
| | | 38,119,649 | | *GSDMA* | 4.404 | 2.19E-05 | 0.002 |
| rs36038753 | g.38035370.T>G | 37,828,296 | | *PGAP3* | 3.739 | 2.75E-04 | 0.002 |
| | | 37,927,002 | | *IKZF3* | -3.528 | 5.80E-04 | 0.001 |
| | | 38,018,804 | | *IKZF3* | 3.616 | 4.26E-04 | 0.046 |
| | | 38,077,191 | | | 4.332 | 2.93E-05 | 0.034 |
| | | 38,080,829 | | *ORMDL3* | 3.435 | 7.95E-04 | 0.014 |
| | | 38,081,812 | | *ORMDL3* | 3.459 | 7.34E-04 | 2.01E-04 |

| | | | | | | | |
|---|---|---|---|---|---|---|---|
| | | 38,081,829 | | *ORMDL3* | 3.730 | 2.85E-04 | 2.01E-04 |
| | | 38,081,995 | | *ORMDL3* | 5.842 | 3.92E-08 | 2.01E-04 |
| | | 38,082,022 | | *ORMDL3* | 6.694 | 5.90E-10 | 6.40E-04 |
| | | 38,082,035 | | *ORMDL3* | 6.772 | 3.96E-10 | 6.40E-04 |
| | | 38,082,073 | | *ORMDL3* | 6.832 | 2.91E-10 | 6.40E-04 |
| | | 38,082,206 | | *ORMDL3* | 3.847 | 1.86E-04 | 0.004 |
| | | 38,082,235 | | *ORMDL3* | 4.278 | 3.62E-05 | 0.004 |
| | | 38,082,346 | | *ORMDL3* | 4.590 | 1.03E-05 | 0.011 |
| | | 38,082,448 | | *ORMDL3* | 3.644 | 3.87E-04 | 0.011 |
| | | 38,082,658 | | *ORMDL3* | 4.266 | 3.79E-05 | 0.011 |
| | | 38,082,778 | | *ORMDL3* | 4.147 | 6.05E-05 | 0.011 |
| | | 38,108,710 | | | -4.669 | 7.42E-06 | 0.039 |
| | | 38,119,198 | *GSDMA* | | 4.246 | 4.11E-05 | 0.009 |
| | | 38,119,207 | *GSDMA* | | 5.331 | 4.19E-07 | 0.009 |
| | | 38,119,649 | | *GSDMA* | 4.404 | 2.19E-05 | 0.002 |
| rs112750506 | g.38035410.G>A | 37,856,732 | | *ERBB2* | -3.668 | 3.56E-04 | 0.009 |
| | | 37,886,749 | | *MIEN1* | 3.945 | 1.30E-04 | 0.002 |
| | | 37,896,063 | | | 3.476 | 6.92E-04 | 0.009 |
| | | 38,020,485 | *IKZF3* | | 3.962 | 1.22E-04 | 0.005 |
| | | 38,080,829 | | *ORMDL3* | 3.995 | 1.08E-04 | 0.014 |
| | | 38,096,333 | | | -3.664 | 3.60E-04 | 0.033 |
| | | 38,096,571 | | | -4.780 | 4.67E-06 | 0.033 |
| | | 38,096,578 | | | -5.642 | 1.00E-07 | 0.033 |
| | | 38,096,648 | | | -3.887 | 1.61E-04 | 0.033 |
| rs35569035 | g.38035624.T>C | 37,828,296 | | *PGAP3* | 3.672 | 3.50E-04 | 0.002 |
| | | 37,927,002 | | *IKZF3* | -3.670 | 3.52E-04 | 0.001 |
| | | 38,018,804 | | *IKZF3* | 3.625 | 4.13E-04 | 0.046 |
| | | 38,077,191 | | | 4.354 | 2.69E-05 | 0.034 |
| | | 38,081,829 | | *ORMDL3* | 3.578 | 4.87E-04 | 2.01E-04 |
| | | 38,081,995 | | *ORMDL3* | 5.804 | 4.69E-08 | 2.01E-04 |
| | | 38,082,022 | | *ORMDL3* | 6.609 | 9.07E-10 | 6.40E-04 |
| | | 38,082,035 | | *ORMDL3* | 6.865 | 2.46E-10 | 6.40E-04 |
| | | 38,082,073 | | *ORMDL3* | 6.788 | 3.65E-10 | 6.40E-04 |
| | | 38,082,206 | | *ORMDL3* | 4.060 | 8.41E-05 | 0.004 |
| | | 38,082,235 | | *ORMDL3* | 4.322 | 3.05E-05 | 0.004 |
| | | 38,082,346 | | *ORMDL3* | 4.864 | 3.26E-06 | 0.011 |
| | | 38,082,448 | | *ORMDL3* | 3.644 | 3.87E-04 | 0.011 |
| | | 38,082,658 | | *ORMDL3* | 4.227 | 4.43E-05 | 0.011 |
| | | 38,082,778 | | *ORMDL3* | 4.406 | 2.18E-05 | 0.011 |
| | | 38,108,710 | | | -4.665 | 7.57E-06 | 0.039 |
| | | 38,119,198 | *GSDMA* | | 3.992 | 1.09E-04 | 0.009 |

| | | | | | | | |
|---|---|---|---|---|---|---|---|
| | | 38,119,207 | *GSDMA* | | 5.113 | 1.11E-06 | 0.009 |
| | | 38,119,649 | | *GSDMA* | 4.237 | 4.26E-05 | 0.002 |
| rs9910826 | g.38035648.G>A | 37,828,296 | | *PGAP3* | 3.672 | 3.50E-04 | 0.002 |
| | | 37,927,002 | | *IKZF3* | -3.670 | 3.52E-04 | 0.001 |
| | | 38,018,804 | | *IKZF3* | 3.625 | 4.13E-04 | 0.046 |
| | | 38,077,191 | | | 4.354 | 2.69E-05 | 0.034 |
| | | 38,081,829 | | *ORMDL3* | 3.578 | 4.87E-04 | 2.01E-04 |
| | | 38,081,995 | | *ORMDL3* | 5.804 | 4.69E-08 | 2.01E-04 |
| | | 38,082,022 | | *ORMDL3* | 6.609 | 9.07E-10 | 6.40E-04 |
| | | 38,082,035 | | *ORMDL3* | 6.865 | 2.46E-10 | 6.40E-04 |
| | | 38,082,073 | | *ORMDL3* | 6.788 | 3.65E-10 | 6.40E-04 |
| | | 38,082,206 | | *ORMDL3* | 4.060 | 8.41E-05 | 0.004 |
| | | 38,082,235 | | *ORMDL3* | 4.322 | 3.05E-05 | 0.004 |
| | | 38,082,346 | | *ORMDL3* | 4.864 | 3.26E-06 | 0.011 |
| | | 38,082,448 | | *ORMDL3* | 3.644 | 3.87E-04 | 0.011 |
| | | 38,082,658 | | *ORMDL3* | 4.227 | 4.43E-05 | 0.011 |
| | | 38,082,778 | | *ORMDL3* | 4.406 | 2.18E-05 | 0.011 |
| | | 38,108,710 | | | -4.665 | 7.57E-06 | 0.039 |
| | | 38,119,198 | *GSDMA* | | 3.992 | 1.09E-04 | 0.009 |
| | | 38,119,207 | *GSDMA* | | 5.113 | 1.11E-06 | 0.009 |
| | | 38,119,649 | | *GSDMA* | 4.237 | 4.26E-05 | 0.002 |
| rs199882592 | g.38035706_38035707del | 37,856,732 | | *ERBB2* | -3.668 | 3.56E-04 | 0.009 |
| | | 37,886,749 | | *MIEN1* | 3.945 | 1.30E-04 | 0.002 |
| | | 37,896,063 | | | 3.476 | 6.92E-04 | 0.009 |
| | | 38,020,485 | *IKZF3* | | 3.962 | 1.22E-04 | 0.005 |
| | | 38,080,829 | | *ORMDL3* | 3.995 | 1.08E-04 | 0.014 |
| | | 38,096,333 | | | -3.664 | 3.60E-04 | 0.033 |
| | | 38,096,571 | | | -4.780 | 4.67E-06 | 0.033 |
| | | 38,096,578 | | | -5.642 | 1.00E-07 | 0.033 |
| | | 38,096,648 | | | -3.887 | 1.61E-04 | 0.033 |
| rs34074973 | g.38035766_38035769del | 37,828,296 | | *PGAP3* | 3.739 | 2.75E-04 | 0.002 |
| | | 37,927,002 | | *IKZF3* | -3.528 | 5.80E-04 | 0.001 |
| | | 38,018,804 | | *IKZF3* | 3.616 | 4.26E-04 | 0.046 |
| | | 38,077,191 | | | 4.332 | 2.93E-05 | 0.034 |
| | | 38,080,829 | | *ORMDL3* | 3.435 | 7.95E-04 | 0.014 |
| | | 38,081,812 | | *ORMDL3* | 3.459 | 7.34E-04 | 2.01E-04 |
| | | 38,081,829 | | *ORMDL3* | 3.730 | 2.85E-04 | 2.01E-04 |
| | | 38,081,995 | | *ORMDL3* | 5.842 | 3.92E-08 | 2.01E-04 |
| | | 38,082,022 | | *ORMDL3* | 6.694 | 5.90E-10 | 6.40E-04 |

| | | | | | | | |
|---|---|---|---|---|---|---|---|
| | | 38,082,035 | | *ORMDL3* | 6.772 | 3.96E-10 | 6.40E-04 |
| | | 38,082,073 | | *ORMDL3* | 6.832 | 2.91E-10 | 6.40E-04 |
| | | 38,082,206 | | *ORMDL3* | 3.847 | 1.86E-04 | 0.004 |
| | | 38,082,235 | | *ORMDL3* | 4.278 | 3.62E-05 | 0.004 |
| | | 38,082,346 | | *ORMDL3* | 4.590 | 1.03E-05 | 0.011 |
| | | 38,082,448 | | *ORMDL3* | 3.644 | 3.87E-04 | 0.011 |
| | | 38,082,658 | | *ORMDL3* | 4.266 | 3.79E-05 | 0.011 |
| | | 38,082,778 | | *ORMDL3* | 4.147 | 6.05E-05 | 0.011 |
| | | 38,108,710 | | | -4.669 | 7.42E-06 | 0.039 |
| | | 38,119,198 | *GSDMA* | | 4.246 | 4.11E-05 | 0.009 |
| | | 38,119,207 | *GSDMA* | | 5.331 | 4.19E-07 | 0.009 |
| | | 38,119,649 | | *GSDMA* | 4.404 | 2.19E-05 | 0.002 |
| rs9904624 | g.38036586.G>A | 37,828,296 | | *PGAP3* | 3.739 | 2.75E-04 | 0.002 |
| | | 37,927,002 | | *IKZF3* | -3.528 | 5.80E-04 | 0.001 |
| | | 38,018,804 | | *IKZF3* | 3.616 | 4.26E-04 | 0.046 |
| | | 38,077,191 | | | 4.332 | 2.93E-05 | 0.034 |
| | | 38,080,829 | | *ORMDL3* | 3.435 | 7.95E-04 | 0.014 |
| | | 38,081,812 | | *ORMDL3* | 3.459 | 7.34E-04 | 2.01E-04 |
| | | 38,081,829 | | *ORMDL3* | 3.730 | 2.85E-04 | 2.01E-04 |
| | | 38,081,995 | | *ORMDL3* | 5.842 | 3.92E-08 | 2.01E-04 |
| | | 38,082,022 | | *ORMDL3* | 6.694 | 5.90E-10 | 6.40E-04 |
| | | 38,082,035 | | *ORMDL3* | 6.772 | 3.96E-10 | 6.40E-04 |
| | | 38,082,073 | | *ORMDL3* | 6.832 | 2.91E-10 | 6.40E-04 |
| | | 38,082,206 | | *ORMDL3* | 3.847 | 1.86E-04 | 0.004 |
| | | 38,082,235 | | *ORMDL3* | 4.278 | 3.62E-05 | 0.004 |
| | | 38,082,346 | | *ORMDL3* | 4.590 | 1.03E-05 | 0.011 |
| | | 38,082,448 | | *ORMDL3* | 3.644 | 3.87E-04 | 0.011 |
| | | 38,082,658 | | *ORMDL3* | 4.266 | 3.79E-05 | 0.011 |
| | | 38,082,778 | | *ORMDL3* | 4.147 | 6.05E-05 | 0.011 |
| | | 38,108,710 | | | -4.669 | 7.42E-06 | 0.039 |
| | | 38,119,198 | *GSDMA* | | 4.246 | 4.11E-05 | 0.009 |
| | | 38,119,207 | *GSDMA* | | 5.331 | 4.19E-07 | 0.009 |
| | | 38,119,649 | | *GSDMA* | 4.404 | 2.19E-05 | 0.002 |
| rs112750995 | g.38037559.T>A | 37,856,732 | | *ERBB2* | -3.668 | 3.56E-04 | 0.009 |
| | | 37,886,749 | | *MIEN1* | 3.945 | 1.30E-04 | 0.002 |
| | | 37,896,063 | | | 3.476 | 6.92E-04 | 0.009 |
| | | 38,020,485 | *IKZF3* | | 3.962 | 1.22E-04 | 0.005 |
| | | 38,080,829 | | *ORMDL3* | 3.995 | 1.08E-04 | 0.014 |
| | | 38,096,333 | | | -3.664 | 3.60E-04 | 0.033 |
| | | 38,096,571 | | | -4.780 | 4.67E-06 | 0.033 |
| | | 38,096,578 | | | -5.642 | 1.00E-07 | 0.033 |

| | | | | | | | |
|---|---|---|---|---|---|---|---|
| | | 38,096,648 | | | -3.887 | 1.61E-04 | 0.033 |
| rs74978817 | g.38037989.G>A | 37,856,732 | | *ERBB2* | -3.668 | 3.56E-04 | 0.009 |
| | | 37,886,749 | | *MIEN1* | 3.945 | 1.30E-04 | 0.002 |
| | | 37,896,063 | | | 3.476 | 6.92E-04 | 0.009 |
| | | 38,020,485 | *IKZF3* | | 3.962 | 1.22E-04 | 0.005 |
| | | 38,080,829 | | *ORMDL3* | 3.995 | 1.08E-04 | 0.014 |
| | | 38,096,333 | | | -3.664 | 3.60E-04 | 0.033 |
| | | 38,096,571 | | | -4.780 | 4.67E-06 | 0.033 |
| | | 38,096,578 | | | -5.642 | 1.00E-07 | 0.033 |
| | | 38,096,648 | | | -3.887 | 1.61E-04 | 0.033 |
| rs4795398 | g.38038179.T>C | 37,828,296 | | *PGAP3* | 3.739 | 2.75E-04 | 0.002 |
| | | 37,927,002 | | *IKZF3* | -3.528 | 5.80E-04 | 0.001 |
| | | 38,018,804 | | *IKZF3* | 3.616 | 4.26E-04 | 0.046 |
| | | 38,077,191 | | | 4.332 | 2.93E-05 | 0.034 |
| | | 38,080,829 | | *ORMDL3* | 3.435 | 7.95E-04 | 0.014 |
| | | 38,081,812 | | *ORMDL3* | 3.459 | 7.34E-04 | 2.01E-04 |
| | | 38,081,829 | | *ORMDL3* | 3.730 | 2.85E-04 | 2.01E-04 |
| | | 38,081,995 | | *ORMDL3* | 5.842 | 3.92E-08 | 2.01E-04 |
| | | 38,082,022 | | *ORMDL3* | 6.694 | 5.90E-10 | 6.40E-04 |
| | | 38,082,035 | | *ORMDL3* | 6.772 | 3.96E-10 | 6.40E-04 |
| | | 38,082,073 | | *ORMDL3* | 6.832 | 2.91E-10 | 6.40E-04 |
| | | 38,082,206 | | *ORMDL3* | 3.847 | 1.86E-04 | 0.004 |
| | | 38,082,235 | | *ORMDL3* | 4.278 | 3.62E-05 | 0.004 |
| | | 38,082,346 | | *ORMDL3* | 4.590 | 1.03E-05 | 0.011 |
| | | 38,082,448 | | *ORMDL3* | 3.644 | 3.87E-04 | 0.011 |
| | | 38,082,658 | | *ORMDL3* | 4.266 | 3.79E-05 | 0.011 |
| | | 38,082,778 | | *ORMDL3* | 4.147 | 6.05E-05 | 0.011 |
| | | 38,108,710 | | | -4.669 | 7.42E-06 | 0.039 |
| | | 38,119,198 | *GSDMA* | | 4.246 | 4.11E-05 | 0.009 |
| | | 38,119,207 | *GSDMA* | | 5.331 | 4.19E-07 | 0.009 |
| | | 38,119,649 | | *GSDMA* | 4.404 | 2.19E-05 | 0.002 |
| rs12939565 | g.38038389.T>A | 37,927,002 | | *IKZF3* | -3.588 | 4.70E-04 | 0.001 |
| | | 38,018,804 | | *IKZF3* | 4.291 | 3.45E-05 | 0.046 |
| | | 38,049,272 | | | 3.531 | 5.74E-04 | 0.046 |
| | | 38,077,191 | | | 4.101 | 7.22E-05 | 0.034 |
| | | 38,080,829 | | *ORMDL3* | 5.326 | 4.28E-07 | 0.014 |
| | | 38,081,812 | | *ORMDL3* | 3.387 | 9.35E-04 | 2.01E-04 |
| | | 38,081,995 | | *ORMDL3* | 4.489 | 1.56E-05 | 2.01E-04 |
| | | 38,082,022 | | *ORMDL3* | 5.027 | 1.62E-06 | 6.40E-04 |
| | | 38,082,035 | | *ORMDL3* | 6.122 | 1.02E-08 | 6.40E-04 |
| | | 38,082,073 | | *ORMDL3* | 5.874 | 3.36E-08 | 6.40E-04 |

| | | | | | | | |
|---|---|---|---|---|---|---|---|
| | | 38,082,235 | | *ORMDL3* | 3.957 | 1.24E-04 | 0.004 |
| | | 38,082,346 | | *ORMDL3* | 4.643 | 8.28E-06 | 0.011 |
| | | 38,082,448 | | *ORMDL3* | 4.146 | 6.05E-05 | 0.011 |
| | | 38,082,658 | | *ORMDL3* | 4.337 | 2.87E-05 | 0.011 |
| | | 38,082,778 | | *ORMDL3* | 3.671 | 3.51E-04 | 0.011 |
| | | 38,108,710 | | | -4.040 | 9.10E-05 | 0.039 |
| | | 38,119,198 | *GSDMA* | | 4.199 | 4.94E-05 | 0.009 |
| | | 38,119,207 | *GSDMA* | | 5.693 | 7.91E-08 | 0.009 |
| | | 38,119,649 | | *GSDMA* | 4.145 | 6.08E-05 | 0.002 |
| rs12939566 | g.38038390.T>A | 37,828,296 | | *PGAP3* | 3.739 | 2.75E-04 | 0.002 |
| | | 37,927,002 | | *IKZF3* | -3.528 | 5.80E-04 | 0.001 |
| | | 38,018,804 | | *IKZF3* | 3.616 | 4.26E-04 | 0.046 |
| | | 38,077,191 | | | 4.332 | 2.93E-05 | 0.034 |
| | | 38,080,829 | | *ORMDL3* | 3.435 | 7.95E-04 | 0.014 |
| | | 38,081,812 | | *ORMDL3* | 3.459 | 7.34E-04 | 2.01E-04 |
| | | 38,081,829 | | *ORMDL3* | 3.730 | 2.85E-04 | 2.01E-04 |
| | | 38,081,995 | | *ORMDL3* | 5.842 | 3.92E-08 | 2.01E-04 |
| | | 38,082,022 | | *ORMDL3* | 6.694 | 5.90E-10 | 6.40E-04 |
| | | 38,082,035 | | *ORMDL3* | 6.772 | 3.96E-10 | 6.40E-04 |
| | | 38,082,073 | | *ORMDL3* | 6.832 | 2.91E-10 | 6.40E-04 |
| | | 38,082,206 | | *ORMDL3* | 3.847 | 1.86E-04 | 0.004 |
| | | 38,082,235 | | *ORMDL3* | 4.278 | 3.62E-05 | 0.004 |
| | | 38,082,346 | | *ORMDL3* | 4.590 | 1.03E-05 | 0.011 |
| | | 38,082,448 | | *ORMDL3* | 3.644 | 3.87E-04 | 0.011 |
| | | 38,082,658 | | *ORMDL3* | 4.266 | 3.79E-05 | 0.011 |
| | | 38,082,778 | | *ORMDL3* | 4.147 | 6.05E-05 | 0.011 |
| | | 38,108,710 | | | -4.669 | 7.42E-06 | 0.039 |
| | | 38,119,198 | *GSDMA* | | 4.246 | 4.11E-05 | 0.009 |
| | | 38,119,207 | *GSDMA* | | 5.331 | 4.19E-07 | 0.009 |
| | | 38,119,649 | | *GSDMA* | 4.404 | 2.19E-05 | 0.002 |
| rs148094956 | g.38039561_38039565del | 37,927,002 | | *IKZF3* | -3.588 | 4.70E-04 | 0.001 |
| | | 38,018,804 | | *IKZF3* | 4.291 | 3.45E-05 | 0.046 |
| | | 38,049,272 | | | 3.531 | 5.74E-04 | 0.046 |
| | | 38,077,191 | | | 4.101 | 7.22E-05 | 0.034 |
| | | 38,080,829 | | *ORMDL3* | 5.326 | 4.28E-07 | 0.014 |
| | | 38,081,812 | | *ORMDL3* | 3.387 | 9.35E-04 | 2.01E-04 |
| | | 38,081,995 | | *ORMDL3* | 4.489 | 1.56E-05 | 2.01E-04 |
| | | 38,082,022 | | *ORMDL3* | 5.027 | 1.62E-06 | 6.40E-04 |
| | | 38,082,035 | | *ORMDL3* | 6.122 | 1.02E-08 | 6.40E-04 |
| | | 38,082,073 | | *ORMDL3* | 5.874 | 3.36E-08 | 6.40E-04 |

| | | | | | | | |
|---|---|---|---|---|---|---|---|
| | | 38,082,235 | | *ORMDL3* | 3.957 | 1.24E-04 | 0.004 |
| | | 38,082,346 | | *ORMDL3* | 4.643 | 8.28E-06 | 0.011 |
| | | 38,082,448 | | *ORMDL3* | 4.146 | 6.05E-05 | 0.011 |
| | | 38,082,658 | | *ORMDL3* | 4.337 | 2.87E-05 | 0.011 |
| | | 38,082,778 | | *ORMDL3* | 3.671 | 3.51E-04 | 0.011 |
| | | 38,108,710 | | | -4.040 | 9.10E-05 | 0.039 |
| | | 38,119,198 | *GSDMA* | | 4.199 | 4.94E-05 | 0.009 |
| | | 38,119,207 | *GSDMA* | | 5.693 | 7.91E-08 | 0.009 |
| | | 38,119,649 | | *GSDMA* | 4.145 | 6.08E-05 | 0.002 |
| rs112569955 | g.38039806.G>A | 37,856,732 | | *ERBB2* | -3.668 | 3.56E-04 | 0.009 |
| | | 37,886,749 | | *MIEN1* | 3.945 | 1.30E-04 | 0.002 |
| | | 37,896,063 | | | 3.476 | 6.92E-04 | 0.009 |
| | | 38,020,485 | *IKZF3* | | 3.962 | 1.22E-04 | 0.005 |
| | | 38,080,829 | | *ORMDL3* | 3.995 | 1.08E-04 | 0.014 |
| | | 38,096,333 | | | -3.664 | 3.60E-04 | 0.033 |
| | | 38,096,571 | | | -4.780 | 4.67E-06 | 0.033 |
| | | 38,096,578 | | | -5.642 | 1.00E-07 | 0.033 |
| | | 38,096,648 | | | -3.887 | 1.61E-04 | 0.033 |
| rs12232497 | g.38040119.C>T | 37,828,296 | | *PGAP3* | 3.739 | 2.75E-04 | 0.002 |
| | | 37,927,002 | | *IKZF3* | -3.528 | 5.80E-04 | 0.001 |
| | | 38,018,804 | | *IKZF3* | 3.616 | 4.26E-04 | 0.046 |
| | | 38,077,191 | | | 4.332 | 2.93E-05 | 0.034 |
| | | 38,080,829 | | *ORMDL3* | 3.435 | 7.95E-04 | 0.014 |
| | | 38,081,812 | | *ORMDL3* | 3.459 | 7.34E-04 | 2.01E-04 |
| | | 38,081,829 | | *ORMDL3* | 3.730 | 2.85E-04 | 2.01E-04 |
| | | 38,081,995 | | *ORMDL3* | 5.842 | 3.92E-08 | 2.01E-04 |
| | | 38,082,022 | | *ORMDL3* | 6.694 | 5.90E-10 | 6.40E-04 |
| | | 38,082,035 | | *ORMDL3* | 6.772 | 3.96E-10 | 6.40E-04 |
| | | 38,082,073 | | *ORMDL3* | 6.832 | 2.91E-10 | 6.40E-04 |
| | | 38,082,206 | | *ORMDL3* | 3.847 | 1.86E-04 | 0.004 |
| | | 38,082,235 | | *ORMDL3* | 4.278 | 3.62E-05 | 0.004 |
| | | 38,082,346 | | *ORMDL3* | 4.590 | 1.03E-05 | 0.011 |
| | | 38,082,448 | | *ORMDL3* | 3.644 | 3.87E-04 | 0.011 |
| | | 38,082,658 | | *ORMDL3* | 4.266 | 3.79E-05 | 0.011 |
| | | 38,082,778 | | *ORMDL3* | 4.147 | 6.05E-05 | 0.011 |
| | | 38,108,710 | | | -4.669 | 7.42E-06 | 0.039 |
| | | 38,119,198 | *GSDMA* | | 4.246 | 4.11E-05 | 0.009 |
| | | 38,119,207 | *GSDMA* | | 5.331 | 4.19E-07 | 0.009 |
| | | 38,119,649 | | *GSDMA* | 4.404 | 2.19E-05 | 0.002 |
| rs112928243 | g.38040193.A>G | 37,856,732 | | *ERBB2* | -3.668 | 3.56E-04 | 0.009 |
| | | 37,886,749 | | *MIEN1* | 3.945 | 1.30E-04 | 0.002 |

| | | | | | | | |
|---|---|---|---|---|---|---|---|
| | | 37,896,063 | | | 3.476 | 6.92E-04 | 0.009 |
| | | 38,020,485 | *IKZF3* | | 3.962 | 1.22E-04 | 0.005 |
| | | 38,080,829 | | *ORMDL3* | 3.995 | 1.08E-04 | 0.014 |
| | | 38,096,333 | | | -3.664 | 3.60E-04 | 0.033 |
| | | 38,096,571 | | | -4.780 | 4.67E-06 | 0.033 |
| | | 38,096,578 | | | -5.642 | 1.00E-07 | 0.033 |
| | | 38,096,648 | | | -3.887 | 1.61E-04 | 0.033 |
| rs12232498 | g.38040363.C>T | 37,828,296 | | *PGAP3* | 3.735 | 2.79E-04 | 0.002 |
| | | 37,927,002 | | *IKZF3* | -3.536 | 5.64E-04 | 0.001 |
| | | 38,018,804 | | *IKZF3* | 3.614 | 4.30E-04 | 0.046 |
| | | 38,077,191 | | | 4.331 | 2.95E-05 | 0.034 |
| | | 38,080,829 | | *ORMDL3* | 3.431 | 8.06E-04 | 0.014 |
| | | 38,081,812 | | *ORMDL3* | 3.450 | 7.55E-04 | 2.01E-04 |
| | | 38,081,829 | | *ORMDL3* | 3.731 | 2.84E-04 | 2.01E-04 |
| | | 38,081,995 | | *ORMDL3* | 5.848 | 3.80E-08 | 2.01E-04 |
| | | 38,082,022 | | *ORMDL3* | 6.702 | 5.67E-10 | 6.40E-04 |
| | | 38,082,035 | | *ORMDL3* | 6.780 | 3.81E-10 | 6.40E-04 |
| | | 38,082,073 | | *ORMDL3* | 6.815 | 3.18E-10 | 6.40E-04 |
| | | 38,082,206 | | *ORMDL3* | 3.848 | 1.86E-04 | 0.004 |
| | | 38,082,235 | | *ORMDL3* | 4.277 | 3.63E-05 | 0.004 |
| | | 38,082,346 | | *ORMDL3* | 4.592 | 1.02E-05 | 0.011 |
| | | 38,082,448 | | *ORMDL3* | 3.647 | 3.83E-04 | 0.011 |
| | | 38,082,658 | | *ORMDL3* | 4.261 | 3.87E-05 | 0.011 |
| | | 38,082,778 | | *ORMDL3* | 4.146 | 6.05E-05 | 0.011 |
| | | 38,108,710 | | | -4.670 | 7.41E-06 | 0.039 |
| | | 38,119,198 | *GSDMA* | | 4.251 | 4.03E-05 | 0.009 |
| | | 38,119,207 | *GSDMA* | | 5.327 | 4.26E-07 | 0.009 |
| | | 38,119,649 | | *GSDMA* | 4.405 | 2.19E-05 | 0.002 |
| rs12941333 | g.38040534.T>C | 37,828,296 | | *PGAP3* | 3.739 | 2.75E-04 | 0.002 |
| | | 37,927,002 | | *IKZF3* | -3.528 | 5.80E-04 | 0.001 |
| | | 38,018,804 | | *IKZF3* | 3.616 | 4.26E-04 | 0.046 |
| | | 38,077,191 | | | 4.332 | 2.93E-05 | 0.034 |
| | | 38,080,829 | | *ORMDL3* | 3.435 | 7.95E-04 | 0.014 |
| | | 38,081,812 | | *ORMDL3* | 3.459 | 7.34E-04 | 2.01E-04 |
| | | 38,081,829 | | *ORMDL3* | 3.730 | 2.85E-04 | 2.01E-04 |
| | | 38,081,995 | | *ORMDL3* | 5.842 | 3.92E-08 | 2.01E-04 |
| | | 38,082,022 | | *ORMDL3* | 6.694 | 5.90E-10 | 6.40E-04 |
| | | 38,082,035 | | *ORMDL3* | 6.772 | 3.96E-10 | 6.40E-04 |
| | | 38,082,073 | | *ORMDL3* | 6.832 | 2.91E-10 | 6.40E-04 |
| | | 38,082,206 | | *ORMDL3* | 3.847 | 1.86E-04 | 0.004 |
| | | 38,082,235 | | *ORMDL3* | 4.278 | 3.62E-05 | 0.004 |

| | | | | | | | |
|---|---|---|---|---|---|---|---|
| | | 38,082,346 | | *ORMDL3* | 4.590 | 1.03E-05 | 0.011 |
| | | 38,082,448 | | *ORMDL3* | 3.644 | 3.87E-04 | 0.011 |
| | | 38,082,658 | | *ORMDL3* | 4.266 | 3.79E-05 | 0.011 |
| | | 38,082,778 | | *ORMDL3* | 4.147 | 6.05E-05 | 0.011 |
| | | 38,108,710 | | | -4.669 | 7.42E-06 | 0.039 |
| | | 38,119,198 | *GSDMA* | | 4.246 | 4.11E-05 | 0.009 |
| | | 38,119,207 | *GSDMA* | | 5.331 | 4.19E-07 | 0.009 |
| | | 38,119,649 | | *GSDMA* | 4.404 | 2.19E-05 | 0.002 |
| rs2872507 | g.38040763.A>G | 37,828,296 | | *PGAP3* | 3.739 | 2.75E-04 | 0.002 |
| | | 37,927,002 | | *IKZF3* | -3.528 | 5.80E-04 | 0.001 |
| | | 38,018,804 | | *IKZF3* | 3.616 | 4.26E-04 | 0.046 |
| | | 38,077,191 | | | 4.332 | 2.93E-05 | 0.034 |
| | | 38,080,829 | | *ORMDL3* | 3.435 | 7.95E-04 | 0.014 |
| | | 38,081,812 | | *ORMDL3* | 3.459 | 7.34E-04 | 2.01E-04 |
| | | 38,081,829 | | *ORMDL3* | 3.730 | 2.85E-04 | 2.01E-04 |
| | | 38,081,995 | | *ORMDL3* | 5.842 | 3.92E-08 | 2.01E-04 |
| | | 38,082,022 | | *ORMDL3* | 6.694 | 5.90E-10 | 6.40E-04 |
| | | 38,082,035 | | *ORMDL3* | 6.772 | 3.96E-10 | 6.40E-04 |
| | | 38,082,073 | | *ORMDL3* | 6.832 | 2.91E-10 | 6.40E-04 |
| | | 38,082,206 | | *ORMDL3* | 3.847 | 1.86E-04 | 0.004 |
| | | 38,082,235 | | *ORMDL3* | 4.278 | 3.62E-05 | 0.004 |
| | | 38,082,346 | | *ORMDL3* | 4.590 | 1.03E-05 | 0.011 |
| | | 38,082,448 | | *ORMDL3* | 3.644 | 3.87E-04 | 0.011 |
| | | 38,082,658 | | *ORMDL3* | 4.266 | 3.79E-05 | 0.011 |
| | | 38,082,778 | | *ORMDL3* | 4.147 | 6.05E-05 | 0.011 |
| | | 38,108,710 | | | -4.669 | 7.42E-06 | 0.039 |
| | | 38,119,198 | *GSDMA* | | 4.246 | 4.11E-05 | 0.009 |
| | | 38,119,207 | *GSDMA* | | 5.331 | 4.19E-07 | 0.009 |
| | | 38,119,649 | | *GSDMA* | 4.404 | 2.19E-05 | 0.002 |
| rs113731681 | g.38040895.T>G | 37,856,732 | | *ERBB2* | -3.668 | 3.56E-04 | 0.009 |
| | | 37,886,749 | | *MIEN1* | 3.945 | 1.30E-04 | 0.002 |
| | | 37,896,063 | | | 3.476 | 6.92E-04 | 0.009 |
| | | 38,020,485 | *IKZF3* | | 3.962 | 1.22E-04 | 0.005 |
| | | 38,080,829 | | *ORMDL3* | 3.995 | 1.08E-04 | 0.014 |
| | | 38,096,333 | | | -3.664 | 3.60E-04 | 0.033 |
| | | 38,096,571 | | | -4.780 | 4.67E-06 | 0.033 |
| | | 38,096,578 | | | -5.642 | 1.00E-07 | 0.033 |
| | | 38,096,648 | | | -3.887 | 1.61E-04 | 0.033 |
| rs112931462 | g.38041246.A>G | 37,856,732 | | *ERBB2* | -3.668 | 3.56E-04 | 0.009 |
| | | 37,886,749 | | *MIEN1* | 3.945 | 1.30E-04 | 0.002 |
| | | 37,896,063 | | | 3.476 | 6.92E-04 | 0.009 |

| | | | | | | | |
|---|---|---|---|---|---|---|---|
| | | 38,020,485 | *IKZF3* | | 3.962 | 1.22E-04 | 0.005 |
| | | 38,080,829 | | *ORMDL3* | 3.995 | 1.08E-04 | 0.014 |
| | | 38,096,333 | | | -3.664 | 3.60E-04 | 0.033 |
| | | 38,096,571 | | | -4.780 | 4.67E-06 | 0.033 |
| | | 38,096,578 | | | -5.642 | 1.00E-07 | 0.033 |
| | | 38,096,648 | | | -3.887 | 1.61E-04 | 0.033 |
| rs12948927 | g.38041484.T>G | 37,856,732 | | *ERBB2* | -3.668 | 3.56E-04 | 0.009 |
| | | 37,886,749 | | *MIEN1* | 3.945 | 1.30E-04 | 0.002 |
| | | 37,896,063 | | | 3.476 | 6.92E-04 | 0.009 |
| | | 38,020,485 | *IKZF3* | | 3.962 | 1.22E-04 | 0.005 |
| | | 38,080,829 | | *ORMDL3* | 3.995 | 1.08E-04 | 0.014 |
| | | 38,096,333 | | | -3.664 | 3.60E-04 | 0.033 |
| | | 38,096,571 | | | -4.780 | 4.67E-06 | 0.033 |
| | | 38,096,578 | | | -5.642 | 1.00E-07 | 0.033 |
| | | 38,096,648 | | | -3.887 | 1.61E-04 | 0.033 |
| rs9908132 | g.38042777.A>T | 37,927,002 | | *IKZF3* | -3.588 | 4.70E-04 | 0.001 |
| | | 38,018,804 | | *IKZF3* | 4.291 | 3.45E-05 | 0.046 |
| | | 38,049,272 | | | 3.531 | 5.74E-04 | 0.046 |
| | | 38,077,191 | | | 4.101 | 7.22E-05 | 0.034 |
| | | 38,080,829 | | *ORMDL3* | 5.326 | 4.28E-07 | 0.014 |
| | | 38,081,812 | | *ORMDL3* | 3.387 | 9.35E-04 | 2.01E-04 |
| | | 38,081,995 | | *ORMDL3* | 4.489 | 1.56E-05 | 2.01E-04 |
| | | 38,082,022 | | *ORMDL3* | 5.027 | 1.62E-06 | 6.40E-04 |
| | | 38,082,035 | | *ORMDL3* | 6.122 | 1.02E-08 | 6.40E-04 |
| | | 38,082,073 | | *ORMDL3* | 5.874 | 3.36E-08 | 6.40E-04 |
| | | 38,082,235 | | *ORMDL3* | 3.957 | 1.24E-04 | 0.004 |
| | | 38,082,346 | | *ORMDL3* | 4.643 | 8.28E-06 | 0.011 |
| | | 38,082,448 | | *ORMDL3* | 4.146 | 6.05E-05 | 0.011 |
| | | 38,082,658 | | *ORMDL3* | 4.337 | 2.87E-05 | 0.011 |
| | | 38,082,778 | | *ORMDL3* | 3.671 | 3.51E-04 | 0.011 |
| | | 38,108,710 | | | -4.040 | 9.10E-05 | 0.039 |
| | | 38,119,198 | *GSDMA* | | 4.199 | 4.94E-05 | 0.009 |
| | | 38,119,207 | *GSDMA* | | 5.693 | 7.91E-08 | 0.009 |
| | | 38,119,649 | | *GSDMA* | 4.145 | 6.08E-05 | 0.002 |
| rs9901146 | g.38043343.A>G | 37,927,002 | | *IKZF3* | -3.588 | 4.70E-04 | 0.001 |
| | | 38,018,804 | | *IKZF3* | 4.291 | 3.45E-05 | 0.046 |
| | | 38,049,272 | | | 3.531 | 5.74E-04 | 0.046 |
| | | 38,077,191 | | | 4.101 | 7.22E-05 | 0.034 |
| | | 38,080,829 | | *ORMDL3* | 5.326 | 4.28E-07 | 0.014 |
| | | 38,081,812 | | *ORMDL3* | 3.387 | 9.35E-04 | 2.01E-04 |
| | | 38,081,995 | | *ORMDL3* | 4.489 | 1.56E-05 | 2.01E-04 |

| | | | | | | | |
|---|---|---|---|---|---|---|---|
| | | 38,082,022 | | *ORMDL3* | 5.027 | 1.62E-06 | 6.40E-04 |
| | | 38,082,035 | | *ORMDL3* | 6.122 | 1.02E-08 | 6.40E-04 |
| | | 38,082,073 | | *ORMDL3* | 5.874 | 3.36E-08 | 6.40E-04 |
| | | 38,082,235 | | *ORMDL3* | 3.957 | 1.24E-04 | 0.004 |
| | | 38,082,346 | | *ORMDL3* | 4.643 | 8.28E-06 | 0.011 |
| | | 38,082,448 | | *ORMDL3* | 4.146 | 6.05E-05 | 0.011 |
| | | 38,082,658 | | *ORMDL3* | 4.337 | 2.87E-05 | 0.011 |
| | | 38,082,778 | | *ORMDL3* | 3.671 | 3.51E-04 | 0.011 |
| | | 38,108,710 | | | -4.040 | 9.10E-05 | 0.039 |
| | | 38,119,198 | *GSDMA* | | 4.199 | 4.94E-05 | 0.009 |
| | | 38,119,207 | *GSDMA* | | 5.693 | 7.91E-08 | 0.009 |
| | | 38,119,649 | | *GSDMA* | 4.145 | 6.08E-05 | 0.002 |
| rs12936409 | g.38043649.T>C | 37,828,296 | | *PGAP3* | 3.739 | 2.75E-04 | 0.002 |
| | | 37,927,002 | | *IKZF3* | -3.528 | 5.80E-04 | 0.001 |
| | | 38,018,804 | | *IKZF3* | 3.616 | 4.26E-04 | 0.046 |
| | | 38,077,191 | | | 4.332 | 2.93E-05 | 0.034 |
| | | 38,080,829 | | *ORMDL3* | 3.435 | 7.95E-04 | 0.014 |
| | | 38,081,812 | | *ORMDL3* | 3.459 | 7.34E-04 | 2.01E-04 |
| | | 38,081,829 | | *ORMDL3* | 3.730 | 2.85E-04 | 2.01E-04 |
| | | 38,081,995 | | *ORMDL3* | 5.842 | 3.92E-08 | 2.01E-04 |
| | | 38,082,022 | | *ORMDL3* | 6.694 | 5.90E-10 | 6.40E-04 |
| | | 38,082,035 | | *ORMDL3* | 6.772 | 3.96E-10 | 6.40E-04 |
| | | 38,082,073 | | *ORMDL3* | 6.832 | 2.91E-10 | 6.40E-04 |
| | | 38,082,206 | | *ORMDL3* | 3.847 | 1.86E-04 | 0.004 |
| | | 38,082,235 | | *ORMDL3* | 4.278 | 3.62E-05 | 0.004 |
| | | 38,082,346 | | *ORMDL3* | 4.590 | 1.03E-05 | 0.011 |
| | | 38,082,448 | | *ORMDL3* | 3.644 | 3.87E-04 | 0.011 |
| | | 38,082,658 | | *ORMDL3* | 4.266 | 3.79E-05 | 0.011 |
| | | 38,082,778 | | *ORMDL3* | 4.147 | 6.05E-05 | 0.011 |
| | | 38,108,710 | | | -4.669 | 7.42E-06 | 0.039 |
| | | 38,119,198 | *GSDMA* | | 4.246 | 4.11E-05 | 0.009 |
| | | 38,119,207 | *GSDMA* | | 5.331 | 4.19E-07 | 0.009 |
| | | 38,119,649 | | *GSDMA* | 4.404 | 2.19E-05 | 0.002 |
| rs112557679 | g.38044164.G>A | 37,856,732 | | *ERBB2* | -3.668 | 3.56E-04 | 0.009 |
| | | 37,886,749 | | *MIEN1* | 3.945 | 1.30E-04 | 0.002 |
| | | 37,896,063 | | | 3.476 | 6.92E-04 | 0.009 |
| | | 38,020,485 | *IKZF3* | | 3.962 | 1.22E-04 | 0.005 |
| | | 38,080,829 | | *ORMDL3* | 3.995 | 1.08E-04 | 0.014 |
| | | 38,096,333 | | | -3.664 | 3.60E-04 | 0.033 |
| | | 38,096,571 | | | -4.780 | 4.67E-06 | 0.033 |
| | | 38,096,578 | | | -5.642 | 1.00E-07 | 0.033 |

| | | | | | | | |
|---|---|---|---|---|---|---|---|
| | | 38,096,648 | | | -3.887 | 1.61E-04 | 0.033 |
| rs111317272 | g.38044525.G>C | 37,856,732 | | *ERBB2* | -3.668 | 3.56E-04 | 0.009 |
| | | 37,886,749 | | *MIEN1* | 3.945 | 1.30E-04 | 0.002 |
| | | 37,896,063 | | | 3.476 | 6.92E-04 | 0.009 |
| | | 38,020,485 | *IKZF3* | | 3.962 | 1.22E-04 | 0.005 |
| | | 38,080,829 | | *ORMDL3* | 3.995 | 1.08E-04 | 0.014 |
| | | 38,096,333 | | | -3.664 | 3.60E-04 | 0.033 |
| | | 38,096,571 | | | -4.780 | 4.67E-06 | 0.033 |
| | | 38,096,578 | | | -5.642 | 1.00E-07 | 0.033 |
| | | 38,096,648 | | | -3.887 | 1.61E-04 | 0.033 |
| rs33938760 | | 37,927,002 | | *IKZF3* | -3.625 | 4.14E-04 | 0.001 |
| | | 38,018,804 | | *IKZF3* | 4.307 | 3.24E-05 | 0.046 |
| | | 38,043,310 | | | -3.369 | 9.95E-04 | 0.046 |
| | | 38,049,272 | | | 3.612 | 4.33E-04 | 0.046 |
| | | 38,077,191 | | | 3.946 | 1.29E-04 | 0.034 |
| | | 38,080,829 | | *ORMDL3* | 5.142 | 9.74E-07 | 0.014 |
| | | 38,081,995 | | *ORMDL3* | 4.367 | 2.55E-05 | 2.01E-04 |
| | | 38,082,022 | | *ORMDL3* | 4.972 | 2.05E-06 | 6.40E-04 |
| | | 38,082,035 | | *ORMDL3* | 6.234 | 5.87E-09 | 6.40E-04 |
| | | 38,082,073 | | *ORMDL3* | 5.796 | 4.86E-08 | 6.40E-04 |
| | | 38,082,235 | | *ORMDL3* | 3.778 | 2.40E-04 | 0.004 |
| | | 38,082,346 | | *ORMDL3* | 4.701 | 6.51E-06 | 0.011 |
| | | 38,082,448 | | *ORMDL3* | 4.086 | 7.63E-05 | 0.011 |
| | | 38,082,658 | | *ORMDL3* | 4.277 | 3.64E-05 | 0.011 |
| | | 38,082,778 | | *ORMDL3* | 3.600 | 4.52E-04 | 0.011 |
| | | 38,108,710 | | | -3.862 | 1.77E-04 | 0.039 |
| | | 38,119,198 | *GSDMA* | | 4.159 | 5.76E-05 | 0.009 |
| | | 38,119,207 | *GSDMA* | | 5.636 | 1.03E-07 | 0.009 |
| | | 38,119,649 | | *GSDMA* | 4.055 | 8.57E-05 | 0.002 |
| rs113486187 | g.38045043.A>G | 37,856,732 | | *ERBB2* | -3.668 | 3.56E-04 | 0.009 |
| | | 37,886,749 | | *MIEN1* | 3.945 | 1.30E-04 | 0.002 |
| | | 37,896,063 | | | 3.476 | 6.92E-04 | 0.009 |
| | | 38,020,485 | *IKZF3* | | 3.962 | 1.22E-04 | 0.005 |
| | | 38,080,829 | | *ORMDL3* | 3.995 | 1.08E-04 | 0.014 |
| | | 38,096,333 | | | -3.664 | 3.60E-04 | 0.033 |
| | | 38,096,571 | | | -4.780 | 4.67E-06 | 0.033 |
| | | 38,096,578 | | | -5.642 | 1.00E-07 | 0.033 |
| | | 38,096,648 | | | -3.887 | 1.61E-04 | 0.033 |
| rs12103884 | g.38045725.T>C | 37,927,002 | | *IKZF3* | -3.588 | 4.70E-04 | 0.001 |
| | | 38,018,804 | | *IKZF3* | 4.291 | 3.45E-05 | 0.046 |
| | | 38,049,272 | | | 3.531 | 5.74E-04 | 0.046 |

| | | | | | | | |
|---|---|---|---|---|---|---|---|
| | | 38,077,191 | | | 4.101 | 7.22E-05 | 0.034 |
| | | 38,080,829 | | *ORMDL3* | 5.326 | 4.28E-07 | 0.014 |
| | | 38,081,812 | | *ORMDL3* | 3.387 | 9.35E-04 | 2.01E-04 |
| | | 38,081,995 | | *ORMDL3* | 4.489 | 1.56E-05 | 2.01E-04 |
| | | 38,082,022 | | *ORMDL3* | 5.027 | 1.62E-06 | 6.40E-04 |
| | | 38,082,035 | | *ORMDL3* | 6.122 | 1.02E-08 | 6.40E-04 |
| | | 38,082,073 | | *ORMDL3* | 5.874 | 3.36E-08 | 6.40E-04 |
| | | 38,082,235 | | *ORMDL3* | 3.957 | 1.24E-04 | 0.004 |
| | | 38,082,346 | | *ORMDL3* | 4.643 | 8.28E-06 | 0.011 |
| | | 38,082,448 | | *ORMDL3* | 4.146 | 6.05E-05 | 0.011 |
| | | 38,082,658 | | *ORMDL3* | 4.337 | 2.87E-05 | 0.011 |
| | | 38,082,778 | | *ORMDL3* | 3.671 | 3.51E-04 | 0.011 |
| | | 38,108,710 | | | -4.040 | 9.10E-05 | 0.039 |
| | | 38,119,198 | *GSDMA* | | 4.199 | 4.94E-05 | 0.009 |
| | | 38,119,207 | *GSDMA* | | 5.693 | 7.91E-08 | 0.009 |
| | | 38,119,649 | | *GSDMA* | 4.145 | 6.08E-05 | 0.002 |
| rs112049571 | g.38047658.A>G | 37,856,732 | | *ERBB2* | -3.668 | 3.56E-04 | 0.009 |
| | | 37,886,749 | | *MIEN1* | 3.945 | 1.30E-04 | 0.002 |
| | | 37,896,063 | | | 3.476 | 6.92E-04 | 0.009 |
| | | 38,020,485 | *IKZF3* | | 3.962 | 1.22E-04 | 0.005 |
| | | 38,080,829 | | *ORMDL3* | 3.995 | 1.08E-04 | 0.014 |
| | | 38,096,333 | | | -3.664 | 3.60E-04 | 0.033 |
| | | 38,096,571 | | | -4.780 | 4.67E-06 | 0.033 |
| | | 38,096,578 | | | -5.642 | 1.00E-07 | 0.033 |
| | | 38,096,648 | | | -3.887 | 1.61E-04 | 0.033 |
| rs9906951 | g.38048244.C>T | 37,927,002 | | *IKZF3* | -3.588 | 4.70E-04 | 0.001 |
| | | 38,018,804 | | *IKZF3* | 4.291 | 3.45E-05 | 0.046 |
| | | 38,049,272 | | | 3.531 | 5.74E-04 | 0.046 |
| | | 38,077,191 | | | 4.101 | 7.22E-05 | 0.034 |
| | | 38,080,829 | | *ORMDL3* | 5.326 | 4.28E-07 | 0.014 |
| | | 38,081,812 | | *ORMDL3* | 3.387 | 9.35E-04 | 2.01E-04 |
| | | 38,081,995 | | *ORMDL3* | 4.489 | 1.56E-05 | 2.01E-04 |
| | | 38,082,022 | | *ORMDL3* | 5.027 | 1.62E-06 | 6.40E-04 |
| | | 38,082,035 | | *ORMDL3* | 6.122 | 1.02E-08 | 6.40E-04 |
| | | 38,082,073 | | *ORMDL3* | 5.874 | 3.36E-08 | 6.40E-04 |
| | | 38,082,235 | | *ORMDL3* | 3.957 | 1.24E-04 | 0.004 |
| | | 38,082,346 | | *ORMDL3* | 4.643 | 8.28E-06 | 0.011 |
| | | 38,082,448 | | *ORMDL3* | 4.146 | 6.05E-05 | 0.011 |
| | | 38,082,658 | | *ORMDL3* | 4.337 | 2.87E-05 | 0.011 |
| | | 38,082,778 | | *ORMDL3* | 3.671 | 3.51E-04 | 0.011 |
| | | 38,108,710 | | | -4.040 | 9.10E-05 | 0.039 |

| | | | | | | | |
|---|---|---|---|---|---|---|---|
| | | 38,119,198 | *GSDMA* | | 4.199 | 4.94E-05 | 0.009 |
| | | 38,119,207 | *GSDMA* | | 5.693 | 7.91E-08 | 0.009 |
| | | 38,119,649 | | *GSDMA* | 4.145 | 6.08E-05 | 0.002 |
| rs75760678 | g.38048721.T>C | 37,856,732 | | *ERBB2* | -3.668 | 3.56E-04 | 0.009 |
| | | 37,886,749 | | *MIEN1* | 3.945 | 1.30E-04 | 0.002 |
| | | 37,896,063 | | | 3.476 | 6.92E-04 | 0.009 |
| | | 38,020,485 | *IKZF3* | | 3.962 | 1.22E-04 | 0.005 |
| | | 38,080,829 | | *ORMDL3* | 3.995 | 1.08E-04 | 0.014 |
| | | 38,096,333 | | | -3.664 | 3.60E-04 | 0.033 |
| | | 38,096,571 | | | -4.780 | 4.67E-06 | 0.033 |
| | | 38,096,578 | | | -5.642 | 1.00E-07 | 0.033 |
| | | 38,096,648 | | | -3.887 | 1.61E-04 | 0.033 |
| rs12950209 | g.38049102.C>T | 37,927,002 | | *IKZF3* | -3.588 | 4.70E-04 | 0.001 |
| | | 38,018,804 | | *IKZF3* | 4.291 | 3.45E-05 | 0.046 |
| | | 38,049,272 | | | 3.531 | 5.74E-04 | 0.046 |
| | | 38,077,191 | | | 4.101 | 7.22E-05 | 0.034 |
| | | 38,080,829 | | *ORMDL3* | 5.326 | 4.28E-07 | 0.014 |
| | | 38,081,812 | | *ORMDL3* | 3.387 | 9.35E-04 | 2.01E-04 |
| | | 38,081,995 | | *ORMDL3* | 4.489 | 1.56E-05 | 2.01E-04 |
| | | 38,082,022 | | *ORMDL3* | 5.027 | 1.62E-06 | 6.40E-04 |
| | | 38,082,035 | | *ORMDL3* | 6.122 | 1.02E-08 | 6.40E-04 |
| | | 38,082,073 | | *ORMDL3* | 5.874 | 3.36E-08 | 6.40E-04 |
| | | 38,082,235 | | *ORMDL3* | 3.957 | 1.24E-04 | 0.004 |
| | | 38,082,346 | | *ORMDL3* | 4.643 | 8.28E-06 | 0.011 |
| | | 38,082,448 | | *ORMDL3* | 4.146 | 6.05E-05 | 0.011 |
| | | 38,082,658 | | *ORMDL3* | 4.337 | 2.87E-05 | 0.011 |
| | | 38,082,778 | | *ORMDL3* | 3.671 | 3.51E-04 | 0.011 |
| | | 38,108,710 | | | -4.040 | 9.10E-05 | 0.039 |
| | | 38,119,198 | *GSDMA* | | 4.199 | 4.94E-05 | 0.009 |
| | | 38,119,207 | *GSDMA* | | 5.693 | 7.91E-08 | 0.009 |
| | | 38,119,649 | | *GSDMA* | 4.145 | 6.08E-05 | 0.002 |
| rs12950743 | g.38049233.C>T | 37,927,002 | | *IKZF3* | -3.588 | 4.70E-04 | 0.001 |
| | | 38,018,804 | | *IKZF3* | 4.291 | 3.45E-05 | 0.046 |
| | | 38,049,272 | | | 3.531 | 5.74E-04 | 0.046 |
| | | 38,077,191 | | | 4.101 | 7.22E-05 | 0.034 |
| | | 38,080,829 | | *ORMDL3* | 5.326 | 4.28E-07 | 0.014 |
| | | 38,081,812 | | *ORMDL3* | 3.387 | 9.35E-04 | 2.01E-04 |
| | | 38,081,995 | | *ORMDL3* | 4.489 | 1.56E-05 | 2.01E-04 |
| | | 38,082,022 | | *ORMDL3* | 5.027 | 1.62E-06 | 6.40E-04 |
| | | 38,082,035 | | *ORMDL3* | 6.122 | 1.02E-08 | 6.40E-04 |
| | | 38,082,073 | | *ORMDL3* | 5.874 | 3.36E-08 | 6.40E-04 |

| | | | | | | | |
|---|---|---|---|---|---|---|---|
| | | 38,082,235 | | *ORMDL3* | 3.957 | 1.24E-04 | 0.004 |
| | | 38,082,346 | | *ORMDL3* | 4.643 | 8.28E-06 | 0.011 |
| | | 38,082,448 | | *ORMDL3* | 4.146 | 6.05E-05 | 0.011 |
| | | 38,082,658 | | *ORMDL3* | 4.337 | 2.87E-05 | 0.011 |
| | | 38,082,778 | | *ORMDL3* | 3.671 | 3.51E-04 | 0.011 |
| | | 38,108,710 | | | -4.040 | 9.10E-05 | 0.039 |
| | | 38,119,198 | *GSDMA* | | 4.199 | 4.94E-05 | 0.009 |
| | | 38,119,207 | *GSDMA* | | 5.693 | 7.91E-08 | 0.009 |
| | | 38,119,649 | | *GSDMA* | 4.145 | 6.08E-05 | 0.002 |
| rs7359623 | g.38049589.T>C | 37,927,002 | | *IKZF3* | -3.544 | 5.48E-04 | 0.001 |
| | | 38,018,804 | | *IKZF3* | 4.236 | 4.28E-05 | 0.046 |
| | | 38,043,310 | | | -3.418 | 8.43E-04 | 0.046 |
| | | 38,049,272 | | | 3.443 | 7.75E-04 | 0.046 |
| | | 38,077,191 | | | 4.237 | 4.27E-05 | 0.034 |
| | | 38,080,829 | | *ORMDL3* | 5.264 | 5.67E-07 | 0.014 |
| | | 38,081,492 | | *ORMDL3* | 3.398 | 9.01E-04 | 2.01E-04 |
| | | 38,081,530 | | *ORMDL3* | 3.403 | 8.86E-04 | 2.01E-04 |
| | | 38,081,995 | | *ORMDL3* | 4.362 | 2.60E-05 | 2.01E-04 |
| | | 38,082,022 | | *ORMDL3* | 5.120 | 1.08E-06 | 6.40E-04 |
| | | 38,082,035 | | *ORMDL3* | 6.111 | 1.07E-08 | 6.40E-04 |
| | | 38,082,073 | | *ORMDL3* | 6.116 | 1.05E-08 | 6.40E-04 |
| | | 38,082,235 | | *ORMDL3* | 3.900 | 1.54E-04 | 0.004 |
| | | 38,082,346 | | *ORMDL3* | 4.755 | 5.18E-06 | 0.011 |
| | | 38,082,448 | | *ORMDL3* | 4.108 | 7.00E-05 | 0.011 |
| | | 38,082,658 | | *ORMDL3* | 4.079 | 7.85E-05 | 0.011 |
| | | 38,082,778 | | *ORMDL3* | 3.751 | 2.64E-04 | 0.011 |
| | | 38,084,256 | *ORMDL3* | | -3.421 | 8.35E-04 | 0.005 |
| | | 38,108,710 | | | -4.168 | 5.56E-05 | 0.039 |
| | | 38,119,198 | *GSDMA* | | 4.234 | 4.31E-05 | 0.009 |
| | | 38,119,207 | *GSDMA* | | 5.672 | 8.72E-08 | 0.009 |
| | | 38,119,649 | | *GSDMA* | 4.113 | 6.87E-05 | 0.002 |
| rs68122720 | | 37,828,296 | | *PGAP3* | 3.739 | 2.75E-04 | 0.002 |
| | | 37,927,002 | | *IKZF3* | -3.528 | 5.80E-04 | 0.001 |
| | | 38,018,804 | | *IKZF3* | 3.616 | 4.26E-04 | 0.046 |
| | | 38,077,191 | | | 4.332 | 2.93E-05 | 0.034 |
| | | 38,080,829 | | *ORMDL3* | 3.435 | 7.95E-04 | 0.014 |
| | | 38,081,812 | | *ORMDL3* | 3.459 | 7.34E-04 | 2.01E-04 |
| | | 38,081,829 | | *ORMDL3* | 3.730 | 2.85E-04 | 2.01E-04 |
| | | 38,081,995 | | *ORMDL3* | 5.842 | 3.92E-08 | 2.01E-04 |
| | | 38,082,022 | | *ORMDL3* | 6.694 | 5.90E-10 | 6.40E-04 |
| | | 38,082,035 | | *ORMDL3* | 6.772 | 3.96E-10 | 6.40E-04 |

| | | | | | | | |
|---|---|---|---|---|---|---|---|
| | | 38,082,073 | | *ORMDL3* | 6.832 | 2.91E-10 | 6.40E-04 |
| | | 38,082,206 | | *ORMDL3* | 3.847 | 1.86E-04 | 0.004 |
| | | 38,082,235 | | *ORMDL3* | 4.278 | 3.62E-05 | 0.004 |
| | | 38,082,346 | | *ORMDL3* | 4.590 | 1.03E-05 | 0.011 |
| | | 38,082,448 | | *ORMDL3* | 3.644 | 3.87E-04 | 0.011 |
| | | 38,082,658 | | *ORMDL3* | 4.266 | 3.79E-05 | 0.011 |
| | | 38,082,778 | | *ORMDL3* | 4.147 | 6.05E-05 | 0.011 |
| | | 38,108,710 | | | -4.669 | 7.42E-06 | 0.039 |
| | | 38,119,198 | *GSDMA* | | 4.246 | 4.11E-05 | 0.009 |
| | | 38,119,207 | *GSDMA* | | 5.331 | 4.19E-07 | 0.009 |
| | | 38,119,649 | | *GSDMA* | 4.404 | 2.19E-05 | 0.002 |
| rs17676923 | g.38050125.T>C | 37,856,732 | | *ERBB2* | -3.585 | 4.76E-04 | 0.009 |
| | | 37,886,749 | | *MIEN1* | 3.926 | 1.39E-04 | 0.002 |
| | | 37,896,006 | | | 3.373 | 9.81E-04 | 0.027 |
| | | 37,896,063 | | | 3.488 | 6.65E-04 | 0.009 |
| | | 38,020,485 | *IKZF3* | | 3.946 | 1.29E-04 | 0.005 |
| | | 38,080,829 | | *ORMDL3* | 3.972 | 1.17E-04 | 0.014 |
| | | 38,096,333 | | | -3.696 | 3.21E-04 | 0.033 |
| | | 38,096,571 | | | -4.793 | 4.42E-06 | 0.033 |
| | | 38,096,578 | | | -5.637 | 1.03E-07 | 0.033 |
| | | 38,096,648 | | | -3.898 | 1.55E-04 | 0.033 |
| rs35336365 | g.38050715.G>A | 37,856,732 | | *ERBB2* | -3.668 | 3.56E-04 | 0.009 |
| | | 37,886,749 | | *MIEN1* | 3.945 | 1.30E-04 | 0.002 |
| | | 37,896,063 | | | 3.476 | 6.92E-04 | 0.009 |
| | | 38,020,485 | *IKZF3* | | 3.962 | 1.22E-04 | 0.005 |
| | | 38,080,829 | | *ORMDL3* | 3.995 | 1.08E-04 | 0.014 |
| | | 38,096,333 | | | -3.664 | 3.60E-04 | 0.033 |
| | | 38,096,571 | | | -4.780 | 4.67E-06 | 0.033 |
| | | 38,096,578 | | | -5.642 | 1.00E-07 | 0.033 |
| | | 38,096,648 | | | -3.887 | 1.61E-04 | 0.033 |
| rs8067378 | g.38051348.G>A | 37,927,002 | | *IKZF3* | -3.588 | 4.70E-04 | 0.001 |
| | | 38,018,804 | | *IKZF3* | 4.291 | 3.45E-05 | 0.046 |
| | | 38,049,272 | | | 3.531 | 5.74E-04 | 0.046 |
| | | 38,077,191 | | | 4.101 | 7.22E-05 | 0.034 |
| | | 38,080,829 | | *ORMDL3* | 5.326 | 4.28E-07 | 0.014 |
| | | 38,081,812 | | *ORMDL3* | 3.387 | 9.35E-04 | 2.01E-04 |
| | | 38,081,995 | | *ORMDL3* | 4.489 | 1.56E-05 | 2.01E-04 |
| | | 38,082,022 | | *ORMDL3* | 5.027 | 1.62E-06 | 6.40E-04 |
| | | 38,082,035 | | *ORMDL3* | 6.122 | 1.02E-08 | 6.40E-04 |
| | | 38,082,073 | | *ORMDL3* | 5.874 | 3.36E-08 | 6.40E-04 |
| | | 38,082,235 | | *ORMDL3* | 3.957 | 1.24E-04 | 0.004 |

| | | | | | | | |
|---|---|---|---|---|---|---|---|
| | | 38,082,346 | | *ORMDL3* | 4.643 | 8.28E-06 | 0.011 |
| | | 38,082,448 | | *ORMDL3* | 4.146 | 6.05E-05 | 0.011 |
| | | 38,082,658 | | *ORMDL3* | 4.337 | 2.87E-05 | 0.011 |
| | | 38,082,778 | | *ORMDL3* | 3.671 | 3.51E-04 | 0.011 |
| | | 38,108,710 | | | -4.040 | 9.10E-05 | 0.039 |
| | | 38,119,198 | *GSDMA* | | 4.199 | 4.94E-05 | 0.009 |
| | | 38,119,207 | *GSDMA* | | 5.693 | 7.91E-08 | 0.009 |
| | | 38,119,649 | | *GSDMA* | 4.145 | 6.08E-05 | 0.002 |
| rs189027840 | g.38052195.A>T | 37,856,732 | | *ERBB2* | -3.680 | 3.40E-04 | 0.009 |
| | | 37,886,749 | | *MIEN1* | 3.934 | 1.35E-04 | 0.002 |
| | | 37,896,006 | | | 3.400 | 8.96E-04 | 0.027 |
| | | 37,896,063 | | | 3.467 | 7.15E-04 | 0.009 |
| | | 38,020,485 | *IKZF3* | | 3.948 | 1.28E-04 | 0.005 |
| | | 38,080,829 | | *ORMDL3* | 3.980 | 1.14E-04 | 0.014 |
| | | 38,096,333 | | | -3.724 | 2.91E-04 | 0.033 |
| | | 38,096,571 | | | -4.804 | 4.22E-06 | 0.033 |
| | | 38,096,578 | | | -5.715 | 7.14E-08 | 0.033 |
| | | 38,096,648 | | | -3.867 | 1.74E-04 | 0.033 |
| rs12453507 | g.38053207.G>C | 37,927,002 | | *IKZF3* | -3.462 | 7.27E-04 | 0.001 |
| | | 38,018,804 | | *IKZF3* | 3.928 | 1.38E-04 | 0.046 |
| | | 38,049,272 | | | 3.863 | 1.76E-04 | 0.046 |
| | | 38,077,191 | | | 4.097 | 7.31E-05 | 0.034 |
| | | 38,080,829 | | *ORMDL3* | 5.660 | 9.22E-08 | 0.014 |
| | | 38,081,492 | | *ORMDL3* | 3.905 | 1.51E-04 | 2.01E-04 |
| | | 38,081,812 | | *ORMDL3* | 4.013 | 1.01E-04 | 2.01E-04 |
| | | 38,081,829 | | *ORMDL3* | 3.393 | 9.16E-04 | 2.01E-04 |
| | | 38,081,995 | | *ORMDL3* | 4.821 | 3.92E-06 | 2.01E-04 |
| | | 38,082,022 | | *ORMDL3* | 5.398 | 3.09E-07 | 6.40E-04 |
| | | 38,082,035 | | *ORMDL3* | 5.607 | 1.18E-07 | 6.40E-04 |
| | | 38,082,073 | | *ORMDL3* | 5.832 | 4.11E-08 | 6.40E-04 |
| | | 38,082,235 | | *ORMDL3* | 3.750 | 2.65E-04 | 0.004 |
| | | 38,082,346 | | *ORMDL3* | 4.646 | 8.20E-06 | 0.011 |
| | | 38,082,448 | | *ORMDL3* | 3.899 | 1.54E-04 | 0.011 |
| | | 38,082,658 | | *ORMDL3* | 4.688 | 6.88E-06 | 0.011 |
| | | 38,082,778 | | *ORMDL3* | 3.986 | 1.12E-04 | 0.011 |
| | | 38,108,710 | | | -4.205 | 4.83E-05 | 0.039 |
| | | 38,119,198 | *GSDMA* | | 4.018 | 9.86E-05 | 0.009 |
| | | 38,119,207 | *GSDMA* | | 5.704 | 7.52E-08 | 0.009 |
| | | 38,119,649 | | *GSDMA* | 4.452 | 1.81E-05 | 0.002 |
| rs2123685 | g.38053889.C>T | 37,856,732 | | *ERBB2* | -3.668 | 3.56E-04 | 0.009 |
| | | 37,886,749 | | *MIEN1* | 3.945 | 1.30E-04 | 0.002 |

| | | | | | | | |
|---|---|---|---|---|---|---|---|
| | | 37,896,063 | | | 3.476 | 6.92E-04 | 0.009 |
| | | 38,020,485 | *IKZF3* | | 3.962 | 1.22E-04 | 0.005 |
| | | 38,080,829 | | *ORMDL3* | 3.995 | 1.08E-04 | 0.014 |
| | | 38,096,333 | | | -3.664 | 3.60E-04 | 0.033 |
| | | 38,096,571 | | | -4.780 | 4.67E-06 | 0.033 |
| | | 38,096,578 | | | -5.642 | 1.00E-07 | 0.033 |
| | | 38,096,648 | | | -3.887 | 1.61E-04 | 0.033 |
| rs34170568 | g.38055921_38055922ins | 38,018,804 | | *IKZF3* | 3.902 | 1.53E-04 | 0.046 |
| | | 38,077,051 | | | 3.733 | 2.82E-04 | 0.034 |
| | | 38,077,191 | | | 3.664 | 3.60E-04 | 0.034 |
| | | 38,080,829 | | *ORMDL3* | 3.923 | 1.41E-04 | 0.014 |
| | | 38,081,829 | | *ORMDL3* | 3.668 | 3.55E-04 | 2.01E-04 |
| | | 38,081,995 | | *ORMDL3* | 5.423 | 2.76E-07 | 2.01E-04 |
| | | 38,082,022 | | *ORMDL3* | 6.265 | 5.04E-09 | 6.40E-04 |
| | | 38,082,035 | | *ORMDL3* | 5.503 | 1.91E-07 | 6.40E-04 |
| | | 38,082,073 | | *ORMDL3* | 5.823 | 4.29E-08 | 6.40E-04 |
| | | 38,082,206 | | *ORMDL3* | 3.480 | 6.82E-04 | 0.004 |
| | | 38,082,235 | | *ORMDL3* | 3.784 | 2.35E-04 | 0.004 |
| | | 38,082,346 | | *ORMDL3* | 4.313 | 3.15E-05 | 0.011 |
| | | 38,082,658 | | *ORMDL3* | 3.958 | 1.24E-04 | 0.011 |
| | | 38,082,778 | | *ORMDL3* | 3.959 | 1.23E-04 | 0.011 |
| | | 38,083,308 | | *ORMDL3* | 3.409 | 8.69E-04 | 0.011 |
| | | 38,108,710 | | | -5.002 | 1.80E-06 | 0.039 |
| | | 38,119,198 | *GSDMA* | | 4.751 | 5.28E-06 | 0.009 |
| | | 38,119,207 | *GSDMA* | | 5.838 | 4.00E-08 | 0.009 |
| | | 38,119,649 | | *GSDMA* | 4.646 | 8.20E-06 | 0.002 |
| | | 38,120,234 | | *GSDMA* | 3.733 | 2.82E-04 | 0.011 |
| rs11651596 | g.38056116.C>T | 37,828,296 | | *PGAP3* | 3.529 | 5.77E-04 | 0.002 |
| | | 38,018,804 | | *IKZF3* | 3.445 | 7.69E-04 | 0.046 |
| | | 38,077,051 | | | 3.572 | 4.97E-04 | 0.034 |
| | | 38,077,191 | | | 4.013 | 1.01E-04 | 0.034 |
| | | 38,080,829 | | *ORMDL3* | 3.756 | 2.60E-04 | 0.014 |
| | | 38,081,829 | | *ORMDL3* | 3.981 | 1.13E-04 | 2.01E-04 |
| | | 38,081,995 | | *ORMDL3* | 5.846 | 3.84E-08 | 2.01E-04 |
| | | 38,082,022 | | *ORMDL3* | 6.711 | 5.39E-10 | 6.40E-04 |
| | | 38,082,035 | | *ORMDL3* | 6.267 | 4.99E-09 | 6.40E-04 |
| | | 38,082,073 | | *ORMDL3* | 6.097 | 1.15E-08 | 6.40E-04 |
| | | 38,082,206 | | *ORMDL3* | 3.756 | 2.60E-04 | 0.004 |
| | | 38,082,235 | | *ORMDL3* | 4.112 | 6.90E-05 | 0.004 |
| | | 38,082,346 | | *ORMDL3* | 4.364 | 2.58E-05 | 0.011 |

| | | | | | | | |
|---|---|---|---|---|---|---|---|
| | | 38,082,448 | | *ORMDL3* | 3.578 | 4.87E-04 | 0.011 |
| | | 38,082,658 | | *ORMDL3* | 4.529 | 1.32E-05 | 0.011 |
| | | 38,082,778 | | *ORMDL3* | 4.241 | 4.19E-05 | 0.011 |
| | | 38,108,710 | | | -4.853 | 3.43E-06 | 0.039 |
| | | 38,119,198 | *GSDMA* | | 4.226 | 4.44E-05 | 0.009 |
| | | 38,119,207 | *GSDMA* | | 5.401 | 3.05E-07 | 0.009 |
| | | 38,119,649 | | *GSDMA* | 4.702 | 6.48E-06 | 0.002 |
| | | 38,120,234 | | *GSDMA* | 3.575 | 4.93E-04 | 0.011 |
| rs79457595 | g.38056247.T>C | 37,856,732 | | *ERBB2* | -3.668 | 3.56E-04 | 0.009 |
| | | 37,886,749 | | *MIEN1* | 3.945 | 1.30E-04 | 0.002 |
| | | 37,896,063 | | | 3.476 | 6.92E-04 | 0.009 |
| | | 38,020,485 | *IKZF3* | | 3.962 | 1.22E-04 | 0.005 |
| | | 38,080,829 | | *ORMDL3* | 3.995 | 1.08E-04 | 0.014 |
| | | 38,096,333 | | | -3.664 | 3.60E-04 | 0.033 |
| | | 38,096,571 | | | -4.780 | 4.67E-06 | 0.033 |
| | | 38,096,578 | | | -5.642 | 1.00E-07 | 0.033 |
| | | 38,096,648 | | | -3.887 | 1.61E-04 | 0.033 |
| rs12949100 | g.38057189.A>G | 37,828,296 | | *PGAP3* | 3.531 | 5.73E-04 | 0.002 |
| | | 38,018,804 | | *IKZF3* | 3.442 | 7.77E-04 | 0.046 |
| | | 38,077,051 | | | 3.577 | 4.88E-04 | 0.034 |
| | | 38,077,191 | | | 4.035 | 9.28E-05 | 0.034 |
| | | 38,080,829 | | *ORMDL3* | 3.760 | 2.56E-04 | 0.014 |
| | | 38,081,829 | | *ORMDL3* | 3.975 | 1.16E-04 | 2.01E-04 |
| | | 38,081,995 | | *ORMDL3* | 5.846 | 3.84E-08 | 2.01E-04 |
| | | 38,082,022 | | *ORMDL3* | 6.706 | 5.54E-10 | 6.40E-04 |
| | | 38,082,035 | | *ORMDL3* | 6.270 | 4.94E-09 | 6.40E-04 |
| | | 38,082,073 | | *ORMDL3* | 6.100 | 1.13E-08 | 6.40E-04 |
| | | 38,082,206 | | *ORMDL3* | 3.773 | 2.44E-04 | 0.004 |
| | | 38,082,235 | | *ORMDL3* | 4.123 | 6.62E-05 | 0.004 |
| | | 38,082,346 | | *ORMDL3* | 4.378 | 2.44E-05 | 0.011 |
| | | 38,082,448 | | *ORMDL3* | 3.582 | 4.81E-04 | 0.011 |
| | | 38,082,658 | | *ORMDL3* | 4.530 | 1.32E-05 | 0.011 |
| | | 38,082,778 | | *ORMDL3* | 4.239 | 4.23E-05 | 0.011 |
| | | 38,108,710 | | | -4.859 | 3.34E-06 | 0.039 |
| | | 38,119,198 | *GSDMA* | | 4.218 | 4.59E-05 | 0.009 |
| | | 38,119,207 | *GSDMA* | | 5.390 | 3.21E-07 | 0.009 |
| | | 38,119,649 | | *GSDMA* | 4.712 | 6.21E-06 | 0.002 |
| | | 38,120,234 | | *GSDMA* | 3.566 | 5.08E-04 | 0.011 |
| rs8069176 | g.38057197.A>G | 37,828,296 | | *PGAP3* | 3.531 | 5.73E-04 | 0.002 |
| | | 38,018,804 | | *IKZF3* | 3.442 | 7.77E-04 | 0.046 |
| | | 38,077,051 | | | 3.577 | 4.88E-04 | 0.034 |

| | | | | | | | |
|---|---|---|---|---|---|---|---|
| | | 38,077,191 | | | 4.035 | 9.28E-05 | 0.034 |
| | | 38,080,829 | | *ORMDL3* | 3.760 | 2.56E-04 | 0.014 |
| | | 38,081,829 | | *ORMDL3* | 3.975 | 1.16E-04 | 2.01E-04 |
| | | 38,081,995 | | *ORMDL3* | 5.846 | 3.84E-08 | 2.01E-04 |
| | | 38,082,022 | | *ORMDL3* | 6.706 | 5.54E-10 | 6.40E-04 |
| | | 38,082,035 | | *ORMDL3* | 6.270 | 4.94E-09 | 6.40E-04 |
| | | 38,082,073 | | *ORMDL3* | 6.100 | 1.13E-08 | 6.40E-04 |
| | | 38,082,206 | | *ORMDL3* | 3.773 | 2.44E-04 | 0.004 |
| | | 38,082,235 | | *ORMDL3* | 4.123 | 6.62E-05 | 0.004 |
| | | 38,082,346 | | *ORMDL3* | 4.378 | 2.44E-05 | 0.011 |
| | | 38,082,448 | | *ORMDL3* | 3.582 | 4.81E-04 | 0.011 |
| | | 38,082,658 | | *ORMDL3* | 4.530 | 1.32E-05 | 0.011 |
| | | 38,082,778 | | *ORMDL3* | 4.239 | 4.23E-05 | 0.011 |
| | | 38,108,710 | | | -4.859 | 3.34E-06 | 0.039 |
| | | 38,119,198 | *GSDMA* | | 4.218 | 4.59E-05 | 0.009 |
| | | 38,119,207 | *GSDMA* | | 5.390 | 3.21E-07 | 0.009 |
| | | 38,119,649 | | *GSDMA* | 4.712 | 6.21E-06 | 0.002 |
| | | 38,120,234 | | *GSDMA* | 3.566 | 5.08E-04 | 0.011 |
| rs151192757 | g.38057566.T>C | 37,856,732 | | *ERBB2* | -3.668 | 3.56E-04 | 0.009 |
| | | 37,886,749 | | *MIEN1* | 3.945 | 1.30E-04 | 0.002 |
| | | 37,896,063 | | | 3.476 | 6.92E-04 | 0.009 |
| | | 38,020,485 | *IKZF3* | | 3.962 | 1.22E-04 | 0.005 |
| | | 38,080,829 | | *ORMDL3* | 3.995 | 1.08E-04 | 0.014 |
| | | 38,096,333 | | | -3.664 | 3.60E-04 | 0.033 |
| | | 38,096,571 | | | -4.780 | 4.67E-06 | 0.033 |
| | | 38,096,578 | | | -5.642 | 1.00E-07 | 0.033 |
| | | 38,096,648 | | | -3.887 | 1.61E-04 | 0.033 |
| rs113348879 | g.38057605.T>A | 37,856,732 | | *ERBB2* | -3.668 | 3.56E-04 | 0.009 |
| | | 37,886,749 | | *MIEN1* | 3.945 | 1.30E-04 | 0.002 |
| | | 37,896,063 | | | 3.476 | 6.92E-04 | 0.009 |
| | | 38,020,485 | *IKZF3* | | 3.962 | 1.22E-04 | 0.005 |
| | | 38,080,829 | | *ORMDL3* | 3.995 | 1.08E-04 | 0.014 |
| | | 38,096,333 | | | -3.664 | 3.60E-04 | 0.033 |
| | | 38,096,571 | | | -4.780 | 4.67E-06 | 0.033 |
| | | 38,096,578 | | | -5.642 | 1.00E-07 | 0.033 |
| | | 38,096,648 | | | -3.887 | 1.61E-04 | 0.033 |
| rs35503505 | g.38057780_38057783del | 37,910,902 | | | 3.424 | 8.26E-04 | 0.035 |
| | | 38,024,331 | | | -3.628 | 4.09E-04 | 0.017 |
| | | 38,082,073 | | *ORMDL3* | 3.925 | 1.40E-04 | 6.40E-04 |
| | | 38,108,710 | | | -4.522 | 1.37E-05 | 0.039 |

| | | | | | | | |
|---|---|---|---|---|---|---|---|
| | | 38,119,198 | *GSDMA* | | 4.158 | 5.79E-05 | 0.009 |
| | | 38,119,207 | *GSDMA* | | 4.858 | 3.35E-06 | 0.009 |
| | | 38,119,649 | | *GSDMA* | 4.851 | 3.46E-06 | 0.002 |
| | | 38,120,370 | | *GSDMA* | 3.466 | 7.16E-04 | 0.011 |
| rs11657449 | g.38057841.C>G | 38,024,331 | | | -3.737 | 2.78E-04 | 0.017 |
| | | 38,072,514 | | *GSDMB* | 3.655 | 3.72E-04 | 0.046 |
| | | 38,082,073 | | *ORMDL3* | 3.865 | 1.74E-04 | 6.40E-04 |
| | | 38,108,710 | | | -4.234 | 4.32E-05 | 0.039 |
| | | 38,119,198 | *GSDMA* | | 4.110 | 6.95E-05 | 0.009 |
| | | 38,119,207 | *GSDMA* | | 4.649 | 8.09E-06 | 0.009 |
| | | 38,119,649 | | *GSDMA* | 4.616 | 9.25E-06 | 0.002 |
| rs4795399 | g.38061439.C>T | 37,828,296 | | *PGAP3* | 3.529 | 5.77E-04 | 0.002 |
| | | 38,018,804 | | *IKZF3* | 3.445 | 7.69E-04 | 0.046 |
| | | 38,077,051 | | | 3.572 | 4.97E-04 | 0.034 |
| | | 38,077,191 | | | 4.013 | 1.01E-04 | 0.034 |
| | | 38,080,829 | | *ORMDL3* | 3.756 | 2.60E-04 | 0.014 |
| | | 38,081,829 | | *ORMDL3* | 3.981 | 1.13E-04 | 2.01E-04 |
| | | 38,081,995 | | *ORMDL3* | 5.846 | 3.84E-08 | 2.01E-04 |
| | | 38,082,022 | | *ORMDL3* | 6.711 | 5.39E-10 | 6.40E-04 |
| | | 38,082,035 | | *ORMDL3* | 6.267 | 4.99E-09 | 6.40E-04 |
| | | 38,082,073 | | *ORMDL3* | 6.097 | 1.15E-08 | 6.40E-04 |
| | | 38,082,206 | | *ORMDL3* | 3.756 | 2.60E-04 | 0.004 |
| | | 38,082,235 | | *ORMDL3* | 4.112 | 6.90E-05 | 0.004 |
| | | 38,082,346 | | *ORMDL3* | 4.364 | 2.58E-05 | 0.011 |
| | | 38,082,448 | | *ORMDL3* | 3.578 | 4.87E-04 | 0.011 |
| | | 38,082,658 | | *ORMDL3* | 4.529 | 1.32E-05 | 0.011 |
| | | 38,082,778 | | *ORMDL3* | 4.241 | 4.19E-05 | 0.011 |
| | | 38,108,710 | | | -4.853 | 3.43E-06 | 0.039 |
| | | 38,119,198 | *GSDMA* | | 4.226 | 4.44E-05 | 0.009 |
| | | 38,119,207 | *GSDMA* | | 5.401 | 3.05E-07 | 0.009 |
| | | 38,119,649 | | *GSDMA* | 4.702 | 6.48E-06 | 0.002 |
| | | 38,120,234 | | *GSDMA* | 3.575 | 4.93E-04 | 0.011 |
| rs2305480 | g.38062196.T>C (Pro.311.Ser) | 37,828,296 | | *PGAP3* | 3.529 | 5.77E-04 | 0.002 |
| | | 38,018,804 | | *IKZF3* | 3.445 | 7.69E-04 | 0.046 |
| | | 38,077,051 | | | 3.572 | 4.97E-04 | 0.034 |
| | | 38,077,191 | | | 4.013 | 1.01E-04 | 0.034 |
| | | 38,080,829 | | *ORMDL3* | 3.756 | 2.60E-04 | 0.014 |
| | | 38,081,829 | | *ORMDL3* | 3.981 | 1.13E-04 | 2.01E-04 |
| | | 38,081,995 | | *ORMDL3* | 5.846 | 3.84E-08 | 2.01E-04 |
| | | 38,082,022 | | *ORMDL3* | 6.711 | 5.39E-10 | 6.40E-04 |

| | | | | | | | |
|---|---|---|---|---|---|---|---|
| | | 38,082,035 | | *ORMDL3* | 6.267 | 4.99E-09 | 6.40E-04 |
| | | 38,082,073 | | *ORMDL3* | 6.097 | 1.15E-08 | 6.40E-04 |
| | | 38,082,206 | | *ORMDL3* | 3.756 | 2.60E-04 | 0.004 |
| | | 38,082,235 | | *ORMDL3* | 4.112 | 6.90E-05 | 0.004 |
| | | 38,082,346 | | *ORMDL3* | 4.364 | 2.58E-05 | 0.011 |
| | | 38,082,448 | | *ORMDL3* | 3.578 | 4.87E-04 | 0.011 |
| | | 38,082,658 | | *ORMDL3* | 4.529 | 1.32E-05 | 0.011 |
| | | 38,082,778 | | *ORMDL3* | 4.241 | 4.19E-05 | 0.011 |
| | | 38,108,710 | | | -4.853 | 3.43E-06 | 0.039 |
| | | 38,119,198 | *GSDMA* | | 4.226 | 4.44E-05 | 0.009 |
| | | 38,119,207 | *GSDMA* | | 5.401 | 3.05E-07 | 0.009 |
| | | 38,119,649 | | *GSDMA* | 4.702 | 6.48E-06 | 0.002 |
| | | 38,120,234 | | *GSDMA* | 3.575 | 4.93E-04 | 0.011 |
| rs2305479 | g.38062217.A>G (Gly.304.Arg) | 38,018,804 | | *IKZF3* | 4.097 | 7.31E-05 | 0.046 |
| | | 38,049,272 | | | 3.672 | 3.50E-04 | 0.046 |
| | | 38,077,191 | | | 3.770 | 2.47E-04 | 0.034 |
| | | 38,080,829 | | *ORMDL3* | 5.679 | 8.44E-08 | 0.014 |
| | | 38,081,995 | | *ORMDL3* | 4.445 | 1.87E-05 | 2.01E-04 |
| | | 38,082,022 | | *ORMDL3* | 4.985 | 1.94E-06 | 6.40E-04 |
| | | 38,082,035 | | *ORMDL3* | 5.617 | 1.13E-07 | 6.40E-04 |
| | | 38,082,073 | | *ORMDL3* | 5.181 | 8.20E-07 | 6.40E-04 |
| | | 38,082,235 | | *ORMDL3* | 3.771 | 2.46E-04 | 0.004 |
| | | 38,082,346 | | *ORMDL3* | 4.394 | 2.29E-05 | 0.011 |
| | | 38,082,448 | | *ORMDL3* | 4.059 | 8.46E-05 | 0.011 |
| | | 38,082,658 | | *ORMDL3* | 4.561 | 1.16E-05 | 0.011 |
| | | 38,082,778 | | *ORMDL3* | 3.728 | 2.87E-04 | 0.011 |
| | | 38,108,710 | | | -4.170 | 5.53E-05 | 0.039 |
| | | 38,119,198 | *GSDMA* | | 4.150 | 5.97E-05 | 0.009 |
| | | 38,119,207 | *GSDMA* | | 5.724 | 6.83E-08 | 0.009 |
| | | 38,119,649 | | *GSDMA* | 4.392 | 2.31E-05 | 0.002 |
| rs201674348 | g.38062297_38062297ins CTGCACTCATAGCA | 37,856,732 | | *ERBB2* | -3.668 | 3.56E-04 | 0.009 |
| | | 37,886,749 | | *MIEN1* | 3.945 | 1.30E-04 | 0.002 |
| | | 37,896,063 | | | 3.476 | 6.92E-04 | 0.009 |
| | | 38,020,485 | *IKZF3* | | 3.962 | 1.22E-04 | 0.005 |
| | | 38,080,829 | | *ORMDL3* | 3.995 | 1.08E-04 | 0.014 |
| | | 38,096,333 | | | -3.664 | 3.60E-04 | 0.033 |
| | | 38,096,571 | | | -4.780 | 4.67E-06 | 0.033 |
| | | 38,096,578 | | | -5.642 | 1.00E-07 | 0.033 |
| | | 38,096,648 | | | -3.887 | 1.61E-04 | 0.033 |

| | | | | | | | |
|---|---|---|---|---|---|---|---|
| rs35104165 | g.38062503.C>T (Asp.250.Gly) | 37,856,732 | | *ERBB2* | -3.668 | 3.56E-04 | 0.009 |
| | | 37,886,749 | | *MIEN1* | 3.945 | 1.30E-04 | 0.002 |
| | | 37,896,063 | | | 3.476 | 6.92E-04 | 0.009 |
| | | 38,020,485 | *IKZF3* | | 3.962 | 1.22E-04 | 0.005 |
| | | 38,080,829 | | *ORMDL3* | 3.995 | 1.08E-04 | 0.014 |
| | | 38,096,333 | | | -3.664 | 3.60E-04 | 0.033 |
| | | 38,096,571 | | | -4.780 | 4.67E-06 | 0.033 |
| | | 38,096,578 | | | -5.642 | 1.00E-07 | 0.033 |
| | | 38,096,648 | | | -3.887 | 1.61E-04 | 0.033 |
| rs111655187 | g.38062738.A>C | 37,856,732 | | *ERBB2* | -3.668 | 3.56E-04 | 0.009 |
| | | 37,886,749 | | *MIEN1* | 3.945 | 1.30E-04 | 0.002 |
| | | 37,896,063 | | | 3.476 | 6.92E-04 | 0.009 |
| | | 38,020,485 | *IKZF3* | | 3.962 | 1.22E-04 | 0.005 |
| | | 38,080,829 | | *ORMDL3* | 3.995 | 1.08E-04 | 0.014 |
| | | 38,096,333 | | | -3.664 | 3.60E-04 | 0.033 |
| | | 38,096,571 | | | -4.780 | 4.67E-06 | 0.033 |
| | | 38,096,578 | | | -5.642 | 1.00E-07 | 0.033 |
| | | 38,096,648 | | | -3.887 | 1.61E-04 | 0.033 |
| rs35196450 | g.38062942_38062942insC | 37,828,296 | | *PGAP3* | 3.529 | 5.77E-04 | 0.002 |
| | | 38,018,804 | | *IKZF3* | 3.445 | 7.69E-04 | 0.046 |
| | | 38,077,051 | | | 3.572 | 4.97E-04 | 0.034 |
| | | 38,077,191 | | | 4.013 | 1.01E-04 | 0.034 |
| | | 38,080,829 | | *ORMDL3* | 3.756 | 2.60E-04 | 0.014 |
| | | 38,081,829 | | *ORMDL3* | 3.981 | 1.13E-04 | 2.01E-04 |
| | | 38,081,995 | | *ORMDL3* | 5.846 | 3.84E-08 | 2.01E-04 |
| | | 38,082,022 | | *ORMDL3* | 6.711 | 5.39E-10 | 6.40E-04 |
| | | 38,082,035 | | *ORMDL3* | 6.267 | 4.99E-09 | 6.40E-04 |
| | | 38,082,073 | | *ORMDL3* | 6.097 | 1.15E-08 | 6.40E-04 |
| | | 38,082,206 | | *ORMDL3* | 3.756 | 2.60E-04 | 0.004 |
| | | 38,082,235 | | *ORMDL3* | 4.112 | 6.90E-05 | 0.004 |
| | | 38,082,346 | | *ORMDL3* | 4.364 | 2.58E-05 | 0.011 |
| | | 38,082,448 | | *ORMDL3* | 3.578 | 4.87E-04 | 0.011 |
| | | 38,082,658 | | *ORMDL3* | 4.529 | 1.32E-05 | 0.011 |
| | | 38,082,778 | | *ORMDL3* | 4.241 | 4.19E-05 | 0.011 |
| | | 38,108,710 | | | -4.853 | 3.43E-06 | 0.039 |
| | | 38,119,198 | *GSDMA* | | 4.226 | 4.44E-05 | 0.009 |
| | | 38,119,207 | *GSDMA* | | 5.401 | 3.05E-07 | 0.009 |
| | | 38,119,649 | | *GSDMA* | 4.702 | 6.48E-06 | 0.002 |
| | | 38,120,234 | | *GSDMA* | 3.575 | 4.93E-04 | 0.011 |

| | | | | | | | |
|---|---|---|---|---|---|---|---|
| rs56750287 | g.38062944.C>A | 37,828,296 | | *PGAP3* | 3.529 | 5.77E-04 | 0.002 |
| | | 38,018,804 | | *IKZF3* | 3.445 | 7.69E-04 | 0.046 |
| | | 38,077,051 | | | 3.572 | 4.97E-04 | 0.034 |
| | | 38,077,191 | | | 4.013 | 1.01E-04 | 0.034 |
| | | 38,080,829 | | *ORMDL3* | 3.756 | 2.60E-04 | 0.014 |
| | | 38,081,829 | | *ORMDL3* | 3.981 | 1.13E-04 | 2.01E-04 |
| | | 38,081,995 | | *ORMDL3* | 5.846 | 3.84E-08 | 2.01E-04 |
| | | 38,082,022 | | *ORMDL3* | 6.711 | 5.39E-10 | 6.40E-04 |
| | | 38,082,035 | | *ORMDL3* | 6.267 | 4.99E-09 | 6.40E-04 |
| | | 38,082,073 | | *ORMDL3* | 6.097 | 1.15E-08 | 6.40E-04 |
| | | 38,082,206 | | *ORMDL3* | 3.756 | 2.60E-04 | 0.004 |
| | | 38,082,235 | | *ORMDL3* | 4.112 | 6.90E-05 | 0.004 |
| | | 38,082,346 | | *ORMDL3* | 4.364 | 2.58E-05 | 0.011 |
| | | 38,082,448 | | *ORMDL3* | 3.578 | 4.87E-04 | 0.011 |
| | | 38,082,658 | | *ORMDL3* | 4.529 | 1.32E-05 | 0.011 |
| | | 38,082,778 | | *ORMDL3* | 4.241 | 4.19E-05 | 0.011 |
| | | 38,108,710 | | | -4.853 | 3.43E-06 | 0.039 |
| | | 38,119,198 | *GSDMA* | | 4.226 | 4.44E-05 | 0.009 |
| | | 38,119,207 | *GSDMA* | | 5.401 | 3.05E-07 | 0.009 |
| | | 38,119,649 | | *GSDMA* | 4.702 | 6.48E-06 | 0.002 |
| | | 38,120,234 | | *GSDMA* | 3.575 | 4.93E-04 | 0.011 |
| rs11078926 | g.38062976.A>G | 37,828,296 | | *PGAP3* | 3.529 | 5.77E-04 | 0.002 |
| | | 38,018,804 | | *IKZF3* | 3.445 | 7.69E-04 | 0.046 |
| | | 38,077,051 | | | 3.572 | 4.97E-04 | 0.034 |
| | | 38,077,191 | | | 4.013 | 1.01E-04 | 0.034 |
| | | 38,080,829 | | *ORMDL3* | 3.756 | 2.60E-04 | 0.014 |
| | | 38,081,829 | | *ORMDL3* | 3.981 | 1.13E-04 | 2.01E-04 |
| | | 38,081,995 | | *ORMDL3* | 5.846 | 3.84E-08 | 2.01E-04 |
| | | 38,082,022 | | *ORMDL3* | 6.711 | 5.39E-10 | 6.40E-04 |
| | | 38,082,035 | | *ORMDL3* | 6.267 | 4.99E-09 | 6.40E-04 |
| | | 38,082,073 | | *ORMDL3* | 6.097 | 1.15E-08 | 6.40E-04 |
| | | 38,082,206 | | *ORMDL3* | 3.756 | 2.60E-04 | 0.004 |
| | | 38,082,235 | | *ORMDL3* | 4.112 | 6.90E-05 | 0.004 |
| | | 38,082,346 | | *ORMDL3* | 4.364 | 2.58E-05 | 0.011 |
| | | 38,082,448 | | *ORMDL3* | 3.578 | 4.87E-04 | 0.011 |
| | | 38,082,658 | | *ORMDL3* | 4.529 | 1.32E-05 | 0.011 |
| | | 38,082,778 | | *ORMDL3* | 4.241 | 4.19E-05 | 0.011 |
| | | 38,108,710 | | | -4.853 | 3.43E-06 | 0.039 |
| | | 38,119,198 | *GSDMA* | | 4.226 | 4.44E-05 | 0.009 |
| | | 38,119,207 | *GSDMA* | | 5.401 | 3.05E-07 | 0.009 |
| | | 38,119,649 | | *GSDMA* | 4.702 | 6.48E-06 | 0.002 |

| | | | | | | | |
|---|---|---|---|---|---|---|---|
| | | 38,120,234 | | *GSDMA* | 3.575 | 4.93E-04 | 0.011 |
| rs883770 | g.38063381.T>C | 38,018,804 | | *IKZF3* | 4.097 | 7.31E-05 | 0.046 |
| | | 38,049,272 | | | 3.672 | 3.50E-04 | 0.046 |
| | | 38,077,191 | | | 3.770 | 2.47E-04 | 0.034 |
| | | 38,080,829 | | *ORMDL3* | 5.679 | 8.44E-08 | 0.014 |
| | | 38,081,995 | | *ORMDL3* | 4.445 | 1.87E-05 | 2.01E-04 |
| | | 38,082,022 | | *ORMDL3* | 4.985 | 1.94E-06 | 6.40E-04 |
| | | 38,082,035 | | *ORMDL3* | 5.617 | 1.13E-07 | 6.40E-04 |
| | | 38,082,073 | | *ORMDL3* | 5.181 | 8.20E-07 | 6.40E-04 |
| | | 38,082,235 | | *ORMDL3* | 3.771 | 2.46E-04 | 0.004 |
| | | 38,082,346 | | *ORMDL3* | 4.394 | 2.29E-05 | 0.011 |
| | | 38,082,448 | | *ORMDL3* | 4.059 | 8.46E-05 | 0.011 |
| | | 38,082,658 | | *ORMDL3* | 4.561 | 1.16E-05 | 0.011 |
| | | 38,082,778 | | *ORMDL3* | 3.728 | 2.87E-04 | 0.011 |
| | | 38,108,710 | | | -4.170 | 5.53E-05 | 0.039 |
| | | 38,119,198 | *GSDMA* | | 4.150 | 5.97E-05 | 0.009 |
| | | 38,119,207 | *GSDMA* | | 5.724 | 6.83E-08 | 0.009 |
| | | 38,119,649 | | *GSDMA* | 4.392 | 2.31E-05 | 0.002 |
| rs62067034 | g.38063738.T>C | 38,018,804 | | *IKZF3* | 4.097 | 7.31E-05 | 0.046 |
| | | 38,049,272 | | | 3.672 | 3.50E-04 | 0.046 |
| | | 38,077,191 | | | 3.770 | 2.47E-04 | 0.034 |
| | | 38,080,829 | | *ORMDL3* | 5.679 | 8.44E-08 | 0.014 |
| | | 38,081,995 | | *ORMDL3* | 4.445 | 1.87E-05 | 2.01E-04 |
| | | 38,082,022 | | *ORMDL3* | 4.985 | 1.94E-06 | 6.40E-04 |
| | | 38,082,035 | | *ORMDL3* | 5.617 | 1.13E-07 | 6.40E-04 |
| | | 38,082,073 | | *ORMDL3* | 5.181 | 8.20E-07 | 6.40E-04 |
| | | 38,082,235 | | *ORMDL3* | 3.771 | 2.46E-04 | 0.004 |
| | | 38,082,346 | | *ORMDL3* | 4.394 | 2.29E-05 | 0.011 |
| | | 38,082,448 | | *ORMDL3* | 4.059 | 8.46E-05 | 0.011 |
| | | 38,082,658 | | *ORMDL3* | 4.561 | 1.16E-05 | 0.011 |
| | | 38,082,778 | | *ORMDL3* | 3.728 | 2.87E-04 | 0.011 |
| | | 38,108,710 | | | -4.170 | 5.53E-05 | 0.039 |
| | | 38,119,198 | *GSDMA* | | 4.150 | 5.97E-05 | 0.009 |
| | | 38,119,207 | *GSDMA* | | 5.724 | 6.83E-08 | 0.009 |
| | | 38,119,649 | | *GSDMA* | 4.392 | 2.31E-05 | 0.002 |
| rs12949547 | g.38063826.A>C | 37,856,732 | | *ERBB2* | -3.668 | 3.56E-04 | 0.009 |
| | | 37,886,749 | | *MIEN1* | 3.945 | 1.30E-04 | 0.002 |
| | | 37,896,063 | | | 3.476 | 6.92E-04 | 0.009 |
| | | 38,020,485 | *IKZF3* | | 3.962 | 1.22E-04 | 0.005 |
| | | 38,080,829 | | *ORMDL3* | 3.995 | 1.08E-04 | 0.014 |
| | | 38,096,333 | | | -3.664 | 3.60E-04 | 0.033 |

| | | | | | | | |
|---|---|---|---|---|---|---|---|
| | | 38,096,571 | | | -4.780 | 4.67E-06 | 0.033 |
| | | 38,096,578 | | | -5.642 | 1.00E-07 | 0.033 |
| | | 38,096,648 | | | -3.887 | 1.61E-04 | 0.033 |
| rs113263629 | g.38063849.T>A | 37,856,732 | | *ERBB2* | -3.668 | 3.56E-04 | 0.009 |
| | | 37,886,749 | | *MIEN1* | 3.945 | 1.30E-04 | 0.002 |
| | | 37,896,063 | | | 3.476 | 6.92E-04 | 0.009 |
| | | 38,020,485 | *IKZF3* | | 3.962 | 1.22E-04 | 0.005 |
| | | 38,080,829 | | *ORMDL3* | 3.995 | 1.08E-04 | 0.014 |
| | | 38,096,333 | | | -3.664 | 3.60E-04 | 0.033 |
| | | 38,096,571 | | | -4.780 | 4.67E-06 | 0.033 |
| | | 38,096,578 | | | -5.642 | 1.00E-07 | 0.033 |
| | | 38,096,648 | | | -3.887 | 1.61E-04 | 0.033 |
| rs36000226 | g.38063929.C>T | 38,018,804 | | *IKZF3* | 4.097 | 7.31E-05 | 0.046 |
| | | 38,049,272 | | | 3.672 | 3.50E-04 | 0.046 |
| | | 38,077,191 | | | 3.770 | 2.47E-04 | 0.034 |
| | | 38,080,829 | | *ORMDL3* | 5.679 | 8.44E-08 | 0.014 |
| | | 38,081,995 | | *ORMDL3* | 4.445 | 1.87E-05 | 2.01E-04 |
| | | 38,082,022 | | *ORMDL3* | 4.985 | 1.94E-06 | 6.40E-04 |
| | | 38,082,035 | | *ORMDL3* | 5.617 | 1.13E-07 | 6.40E-04 |
| | | 38,082,073 | | *ORMDL3* | 5.181 | 8.20E-07 | 6.40E-04 |
| | | 38,082,235 | | *ORMDL3* | 3.771 | 2.46E-04 | 0.004 |
| | | 38,082,346 | | *ORMDL3* | 4.394 | 2.29E-05 | 0.011 |
| | | 38,082,448 | | *ORMDL3* | 4.059 | 8.46E-05 | 0.011 |
| | | 38,082,658 | | *ORMDL3* | 4.561 | 1.16E-05 | 0.011 |
| | | 38,082,778 | | *ORMDL3* | 3.728 | 2.87E-04 | 0.011 |
| | | 38,108,710 | | | -4.170 | 5.53E-05 | 0.039 |
| | | 38,119,198 | *GSDMA* | | 4.150 | 5.97E-05 | 0.009 |
| | | 38,119,207 | *GSDMA* | | 5.724 | 6.83E-08 | 0.009 |
| | | 38,119,649 | | *GSDMA* | 4.392 | 2.31E-05 | 0.002 |
| rs36084703 | g.38063980_38063981del | 38,018,804 | | *IKZF3* | 4.097 | 7.31E-05 | 0.046 |
| | | 38,049,272 | | | 3.672 | 3.50E-04 | 0.046 |
| | | 38,077,191 | | | 3.770 | 2.47E-04 | 0.034 |
| | | 38,080,829 | | *ORMDL3* | 5.679 | 8.44E-08 | 0.014 |
| | | 38,081,995 | | *ORMDL3* | 4.445 | 1.87E-05 | 2.01E-04 |
| | | 38,082,022 | | *ORMDL3* | 4.985 | 1.94E-06 | 6.40E-04 |
| | | 38,082,035 | | *ORMDL3* | 5.617 | 1.13E-07 | 6.40E-04 |
| | | 38,082,073 | | *ORMDL3* | 5.181 | 8.20E-07 | 6.40E-04 |
| | | 38,082,235 | | *ORMDL3* | 3.771 | 2.46E-04 | 0.004 |
| | | 38,082,346 | | *ORMDL3* | 4.394 | 2.29E-05 | 0.011 |
| | | 38,082,448 | | *ORMDL3* | 4.059 | 8.46E-05 | 0.011 |

| | | | | | | | |
|---|---|---|---|---|---|---|---|
| | | 38,082,658 | | *ORMDL3* | 4.561 | 1.16E-05 | 0.011 |
| | | 38,082,778 | | *ORMDL3* | 3.728 | 2.87E-04 | 0.011 |
| | | 38,108,710 | | | -4.170 | 5.53E-05 | 0.039 |
| | | 38,119,198 | *GSDMA* | | 4.150 | 5.97E-05 | 0.009 |
| | | 38,119,207 | *GSDMA* | | 5.724 | 6.83E-08 | 0.009 |
| | | 38,119,649 | | *GSDMA* | 4.392 | 2.31E-05 | 0.002 |
| rs11078927 | g.38064405.T>C | 37,828,296 | | *PGAP3* | 3.529 | 5.77E-04 | 0.002 |
| | | 38,018,804 | | *IKZF3* | 3.445 | 7.69E-04 | 0.046 |
| | | 38,077,051 | | | 3.572 | 4.97E-04 | 0.034 |
| | | 38,077,191 | | | 4.013 | 1.01E-04 | 0.034 |
| | | 38,080,829 | | *ORMDL3* | 3.756 | 2.60E-04 | 0.014 |
| | | 38,081,829 | | *ORMDL3* | 3.981 | 1.13E-04 | 2.01E-04 |
| | | 38,081,995 | | *ORMDL3* | 5.846 | 3.84E-08 | 2.01E-04 |
| | | 38,082,022 | | *ORMDL3* | 6.711 | 5.39E-10 | 6.40E-04 |
| | | 38,082,035 | | *ORMDL3* | 6.267 | 4.99E-09 | 6.40E-04 |
| | | 38,082,073 | | *ORMDL3* | 6.097 | 1.15E-08 | 6.40E-04 |
| | | 38,082,206 | | *ORMDL3* | 3.756 | 2.60E-04 | 0.004 |
| | | 38,082,235 | | *ORMDL3* | 4.112 | 6.90E-05 | 0.004 |
| | | 38,082,346 | | *ORMDL3* | 4.364 | 2.58E-05 | 0.011 |
| | | 38,082,448 | | *ORMDL3* | 3.578 | 4.87E-04 | 0.011 |
| | | 38,082,658 | | *ORMDL3* | 4.529 | 1.32E-05 | 0.011 |
| | | 38,082,778 | | *ORMDL3* | 4.241 | 4.19E-05 | 0.011 |
| | | 38,108,710 | | | -4.853 | 3.43E-06 | 0.039 |
| | | 38,119,198 | *GSDMA* | | 4.226 | 4.44E-05 | 0.009 |
| | | 38,119,207 | *GSDMA* | | 5.401 | 3.05E-07 | 0.009 |
| | | 38,119,649 | | *GSDMA* | 4.702 | 6.48E-06 | 0.002 |
| | | 38,120,234 | | *GSDMA* | 3.575 | 4.93E-04 | 0.011 |
| rs11078928 | g.38064469.C>T<br>(Leu.224.Leu) | 37,828,296 | | *PGAP3* | 3.529 | 5.77E-04 | 0.002 |
| | | 38,018,804 | | *IKZF3* | 3.445 | 7.69E-04 | 0.046 |
| | | 38,077,051 | | | 3.572 | 4.97E-04 | 0.034 |
| | | 38,077,191 | | | 4.013 | 1.01E-04 | 0.034 |
| | | 38,080,829 | | *ORMDL3* | 3.756 | 2.60E-04 | 0.014 |
| | | 38,081,829 | | *ORMDL3* | 3.981 | 1.13E-04 | 2.01E-04 |
| | | 38,081,995 | | *ORMDL3* | 5.846 | 3.84E-08 | 2.01E-04 |
| | | 38,082,022 | | *ORMDL3* | 6.711 | 5.39E-10 | 6.40E-04 |
| | | 38,082,035 | | *ORMDL3* | 6.267 | 4.99E-09 | 6.40E-04 |
| | | 38,082,073 | | *ORMDL3* | 6.097 | 1.15E-08 | 6.40E-04 |
| | | 38,082,206 | | *ORMDL3* | 3.756 | 2.60E-04 | 0.004 |
| | | 38,082,235 | | *ORMDL3* | 4.112 | 6.90E-05 | 0.004 |
| | | 38,082,346 | | *ORMDL3* | 4.364 | 2.58E-05 | 0.011 |

| | | | | | | | |
|---|---|---|---|---|---|---|---|
| | | 38,082,448 | | *ORMDL3* | 3.578 | 4.87E-04 | 0.011 |
| | | 38,082,658 | | *ORMDL3* | 4.529 | 1.32E-05 | 0.011 |
| | | 38,082,778 | | *ORMDL3* | 4.241 | 4.19E-05 | 0.011 |
| | | 38,108,710 | | | -4.853 | 3.43E-06 | 0.039 |
| | | 38,119,198 | *GSDMA* | | 4.226 | 4.44E-05 | 0.009 |
| | | 38,119,207 | *GSDMA* | | 5.401 | 3.05E-07 | 0.009 |
| | | 38,119,649 | | *GSDMA* | 4.702 | 6.48E-06 | 0.002 |
| | | 38,120,234 | | *GSDMA* | 3.575 | 4.93E-04 | 0.011 |
| rs111826178 | g.38064924.T>C | 37,856,732 | | *ERBB2* | -3.668 | 3.56E-04 | 0.009 |
| | | 37,886,749 | | *MIEN1* | 3.945 | 1.30E-04 | 0.002 |
| | | 37,896,063 | | | 3.476 | 6.92E-04 | 0.009 |
| | | 38,020,485 | *IKZF3* | | 3.962 | 1.22E-04 | 0.005 |
| | | 38,080,829 | | *ORMDL3* | 3.995 | 1.08E-04 | 0.014 |
| | | 38,096,333 | | | -3.664 | 3.60E-04 | 0.033 |
| | | 38,096,571 | | | -4.780 | 4.67E-06 | 0.033 |
| | | 38,096,578 | | | -5.642 | 1.00E-07 | 0.033 |
| | | 38,096,648 | | | -3.887 | 1.61E-04 | 0.033 |
| rs113657930 | g.38065485.G>C | 37,856,732 | | *ERBB2* | -3.668 | 3.56E-04 | 0.009 |
| | | 37,886,749 | | *MIEN1* | 3.945 | 1.30E-04 | 0.002 |
| | | 37,896,063 | | | 3.476 | 6.92E-04 | 0.009 |
| | | 38,020,485 | *IKZF3* | | 3.962 | 1.22E-04 | 0.005 |
| | | 38,080,829 | | *ORMDL3* | 3.995 | 1.08E-04 | 0.014 |
| | | 38,096,333 | | | -3.664 | 3.60E-04 | 0.033 |
| | | 38,096,571 | | | -4.780 | 4.67E-06 | 0.033 |
| | | 38,096,578 | | | -5.642 | 1.00E-07 | 0.033 |
| | | 38,096,648 | | | -3.887 | 1.61E-04 | 0.033 |
| rs2290400 | g.38066240.G>A | 38,018,804 | | *IKZF3* | 4.097 | 7.31E-05 | 0.046 |
| | | 38,049,272 | | | 3.672 | 3.50E-04 | 0.046 |
| | | 38,077,191 | | | 3.770 | 2.47E-04 | 0.034 |
| | | 38,080,829 | | *ORMDL3* | 5.679 | 8.44E-08 | 0.014 |
| | | 38,081,995 | | *ORMDL3* | 4.445 | 1.87E-05 | 2.01E-04 |
| | | 38,082,022 | | *ORMDL3* | 4.985 | 1.94E-06 | 6.40E-04 |
| | | 38,082,035 | | *ORMDL3* | 5.617 | 1.13E-07 | 6.40E-04 |
| | | 38,082,073 | | *ORMDL3* | 5.181 | 8.20E-07 | 6.40E-04 |
| | | 38,082,235 | | *ORMDL3* | 3.771 | 2.46E-04 | 0.004 |
| | | 38,082,346 | | *ORMDL3* | 4.394 | 2.29E-05 | 0.011 |
| | | 38,082,448 | | *ORMDL3* | 4.059 | 8.46E-05 | 0.011 |
| | | 38,082,658 | | *ORMDL3* | 4.561 | 1.16E-05 | 0.011 |
| | | 38,082,778 | | *ORMDL3* | 3.728 | 2.87E-04 | 0.011 |
| | | 38,108,710 | | | -4.170 | 5.53E-05 | 0.039 |
| | | 38,119,198 | *GSDMA* | | 4.150 | 5.97E-05 | 0.009 |

| | | | | | | | |
|---|---|---|---|---|---|---|---|
| | | 38,119,207 | *GSDMA* | | 5.724 | 6.83E-08 | 0.009 |
| | | 38,119,649 | | *GSDMA* | 4.392 | 2.31E-05 | 0.002 |
| rs1008723 | g.38066267.T>G | 38,018,804 | | *IKZF3* | 4.097 | 7.31E-05 | 0.046 |
| | | 38,049,272 | | | 3.672 | 3.50E-04 | 0.046 |
| | | 38,077,191 | | | 3.770 | 2.47E-04 | 0.034 |
| | | 38,080,829 | | *ORMDL3* | 5.679 | 8.44E-08 | 0.014 |
| | | 38,081,995 | | *ORMDL3* | 4.445 | 1.87E-05 | 2.01E-04 |
| | | 38,082,022 | | *ORMDL3* | 4.985 | 1.94E-06 | 6.40E-04 |
| | | 38,082,035 | | *ORMDL3* | 5.617 | 1.13E-07 | 6.40E-04 |
| | | 38,082,073 | | *ORMDL3* | 5.181 | 8.20E-07 | 6.40E-04 |
| | | 38,082,235 | | *ORMDL3* | 3.771 | 2.46E-04 | 0.004 |
| | | 38,082,346 | | *ORMDL3* | 4.394 | 2.29E-05 | 0.011 |
| | | 38,082,448 | | *ORMDL3* | 4.059 | 8.46E-05 | 0.011 |
| | | 38,082,658 | | *ORMDL3* | 4.561 | 1.16E-05 | 0.011 |
| | | 38,082,778 | | *ORMDL3* | 3.728 | 2.87E-04 | 0.011 |
| | | 38,108,710 | | | -4.170 | 5.53E-05 | 0.039 |
| | | 38,119,198 | *GSDMA* | | 4.150 | 5.97E-05 | 0.009 |
| | | 38,119,207 | *GSDMA* | | 5.724 | 6.83E-08 | 0.009 |
| | | 38,119,649 | | *GSDMA* | 4.392 | 2.31E-05 | 0.002 |
| rs56380902 | g.38066372.C>T | 38,018,804 | | *IKZF3* | 4.097 | 7.31E-05 | 0.046 |
| | | 38,049,272 | | | 3.672 | 3.50E-04 | 0.046 |
| | | 38,077,191 | | | 3.770 | 2.47E-04 | 0.034 |
| | | 38,080,829 | | *ORMDL3* | 5.679 | 8.44E-08 | 0.014 |
| | | 38,081,995 | | *ORMDL3* | 4.445 | 1.87E-05 | 2.01E-04 |
| | | 38,082,022 | | *ORMDL3* | 4.985 | 1.94E-06 | 6.40E-04 |
| | | 38,082,035 | | *ORMDL3* | 5.617 | 1.13E-07 | 6.40E-04 |
| | | 38,082,073 | | *ORMDL3* | 5.181 | 8.20E-07 | 6.40E-04 |
| | | 38,082,235 | | *ORMDL3* | 3.771 | 2.46E-04 | 0.004 |
| | | 38,082,346 | | *ORMDL3* | 4.394 | 2.29E-05 | 0.011 |
| | | 38,082,448 | | *ORMDL3* | 4.059 | 8.46E-05 | 0.011 |
| | | 38,082,658 | | *ORMDL3* | 4.561 | 1.16E-05 | 0.011 |
| | | 38,082,778 | | *ORMDL3* | 3.728 | 2.87E-04 | 0.011 |
| | | 38,108,710 | | | -4.170 | 5.53E-05 | 0.039 |
| | | 38,119,198 | *GSDMA* | | 4.150 | 5.97E-05 | 0.009 |
| | | 38,119,207 | *GSDMA* | | 5.724 | 6.83E-08 | 0.009 |
| | | 38,119,649 | | *GSDMA* | 4.392 | 2.31E-05 | 0.002 |
| rs4795400 | g.38067020.T>C | 37,828,296 | | *PGAP3* | 3.529 | 5.77E-04 | 0.002 |
| | | 38,018,804 | | *IKZF3* | 3.445 | 7.69E-04 | 0.046 |
| | | 38,077,051 | | | 3.572 | 4.97E-04 | 0.034 |
| | | 38,077,191 | | | 4.013 | 1.01E-04 | 0.034 |
| | | 38,080,829 | | *ORMDL3* | 3.756 | 2.60E-04 | 0.014 |

| | | | | | | | |
|---|---|---|---|---|---|---|---|
| | | 38,081,829 | | *ORMDL3* | 3.981 | 1.13E-04 | 2.01E-04 |
| | | 38,081,995 | | *ORMDL3* | 5.846 | 3.84E-08 | 2.01E-04 |
| | | 38,082,022 | | *ORMDL3* | 6.711 | 5.39E-10 | 6.40E-04 |
| | | 38,082,035 | | *ORMDL3* | 6.267 | 4.99E-09 | 6.40E-04 |
| | | 38,082,073 | | *ORMDL3* | 6.097 | 1.15E-08 | 6.40E-04 |
| | | 38,082,206 | | *ORMDL3* | 3.756 | 2.60E-04 | 0.004 |
| | | 38,082,235 | | *ORMDL3* | 4.112 | 6.90E-05 | 0.004 |
| | | 38,082,346 | | *ORMDL3* | 4.364 | 2.58E-05 | 0.011 |
| | | 38,082,448 | | *ORMDL3* | 3.578 | 4.87E-04 | 0.011 |
| | | 38,082,658 | | *ORMDL3* | 4.529 | 1.32E-05 | 0.011 |
| | | 38,082,778 | | *ORMDL3* | 4.241 | 4.19E-05 | 0.011 |
| | | 38,108,710 | | | -4.853 | 3.43E-06 | 0.039 |
| | | 38,119,198 | *GSDMA* | | 4.226 | 4.44E-05 | 0.009 |
| | | 38,119,207 | *GSDMA* | | 5.401 | 3.05E-07 | 0.009 |
| | | 38,119,649 | | *GSDMA* | 4.702 | 6.48E-06 | 0.002 |
| | | 38,120,234 | | *GSDMA* | 3.575 | 4.93E-04 | 0.011 |
| rs183631218 | g.38067532.C>T | 37,856,732 | | *ERBB2* | -3.668 | 3.56E-04 | 0.009 |
| | | 37,886,749 | | *MIEN1* | 3.945 | 1.30E-04 | 0.002 |
| | | 37,896,063 | | | 3.476 | 6.92E-04 | 0.009 |
| | | 38,020,485 | *IKZF3* | | 3.962 | 1.22E-04 | 0.005 |
| | | 38,080,829 | | *ORMDL3* | 3.995 | 1.08E-04 | 0.014 |
| | | 38,096,333 | | | -3.664 | 3.60E-04 | 0.033 |
| | | 38,096,571 | | | -4.780 | 4.67E-06 | 0.033 |
| | | 38,096,578 | | | -5.642 | 1.00E-07 | 0.033 |
| | | 38,096,648 | | | -3.887 | 1.61E-04 | 0.033 |
| rs4795401 | g.38067533.G>A | 38,018,804 | | *IKZF3* | 4.097 | 7.31E-05 | 0.046 |
| | | 38,049,272 | | | 3.672 | 3.50E-04 | 0.046 |
| | | 38,077,191 | | | 3.770 | 2.47E-04 | 0.034 |
| | | 38,080,829 | | *ORMDL3* | 5.679 | 8.44E-08 | 0.014 |
| | | 38,081,995 | | *ORMDL3* | 4.445 | 1.87E-05 | 2.01E-04 |
| | | 38,082,022 | | *ORMDL3* | 4.985 | 1.94E-06 | 6.40E-04 |
| | | 38,082,035 | | *ORMDL3* | 5.617 | 1.13E-07 | 6.40E-04 |
| | | 38,082,073 | | *ORMDL3* | 5.181 | 8.20E-07 | 6.40E-04 |
| | | 38,082,235 | | *ORMDL3* | 3.771 | 2.46E-04 | 0.004 |
| | | 38,082,346 | | *ORMDL3* | 4.394 | 2.29E-05 | 0.011 |
| | | 38,082,448 | | *ORMDL3* | 4.059 | 8.46E-05 | 0.011 |
| | | 38,082,658 | | *ORMDL3* | 4.561 | 1.16E-05 | 0.011 |
| | | 38,082,778 | | *ORMDL3* | 3.728 | 2.87E-04 | 0.011 |
| | | 38,108,710 | | | -4.170 | 5.53E-05 | 0.039 |
| | | 38,119,198 | *GSDMA* | | 4.150 | 5.97E-05 | 0.009 |
| | | 38,119,207 | *GSDMA* | | 5.724 | 6.83E-08 | 0.009 |

| | | | | | | | | | |
|---|---|---|---|---|---|---|---|---|---|
| | | | 38,119,649 | | *GSDMA* | 4.392 | 2.31E-05 | | 0.002 |
| | rs869402 | g.38068043.T>C | 37,887,061 | *MIEN1* | | 3.704 | 3.12E-04 | 5.35E-04 | 0.002 |
| | | | 38,018,804 | | *IKZF3* | 3.417 | 8.44E-04 | 5.35E-04 | 0.046 |
| | | | 38,077,191 | | | 3.827 | 2.00E-04 | 5.35E-04 | 0.034 |
| | | | 38,077,464 | | *ORMDL3* | 3.639 | 3.93E-04 | 5.35E-04 | 0.014 |
| | | | 38,080,829 | | *ORMDL3* | 5.536 | 1.64E-07 | 5.35E-04 | 0.014 |
| | | | 38,081,492 | | *ORMDL3* | 3.585 | 4.75E-04 | 5.35E-04 | 2.01E-04 |
| | | | 38,081,995 | | *ORMDL3* | 4.416 | 2.10E-05 | 5.35E-04 | 2.01E-04 |
| | | | 38,082,022 | | *ORMDL3* | 5.112 | 1.11E-06 | 5.35E-04 | 6.40E-04 |
| | | | 38,082,035 | | *ORMDL3* | 5.307 | 4.66E-07 | 5.35E-04 | 6.40E-04 |
| | | | 38,082,073 | | *ORMDL3* | 5.048 | 1.47E-06 | 5.35E-04 | 6.40E-04 |
| | | | 38,082,235 | | *ORMDL3* | 4.077 | 7.89E-05 | 5.35E-04 | 0.004 |
| | | | 38,082,346 | | *ORMDL3* | 4.493 | 1.53E-05 | 5.35E-04 | 0.011 |
| | | | 38,082,448 | | *ORMDL3* | 3.848 | 1.86E-04 | 5.35E-04 | 0.011 |
| | | | 38,082,658 | | *ORMDL3* | 5.521 | 1.76E-07 | 5.35E-04 | 0.011 |
| | | | 38,108,464 | | | -3.396 | 9.08E-04 | 5.35E-04 | 0.007 |
| | | | 38,108,710 | | | -4.944 | 2.31E-06 | 5.35E-04 | 0.039 |
| | | | 38,119,198 | *GSDMA* | | 4.736 | 5.63E-06 | 5.35E-04 | 0.009 |
| | | | 38,119,207 | *GSDMA* | | 6.586 | 1.02E-09 | 5.35E-04 | 0.009 |
| | | | 38,119,649 | | *GSDMA* | 5.298 | 4.85E-07 | 5.35E-04 | 0.002 |
| | | | 38,119,760 | | *GSDMA* | 5.134 | 1.01E-06 | 5.35E-04 | 0.011 |
| Fifth block | rs870829 | g.38068382.T>G | 38,024,048 | | | -3.457 | 7.39E-04 | 7.23E-04 | 0.004 |
| | | | 38,024,063 | | | -3.508 | 6.22E-04 | 7.23E-04 | 0.004 |
| | | | 38,024,146 | | | -3.530 | 5.75E-04 | 7.23E-04 | 0.004 |
| | | | 38,024,331 | | | -3.386 | 9.39E-04 | 7.23E-04 | 0.017 |
| | | | 38,077,464 | | *ORMDL3* | 4.137 | 6.28E-05 | 7.23E-04 | 0.014 |
| | | | 38,080,829 | | *ORMDL3* | 4.882 | 3.02E-06 | 7.23E-04 | 0.014 |
| | | | 38,108,464 | | | -3.472 | 7.01E-04 | 7.23E-04 | 0.007 |
| | | | 38,108,710 | | | -4.310 | 3.19E-05 | 7.23E-04 | 0.039 |
| | | | 38,119,198 | *GSDMA* | | 5.612 | 1.16E-07 | 7.23E-04 | 0.009 |
| | | | 38,119,207 | *GSDMA* | | 7.125 | 6.38E-11 | 7.23E-04 | 0.009 |
| | | | 38,119,649 | | *GSDMA* | 4.403 | 2.20E-05 | 7.23E-04 | 0.002 |
| | | | 38,119,760 | | *GSDMA* | 4.196 | 5.01E-05 | 7.23E-04 | 0.011 |
| | rs1011082 | g.38068514.A>G | 37,887,061 | *MIEN1* | | 4.098 | 7.30E-05 | 7.66E-04 | 0.002 |
| | | | 38,018,804 | | *IKZF3* | 3.467 | 7.14E-04 | 7.66E-04 | 0.046 |
| | | | 38,024,063 | | | -3.393 | 9.18E-04 | 7.66E-04 | 0.004 |
| | | | 38,077,191 | | | 3.710 | 3.06E-04 | 7.66E-04 | 0.034 |
| | | | 38,077,464 | | *ORMDL3* | 3.569 | 5.03E-04 | 7.66E-04 | 0.014 |
| | | | 38,080,829 | | *ORMDL3* | 5.568 | 1.42E-07 | 7.66E-04 | 0.014 |
| | | | 38,081,492 | | *ORMDL3* | 3.647 | 3.82E-04 | 7.66E-04 | 2.01E-04 |
| | | | 38,081,829 | | *ORMDL3* | 3.381 | 9.54E-04 | 7.66E-04 | 2.01E-04 |

| | | | | | | | | |
|---|---|---|---|---|---|---|---|---|
| | | 38,081,995 | | *ORMDL3* | 4.302 | 3.30E-05 | 7.66E-04 | 2.01E-04 |
| | | 38,082,022 | | *ORMDL3* | 5.086 | 1.25E-06 | 7.66E-04 | 6.40E-04 |
| | | 38,082,035 | | *ORMDL3* | 5.500 | 1.94E-07 | 7.66E-04 | 6.40E-04 |
| | | 38,082,073 | | *ORMDL3* | 4.964 | 2.13E-06 | 7.66E-04 | 6.40E-04 |
| | | 38,082,235 | | *ORMDL3* | 4.057 | 8.52E-05 | 7.66E-04 | 0.004 |
| | | 38,082,346 | | *ORMDL3* | 4.502 | 1.48E-05 | 7.66E-04 | 0.011 |
| | | 38,082,448 | | *ORMDL3* | 3.913 | 1.46E-04 | 7.66E-04 | 0.011 |
| | | 38,082,658 | | *ORMDL3* | 5.465 | 2.28E-07 | 7.66E-04 | 0.011 |
| | | 38,108,464 | | | -3.506 | 6.26E-04 | 7.66E-04 | 0.007 |
| | | 38,108,710 | | | -4.708 | 6.33E-06 | 7.66E-04 | 0.039 |
| | | 38,119,198 | *GSDMA* | | 4.693 | 6.73E-06 | 7.66E-04 | 0.009 |
| | | 38,119,207 | *GSDMA* | | 6.474 | 1.79E-09 | 7.66E-04 | 0.009 |
| | | 38,119,649 | | *GSDMA* | 5.227 | 6.68E-07 | 7.66E-04 | 0.002 |
| | | 38,119,760 | | *GSDMA* | 5.152 | 9.34E-07 | 7.66E-04 | 0.011 |
| rs921650 | g.38069076.C>T | 37,887,061 | *MIEN1* | | 3.704 | 3.12E-04 | 5.34E-04 | 0.002 |
| | | 38,018,804 | | *IKZF3* | 3.417 | 8.44E-04 | 5.34E-04 | 0.046 |
| | | 38,077,191 | | | 3.827 | 2.00E-04 | 5.34E-04 | 0.034 |
| | | 38,077,464 | | *ORMDL3* | 3.639 | 3.93E-04 | 5.34E-04 | 0.014 |
| | | 38,080,829 | | *ORMDL3* | 5.536 | 1.64E-07 | 5.34E-04 | 0.014 |
| | | 38,081,492 | | *ORMDL3* | 3.585 | 4.75E-04 | 5.34E-04 | 2.01E-04 |
| | | 38,081,995 | | *ORMDL3* | 4.416 | 2.10E-05 | 5.34E-04 | 2.01E-04 |
| | | 38,082,022 | | *ORMDL3* | 5.112 | 1.11E-06 | 5.34E-04 | 6.40E-04 |
| | | 38,082,035 | | *ORMDL3* | 5.307 | 4.66E-07 | 5.34E-04 | 6.40E-04 |
| | | 38,082,073 | | *ORMDL3* | 5.048 | 1.47E-06 | 5.34E-04 | 6.40E-04 |
| | | 38,082,235 | | *ORMDL3* | 4.077 | 7.89E-05 | 5.34E-04 | 0.004 |
| | | 38,082,346 | | *ORMDL3* | 4.493 | 1.53E-05 | 5.34E-04 | 0.011 |
| | | 38,082,448 | | *ORMDL3* | 3.848 | 1.86E-04 | 5.34E-04 | 0.011 |
| | | 38,082,658 | | *ORMDL3* | 5.521 | 1.76E-07 | 5.34E-04 | 0.011 |
| | | 38,108,464 | | | -3.396 | 9.08E-04 | 5.34E-04 | 0.007 |
| | | 38,108,710 | | | -4.944 | 2.31E-06 | 5.34E-04 | 0.039 |
| | | 38,119,198 | *GSDMA* | | 4.736 | 5.63E-06 | 5.34E-04 | 0.009 |
| | | 38,119,207 | *GSDMA* | | 6.586 | 1.02E-09 | 5.34E-04 | 0.009 |
| | | 38,119,649 | | *GSDMA* | 5.298 | 4.85E-07 | 5.34E-04 | 0.002 |
| | | 38,119,760 | | *GSDMA* | 5.134 | 1.01E-06 | 5.34E-04 | 0.011 |
| rs921649 | g.38069274.G>A | 37,887,061 | *MIEN1* | | 3.704 | 3.12E-04 | 5.34E-04 | 0.002 |
| | | 38,018,804 | | *IKZF3* | 3.417 | 8.44E-04 | 5.34E-04 | 0.046 |
| | | 38,077,191 | | | 3.827 | 2.00E-04 | 5.34E-04 | 0.034 |
| | | 38,077,464 | | *ORMDL3* | 3.639 | 3.93E-04 | 5.34E-04 | 0.014 |
| | | 38,080,829 | | *ORMDL3* | 5.536 | 1.64E-07 | 5.34E-04 | 0.014 |
| | | 38,081,492 | | *ORMDL3* | 3.585 | 4.75E-04 | 5.34E-04 | 2.01E-04 |
| | | 38,081,995 | | *ORMDL3* | 4.416 | 2.10E-05 | 5.34E-04 | 2.01E-04 |

| | | | | | | | | |
|---|---|---|---|---|---|---|---|---|
| | | 38,082,022 | | *ORMDL3* | 5.112 | 1.11E-06 | 5.34E-04 | 6.40E-04 |
| | | 38,082,035 | | *ORMDL3* | 5.307 | 4.66E-07 | 5.34E-04 | 6.40E-04 |
| | | 38,082,073 | | *ORMDL3* | 5.048 | 1.47E-06 | 5.34E-04 | 6.40E-04 |
| | | 38,082,235 | | *ORMDL3* | 4.077 | 7.89E-05 | 5.34E-04 | 0.004 |
| | | 38,082,346 | | *ORMDL3* | 4.493 | 1.53E-05 | 5.34E-04 | 0.011 |
| | | 38,082,448 | | *ORMDL3* | 3.848 | 1.86E-04 | 5.34E-04 | 0.011 |
| | | 38,082,658 | | *ORMDL3* | 5.521 | 1.76E-07 | 5.34E-04 | 0.011 |
| | | 38,108,464 | | | -3.396 | 9.08E-04 | 5.34E-04 | 0.007 |
| | | 38,108,710 | | | -4.944 | 2.31E-06 | 5.34E-04 | 0.039 |
| | | 38,119,198 | *GSDMA* | | 4.736 | 5.63E-06 | 5.34E-04 | 0.009 |
| | | 38,119,207 | *GSDMA* | | 6.586 | 1.02E-09 | 5.34E-04 | 0.009 |
| | | 38,119,649 | | *GSDMA* | 5.298 | 4.85E-07 | 5.34E-04 | 0.002 |
| | | 38,119,760 | | *GSDMA* | 5.134 | 1.01E-06 | 5.34E-04 | 0.011 |
| rs5820308 | g.38069364_38069369del | 37,887,061 | *MIEN1* | | 4.496 | 1.52E-05 | 8.13E-04 | 0.002 |
| | | 38,077,191 | | | 4.060 | 8.44E-05 | 8.13E-04 | 0.034 |
| | | 38,077,464 | | *ORMDL3* | 4.269 | 3.76E-05 | 8.13E-04 | 0.014 |
| | | 38,080,829 | | *ORMDL3* | 3.670 | 3.53E-04 | 8.13E-04 | 0.014 |
| | | 38,081,812 | | *ORMDL3* | 3.450 | 7.57E-04 | 8.13E-04 | 2.01E-04 |
| | | 38,081,829 | | *ORMDL3* | 4.212 | 4.70E-05 | 8.13E-04 | 2.01E-04 |
| | | 38,081,995 | | *ORMDL3* | 5.759 | 5.79E-08 | 8.13E-04 | 2.01E-04 |
| | | 38,082,022 | | *ORMDL3* | 6.804 | 3.36E-10 | 8.13E-04 | 6.40E-04 |
| | | 38,082,035 | | *ORMDL3* | 5.893 | 3.07E-08 | 8.13E-04 | 6.40E-04 |
| | | 38,082,073 | | *ORMDL3* | 5.907 | 2.87E-08 | 8.13E-04 | 6.40E-04 |
| | | 38,082,206 | | *ORMDL3* | 3.944 | 1.30E-04 | 8.13E-04 | 0.004 |
| | | 38,082,235 | | *ORMDL3* | 4.422 | 2.05E-05 | 8.13E-04 | 0.004 |
| | | 38,082,346 | | *ORMDL3* | 4.459 | 1.76E-05 | 8.13E-04 | 0.011 |
| | | 38,082,448 | | *ORMDL3* | 3.372 | 9.83E-04 | 8.13E-04 | 0.011 |
| | | 38,082,658 | | *ORMDL3* | 5.515 | 1.81E-07 | 8.13E-04 | 0.011 |
| | | 38,108,710 | | | -5.683 | 8.27E-08 | 8.13E-04 | 0.039 |
| | | 38,119,198 | *GSDMA* | | 4.828 | 3.81E-06 | 8.13E-04 | 0.009 |
| | | 38,119,207 | *GSDMA* | | 6.261 | 5.14E-09 | 8.13E-04 | 0.009 |
| | | 38,119,649 | | *GSDMA* | 5.655 | 9.45E-08 | 8.13E-04 | 0.002 |
| | | 38,119,760 | | *GSDMA* | 4.753 | 5.23E-06 | 8.13E-04 | 0.011 |
| | | 38,120,234 | | *GSDMA* | 3.403 | 8.87E-04 | 8.13E-04 | 0.011 |
| rs6503524 | g.38069809.C>T | 37,887,061 | *MIEN1* | | 3.704 | 3.12E-04 | 5.34E-04 | 0.002 |
| | | 38,018,804 | | *IKZF3* | 3.417 | 8.44E-04 | 5.34E-04 | 0.046 |
| | | 38,077,191 | | | 3.827 | 2.00E-04 | 5.34E-04 | 0.034 |
| | | 38,077,464 | | *ORMDL3* | 3.639 | 3.93E-04 | 5.34E-04 | 0.014 |
| | | 38,080,829 | | *ORMDL3* | 5.536 | 1.64E-07 | 5.34E-04 | 0.014 |
| | | 38,081,492 | | *ORMDL3* | 3.585 | 4.75E-04 | 5.34E-04 | 2.01E-04 |

| | | | | | | | | |
|---|---|---|---|---|---|---|---|---|
| | | 38,081,995 | | *ORMDL3* | 4.416 | 2.10E-05 | 5.34E-04 | 2.01E-04 |
| | | 38,082,022 | | *ORMDL3* | 5.112 | 1.11E-06 | 5.34E-04 | 6.40E-04 |
| | | 38,082,035 | | *ORMDL3* | 5.307 | 4.66E-07 | 5.34E-04 | 6.40E-04 |
| | | 38,082,073 | | *ORMDL3* | 5.048 | 1.47E-06 | 5.34E-04 | 6.40E-04 |
| | | 38,082,235 | | *ORMDL3* | 4.077 | 7.89E-05 | 5.34E-04 | 0.004 |
| | | 38,082,346 | | *ORMDL3* | 4.493 | 1.53E-05 | 5.34E-04 | 0.011 |
| | | 38,082,448 | | *ORMDL3* | 3.848 | 1.86E-04 | 5.34E-04 | 0.011 |
| | | 38,082,658 | | *ORMDL3* | 5.521 | 1.76E-07 | 5.34E-04 | 0.011 |
| | | 38,108,464 | | | -3.396 | 9.08E-04 | 5.34E-04 | 0.007 |
| | | 38,108,710 | | | -4.944 | 2.31E-06 | 5.34E-04 | 0.039 |
| | | 38,119,198 | *GSDMA* | | 4.736 | 5.63E-06 | 5.34E-04 | 0.009 |
| | | 38,119,207 | *GSDMA* | | 6.586 | 1.02E-09 | 5.34E-04 | 0.009 |
| | | 38,119,649 | | *GSDMA* | 5.298 | 4.85E-07 | 5.34E-04 | 0.002 |
| | | 38,119,760 | | *GSDMA* | 5.134 | 1.01E-06 | 5.34E-04 | 0.011 |
| rs7216389 | g.38069949.C>T | 37,887,061 | *MIEN1* | | 3.704 | 3.12E-04 | 5.34E-04 | 0.002 |
| | | 38,018,804 | | *IKZF3* | 3.417 | 8.44E-04 | 5.34E-04 | 0.046 |
| | | 38,077,191 | | | 3.827 | 2.00E-04 | 5.34E-04 | 0.034 |
| | | 38,077,464 | | *ORMDL3* | 3.639 | 3.93E-04 | 5.34E-04 | 0.014 |
| | | 38,080,829 | | *ORMDL3* | 5.536 | 1.64E-07 | 5.34E-04 | 0.014 |
| | | 38,081,492 | | *ORMDL3* | 3.585 | 4.75E-04 | 5.34E-04 | 2.01E-04 |
| | | 38,081,995 | | *ORMDL3* | 4.416 | 2.10E-05 | 5.34E-04 | 2.01E-04 |
| | | 38,082,022 | | *ORMDL3* | 5.112 | 1.11E-06 | 5.34E-04 | 6.40E-04 |
| | | 38,082,035 | | *ORMDL3* | 5.307 | 4.66E-07 | 5.34E-04 | 6.40E-04 |
| | | 38,082,073 | | *ORMDL3* | 5.048 | 1.47E-06 | 5.34E-04 | 6.40E-04 |
| | | 38,082,235 | | *ORMDL3* | 4.077 | 7.89E-05 | 5.34E-04 | 0.004 |
| | | 38,082,346 | | *ORMDL3* | 4.493 | 1.53E-05 | 5.34E-04 | 0.011 |
| | | 38,082,448 | | *ORMDL3* | 3.848 | 1.86E-04 | 5.34E-04 | 0.011 |
| | | 38,082,658 | | *ORMDL3* | 5.521 | 1.76E-07 | 5.34E-04 | 0.011 |
| | | 38,108,464 | | | -3.396 | 9.08E-04 | 5.34E-04 | 0.007 |
| | | 38,108,710 | | | -4.944 | 2.31E-06 | 5.34E-04 | 0.039 |
| | | 38,119,198 | *GSDMA* | | 4.736 | 5.63E-06 | 5.34E-04 | 0.009 |
| | | 38,119,207 | *GSDMA* | | 6.586 | 1.02E-09 | 5.34E-04 | 0.009 |
| | | 38,119,649 | | *GSDMA* | 5.298 | 4.85E-07 | 5.34E-04 | 0.002 |
| | | 38,119,760 | | *GSDMA* | 5.134 | 1.01E-06 | 5.34E-04 | 0.011 |
| rs7216558 | g.38070071.C>T | 37,887,061 | *MIEN1* | | 3.704 | 3.12E-04 | 5.34E-04 | 0.002 |
| | | 38,018,804 | | *IKZF3* | 3.417 | 8.44E-04 | 5.34E-04 | 0.046 |
| | | 38,077,191 | | | 3.827 | 2.00E-04 | 5.34E-04 | 0.034 |
| | | 38,077,464 | | *ORMDL3* | 3.639 | 3.93E-04 | 5.34E-04 | 0.014 |
| | | 38,080,829 | | *ORMDL3* | 5.536 | 1.64E-07 | 5.34E-04 | 0.014 |
| | | 38,081,492 | | *ORMDL3* | 3.585 | 4.75E-04 | 5.34E-04 | 2.01E-04 |
| | | 38,081,995 | | *ORMDL3* | 4.416 | 2.10E-05 | 5.34E-04 | 2.01E-04 |

| | | | | | | | | |
|---|---|---|---|---|---|---|---|---|
| | | 38,082,022 | | *ORMDL3* | 5.112 | 1.11E-06 | 5.34E-04 | 6.40E-04 |
| | | 38,082,035 | | *ORMDL3* | 5.307 | 4.66E-07 | 5.34E-04 | 6.40E-04 |
| | | 38,082,073 | | *ORMDL3* | 5.048 | 1.47E-06 | 5.34E-04 | 6.40E-04 |
| | | 38,082,235 | | *ORMDL3* | 4.077 | 7.89E-05 | 5.34E-04 | 0.004 |
| | | 38,082,346 | | *ORMDL3* | 4.493 | 1.53E-05 | 5.34E-04 | 0.011 |
| | | 38,082,448 | | *ORMDL3* | 3.848 | 1.86E-04 | 5.34E-04 | 0.011 |
| | | 38,082,658 | | *ORMDL3* | 5.521 | 1.76E-07 | 5.34E-04 | 0.011 |
| | | 38,108,464 | | | -3.396 | 9.08E-04 | 5.34E-04 | 0.007 |
| | | 38,108,710 | | | -4.944 | 2.31E-06 | 5.34E-04 | 0.039 |
| | | 38,119,198 | *GSDMA* | | 4.736 | 5.63E-06 | 5.34E-04 | 0.009 |
| | | 38,119,207 | *GSDMA* | | 6.586 | 1.02E-09 | 5.34E-04 | 0.009 |
| | | 38,119,649 | | *GSDMA* | 5.298 | 4.85E-07 | 5.34E-04 | 0.002 |
| | | 38,119,760 | | *GSDMA* | 5.134 | 1.01E-06 | 5.34E-04 | 0.011 |
| rs7221605 | g.38070789.C>T | 37,887,061 | *MIEN1* | | 3.963 | 1.21E-04 | | 0.002 |
| | | 37,894,590 | | | 3.408 | 8.73E-04 | | 0.013 |
| | | 37,896,827 | | | 3.416 | 8.48E-04 | | 0.008 |
| | | 38,077,191 | | | 3.920 | 1.43E-04 | | 0.034 |
| | | 38,077,464 | | *ORMDL3* | 3.848 | 1.86E-04 | | 0.014 |
| | | 38,080,829 | | *ORMDL3* | 5.980 | 2.02E-08 | | 0.014 |
| | | 38,081,812 | | *ORMDL3* | 3.721 | 2.95E-04 | | 2.01E-04 |
| | | 38,081,829 | | *ORMDL3* | 3.444 | 7.73E-04 | | 2.01E-04 |
| | | 38,081,995 | | *ORMDL3* | 4.405 | 2.19E-05 | | 2.01E-04 |
| | | 38,082,022 | | *ORMDL3* | 4.799 | 4.31E-06 | | 6.40E-04 |
| | | 38,082,035 | | *ORMDL3* | 5.305 | 4.71E-07 | | 6.40E-04 |
| | | 38,082,073 | | *ORMDL3* | 5.047 | 1.48E-06 | | 6.40E-04 |
| | | 38,082,235 | | *ORMDL3* | 4.024 | 9.67E-05 | | 0.004 |
| | | 38,082,346 | | *ORMDL3* | 4.398 | 2.25E-05 | | 0.011 |
| | | 38,082,448 | | *ORMDL3* | 3.951 | 1.27E-04 | | 0.011 |
| | | 38,082,658 | | *ORMDL3* | 5.500 | 1.94E-07 | | 0.011 |
| | | 38,108,710 | | | -5.118 | 1.09E-06 | | 0.039 |
| | | 38,119,198 | *GSDMA* | | 4.691 | 6.77E-06 | | 0.009 |
| | | 38,119,207 | *GSDMA* | | 6.587 | 1.02E-09 | | 0.009 |
| | | 38,119,649 | | *GSDMA* | 5.445 | 2.49E-07 | | 0.002 |
| | | 38,119,760 | | *GSDMA* | 5.302 | 4.78E-07 | | 0.011 |
| rs150597688 | g.38071086_38071086ins TT | 37,887,061 | *MIEN1* | | 3.963 | 1.21E-04 | | 0.002 |
| | | 37,894,590 | | | 3.408 | 8.73E-04 | | 0.013 |
| | | 37,896,827 | | | 3.416 | 8.48E-04 | | 0.008 |
| | | 38,077,191 | | | 3.920 | 1.43E-04 | | 0.034 |
| | | 38,077,464 | | *ORMDL3* | 3.848 | 1.86E-04 | | 0.014 |
| | | 38,080,829 | | *ORMDL3* | 5.980 | 2.02E-08 | | 0.014 |

| | | | | | | | |
|---|---|---|---|---|---|---|---|
| | | 38,081,812 | | *ORMDL3* | 3.721 | 2.95E-04 | 2.01E-04 |
| | | 38,081,829 | | *ORMDL3* | 3.444 | 7.73E-04 | 2.01E-04 |
| | | 38,081,995 | | *ORMDL3* | 4.405 | 2.19E-05 | 2.01E-04 |
| | | 38,082,022 | | *ORMDL3* | 4.799 | 4.31E-06 | 6.40E-04 |
| | | 38,082,035 | | *ORMDL3* | 5.305 | 4.71E-07 | 6.40E-04 |
| | | 38,082,073 | | *ORMDL3* | 5.047 | 1.48E-06 | 6.40E-04 |
| | | 38,082,235 | | *ORMDL3* | 4.024 | 9.67E-05 | 0.004 |
| | | 38,082,346 | | *ORMDL3* | 4.398 | 2.25E-05 | 0.011 |
| | | 38,082,448 | | *ORMDL3* | 3.951 | 1.27E-04 | 0.011 |
| | | 38,082,658 | | *ORMDL3* | 5.500 | 1.94E-07 | 0.011 |
| | | 38,108,710 | | | -5.118 | 1.09E-06 | 0.039 |
| | | 38,119,198 | *GSDMA* | | 4.691 | 6.77E-06 | 0.009 |
| | | 38,119,207 | *GSDMA* | | 6.587 | 1.02E-09 | 0.009 |
| | | 38,119,649 | | *GSDMA* | 5.445 | 2.49E-07 | 0.002 |
| | | 38,119,760 | | *GSDMA* | 5.302 | 4.78E-07 | 0.011 |
| rs143385463 | g.38071855_38071858del | 37,887,061 | *MIEN1* | | 3.963 | 1.21E-04 | 0.002 |
| | | 37,894,590 | | | 3.408 | 8.73E-04 | 0.013 |
| | | 37,896,827 | | | 3.416 | 8.48E-04 | 0.008 |
| | | 38,077,191 | | | 3.920 | 1.43E-04 | 0.034 |
| | | 38,077,464 | | *ORMDL3* | 3.848 | 1.86E-04 | 0.014 |
| | | 38,080,829 | | *ORMDL3* | 5.980 | 2.02E-08 | 0.014 |
| | | 38,081,812 | | *ORMDL3* | 3.721 | 2.95E-04 | 2.01E-04 |
| | | 38,081,829 | | *ORMDL3* | 3.444 | 7.73E-04 | 2.01E-04 |
| | | 38,081,995 | | *ORMDL3* | 4.405 | 2.19E-05 | 2.01E-04 |
| | | 38,082,022 | | *ORMDL3* | 4.799 | 4.31E-06 | 6.40E-04 |
| | | 38,082,035 | | *ORMDL3* | 5.305 | 4.71E-07 | 6.40E-04 |
| | | 38,082,073 | | *ORMDL3* | 5.047 | 1.48E-06 | 6.40E-04 |
| | | 38,082,235 | | *ORMDL3* | 4.024 | 9.67E-05 | 0.004 |
| | | 38,082,346 | | *ORMDL3* | 4.398 | 2.25E-05 | 0.011 |
| | | 38,082,448 | | *ORMDL3* | 3.951 | 1.27E-04 | 0.011 |
| | | 38,082,658 | | *ORMDL3* | 5.500 | 1.94E-07 | 0.011 |
| | | 38,108,710 | | | -5.118 | 1.09E-06 | 0.039 |
| | | 38,119,198 | *GSDMA* | | 4.691 | 6.77E-06 | 0.009 |
| | | 38,119,207 | *GSDMA* | | 6.587 | 1.02E-09 | 0.009 |
| | | 38,119,649 | | *GSDMA* | 5.445 | 2.49E-07 | 0.002 |
| | | 38,119,760 | | *GSDMA* | 5.302 | 4.78E-07 | 0.011 |
| rs1031458 | g.38072173.G>T | 37,887,061 | *MIEN1* | | 3.963 | 1.21E-04 | 0.002 |
| | | 37,894,590 | | | 3.408 | 8.73E-04 | 0.013 |
| | | 37,896,827 | | | 3.416 | 8.48E-04 | 0.008 |
| | | 38,077,191 | | | 3.920 | 1.43E-04 | 0.034 |

| | | | | | | | |
|---|---|---|---|---|---|---|---|
| | | 38,077,464 | | *ORMDL3* | 3.848 | 1.86E-04 | 0.014 |
| | | 38,080,829 | | *ORMDL3* | 5.980 | 2.02E-08 | 0.014 |
| | | 38,081,812 | | *ORMDL3* | 3.721 | 2.95E-04 | 2.01E-04 |
| | | 38,081,829 | | *ORMDL3* | 3.444 | 7.73E-04 | 2.01E-04 |
| | | 38,081,995 | | *ORMDL3* | 4.405 | 2.19E-05 | 2.01E-04 |
| | | 38,082,022 | | *ORMDL3* | 4.799 | 4.31E-06 | 6.40E-04 |
| | | 38,082,035 | | *ORMDL3* | 5.305 | 4.71E-07 | 6.40E-04 |
| | | 38,082,073 | | *ORMDL3* | 5.047 | 1.48E-06 | 6.40E-04 |
| | | 38,082,235 | | *ORMDL3* | 4.024 | 9.67E-05 | 0.004 |
| | | 38,082,346 | | *ORMDL3* | 4.398 | 2.25E-05 | 0.011 |
| | | 38,082,448 | | *ORMDL3* | 3.951 | 1.27E-04 | 0.011 |
| | | 38,082,658 | | *ORMDL3* | 5.500 | 1.94E-07 | 0.011 |
| | | 38,108,710 | | | -5.118 | 1.09E-06 | 0.039 |
| | | 38,119,198 | *GSDMA* | | 4.691 | 6.77E-06 | 0.009 |
| | | 38,119,207 | *GSDMA* | | 6.587 | 1.02E-09 | 0.009 |
| | | 38,119,649 | | *GSDMA* | 5.445 | 2.49E-07 | 0.002 |
| | | 38,119,760 | | *GSDMA* | 5.302 | 4.78E-07 | 0.011 |
| rs1031459 | g.38072245.C>G | 37,887,061 | *MIEN1* | | 3.937 | 1.34E-04 | 0.002 |
| | | 38,024,063 | | | -3.442 | 7.78E-04 | 0.004 |
| | | 38,024,331 | | | -3.505 | 6.28E-04 | 0.017 |
| | | 38,077,464 | | *ORMDL3* | 5.531 | 1.68E-07 | 0.014 |
| | | 38,080,829 | | *ORMDL3* | 3.548 | 5.41E-04 | 0.014 |
| | | 38,108,710 | | | -5.361 | 3.66E-07 | 0.039 |
| | | 38,119,071 | *GSDMA* | | 3.410 | 8.67E-04 | 0.009 |
| | | 38,119,132 | *GSDMA* | | 3.685 | 3.35E-04 | 0.009 |
| | | 38,119,198 | *GSDMA* | | 5.422 | 2.78E-07 | 0.009 |
| | | 38,119,207 | *GSDMA* | | 6.375 | 2.93E-09 | 0.009 |
| | | 38,119,649 | | *GSDMA* | 4.926 | 2.50E-06 | 0.002 |
| | | 38,119,760 | | *GSDMA* | 4.492 | 1.54E-05 | 0.011 |
| rs1031460 | g.38072247.T>G | 37,887,061 | *MIEN1* | | 3.973 | 1.17E-04 | 0.002 |
| | | 37,894,590 | | | 3.381 | 9.54E-04 | 0.013 |
| | | 37,896,827 | | | 3.421 | 8.35E-04 | 0.008 |
| | | 38,077,191 | | | 3.890 | 1.59E-04 | 0.034 |
| | | 38,077,464 | | *ORMDL3* | 3.865 | 1.74E-04 | 0.014 |
| | | 38,080,829 | | *ORMDL3* | 5.944 | 2.41E-08 | 0.014 |
| | | 38,081,812 | | *ORMDL3* | 3.744 | 2.71E-04 | 2.01E-04 |
| | | 38,081,829 | | *ORMDL3* | 3.454 | 7.46E-04 | 2.01E-04 |
| | | 38,081,995 | | *ORMDL3* | 4.366 | 2.55E-05 | 2.01E-04 |
| | | 38,082,022 | | *ORMDL3* | 4.767 | 4.94E-06 | 6.40E-04 |
| | | 38,082,035 | | *ORMDL3* | 5.322 | 4.36E-07 | 6.40E-04 |
| | | 38,082,073 | | *ORMDL3* | 5.052 | 1.45E-06 | 6.40E-04 |

| | | | | | | | |
|---|---|---|---|---|---|---|---|
| | | 38,082,235 | | *ORMDL3* | 4.149 | 5.99E-05 | 0.004 |
| | | 38,082,346 | | *ORMDL3* | 4.375 | 2.47E-05 | 0.011 |
| | | 38,082,448 | | *ORMDL3* | 4.006 | 1.03E-04 | 0.011 |
| | | 38,082,658 | | *ORMDL3* | 5.526 | 1.72E-07 | 0.011 |
| | | 38,108,710 | | | -5.085 | 1.25E-06 | 0.039 |
| | | 38,119,198 | *GSDMA* | | 4.689 | 6.85E-06 | 0.009 |
| | | 38,119,207 | *GSDMA* | | 6.566 | 1.13E-09 | 0.009 |
| | | 38,119,649 | | *GSDMA* | 5.412 | 2.89E-07 | 0.002 |
| | | 38,119,760 | | *GSDMA* | 5.325 | 4.31E-07 | 0.011 |
| rs8065777 | g.38072402.C>T | 37,887,061 | *MIEN1* | | 3.963 | 1.21E-04 | 0.002 |
| | | 37,894,590 | | | 3.408 | 8.73E-04 | 0.013 |
| | | 37,896,827 | | | 3.416 | 8.48E-04 | 0.008 |
| | | 38,077,191 | | | 3.920 | 1.43E-04 | 0.034 |
| | | 38,077,464 | | *ORMDL3* | 3.848 | 1.86E-04 | 0.014 |
| | | 38,080,829 | | *ORMDL3* | 5.980 | 2.02E-08 | 0.014 |
| | | 38,081,812 | | *ORMDL3* | 3.721 | 2.95E-04 | 2.01E-04 |
| | | 38,081,829 | | *ORMDL3* | 3.444 | 7.73E-04 | 2.01E-04 |
| | | 38,081,995 | | *ORMDL3* | 4.405 | 2.19E-05 | 2.01E-04 |
| | | 38,082,022 | | *ORMDL3* | 4.799 | 4.31E-06 | 6.40E-04 |
| | | 38,082,035 | | *ORMDL3* | 5.305 | 4.71E-07 | 6.40E-04 |
| | | 38,082,073 | | *ORMDL3* | 5.047 | 1.48E-06 | 6.40E-04 |
| | | 38,082,235 | | *ORMDL3* | 4.024 | 9.67E-05 | 0.004 |
| | | 38,082,346 | | *ORMDL3* | 4.398 | 2.25E-05 | 0.011 |
| | | 38,082,448 | | *ORMDL3* | 3.951 | 1.27E-04 | 0.011 |
| | | 38,082,658 | | *ORMDL3* | 5.500 | 1.94E-07 | 0.011 |
| | | 38,108,710 | | | -5.118 | 1.09E-06 | 0.039 |
| | | 38,119,198 | *GSDMA* | | 4.691 | 6.77E-06 | 0.009 |
| | | 38,119,207 | *GSDMA* | | 6.587 | 1.02E-09 | 0.009 |
| | | 38,119,649 | | *GSDMA* | 5.445 | 2.49E-07 | 0.002 |
| | | 38,119,760 | | *GSDMA* | 5.302 | 4.78E-07 | 0.011 |
| rs112599791 | g.38072475.A>G | 37,856,732 | | *ERBB2* | -3.668 | 3.56E-04 | 0.009 |
| | | 37,886,749 | | *MIEN1* | 3.945 | 1.30E-04 | 0.002 |
| | | 37,896,063 | | | 3.476 | 6.92E-04 | 0.009 |
| | | 38,020,485 | *IKZF3* | | 3.962 | 1.22E-04 | 0.005 |
| | | 38,080,829 | | *ORMDL3* | 3.995 | 1.08E-04 | 0.014 |
| | | 38,096,333 | | | -3.664 | 3.60E-04 | 0.033 |
| | | 38,096,571 | | | -4.780 | 4.67E-06 | 0.033 |
| | | 38,096,578 | | | -5.642 | 1.00E-07 | 0.033 |
| | | 38,096,648 | | | -3.887 | 1.61E-04 | 0.033 |
| rs2872516 | g.38072727.C>T | 37,828,296 | | *PGAP3* | 3.534 | 5.67E-04 | 0.002 |
| | | 37,887,061 | *MIEN1* | | 4.828 | 3.81E-06 | 0.002 |

| | | | | | | | | |
|---|---|---|---|---|---|---|---|---|
| | | 37,894,590 | | | 3.577 | 4.89E-04 | | 0.013 |
| | | 37,898,302 | | | -3.502 | 6.33E-04 | | 0.008 |
| | | 38,077,191 | | | 4.169 | 5.54E-05 | | 0.034 |
| | | 38,077,277 | | | 3.429 | 8.13E-04 | | 0.014 |
| | | 38,077,464 | | *ORMDL3* | 4.537 | 1.28E-05 | | 0.014 |
| | | 38,080,829 | | *ORMDL3* | 4.058 | 8.50E-05 | | 0.014 |
| | | 38,081,812 | | *ORMDL3* | 3.862 | 1.76E-04 | | 2.01E-04 |
| | | 38,081,829 | | *ORMDL3* | 4.343 | 2.80E-05 | | 2.01E-04 |
| | | 38,081,995 | | *ORMDL3* | 5.792 | 4.96E-08 | | 2.01E-04 |
| | | 38,082,022 | | *ORMDL3* | 6.441 | 2.10E-09 | | 6.40E-04 |
| | | 38,082,035 | | *ORMDL3* | 5.916 | 2.76E-08 | | 6.40E-04 |
| | | 38,082,073 | | *ORMDL3* | 5.945 | 2.40E-08 | | 6.40E-04 |
| | | 38,082,206 | | *ORMDL3* | 3.921 | 1.42E-04 | | 0.004 |
| | | 38,082,235 | | *ORMDL3* | 4.393 | 2.30E-05 | | 0.004 |
| | | 38,082,346 | | *ORMDL3* | 4.373 | 2.49E-05 | | 0.011 |
| | | 38,082,448 | | *ORMDL3* | 3.484 | 6.73E-04 | | 0.011 |
| | | 38,082,658 | | *ORMDL3* | 5.511 | 1.85E-07 | | 0.011 |
| | | 38,108,710 | | | -5.927 | 2.60E-08 | | 0.039 |
| | | 38,119,198 | *GSDMA* | | 4.803 | 4.23E-06 | | 0.009 |
| | | 38,119,207 | *GSDMA* | | 6.298 | 4.30E-09 | | 0.009 |
| | | 38,119,649 | | *GSDMA* | 5.854 | 3.70E-08 | | 0.002 |
| | | 38,119,760 | | *GSDMA* | 4.940 | 2.35E-06 | | 0.011 |
| | | 38,120,234 | | *GSDMA* | 3.435 | 7.96E-04 | | 0.011 |
| rs113894104 | g.38072955.T>C | 37,856,732 | | *ERBB2* | -3.668 | 3.56E-04 | | 0.009 |
| | | 37,886,749 | | *MIEN1* | 3.945 | 1.30E-04 | | 0.002 |
| | | 37,896,063 | | | 3.476 | 6.92E-04 | | 0.009 |
| | | 38,020,485 | *IKZF3* | | 3.962 | 1.22E-04 | | 0.005 |
| | | 38,080,829 | | *ORMDL3* | 3.995 | 1.08E-04 | | 0.014 |
| | | 38,096,333 | | | -3.664 | 3.60E-04 | | 0.033 |
| | | 38,096,571 | | | -4.780 | 4.67E-06 | | 0.033 |
| | | 38,096,578 | | | -5.642 | 1.00E-07 | | 0.033 |
| | | 38,096,648 | | | -3.887 | 1.61E-04 | | 0.033 |
| rs9303279 | g.38073968.G>C | 37,887,061 | *MIEN1* | | 4.496 | 1.52E-05 | 6.13E-04 | 0.002 |
| | | 38,077,191 | | | 4.060 | 8.44E-05 | 6.13E-04 | 0.034 |
| | | 38,077,464 | | *ORMDL3* | 4.269 | 3.76E-05 | 6.13E-04 | 0.014 |
| | | 38,080,829 | | *ORMDL3* | 3.670 | 3.53E-04 | 6.13E-04 | 0.014 |
| | | 38,081,812 | | *ORMDL3* | 3.450 | 7.57E-04 | 6.13E-04 | 2.01E-04 |
| | | 38,081,829 | | *ORMDL3* | 4.212 | 4.70E-05 | 6.13E-04 | 2.01E-04 |
| | | 38,081,995 | | *ORMDL3* | 5.759 | 5.79E-08 | 6.13E-04 | 2.01E-04 |
| | | 38,082,022 | | *ORMDL3* | 6.804 | 3.36E-10 | 6.13E-04 | 6.40E-04 |
| | | 38,082,035 | | *ORMDL3* | 5.893 | 3.07E-08 | 6.13E-04 | 6.40E-04 |

| | | | | | | | | |
|---|---|---|---|---|---|---|---|---|
| | | 38,082,073 | | *ORMDL3* | 5.907 | 2.87E-08 | 6.13E-04 | 6.40E-04 |
| | | 38,082,206 | | *ORMDL3* | 3.944 | 1.30E-04 | 6.13E-04 | 0.004 |
| | | 38,082,235 | | *ORMDL3* | 4.422 | 2.05E-05 | 6.13E-04 | 0.004 |
| | | 38,082,346 | | *ORMDL3* | 4.459 | 1.76E-05 | 6.13E-04 | 0.011 |
| | | 38,082,448 | | *ORMDL3* | 3.372 | 9.83E-04 | 6.13E-04 | 0.011 |
| | | 38,082,658 | | *ORMDL3* | 5.515 | 1.81E-07 | 6.13E-04 | 0.011 |
| | | 38,108,710 | | | -5.683 | 8.27E-08 | 6.13E-04 | 0.039 |
| | | 38,119,198 | *GSDMA* | | 4.828 | 3.81E-06 | 6.13E-04 | 0.009 |
| | | 38,119,207 | *GSDMA* | | 6.261 | 5.14E-09 | 6.13E-04 | 0.009 |
| | | 38,119,649 | | *GSDMA* | 5.655 | 9.45E-08 | 6.13E-04 | 0.002 |
| | | 38,119,760 | | *GSDMA* | 4.753 | 5.23E-06 | 6.13E-04 | 0.011 |
| | | 38,120,234 | | *GSDMA* | 3.403 | 8.87E-04 | 6.13E-04 | 0.011 |
| rs9303280 | g.38074031.T>C | 37,887,061 | *MIEN1* | | 3.963 | 1.21E-04 | 8.49E-04 | 0.002 |
| | | 37,894,590 | | | 3.408 | 8.73E-04 | 8.49E-04 | 0.013 |
| | | 37,896,827 | | | 3.416 | 8.48E-04 | 8.49E-04 | 0.008 |
| | | 38,077,191 | | | 3.920 | 1.43E-04 | 8.49E-04 | 0.034 |
| | | 38,077,464 | | *ORMDL3* | 3.848 | 1.86E-04 | 8.49E-04 | 0.014 |
| | | 38,080,829 | | *ORMDL3* | 5.980 | 2.02E-08 | 8.49E-04 | 0.014 |
| | | 38,081,812 | | *ORMDL3* | 3.721 | 2.95E-04 | 8.49E-04 | 2.01E-04 |
| | | 38,081,829 | | *ORMDL3* | 3.444 | 7.73E-04 | 8.49E-04 | 2.01E-04 |
| | | 38,081,995 | | *ORMDL3* | 4.405 | 2.19E-05 | 8.49E-04 | 2.01E-04 |
| | | 38,082,022 | | *ORMDL3* | 4.799 | 4.31E-06 | 8.49E-04 | 6.40E-04 |
| | | 38,082,035 | | *ORMDL3* | 5.305 | 4.71E-07 | 8.49E-04 | 6.40E-04 |
| | | 38,082,073 | | *ORMDL3* | 5.047 | 1.48E-06 | 8.49E-04 | 6.40E-04 |
| | | 38,082,235 | | *ORMDL3* | 4.024 | 9.67E-05 | 8.49E-04 | 0.004 |
| | | 38,082,346 | | *ORMDL3* | 4.398 | 2.25E-05 | 8.49E-04 | 0.011 |
| | | 38,082,448 | | *ORMDL3* | 3.951 | 1.27E-04 | 8.49E-04 | 0.011 |
| | | 38,082,658 | | *ORMDL3* | 5.500 | 1.94E-07 | 8.49E-04 | 0.011 |
| | | 38,108,710 | | | -5.118 | 1.09E-06 | 8.49E-04 | 0.039 |
| | | 38,119,198 | *GSDMA* | | 4.691 | 6.77E-06 | 8.49E-04 | 0.009 |
| | | 38,119,207 | *GSDMA* | | 6.587 | 1.02E-09 | 8.49E-04 | 0.009 |
| | | 38,119,649 | | *GSDMA* | 5.445 | 2.49E-07 | 8.49E-04 | 0.002 |
| | | 38,119,760 | | *GSDMA* | 5.302 | 4.78E-07 | 8.49E-04 | 0.011 |
| rs9303281 | g.38074046.G>A | 37,887,061 | *MIEN1* | | 3.704 | 3.12E-04 | 2.60E-04 | 0.002 |
| | | 38,018,804 | | *IKZF3* | 3.417 | 8.44E-04 | 2.60E-04 | 0.046 |
| | | 38,077,191 | | | 3.827 | 2.00E-04 | 2.60E-04 | 0.034 |
| | | 38,077,464 | | *ORMDL3* | 3.639 | 3.93E-04 | 2.60E-04 | 0.014 |
| | | 38,080,829 | | *ORMDL3* | 5.536 | 1.64E-07 | 2.60E-04 | 0.014 |
| | | 38,081,492 | | *ORMDL3* | 3.585 | 4.75E-04 | 2.60E-04 | 2.01E-04 |
| | | 38,081,995 | | *ORMDL3* | 4.416 | 2.10E-05 | 2.60E-04 | 2.01E-04 |
| | | 38,082,022 | | *ORMDL3* | 5.112 | 1.11E-06 | 2.60E-04 | 6.40E-04 |

| | | | | | | | | |
|---|---|---|---|---|---|---|---|---|
| | | 38,082,035 | | *ORMDL3* | 5.307 | 4.66E-07 | 2.60E-04 | 6.40E-04 |
| | | 38,082,073 | | *ORMDL3* | 5.048 | 1.47E-06 | 2.60E-04 | 6.40E-04 |
| | | 38,082,235 | | *ORMDL3* | 4.077 | 7.89E-05 | 2.60E-04 | 0.004 |
| | | 38,082,346 | | *ORMDL3* | 4.493 | 1.53E-05 | 2.60E-04 | 0.011 |
| | | 38,082,448 | | *ORMDL3* | 3.848 | 1.86E-04 | 2.60E-04 | 0.011 |
| | | 38,082,658 | | *ORMDL3* | 5.521 | 1.76E-07 | 2.60E-04 | 0.011 |
| | | 38,108,464 | | | -3.396 | 9.08E-04 | 2.60E-04 | 0.007 |
| | | 38,108,710 | | | -4.944 | 2.31E-06 | 2.60E-04 | 0.039 |
| | | 38,119,198 | *GSDMA* | | 4.736 | 5.63E-06 | 2.60E-04 | 0.009 |
| | | 38,119,207 | *GSDMA* | | 6.586 | 1.02E-09 | 2.60E-04 | 0.009 |
| | | 38,119,649 | | *GSDMA* | 5.298 | 4.85E-07 | 2.60E-04 | 0.002 |
| | | 38,119,760 | | *GSDMA* | 5.134 | 1.01E-06 | 2.60E-04 | 0.011 |
| rs7219923 | g.38074518.C>T | 37,887,061 | *MIEN1* | | 3.704 | 3.12E-04 | 2.75E-04 | 0.002 |
| | | 38,018,804 | | *IKZF3* | 3.417 | 8.44E-04 | 2.75E-04 | 0.046 |
| | | 38,077,191 | | | 3.827 | 2.00E-04 | 2.75E-04 | 0.034 |
| | | 38,077,464 | | *ORMDL3* | 3.639 | 3.93E-04 | 2.75E-04 | 0.014 |
| | | 38,080,829 | | *ORMDL3* | 5.536 | 1.64E-07 | 2.75E-04 | 0.014 |
| | | 38,081,492 | | *ORMDL3* | 3.585 | 4.75E-04 | 2.75E-04 | 2.01E-04 |
| | | 38,081,995 | | *ORMDL3* | 4.416 | 2.10E-05 | 2.75E-04 | 2.01E-04 |
| | | 38,082,022 | | *ORMDL3* | 5.112 | 1.11E-06 | 2.75E-04 | 6.40E-04 |
| | | 38,082,035 | | *ORMDL3* | 5.307 | 4.66E-07 | 2.75E-04 | 6.40E-04 |
| | | 38,082,073 | | *ORMDL3* | 5.048 | 1.47E-06 | 2.75E-04 | 6.40E-04 |
| | | 38,082,235 | | *ORMDL3* | 4.077 | 7.89E-05 | 2.75E-04 | 0.004 |
| | | 38,082,346 | | *ORMDL3* | 4.493 | 1.53E-05 | 2.75E-04 | 0.011 |
| | | 38,082,448 | | *ORMDL3* | 3.848 | 1.86E-04 | 2.75E-04 | 0.011 |
| | | 38,082,658 | | *ORMDL3* | 5.521 | 1.76E-07 | 2.75E-04 | 0.011 |
| | | 38,108,464 | | | -3.396 | 9.08E-04 | 2.75E-04 | 0.007 |
| | | 38,108,710 | | | -4.944 | 2.31E-06 | 2.75E-04 | 0.039 |
| | | 38,119,198 | *GSDMA* | | 4.736 | 5.63E-06 | 2.75E-04 | 0.009 |
| | | 38,119,207 | *GSDMA* | | 6.586 | 1.02E-09 | 2.75E-04 | 0.009 |
| | | 38,119,649 | | *GSDMA* | 5.298 | 4.85E-07 | 2.75E-04 | 0.002 |
| | | 38,119,760 | | *GSDMA* | 5.134 | 1.01E-06 | 2.75E-04 | 0.011 |
| rs111379006 | g.38075100.A>G | 37,856,732 | | *ERBB2* | -3.668 | 3.56E-04 | | 0.009 |
| | | 37,886,749 | | *MIEN1* | 3.945 | 1.30E-04 | | 0.002 |
| | | 37,896,063 | | | 3.476 | 6.92E-04 | | 0.009 |
| | | 38,020,485 | *IKZF3* | | 3.962 | 1.22E-04 | | 0.005 |
| | | 38,080,829 | | *ORMDL3* | 3.995 | 1.08E-04 | | 0.014 |
| | | 38,096,333 | | | -3.664 | 3.60E-04 | | 0.033 |
| | | 38,096,571 | | | -4.780 | 4.67E-06 | | 0.033 |
| | | 38,096,578 | | | -5.642 | 1.00E-07 | | 0.033 |
| | | 38,096,648 | | | -3.887 | 1.61E-04 | | 0.033 |

| | | | | | | | | |
|---|---|---|---|---|---|---|---|---|
| rs7224129 | g.38075426.G>A | 37,887,061 | *MIEN1* | | 3.887 | 1.61E-04 | 3.47E-04 | 0.002 |
| | | 38,018,804 | | *IKZF3* | 3.402 | 8.89E-04 | 3.47E-04 | 0.046 |
| | | 38,077,191 | | | 3.731 | 2.84E-04 | 3.47E-04 | 0.034 |
| | | 38,077,464 | | *ORMDL3* | 3.804 | 2.18E-04 | 3.47E-04 | 0.014 |
| | | 38,080,829 | | *ORMDL3* | 5.629 | 1.06E-07 | 3.47E-04 | 0.014 |
| | | 38,081,492 | | *ORMDL3* | 3.436 | 7.93E-04 | 3.47E-04 | 2.01E-04 |
| | | 38,081,812 | | *ORMDL3* | 3.668 | 3.56E-04 | 3.47E-04 | 2.01E-04 |
| | | 38,081,829 | | *ORMDL3* | 3.371 | 9.87E-04 | 3.47E-04 | 2.01E-04 |
| | | 38,081,995 | | *ORMDL3* | 4.471 | 1.68E-05 | 3.47E-04 | 2.01E-04 |
| | | 38,082,022 | | *ORMDL3* | 4.825 | 3.85E-06 | 3.47E-04 | 6.40E-04 |
| | | 38,082,035 | | *ORMDL3* | 5.211 | 7.17E-07 | 3.47E-04 | 6.40E-04 |
| | | 38,082,073 | | *ORMDL3* | 5.075 | 1.31E-06 | 3.47E-04 | 6.40E-04 |
| | | 38,082,235 | | *ORMDL3* | 4.182 | 5.28E-05 | 3.47E-04 | 0.004 |
| | | 38,082,346 | | *ORMDL3* | 4.438 | 1.92E-05 | 3.47E-04 | 0.011 |
| | | 38,082,448 | | *ORMDL3* | 3.862 | 1.76E-04 | 3.47E-04 | 0.011 |
| | | 38,082,658 | | *ORMDL3* | 5.439 | 2.56E-07 | 3.47E-04 | 0.011 |
| | | 38,108,710 | | | -5.129 | 1.03E-06 | 3.47E-04 | 0.039 |
| | | 38,119,198 | *GSDMA* | | 4.743 | 5.45E-06 | 3.47E-04 | 0.009 |
| | | 38,119,207 | *GSDMA* | | 6.742 | 4.61E-10 | 3.47E-04 | 0.009 |
| | | 38,119,649 | | *GSDMA* | 5.439 | 2.57E-07 | 3.47E-04 | 0.002 |
| | | 38,119,760 | | *GSDMA* | 5.187 | 7.98E-07 | 3.47E-04 | 0.011 |
| rs8074437 | g.38076137.T>G | 37,887,061 | *MIEN1* | | 3.887 | 1.61E-04 | 3.47E-04 | 0.002 |
| | | 38,018,804 | | *IKZF3* | 3.402 | 8.89E-04 | 3.47E-04 | 0.046 |
| | | 38,077,191 | | | 3.731 | 2.84E-04 | 3.47E-04 | 0.034 |
| | | 38,077,464 | | *ORMDL3* | 3.804 | 2.18E-04 | 3.47E-04 | 0.014 |
| | | 38,080,829 | | *ORMDL3* | 5.629 | 1.06E-07 | 3.47E-04 | 0.014 |
| | | 38,081,492 | | *ORMDL3* | 3.436 | 7.93E-04 | 3.47E-04 | 2.01E-04 |
| | | 38,081,812 | | *ORMDL3* | 3.668 | 3.56E-04 | 3.47E-04 | 2.01E-04 |
| | | 38,081,829 | | *ORMDL3* | 3.371 | 9.87E-04 | 3.47E-04 | 2.01E-04 |
| | | 38,081,995 | | *ORMDL3* | 4.471 | 1.68E-05 | 3.47E-04 | 2.01E-04 |
| | | 38,082,022 | | *ORMDL3* | 4.825 | 3.85E-06 | 3.47E-04 | 6.40E-04 |
| | | 38,082,035 | | *ORMDL3* | 5.211 | 7.17E-07 | 3.47E-04 | 6.40E-04 |
| | | 38,082,073 | | *ORMDL3* | 5.075 | 1.31E-06 | 3.47E-04 | 6.40E-04 |
| | | 38,082,235 | | *ORMDL3* | 4.182 | 5.28E-05 | 3.47E-04 | 0.004 |
| | | 38,082,346 | | *ORMDL3* | 4.438 | 1.92E-05 | 3.47E-04 | 0.011 |
| | | 38,082,448 | | *ORMDL3* | 3.862 | 1.76E-04 | 3.47E-04 | 0.011 |
| | | 38,082,658 | | *ORMDL3* | 5.439 | 2.56E-07 | 3.47E-04 | 0.011 |
| | | 38,108,710 | | | -5.129 | 1.03E-06 | 3.47E-04 | 0.039 |
| | | 38,119,198 | *GSDMA* | | 4.743 | 5.45E-06 | 3.47E-04 | 0.009 |
| | | 38,119,207 | *GSDMA* | | 6.742 | 4.61E-10 | 3.47E-04 | 0.009 |
| | | 38,119,649 | | *GSDMA* | 5.439 | 2.57E-07 | 3.47E-04 | 0.002 |

| | | | | | | | | |
|---|---|---|---|---|---|---|---|---|
| | | 38,119,760 | | *GSDMA* | 5.187 | 7.98E-07 | 3.47E-04 | 0.011 |
| rs71971950 | g.38076198_38076201del | 37,887,061 | *MIEN1* | | 3.869 | 1.72E-04 | 3.77E-04 | 0.002 |
| | | 38,018,804 | | *IKZF3* | 3.544 | 5.48E-04 | 3.77E-04 | 0.046 |
| | | 38,077,191 | | | 3.879 | 1.66E-04 | 3.77E-04 | 0.034 |
| | | 38,077,464 | | *ORMDL3* | 3.821 | 2.05E-04 | 3.77E-04 | 0.014 |
| | | 38,080,829 | | *ORMDL3* | 5.496 | 1.97E-07 | 3.77E-04 | 0.014 |
| | | 38,081,492 | | *ORMDL3* | 3.538 | 5.60E-04 | 3.77E-04 | 2.01E-04 |
| | | 38,081,812 | | *ORMDL3* | 3.721 | 2.94E-04 | 3.77E-04 | 2.01E-04 |
| | | 38,081,995 | | *ORMDL3* | 4.434 | 1.95E-05 | 3.77E-04 | 2.01E-04 |
| | | 38,082,022 | | *ORMDL3* | 4.796 | 4.37E-06 | 3.77E-04 | 6.40E-04 |
| | | 38,082,035 | | *ORMDL3* | 5.294 | 4.94E-07 | 3.77E-04 | 6.40E-04 |
| | | 38,082,073 | | *ORMDL3* | 4.999 | 1.83E-06 | 3.77E-04 | 6.40E-04 |
| | | 38,082,235 | | *ORMDL3* | 4.087 | 7.59E-05 | 3.77E-04 | 0.004 |
| | | 38,082,346 | | *ORMDL3* | 4.323 | 3.04E-05 | 3.77E-04 | 0.011 |
| | | 38,082,448 | | *ORMDL3* | 3.726 | 2.89E-04 | 3.77E-04 | 0.011 |
| | | 38,082,658 | | *ORMDL3* | 5.321 | 4.39E-07 | 3.77E-04 | 0.011 |
| | | 38,108,710 | | | -5.015 | 1.70E-06 | 3.77E-04 | 0.039 |
| | | 38,119,198 | *GSDMA* | | 4.761 | 5.05E-06 | 3.77E-04 | 0.009 |
| | | 38,119,207 | *GSDMA* | | 6.721 | 5.13E-10 | 3.77E-04 | 0.009 |
| | | 38,119,649 | | *GSDMA* | 5.488 | 2.05E-07 | 3.77E-04 | 0.002 |
| | | 38,119,760 | | *GSDMA* | 5.029 | 1.60E-06 | 3.77E-04 | 0.011 |
| rs3859186 | g.38076329.T>C | 37,856,732 | | *ERBB2* | -3.668 | 3.56E-04 | | 0.009 |
| | | 37,886,749 | | *MIEN1* | 3.945 | 1.30E-04 | | 0.002 |
| | | 37,896,063 | | | 3.476 | 6.92E-04 | | 0.009 |
| | | 38,020,485 | *IKZF3* | | 3.962 | 1.22E-04 | | 0.005 |
| | | 38,080,829 | | *ORMDL3* | 3.995 | 1.08E-04 | | 0.014 |
| | | 38,096,333 | | | -3.664 | 3.60E-04 | | 0.033 |
| | | 38,096,571 | | | -4.780 | 4.67E-06 | | 0.033 |
| | | 38,096,578 | | | -5.642 | 1.00E-07 | | 0.033 |
| | | 38,096,648 | | | -3.887 | 1.61E-04 | | 0.033 |
| rs112260932 | g.38077185.C>T | 37,856,732 | | *ERBB2* | -3.668 | 3.56E-04 | | 0.009 |
| | | 37,886,749 | | *MIEN1* | 3.945 | 1.30E-04 | | 0.002 |
| | | 37,896,063 | | | 3.476 | 6.92E-04 | | 0.009 |
| | | 38,020,485 | *IKZF3* | | 3.962 | 1.22E-04 | | 0.005 |
| | | 38,080,829 | | *ORMDL3* | 3.995 | 1.08E-04 | | 0.014 |
| | | 38,096,333 | | | -3.664 | 3.60E-04 | | 0.033 |
| | | 38,096,571 | | | -4.780 | 4.67E-06 | | 0.033 |
| | | 38,096,578 | | | -5.642 | 1.00E-07 | | 0.033 |
| | | 38,096,648 | | | -3.887 | 1.61E-04 | | 0.033 |
| rs3169574 | g.38077485.T>C | 37,856,732 | | *ERBB2* | -3.668 | 3.56E-04 | | 0.009 |

| | | | | | | | | |
|---|---|---|---|---|---|---|---|---|
| | | 37,886,749 | | *MIEN1* | 3.945 | 1.30E-04 | | 0.002 |
| | | 37,896,063 | | | 3.476 | 6.92E-04 | | 0.009 |
| | | 38,020,485 | *IKZF3* | | 3.962 | 1.22E-04 | | 0.005 |
| | | 38,080,829 | | *ORMDL3* | 3.995 | 1.08E-04 | | 0.014 |
| | | 38,096,333 | | | -3.664 | 3.60E-04 | | 0.033 |
| | | 38,096,571 | | | -4.780 | 4.67E-06 | | 0.033 |
| | | 38,096,578 | | | -5.642 | 1.00E-07 | | 0.033 |
| | | 38,096,648 | | | -3.887 | 1.61E-04 | | 0.033 |
| rs78199107 | g.38077750.C>T | 37,856,732 | | *ERBB2* | -3.668 | 3.56E-04 | | 0.009 |
| | | 37,886,749 | | *MIEN1* | 3.945 | 1.30E-04 | | 0.002 |
| | | 37,896,063 | | | 3.476 | 6.92E-04 | | 0.009 |
| | | 38,020,485 | *IKZF3* | | 3.962 | 1.22E-04 | | 0.005 |
| | | 38,080,829 | | *ORMDL3* | 3.995 | 1.08E-04 | | 0.014 |
| | | 38,096,333 | | | -3.664 | 3.60E-04 | | 0.033 |
| | | 38,096,571 | | | -4.780 | 4.67E-06 | | 0.033 |
| | | 38,096,578 | | | -5.642 | 1.00E-07 | | 0.033 |
| | | 38,096,648 | | | -3.887 | 1.61E-04 | | 0.033 |
| rs4065275 | g.38080865.A>G | 37,887,061 | *MIEN1* | | 4.066 | 8.25E-05 | 8.19E-04 | 0.002 |
| | | 37,894,590 | | | 3.493 | 6.53E-04 | 8.19E-04 | 0.013 |
| | | 37,896,827 | | | 3.519 | 5.98E-04 | 8.19E-04 | 0.008 |
| | | 38,077,191 | | | 4.093 | 7.42E-05 | 8.19E-04 | 0.034 |
| | | 38,077,277 | | | 3.541 | 5.54E-04 | 8.19E-04 | 0.014 |
| | | 38,077,464 | | *ORMDL3* | 3.913 | 1.46E-04 | 8.19E-04 | 0.014 |
| | | 38,080,829 | | *ORMDL3* | 6.098 | 1.14E-08 | 8.19E-04 | 0.014 |
| | | 38,081,492 | | *ORMDL3* | 3.739 | 2.76E-04 | 8.19E-04 | 2.01E-04 |
| | | 38,081,812 | | *ORMDL3* | 4.374 | 2.48E-05 | 8.19E-04 | 2.01E-04 |
| | | 38,081,829 | | *ORMDL3* | 3.930 | 1.38E-04 | 8.19E-04 | 2.01E-04 |
| | | 38,081,995 | | *ORMDL3* | 4.684 | 6.99E-06 | 8.19E-04 | 2.01E-04 |
| | | 38,082,022 | | *ORMDL3* | 5.332 | 4.17E-07 | 8.19E-04 | 6.40E-04 |
| | | 38,082,035 | | *ORMDL3* | 5.503 | 1.91E-07 | 8.19E-04 | 6.40E-04 |
| | | 38,082,073 | | *ORMDL3* | 5.472 | 2.21E-07 | 8.19E-04 | 6.40E-04 |
| | | 38,082,235 | | *ORMDL3* | 4.155 | 5.86E-05 | 8.19E-04 | 0.004 |
| | | 38,082,346 | | *ORMDL3* | 4.515 | 1.41E-05 | 8.19E-04 | 0.011 |
| | | 38,082,448 | | *ORMDL3* | 4.041 | 9.06E-05 | 8.19E-04 | 0.011 |
| | | 38,082,658 | | *ORMDL3* | 5.781 | 5.23E-08 | 8.19E-04 | 0.011 |
| | | 38,108,710 | | | -5.233 | 6.49E-07 | 8.19E-04 | 0.039 |
| | | 38,119,198 | *GSDMA* | | 4.562 | 1.16E-05 | 8.19E-04 | 0.009 |
| | | 38,119,207 | *GSDMA* | | 6.442 | 2.10E-09 | 8.19E-04 | 0.009 |
| | | 38,119,649 | | *GSDMA* | 5.313 | 4.53E-07 | 8.19E-04 | 0.002 |
| | | 38,119,760 | | *GSDMA* | 5.332 | 4.18E-07 | 8.19E-04 | 0.011 |
| rs8076131 | g.38080912.G>A | 37,828,296 | | *PGAP3* | 3.578 | 4.88E-04 | 5.13E-04 | 0.002 |

| | | | | | | | | |
|---|---|---|---|---|---|---|---|---|
| | | 37,887,061 | *MIEN1* | | 4.863 | 3.28E-06 | 5.13E-04 | 0.002 |
| | | 38,077,191 | | | 4.199 | 4.94E-05 | 5.13E-04 | 0.034 |
| | | 38,077,277 | | | 3.563 | 5.13E-04 | 5.13E-04 | 0.014 |
| | | 38,077,323 | | *ORMDL3* | 3.450 | 7.56E-04 | 5.13E-04 | 0.014 |
| | | 38,077,464 | | *ORMDL3* | 4.561 | 1.16E-05 | 5.13E-04 | 0.014 |
| | | 38,080,829 | | *ORMDL3* | 3.969 | 1.19E-04 | 5.13E-04 | 0.014 |
| | | 38,081,812 | | *ORMDL3* | 4.505 | 1.46E-05 | 5.13E-04 | 2.01E-04 |
| | | 38,081,829 | | *ORMDL3* | 4.812 | 4.08E-06 | 5.13E-04 | 2.01E-04 |
| | | 38,081,995 | | *ORMDL3* | 6.156 | 8.62E-09 | 5.13E-04 | 2.01E-04 |
| | | 38,082,018 | | *ORMDL3* | 3.714 | 3.02E-04 | 5.13E-04 | 6.40E-04 |
| | | 38,082,022 | | *ORMDL3* | 7.092 | 7.61E-11 | 5.13E-04 | 6.40E-04 |
| | | 38,082,035 | | *ORMDL3* | 6.037 | 1.54E-08 | 5.13E-04 | 6.40E-04 |
| | | 38,082,073 | | *ORMDL3* | 6.438 | 2.14E-09 | 5.13E-04 | 6.40E-04 |
| | | 38,082,206 | | *ORMDL3* | 4.138 | 6.24E-05 | 5.13E-04 | 0.004 |
| | | 38,082,235 | | *ORMDL3* | 4.642 | 8.32E-06 | 5.13E-04 | 0.004 |
| | | 38,082,346 | | *ORMDL3* | 4.510 | 1.43E-05 | 5.13E-04 | 0.011 |
| | | 38,082,448 | | *ORMDL3* | 3.512 | 6.12E-04 | 5.13E-04 | 0.011 |
| | | 38,082,658 | | *ORMDL3* | 5.752 | 6.01E-08 | 5.13E-04 | 0.011 |
| | | 38,082,778 | | *ORMDL3* | 3.489 | 6.62E-04 | 5.13E-04 | 0.011 |
| | | 38,108,710 | | | -6.052 | 1.43E-08 | 5.13E-04 | 0.039 |
| | | 38,119,198 | *GSDMA* | | 4.695 | 6.67E-06 | 5.13E-04 | 0.009 |
| | | 38,119,207 | *GSDMA* | | 6.254 | 5.33E-09 | 5.13E-04 | 0.009 |
| | | 38,119,649 | | *GSDMA* | 5.694 | 7.86E-08 | 5.13E-04 | 0.002 |
| | | 38,119,760 | | *GSDMA* | 4.894 | 2.87E-06 | 5.13E-04 | 0.011 |
| rs12603332 | g.38082807.T>C | 37,887,061 | *MIEN1* | | 4.064 | 8.29E-05 | 4.23E-04 | 0.002 |
| | | 37,894,590 | | | 3.495 | 6.48E-04 | 4.23E-04 | 0.013 |
| | | 37,896,827 | | | 3.519 | 5.97E-04 | 4.23E-04 | 0.008 |
| | | 38,077,191 | | | 4.089 | 7.53E-05 | 4.23E-04 | 0.034 |
| | | 38,077,277 | | | 3.536 | 5.64E-04 | 4.23E-04 | 0.014 |
| | | 38,077,464 | | *ORMDL3* | 3.915 | 1.45E-04 | 4.23E-04 | 0.014 |
| | | 38,080,829 | | *ORMDL3* | 6.103 | 1.12E-08 | 4.23E-04 | 0.014 |
| | | 38,081,492 | | *ORMDL3* | 3.740 | 2.75E-04 | 4.23E-04 | 2.01E-04 |
| | | 38,081,812 | | *ORMDL3* | 4.374 | 2.48E-05 | 4.23E-04 | 2.01E-04 |
| | | 38,081,829 | | *ORMDL3* | 3.930 | 1.37E-04 | 4.23E-04 | 2.01E-04 |
| | | 38,081,995 | | *ORMDL3* | 4.683 | 7.00E-06 | 4.23E-04 | 2.01E-04 |
| | | 38,082,022 | | *ORMDL3* | 5.328 | 4.24E-07 | 4.23E-04 | 6.40E-04 |
| | | 38,082,035 | | *ORMDL3* | 5.496 | 1.97E-07 | 4.23E-04 | 6.40E-04 |
| | | 38,082,073 | | *ORMDL3* | 5.475 | 2.18E-07 | 4.23E-04 | 6.40E-04 |
| | | 38,082,235 | | *ORMDL3* | 4.151 | 5.95E-05 | 4.23E-04 | 0.004 |
| | | 38,082,346 | | *ORMDL3* | 4.505 | 1.46E-05 | 4.23E-04 | 0.011 |
| | | 38,082,448 | | *ORMDL3* | 4.038 | 9.17E-05 | 4.23E-04 | 0.011 |

| | | | | | | | | |
|---|---|---|---|---|---|---|---|---|
| | | 38,082,658 | | *ORMDL3* | 5.779 | 5.27E-08 | 4.23E-04 | 0.011 |
| | | 38,108,710 | | | -5.235 | 6.43E-07 | 4.23E-04 | 0.039 |
| | | 38,119,198 | *GSDMA* | | 4.559 | 1.17E-05 | 4.23E-04 | 0.009 |
| | | 38,119,207 | *GSDMA* | | 6.439 | 2.13E-09 | 4.23E-04 | 0.009 |
| | | 38,119,649 | | *GSDMA* | 5.314 | 4.53E-07 | 4.23E-04 | 0.002 |
| | | 38,119,760 | | *GSDMA* | 5.336 | 4.10E-07 | 4.23E-04 | 0.011 |
| rs3744246 | g.38084350.T>C | 37,834,869 | | *PGAP3* | 3.413 | 8.58E-04 | | 0.020 |
| | | 37,887,061 | *MIEN1* | | 3.599 | 4.54E-04 | | 0.002 |
| | | 38,077,098 | | | 3.695 | 3.23E-04 | | 0.034 |
| | | 38,077,464 | | *ORMDL3* | 6.824 | 3.04E-10 | | 0.014 |
| | | 38,077,703 | | *ORMDL3* | 3.681 | 3.39E-04 | | 0.014 |
| | | 38,081,492 | | *ORMDL3* | 3.959 | 1.23E-04 | | 2.01E-04 |
| | | 38,081,812 | | *ORMDL3* | 4.248 | 4.08E-05 | | 2.01E-04 |
| | | 38,081,829 | | *ORMDL3* | 4.077 | 7.89E-05 | | 2.01E-04 |
| | | 38,081,995 | | *ORMDL3* | 5.489 | 2.04E-07 | | 2.01E-04 |
| | | 38,082,018 | | *ORMDL3* | 3.807 | 2.16E-04 | | 6.40E-04 |
| | | 38,082,022 | | *ORMDL3* | 5.653 | 9.52E-08 | | 6.40E-04 |
| | | 38,082,035 | | *ORMDL3* | 5.283 | 5.20E-07 | | 6.40E-04 |
| | | 38,082,073 | | *ORMDL3* | 4.996 | 1.85E-06 | | 6.40E-04 |
| | | 38,082,206 | | *ORMDL3* | 4.842 | 3.58E-06 | | 0.004 |
| | | 38,082,235 | | *ORMDL3* | 4.637 | 8.50E-06 | | 0.004 |
| | | 38,082,346 | | *ORMDL3* | 4.928 | 2.48E-06 | | 0.011 |
| | | 38,082,448 | | *ORMDL3* | 3.940 | 1.32E-04 | | 0.011 |
| | | 38,082,658 | | *ORMDL3* | 6.406 | 2.51E-09 | | 0.011 |
| | | 38,084,331 | *ORMDL3* | | 4.318 | 3.10E-05 | | 0.005 |
| | | 38,084,372 | *ORMDL3* | | 4.152 | 5.93E-05 | | 0.005 |
| | | 38,084,377 | *ORMDL3* | | 5.051 | 1.46E-06 | | 0.005 |
| | | 38,084,450 | *ORMDL3* | | 3.997 | 1.07E-04 | | 0.005 |
| | | 38,084,459 | *ORMDL3* | | 3.602 | 4.48E-04 | | 0.005 |
| | | 38,084,474 | *ORMDL3* | | 4.184 | 5.23E-05 | | 0.005 |
| | | 38,108,710 | | | -3.408 | 8.72E-04 | | 0.039 |
| | | 38,119,649 | | *GSDMA* | 3.831 | 1.98E-04 | | 0.002 |
| | | 38,119,760 | | *GSDMA* | 5.087 | 1.24E-06 | | 0.011 |
| rs4795402 | g.38085385.A>C | 37,834,869 | | *PGAP3* | 3.420 | 8.38E-04 | | 0.020 |
| | | 37,910,990 | | | 3.504 | 6.30E-04 | | 0.035 |
| | | 38,077,464 | | *ORMDL3* | 5.523 | 1.75E-07 | | 0.014 |
| | | 38,080,829 | | *ORMDL3* | 3.701 | 3.16E-04 | | 0.014 |
| | | 38,081,492 | | *ORMDL3* | 4.362 | 2.60E-05 | | 2.01E-04 |
| | | 38,081,812 | | *ORMDL3* | 3.938 | 1.33E-04 | | 2.01E-04 |
| | | 38,081,995 | | *ORMDL3* | 3.828 | 2.00E-04 | | 2.01E-04 |
| | | 38,082,022 | | *ORMDL3* | 3.787 | 2.32E-04 | | 6.40E-04 |

| | | | | | | | |
|---|---|---|---|---|---|---|---|
| | | 38,082,035 | | *ORMDL3* | 4.472 | 1.67E-05 | 6.40E-04 |
| | | 38,082,073 | | *ORMDL3* | 3.950 | 1.27E-04 | 6.40E-04 |
| | | 38,082,206 | | *ORMDL3* | 3.626 | 4.12E-04 | 0.004 |
| | | 38,082,235 | | *ORMDL3* | 4.047 | 8.84E-05 | 0.004 |
| | | 38,082,346 | | *ORMDL3* | 4.737 | 5.60E-06 | 0.011 |
| | | 38,082,448 | | *ORMDL3* | 4.306 | 3.24E-05 | 0.011 |
| | | 38,082,658 | | *ORMDL3* | 6.072 | 1.30E-08 | 0.011 |
| | | 38,084,377 | *ORMDL3* | | 4.428 | 2.00E-05 | 0.005 |
| | | 38,084,474 | *ORMDL3* | | 3.635 | 3.99E-04 | 0.005 |
| | | 38,119,649 | | *GSDMA* | 3.408 | 8.70E-04 | 0.002 |
| | | 38,119,760 | | *GSDMA* | 5.284 | 5.18E-07 | 0.011 |
| rs4795403 | g.38085722.T>C | 37,834,869 | | *PGAP3* | 3.413 | 8.58E-04 | 0.020 |
| | | 37,887,061 | *MIEN1* | | 3.599 | 4.54E-04 | 0.002 |
| | | 38,077,098 | | | 3.695 | 3.23E-04 | 0.034 |
| | | 38,077,464 | | *ORMDL3* | 6.824 | 3.04E-10 | 0.014 |
| | | 38,077,703 | | *ORMDL3* | 3.681 | 3.39E-04 | 0.014 |
| | | 38,081,492 | | *ORMDL3* | 3.959 | 1.23E-04 | 2.01E-04 |
| | | 38,081,812 | | *ORMDL3* | 4.248 | 4.08E-05 | 2.01E-04 |
| | | 38,081,829 | | *ORMDL3* | 4.077 | 7.89E-05 | 2.01E-04 |
| | | 38,081,995 | | *ORMDL3* | 5.489 | 2.04E-07 | 2.01E-04 |
| | | 38,082,018 | | *ORMDL3* | 3.807 | 2.16E-04 | 6.40E-04 |
| | | 38,082,022 | | *ORMDL3* | 5.653 | 9.52E-08 | 6.40E-04 |
| | | 38,082,035 | | *ORMDL3* | 5.283 | 5.20E-07 | 6.40E-04 |
| | | 38,082,073 | | *ORMDL3* | 4.996 | 1.85E-06 | 6.40E-04 |
| | | 38,082,206 | | *ORMDL3* | 4.842 | 3.58E-06 | 0.004 |
| | | 38,082,235 | | *ORMDL3* | 4.637 | 8.50E-06 | 0.004 |
| | | 38,082,346 | | *ORMDL3* | 4.928 | 2.48E-06 | 0.011 |
| | | 38,082,448 | | *ORMDL3* | 3.940 | 1.32E-04 | 0.011 |
| | | 38,082,658 | | *ORMDL3* | 6.406 | 2.51E-09 | 0.011 |
| | | 38,084,331 | *ORMDL3* | | 4.318 | 3.10E-05 | 0.005 |
| | | 38,084,372 | *ORMDL3* | | 4.152 | 5.93E-05 | 0.005 |
| | | 38,084,377 | *ORMDL3* | | 5.051 | 1.46E-06 | 0.005 |
| | | 38,084,450 | *ORMDL3* | | 3.997 | 1.07E-04 | 0.005 |
| | | 38,084,459 | *ORMDL3* | | 3.602 | 4.48E-04 | 0.005 |
| | | 38,084,474 | *ORMDL3* | | 4.184 | 5.23E-05 | 0.005 |
| | | 38,108,710 | | | -3.408 | 8.72E-04 | 0.039 |
| | | 38,119,649 | | *GSDMA* | 3.831 | 1.98E-04 | 0.002 |
| | | 38,119,760 | | *GSDMA* | 5.087 | 1.24E-06 | 0.011 |
| rs4795404 | g.38085791.A>C | 37,834,869 | | *PGAP3* | 3.413 | 8.58E-04 | 0.020 |
| | | 37,887,061 | *MIEN1* | | 3.599 | 4.54E-04 | 0.002 |
| | | 38,077,098 | | | 3.695 | 3.23E-04 | 0.034 |

| | | | | | | | |
|---|---|---|---|---|---|---|---|
| | | 38,077,464 | | *ORMDL3* | 6.824 | 3.04E-10 | 0.014 |
| | | 38,077,703 | | *ORMDL3* | 3.681 | 3.39E-04 | 0.014 |
| | | 38,081,492 | | *ORMDL3* | 3.959 | 1.23E-04 | 2.01E-04 |
| | | 38,081,812 | | *ORMDL3* | 4.248 | 4.08E-05 | 2.01E-04 |
| | | 38,081,829 | | *ORMDL3* | 4.077 | 7.89E-05 | 2.01E-04 |
| | | 38,081,995 | | *ORMDL3* | 5.489 | 2.04E-07 | 2.01E-04 |
| | | 38,082,018 | | *ORMDL3* | 3.807 | 2.16E-04 | 6.40E-04 |
| | | 38,082,022 | | *ORMDL3* | 5.653 | 9.52E-08 | 6.40E-04 |
| | | 38,082,035 | | *ORMDL3* | 5.283 | 5.20E-07 | 6.40E-04 |
| | | 38,082,073 | | *ORMDL3* | 4.996 | 1.85E-06 | 6.40E-04 |
| | | 38,082,206 | | *ORMDL3* | 4.842 | 3.58E-06 | 0.004 |
| | | 38,082,235 | | *ORMDL3* | 4.637 | 8.50E-06 | 0.004 |
| | | 38,082,346 | | *ORMDL3* | 4.928 | 2.48E-06 | 0.011 |
| | | 38,082,448 | | *ORMDL3* | 3.940 | 1.32E-04 | 0.011 |
| | | 38,082,658 | | *ORMDL3* | 6.406 | 2.51E-09 | 0.011 |
| | | 38,084,331 | *ORMDL3* | | 4.318 | 3.10E-05 | 0.005 |
| | | 38,084,372 | *ORMDL3* | | 4.152 | 5.93E-05 | 0.005 |
| | | 38,084,377 | *ORMDL3* | | 5.051 | 1.46E-06 | 0.005 |
| | | 38,084,450 | *ORMDL3* | | 3.997 | 1.07E-04 | 0.005 |
| | | 38,084,459 | *ORMDL3* | | 3.602 | 4.48E-04 | 0.005 |
| | | 38,084,474 | *ORMDL3* | | 4.184 | 5.23E-05 | 0.005 |
| | | 38,108,710 | | | -3.408 | 8.72E-04 | 0.039 |
| | | 38,119,649 | | *GSDMA* | 3.831 | 1.98E-04 | 0.002 |
| | | 38,119,760 | | *GSDMA* | 5.087 | 1.24E-06 | 0.011 |
| rs145604770 | g.38086736.A>G | 37,856,732 | | *ERBB2* | -3.668 | 3.56E-04 | 0.009 |
| | | 37,886,749 | | *MIEN1* | 3.945 | 1.30E-04 | 0.002 |
| | | 37,896,063 | | | 3.476 | 6.92E-04 | 0.009 |
| | | 38,020,485 | *IKZF3* | | 3.962 | 1.22E-04 | 0.005 |
| | | 38,080,829 | | *ORMDL3* | 3.995 | 1.08E-04 | 0.014 |
| | | 38,096,333 | | | -3.664 | 3.60E-04 | 0.033 |
| | | 38,096,571 | | | -4.780 | 4.67E-06 | 0.033 |
| | | 38,096,578 | | | -5.642 | 1.00E-07 | 0.033 |
| | | 38,096,648 | | | -3.887 | 1.61E-04 | 0.033 |
| rs7224908 | g.38086854.A>G | 37,886,797 | *MIEN1* | | 3.377 | 9.66E-04 | 0.002 |
| | | 37,887,061 | *MIEN1* | | 3.403 | 8.87E-04 | 0.002 |
| | | 38,077,098 | | | 3.660 | 3.66E-04 | 0.034 |
| | | 38,077,464 | | *ORMDL3* | 6.953 | 1.56E-10 | 0.014 |
| | | 38,077,703 | | *ORMDL3* | 3.639 | 3.93E-04 | 0.014 |
| | | 38,081,492 | | *ORMDL3* | 3.890 | 1.59E-04 | 2.01E-04 |
| | | 38,081,812 | | *ORMDL3* | 4.242 | 4.18E-05 | 2.01E-04 |
| | | 38,081,829 | | *ORMDL3* | 3.998 | 1.06E-04 | 2.01E-04 |

| | | | | | | | |
|---|---|---|---|---|---|---|---|
| | | 38,081,995 | | *ORMDL3* | 5.480 | 2.12E-07 | 2.01E-04 |
| | | 38,082,018 | | *ORMDL3* | 3.925 | 1.40E-04 | 6.40E-04 |
| | | 38,082,022 | | *ORMDL3* | 5.825 | 4.24E-08 | 6.40E-04 |
| | | 38,082,035 | | *ORMDL3* | 5.334 | 4.12E-07 | 6.40E-04 |
| | | 38,082,073 | | *ORMDL3* | 4.999 | 1.83E-06 | 6.40E-04 |
| | | 38,082,206 | | *ORMDL3* | 4.834 | 3.71E-06 | 0.004 |
| | | 38,082,235 | | *ORMDL3* | 4.683 | 7.01E-06 | 0.004 |
| | | 38,082,346 | | *ORMDL3* | 5.052 | 1.45E-06 | 0.011 |
| | | 38,082,448 | | *ORMDL3* | 3.915 | 1.45E-04 | 0.011 |
| | | 38,082,658 | | *ORMDL3* | 6.539 | 1.29E-09 | 0.011 |
| | | 38,084,331 | *ORMDL3* | | 4.337 | 2.87E-05 | 0.005 |
| | | 38,084,372 | *ORMDL3* | | 4.222 | 4.52E-05 | 0.005 |
| | | 38,084,377 | *ORMDL3* | | 5.073 | 1.32E-06 | 0.005 |
| | | 38,084,450 | *ORMDL3* | | 3.928 | 1.38E-04 | 0.005 |
| | | 38,084,459 | *ORMDL3* | | 3.559 | 5.21E-04 | 0.005 |
| | | 38,084,474 | *ORMDL3* | | 4.034 | 9.29E-05 | 0.005 |
| | | 38,108,710 | | | -3.514 | 6.08E-04 | 0.039 |
| | | 38,119,649 | | *GSDMA* | 3.823 | 2.04E-04 | 0.002 |
| | | 38,119,760 | | *GSDMA* | 5.282 | 5.23E-07 | 0.011 |
| rs12946393 | g.38087429.T>G | 37,886,797 | *MIEN1* | | 3.377 | 9.66E-04 | 0.002 |
| | | 37,887,061 | *MIEN1* | | 3.403 | 8.87E-04 | 0.002 |
| | | 38,077,098 | | | 3.660 | 3.66E-04 | 0.034 |
| | | 38,077,464 | | *ORMDL3* | 6.953 | 1.56E-10 | 0.014 |
| | | 38,077,703 | | *ORMDL3* | 3.639 | 3.93E-04 | 0.014 |
| | | 38,081,492 | | *ORMDL3* | 3.890 | 1.59E-04 | 2.01E-04 |
| | | 38,081,812 | | *ORMDL3* | 4.242 | 4.18E-05 | 2.01E-04 |
| | | 38,081,829 | | *ORMDL3* | 3.998 | 1.06E-04 | 2.01E-04 |
| | | 38,081,995 | | *ORMDL3* | 5.480 | 2.12E-07 | 2.01E-04 |
| | | 38,082,018 | | *ORMDL3* | 3.925 | 1.40E-04 | 6.40E-04 |
| | | 38,082,022 | | *ORMDL3* | 5.825 | 4.24E-08 | 6.40E-04 |
| | | 38,082,035 | | *ORMDL3* | 5.334 | 4.12E-07 | 6.40E-04 |
| | | 38,082,073 | | *ORMDL3* | 4.999 | 1.83E-06 | 6.40E-04 |
| | | 38,082,206 | | *ORMDL3* | 4.834 | 3.71E-06 | 0.004 |
| | | 38,082,235 | | *ORMDL3* | 4.683 | 7.01E-06 | 0.004 |
| | | 38,082,346 | | *ORMDL3* | 5.052 | 1.45E-06 | 0.011 |
| | | 38,082,448 | | *ORMDL3* | 3.915 | 1.45E-04 | 0.011 |
| | | 38,082,658 | | *ORMDL3* | 6.539 | 1.29E-09 | 0.011 |
| | | 38,084,331 | *ORMDL3* | | 4.337 | 2.87E-05 | 0.005 |
| | | 38,084,372 | *ORMDL3* | | 4.222 | 4.52E-05 | 0.005 |
| | | 38,084,377 | *ORMDL3* | | 5.073 | 1.32E-06 | 0.005 |
| | | 38,084,450 | *ORMDL3* | | 3.928 | 1.38E-04 | 0.005 |

| | | | | | | | | |
|---|---|---|---|---|---|---|---|---|
| | | | 38,084,459 | *ORMDL3* | | 3.559 | 5.21E-04 | 0.005 |
| | | | 38,084,474 | *ORMDL3* | | 4.034 | 9.29E-05 | 0.005 |
| | | | 38,108,710 | | | -3.514 | 6.08E-04 | 0.039 |
| | | | 38,119,649 | | *GSDMA* | 3.823 | 2.04E-04 | 0.002 |
| | | | 38,119,760 | | *GSDMA* | 5.282 | 5.23E-07 | 0.011 |
| | rs35557848 | g.38087439.T>C | 37,846,697 | | *ERBB2* | 3.602 | 4.49E-04 | 0.027 |
| | | | 38,024,331 | | | -3.374 | 9.77E-04 | 0.017 |
| | | | 38,083,308 | | *ORMDL3* | 4.527 | 1.34E-05 | 0.011 |
| | | | 38,108,710 | | | -3.404 | 8.83E-04 | 0.039 |
| | | | 38,119,198 | *GSDMA* | | 3.389 | 9.29E-04 | 0.009 |
| | | | 38,119,207 | *GSDMA* | | 4.795 | 4.37E-06 | 0.009 |
| Sixth block | rs112191651 | g.38088150.T>C | 37,856,732 | | *ERBB2* | -3.668 | 3.56E-04 | 0.009 |
| | | | 37,886,749 | | *MIEN1* | 3.945 | 1.30E-04 | 0.002 |
| | | | 37,896,063 | | | 3.476 | 6.92E-04 | 0.009 |
| | | | 38,020,485 | *IKZF3* | | 3.962 | 1.22E-04 | 0.005 |
| | | | 38,080,829 | | *ORMDL3* | 3.995 | 1.08E-04 | 0.014 |
| | | | 38,096,333 | | | -3.664 | 3.60E-04 | 0.033 |
| | | | 38,096,571 | | | -4.780 | 4.67E-06 | 0.033 |
| | | | 38,096,578 | | | -5.642 | 1.00E-07 | 0.033 |
| | | | 38,096,648 | | | -3.887 | 1.61E-04 | 0.033 |
| | rs4795405 | g.38088417.T>C | 37,828,296 | | *PGAP3* | 3.375 | 9.73E-04 | 0.002 |
| | | | 37,887,061 | *MIEN1* | | 4.568 | 1.13E-05 | 0.002 |
| | | | 37,894,590 | | | 3.502 | 6.34E-04 | 0.013 |
| | | | 37,896,827 | | | 3.398 | 9.02E-04 | 0.008 |
| | | | 38,077,191 | | | 4.147 | 6.03E-05 | 0.034 |
| | | | 38,077,277 | | | 3.677 | 3.44E-04 | 0.014 |
| | | | 38,077,323 | | *ORMDL3* | 3.646 | 3.85E-04 | 0.014 |
| | | | 38,077,464 | | *ORMDL3* | 4.619 | 9.16E-06 | 0.014 |
| | | | 38,080,829 | | *ORMDL3* | 4.020 | 9.80E-05 | 0.014 |
| | | | 38,081,812 | | *ORMDL3* | 4.417 | 2.08E-05 | 2.01E-04 |
| | | | 38,081,829 | | *ORMDL3* | 4.725 | 5.89E-06 | 2.01E-04 |
| | | | 38,081,995 | | *ORMDL3* | 6.008 | 1.77E-08 | 2.01E-04 |
| | | | 38,082,018 | | *ORMDL3* | 3.735 | 2.80E-04 | 6.40E-04 |
| | | | 38,082,022 | | *ORMDL3* | 7.347 | 1.99E-11 | 6.40E-04 |
| | | | 38,082,035 | | *ORMDL3* | 6.159 | 8.51E-09 | 6.40E-04 |
| | | | 38,082,073 | | *ORMDL3* | 6.360 | 3.16E-09 | 6.40E-04 |
| | | | 38,082,206 | | *ORMDL3* | 3.899 | 1.54E-04 | 0.004 |
| | | | 38,082,235 | | *ORMDL3* | 4.506 | 1.45E-05 | 0.004 |
| | | | 38,082,346 | | *ORMDL3* | 4.604 | 9.74E-06 | 0.011 |
| | | | 38,082,448 | | *ORMDL3* | 3.512 | 6.11E-04 | 0.011 |
| | | | 38,082,658 | | *ORMDL3* | 5.914 | 2.77E-08 | 0.011 |

| | | | | | | | |
|---|---|---|---|---|---|---|---|
| | | 38,082,778 | | *ORMDL3* | 3.690 | 3.29E-04 | 0.011 |
| | | 38,108,710 | | | -6.125 | 1.00E-08 | 0.039 |
| | | 38,119,198 | *GSDMA* | | 4.590 | 1.03E-05 | 0.009 |
| | | 38,119,207 | *GSDMA* | | 5.962 | 2.21E-08 | 0.009 |
| | | 38,119,649 | | *GSDMA* | 5.612 | 1.15E-07 | 0.002 |
| | | 38,119,760 | | *GSDMA* | 5.155 | 9.20E-07 | 0.011 |
| rs56199421 | g.38090808.C>T | 37,894,590 | | | 4.019 | 9.86E-05 | 0.013 |
| | | 38,077,051 | | | 3.375 | 9.73E-04 | 0.034 |
| | | 38,077,464 | | *ORMDL3* | 5.121 | 1.07E-06 | 0.014 |
| | | 38,081,812 | | *ORMDL3* | 3.850 | 1.85E-04 | 2.01E-04 |
| | | 38,081,829 | | *ORMDL3* | 4.098 | 7.28E-05 | 2.01E-04 |
| | | 38,081,995 | | *ORMDL3* | 3.936 | 1.34E-04 | 2.01E-04 |
| | | 38,082,022 | | *ORMDL3* | 4.732 | 5.72E-06 | 6.40E-04 |
| | | 38,082,035 | | *ORMDL3* | 4.060 | 8.43E-05 | 6.40E-04 |
| | | 38,082,073 | | *ORMDL3* | 4.197 | 4.97E-05 | 6.40E-04 |
| | | 38,082,658 | | *ORMDL3* | 5.273 | 5.44E-07 | 0.011 |
| | | 38,108,710 | | | -6.702 | 5.66E-10 | 0.039 |
| | | 38,119,071 | *GSDMA* | | 5.224 | 6.76E-07 | 0.009 |
| | | 38,119,132 | *GSDMA* | | 3.488 | 6.64E-04 | 0.009 |
| | | 38,119,198 | *GSDMA* | | 7.503 | 8.71E-12 | 0.009 |
| | | 38,119,207 | *GSDMA* | | 8.224 | 1.76E-13 | 0.009 |
| | | 38,119,649 | | *GSDMA* | 6.490 | 1.65E-09 | 0.002 |
| | | 38,119,760 | | *GSDMA* | 4.168 | 5.57E-05 | 0.011 |
| | | 38,120,234 | | *GSDMA* | 3.509 | 6.19E-04 | 0.011 |
| | | 38,120,370 | | *GSDMA* | 3.875 | 1.69E-04 | 0.011 |
| | | 38,128,699 | | *GSDMA* | -4.177 | 5.38E-05 | 0.006 |
| rs8065244 | g.38091228.G>C | 37,879,672 | | *ERBB2* | 3.591 | 4.66E-04 | 0.002 |
| | | 37,910,684 | | | 3.413 | 8.56E-04 | 0.013 |
| | | 38,083,874 | | *ORMDL3* | 4.119 | 6.73E-05 | 0.011 |
| | | 38,096,333 | | | -5.290 | 5.05E-07 | 0.033 |
| | | 38,096,571 | | | -10.420 | 7.42E-19 | 0.033 |
| | | 38,096,578 | | | -9.855 | 1.87E-17 | 0.033 |
| | | 38,096,648 | | | -4.993 | 1.87E-06 | 0.033 |
| rs76285844 | g.38092183.C>T | 37,828,195 | | *PGAP3* | 3.496 | 6.48E-04 | |
| | | 37,976,456 | | *IKZF3* | -5.057 | 1.42E-06 | 0.039 |
| | | 38,109,639 | | | 3.616 | 4.26E-04 | 0.002 |
| rs8079416 | g.38092713.C>T | 37,834,869 | | *PGAP3* | 3.453 | 7.49E-04 | 0.020 |
| | | 37,910,697 | | | 3.491 | 6.58E-04 | 0.035 |
| | | 38,077,277 | | | 4.203 | 4.86E-05 | 0.014 |
| | | 38,077,464 | | *ORMDL3* | 4.071 | 8.07E-05 | 0.014 |
| | | 38,080,829 | | *ORMDL3* | 3.663 | 3.61E-04 | 0.014 |

| | | | | | | | |
|---|---|---|---|---|---|---|---|
| | | 38,081,530 | | *ORMDL3* | 3.708 | 3.08E-04 | 2.01E-04 |
| | | 38,081,812 | | *ORMDL3* | 4.038 | 9.16E-05 | 2.01E-04 |
| | | 38,082,022 | | *ORMDL3* | 3.625 | 4.13E-04 | 6.40E-04 |
| | | 38,082,073 | | *ORMDL3* | 4.886 | 2.98E-06 | 6.40E-04 |
| | | 38,082,346 | | *ORMDL3* | 3.433 | 8.01E-04 | 0.011 |
| | | 38,082,658 | | *ORMDL3* | 4.551 | 1.21E-05 | 0.011 |
| | | 38,108,710 | | | -6.111 | 1.07E-08 | 0.039 |
| | | 38,119,071 | *GSDMA* | | 4.281 | 3.59E-05 | 0.009 |
| | | 38,119,198 | *GSDMA* | | 6.240 | 5.72E-09 | 0.009 |
| | | 38,119,207 | *GSDMA* | | 7.498 | 8.96E-12 | 0.009 |
| | | 38,119,649 | | *GSDMA* | 6.524 | 1.39E-09 | 0.002 |
| | | 38,119,760 | | *GSDMA* | 4.665 | 7.58E-06 | 0.011 |
| | | 38,120,234 | | *GSDMA* | 3.402 | 8.90E-04 | 0.011 |
| | | 38,128,699 | | *GSDMA* | -4.073 | 8.01E-05 | 0.006 |
| rs35123741 | g.38092930.G>A | 37,834,869 | | *PGAP3* | 3.453 | 7.49E-04 | 0.020 |
| | | 37,910,697 | | | 3.491 | 6.58E-04 | 0.035 |
| | | 38,077,277 | | | 4.203 | 4.86E-05 | 0.014 |
| | | 38,077,464 | | *ORMDL3* | 4.071 | 8.07E-05 | 0.014 |
| | | 38,080,829 | | *ORMDL3* | 3.663 | 3.61E-04 | 0.014 |
| | | 38,081,530 | | *ORMDL3* | 3.708 | 3.08E-04 | 2.01E-04 |
| | | 38,081,812 | | *ORMDL3* | 4.038 | 9.16E-05 | 2.01E-04 |
| | | 38,082,022 | | *ORMDL3* | 3.625 | 4.13E-04 | 6.40E-04 |
| | | 38,082,073 | | *ORMDL3* | 4.886 | 2.98E-06 | 6.40E-04 |
| | | 38,082,346 | | *ORMDL3* | 3.433 | 8.01E-04 | 0.011 |
| | | 38,082,658 | | *ORMDL3* | 4.551 | 1.21E-05 | 0.011 |
| | | 38,108,710 | | | -6.111 | 1.07E-08 | 0.039 |
| | | 38,119,071 | *GSDMA* | | 4.281 | 3.59E-05 | 0.009 |
| | | 38,119,198 | *GSDMA* | | 6.240 | 5.72E-09 | 0.009 |
| | | 38,119,207 | *GSDMA* | | 7.498 | 8.96E-12 | 0.009 |
| | | 38,119,649 | | *GSDMA* | 6.524 | 1.39E-09 | 0.002 |
| | | 38,119,760 | | *GSDMA* | 4.665 | 7.58E-06 | 0.011 |
| | | 38,120,234 | | *GSDMA* | 3.402 | 8.90E-04 | 0.011 |
| | | 38,128,699 | | *GSDMA* | -4.073 | 8.01E-05 | 0.006 |
| rs72832957 | g.38093085.G>A | 37,911,537 | | | -3.392 | 9.19E-04 | 0.035 |
| | | 37,911,661 | | | -3.420 | 8.38E-04 | 0.035 |
| | | 38,083,874 | | *ORMDL3* | 3.528 | 5.80E-04 | 0.011 |
| | | 38,096,333 | | | -5.589 | 1.28E-07 | 0.033 |
| | | 38,096,571 | | | -10.118 | 4.18E-18 | 0.033 |
| | | 38,096,578 | | | -9.592 | 8.40E-17 | 0.033 |
| | | 38,096,648 | | | -4.297 | 3.37E-05 | 0.033 |
| rs7219492 | g.38093113.G>A | 37,911,661 | | | -3.459 | 7.34E-04 | 0.035 |

| | | | | | | | |
|---|---|---|---|---|---|---|---|
| | | 38,024,634 | | | 3.786 | 2.32E-04 | 0.046 |
| | | 38,083,874 | | *ORMDL3* | 4.282 | 3.56E-05 | 0.011 |
| | | 38,096,333 | | | -4.532 | 1.31E-05 | 0.033 |
| | | 38,096,571 | | | -6.654 | 7.21E-10 | 0.033 |
| | | 38,096,578 | | | -6.319 | 3.87E-09 | 0.033 |
| rs12601749 | g.38093315.A>G | 37,894,590 | | | 4.022 | 9.72E-05 | 0.013 |
| | | 38,077,051 | | | 3.381 | 9.53E-04 | 0.034 |
| | | 38,077,464 | | *ORMDL3* | 5.159 | 9.02E-07 | 0.014 |
| | | 38,081,812 | | *ORMDL3* | 3.876 | 1.68E-04 | 2.01E-04 |
| | | 38,081,829 | | *ORMDL3* | 4.103 | 7.15E-05 | 2.01E-04 |
| | | 38,081,995 | | *ORMDL3* | 3.904 | 1.51E-04 | 2.01E-04 |
| | | 38,082,022 | | *ORMDL3* | 4.713 | 6.20E-06 | 6.40E-04 |
| | | 38,082,035 | | *ORMDL3* | 4.030 | 9.43E-05 | 6.40E-04 |
| | | 38,082,073 | | *ORMDL3* | 4.244 | 4.14E-05 | 6.40E-04 |
| | | 38,082,658 | | *ORMDL3* | 5.266 | 5.61E-07 | 0.011 |
| | | 38,108,710 | | | -6.712 | 5.38E-10 | 0.039 |
| | | 38,119,071 | *GSDMA* | | 5.210 | 7.21E-07 | 0.009 |
| | | 38,119,132 | *GSDMA* | | 3.479 | 6.86E-04 | 0.009 |
| | | 38,119,198 | *GSDMA* | | 7.500 | 8.85E-12 | 0.009 |
| | | 38,119,207 | *GSDMA* | | 8.154 | 2.59E-13 | 0.009 |
| | | 38,119,649 | | *GSDMA* | 6.482 | 1.72E-09 | 0.002 |
| | | 38,119,760 | | *GSDMA* | 4.222 | 4.52E-05 | 0.011 |
| | | 38,120,234 | | *GSDMA* | 3.485 | 6.72E-04 | 0.011 |
| | | 38,120,370 | | *GSDMA* | 3.846 | 1.87E-04 | 0.011 |
| | | 38,128,699 | | *GSDMA* | -4.253 | 4.00E-05 | 0.006 |
| rs12603481 | g.38093339.A>G | 37,834,869 | | *PGAP3* | 3.453 | 7.49E-04 | 0.020 |
| | | 37,910,697 | | | 3.491 | 6.58E-04 | 0.035 |
| | | 38,077,277 | | | 4.203 | 4.86E-05 | 0.014 |
| | | 38,077,464 | | *ORMDL3* | 4.071 | 8.07E-05 | 0.014 |
| | | 38,080,829 | | *ORMDL3* | 3.663 | 3.61E-04 | 0.014 |
| | | 38,081,530 | | *ORMDL3* | 3.708 | 3.08E-04 | 2.01E-04 |
| | | 38,081,812 | | *ORMDL3* | 4.038 | 9.16E-05 | 2.01E-04 |
| | | 38,082,022 | | *ORMDL3* | 3.625 | 4.13E-04 | 6.40E-04 |
| | | 38,082,073 | | *ORMDL3* | 4.886 | 2.98E-06 | 6.40E-04 |
| | | 38,082,346 | | *ORMDL3* | 3.433 | 8.01E-04 | 0.011 |
| | | 38,082,658 | | *ORMDL3* | 4.551 | 1.21E-05 | 0.011 |
| | | 38,108,710 | | | -6.111 | 1.07E-08 | 0.039 |
| | | 38,119,071 | *GSDMA* | | 4.281 | 3.59E-05 | 0.009 |
| | | 38,119,198 | *GSDMA* | | 6.240 | 5.72E-09 | 0.009 |
| | | 38,119,207 | *GSDMA* | | 7.498 | 8.96E-12 | 0.009 |
| | | 38,119,649 | | *GSDMA* | 6.524 | 1.39E-09 | 0.002 |

| | | | | | | | |
|---|---|---|---|---|---|---|---|
| | | 38,119,760 | | *GSDMA* | 4.665 | 7.58E-06 | 0.011 |
| | | 38,120,234 | | *GSDMA* | 3.402 | 8.90E-04 | 0.011 |
| | | 38,128,699 | | *GSDMA* | -4.073 | 8.01E-05 | 0.006 |
| rs72832958 | g.38093881.A>C | 37,911,537 | | | -3.392 | 9.19E-04 | 0.035 |
| | | 37,911,661 | | | -3.420 | 8.38E-04 | 0.035 |
| | | 38,083,874 | | *ORMDL3* | 3.528 | 5.80E-04 | 0.011 |
| | | 38,096,333 | | | -5.589 | 1.28E-07 | 0.033 |
| | | 38,096,571 | | | -10.118 | 4.18E-18 | 0.033 |
| | | 38,096,578 | | | -9.592 | 8.40E-17 | 0.033 |
| | | 38,096,648 | | | -4.297 | 3.37E-05 | 0.033 |
| rs113778191 | g.38094183.T>G | 37,911,537 | | | -3.392 | 9.19E-04 | 0.035 |
| | | 37,911,661 | | | -3.420 | 8.38E-04 | 0.035 |
| | | 38,083,874 | | *ORMDL3* | 3.528 | 5.80E-04 | 0.011 |
| | | 38,096,333 | | | -5.589 | 1.28E-07 | 0.033 |
| | | 38,096,571 | | | -10.118 | 4.18E-18 | 0.033 |
| | | 38,096,578 | | | -9.592 | 8.40E-17 | 0.033 |
| | | 38,096,648 | | | -4.297 | 3.37E-05 | 0.033 |
| rs6503525 | g.38095174.C>G | 37,834,869 | | *PGAP3* | 3.453 | 7.49E-04 | 0.020 |
| | | 37,910,697 | | | 3.491 | 6.58E-04 | 0.035 |
| | | 38,077,277 | | | 4.203 | 4.86E-05 | 0.014 |
| | | 38,077,464 | | *ORMDL3* | 4.071 | 8.07E-05 | 0.014 |
| | | 38,080,829 | | *ORMDL3* | 3.663 | 3.61E-04 | 0.014 |
| | | 38,081,530 | | *ORMDL3* | 3.708 | 3.08E-04 | 2.01E-04 |
| | | 38,081,812 | | *ORMDL3* | 4.038 | 9.16E-05 | 2.01E-04 |
| | | 38,082,022 | | *ORMDL3* | 3.625 | 4.13E-04 | 6.40E-04 |
| | | 38,082,073 | | *ORMDL3* | 4.886 | 2.98E-06 | 6.40E-04 |
| | | 38,082,346 | | *ORMDL3* | 3.433 | 8.01E-04 | 0.011 |
| | | 38,082,658 | | *ORMDL3* | 4.551 | 1.21E-05 | 0.011 |
| | | 38,108,710 | | | -6.111 | 1.07E-08 | 0.039 |
| | | 38,119,071 | *GSDMA* | | 4.281 | 3.59E-05 | 0.009 |
| | | 38,119,198 | *GSDMA* | | 6.240 | 5.72E-09 | 0.009 |
| | | 38,119,207 | *GSDMA* | | 7.498 | 8.96E-12 | 0.009 |
| | | 38,119,649 | | *GSDMA* | 6.524 | 1.39E-09 | 0.002 |
| | | 38,119,760 | | *GSDMA* | 4.665 | 7.58E-06 | 0.011 |
| | | 38,120,234 | | *GSDMA* | 3.402 | 8.90E-04 | 0.011 |
| | | 38,128,699 | | *GSDMA* | -4.073 | 8.01E-05 | 0.006 |
| rs72832962 | g.38096501.C>A | 37,911,537 | | | -3.392 | 9.19E-04 | 0.035 |
| | | 37,911,661 | | | -3.420 | 8.38E-04 | 0.035 |
| | | 38,083,874 | | *ORMDL3* | 3.528 | 5.80E-04 | 0.011 |
| | | 38,096,333 | | | -5.589 | 1.28E-07 | 0.033 |
| | | 38,096,571 | | | -10.118 | 4.18E-18 | 0.033 |

| | | | | | | | |
|---|---|---|---|---|---|---|---|
| | | 38,096,578 | | | -9.592 | 8.40E-17 | 0.033 |
| | | 38,096,648 | | | -4.297 | 3.37E-05 | 0.033 |
| rs75541765 | g.38096881.C>T | 37,911,537 | | | -3.392 | 9.19E-04 | 0.035 |
| | | 37,911,661 | | | -3.420 | 8.38E-04 | 0.035 |
| | | 38,083,874 | | *ORMDL3* | 3.528 | 5.80E-04 | 0.011 |
| | | 38,096,333 | | | -5.589 | 1.28E-07 | 0.033 |
| | | 38,096,571 | | | -10.118 | 4.18E-18 | 0.033 |
| | | 38,096,578 | | | -9.592 | 8.40E-17 | 0.033 |
| | | 38,096,648 | | | -4.297 | 3.37E-05 | 0.033 |
| rs72832964 | g.38097008.C>T | 37,911,537 | | | -3.392 | 9.19E-04 | 0.035 |
| | | 37,911,661 | | | -3.420 | 8.38E-04 | 0.035 |
| | | 38,083,874 | | *ORMDL3* | 3.528 | 5.80E-04 | 0.011 |
| | | 38,096,333 | | | -5.589 | 1.28E-07 | 0.033 |
| | | 38,096,571 | | | -10.118 | 4.18E-18 | 0.033 |
| | | 38,096,578 | | | -9.592 | 8.40E-17 | 0.033 |
| | | 38,096,648 | | | -4.297 | 3.37E-05 | 0.033 |
| rs72832965 | g.38097143.C>T | 37,911,537 | | | -3.392 | 9.19E-04 | 0.035 |
| | | 37,911,661 | | | -3.420 | 8.38E-04 | 0.035 |
| | | 38,083,874 | | *ORMDL3* | 3.528 | 5.80E-04 | 0.011 |
| | | 38,096,333 | | | -5.589 | 1.28E-07 | 0.033 |
| | | 38,096,571 | | | -10.118 | 4.18E-18 | 0.033 |
| | | 38,096,578 | | | -9.592 | 8.40E-17 | 0.033 |
| | | 38,096,648 | | | -4.297 | 3.37E-05 | 0.033 |
| rs7216564 | g.38097172.A>T | 37,834,869 | | *PGAP3* | 3.453 | 7.49E-04 | 0.020 |
| | | 37,910,697 | | | 3.491 | 6.58E-04 | 0.035 |
| | | 38,077,277 | | | 4.203 | 4.86E-05 | 0.014 |
| | | 38,077,464 | | *ORMDL3* | 4.071 | 8.07E-05 | 0.014 |
| | | 38,080,829 | | *ORMDL3* | 3.663 | 3.61E-04 | 0.014 |
| | | 38,081,530 | | *ORMDL3* | 3.708 | 3.08E-04 | 2.01E-04 |
| | | 38,081,812 | | *ORMDL3* | 4.038 | 9.16E-05 | 2.01E-04 |
| | | 38,082,022 | | *ORMDL3* | 3.625 | 4.13E-04 | 6.40E-04 |
| | | 38,082,073 | | *ORMDL3* | 4.886 | 2.98E-06 | 6.40E-04 |
| | | 38,082,346 | | *ORMDL3* | 3.433 | 8.01E-04 | 0.011 |
| | | 38,082,658 | | *ORMDL3* | 4.551 | 1.21E-05 | 0.011 |
| | | 38,108,710 | | | -6.111 | 1.07E-08 | 0.039 |
| | | 38,119,071 | *GSDMA* | | 4.281 | 3.59E-05 | 0.009 |
| | | 38,119,198 | *GSDMA* | | 6.240 | 5.72E-09 | 0.009 |
| | | 38,119,207 | *GSDMA* | | 7.498 | 8.96E-12 | 0.009 |
| | | 38,119,649 | | *GSDMA* | 6.524 | 1.39E-09 | 0.002 |
| | | 38,119,760 | | *GSDMA* | 4.665 | 7.58E-06 | 0.011 |
| | | 38,120,234 | | *GSDMA* | 3.402 | 8.90E-04 | 0.011 |

| | | | | | | | |
|---|---|---|---|---|---|---|---|
| | | 38,128,699 | | *GSDMA* | -4.073 | 8.01E-05 | 0.006 |
| rs79785426 | g.38098027.A>G | 37,911,537 | | | -3.392 | 9.19E-04 | 0.035 |
| | | 37,911,661 | | | -3.420 | 8.38E-04 | 0.035 |
| | | 38,083,874 | | *ORMDL3* | 3.528 | 5.80E-04 | 0.011 |
| | | 38,096,333 | | | -5.589 | 1.28E-07 | 0.033 |
| | | 38,096,571 | | | -10.118 | 4.18E-18 | 0.033 |
| | | 38,096,578 | | | -9.592 | 8.40E-17 | 0.033 |
| | | 38,096,648 | | | -4.297 | 3.37E-05 | 0.033 |
| rs144007425 | g.38098781_38098781ins G | 37,834,869 | | *PGAP3* | 3.453 | 7.49E-04 | 0.020 |
| | | 37,910,697 | | | 3.491 | 6.58E-04 | 0.035 |
| | | 38,077,277 | | | 4.203 | 4.86E-05 | 0.014 |
| | | 38,077,464 | | *ORMDL3* | 4.071 | 8.07E-05 | 0.014 |
| | | 38,080,829 | | *ORMDL3* | 3.663 | 3.61E-04 | 0.014 |
| | | 38,081,530 | | *ORMDL3* | 3.708 | 3.08E-04 | 2.01E-04 |
| | | 38,081,812 | | *ORMDL3* | 4.038 | 9.16E-05 | 2.01E-04 |
| | | 38,082,022 | | *ORMDL3* | 3.625 | 4.13E-04 | 6.40E-04 |
| | | 38,082,073 | | *ORMDL3* | 4.886 | 2.98E-06 | 6.40E-04 |
| | | 38,082,346 | | *ORMDL3* | 3.433 | 8.01E-04 | 0.011 |
| | | 38,082,658 | | *ORMDL3* | 4.551 | 1.21E-05 | 0.011 |
| | | 38,108,710 | | | -6.111 | 1.07E-08 | 0.039 |
| | | 38,119,071 | *GSDMA* | | 4.281 | 3.59E-05 | 0.009 |
| | | 38,119,198 | *GSDMA* | | 6.240 | 5.72E-09 | 0.009 |
| | | 38,119,207 | *GSDMA* | | 7.498 | 8.96E-12 | 0.009 |
| | | 38,119,649 | | *GSDMA* | 6.524 | 1.39E-09 | 0.002 |
| | | 38,119,760 | | *GSDMA* | 4.665 | 7.58E-06 | 0.011 |
| | | 38,120,234 | | *GSDMA* | 3.402 | 8.90E-04 | 0.011 |
| | | 38,128,699 | | *GSDMA* | -4.073 | 8.01E-05 | 0.006 |
| rs8065126 | g.38099035.T>C | 37,887,061 | *MIEN1* | | 3.516 | 6.05E-04 | 0.002 |
| | | 37,894,590 | | | 3.678 | 3.44E-04 | 0.013 |
| | | 37,910,697 | | | 3.727 | 2.88E-04 | 0.035 |
| | | 38,024,331 | | | -3.479 | 6.86E-04 | 0.017 |
| | | 38,077,464 | | *ORMDL3* | 5.568 | 1.41E-07 | 0.014 |
| | | 38,080,829 | | *ORMDL3* | 3.433 | 8.02E-04 | 0.014 |
| | | 38,081,995 | | *ORMDL3* | 3.739 | 2.76E-04 | 2.01E-04 |
| | | 38,082,022 | | *ORMDL3* | 4.789 | 4.49E-06 | 6.40E-04 |
| | | 38,082,035 | | *ORMDL3* | 3.948 | 1.28E-04 | 6.40E-04 |
| | | 38,082,073 | | *ORMDL3* | 4.447 | 1.85E-05 | 6.40E-04 |
| | | 38,082,658 | | *ORMDL3* | 4.810 | 4.12E-06 | 0.011 |
| | | 38,108,710 | | | -7.597 | 5.27E-12 | 0.039 |
| | | 38,119,071 | *GSDMA* | | 4.264 | 3.83E-05 | 0.009 |

| | | | | | | | |
|---|---|---|---|---|---|---|---|
| | | 38,119,198 | *GSDMA* | | 7.078 | 8.16E-11 | 0.009 |
| | | 38,119,207 | *GSDMA* | | 7.857 | 1.30E-12 | 0.009 |
| | | 38,119,649 | | *GSDMA* | 6.009 | 1.76E-08 | 0.002 |
| | | 38,119,760 | | *GSDMA* | 5.350 | 3.84E-07 | 0.011 |
| | | 38,128,699 | | *GSDMA* | -4.346 | 2.77E-05 | 0.006 |
| rs72832966 | g.38100129.A>G | 37,911,537 | | | -3.392 | 9.19E-04 | 0.035 |
| | | 37,911,661 | | | -3.420 | 8.38E-04 | 0.035 |
| | | 38,083,874 | | *ORMDL3* | 3.528 | 5.80E-04 | 0.011 |
| | | 38,096,333 | | | -5.589 | 1.28E-07 | 0.033 |
| | | 38,096,571 | | | -10.118 | 4.18E-18 | 0.033 |
| | | 38,096,578 | | | -9.592 | 8.40E-17 | 0.033 |
| | | 38,096,648 | | | -4.297 | 3.37E-05 | 0.033 |
| rs4795406 | g.38100134.C>G | 37,834,869 | | *PGAP3* | 3.453 | 7.49E-04 | 0.020 |
| | | 37,910,697 | | | 3.491 | 6.58E-04 | 0.035 |
| | | 38,077,277 | | | 4.203 | 4.86E-05 | 0.014 |
| | | 38,077,464 | | *ORMDL3* | 4.071 | 8.07E-05 | 0.014 |
| | | 38,080,829 | | *ORMDL3* | 3.663 | 3.61E-04 | 0.014 |
| | | 38,081,530 | | *ORMDL3* | 3.708 | 3.08E-04 | 2.01E-04 |
| | | 38,081,812 | | *ORMDL3* | 4.038 | 9.16E-05 | 2.01E-04 |
| | | 38,082,022 | | *ORMDL3* | 3.625 | 4.13E-04 | 6.40E-04 |
| | | 38,082,073 | | *ORMDL3* | 4.886 | 2.98E-06 | 6.40E-04 |
| | | 38,082,346 | | *ORMDL3* | 3.433 | 8.01E-04 | 0.011 |
| | | 38,082,658 | | *ORMDL3* | 4.551 | 1.21E-05 | 0.011 |
| | | 38,108,710 | | | -6.111 | 1.07E-08 | 0.039 |
| | | 38,119,071 | *GSDMA* | | 4.281 | 3.59E-05 | 0.009 |
| | | 38,119,198 | *GSDMA* | | 6.240 | 5.72E-09 | 0.009 |
| | | 38,119,207 | *GSDMA* | | 7.498 | 8.96E-12 | 0.009 |
| | | 38,119,649 | | *GSDMA* | 6.524 | 1.39E-09 | 0.002 |
| | | 38,119,760 | | *GSDMA* | 4.665 | 7.58E-06 | 0.011 |
| | | 38,120,234 | | *GSDMA* | 3.402 | 8.90E-04 | 0.011 |
| | | 38,128,699 | | *GSDMA* | -4.073 | 8.01E-05 | 0.006 |
| rs74717022 | g.38100633.T>C (Ser.158.Ser) | 37,911,537 | | | -3.392 | 9.19E-04 | 0.035 |
| | | 37,911,661 | | | -3.420 | 8.38E-04 | 0.035 |
| | | 38,083,874 | | *ORMDL3* | 3.528 | 5.80E-04 | 0.011 |
| | | 38,096,333 | | | -5.589 | 1.28E-07 | 0.033 |
| | | 38,096,571 | | | -10.118 | 4.18E-18 | 0.033 |
| | | 38,096,578 | | | -9.592 | 8.40E-17 | 0.033 |
| | | 38,096,648 | | | -4.297 | 3.37E-05 | 0.033 |
| rs3848395 | g.38101468.T>C | 37,887,061 | *MIEN1* | | 3.516 | 6.05E-04 | 0.002 |
| | | 37,894,590 | | | 3.678 | 3.44E-04 | 0.013 |

| | | | | | | | |
|---|---|---|---|---|---|---|---|
| | | 37,910,697 | | | 3.727 | 2.88E-04 | 0.035 |
| | | 38,024,331 | | | -3.479 | 6.86E-04 | 0.017 |
| | | 38,077,464 | | *ORMDL3* | 5.568 | 1.41E-07 | 0.014 |
| | | 38,080,829 | | *ORMDL3* | 3.433 | 8.02E-04 | 0.014 |
| | | 38,081,995 | | *ORMDL3* | 3.739 | 2.76E-04 | 2.01E-04 |
| | | 38,082,022 | | *ORMDL3* | 4.789 | 4.49E-06 | 6.40E-04 |
| | | 38,082,035 | | *ORMDL3* | 3.948 | 1.28E-04 | 6.40E-04 |
| | | 38,082,073 | | *ORMDL3* | 4.447 | 1.85E-05 | 6.40E-04 |
| | | 38,082,658 | | *ORMDL3* | 4.810 | 4.12E-06 | 0.011 |
| | | 38,108,710 | | | -7.597 | 5.27E-12 | 0.039 |
| | | 38,119,071 | *GSDMA* | | 4.264 | 3.83E-05 | 0.009 |
| | | 38,119,198 | *GSDMA* | | 7.078 | 8.16E-11 | 0.009 |
| | | 38,119,207 | *GSDMA* | | 7.857 | 1.30E-12 | 0.009 |
| | | 38,119,649 | | *GSDMA* | 6.009 | 1.76E-08 | 0.002 |
| | | 38,119,760 | | *GSDMA* | 5.350 | 3.84E-07 | 0.011 |
| | | 38,128,699 | | *GSDMA* | -4.346 | 2.77E-05 | 0.006 |
| rs4065985 | g.38101932.G>C | 37,894,590 | | | 4.022 | 9.72E-05 | 0.013 |
| | | 38,077,051 | | | 3.381 | 9.53E-04 | 0.034 |
| | | 38,077,464 | | *ORMDL3* | 5.159 | 9.02E-07 | 0.014 |
| | | 38,081,812 | | *ORMDL3* | 3.876 | 1.68E-04 | 2.01E-04 |
| | | 38,081,829 | | *ORMDL3* | 4.103 | 7.15E-05 | 2.01E-04 |
| | | 38,081,995 | | *ORMDL3* | 3.904 | 1.51E-04 | 2.01E-04 |
| | | 38,082,022 | | *ORMDL3* | 4.713 | 6.20E-06 | 6.40E-04 |
| | | 38,082,035 | | *ORMDL3* | 4.030 | 9.43E-05 | 6.40E-04 |
| | | 38,082,073 | | *ORMDL3* | 4.244 | 4.14E-05 | 6.40E-04 |
| | | 38,082,658 | | *ORMDL3* | 5.266 | 5.61E-07 | 0.011 |
| | | 38,108,710 | | | -6.712 | 5.38E-10 | 0.039 |
| | | 38,119,071 | *GSDMA* | | 5.210 | 7.21E-07 | 0.009 |
| | | 38,119,132 | *GSDMA* | | 3.479 | 6.86E-04 | 0.009 |
| | | 38,119,198 | *GSDMA* | | 7.500 | 8.85E-12 | 0.009 |
| | | 38,119,207 | *GSDMA* | | 8.154 | 2.59E-13 | 0.009 |
| | | 38,119,649 | | *GSDMA* | 6.482 | 1.72E-09 | 0.002 |
| | | 38,119,760 | | *GSDMA* | 4.222 | 4.52E-05 | 0.011 |
| | | 38,120,234 | | *GSDMA* | 3.485 | 6.72E-04 | 0.011 |
| | | 38,120,370 | | *GSDMA* | 3.846 | 1.87E-04 | 0.011 |
| | | 38,128,699 | | *GSDMA* | -4.253 | 4.00E-05 | 0.006 |
| rs4065986 | g.38102641.A>G | 37,834,869 | | *PGAP3* | 3.453 | 7.49E-04 | 0.020 |
| | | 37,910,697 | | | 3.491 | 6.58E-04 | 0.035 |
| | | 38,077,277 | | | 4.203 | 4.86E-05 | 0.014 |
| | | 38,077,464 | | *ORMDL3* | 4.071 | 8.07E-05 | 0.014 |
| | | 38,080,829 | | *ORMDL3* | 3.663 | 3.61E-04 | 0.014 |

| | | | | | | | |
|---|---|---|---|---|---|---|---|
| | | 38,081,530 | | *ORMDL3* | 3.708 | 3.08E-04 | 2.01E-04 |
| | | 38,081,812 | | *ORMDL3* | 4.038 | 9.16E-05 | 2.01E-04 |
| | | 38,082,022 | | *ORMDL3* | 3.625 | 4.13E-04 | 6.40E-04 |
| | | 38,082,073 | | *ORMDL3* | 4.886 | 2.98E-06 | 6.40E-04 |
| | | 38,082,346 | | *ORMDL3* | 3.433 | 8.01E-04 | 0.011 |
| | | 38,082,658 | | *ORMDL3* | 4.551 | 1.21E-05 | 0.011 |
| | | 38,108,710 | | | -6.111 | 1.07E-08 | 0.039 |
| | | 38,119,071 | *GSDMA* | | 4.281 | 3.59E-05 | 0.009 |
| | | 38,119,198 | *GSDMA* | | 6.240 | 5.72E-09 | 0.009 |
| | | 38,119,207 | *GSDMA* | | 7.498 | 8.96E-12 | 0.009 |
| | | 38,119,649 | | *GSDMA* | 6.524 | 1.39E-09 | 0.002 |
| | | 38,119,760 | | *GSDMA* | 4.665 | 7.58E-06 | 0.011 |
| | | 38,120,234 | | *GSDMA* | 3.402 | 8.90E-04 | 0.011 |
| | | 38,128,699 | | *GSDMA* | -4.073 | 8.01E-05 | 0.006 |
| rs3893044 | g.38103016.A>G | 37,894,590 | | | 4.022 | 9.72E-05 | 0.013 |
| | | 38,077,051 | | | 3.381 | 9.53E-04 | 0.034 |
| | | 38,077,464 | | *ORMDL3* | 5.159 | 9.02E-07 | 0.014 |
| | | 38,081,812 | | *ORMDL3* | 3.876 | 1.68E-04 | 2.01E-04 |
| | | 38,081,829 | | *ORMDL3* | 4.103 | 7.15E-05 | 2.01E-04 |
| | | 38,081,995 | | *ORMDL3* | 3.904 | 1.51E-04 | 2.01E-04 |
| | | 38,082,022 | | *ORMDL3* | 4.713 | 6.20E-06 | 6.40E-04 |
| | | 38,082,035 | | *ORMDL3* | 4.030 | 9.43E-05 | 6.40E-04 |
| | | 38,082,073 | | *ORMDL3* | 4.244 | 4.14E-05 | 6.40E-04 |
| | | 38,082,658 | | *ORMDL3* | 5.266 | 5.61E-07 | 0.011 |
| | | 38,108,710 | | | -6.712 | 5.38E-10 | 0.039 |
| | | 38,119,071 | *GSDMA* | | 5.210 | 7.21E-07 | 0.009 |
| | | 38,119,132 | *GSDMA* | | 3.479 | 6.86E-04 | 0.009 |
| | | 38,119,198 | *GSDMA* | | 7.500 | 8.85E-12 | 0.009 |
| | | 38,119,207 | *GSDMA* | | 8.154 | 2.59E-13 | 0.009 |
| | | 38,119,649 | | *GSDMA* | 6.482 | 1.72E-09 | 0.002 |
| | | 38,119,760 | | *GSDMA* | 4.222 | 4.52E-05 | 0.011 |
| | | 38,120,234 | | *GSDMA* | 3.485 | 6.72E-04 | 0.011 |
| | | 38,120,370 | | *GSDMA* | 3.846 | 1.87E-04 | 0.011 |
| | | 38,128,699 | | *GSDMA* | -4.253 | 4.00E-05 | 0.006 |
| rs62068170 | g.38103210.G>A | 37,894,590 | | | 4.022 | 9.72E-05 | 0.013 |
| | | 38,077,051 | | | 3.381 | 9.53E-04 | 0.034 |
| | | 38,077,464 | | *ORMDL3* | 5.159 | 9.02E-07 | 0.014 |
| | | 38,081,812 | | *ORMDL3* | 3.876 | 1.68E-04 | 2.01E-04 |
| | | 38,081,829 | | *ORMDL3* | 4.103 | 7.15E-05 | 2.01E-04 |
| | | 38,081,995 | | *ORMDL3* | 3.904 | 1.51E-04 | 2.01E-04 |
| | | 38,082,022 | | *ORMDL3* | 4.713 | 6.20E-06 | 6.40E-04 |

| | | | | | | | |
|---|---|---|---|---|---|---|---|
| | | 38,082,035 | | *ORMDL3* | 4.030 | 9.43E-05 | 6.40E-04 |
| | | 38,082,073 | | *ORMDL3* | 4.244 | 4.14E-05 | 6.40E-04 |
| | | 38,082,658 | | *ORMDL3* | 5.266 | 5.61E-07 | 0.011 |
| | | 38,108,710 | | | -6.712 | 5.38E-10 | 0.039 |
| | | 38,119,071 | *GSDMA* | | 5.210 | 7.21E-07 | 0.009 |
| | | 38,119,132 | *GSDMA* | | 3.479 | 6.86E-04 | 0.009 |
| | | 38,119,198 | *GSDMA* | | 7.500 | 8.85E-12 | 0.009 |
| | | 38,119,207 | *GSDMA* | | 8.154 | 2.59E-13 | 0.009 |
| | | 38,119,649 | | *GSDMA* | 6.482 | 1.72E-09 | 0.002 |
| | | 38,119,760 | | *GSDMA* | 4.222 | 4.52E-05 | 0.011 |
| | | 38,120,234 | | *GSDMA* | 3.485 | 6.72E-04 | 0.011 |
| | | 38,120,370 | | *GSDMA* | 3.846 | 1.87E-04 | 0.011 |
| | | 38,128,699 | | *GSDMA* | -4.253 | 4.00E-05 | 0.006 |
| rs62068171 | g.38103242.G>A | 37,894,590 | | | 4.022 | 9.72E-05 | 0.013 |
| | | 38,077,051 | | | 3.381 | 9.53E-04 | 0.034 |
| | | 38,077,464 | | *ORMDL3* | 5.159 | 9.02E-07 | 0.014 |
| | | 38,081,812 | | *ORMDL3* | 3.876 | 1.68E-04 | 2.01E-04 |
| | | 38,081,829 | | *ORMDL3* | 4.103 | 7.15E-05 | 2.01E-04 |
| | | 38,081,995 | | *ORMDL3* | 3.904 | 1.51E-04 | 2.01E-04 |
| | | 38,082,022 | | *ORMDL3* | 4.713 | 6.20E-06 | 6.40E-04 |
| | | 38,082,035 | | *ORMDL3* | 4.030 | 9.43E-05 | 6.40E-04 |
| | | 38,082,073 | | *ORMDL3* | 4.244 | 4.14E-05 | 6.40E-04 |
| | | 38,082,658 | | *ORMDL3* | 5.266 | 5.61E-07 | 0.011 |
| | | 38,108,710 | | | -6.712 | 5.38E-10 | 0.039 |
| | | 38,119,071 | *GSDMA* | | 5.210 | 7.21E-07 | 0.009 |
| | | 38,119,132 | *GSDMA* | | 3.479 | 6.86E-04 | 0.009 |
| | | 38,119,198 | *GSDMA* | | 7.500 | 8.85E-12 | 0.009 |
| | | 38,119,207 | *GSDMA* | | 8.154 | 2.59E-13 | 0.009 |
| | | 38,119,649 | | *GSDMA* | 6.482 | 1.72E-09 | 0.002 |
| | | 38,119,760 | | *GSDMA* | 4.222 | 4.52E-05 | 0.011 |
| | | 38,120,234 | | *GSDMA* | 3.485 | 6.72E-04 | 0.011 |
| | | 38,120,370 | | *GSDMA* | 3.846 | 1.87E-04 | 0.011 |
| | | 38,128,699 | | *GSDMA* | -4.253 | 4.00E-05 | 0.006 |
| rs8080734 | g.38103285.G>A | 37,894,590 | | | 3.997 | 1.07E-04 | 0.013 |
| | | 38,077,464 | | *ORMDL3* | 5.117 | 1.09E-06 | 0.014 |
| | | 38,081,812 | | *ORMDL3* | 3.921 | 1.42E-04 | 2.01E-04 |
| | | 38,081,829 | | *ORMDL3* | 4.003 | 1.05E-04 | 2.01E-04 |
| | | 38,081,995 | | *ORMDL3* | 3.725 | 2.90E-04 | 2.01E-04 |
| | | 38,082,022 | | *ORMDL3* | 4.569 | 1.12E-05 | 6.40E-04 |
| | | 38,082,035 | | *ORMDL3* | 3.850 | 1.84E-04 | 6.40E-04 |
| | | 38,082,073 | | *ORMDL3* | 4.326 | 3.00E-05 | 6.40E-04 |

| | | | | | | | |
|---|---|---|---|---|---|---|---|
| | | 38,082,658 | | *ORMDL3* | 5.208 | 7.28E-07 | 0.011 |
| | | 38,108,710 | | | -6.537 | 1.30E-09 | 0.039 |
| | | 38,119,071 | *GSDMA* | | 5.226 | 6.70E-07 | 0.009 |
| | | 38,119,132 | *GSDMA* | | 3.484 | 6.74E-04 | 0.009 |
| | | 38,119,198 | *GSDMA* | | 7.483 | 9.67E-12 | 0.009 |
| | | 38,119,207 | *GSDMA* | | 8.337 | 9.47E-14 | 0.009 |
| | | 38,119,649 | | *GSDMA* | 6.373 | 2.96E-09 | 0.002 |
| | | 38,119,760 | | *GSDMA* | 4.080 | 7.81E-05 | 0.011 |
| | | 38,120,234 | | *GSDMA* | 3.480 | 6.82E-04 | 0.011 |
| | | 38,120,370 | | *GSDMA* | 3.868 | 1.73E-04 | 0.011 |
| | | 38,128,699 | | *GSDMA* | -4.265 | 3.82E-05 | 0.006 |
| rs8071050 | g.38106599.A>G | 37,910,697 | | | 3.818 | 2.08E-04 | 0.035 |
| | | 38,049,272 | | | 3.434 | 7.99E-04 | 0.046 |
| | | 38,077,277 | | | 3.521 | 5.93E-04 | 0.014 |
| | | 38,077,464 | | *ORMDL3* | 4.150 | 5.97E-05 | 0.014 |
| | | 38,080,829 | | *ORMDL3* | 4.211 | 4.71E-05 | 0.014 |
| | | 38,082,022 | | *ORMDL3* | 3.675 | 3.46E-04 | 6.40E-04 |
| | | 38,082,073 | | *ORMDL3* | 4.960 | 2.17E-06 | 6.40E-04 |
| | | 38,082,346 | | *ORMDL3* | 3.441 | 7.81E-04 | 0.011 |
| | | 38,082,658 | | *ORMDL3* | 4.101 | 7.20E-05 | 0.011 |
| | | 38,108,710 | | | -6.715 | 5.30E-10 | 0.039 |
| | | 38,119,071 | *GSDMA* | | 3.416 | 8.50E-04 | 0.009 |
| | | 38,119,198 | *GSDMA* | | 5.729 | 6.68E-08 | 0.009 |
| | | 38,119,207 | *GSDMA* | | 6.977 | 1.38E-10 | 0.009 |
| | | 38,119,649 | | *GSDMA* | 6.069 | 1.31E-08 | 0.002 |
| | | 38,119,760 | | *GSDMA* | 5.659 | 9.25E-08 | 0.011 |
| | | 38,128,699 | | *GSDMA* | -3.994 | 1.08E-04 | 0.006 |
| rs4795408 | g.38107627.A>G | 37,834,869 | | *PGAP3* | 3.453 | 7.49E-04 | 0.020 |
| | | 37,910,697 | | | 3.491 | 6.58E-04 | 0.035 |
| | | 38,077,277 | | | 4.203 | 4.86E-05 | 0.014 |
| | | 38,077,464 | | *ORMDL3* | 4.071 | 8.07E-05 | 0.014 |
| | | 38,080,829 | | *ORMDL3* | 3.663 | 3.61E-04 | 0.014 |
| | | 38,081,530 | | *ORMDL3* | 3.708 | 3.08E-04 | 2.01E-04 |
| | | 38,081,812 | | *ORMDL3* | 4.038 | 9.16E-05 | 2.01E-04 |
| | | 38,082,022 | | *ORMDL3* | 3.625 | 4.13E-04 | 6.40E-04 |
| | | 38,082,073 | | *ORMDL3* | 4.886 | 2.98E-06 | 6.40E-04 |
| | | 38,082,346 | | *ORMDL3* | 3.433 | 8.01E-04 | 0.011 |
| | | 38,082,658 | | *ORMDL3* | 4.551 | 1.21E-05 | 0.011 |
| | | 38,108,710 | | | -6.111 | 1.07E-08 | 0.039 |
| | | 38,119,071 | *GSDMA* | | 4.281 | 3.59E-05 | 0.009 |
| | | 38,119,198 | *GSDMA* | | 6.240 | 5.72E-09 | 0.009 |

| | | | | | | | |
|---|---|---|---|---|---|---|---|
| | | 38,119,207 | *GSDMA* | | 7.498 | 8.96E-12 | 0.009 |
| | | 38,119,649 | | *GSDMA* | 6.524 | 1.39E-09 | 0.002 |
| | | 38,119,760 | | *GSDMA* | 4.665 | 7.58E-06 | 0.011 |
| | | 38,120,234 | | *GSDMA* | 3.402 | 8.90E-04 | 0.011 |
| | | 38,128,699 | | *GSDMA* | -4.073 | 8.01E-05 | 0.006 |
| rs56946324 | g.38107966.A>C | 37,910,144 | | | 3.390 | 9.25E-04 | 0.043 |
| | | 37,911,661 | | | -3.474 | 6.98E-04 | 0.035 |
| | | 38,024,634 | | | 3.677 | 3.44E-04 | 0.046 |
| | | 38,083,874 | | *ORMDL3* | 4.167 | 5.60E-05 | 0.011 |
| | | 38,096,333 | | | -4.384 | 2.38E-05 | 0.033 |
| | | 38,096,571 | | | -6.389 | 2.73E-09 | 0.033 |
| | | 38,096,578 | | | -6.182 | 7.57E-09 | 0.033 |
| rs10589831 | g.38108023_38108025del | 37,887,061 | *MIEN1* | | 3.573 | 4.96E-04 | 0.002 |
| | | 37,894,590 | | | 3.763 | 2.53E-04 | 0.013 |
| | | 37,910,697 | | | 3.671 | 3.51E-04 | 0.035 |
| | | 38,024,331 | | | -3.468 | 7.12E-04 | 0.017 |
| | | 38,077,464 | | *ORMDL3* | 5.497 | 1.97E-07 | 0.014 |
| | | 38,080,829 | | *ORMDL3* | 3.499 | 6.41E-04 | 0.014 |
| | | 38,081,995 | | *ORMDL3* | 3.928 | 1.38E-04 | 2.01E-04 |
| | | 38,082,022 | | *ORMDL3* | 4.951 | 2.25E-06 | 6.40E-04 |
| | | 38,082,035 | | *ORMDL3* | 4.180 | 5.31E-05 | 6.40E-04 |
| | | 38,082,073 | | *ORMDL3* | 4.405 | 2.19E-05 | 6.40E-04 |
| | | 38,082,658 | | *ORMDL3* | 4.901 | 2.78E-06 | 0.011 |
| | | 38,108,710 | | | -7.708 | 2.90E-12 | 0.039 |
| | | 38,119,071 | *GSDMA* | | 4.319 | 3.08E-05 | 0.009 |
| | | 38,119,198 | *GSDMA* | | 7.121 | 6.53E-11 | 0.009 |
| | | 38,119,207 | *GSDMA* | | 7.809 | 1.68E-12 | 0.009 |
| | | 38,119,649 | | *GSDMA* | 6.184 | 7.51E-09 | 0.002 |
| | | 38,119,760 | | *GSDMA* | 5.425 | 2.74E-07 | 0.011 |
| | | 38,120,234 | | *GSDMA* | 3.383 | 9.48E-04 | 0.011 |
| | | 38,128,699 | | *GSDMA* | -4.328 | 2.98E-05 | 0.006 |
| rs9889716 | g.38108298.A>G | 37,894,590 | | | 3.809 | 2.14E-04 | 0.013 |
| | | 37,910,697 | | | 3.684 | 3.36E-04 | 0.035 |
| | | 38,024,331 | | | -3.654 | 3.74E-04 | 0.017 |
| | | 38,077,464 | | *ORMDL3* | 5.537 | 1.63E-07 | 0.014 |
| | | 38,081,995 | | *ORMDL3* | 3.806 | 2.16E-04 | 2.01E-04 |
| | | 38,082,022 | | *ORMDL3* | 4.689 | 6.84E-06 | 6.40E-04 |
| | | 38,082,035 | | *ORMDL3* | 3.679 | 3.41E-04 | 6.40E-04 |
| | | 38,082,073 | | *ORMDL3* | 4.435 | 1.94E-05 | 6.40E-04 |
| | | 38,082,658 | | *ORMDL3* | 4.593 | 1.02E-05 | 0.011 |

| | | | | | | | |
|---|---|---|---|---|---|---|---|
| | | 38,108,710 | | | -7.896 | 1.06E-12 | 0.039 |
| | | 38,119,071 | *GSDMA* | | 4.118 | 6.74E-05 | 0.009 |
| | | 38,119,198 | *GSDMA* | | 6.731 | 4.88E-10 | 0.009 |
| | | 38,119,207 | *GSDMA* | | 7.631 | 4.40E-12 | 0.009 |
| | | 38,119,649 | | *GSDMA* | 6.143 | 9.16E-09 | 0.002 |
| | | 38,119,760 | | *GSDMA* | 5.349 | 3.86E-07 | 0.011 |
| | | 38,128,699 | | *GSDMA* | -4.519 | 1.38E-05 | 0.006 |
| rs9895948 | g.38108363.T>C | 37,887,061 | *MIEN1* | | 3.573 | 4.96E-04 | 0.002 |
| | | 37,894,590 | | | 3.763 | 2.53E-04 | 0.013 |
| | | 37,910,697 | | | 3.671 | 3.51E-04 | 0.035 |
| | | 38,024,331 | | | -3.468 | 7.12E-04 | 0.017 |
| | | 38,077,464 | | *ORMDL3* | 5.497 | 1.97E-07 | 0.014 |
| | | 38,080,829 | | *ORMDL3* | 3.499 | 6.41E-04 | 0.014 |
| | | 38,081,995 | | *ORMDL3* | 3.928 | 1.38E-04 | 2.01E-04 |
| | | 38,082,022 | | *ORMDL3* | 4.951 | 2.25E-06 | 6.40E-04 |
| | | 38,082,035 | | *ORMDL3* | 4.180 | 5.31E-05 | 6.40E-04 |
| | | 38,082,073 | | *ORMDL3* | 4.405 | 2.19E-05 | 6.40E-04 |
| | | 38,082,658 | | *ORMDL3* | 4.901 | 2.78E-06 | 0.011 |
| | | 38,108,710 | | | -7.708 | 2.90E-12 | 0.039 |
| | | 38,119,071 | *GSDMA* | | 4.319 | 3.08E-05 | 0.009 |
| | | 38,119,198 | *GSDMA* | | 7.121 | 6.53E-11 | 0.009 |
| | | 38,119,207 | *GSDMA* | | 7.809 | 1.68E-12 | 0.009 |
| | | 38,119,649 | | *GSDMA* | 6.184 | 7.51E-09 | 0.002 |
| | | 38,119,760 | | *GSDMA* | 5.425 | 2.74E-07 | 0.011 |
| | | 38,120,234 | | *GSDMA* | 3.383 | 9.48E-04 | 0.011 |
| | | 38,128,699 | | *GSDMA* | -4.328 | 2.98E-05 | 0.006 |
| rs7223318 | g.38108553.G>C | 37,887,061 | *MIEN1* | | 3.573 | 4.96E-04 | 0.002 |
| | | 37,894,590 | | | 3.763 | 2.53E-04 | 0.013 |
| | | 37,910,697 | | | 3.671 | 3.51E-04 | 0.035 |
| | | 38,024,331 | | | -3.468 | 7.12E-04 | 0.017 |
| | | 38,077,464 | | *ORMDL3* | 5.497 | 1.97E-07 | 0.014 |
| | | 38,080,829 | | *ORMDL3* | 3.499 | 6.41E-04 | 0.014 |
| | | 38,081,995 | | *ORMDL3* | 3.928 | 1.38E-04 | 2.01E-04 |
| | | 38,082,022 | | *ORMDL3* | 4.951 | 2.25E-06 | 6.40E-04 |
| | | 38,082,035 | | *ORMDL3* | 4.180 | 5.31E-05 | 6.40E-04 |
| | | 38,082,073 | | *ORMDL3* | 4.405 | 2.19E-05 | 6.40E-04 |
| | | 38,082,658 | | *ORMDL3* | 4.901 | 2.78E-06 | 0.011 |
| | | 38,108,710 | | | -7.708 | 2.90E-12 | 0.039 |
| | | 38,119,071 | *GSDMA* | | 4.319 | 3.08E-05 | 0.009 |
| | | 38,119,198 | *GSDMA* | | 7.121 | 6.53E-11 | 0.009 |
| | | 38,119,207 | *GSDMA* | | 7.809 | 1.68E-12 | 0.009 |

| | | | | | | | |
|---|---|---|---|---|---|---|---|
| | | 38,119,649 | | *GSDMA* | 6.184 | 7.51E-09 | 0.002 |
| | | 38,119,760 | | *GSDMA* | 5.425 | 2.74E-07 | 0.011 |
| | | 38,120,234 | | *GSDMA* | 3.383 | 9.48E-04 | 0.011 |
| | | 38,128,699 | | *GSDMA* | -4.328 | 2.98E-05 | 0.006 |
| rs72832971 | g.38108619.T>C | 37,911,537 | | | -3.387 | 9.34E-04 | 0.035 |
| | | 37,911,661 | | | -3.423 | 8.28E-04 | 0.035 |
| | | 38,083,874 | | *ORMDL3* | 3.527 | 5.81E-04 | 0.011 |
| | | 38,096,333 | | | -5.590 | 1.28E-07 | 0.033 |
| | | 38,096,571 | | | -10.113 | 4.31E-18 | 0.033 |
| | | 38,096,578 | | | -9.581 | 8.92E-17 | 0.033 |
| | | 38,096,648 | | | -4.294 | 3.40E-05 | 0.033 |
| rs7209742 | g.38108708.A>G | 37,887,061 | *MIEN1* | | 3.573 | 4.96E-04 | 0.002 |
| | | 37,894,590 | | | 3.763 | 2.53E-04 | 0.013 |
| | | 37,910,697 | | | 3.671 | 3.51E-04 | 0.035 |
| | | 38,024,331 | | | -3.468 | 7.12E-04 | 0.017 |
| | | 38,077,464 | | *ORMDL3* | 5.497 | 1.97E-07 | 0.014 |
| | | 38,080,829 | | *ORMDL3* | 3.499 | 6.41E-04 | 0.014 |
| | | 38,081,995 | | *ORMDL3* | 3.928 | 1.38E-04 | 2.01E-04 |
| | | 38,082,022 | | *ORMDL3* | 4.951 | 2.25E-06 | 6.40E-04 |
| | | 38,082,035 | | *ORMDL3* | 4.180 | 5.31E-05 | 6.40E-04 |
| | | 38,082,073 | | *ORMDL3* | 4.405 | 2.19E-05 | 6.40E-04 |
| | | 38,082,658 | | *ORMDL3* | 4.901 | 2.78E-06 | 0.011 |
| | | 38,108,710 | | | -7.708 | 2.90E-12 | 0.039 |
| | | 38,119,071 | *GSDMA* | | 4.319 | 3.08E-05 | 0.009 |
| | | 38,119,198 | *GSDMA* | | 7.121 | 6.53E-11 | 0.009 |
| | | 38,119,207 | *GSDMA* | | 7.809 | 1.68E-12 | 0.009 |
| | | 38,119,649 | | *GSDMA* | 6.184 | 7.51E-09 | 0.002 |
| | | 38,119,760 | | *GSDMA* | 5.425 | 2.74E-07 | 0.011 |
| | | 38,120,234 | | *GSDMA* | 3.383 | 9.48E-04 | 0.011 |
| | | 38,128,699 | | *GSDMA* | -4.328 | 2.98E-05 | 0.006 |
| rs28618095 | g.38109075.T>C | 37,887,061 | *MIEN1* | | 3.463 | 7.24E-04 | 0.002 |
| | | 37,894,590 | | | 3.476 | 6.92E-04 | 0.013 |
| | | 37,910,697 | | | 3.595 | 4.59E-04 | 0.035 |
| | | 38,024,331 | | | -4.001 | 1.05E-04 | 0.017 |
| | | 38,077,464 | | *ORMDL3* | 5.453 | 2.41E-07 | 0.014 |
| | | 38,081,995 | | *ORMDL3* | 4.113 | 6.87E-05 | 2.01E-04 |
| | | 38,082,022 | | *ORMDL3* | 5.053 | 1.44E-06 | 6.40E-04 |
| | | 38,082,035 | | *ORMDL3* | 4.209 | 4.76E-05 | 6.40E-04 |
| | | 38,082,073 | | *ORMDL3* | 4.426 | 2.01E-05 | 6.40E-04 |
| | | 38,082,658 | | *ORMDL3* | 4.991 | 1.89E-06 | 0.011 |
| | | 38,108,710 | | | -7.692 | 3.16E-12 | 0.039 |

| | | | | | | | |
|---|---|---|---|---|---|---|---|
| | | 38,119,071 | *GSDMA* | | 4.508 | 1.44E-05 | 0.009 |
| | | 38,119,198 | *GSDMA* | | 7.160 | 5.32E-11 | 0.009 |
| | | 38,119,207 | *GSDMA* | | 8.027 | 5.17E-13 | 0.009 |
| | | 38,119,649 | | *GSDMA* | 6.304 | 4.17E-09 | 0.002 |
| | | 38,119,760 | | *GSDMA* | 5.429 | 2.69E-07 | 0.011 |
| | | 38,120,234 | | *GSDMA* | 3.499 | 6.39E-04 | 0.011 |
| | | 38,128,699 | | *GSDMA* | -4.444 | 1.87E-05 | 0.006 |
| rs56301252 | g.38109155.T>G | 37,887,061 | *MIEN1* | | 3.573 | 4.96E-04 | 0.002 |
| | | 37,894,590 | | | 3.763 | 2.53E-04 | 0.013 |
| | | 37,910,697 | | | 3.671 | 3.51E-04 | 0.035 |
| | | 38,024,331 | | | -3.468 | 7.12E-04 | 0.017 |
| | | 38,077,464 | | *ORMDL3* | 5.497 | 1.97E-07 | 0.014 |
| | | 38,080,829 | | *ORMDL3* | 3.499 | 6.41E-04 | 0.014 |
| | | 38,081,995 | | *ORMDL3* | 3.928 | 1.38E-04 | 2.01E-04 |
| | | 38,082,022 | | *ORMDL3* | 4.951 | 2.25E-06 | 6.40E-04 |
| | | 38,082,035 | | *ORMDL3* | 4.180 | 5.31E-05 | 6.40E-04 |
| | | 38,082,073 | | *ORMDL3* | 4.405 | 2.19E-05 | 6.40E-04 |
| | | 38,082,658 | | *ORMDL3* | 4.901 | 2.78E-06 | 0.011 |
| | | 38,108,710 | | | -7.708 | 2.90E-12 | 0.039 |
| | | 38,119,071 | *GSDMA* | | 4.319 | 3.08E-05 | 0.009 |
| | | 38,119,198 | *GSDMA* | | 7.121 | 6.53E-11 | 0.009 |
| | | 38,119,207 | *GSDMA* | | 7.809 | 1.68E-12 | 0.009 |
| | | 38,119,649 | | *GSDMA* | 6.184 | 7.51E-09 | 0.002 |
| | | 38,119,760 | | *GSDMA* | 5.425 | 2.74E-07 | 0.011 |
| | | 38,120,234 | | *GSDMA* | 3.383 | 9.48E-04 | 0.011 |
| | | 38,128,699 | | *GSDMA* | -4.328 | 2.98E-05 | 0.006 |
| rs55927420 | g.38109251.G>T | 37,887,061 | *MIEN1* | | 3.573 | 4.96E-04 | 0.002 |
| | | 37,894,590 | | | 3.763 | 2.53E-04 | 0.013 |
| | | 37,910,697 | | | 3.671 | 3.51E-04 | 0.035 |
| | | 38,024,331 | | | -3.468 | 7.12E-04 | 0.017 |
| | | 38,077,464 | | *ORMDL3* | 5.497 | 1.97E-07 | 0.014 |
| | | 38,080,829 | | *ORMDL3* | 3.499 | 6.41E-04 | 0.014 |
| | | 38,081,995 | | *ORMDL3* | 3.928 | 1.38E-04 | 2.01E-04 |
| | | 38,082,022 | | *ORMDL3* | 4.951 | 2.25E-06 | 6.40E-04 |
| | | 38,082,035 | | *ORMDL3* | 4.180 | 5.31E-05 | 6.40E-04 |
| | | 38,082,073 | | *ORMDL3* | 4.405 | 2.19E-05 | 6.40E-04 |
| | | 38,082,658 | | *ORMDL3* | 4.901 | 2.78E-06 | 0.011 |
| | | 38,108,710 | | | -7.708 | 2.90E-12 | 0.039 |
| | | 38,119,071 | *GSDMA* | | 4.319 | 3.08E-05 | 0.009 |
| | | 38,119,198 | *GSDMA* | | 7.121 | 6.53E-11 | 0.009 |
| | | 38,119,207 | *GSDMA* | | 7.809 | 1.68E-12 | 0.009 |

| | | | | | | | |
|---|---|---|---|---|---|---|---|
| | | 38,119,649 | | *GSDMA* | 6.184 | 7.51E-09 | 0.002 |
| | | 38,119,760 | | *GSDMA* | 5.425 | 2.74E-07 | 0.011 |
| | | 38,120,234 | | *GSDMA* | 3.383 | 9.48E-04 | 0.011 |
| | | 38,128,699 | | *GSDMA* | -4.328 | 2.98E-05 | 0.006 |
| rs56410675 | g.38109254.A>G | 37,887,061 | *MIEN1* | | 3.573 | 4.96E-04 | 0.002 |
| | | 37,894,590 | | | 3.763 | 2.53E-04 | 0.013 |
| | | 37,910,697 | | | 3.671 | 3.51E-04 | 0.035 |
| | | 38,024,331 | | | -3.468 | 7.12E-04 | 0.017 |
| | | 38,077,464 | | *ORMDL3* | 5.497 | 1.97E-07 | 0.014 |
| | | 38,080,829 | | *ORMDL3* | 3.499 | 6.41E-04 | 0.014 |
| | | 38,081,995 | | *ORMDL3* | 3.928 | 1.38E-04 | 2.01E-04 |
| | | 38,082,022 | | *ORMDL3* | 4.951 | 2.25E-06 | 6.40E-04 |
| | | 38,082,035 | | *ORMDL3* | 4.180 | 5.31E-05 | 6.40E-04 |
| | | 38,082,073 | | *ORMDL3* | 4.405 | 2.19E-05 | 6.40E-04 |
| | | 38,082,658 | | *ORMDL3* | 4.901 | 2.78E-06 | 0.011 |
| | | 38,108,710 | | | -7.708 | 2.90E-12 | 0.039 |
| | | 38,119,071 | *GSDMA* | | 4.319 | 3.08E-05 | 0.009 |
| | | 38,119,198 | *GSDMA* | | 7.121 | 6.53E-11 | 0.009 |
| | | 38,119,207 | *GSDMA* | | 7.809 | 1.68E-12 | 0.009 |
| | | 38,119,649 | | *GSDMA* | 6.184 | 7.51E-09 | 0.002 |
| | | 38,119,760 | | *GSDMA* | 5.425 | 2.74E-07 | 0.011 |
| | | 38,120,234 | | *GSDMA* | 3.383 | 9.48E-04 | 0.011 |
| | | 38,128,699 | | *GSDMA* | -4.328 | 2.98E-05 | 0.006 |
| rs76137456 | g.38109590.G>C | 37,879,672 | | *ERBB2* | 3.542 | 5.52E-04 | 0.002 |
| | | 37,911,661 | | | -3.439 | 7.86E-04 | 0.035 |
| | | 38,083,874 | | *ORMDL3* | 3.391 | 9.23E-04 | 0.011 |
| | | 38,096,333 | | | -6.049 | 1.45E-08 | 0.033 |
| | | 38,096,571 | | | -10.722 | 1.32E-19 | 0.033 |
| | | 38,096,578 | | | -10.218 | 2.36E-18 | 0.033 |
| | | 38,096,648 | | | -4.123 | 6.61E-05 | 0.033 |
| rs60667221 | g.38110390.T>A | 37,894,590 | | | 3.766 | 2.50E-04 | 0.013 |
| | | 37,910,697 | | | 3.575 | 4.92E-04 | 0.035 |
| | | 38,077,464 | | *ORMDL3* | 5.290 | 5.04E-07 | 0.014 |
| | | 38,080,829 | | *ORMDL3* | 3.407 | 8.75E-04 | 0.014 |
| | | 38,081,995 | | *ORMDL3* | 3.637 | 3.97E-04 | 2.01E-04 |
| | | 38,082,022 | | *ORMDL3* | 4.853 | 3.43E-06 | 6.40E-04 |
| | | 38,082,035 | | *ORMDL3* | 3.788 | 2.31E-04 | 6.40E-04 |
| | | 38,082,073 | | *ORMDL3* | 4.745 | 5.41E-06 | 6.40E-04 |
| | | 38,082,658 | | *ORMDL3* | 4.634 | 8.61E-06 | 0.011 |
| | | 38,108,710 | | | -8.226 | 1.74E-13 | 0.039 |
| | | 38,119,071 | *GSDMA* | | 4.340 | 2.84E-05 | 0.009 |

| | | | | | | | |
|---|---|---|---|---|---|---|---|
| | | 38,119,198 | *GSDMA* | | 7.214 | 4.01E-11 | 0.009 |
| | | 38,119,207 | *GSDMA* | | 7.866 | 1.24E-12 | 0.009 |
| | | 38,119,649 | | *GSDMA* | 6.033 | 1.57E-08 | 0.002 |
| | | 38,119,760 | | *GSDMA* | 5.590 | 1.28E-07 | 0.011 |
| | | 38,120,234 | | *GSDMA* | 3.425 | 8.23E-04 | 0.011 |
| | | 38,120,370 | | *GSDMA* | 3.512 | 6.12E-04 | 0.011 |
| | | 38,128,699 | | *GSDMA* | -4.310 | 3.20E-05 | 0.006 |
| rs72832972 | g.38110575.T>C | 37,844,283 | *ERBB2* | *PGAP3* | 3.480 | 6.83E-04 | 7.82E-05 |
| | | 37,895,143 | | | -3.656 | 3.71E-04 | 0.004 |
| rs17609240 | g.38110689.T>G | 37,894,590 | | | 3.766 | 2.50E-04 | 0.013 |
| | | 37,910,697 | | | 3.575 | 4.92E-04 | 0.035 |
| | | 38,077,464 | | *ORMDL3* | 5.290 | 5.04E-07 | 0.014 |
| | | 38,080,829 | | *ORMDL3* | 3.407 | 8.75E-04 | 0.014 |
| | | 38,081,995 | | *ORMDL3* | 3.637 | 3.97E-04 | 2.01E-04 |
| | | 38,082,022 | | *ORMDL3* | 4.853 | 3.43E-06 | 6.40E-04 |
| | | 38,082,035 | | *ORMDL3* | 3.788 | 2.31E-04 | 6.40E-04 |
| | | 38,082,073 | | *ORMDL3* | 4.745 | 5.41E-06 | 6.40E-04 |
| | | 38,082,658 | | *ORMDL3* | 4.634 | 8.61E-06 | 0.011 |
| | | 38,108,710 | | | -8.226 | 1.74E-13 | 0.039 |
| | | 38,119,071 | *GSDMA* | | 4.340 | 2.84E-05 | 0.009 |
| | | 38,119,198 | *GSDMA* | | 7.214 | 4.01E-11 | 0.009 |
| | | 38,119,207 | *GSDMA* | | 7.866 | 1.24E-12 | 0.009 |
| | | 38,119,649 | | *GSDMA* | 6.033 | 1.57E-08 | 0.002 |
| | | 38,119,760 | | *GSDMA* | 5.590 | 1.28E-07 | 0.011 |
| | | 38,120,234 | | *GSDMA* | 3.425 | 8.23E-04 | 0.011 |
| | | 38,120,370 | | *GSDMA* | 3.512 | 6.12E-04 | 0.011 |
| | | 38,128,699 | | *GSDMA* | -4.310 | 3.20E-05 | 0.006 |
| rs8076474 | g.38111234.G>C | 37,894,590 | | | 3.766 | 2.50E-04 | 0.013 |
| | | 37,910,697 | | | 3.575 | 4.92E-04 | 0.035 |
| | | 38,077,464 | | *ORMDL3* | 5.290 | 5.04E-07 | 0.014 |
| | | 38,080,829 | | *ORMDL3* | 3.407 | 8.75E-04 | 0.014 |
| | | 38,081,995 | | *ORMDL3* | 3.637 | 3.97E-04 | 2.01E-04 |
| | | 38,082,022 | | *ORMDL3* | 4.853 | 3.43E-06 | 6.40E-04 |
| | | 38,082,035 | | *ORMDL3* | 3.788 | 2.31E-04 | 6.40E-04 |
| | | 38,082,073 | | *ORMDL3* | 4.745 | 5.41E-06 | 6.40E-04 |
| | | 38,082,658 | | *ORMDL3* | 4.634 | 8.61E-06 | 0.011 |
| | | 38,108,710 | | | -8.226 | 1.74E-13 | 0.039 |
| | | 38,119,071 | *GSDMA* | | 4.340 | 2.84E-05 | 0.009 |
| | | 38,119,198 | *GSDMA* | | 7.214 | 4.01E-11 | 0.009 |
| | | 38,119,207 | *GSDMA* | | 7.866 | 1.24E-12 | 0.009 |
| | | 38,119,649 | | *GSDMA* | 6.033 | 1.57E-08 | 0.002 |

| | | | | | | | |
|---|---|---|---|---|---|---|---|
| | | 38,119,760 | | *GSDMA* | 5.590 | 1.28E-07 | 0.011 |
| | | 38,120,234 | | *GSDMA* | 3.425 | 8.23E-04 | 0.011 |
| | | 38,120,370 | | *GSDMA* | 3.512 | 6.12E-04 | 0.011 |
| | | 38,128,699 | | *GSDMA* | -4.310 | 3.20E-05 | 0.006 |
| rs1007654 | g.38111354.A>G | 37,894,590 | | | 3.766 | 2.50E-04 | 0.013 |
| | | 37,910,697 | | | 3.575 | 4.92E-04 | 0.035 |
| | | 38,077,464 | | *ORMDL3* | 5.290 | 5.04E-07 | 0.014 |
| | | 38,080,829 | | *ORMDL3* | 3.407 | 8.75E-04 | 0.014 |
| | | 38,081,995 | | *ORMDL3* | 3.637 | 3.97E-04 | 2.01E-04 |
| | | 38,082,022 | | *ORMDL3* | 4.853 | 3.43E-06 | 6.40E-04 |
| | | 38,082,035 | | *ORMDL3* | 3.788 | 2.31E-04 | 6.40E-04 |
| | | 38,082,073 | | *ORMDL3* | 4.745 | 5.41E-06 | 6.40E-04 |
| | | 38,082,658 | | *ORMDL3* | 4.634 | 8.61E-06 | 0.011 |
| | | 38,108,710 | | | -8.226 | 1.74E-13 | 0.039 |
| | | 38,119,071 | *GSDMA* | | 4.340 | 2.84E-05 | 0.009 |
| | | 38,119,198 | *GSDMA* | | 7.214 | 4.01E-11 | 0.009 |
| | | 38,119,207 | *GSDMA* | | 7.866 | 1.24E-12 | 0.009 |
| | | 38,119,649 | | *GSDMA* | 6.033 | 1.57E-08 | 0.002 |
| | | 38,119,760 | | *GSDMA* | 5.590 | 1.28E-07 | 0.011 |
| | | 38,120,234 | | *GSDMA* | 3.425 | 8.23E-04 | 0.011 |
| | | 38,120,370 | | *GSDMA* | 3.512 | 6.12E-04 | 0.011 |
| | | 38,128,699 | | *GSDMA* | -4.310 | 3.20E-05 | 0.006 |
| rs1007655 | g.38111419.G>A | 37,894,590 | | | 3.766 | 2.50E-04 | 0.013 |
| | | 37,910,697 | | | 3.575 | 4.92E-04 | 0.035 |
| | | 38,077,464 | | *ORMDL3* | 5.290 | 5.04E-07 | 0.014 |
| | | 38,080,829 | | *ORMDL3* | 3.407 | 8.75E-04 | 0.014 |
| | | 38,081,995 | | *ORMDL3* | 3.637 | 3.97E-04 | 2.01E-04 |
| | | 38,082,022 | | *ORMDL3* | 4.853 | 3.43E-06 | 6.40E-04 |
| | | 38,082,035 | | *ORMDL3* | 3.788 | 2.31E-04 | 6.40E-04 |
| | | 38,082,073 | | *ORMDL3* | 4.745 | 5.41E-06 | 6.40E-04 |
| | | 38,082,658 | | *ORMDL3* | 4.634 | 8.61E-06 | 0.011 |
| | | 38,108,710 | | | -8.226 | 1.74E-13 | 0.039 |
| | | 38,119,071 | *GSDMA* | | 4.340 | 2.84E-05 | 0.009 |
| | | 38,119,198 | *GSDMA* | | 7.214 | 4.01E-11 | 0.009 |
| | | 38,119,207 | *GSDMA* | | 7.866 | 1.24E-12 | 0.009 |
| | | 38,119,649 | | *GSDMA* | 6.033 | 1.57E-08 | 0.002 |
| | | 38,119,760 | | *GSDMA* | 5.590 | 1.28E-07 | 0.011 |
| | | 38,120,234 | | *GSDMA* | 3.425 | 8.23E-04 | 0.011 |
| | | 38,120,370 | | *GSDMA* | 3.512 | 6.12E-04 | 0.011 |
| | | 38,128,699 | | *GSDMA* | -4.310 | 3.20E-05 | 0.006 |
| rs1563103 | g.38111740.G>A | 37,894,590 | | | 3.766 | 2.50E-04 | 0.013 |

| | | | | | | | |
|---|---|---|---|---|---|---|---|
| | | 37,910,697 | | | 3.575 | 4.92E-04 | 0.035 |
| | | 38,077,464 | | *ORMDL3* | 5.290 | 5.04E-07 | 0.014 |
| | | 38,080,829 | | *ORMDL3* | 3.407 | 8.75E-04 | 0.014 |
| | | 38,081,995 | | *ORMDL3* | 3.637 | 3.97E-04 | 2.01E-04 |
| | | 38,082,022 | | *ORMDL3* | 4.853 | 3.43E-06 | 6.40E-04 |
| | | 38,082,035 | | *ORMDL3* | 3.788 | 2.31E-04 | 6.40E-04 |
| | | 38,082,073 | | *ORMDL3* | 4.745 | 5.41E-06 | 6.40E-04 |
| | | 38,082,658 | | *ORMDL3* | 4.634 | 8.61E-06 | 0.011 |
| | | 38,108,710 | | | -8.226 | 1.74E-13 | 0.039 |
| | | 38,119,071 | *GSDMA* | | 4.340 | 2.84E-05 | 0.009 |
| | | 38,119,198 | *GSDMA* | | 7.214 | 4.01E-11 | 0.009 |
| | | 38,119,207 | *GSDMA* | | 7.866 | 1.24E-12 | 0.009 |
| | | 38,119,649 | | *GSDMA* | 6.033 | 1.57E-08 | 0.002 |
| | | 38,119,760 | | *GSDMA* | 5.590 | 1.28E-07 | 0.011 |
| | | 38,120,234 | | *GSDMA* | 3.425 | 8.23E-04 | 0.011 |
| | | 38,120,370 | | *GSDMA* | 3.512 | 6.12E-04 | 0.011 |
| | | 38,128,699 | | *GSDMA* | -4.310 | 3.20E-05 | 0.006 |
| rs2313640 | g.38111845.G>A | 37,894,590 | | | 3.766 | 2.50E-04 | 0.013 |
| | | 37,910,697 | | | 3.575 | 4.92E-04 | 0.035 |
| | | 38,077,464 | | *ORMDL3* | 5.290 | 5.04E-07 | 0.014 |
| | | 38,080,829 | | *ORMDL3* | 3.407 | 8.75E-04 | 0.014 |
| | | 38,081,995 | | *ORMDL3* | 3.637 | 3.97E-04 | 2.01E-04 |
| | | 38,082,022 | | *ORMDL3* | 4.853 | 3.43E-06 | 6.40E-04 |
| | | 38,082,035 | | *ORMDL3* | 3.788 | 2.31E-04 | 6.40E-04 |
| | | 38,082,073 | | *ORMDL3* | 4.745 | 5.41E-06 | 6.40E-04 |
| | | 38,082,658 | | *ORMDL3* | 4.634 | 8.61E-06 | 0.011 |
| | | 38,108,710 | | | -8.226 | 1.74E-13 | 0.039 |
| | | 38,119,071 | *GSDMA* | | 4.340 | 2.84E-05 | 0.009 |
| | | 38,119,198 | *GSDMA* | | 7.214 | 4.01E-11 | 0.009 |
| | | 38,119,207 | *GSDMA* | | 7.866 | 1.24E-12 | 0.009 |
| | | 38,119,649 | | *GSDMA* | 6.033 | 1.57E-08 | 0.002 |
| | | 38,119,760 | | *GSDMA* | 5.590 | 1.28E-07 | 0.011 |
| | | 38,120,234 | | *GSDMA* | 3.425 | 8.23E-04 | 0.011 |
| | | 38,120,370 | | *GSDMA* | 3.512 | 6.12E-04 | 0.011 |
| | | 38,128,699 | | *GSDMA* | -4.310 | 3.20E-05 | 0.006 |
| rs8068522 | g.38112076.T>C | 37,894,590 | | | 3.422 | 8.32E-04 | 0.013 |
| | | 37,910,697 | | | 3.543 | 5.49E-04 | 0.035 |
| | | 38,077,464 | | *ORMDL3* | 5.311 | 4.58E-07 | 0.014 |
| | | 38,081,995 | | *ORMDL3* | 3.616 | 4.26E-04 | 2.01E-04 |
| | | 38,082,022 | | *ORMDL3* | 4.714 | 6.16E-06 | 6.40E-04 |
| | | 38,082,035 | | *ORMDL3* | 3.629 | 4.07E-04 | 6.40E-04 |

| | | | | | | | |
|---|---|---|---|---|---|---|---|
| | | 38,082,073 | | *ORMDL3* | 4.576 | 1.09E-05 | 6.40E-04 |
| | | 38,082,658 | | *ORMDL3* | 4.575 | 1.10E-05 | 0.011 |
| | | 38,108,710 | | | -8.022 | 5.31E-13 | 0.039 |
| | | 38,119,071 | *GSDMA* | | 4.343 | 2.80E-05 | 0.009 |
| | | 38,119,198 | *GSDMA* | | 7.203 | 4.26E-11 | 0.009 |
| | | 38,119,207 | *GSDMA* | | 7.742 | 2.43E-12 | 0.009 |
| | | 38,119,649 | | *GSDMA* | 6.050 | 1.45E-08 | 0.002 |
| | | 38,119,760 | | *GSDMA* | 5.507 | 1.88E-07 | 0.011 |
| | | 38,120,234 | | *GSDMA* | 3.503 | 6.32E-04 | 0.011 |
| | | 38,120,370 | | *GSDMA* | 3.647 | 3.82E-04 | 0.011 |
| | | 38,128,699 | | *GSDMA* | -4.338 | 2.86E-05 | 0.006 |
| rs8081437 | g.38112114.G>A | 37,894,590 | | | 3.763 | 2.53E-04 | 0.013 |
| | | 37,910,697 | | | 3.582 | 4.80E-04 | 0.035 |
| | | 38,077,464 | | *ORMDL3* | 5.208 | 7.27E-07 | 0.014 |
| | | 38,081,995 | | *ORMDL3* | 3.562 | 5.16E-04 | 2.01E-04 |
| | | 38,082,022 | | *ORMDL3* | 4.917 | 2.61E-06 | 6.40E-04 |
| | | 38,082,035 | | *ORMDL3* | 3.834 | 1.95E-04 | 6.40E-04 |
| | | 38,082,073 | | *ORMDL3* | 4.861 | 3.31E-06 | 6.40E-04 |
| | | 38,082,658 | | *ORMDL3* | 4.590 | 1.03E-05 | 0.011 |
| | | 38,108,710 | | | -8.167 | 2.41E-13 | 0.039 |
| | | 38,119,071 | *GSDMA* | | 4.046 | 8.90E-05 | 0.009 |
| | | 38,119,198 | *GSDMA* | | 7.181 | 4.78E-11 | 0.009 |
| | | 38,119,207 | *GSDMA* | | 7.760 | 2.20E-12 | 0.009 |
| | | 38,119,649 | | *GSDMA* | 6.010 | 1.75E-08 | 0.002 |
| | | 38,119,760 | | *GSDMA* | 5.250 | 6.03E-07 | 0.011 |
| | | 38,120,370 | | *GSDMA* | 3.599 | 4.53E-04 | 0.011 |
| | | 38,122,024 | | *GSDMA* | -3.404 | 8.83E-04 | 0.011 |
| | | 38,128,699 | | *GSDMA* | -4.371 | 2.50E-05 | 0.006 |
| rs8081462 | g.38112190.C>G | 37,834,869 | | *PGAP3* | 3.453 | 7.49E-04 | 0.020 |
| | | 37,910,697 | | | 3.491 | 6.58E-04 | 0.035 |
| | | 38,077,277 | | | 4.203 | 4.86E-05 | 0.014 |
| | | 38,077,464 | | *ORMDL3* | 4.071 | 8.07E-05 | 0.014 |
| | | 38,080,829 | | *ORMDL3* | 3.663 | 3.61E-04 | 0.014 |
| | | 38,081,530 | | *ORMDL3* | 3.708 | 3.08E-04 | 2.01E-04 |
| | | 38,081,812 | | *ORMDL3* | 4.038 | 9.16E-05 | 2.01E-04 |
| | | 38,082,022 | | *ORMDL3* | 3.625 | 4.13E-04 | 6.40E-04 |
| | | 38,082,073 | | *ORMDL3* | 4.886 | 2.98E-06 | 6.40E-04 |
| | | 38,082,346 | | *ORMDL3* | 3.433 | 8.01E-04 | 0.011 |
| | | 38,082,658 | | *ORMDL3* | 4.551 | 1.21E-05 | 0.011 |
| | | 38,108,710 | | | -6.111 | 1.07E-08 | 0.039 |
| | | 38,119,071 | *GSDMA* | | 4.281 | 3.59E-05 | 0.009 |

| | | | | | | | |
|---|---|---|---|---|---|---|---|
| | | 38,119,198 | *GSDMA* | | 6.240 | 5.72E-09 | 0.009 |
| | | 38,119,207 | *GSDMA* | | 7.498 | 8.96E-12 | 0.009 |
| | | 38,119,649 | | *GSDMA* | 6.524 | 1.39E-09 | 0.002 |
| | | 38,119,760 | | *GSDMA* | 4.665 | 7.58E-06 | 0.011 |
| | | 38,120,234 | | *GSDMA* | 3.402 | 8.90E-04 | 0.011 |
| | | 38,128,699 | | *GSDMA* | -4.073 | 8.01E-05 | 0.006 |
| rs62068174 | g.38112255.G>C | 37,894,590 | | | 3.766 | 2.50E-04 | 0.013 |
| | | 37,910,697 | | | 3.575 | 4.92E-04 | 0.035 |
| | | 38,077,464 | | *ORMDL3* | 5.290 | 5.04E-07 | 0.014 |
| | | 38,080,829 | | *ORMDL3* | 3.407 | 8.75E-04 | 0.014 |
| | | 38,081,995 | | *ORMDL3* | 3.637 | 3.97E-04 | 2.01E-04 |
| | | 38,082,022 | | *ORMDL3* | 4.853 | 3.43E-06 | 6.40E-04 |
| | | 38,082,035 | | *ORMDL3* | 3.788 | 2.31E-04 | 6.40E-04 |
| | | 38,082,073 | | *ORMDL3* | 4.745 | 5.41E-06 | 6.40E-04 |
| | | 38,082,658 | | *ORMDL3* | 4.634 | 8.61E-06 | 0.011 |
| | | 38,108,710 | | | -8.226 | 1.74E-13 | 0.039 |
| | | 38,119,071 | *GSDMA* | | 4.340 | 2.84E-05 | 0.009 |
| | | 38,119,198 | *GSDMA* | | 7.214 | 4.01E-11 | 0.009 |
| | | 38,119,207 | *GSDMA* | | 7.866 | 1.24E-12 | 0.009 |
| | | 38,119,649 | | *GSDMA* | 6.033 | 1.57E-08 | 0.002 |
| | | 38,119,760 | | *GSDMA* | 5.590 | 1.28E-07 | 0.011 |
| | | 38,120,234 | | *GSDMA* | 3.425 | 8.23E-04 | 0.011 |
| | | 38,120,370 | | *GSDMA* | 3.512 | 6.12E-04 | 0.011 |
| | | 38,128,699 | | *GSDMA* | -4.310 | 3.20E-05 | 0.006 |
| rs62068175 | g.38112300.A>C | 37,894,590 | | | 3.766 | 2.50E-04 | 0.013 |
| | | 37,910,697 | | | 3.575 | 4.92E-04 | 0.035 |
| | | 38,077,464 | | *ORMDL3* | 5.290 | 5.04E-07 | 0.014 |
| | | 38,080,829 | | *ORMDL3* | 3.407 | 8.75E-04 | 0.014 |
| | | 38,081,995 | | *ORMDL3* | 3.637 | 3.97E-04 | 2.01E-04 |
| | | 38,082,022 | | *ORMDL3* | 4.853 | 3.43E-06 | 6.40E-04 |
| | | 38,082,035 | | *ORMDL3* | 3.788 | 2.31E-04 | 6.40E-04 |
| | | 38,082,073 | | *ORMDL3* | 4.745 | 5.41E-06 | 6.40E-04 |
| | | 38,082,658 | | *ORMDL3* | 4.634 | 8.61E-06 | 0.011 |
| | | 38,108,710 | | | -8.226 | 1.74E-13 | 0.039 |
| | | 38,119,071 | *GSDMA* | | 4.340 | 2.84E-05 | 0.009 |
| | | 38,119,198 | *GSDMA* | | 7.214 | 4.01E-11 | 0.009 |
| | | 38,119,207 | *GSDMA* | | 7.866 | 1.24E-12 | 0.009 |
| | | 38,119,649 | | *GSDMA* | 6.033 | 1.57E-08 | 0.002 |
| | | 38,119,760 | | *GSDMA* | 5.590 | 1.28E-07 | 0.011 |
| | | 38,120,234 | | *GSDMA* | 3.425 | 8.23E-04 | 0.011 |
| | | 38,120,370 | | *GSDMA* | 3.512 | 6.12E-04 | 0.011 |

| | | | | | | | |
|---|---|---|---|---|---|---|---|
| | | 38,128,699 | | *GSDMA* | -4.310 | 3.20E-05 | 0.006 |
| rs62068176 | g.38112438.A>G | 37,844,069 | *ERBB2* | *PGAP3* | 3.581 | 4.83E-04 | 7.82E-05 |
| | | 37,894,590 | | | 3.679 | 3.42E-04 | 0.013 |
| | | 37,910,697 | | | 3.572 | 4.97E-04 | 0.035 |
| | | 38,077,464 | | *ORMDL3* | 5.199 | 7.58E-07 | 0.014 |
| | | 38,080,829 | | *ORMDL3* | 3.391 | 9.24E-04 | 0.014 |
| | | 38,082,022 | | *ORMDL3* | 4.700 | 6.55E-06 | 6.40E-04 |
| | | 38,082,035 | | *ORMDL3* | 3.704 | 3.12E-04 | 6.40E-04 |
| | | 38,082,073 | | *ORMDL3* | 4.500 | 1.49E-05 | 6.40E-04 |
| | | 38,082,658 | | *ORMDL3* | 4.536 | 1.29E-05 | 0.011 |
| | | 38,108,710 | | | -8.056 | 4.41E-13 | 0.039 |
| | | 38,119,071 | *GSDMA* | | 4.538 | 1.28E-05 | 0.009 |
| | | 38,119,198 | *GSDMA* | | 6.753 | 4.35E-10 | 0.009 |
| | | 38,119,207 | *GSDMA* | | 7.930 | 8.78E-13 | 0.009 |
| | | 38,119,649 | | *GSDMA* | 5.975 | 2.07E-08 | 0.002 |
| | | 38,119,760 | | *GSDMA* | 5.719 | 7.00E-08 | 0.011 |
| | | 38,122,024 | | *GSDMA* | -3.593 | 4.62E-04 | 0.011 |
| | | 38,128,699 | | *GSDMA* | -4.242 | 4.17E-05 | 0.006 |
| rs62068177 | g.38112601.G>A | 37,844,069 | *ERBB2* | *PGAP3* | 3.496 | 6.46E-04 | 7.82E-05 |
| | | 37,910,697 | | | 3.897 | 1.55E-04 | 0.035 |
| | | 38,077,464 | | *ORMDL3* | 5.154 | 9.23E-07 | 0.014 |
| | | 38,081,995 | | *ORMDL3* | 3.440 | 7.82E-04 | 2.01E-04 |
| | | 38,082,022 | | *ORMDL3* | 4.405 | 2.19E-05 | 6.40E-04 |
| | | 38,082,035 | | *ORMDL3* | 3.374 | 9.75E-04 | 6.40E-04 |
| | | 38,082,073 | | *ORMDL3* | 4.761 | 5.07E-06 | 6.40E-04 |
| | | 38,082,658 | | *ORMDL3* | 4.401 | 2.23E-05 | 0.011 |
| | | 38,084,377 | *ORMDL3* | | 3.388 | 9.33E-04 | 0.005 |
| | | 38,108,710 | | | -7.957 | 7.58E-13 | 0.039 |
| | | 38,119,071 | *GSDMA* | | 4.347 | 2.76E-05 | 0.009 |
| | | 38,119,132 | *GSDMA* | | 3.414 | 8.55E-04 | 0.009 |
| | | 38,119,198 | *GSDMA* | | 6.665 | 6.83E-10 | 0.009 |
| | | 38,119,207 | *GSDMA* | | 7.113 | 6.81E-11 | 0.009 |
| | | 38,119,649 | | *GSDMA* | 6.014 | 1.71E-08 | 0.002 |
| | | 38,119,760 | | *GSDMA* | 5.607 | 1.18E-07 | 0.011 |
| | | 38,122,024 | | *GSDMA* | -3.552 | 5.33E-04 | 0.011 |
| | | 38,128,699 | | *GSDMA* | -3.463 | 7.23E-04 | 0.006 |
| rs62068178 | g.38112608.C>T | 37,844,069 | *ERBB2* | *PGAP3* | 3.496 | 6.46E-04 | 7.82E-05 |
| | | 37,910,697 | | | 3.897 | 1.55E-04 | 0.035 |
| | | 38,077,464 | | *ORMDL3* | 5.154 | 9.23E-07 | 0.014 |
| | | 38,081,995 | | *ORMDL3* | 3.440 | 7.82E-04 | 2.01E-04 |
| | | 38,082,022 | | *ORMDL3* | 4.405 | 2.19E-05 | 6.40E-04 |

| | | | | | | | |
|---|---|---|---|---|---|---|---|
| | | 38,082,035 | | *ORMDL3* | 3.374 | 9.75E-04 | 6.40E-04 |
| | | 38,082,073 | | *ORMDL3* | 4.761 | 5.07E-06 | 6.40E-04 |
| | | 38,082,658 | | *ORMDL3* | 4.401 | 2.23E-05 | 0.011 |
| | | 38,084,377 | *ORMDL3* | | 3.388 | 9.33E-04 | 0.005 |
| | | 38,108,710 | | | -7.957 | 7.58E-13 | 0.039 |
| | | 38,119,071 | *GSDMA* | | 4.347 | 2.76E-05 | 0.009 |
| | | 38,119,132 | *GSDMA* | | 3.414 | 8.55E-04 | 0.009 |
| | | 38,119,198 | *GSDMA* | | 6.665 | 6.83E-10 | 0.009 |
| | | 38,119,207 | *GSDMA* | | 7.113 | 6.81E-11 | 0.009 |
| | | 38,119,649 | | *GSDMA* | 6.014 | 1.71E-08 | 0.002 |
| | | 38,119,760 | | *GSDMA* | 5.607 | 1.18E-07 | 0.011 |
| | | 38,122,024 | | *GSDMA* | -3.552 | 5.33E-04 | 0.011 |
| | | 38,128,699 | | *GSDMA* | -3.463 | 7.23E-04 | 0.006 |
| rs62068179 | g.38112617.G>A | 37,844,069 | *ERBB2* | *PGAP3* | 3.496 | 6.46E-04 | 7.82E-05 |
| | | 37,910,697 | | | 3.897 | 1.55E-04 | 0.035 |
| | | 38,077,464 | | *ORMDL3* | 5.154 | 9.23E-07 | 0.014 |
| | | 38,081,995 | | *ORMDL3* | 3.440 | 7.82E-04 | 2.01E-04 |
| | | 38,082,022 | | *ORMDL3* | 4.405 | 2.19E-05 | 6.40E-04 |
| | | 38,082,035 | | *ORMDL3* | 3.374 | 9.75E-04 | 6.40E-04 |
| | | 38,082,073 | | *ORMDL3* | 4.761 | 5.07E-06 | 6.40E-04 |
| | | 38,082,658 | | *ORMDL3* | 4.401 | 2.23E-05 | 0.011 |
| | | 38,084,377 | *ORMDL3* | | 3.388 | 9.33E-04 | 0.005 |
| | | 38,108,710 | | | -7.957 | 7.58E-13 | 0.039 |
| | | 38,119,071 | *GSDMA* | | 4.347 | 2.76E-05 | 0.009 |
| | | 38,119,132 | *GSDMA* | | 3.414 | 8.55E-04 | 0.009 |
| | | 38,119,198 | *GSDMA* | | 6.665 | 6.83E-10 | 0.009 |
| | | 38,119,207 | *GSDMA* | | 7.113 | 6.81E-11 | 0.009 |
| | | 38,119,649 | | *GSDMA* | 6.014 | 1.71E-08 | 0.002 |
| | | 38,119,760 | | *GSDMA* | 5.607 | 1.18E-07 | 0.011 |
| | | 38,122,024 | | *GSDMA* | -3.552 | 5.33E-04 | 0.011 |
| | | 38,128,699 | | *GSDMA* | -3.463 | 7.23E-04 | 0.006 |
| rs67480438 | g.38112774.A>G | 37,887,061 | *MIEN1* | | 3.389 | 9.30E-04 | 0.002 |
| | | 37,894,590 | | | 3.738 | 2.77E-04 | 0.013 |
| | | 37,910,697 | | | 3.613 | 4.31E-04 | 0.035 |
| | | 38,077,464 | | *ORMDL3* | 5.374 | 3.45E-07 | 0.014 |
| | | 38,080,829 | | *ORMDL3* | 3.405 | 8.82E-04 | 0.014 |
| | | 38,081,995 | | *ORMDL3* | 3.464 | 7.21E-04 | 2.01E-04 |
| | | 38,082,022 | | *ORMDL3* | 4.700 | 6.52E-06 | 6.40E-04 |
| | | 38,082,035 | | *ORMDL3* | 3.734 | 2.81E-04 | 6.40E-04 |
| | | 38,082,073 | | *ORMDL3* | 4.565 | 1.14E-05 | 6.40E-04 |
| | | 38,082,658 | | *ORMDL3* | 4.453 | 1.81E-05 | 0.011 |

| | | | | | | | |
|---|---|---|---|---|---|---|---|
| | | 38,108,710 | | | -8.262 | 1.43E-13 | 0.039 |
| | | 38,119,071 | *GSDMA* | | 4.249 | 4.06E-05 | 0.009 |
| | | 38,119,198 | *GSDMA* | | 7.201 | 4.29E-11 | 0.009 |
| | | 38,119,207 | *GSDMA* | | 7.868 | 1.23E-12 | 0.009 |
| | | 38,119,649 | | *GSDMA* | 5.965 | 2.17E-08 | 0.002 |
| | | 38,119,760 | | *GSDMA* | 5.650 | 9.69E-08 | 0.011 |
| | | 38,120,370 | | *GSDMA* | 3.496 | 6.47E-04 | 0.011 |
| | | 38,122,024 | | *GSDMA* | -3.424 | 8.25E-04 | 0.011 |
| | | 38,128,699 | | *GSDMA* | -4.382 | 2.41E-05 | 0.006 |
| rs62068180 | g.38112825.T>C | 37,887,061 | *MIEN1* | | 3.389 | 9.30E-04 | 0.002 |
| | | 37,894,590 | | | 3.738 | 2.77E-04 | 0.013 |
| | | 37,910,697 | | | 3.613 | 4.31E-04 | 0.035 |
| | | 38,077,464 | | *ORMDL3* | 5.374 | 3.45E-07 | 0.014 |
| | | 38,080,829 | | *ORMDL3* | 3.405 | 8.82E-04 | 0.014 |
| | | 38,081,995 | | *ORMDL3* | 3.464 | 7.21E-04 | 2.01E-04 |
| | | 38,082,022 | | *ORMDL3* | 4.700 | 6.52E-06 | 6.40E-04 |
| | | 38,082,035 | | *ORMDL3* | 3.734 | 2.81E-04 | 6.40E-04 |
| | | 38,082,073 | | *ORMDL3* | 4.565 | 1.14E-05 | 6.40E-04 |
| | | 38,082,658 | | *ORMDL3* | 4.453 | 1.81E-05 | 0.011 |
| | | 38,108,710 | | | -8.262 | 1.43E-13 | 0.039 |
| | | 38,119,071 | *GSDMA* | | 4.249 | 4.06E-05 | 0.009 |
| | | 38,119,198 | *GSDMA* | | 7.201 | 4.29E-11 | 0.009 |
| | | 38,119,207 | *GSDMA* | | 7.868 | 1.23E-12 | 0.009 |
| | | 38,119,649 | | *GSDMA* | 5.965 | 2.17E-08 | 0.002 |
| | | 38,119,760 | | *GSDMA* | 5.650 | 9.69E-08 | 0.011 |
| | | 38,120,370 | | *GSDMA* | 3.496 | 6.47E-04 | 0.011 |
| | | 38,122,024 | | *GSDMA* | -3.424 | 8.25E-04 | 0.011 |
| | | 38,128,699 | | *GSDMA* | -4.382 | 2.41E-05 | 0.006 |
| rs62068181 | g.38112832.T>C | 37,887,061 | *MIEN1* | | 3.389 | 9.30E-04 | 0.002 |
| | | 37,894,590 | | | 3.738 | 2.77E-04 | 0.013 |
| | | 37,910,697 | | | 3.613 | 4.31E-04 | 0.035 |
| | | 38,077,464 | | *ORMDL3* | 5.374 | 3.45E-07 | 0.014 |
| | | 38,080,829 | | *ORMDL3* | 3.405 | 8.82E-04 | 0.014 |
| | | 38,081,995 | | *ORMDL3* | 3.464 | 7.21E-04 | 2.01E-04 |
| | | 38,082,022 | | *ORMDL3* | 4.700 | 6.52E-06 | 6.40E-04 |
| | | 38,082,035 | | *ORMDL3* | 3.734 | 2.81E-04 | 6.40E-04 |
| | | 38,082,073 | | *ORMDL3* | 4.565 | 1.14E-05 | 6.40E-04 |
| | | 38,082,658 | | *ORMDL3* | 4.453 | 1.81E-05 | 0.011 |
| | | 38,108,710 | | | -8.262 | 1.43E-13 | 0.039 |
| | | 38,119,071 | *GSDMA* | | 4.249 | 4.06E-05 | 0.009 |
| | | 38,119,198 | *GSDMA* | | 7.201 | 4.29E-11 | 0.009 |

| | | | | | | | |
|---|---|---|---|---|---|---|---|
| | | 38,119,207 | *GSDMA* | | 7.868 | 1.23E-12 | 0.009 |
| | | 38,119,649 | | *GSDMA* | 5.965 | 2.17E-08 | 0.002 |
| | | 38,119,760 | | *GSDMA* | 5.650 | 9.69E-08 | 0.011 |
| | | 38,120,370 | | *GSDMA* | 3.496 | 6.47E-04 | 0.011 |
| | | 38,122,024 | | *GSDMA* | -3.424 | 8.25E-04 | 0.011 |
| | | 38,128,699 | | *GSDMA* | -4.382 | 2.41E-05 | 0.006 |
| rs62068182 | g.38112864.G>A | 37,887,061 | *MIEN1* | | 3.389 | 9.30E-04 | 0.002 |
| | | 37,894,590 | | | 3.738 | 2.77E-04 | 0.013 |
| | | 37,910,697 | | | 3.613 | 4.31E-04 | 0.035 |
| | | 38,077,464 | | *ORMDL3* | 5.374 | 3.45E-07 | 0.014 |
| | | 38,080,829 | | *ORMDL3* | 3.405 | 8.82E-04 | 0.014 |
| | | 38,081,995 | | *ORMDL3* | 3.464 | 7.21E-04 | 2.01E-04 |
| | | 38,082,022 | | *ORMDL3* | 4.700 | 6.52E-06 | 6.40E-04 |
| | | 38,082,035 | | *ORMDL3* | 3.734 | 2.81E-04 | 6.40E-04 |
| | | 38,082,073 | | *ORMDL3* | 4.565 | 1.14E-05 | 6.40E-04 |
| | | 38,082,658 | | *ORMDL3* | 4.453 | 1.81E-05 | 0.011 |
| | | 38,108,710 | | | -8.262 | 1.43E-13 | 0.039 |
| | | 38,119,071 | *GSDMA* | | 4.249 | 4.06E-05 | 0.009 |
| | | 38,119,198 | *GSDMA* | | 7.201 | 4.29E-11 | 0.009 |
| | | 38,119,207 | *GSDMA* | | 7.868 | 1.23E-12 | 0.009 |
| | | 38,119,649 | | *GSDMA* | 5.965 | 2.17E-08 | 0.002 |
| | | 38,119,760 | | *GSDMA* | 5.650 | 9.69E-08 | 0.011 |
| | | 38,120,370 | | *GSDMA* | 3.496 | 6.47E-04 | 0.011 |
| | | 38,122,024 | | *GSDMA* | -3.424 | 8.25E-04 | 0.011 |
| | | 38,128,699 | | *GSDMA* | -4.382 | 2.41E-05 | 0.006 |
| rs62068183 | g.38112918.C>T | 37,894,590 | | | 3.766 | 2.50E-04 | 0.013 |
| | | 37,910,697 | | | 3.575 | 4.92E-04 | 0.035 |
| | | 38,077,464 | | *ORMDL3* | 5.290 | 5.04E-07 | 0.014 |
| | | 38,080,829 | | *ORMDL3* | 3.407 | 8.75E-04 | 0.014 |
| | | 38,081,995 | | *ORMDL3* | 3.637 | 3.97E-04 | 2.01E-04 |
| | | 38,082,022 | | *ORMDL3* | 4.853 | 3.43E-06 | 6.40E-04 |
| | | 38,082,035 | | *ORMDL3* | 3.788 | 2.31E-04 | 6.40E-04 |
| | | 38,082,073 | | *ORMDL3* | 4.745 | 5.41E-06 | 6.40E-04 |
| | | 38,082,658 | | *ORMDL3* | 4.634 | 8.61E-06 | 0.011 |
| | | 38,108,710 | | | -8.226 | 1.74E-13 | 0.039 |
| | | 38,119,071 | *GSDMA* | | 4.340 | 2.84E-05 | 0.009 |
| | | 38,119,198 | *GSDMA* | | 7.214 | 4.01E-11 | 0.009 |
| | | 38,119,207 | *GSDMA* | | 7.866 | 1.24E-12 | 0.009 |
| | | 38,119,649 | | *GSDMA* | 6.033 | 1.57E-08 | 0.002 |
| | | 38,119,760 | | *GSDMA* | 5.590 | 1.28E-07 | 0.011 |
| | | 38,120,234 | | *GSDMA* | 3.425 | 8.23E-04 | 0.011 |

| | | | | | | | |
|---|---|---|---|---|---|---|---|
| | | 38,120,370 | | *GSDMA* | 3.512 | 6.12E-04 | 0.011 |
| | | 38,128,699 | | *GSDMA* | -4.310 | 3.20E-05 | 0.006 |
| rs62068184 | g.38113054.C>T | 37,894,590 | | | 3.766 | 2.50E-04 | 0.013 |
| | | 37,910,697 | | | 3.575 | 4.92E-04 | 0.035 |
| | | 38,077,464 | | *ORMDL3* | 5.290 | 5.04E-07 | 0.014 |
| | | 38,080,829 | | *ORMDL3* | 3.407 | 8.75E-04 | 0.014 |
| | | 38,081,995 | | *ORMDL3* | 3.637 | 3.97E-04 | 2.01E-04 |
| | | 38,082,022 | | *ORMDL3* | 4.853 | 3.43E-06 | 6.40E-04 |
| | | 38,082,035 | | *ORMDL3* | 3.788 | 2.31E-04 | 6.40E-04 |
| | | 38,082,073 | | *ORMDL3* | 4.745 | 5.41E-06 | 6.40E-04 |
| | | 38,082,658 | | *ORMDL3* | 4.634 | 8.61E-06 | 0.011 |
| | | 38,108,710 | | | -8.226 | 1.74E-13 | 0.039 |
| | | 38,119,071 | *GSDMA* | | 4.340 | 2.84E-05 | 0.009 |
| | | 38,119,198 | *GSDMA* | | 7.214 | 4.01E-11 | 0.009 |
| | | 38,119,207 | *GSDMA* | | 7.866 | 1.24E-12 | 0.009 |
| | | 38,119,649 | | *GSDMA* | 6.033 | 1.57E-08 | 0.002 |
| | | 38,119,760 | | *GSDMA* | 5.590 | 1.28E-07 | 0.011 |
| | | 38,120,234 | | *GSDMA* | 3.425 | 8.23E-04 | 0.011 |
| | | 38,120,370 | | *GSDMA* | 3.512 | 6.12E-04 | 0.011 |
| | | 38,128,699 | | *GSDMA* | -4.310 | 3.20E-05 | 0.006 |
| rs4134500 | g.38113270.G>A | 37,894,590 | | | 3.802 | 2.20E-04 | 0.013 |
| | | 37,910,697 | | | 3.602 | 4.49E-04 | 0.035 |
| | | 38,077,464 | | *ORMDL3* | 5.179 | 8.27E-07 | 0.014 |
| | | 38,080,829 | | *ORMDL3* | 3.438 | 7.89E-04 | 0.014 |
| | | 38,081,995 | | *ORMDL3* | 3.654 | 3.74E-04 | 2.01E-04 |
| | | 38,082,022 | | *ORMDL3* | 4.899 | 2.82E-06 | 6.40E-04 |
| | | 38,082,035 | | *ORMDL3* | 3.827 | 2.00E-04 | 6.40E-04 |
| | | 38,082,073 | | *ORMDL3* | 4.776 | 4.74E-06 | 6.40E-04 |
| | | 38,082,658 | | *ORMDL3* | 4.586 | 1.05E-05 | 0.011 |
| | | 38,108,710 | | | -8.180 | 2.24E-13 | 0.039 |
| | | 38,119,071 | *GSDMA* | | 4.307 | 3.23E-05 | 0.009 |
| | | 38,119,198 | *GSDMA* | | 7.195 | 4.44E-11 | 0.009 |
| | | 38,119,207 | *GSDMA* | | 7.841 | 1.42E-12 | 0.009 |
| | | 38,119,649 | | *GSDMA* | 6.023 | 1.64E-08 | 0.002 |
| | | 38,119,760 | | *GSDMA* | 5.633 | 1.05E-07 | 0.011 |
| | | 38,120,370 | | *GSDMA* | 3.467 | 7.14E-04 | 0.011 |
| | | 38,128,699 | | *GSDMA* | -4.379 | 2.43E-05 | 0.006 |
| rs4134499 | g.38113274.G>A | 37,894,590 | | | 3.802 | 2.20E-04 | 0.013 |
| | | 37,910,697 | | | 3.602 | 4.49E-04 | 0.035 |
| | | 38,077,464 | | *ORMDL3* | 5.179 | 8.27E-07 | 0.014 |
| | | 38,080,829 | | *ORMDL3* | 3.438 | 7.89E-04 | 0.014 |

| | | | | | | | |
|---|---|---|---|---|---|---|---|
| | | 38,081,995 | | *ORMDL3* | 3.654 | 3.74E-04 | 2.01E-04 |
| | | 38,082,022 | | *ORMDL3* | 4.899 | 2.82E-06 | 6.40E-04 |
| | | 38,082,035 | | *ORMDL3* | 3.827 | 2.00E-04 | 6.40E-04 |
| | | 38,082,073 | | *ORMDL3* | 4.776 | 4.74E-06 | 6.40E-04 |
| | | 38,082,658 | | *ORMDL3* | 4.586 | 1.05E-05 | 0.011 |
| | | 38,108,710 | | | -8.180 | 2.24E-13 | 0.039 |
| | | 38,119,071 | *GSDMA* | | 4.307 | 3.23E-05 | 0.009 |
| | | 38,119,198 | *GSDMA* | | 7.195 | 4.44E-11 | 0.009 |
| | | 38,119,207 | *GSDMA* | | 7.841 | 1.42E-12 | 0.009 |
| | | 38,119,649 | | *GSDMA* | 6.023 | 1.64E-08 | 0.002 |
| | | 38,119,760 | | *GSDMA* | 5.633 | 1.05E-07 | 0.011 |
| | | 38,120,370 | | *GSDMA* | 3.467 | 7.14E-04 | 0.011 |
| | | 38,128,699 | | *GSDMA* | -4.379 | 2.43E-05 | 0.006 |
| rs4134417 | g.38113278.G>A | 37,894,590 | | | 3.802 | 2.20E-04 | 0.013 |
| | | 37,910,697 | | | 3.602 | 4.49E-04 | 0.035 |
| | | 38,077,464 | | *ORMDL3* | 5.179 | 8.27E-07 | 0.014 |
| | | 38,080,829 | | *ORMDL3* | 3.438 | 7.89E-04 | 0.014 |
| | | 38,081,995 | | *ORMDL3* | 3.654 | 3.74E-04 | 2.01E-04 |
| | | 38,082,022 | | *ORMDL3* | 4.899 | 2.82E-06 | 6.40E-04 |
| | | 38,082,035 | | *ORMDL3* | 3.827 | 2.00E-04 | 6.40E-04 |
| | | 38,082,073 | | *ORMDL3* | 4.776 | 4.74E-06 | 6.40E-04 |
| | | 38,082,658 | | *ORMDL3* | 4.586 | 1.05E-05 | 0.011 |
| | | 38,108,710 | | | -8.180 | 2.24E-13 | 0.039 |
| | | 38,119,071 | *GSDMA* | | 4.307 | 3.23E-05 | 0.009 |
| | | 38,119,198 | *GSDMA* | | 7.195 | 4.44E-11 | 0.009 |
| | | 38,119,207 | *GSDMA* | | 7.841 | 1.42E-12 | 0.009 |
| | | 38,119,649 | | *GSDMA* | 6.023 | 1.64E-08 | 0.002 |
| | | 38,119,760 | | *GSDMA* | 5.633 | 1.05E-07 | 0.011 |
| | | 38,120,370 | | *GSDMA* | 3.467 | 7.14E-04 | 0.011 |
| | | 38,128,699 | | *GSDMA* | -4.379 | 2.43E-05 | 0.006 |
| rs4134498 | g.38113282.G>A | 37,894,590 | | | 3.802 | 2.20E-04 | 0.013 |
| | | 37,910,697 | | | 3.602 | 4.49E-04 | 0.035 |
| | | 38,077,464 | | *ORMDL3* | 5.179 | 8.27E-07 | 0.014 |
| | | 38,080,829 | | *ORMDL3* | 3.438 | 7.89E-04 | 0.014 |
| | | 38,081,995 | | *ORMDL3* | 3.654 | 3.74E-04 | 2.01E-04 |
| | | 38,082,022 | | *ORMDL3* | 4.899 | 2.82E-06 | 6.40E-04 |
| | | 38,082,035 | | *ORMDL3* | 3.827 | 2.00E-04 | 6.40E-04 |
| | | 38,082,073 | | *ORMDL3* | 4.776 | 4.74E-06 | 6.40E-04 |
| | | 38,082,658 | | *ORMDL3* | 4.586 | 1.05E-05 | 0.011 |
| | | 38,108,710 | | | -8.180 | 2.24E-13 | 0.039 |
| | | 38,119,071 | *GSDMA* | | 4.307 | 3.23E-05 | 0.009 |

| | | | | | | | |
|---|---|---|---|---|---|---|---|
| | | 38,119,198 | *GSDMA* | | 7.195 | 4.44E-11 | 0.009 |
| | | 38,119,207 | *GSDMA* | | 7.841 | 1.42E-12 | 0.009 |
| | | 38,119,649 | | *GSDMA* | 6.023 | 1.64E-08 | 0.002 |
| | | 38,119,760 | | *GSDMA* | 5.633 | 1.05E-07 | 0.011 |
| | | 38,120,370 | | *GSDMA* | 3.467 | 7.14E-04 | 0.011 |
| | | 38,128,699 | | *GSDMA* | -4.379 | 2.43E-05 | 0.006 |
| rs4134502 | g.38113286.G>A | 37,894,590 | | | 3.802 | 2.20E-04 | 0.013 |
| | | 37,910,697 | | | 3.602 | 4.49E-04 | 0.035 |
| | | 38,077,464 | | *ORMDL3* | 5.179 | 8.27E-07 | 0.014 |
| | | 38,080,829 | | *ORMDL3* | 3.438 | 7.89E-04 | 0.014 |
| | | 38,081,995 | | *ORMDL3* | 3.654 | 3.74E-04 | 2.01E-04 |
| | | 38,082,022 | | *ORMDL3* | 4.899 | 2.82E-06 | 6.40E-04 |
| | | 38,082,035 | | *ORMDL3* | 3.827 | 2.00E-04 | 6.40E-04 |
| | | 38,082,073 | | *ORMDL3* | 4.776 | 4.74E-06 | 6.40E-04 |
| | | 38,082,658 | | *ORMDL3* | 4.586 | 1.05E-05 | 0.011 |
| | | 38,108,710 | | | -8.180 | 2.24E-13 | 0.039 |
| | | 38,119,071 | *GSDMA* | | 4.307 | 3.23E-05 | 0.009 |
| | | 38,119,198 | *GSDMA* | | 7.195 | 4.44E-11 | 0.009 |
| | | 38,119,207 | *GSDMA* | | 7.841 | 1.42E-12 | 0.009 |
| | | 38,119,649 | | *GSDMA* | 6.023 | 1.64E-08 | 0.002 |
| | | 38,119,760 | | *GSDMA* | 5.633 | 1.05E-07 | 0.011 |
| | | 38,120,370 | | *GSDMA* | 3.467 | 7.14E-04 | 0.011 |
| | | 38,128,699 | | *GSDMA* | -4.379 | 2.43E-05 | 0.006 |
| rs4134501 | g.38113290.G>A | 37,894,590 | | | 3.802 | 2.20E-04 | 0.013 |
| | | 37,910,697 | | | 3.602 | 4.49E-04 | 0.035 |
| | | 38,077,464 | | *ORMDL3* | 5.179 | 8.27E-07 | 0.014 |
| | | 38,080,829 | | *ORMDL3* | 3.438 | 7.89E-04 | 0.014 |
| | | 38,081,995 | | *ORMDL3* | 3.654 | 3.74E-04 | 2.01E-04 |
| | | 38,082,022 | | *ORMDL3* | 4.899 | 2.82E-06 | 6.40E-04 |
| | | 38,082,035 | | *ORMDL3* | 3.827 | 2.00E-04 | 6.40E-04 |
| | | 38,082,073 | | *ORMDL3* | 4.776 | 4.74E-06 | 6.40E-04 |
| | | 38,082,658 | | *ORMDL3* | 4.586 | 1.05E-05 | 0.011 |
| | | 38,108,710 | | | -8.180 | 2.24E-13 | 0.039 |
| | | 38,119,071 | *GSDMA* | | 4.307 | 3.23E-05 | 0.009 |
| | | 38,119,198 | *GSDMA* | | 7.195 | 4.44E-11 | 0.009 |
| | | 38,119,207 | *GSDMA* | | 7.841 | 1.42E-12 | 0.009 |
| | | 38,119,649 | | *GSDMA* | 6.023 | 1.64E-08 | 0.002 |
| | | 38,119,760 | | *GSDMA* | 5.633 | 1.05E-07 | 0.011 |
| | | 38,120,370 | | *GSDMA* | 3.467 | 7.14E-04 | 0.011 |
| | | 38,128,699 | | *GSDMA* | -4.379 | 2.43E-05 | 0.006 |

| | | | | | | | |
|---|---|---|---|---|---|---|---|
| rs201351591 | g.38113956_38113956insG | 37,828,195 | | *PGAP3* | 3.604 | 4.46E-04 | |
| | | 37,976,456 | | *IKZF3* | -4.920 | 2.57E-06 | 0.039 |
| | | 38,077,277 | | | -3.827 | 2.01E-04 | 0.014 |
| | | 38,084,119 | *ORMDL3* | | 3.490 | 6.61E-04 | 0.005 |
| rs11399056 | g.38114046_38114046insA | 37,887,061 | *MIEN1* | | 3.596 | 4.58E-04 | 0.002 |
| | | 37,894,590 | | | 3.490 | 6.60E-04 | 0.013 |
| | | 37,910,697 | | | 3.874 | 1.69E-04 | 0.035 |
| | | 38,077,464 | | *ORMDL3* | 5.344 | 3.96E-07 | 0.014 |
| | | 38,082,022 | | *ORMDL3* | 4.385 | 2.37E-05 | 6.40E-04 |
| | | 38,082,035 | | *ORMDL3* | 3.626 | 4.12E-04 | 6.40E-04 |
| | | 38,082,073 | | *ORMDL3* | 4.589 | 1.04E-05 | 6.40E-04 |
| | | 38,082,658 | | *ORMDL3* | 4.612 | 9.43E-06 | 0.011 |
| | | 38,108,710 | | | -7.685 | 3.29E-12 | 0.039 |
| | | 38,119,071 | *GSDMA* | | 4.112 | 6.92E-05 | 0.009 |
| | | 38,119,198 | *GSDMA* | | 6.853 | 2.62E-10 | 0.009 |
| | | 38,119,207 | *GSDMA* | | 7.273 | 2.94E-11 | 0.009 |
| | | 38,119,649 | | *GSDMA* | 5.794 | 4.91E-08 | 0.002 |
| | | 38,119,760 | | *GSDMA* | 5.968 | 2.14E-08 | 0.011 |
| | | 38,128,699 | | *GSDMA* | -4.047 | 8.84E-05 | 0.006 |
| rs7218742 | g.38114361.A>G | 37,894,590 | | | 3.766 | 2.50E-04 | 0.013 |
| | | 37,910,697 | | | 3.575 | 4.92E-04 | 0.035 |
| | | 38,077,464 | | *ORMDL3* | 5.290 | 5.04E-07 | 0.014 |
| | | 38,080,829 | | *ORMDL3* | 3.407 | 8.75E-04 | 0.014 |
| | | 38,081,995 | | *ORMDL3* | 3.637 | 3.97E-04 | 2.01E-04 |
| | | 38,082,022 | | *ORMDL3* | 4.853 | 3.43E-06 | 6.40E-04 |
| | | 38,082,035 | | *ORMDL3* | 3.788 | 2.31E-04 | 6.40E-04 |
| | | 38,082,073 | | *ORMDL3* | 4.745 | 5.41E-06 | 6.40E-04 |
| | | 38,082,658 | | *ORMDL3* | 4.634 | 8.61E-06 | 0.011 |
| | | 38,108,710 | | | -8.226 | 1.74E-13 | 0.039 |
| | | 38,119,071 | *GSDMA* | | 4.340 | 2.84E-05 | 0.009 |
| | | 38,119,198 | *GSDMA* | | 7.214 | 4.01E-11 | 0.009 |
| | | 38,119,207 | *GSDMA* | | 7.866 | 1.24E-12 | 0.009 |
| | | 38,119,649 | | *GSDMA* | 6.033 | 1.57E-08 | 0.002 |
| | | 38,119,760 | | *GSDMA* | 5.590 | 1.28E-07 | 0.011 |
| | | 38,120,234 | | *GSDMA* | 3.425 | 8.23E-04 | 0.011 |
| | | 38,120,370 | | *GSDMA* | 3.512 | 6.12E-04 | 0.011 |
| | | 38,128,699 | | *GSDMA* | -4.310 | 3.20E-05 | 0.006 |
| rs7218321 | g.38114469.C>T | 37,894,590 | | | 3.766 | 2.50E-04 | 0.013 |
| | | 37,910,697 | | | 3.575 | 4.92E-04 | 0.035 |

| | | | | | | | |
|---|---|---|---|---|---|---|---|
| | | 38,077,464 | | *ORMDL3* | 5.290 | 5.04E-07 | 0.014 |
| | | 38,080,829 | | *ORMDL3* | 3.407 | 8.75E-04 | 0.014 |
| | | 38,081,995 | | *ORMDL3* | 3.637 | 3.97E-04 | 2.01E-04 |
| | | 38,082,022 | | *ORMDL3* | 4.853 | 3.43E-06 | 6.40E-04 |
| | | 38,082,035 | | *ORMDL3* | 3.788 | 2.31E-04 | 6.40E-04 |
| | | 38,082,073 | | *ORMDL3* | 4.745 | 5.41E-06 | 6.40E-04 |
| | | 38,082,658 | | *ORMDL3* | 4.634 | 8.61E-06 | 0.011 |
| | | 38,108,710 | | | -8.226 | 1.74E-13 | 0.039 |
| | | 38,119,071 | *GSDMA* | | 4.340 | 2.84E-05 | 0.009 |
| | | 38,119,198 | *GSDMA* | | 7.214 | 4.01E-11 | 0.009 |
| | | 38,119,207 | *GSDMA* | | 7.866 | 1.24E-12 | 0.009 |
| | | 38,119,649 | | *GSDMA* | 6.033 | 1.57E-08 | 0.002 |
| | | 38,119,760 | | *GSDMA* | 5.590 | 1.28E-07 | 0.011 |
| | | 38,120,234 | | *GSDMA* | 3.425 | 8.23E-04 | 0.011 |
| | | 38,120,370 | | *GSDMA* | 3.512 | 6.12E-04 | 0.011 |
| | | 38,128,699 | | *GSDMA* | -4.310 | 3.20E-05 | 0.006 |
| rs7219080 | g.38114516.A>C | 37,894,590 | | | 3.766 | 2.50E-04 | 0.013 |
| | | 37,910,697 | | | 3.575 | 4.92E-04 | 0.035 |
| | | 38,077,464 | | *ORMDL3* | 5.290 | 5.04E-07 | 0.014 |
| | | 38,080,829 | | *ORMDL3* | 3.407 | 8.75E-04 | 0.014 |
| | | 38,081,995 | | *ORMDL3* | 3.637 | 3.97E-04 | 2.01E-04 |
| | | 38,082,022 | | *ORMDL3* | 4.853 | 3.43E-06 | 6.40E-04 |
| | | 38,082,035 | | *ORMDL3* | 3.788 | 2.31E-04 | 6.40E-04 |
| | | 38,082,073 | | *ORMDL3* | 4.745 | 5.41E-06 | 6.40E-04 |
| | | 38,082,658 | | *ORMDL3* | 4.634 | 8.61E-06 | 0.011 |
| | | 38,108,710 | | | -8.226 | 1.74E-13 | 0.039 |
| | | 38,119,071 | *GSDMA* | | 4.340 | 2.84E-05 | 0.009 |
| | | 38,119,198 | *GSDMA* | | 7.214 | 4.01E-11 | 0.009 |
| | | 38,119,207 | *GSDMA* | | 7.866 | 1.24E-12 | 0.009 |
| | | 38,119,649 | | *GSDMA* | 6.033 | 1.57E-08 | 0.002 |
| | | 38,119,760 | | *GSDMA* | 5.590 | 1.28E-07 | 0.011 |
| | | 38,120,234 | | *GSDMA* | 3.425 | 8.23E-04 | 0.011 |
| | | 38,120,370 | | *GSDMA* | 3.512 | 6.12E-04 | 0.011 |
| | | 38,128,699 | | *GSDMA* | -4.310 | 3.20E-05 | 0.006 |
| rs6503526 | g.38114598.T>C | 38,077,277 | | | 4.016 | 9.95E-05 | 0.014 |
| | | 38,077,464 | | *ORMDL3* | 3.725 | 2.90E-04 | 0.014 |
| | | 38,080,829 | | *ORMDL3* | 3.811 | 2.13E-04 | 0.014 |
| | | 38,081,530 | | *ORMDL3* | 3.547 | 5.43E-04 | 2.01E-04 |
| | | 38,081,812 | | *ORMDL3* | 3.985 | 1.12E-04 | 2.01E-04 |
| | | 38,082,022 | | *ORMDL3* | 3.787 | 2.32E-04 | 6.40E-04 |
| | | 38,082,073 | | *ORMDL3* | 4.605 | 9.70E-06 | 6.40E-04 |

| | | | | | | | |
|---|---|---|---|---|---|---|---|
| | | 38,082,658 | | *ORMDL3* | 4.081 | 7.77E-05 | 0.011 |
| | | 38,108,710 | | | -5.944 | 2.41E-08 | 0.039 |
| | | 38,119,071 | *GSDMA* | | 3.893 | 1.57E-04 | 0.009 |
| | | 38,119,198 | *GSDMA* | | 6.114 | 1.06E-08 | 0.009 |
| | | 38,119,207 | *GSDMA* | | 7.407 | 1.45E-11 | 0.009 |
| | | 38,119,649 | | *GSDMA* | 6.684 | 6.21E-10 | 0.002 |
| | | 38,119,760 | | *GSDMA* | 4.506 | 1.46E-05 | 0.011 |
| | | 38,128,699 | | *GSDMA* | -3.882 | 1.64E-04 | 0.006 |
| rs6503527 | g.38114719.G>A | 37,894,590 | | | 3.766 | 2.50E-04 | 0.013 |
| | | 37,910,697 | | | 3.575 | 4.92E-04 | 0.035 |
| | | 38,077,464 | | *ORMDL3* | 5.290 | 5.04E-07 | 0.014 |
| | | 38,080,829 | | *ORMDL3* | 3.407 | 8.75E-04 | 0.014 |
| | | 38,081,995 | | *ORMDL3* | 3.637 | 3.97E-04 | 2.01E-04 |
| | | 38,082,022 | | *ORMDL3* | 4.853 | 3.43E-06 | 6.40E-04 |
| | | 38,082,035 | | *ORMDL3* | 3.788 | 2.31E-04 | 6.40E-04 |
| | | 38,082,073 | | *ORMDL3* | 4.745 | 5.41E-06 | 6.40E-04 |
| | | 38,082,658 | | *ORMDL3* | 4.634 | 8.61E-06 | 0.011 |
| | | 38,108,710 | | | -8.226 | 1.74E-13 | 0.039 |
| | | 38,119,071 | *GSDMA* | | 4.340 | 2.84E-05 | 0.009 |
| | | 38,119,198 | *GSDMA* | | 7.214 | 4.01E-11 | 0.009 |
| | | 38,119,207 | *GSDMA* | | 7.866 | 1.24E-12 | 0.009 |
| | | 38,119,649 | | *GSDMA* | 6.033 | 1.57E-08 | 0.002 |
| | | 38,119,760 | | *GSDMA* | 5.590 | 1.28E-07 | 0.011 |
| | | 38,120,234 | | *GSDMA* | 3.425 | 8.23E-04 | 0.011 |
| | | 38,120,370 | | *GSDMA* | 3.512 | 6.12E-04 | 0.011 |
| | | 38,128,699 | | *GSDMA* | -4.310 | 3.20E-05 | 0.006 |
| rs3931960 | g.38114977.G>C | 37,910,697 | | | 3.818 | 2.08E-04 | 0.035 |
| | | 38,049,272 | | | 3.434 | 7.99E-04 | 0.046 |
| | | 38,077,277 | | | 3.521 | 5.93E-04 | 0.014 |
| | | 38,077,464 | | *ORMDL3* | 4.150 | 5.97E-05 | 0.014 |
| | | 38,080,829 | | *ORMDL3* | 4.211 | 4.71E-05 | 0.014 |
| | | 38,082,022 | | *ORMDL3* | 3.675 | 3.46E-04 | 6.40E-04 |
| | | 38,082,073 | | *ORMDL3* | 4.960 | 2.17E-06 | 6.40E-04 |
| | | 38,082,346 | | *ORMDL3* | 3.441 | 7.81E-04 | 0.011 |
| | | 38,082,658 | | *ORMDL3* | 4.101 | 7.20E-05 | 0.011 |
| | | 38,108,710 | | | -6.715 | 5.30E-10 | 0.039 |
| | | 38,119,071 | *GSDMA* | | 3.416 | 8.50E-04 | 0.009 |
| | | 38,119,198 | *GSDMA* | | 5.729 | 6.68E-08 | 0.009 |
| | | 38,119,207 | *GSDMA* | | 6.977 | 1.38E-10 | 0.009 |
| | | 38,119,649 | | *GSDMA* | 6.069 | 1.31E-08 | 0.002 |
| | | 38,119,760 | | *GSDMA* | 5.659 | 9.25E-08 | 0.011 |

| | | | | | | | |
|---|---|---|---|---|---|---|---|
| | | 38,128,699 | | *GSDMA* | -3.994 | 1.08E-04 | 0.006 |
| rs55913526 | g.38115299.T>C | 37,886,803 | *MIEN1* | | 3.625 | 4.14E-04 | 0.002 |
| rs140941705 | g.38115333_38115333ins A | 37,879,672 | | *ERBB2* | 3.542 | 5.52E-04 | 0.002 |
| rs7223717 | g.38115333.C>A | 37,910,697 | | | 3.818 | 2.08E-04 | 0.035 |
| | | 37,911,661 | | | -3.439 | 7.86E-04 | 0.035 |
| | | 38,049,272 | | | 3.434 | 7.99E-04 | 0.046 |
| | | 38,077,277 | | | 3.521 | 5.93E-04 | 0.014 |
| | | 38,077,464 | | *ORMDL3* | 4.150 | 5.97E-05 | 0.014 |
| | | 38,080,829 | | *ORMDL3* | 4.211 | 4.71E-05 | 0.014 |
| | | 38,082,022 | | *ORMDL3* | 3.675 | 3.46E-04 | 6.40E-04 |
| | | 38,082,073 | | *ORMDL3* | 4.960 | 2.17E-06 | 6.40E-04 |
| | | 38,082,346 | | *ORMDL3* | 3.441 | 7.81E-04 | 0.011 |
| | | 38,082,658 | | *ORMDL3* | 4.101 | 7.20E-05 | 0.011 |
| | | 38,083,874 | | *ORMDL3* | 3.391 | 9.23E-04 | 0.011 |
| | | 38,096,333 | | | -6.049 | 1.45E-08 | 0.033 |
| | | 38,096,571 | | | -10.722 | 1.32E-19 | 0.033 |
| | | 38,096,578 | | | -10.218 | 2.36E-18 | 0.033 |
| | | 38,096,648 | | | -4.123 | 6.61E-05 | 0.033 |
| | | 38,108,710 | | | -6.715 | 5.30E-10 | 0.039 |
| | | 38,119,071 | *GSDMA* | | 3.416 | 8.50E-04 | 0.009 |
| | | 38,119,198 | *GSDMA* | | 5.729 | 6.68E-08 | 0.009 |
| | | 38,119,207 | *GSDMA* | | 6.977 | 1.38E-10 | 0.009 |
| | | 38,119,649 | | *GSDMA* | 6.069 | 1.31E-08 | 0.002 |
| | | 38,119,760 | | *GSDMA* | 5.659 | 9.25E-08 | 0.011 |
| | | 38,128,699 | | *GSDMA* | -3.994 | 1.08E-04 | 0.006 |
| rs10693935 | g.38115429_38115429ins TG | 37,910,697 | | | 3.818 | 2.08E-04 | 0.035 |
| | | 38,049,272 | | | 3.434 | 7.99E-04 | 0.046 |
| | | 38,077,277 | | | 3.521 | 5.93E-04 | 0.014 |
| | | 38,077,464 | | *ORMDL3* | 4.150 | 5.97E-05 | 0.014 |
| | | 38,080,829 | | *ORMDL3* | 4.211 | 4.71E-05 | 0.014 |
| | | 38,082,022 | | *ORMDL3* | 3.675 | 3.46E-04 | 6.40E-04 |
| | | 38,082,073 | | *ORMDL3* | 4.960 | 2.17E-06 | 6.40E-04 |
| | | 38,082,346 | | *ORMDL3* | 3.441 | 7.81E-04 | 0.011 |
| | | 38,082,658 | | *ORMDL3* | 4.101 | 7.20E-05 | 0.011 |
| | | 38,108,710 | | | -6.715 | 5.30E-10 | 0.039 |
| | | 38,119,071 | *GSDMA* | | 3.416 | 8.50E-04 | 0.009 |
| | | 38,119,198 | *GSDMA* | | 5.729 | 6.68E-08 | 0.009 |
| | | 38,119,207 | *GSDMA* | | 6.977 | 1.38E-10 | 0.009 |
| | | 38,119,649 | | *GSDMA* | 6.069 | 1.31E-08 | 0.002 |

| | | | | | | | |
|---|---|---|---|---|---|---|---|
| | | 38,119,760 | | *GSDMA* | 5.659 | 9.25E-08 | 0.011 |
| | | 38,128,699 | | *GSDMA* | -3.994 | 1.08E-04 | 0.006 |
| rs56340811 | g.38115430.A>C | 37,910,697 | | | 3.818 | 2.08E-04 | 0.035 |
| | | 38,049,272 | | | 3.434 | 7.99E-04 | 0.046 |
| | | 38,077,277 | | | 3.521 | 5.93E-04 | 0.014 |
| | | 38,077,464 | | *ORMDL3* | 4.150 | 5.97E-05 | 0.014 |
| | | 38,080,829 | | *ORMDL3* | 4.211 | 4.71E-05 | 0.014 |
| | | 38,082,022 | | *ORMDL3* | 3.675 | 3.46E-04 | 6.40E-04 |
| | | 38,082,073 | | *ORMDL3* | 4.960 | 2.17E-06 | 6.40E-04 |
| | | 38,082,346 | | *ORMDL3* | 3.441 | 7.81E-04 | 0.011 |
| | | 38,082,658 | | *ORMDL3* | 4.101 | 7.20E-05 | 0.011 |
| | | 38,108,710 | | | -6.715 | 5.30E-10 | 0.039 |
| | | 38,119,071 | *GSDMA* | | 3.416 | 8.50E-04 | 0.009 |
| | | 38,119,198 | *GSDMA* | | 5.729 | 6.68E-08 | 0.009 |
| | | 38,119,207 | *GSDMA* | | 6.977 | 1.38E-10 | 0.009 |
| | | 38,119,649 | | *GSDMA* | 6.069 | 1.31E-08 | 0.002 |
| | | 38,119,760 | | *GSDMA* | 5.659 | 9.25E-08 | 0.011 |
| | | 38,128,699 | | *GSDMA* | -3.994 | 1.08E-04 | 0.006 |
| rs112979959 | g.38115873.G>T | 37,879,672 | | *ERBB2* | 3.542 | 5.52E-04 | 0.002 |
| | | 37,911,661 | | | -3.439 | 7.86E-04 | 0.035 |
| | | 38,083,874 | | *ORMDL3* | 3.391 | 9.23E-04 | 0.011 |
| | | 38,096,333 | | | -6.049 | 1.45E-08 | 0.033 |
| | | 38,096,571 | | | -10.722 | 1.32E-19 | 0.033 |
| | | 38,096,578 | | | -10.218 | 2.36E-18 | 0.033 |
| | | 38,096,648 | | | -4.123 | 6.61E-05 | 0.033 |
| rs112570995 | g.38116212.A>C | 37,879,672 | | *ERBB2* | 3.542 | 5.52E-04 | 0.002 |
| | | 37,911,661 | | | -3.439 | 7.86E-04 | 0.035 |
| | | 38,083,874 | | *ORMDL3* | 3.391 | 9.23E-04 | 0.011 |
| | | 38,096,333 | | | -6.049 | 1.45E-08 | 0.033 |
| | | 38,096,571 | | | -10.722 | 1.32E-19 | 0.033 |
| | | 38,096,578 | | | -10.218 | 2.36E-18 | 0.033 |
| | | 38,096,648 | | | -4.123 | 6.61E-05 | 0.033 |
| rs111522131 | g.38116225.G>A | 37,879,672 | | *ERBB2* | 3.542 | 5.52E-04 | 0.002 |
| | | 37,911,661 | | | -3.439 | 7.86E-04 | 0.035 |
| | | 38,083,874 | | *ORMDL3* | 3.391 | 9.23E-04 | 0.011 |
| | | 38,096,333 | | | -6.049 | 1.45E-08 | 0.033 |
| | | 38,096,571 | | | -10.722 | 1.32E-19 | 0.033 |
| | | 38,096,578 | | | -10.218 | 2.36E-18 | 0.033 |
| | | 38,096,648 | | | -4.123 | 6.61E-05 | 0.033 |
| rs11424020 | g.38116339_38116339insT | 37,894,590 | | | 3.968 | 1.19E-04 | 0.013 |

| | | | | | | | |
|---|---|---|---|---|---|---|---|
| | | 38,077,464 | | *ORMDL3* | 4.101 | 7.21E-05 | 0.014 |
| | | 38,080,829 | | *ORMDL3* | 3.668 | 3.55E-04 | 0.014 |
| | | 38,081,995 | | *ORMDL3* | 3.370 | 9.91E-04 | 2.01E-04 |
| | | 38,082,022 | | *ORMDL3* | 4.959 | 2.17E-06 | 6.40E-04 |
| | | 38,082,035 | | *ORMDL3* | 3.796 | 2.24E-04 | 6.40E-04 |
| | | 38,082,073 | | *ORMDL3* | 4.268 | 3.77E-05 | 6.40E-04 |
| | | 38,082,346 | | *ORMDL3* | 3.427 | 8.16E-04 | 0.011 |
| | | 38,082,658 | | *ORMDL3* | 4.485 | 1.59E-05 | 0.011 |
| | | 38,108,710 | | | -7.540 | 7.15E-12 | 0.039 |
| | | 38,119,071 | *GSDMA* | | 4.067 | 8.19E-05 | 0.009 |
| | | 38,119,198 | *GSDMA* | | 6.857 | 2.57E-10 | 0.009 |
| | | 38,119,207 | *GSDMA* | | 7.594 | 5.37E-12 | 0.009 |
| | | 38,119,649 | | *GSDMA* | 5.804 | 4.69E-08 | 0.002 |
| | | 38,119,760 | | *GSDMA* | 4.681 | 7.07E-06 | 0.011 |
| | | 38,120,234 | | *GSDMA* | 3.505 | 6.27E-04 | 0.011 |
| | | 38,120,370 | | *GSDMA* | 3.595 | 4.60E-04 | 0.011 |
| | | 38,122,024 | | *GSDMA* | -3.814 | 2.10E-04 | 0.011 |
| | | 38,128,699 | | *GSDMA* | -3.672 | 3.51E-04 | 0.006 |
| rs11869855 | g.38117653.A>G | 37,894,590 | | | 3.939 | 1.33E-04 | 0.013 |
| | | 38,077,464 | | *ORMDL3* | 4.176 | 5.41E-05 | 0.014 |
| | | 38,080,829 | | *ORMDL3* | 3.671 | 3.52E-04 | 0.014 |
| | | 38,082,022 | | *ORMDL3* | 4.789 | 4.50E-06 | 6.40E-04 |
| | | 38,082,035 | | *ORMDL3* | 3.737 | 2.78E-04 | 6.40E-04 |
| | | 38,082,073 | | *ORMDL3* | 4.071 | 8.07E-05 | 6.40E-04 |
| | | 38,082,658 | | *ORMDL3* | 4.285 | 3.53E-05 | 0.011 |
| | | 38,108,710 | | | -7.578 | 5.84E-12 | 0.039 |
| | | 38,108,850 | | | -3.379 | 9.61E-04 | 0.036 |
| | | 38,119,071 | *GSDMA* | | 3.967 | 1.20E-04 | 0.009 |
| | | 38,119,198 | *GSDMA* | | 6.843 | 2.75E-10 | 0.009 |
| | | 38,119,207 | *GSDMA* | | 7.599 | 5.22E-12 | 0.009 |
| | | 38,119,649 | | *GSDMA* | 5.731 | 6.63E-08 | 0.002 |
| | | 38,119,760 | | *GSDMA* | 4.739 | 5.55E-06 | 0.011 |
| | | 38,120,234 | | *GSDMA* | 3.425 | 8.24E-04 | 0.011 |
| | | 38,120,370 | | *GSDMA* | 3.577 | 4.89E-04 | 0.011 |
| | | 38,122,024 | | *GSDMA* | -3.898 | 1.55E-04 | 0.011 |
| | | 38,128,699 | | *GSDMA* | -3.745 | 2.70E-04 | 0.006 |
| rs113510790 | g.38118010.T>C | 37,879,672 | | *ERBB2* | 3.584 | 4.78E-04 | 0.002 |
| | | 37,911,661 | | | -3.412 | 8.60E-04 | 0.035 |
| | | 38,096,333 | | | -5.891 | 3.09E-08 | 0.033 |
| | | 38,096,571 | | | -10.724 | 1.30E-19 | 0.033 |
| | | 38,096,578 | | | -10.239 | 2.10E-18 | 0.033 |

| | | | | | | | |
|---|---|---|---|---|---|---|---|
| | | 38,096,648 | | | -4.134 | 6.34E-05 | 0.033 |
| rs564455927 | g.38118235_38118235ins A | 37,910,697 | | | 3.545 | 5.47E-04 | 0.035 |
| | | 38,077,277 | | | 4.209 | 4.75E-05 | 0.014 |
| | | 38,077,464 | | *ORMDL3* | 4.131 | 6.43E-05 | 0.014 |
| | | 38,080,829 | | *ORMDL3* | 3.701 | 3.16E-04 | 0.014 |
| | | 38,081,530 | | *ORMDL3* | 3.763 | 2.53E-04 | 2.01E-04 |
| | | 38,081,812 | | *ORMDL3* | 4.093 | 7.43E-05 | 2.01E-04 |
| | | 38,082,022 | | *ORMDL3* | 3.463 | 7.23E-04 | 6.40E-04 |
| | | 38,082,073 | | *ORMDL3* | 4.710 | 6.26E-06 | 6.40E-04 |
| | | 38,082,658 | | *ORMDL3* | 4.296 | 3.38E-05 | 0.011 |
| | | 38,108,710 | | | -6.021 | 1.66E-08 | 0.039 |
| | | 38,119,071 | *GSDMA* | | 4.270 | 3.74E-05 | 0.009 |
| | | 38,119,198 | *GSDMA* | | 6.144 | 9.15E-09 | 0.009 |
| | | 38,119,207 | *GSDMA* | | 7.345 | 2.02E-11 | 0.009 |
| | | 38,119,649 | | *GSDMA* | 6.358 | 3.18E-09 | 0.002 |
| | | 38,119,760 | | *GSDMA* | 4.644 | 8.25E-06 | 0.011 |
| | | 38,128,699 | | *GSDMA* | -4.147 | 6.05E-05 | 0.006 |
| rs72832987 | g.38118468.C>A | 37,879,672 | | *ERBB2* | 3.542 | 5.52E-04 | 0.002 |
| | | 37,911,661 | | | -3.439 | 7.86E-04 | 0.035 |
| | | 38,083,874 | | *ORMDL3* | 3.391 | 9.23E-04 | 0.011 |
| | | 38,096,333 | | | -6.049 | 1.45E-08 | 0.033 |
| | | 38,096,571 | | | -10.722 | 1.32E-19 | 0.033 |
| | | 38,096,578 | | | -10.218 | 2.36E-18 | 0.033 |
| | | 38,096,648 | | | -4.123 | 6.61E-05 | 0.033 |
| rs72832988 | g.38118541.A>G | 37,879,672 | | *ERBB2* | 3.542 | 5.52E-04 | 0.002 |
| | | 37,911,661 | | | -3.439 | 7.86E-04 | 0.035 |
| | | 38,083,874 | | *ORMDL3* | 3.391 | 9.23E-04 | 0.011 |
| | | 38,096,333 | | | -6.049 | 1.45E-08 | 0.033 |
| | | 38,096,571 | | | -10.722 | 1.32E-19 | 0.033 |
| | | 38,096,578 | | | -10.218 | 2.36E-18 | 0.033 |
| | | 38,096,648 | | | -4.123 | 6.61E-05 | 0.033 |
| rs72832989 | g.38118752.T>C | 37,879,672 | | *ERBB2* | 3.542 | 5.52E-04 | 0.002 |
| | | 37,911,661 | | | -3.439 | 7.86E-04 | 0.035 |
| | | 38,083,874 | | *ORMDL3* | 3.391 | 9.23E-04 | 0.011 |
| | | 38,096,333 | | | -6.049 | 1.45E-08 | 0.033 |
| | | 38,096,571 | | | -10.722 | 1.32E-19 | 0.033 |
| | | 38,096,578 | | | -10.218 | 2.36E-18 | 0.033 |
| | | 38,096,648 | | | -4.123 | 6.61E-05 | 0.033 |
| rs3902025 | g.38119254.C>A | 37,834,869 | | *PGAP3* | 3.604 | 4.44E-04 | 0.020 |
| | | 37,894,590 | | | 4.056 | 8.54E-05 | 0.013 |

| | | | | | | | |
|---|---|---|---|---|---|---|---|
| | | 38,077,051 | | | 3.434 | 7.98E-04 | 0.034 |
| | | 38,077,277 | | | 3.641 | 3.91E-04 | 0.014 |
| | | 38,077,464 | | *ORMDL3* | 5.023 | 1.64E-06 | 0.014 |
| | | 38,081,812 | | *ORMDL3* | 3.918 | 1.43E-04 | 2.01E-04 |
| | | 38,081,829 | | *ORMDL3* | 3.881 | 1.65E-04 | 2.01E-04 |
| | | 38,081,995 | | *ORMDL3* | 3.650 | 3.79E-04 | 2.01E-04 |
| | | 38,082,022 | | *ORMDL3* | 4.631 | 8.71E-06 | 6.40E-04 |
| | | 38,082,035 | | *ORMDL3* | 3.680 | 3.41E-04 | 6.40E-04 |
| | | 38,082,073 | | *ORMDL3* | 4.571 | 1.11E-05 | 6.40E-04 |
| | | 38,082,658 | | *ORMDL3* | 5.018 | 1.68E-06 | 0.011 |
| | | 38,108,710 | | | -7.171 | 5.04E-11 | 0.039 |
| | | 38,119,071 | *GSDMA* | | 5.222 | 6.85E-07 | 0.009 |
| | | 38,119,132 | *GSDMA* | | 3.427 | 8.17E-04 | 0.009 |
| | | 38,119,198 | *GSDMA* | | 7.601 | 5.15E-12 | 0.009 |
| | | 38,119,207 | *GSDMA* | | 8.226 | 1.75E-13 | 0.009 |
| | | 38,119,649 | | *GSDMA* | 6.377 | 2.90E-09 | 0.002 |
| | | 38,119,760 | | *GSDMA* | 4.401 | 2.22E-05 | 0.011 |
| | | 38,120,234 | | *GSDMA* | 3.524 | 5.87E-04 | 0.011 |
| | | 38,120,370 | | *GSDMA* | 4.123 | 6.62E-05 | 0.011 |
| | | 38,128,699 | | *GSDMA* | -4.291 | 3.45E-05 | 0.006 |
| rs72832991 | g.38119448.T>C | 37,879,672 | | *ERBB2* | 3.542 | 5.52E-04 | 0.002 |
| | | 37,911,661 | | | -3.439 | 7.86E-04 | 0.035 |
| | | 38,083,874 | | *ORMDL3* | 3.391 | 9.23E-04 | 0.011 |
| | | 38,096,333 | | | -6.049 | 1.45E-08 | 0.033 |
| | | 38,096,571 | | | -10.722 | 1.32E-19 | 0.033 |
| | | 38,096,578 | | | -10.218 | 2.36E-18 | 0.033 |
| | | 38,096,648 | | | -4.123 | 6.61E-05 | 0.033 |
| rs3902024 | g.38119548.T>C | 37,894,590 | | | 3.939 | 1.33E-04 | 0.013 |
| | | 38,077,464 | | *ORMDL3* | 4.176 | 5.41E-05 | 0.014 |
| | | 38,080,829 | | *ORMDL3* | 3.671 | 3.52E-04 | 0.014 |
| | | 38,082,022 | | *ORMDL3* | 4.789 | 4.50E-06 | 6.40E-04 |
| | | 38,082,035 | | *ORMDL3* | 3.737 | 2.78E-04 | 6.40E-04 |
| | | 38,082,073 | | *ORMDL3* | 4.071 | 8.07E-05 | 6.40E-04 |
| | | 38,082,658 | | *ORMDL3* | 4.285 | 3.53E-05 | 0.011 |
| | | 38,108,710 | | | -7.578 | 5.84E-12 | 0.039 |
| | | 38,108,850 | | | -3.379 | 9.61E-04 | 0.036 |
| | | 38,119,071 | *GSDMA* | | 3.967 | 1.20E-04 | 0.009 |
| | | 38,119,198 | *GSDMA* | | 6.843 | 2.75E-10 | 0.009 |
| | | 38,119,207 | *GSDMA* | | 7.599 | 5.22E-12 | 0.009 |
| | | 38,119,649 | | *GSDMA* | 5.731 | 6.63E-08 | 0.002 |
| | | 38,119,760 | | *GSDMA* | 4.739 | 5.55E-06 | 0.011 |

| | | | | | | | |
|---|---|---|---|---|---|---|---|
| | | 38,120,234 | | *GSDMA* | 3.425 | 8.24E-04 | 0.011 |
| | | 38,120,370 | | *GSDMA* | 3.577 | 4.89E-04 | 0.011 |
| | | 38,122,024 | | *GSDMA* | -3.898 | 1.55E-04 | 0.011 |
| | | 38,128,699 | | *GSDMA* | -3.745 | 2.70E-04 | 0.006 |
| rs55739615 | g.38119638.C>T | 37,834,869 | | *PGAP3* | 3.604 | 4.44E-04 | 0.020 |
| | | 37,894,590 | | | 4.056 | 8.54E-05 | 0.013 |
| | | 38,077,051 | | | 3.434 | 7.98E-04 | 0.034 |
| | | 38,077,277 | | | 3.641 | 3.91E-04 | 0.014 |
| | | 38,077,464 | | *ORMDL3* | 5.023 | 1.64E-06 | 0.014 |
| | | 38,081,812 | | *ORMDL3* | 3.918 | 1.43E-04 | 2.01E-04 |
| | | 38,081,829 | | *ORMDL3* | 3.881 | 1.65E-04 | 2.01E-04 |
| | | 38,081,995 | | *ORMDL3* | 3.650 | 3.79E-04 | 2.01E-04 |
| | | 38,082,022 | | *ORMDL3* | 4.631 | 8.71E-06 | 6.40E-04 |
| | | 38,082,035 | | *ORMDL3* | 3.680 | 3.41E-04 | 6.40E-04 |
| | | 38,082,073 | | *ORMDL3* | 4.571 | 1.11E-05 | 6.40E-04 |
| | | 38,082,658 | | *ORMDL3* | 5.018 | 1.68E-06 | 0.011 |
| | | 38,108,710 | | | -7.171 | 5.04E-11 | 0.039 |
| | | 38,119,071 | *GSDMA* | | 5.222 | 6.85E-07 | 0.009 |
| | | 38,119,132 | *GSDMA* | | 3.427 | 8.17E-04 | 0.009 |
| | | 38,119,198 | *GSDMA* | | 7.601 | 5.15E-12 | 0.009 |
| | | 38,119,207 | *GSDMA* | | 8.226 | 1.75E-13 | 0.009 |
| | | 38,119,649 | | *GSDMA* | 6.377 | 2.90E-09 | 0.002 |
| | | 38,119,760 | | *GSDMA* | 4.401 | 2.22E-05 | 0.011 |
| | | 38,120,234 | | *GSDMA* | 3.524 | 5.87E-04 | 0.011 |
| | | 38,120,370 | | *GSDMA* | 4.123 | 6.62E-05 | 0.011 |
| | | 38,128,699 | | *GSDMA* | -4.291 | 3.45E-05 | 0.006 |
| rs78295195 | g.38119663.G>C | 37,879,672 | | *ERBB2* | 3.542 | 5.52E-04 | 0.002 |
| | | 37,911,661 | | | -3.439 | 7.86E-04 | 0.035 |
| | | 38,083,874 | | *ORMDL3* | 3.391 | 9.23E-04 | 0.011 |
| | | 38,096,333 | | | -6.049 | 1.45E-08 | 0.033 |
| | | 38,096,571 | | | -10.722 | 1.32E-19 | 0.033 |
| | | 38,096,578 | | | -10.218 | 2.36E-18 | 0.033 |
| | | 38,096,648 | | | -4.123 | 6.61E-05 | 0.033 |
| rs56396280 | g.38119708.A>C | 37,894,590 | | | 3.939 | 1.33E-04 | 0.013 |
| | | 38,077,464 | | *ORMDL3* | 4.176 | 5.41E-05 | 0.014 |
| | | 38,080,829 | | *ORMDL3* | 3.671 | 3.52E-04 | 0.014 |
| | | 38,082,022 | | *ORMDL3* | 4.789 | 4.50E-06 | 6.40E-04 |
| | | 38,082,035 | | *ORMDL3* | 3.737 | 2.78E-04 | 6.40E-04 |
| | | 38,082,073 | | *ORMDL3* | 4.071 | 8.07E-05 | 6.40E-04 |
| | | 38,082,658 | | *ORMDL3* | 4.285 | 3.53E-05 | 0.011 |
| | | 38,108,710 | | | -7.578 | 5.84E-12 | 0.039 |

| | | | | | | | |
|---|---|---|---|---|---|---|---|
| | | 38,108,850 | | | -3.379 | 9.61E-04 | 0.036 |
| | | 38,119,071 | *GSDMA* | | 3.967 | 1.20E-04 | 0.009 |
| | | 38,119,198 | *GSDMA* | | 6.843 | 2.75E-10 | 0.009 |
| | | 38,119,207 | *GSDMA* | | 7.599 | 5.22E-12 | 0.009 |
| | | 38,119,649 | | *GSDMA* | 5.731 | 6.63E-08 | 0.002 |
| | | 38,119,760 | | *GSDMA* | 4.739 | 5.55E-06 | 0.011 |
| | | 38,120,234 | | *GSDMA* | 3.425 | 8.24E-04 | 0.011 |
| | | 38,120,370 | | *GSDMA* | 3.577 | 4.89E-04 | 0.011 |
| | | 38,122,024 | | *GSDMA* | -3.898 | 1.55E-04 | 0.011 |
| | | 38,128,699 | | *GSDMA* | -3.745 | 2.70E-04 | 0.006 |
| rs12451084 | g.38119757.T>G | 37,834,869 | | *PGAP3* | 3.453 | 7.49E-04 | 0.020 |
| | | 37,910,697 | | | 3.491 | 6.58E-04 | 0.035 |
| | | 38,077,277 | | | 4.203 | 4.86E-05 | 0.014 |
| | | 38,077,464 | | *ORMDL3* | 4.071 | 8.07E-05 | 0.014 |
| | | 38,080,829 | | *ORMDL3* | 3.663 | 3.61E-04 | 0.014 |
| | | 38,081,530 | | *ORMDL3* | 3.708 | 3.08E-04 | 2.01E-04 |
| | | 38,081,812 | | *ORMDL3* | 4.038 | 9.16E-05 | 2.01E-04 |
| | | 38,082,022 | | *ORMDL3* | 3.625 | 4.13E-04 | 6.40E-04 |
| | | 38,082,073 | | *ORMDL3* | 4.886 | 2.98E-06 | 6.40E-04 |
| | | 38,082,346 | | *ORMDL3* | 3.433 | 8.01E-04 | 0.011 |
| | | 38,082,658 | | *ORMDL3* | 4.551 | 1.21E-05 | 0.011 |
| | | 38,108,710 | | | -6.111 | 1.07E-08 | 0.039 |
| | | 38,119,071 | *GSDMA* | | 4.281 | 3.59E-05 | 0.009 |
| | | 38,119,198 | *GSDMA* | | 6.240 | 5.72E-09 | 0.009 |
| | | 38,119,207 | *GSDMA* | | 7.498 | 8.96E-12 | 0.009 |
| | | 38,119,649 | | *GSDMA* | 6.524 | 1.39E-09 | 0.002 |
| | | 38,119,760 | | *GSDMA* | 4.665 | 7.58E-06 | 0.011 |
| | | 38,120,234 | | *GSDMA* | 3.402 | 8.90E-04 | 0.011 |
| | | 38,128,699 | | *GSDMA* | -4.073 | 8.01E-05 | 0.006 |
| rs12451100 | g.38119831.T>G | 37,834,869 | | *PGAP3* | 3.453 | 7.49E-04 | 0.020 |
| | | 37,910,697 | | | 3.491 | 6.58E-04 | 0.035 |
| | | 38,077,277 | | | 4.203 | 4.86E-05 | 0.014 |
| | | 38,077,464 | | *ORMDL3* | 4.071 | 8.07E-05 | 0.014 |
| | | 38,080,829 | | *ORMDL3* | 3.663 | 3.61E-04 | 0.014 |
| | | 38,081,530 | | *ORMDL3* | 3.708 | 3.08E-04 | 2.01E-04 |
| | | 38,081,812 | | *ORMDL3* | 4.038 | 9.16E-05 | 2.01E-04 |
| | | 38,082,022 | | *ORMDL3* | 3.625 | 4.13E-04 | 6.40E-04 |
| | | 38,082,073 | | *ORMDL3* | 4.886 | 2.98E-06 | 6.40E-04 |
| | | 38,082,346 | | *ORMDL3* | 3.433 | 8.01E-04 | 0.011 |
| | | 38,082,658 | | *ORMDL3* | 4.551 | 1.21E-05 | 0.011 |
| | | 38,108,710 | | | -6.111 | 1.07E-08 | 0.039 |

| | | | | | | | |
|---|---|---|---|---|---|---|---|
| | | 38,119,071 | *GSDMA* | | 4.281 | 3.59E-05 | 0.009 |
| | | 38,119,198 | *GSDMA* | | 6.240 | 5.72E-09 | 0.009 |
| | | 38,119,207 | *GSDMA* | | 7.498 | 8.96E-12 | 0.009 |
| | | 38,119,649 | | *GSDMA* | 6.524 | 1.39E-09 | 0.002 |
| | | 38,119,760 | | *GSDMA* | 4.665 | 7.58E-06 | 0.011 |
| | | 38,120,234 | | *GSDMA* | 3.402 | 8.90E-04 | 0.011 |
| | | 38,128,699 | | *GSDMA* | -4.073 | 8.01E-05 | 0.006 |
| rs111690387 | g.38119882.A>G | 37,879,672 | | *ERBB2* | 3.542 | 5.52E-04 | 0.002 |
| | | 37,911,661 | | | -3.439 | 7.86E-04 | 0.035 |
| | | 38,083,874 | | *ORMDL3* | 3.391 | 9.23E-04 | 0.011 |
| | | 38,096,333 | | | -6.049 | 1.45E-08 | 0.033 |
| | | 38,096,571 | | | -10.722 | 1.32E-19 | 0.033 |
| | | 38,096,578 | | | -10.218 | 2.36E-18 | 0.033 |
| | | 38,096,648 | | | -4.123 | 6.61E-05 | 0.033 |
| rs60701125 | g.38120179.T>C | 37,879,672 | | *ERBB2* | 3.542 | 5.52E-04 | 0.002 |
| | | 37,911,661 | | | -3.439 | 7.86E-04 | 0.035 |
| | | 38,083,874 | | *ORMDL3* | 3.391 | 9.23E-04 | 0.011 |
| | | 38,096,333 | | | -6.049 | 1.45E-08 | 0.033 |
| | | 38,096,571 | | | -10.722 | 1.32E-19 | 0.033 |
| | | 38,096,578 | | | -10.218 | 2.36E-18 | 0.033 |
| | | 38,096,648 | | | -4.123 | 6.61E-05 | 0.033 |
| rs2001476 | g.38120604.T>C | 37,894,590 | | | 3.939 | 1.33E-04 | 0.013 |
| | | 38,077,464 | | *ORMDL3* | 4.176 | 5.41E-05 | 0.014 |
| | | 38,080,829 | | *ORMDL3* | 3.671 | 3.52E-04 | 0.014 |
| | | 38,082,022 | | *ORMDL3* | 4.789 | 4.50E-06 | 6.40E-04 |
| | | 38,082,035 | | *ORMDL3* | 3.737 | 2.78E-04 | 6.40E-04 |
| | | 38,082,073 | | *ORMDL3* | 4.071 | 8.07E-05 | 6.40E-04 |
| | | 38,082,658 | | *ORMDL3* | 4.285 | 3.53E-05 | 0.011 |
| | | 38,108,710 | | | -7.578 | 5.84E-12 | 0.039 |
| | | 38,108,850 | | | -3.379 | 9.61E-04 | 0.036 |
| | | 38,119,071 | *GSDMA* | | 3.967 | 1.20E-04 | 0.009 |
| | | 38,119,198 | *GSDMA* | | 6.843 | 2.75E-10 | 0.009 |
| | | 38,119,207 | *GSDMA* | | 7.599 | 5.22E-12 | 0.009 |
| | | 38,119,649 | | *GSDMA* | 5.731 | 6.63E-08 | 0.002 |
| | | 38,119,760 | | *GSDMA* | 4.739 | 5.55E-06 | 0.011 |
| | | 38,120,234 | | *GSDMA* | 3.425 | 8.24E-04 | 0.011 |
| | | 38,120,370 | | *GSDMA* | 3.577 | 4.89E-04 | 0.011 |
| | | 38,122,024 | | *GSDMA* | -3.898 | 1.55E-04 | 0.011 |
| | | 38,128,699 | | *GSDMA* | -3.745 | 2.70E-04 | 0.006 |
| rs4795409 | g.38120736.T>C | 37,834,869 | | *PGAP3* | 3.453 | 7.49E-04 | 0.020 |
| | | 37,910,697 | | | 3.491 | 6.58E-04 | 0.035 |

| | | | | | | | |
|---|---|---|---|---|---|---|---|
| | | 38,077,277 | | | 4.203 | 4.86E-05 | 0.014 |
| | | 38,077,464 | | *ORMDL3* | 4.071 | 8.07E-05 | 0.014 |
| | | 38,080,829 | | *ORMDL3* | 3.663 | 3.61E-04 | 0.014 |
| | | 38,081,530 | | *ORMDL3* | 3.708 | 3.08E-04 | 2.01E-04 |
| | | 38,081,812 | | *ORMDL3* | 4.038 | 9.16E-05 | 2.01E-04 |
| | | 38,082,022 | | *ORMDL3* | 3.625 | 4.13E-04 | 6.40E-04 |
| | | 38,082,073 | | *ORMDL3* | 4.886 | 2.98E-06 | 6.40E-04 |
| | | 38,082,346 | | *ORMDL3* | 3.433 | 8.01E-04 | 0.011 |
| | | 38,082,658 | | *ORMDL3* | 4.551 | 1.21E-05 | 0.011 |
| | | 38,108,710 | | | -6.111 | 1.07E-08 | 0.039 |
| | | 38,119,071 | *GSDMA* | | 4.281 | 3.59E-05 | 0.009 |
| | | 38,119,198 | *GSDMA* | | 6.240 | 5.72E-09 | 0.009 |
| | | 38,119,207 | *GSDMA* | | 7.498 | 8.96E-12 | 0.009 |
| | | 38,119,649 | | *GSDMA* | 6.524 | 1.39E-09 | 0.002 |
| | | 38,119,760 | | *GSDMA* | 4.665 | 7.58E-06 | 0.011 |
| | | 38,120,234 | | *GSDMA* | 3.402 | 8.90E-04 | 0.011 |
| | | 38,128,699 | | *GSDMA* | -4.073 | 8.01E-05 | 0.006 |
| rs142218801 | g.38120827_38120829del | 37,879,672 | | *ERBB2* | 3.542 | 5.52E-04 | 0.002 |
| | | 37,911,661 | | | -3.439 | 7.86E-04 | 0.035 |
| | | 38,083,874 | | *ORMDL3* | 3.391 | 9.23E-04 | 0.011 |
| | | 38,096,333 | | | -6.049 | 1.45E-08 | 0.033 |
| | | 38,096,571 | | | -10.722 | 1.32E-19 | 0.033 |
| | | 38,096,578 | | | -10.218 | 2.36E-18 | 0.033 |
| | | 38,096,648 | | | -4.123 | 6.61E-05 | 0.033 |
| rs4458030 | g.38121706.C>T | 37,894,590 | | | 3.991 | 1.09E-04 | 0.013 |
| | | 38,024,290 | | | -3.442 | 7.78E-04 | 0.019 |
| | | 38,077,464 | | *ORMDL3* | 4.199 | 4.93E-05 | 0.014 |
| | | 38,080,829 | | *ORMDL3* | 3.430 | 8.10E-04 | 0.014 |
| | | 38,082,022 | | *ORMDL3* | 4.517 | 1.39E-05 | 6.40E-04 |
| | | 38,082,073 | | *ORMDL3* | 4.099 | 7.27E-05 | 6.40E-04 |
| | | 38,082,658 | | *ORMDL3* | 3.973 | 1.17E-04 | 0.011 |
| | | 38,108,710 | | | -7.782 | 1.95E-12 | 0.039 |
| | | 38,119,071 | *GSDMA* | | 3.762 | 2.54E-04 | 0.009 |
| | | 38,119,198 | *GSDMA* | | 6.451 | 2.01E-09 | 0.009 |
| | | 38,119,207 | *GSDMA* | | 7.425 | 1.32E-11 | 0.009 |
| | | 38,119,649 | | *GSDMA* | 5.692 | 7.95E-08 | 0.002 |
| | | 38,119,760 | | *GSDMA* | 4.665 | 7.57E-06 | 0.011 |
| | | 38,120,370 | | *GSDMA* | 3.550 | 5.36E-04 | 0.011 |
| | | 38,122,024 | | *GSDMA* | -3.651 | 3.77E-04 | 0.011 |
| | | 38,128,699 | | *GSDMA* | -3.935 | 1.35E-04 | 0.006 |

| | | | | | | | | |
|---|---|---|---|---|---|---|---|---|
| | rs3894194 | g.38121993.T>C (Arg.18.Gln) | 37,894,590 | | | 3.372 | 9.84E-04 | 0.013 |
| | | | 38,077,277 | | | 3.570 | 5.02E-04 | 0.014 |
| | | | 38,077,464 | | *ORMDL3* | 3.609 | 4.37E-04 | 0.014 |
| | | | 38,081,530 | | *ORMDL3* | 3.918 | 1.43E-04 | 2.01E-04 |
| | | | 38,081,812 | | *ORMDL3* | 4.020 | 9.81E-05 | 2.01E-04 |
| | | | 38,082,022 | | *ORMDL3* | 3.868 | 1.73E-04 | 6.40E-04 |
| | | | 38,082,073 | | *ORMDL3* | 4.637 | 8.51E-06 | 6.40E-04 |
| | | | 38,082,658 | | *ORMDL3* | 3.590 | 4.67E-04 | 0.011 |
| | | | 38,108,710 | | | -6.429 | 2.24E-09 | 0.039 |
| | | | 38,119,071 | *GSDMA* | | 3.897 | 1.55E-04 | 0.009 |
| | | | 38,119,198 | *GSDMA* | | 5.788 | 5.07E-08 | 0.009 |
| | | | 38,119,207 | *GSDMA* | | 6.843 | 2.75E-10 | 0.009 |
| | | | 38,119,649 | | *GSDMA* | 6.508 | 1.51E-09 | 0.002 |
| | | | 38,119,760 | | *GSDMA* | 4.674 | 7.29E-06 | 0.011 |
| | | | 38,120,234 | | *GSDMA* | 3.805 | 2.18E-04 | 0.011 |
| | | | 38,128,699 | | *GSDMA* | -3.792 | 2.28E-04 | 0.006 |
| | rs8069202 | g.38122200.A>G | 37,894,590 | | | 3.372 | 9.84E-04 | 0.013 |
| | | | 38,077,277 | | | 3.570 | 5.02E-04 | 0.014 |
| | | | 38,077,464 | | *ORMDL3* | 3.609 | 4.37E-04 | 0.014 |
| | | | 38,081,530 | | *ORMDL3* | 3.918 | 1.43E-04 | 2.01E-04 |
| | | | 38,081,812 | | *ORMDL3* | 4.020 | 9.81E-05 | 2.01E-04 |
| | | | 38,082,022 | | *ORMDL3* | 3.868 | 1.73E-04 | 6.40E-04 |
| | | | 38,082,073 | | *ORMDL3* | 4.637 | 8.51E-06 | 6.40E-04 |
| | | | 38,082,658 | | *ORMDL3* | 3.590 | 4.67E-04 | 0.011 |
| | | | 38,108,710 | | | -6.429 | 2.24E-09 | 0.039 |
| | | | 38,119,071 | *GSDMA* | | 3.897 | 1.55E-04 | 0.009 |
| | | | 38,119,198 | *GSDMA* | | 5.788 | 5.07E-08 | 0.009 |
| | | | 38,119,207 | *GSDMA* | | 6.843 | 2.75E-10 | 0.009 |
| | | | 38,119,649 | | *GSDMA* | 6.508 | 1.51E-09 | 0.002 |
| | | | 38,119,760 | | *GSDMA* | 4.674 | 7.29E-06 | 0.011 |
| | | | 38,120,234 | | *GSDMA* | 3.805 | 2.18E-04 | 0.011 |
| | | | 38,128,699 | | *GSDMA* | -3.792 | 2.28E-04 | 0.006 |
| | rs4082190 | g.38125891.G>A | 38,077,464 | | *ORMDL3* | 3.910 | 1.48E-04 | 0.014 |
| | | | 38,096,571 | | | -4.991 | 1.89E-06 | 0.033 |
| | | | 38,096,578 | | | -4.278 | 3.63E-05 | 0.033 |
| Seventh block | rs4488488 | g.38126459.A>G | 38,077,464 | | *ORMDL3* | 3.910 | 1.48E-04 | 0.014 |
| | | | 38,096,571 | | | -4.991 | 1.89E-06 | 0.033 |
| | | | 38,096,578 | | | -4.278 | 3.63E-05 | 0.033 |
| | rs150060808 | g.38126911_38126919del | 37,910,697 | | | 3.490 | 6.60E-04 | 0.035 |

| | | | | | | | |
|---|---|---|---|---|---|---|---|
| rs556061928 | g.38126911_38126919del | 37,913,383 | | | 3.882 | 1.64E-04 | 0.033 |
| | | 38,024,331 | | | -3.396 | 9.07E-04 | 0.017 |
| | | 38,049,272 | | | 3.529 | 5.77E-04 | 0.046 |
| | | 38,081,530 | | *ORMDL3* | 4.240 | 4.21E-05 | 2.01E-04 |
| | | 38,081,812 | | *ORMDL3* | 3.471 | 7.05E-04 | 2.01E-04 |
| | | 38,082,073 | | *ORMDL3* | 4.301 | 3.31E-05 | 6.40E-04 |
| | | 38,082,448 | | *ORMDL3* | 3.424 | 8.26E-04 | 0.011 |
| | | 38,082,658 | | *ORMDL3* | 4.294 | 3.40E-05 | 0.011 |
| | | 38,108,710 | | | -4.778 | 4.70E-06 | 0.039 |
| | | 38,108,710 | | | -4.575 | 1.10E-05 | 0.039 |
| | | 38,119,198 | *GSDMA* | | 3.563 | 5.13E-04 | 0.009 |
| | | 38,119,207 | *GSDMA* | | 4.847 | 3.51E-06 | 0.009 |
| | | 38,119,649 | | *GSDMA* | 4.447 | 1.85E-05 | 0.002 |
| | | 38,119,760 | | *GSDMA* | 3.845 | 1.88E-04 | 0.011 |
| | | 38,128,699 | | *GSDMA* | -4.455 | 1.79E-05 | 0.006 |
| | | 38,128,699 | | *GSDMA* | -3.682 | 3.38E-04 | 0.006 |
| rs4239225 | g.38127112.T>C | 37,856,888 | | *ERBB2* | 3.370 | 9.90E-04 | 0.003 |
| | | 37,910,697 | | | 3.520 | 5.95E-04 | 0.035 |
| | | 38,077,323 | | *ORMDL3* | 3.539 | 5.59E-04 | 0.014 |
| | | 38,077,464 | | *ORMDL3* | 3.891 | 1.58E-04 | 0.014 |
| | | 38,082,022 | | *ORMDL3* | 3.743 | 2.72E-04 | 6.40E-04 |
| | | 38,082,073 | | *ORMDL3* | 4.218 | 4.58E-05 | 6.40E-04 |
| | | 38,082,658 | | *ORMDL3* | 3.713 | 3.03E-04 | 0.011 |
| | | 38,108,710 | | | -5.089 | 1.23E-06 | 0.039 |
| | | 38,119,198 | *GSDMA* | | 3.880 | 1.65E-04 | 0.009 |
| | | 38,119,207 | *GSDMA* | | 4.377 | 2.45E-05 | 0.009 |
| | | 38,119,649 | | *GSDMA* | 4.318 | 3.10E-05 | 0.002 |
| | | 38,119,760 | | *GSDMA* | 3.733 | 2.82E-04 | 0.011 |
| | | 38,128,699 | | *GSDMA* | -5.175 | 8.41E-07 | 0.006 |
| rs3916061 | g.38127713.A>G | 37,832,464 | | *PGAP3* | 3.501 | 6.35E-04 | 0.028 |
| | | 37,844,223 | *ERBB2* | *PGAP3* | 3.714 | 3.01E-04 | 7.82E-05 |
| | | 37,844,398 | *PGAP3* | *ERBB2* | 3.476 | 6.92E-04 | 7.82E-05 |
| | | 37,895,953 | | | -4.137 | 6.27E-05 | 0.027 |
| | | 37,915,921 | | | 3.754 | 2.62E-04 | 0.001 |
| | | 38,077,464 | | *ORMDL3* | 12.610 | 2.62E-24 | 0.014 |
| | | 38,084,148 | *ORMDL3* | | 3.624 | 4.15E-04 | 0.005 |
| | | 38,084,331 | *ORMDL3* | | 3.758 | 2.58E-04 | 0.005 |
| | | 38,084,450 | *ORMDL3* | | 3.673 | 3.49E-04 | 0.005 |
| | | 38,119,760 | | *GSDMA* | 3.634 | 4.01E-04 | 0.011 |

| | | | | | | | |
|---|---|---|---|---|---|---|---|
| rs74727658 | g.38128181_38128182del | 37,913,383 | | | 3.882 | 1.64E-04 | 0.033 |
| | | 38,024,331 | | | -3.396 | 9.07E-04 | 0.017 |
| | | 38,049,272 | | | 3.529 | 5.77E-04 | 0.046 |
| | | 38,108,710 | | | -4.778 | 4.70E-06 | 0.039 |
| | | 38,128,699 | | *GSDMA* | -3.682 | 3.38E-04 | 0.006 |
| rs5820314 | g.38128584_38128584insG | 37,844,223 | *ERBB2* | *PGAP3* | 3.633 | 4.02E-04 | 7.82E-05 |
| | | 37,844,398 | *PGAP3* | *ERBB2* | 3.372 | 9.83E-04 | 7.82E-05 |
| | | 38,077,464 | | *ORMDL3* | 7.076 | 8.24E-11 | 0.014 |
| | | 38,084,331 | *ORMDL3* | | 4.133 | 6.38E-05 | 0.005 |
| | | 38,084,404 | *ORMDL3* | | 3.514 | 6.07E-04 | 0.005 |
| | | 38,084,450 | *ORMDL3* | | 3.443 | 7.75E-04 | 0.005 |
| rs3859192 | g.38128648.T>C | 37,856,888 | | *ERBB2* | 3.369 | 9.92E-04 | 0.003 |
| | | 37,910,697 | | | 3.519 | 5.98E-04 | 0.035 |
| | | 38,077,323 | | *ORMDL3* | 3.537 | 5.61E-04 | 0.014 |
| | | 38,077,464 | | *ORMDL3* | 3.891 | 1.59E-04 | 0.014 |
| | | 38,082,022 | | *ORMDL3* | 3.737 | 2.78E-04 | 6.40E-04 |
| | | 38,082,073 | | *ORMDL3* | 4.215 | 4.64E-05 | 6.40E-04 |
| | | 38,082,658 | | *ORMDL3* | 3.712 | 3.04E-04 | 0.011 |
| | | 38,108,710 | | | -5.081 | 1.28E-06 | 0.039 |
| | | 38,119,198 | *GSDMA* | | 3.880 | 1.65E-04 | 0.009 |
| | | 38,119,207 | *GSDMA* | | 4.376 | 2.46E-05 | 0.009 |
| | | 38,119,649 | | *GSDMA* | 4.316 | 3.12E-05 | 0.002 |
| | | 38,119,760 | | *GSDMA* | 3.733 | 2.82E-04 | 0.011 |
| | | 38,128,699 | | *GSDMA* | -5.175 | 8.43E-07 | 0.006 |
| rs3859191 | g.38128714.A>G | 37,856,888 | | *ERBB2* | 3.369 | 9.92E-04 | 0.003 |
| | | 37,910,697 | | | 3.519 | 5.98E-04 | 0.035 |
| | | 38,077,323 | | *ORMDL3* | 3.537 | 5.61E-04 | 0.014 |
| | | 38,077,464 | | *ORMDL3* | 3.891 | 1.59E-04 | 0.014 |
| | | 38,082,022 | | *ORMDL3* | 3.737 | 2.78E-04 | 6.40E-04 |
| | | 38,082,073 | | *ORMDL3* | 4.215 | 4.64E-05 | 6.40E-04 |
| | | 38,082,658 | | *ORMDL3* | 3.712 | 3.04E-04 | 0.011 |
| | | 38,108,710 | | | -5.081 | 1.28E-06 | 0.039 |
| | | 38,119,198 | *GSDMA* | | 3.880 | 1.65E-04 | 0.009 |
| | | 38,119,207 | *GSDMA* | | 4.376 | 2.46E-05 | 0.009 |
| | | 38,119,649 | | *GSDMA* | 4.316 | 3.12E-05 | 0.002 |
| | | 38,119,760 | | *GSDMA* | 3.733 | 2.82E-04 | 0.011 |
| | | 38,128,699 | | *GSDMA* | -5.175 | 8.43E-07 | 0.006 |
| rs8077456 | g.38128765.C>G | 37,913,383 | | | 3.882 | 1.64E-04 | 0.033 |
| | | 38,024,331 | | | -3.396 | 9.07E-04 | 0.017 |

| | | | | | | | |
|---|---|---|---|---|---|---|---|
| | | 38,049,272 | | | 3.529 | 5.77E-04 | 0.046 |
| | | 38,108,710 | | | -4.778 | 4.70E-06 | 0.039 |
| | | 38,128,699 | | *GSDMA* | -3.682 | 3.38E-04 | 0.006 |
| rs10852937 | g.38129097.C>G | 37,844,223 | *ERBB2* | *PGAP3* | 3.637 | 3.97E-04 | 7.82E-05 |
| | | 38,077,464 | | *ORMDL3* | 7.060 | 8.98E-11 | 0.014 |
| | | 38,084,331 | *ORMDL3* | | 4.116 | 6.81E-05 | 0.005 |
| | | 38,084,404 | *ORMDL3* | | 3.518 | 6.00E-04 | 0.005 |
| | | 38,084,450 | *ORMDL3* | | 3.457 | 7.39E-04 | 0.005 |
| rs2313641 | g.38129233.A>T | 37,844,223 | *ERBB2* | *PGAP3* | 3.637 | 3.97E-04 | 7.82E-05 |
| | | 38,077,464 | | *ORMDL3* | 7.060 | 8.98E-11 | 0.014 |
| | | 38,084,331 | *ORMDL3* | | 4.116 | 6.81E-05 | 0.005 |
| | | 38,084,404 | *ORMDL3* | | 3.518 | 6.00E-04 | 0.005 |
| | | 38,084,450 | *ORMDL3* | | 3.457 | 7.39E-04 | 0.005 |
| rs4065876 | g.38129506.A>G | 37,910,697 | | | 3.513 | 6.10E-04 | 0.035 |
| | | 38,077,323 | | *ORMDL3* | 3.548 | 5.40E-04 | 0.014 |
| | | 38,077,464 | | *ORMDL3* | 3.885 | 1.62E-04 | 0.014 |
| | | 38,082,022 | | *ORMDL3* | 3.758 | 2.58E-04 | 6.40E-04 |
| | | 38,082,073 | | *ORMDL3* | 4.207 | 4.79E-05 | 6.40E-04 |
| | | 38,082,658 | | *ORMDL3* | 3.720 | 2.95E-04 | 0.011 |
| | | 38,108,710 | | | -5.079 | 1.29E-06 | 0.039 |
| | | 38,119,198 | *GSDMA* | | 3.881 | 1.65E-04 | 0.009 |
| | | 38,119,207 | *GSDMA* | | 4.375 | 2.47E-05 | 0.009 |
| | | 38,119,649 | | *GSDMA* | 4.305 | 3.26E-05 | 0.002 |
| | | 38,119,760 | | *GSDMA* | 3.726 | 2.89E-04 | 0.011 |
| | | 38,128,699 | | *GSDMA* | -5.161 | 8.97E-07 | 0.006 |
| rs748908389 | g.38129525_38129525ins | 37,910,697 | | | 3.378 | 9.65E-04 | 0.035 |
| | | 38,077,323 | | *ORMDL3* | 3.600 | 4.51E-04 | 0.014 |
| | | 38,077,464 | | *ORMDL3* | 3.702 | 3.14E-04 | 0.014 |
| | | 38,082,022 | | *ORMDL3* | 3.660 | 3.65E-04 | 6.40E-04 |
| | | 38,082,073 | | *ORMDL3* | 4.168 | 5.56E-05 | 6.40E-04 |
| | | 38,082,658 | | *ORMDL3* | 3.764 | 2.52E-04 | 0.011 |
| | | 38,108,710 | | | -5.039 | 1.53E-06 | 0.039 |
| | | 38,119,198 | *GSDMA* | | 3.816 | 2.09E-04 | 0.009 |
| | | 38,119,207 | *GSDMA* | | 4.320 | 3.07E-05 | 0.009 |
| | | 38,119,649 | | *GSDMA* | 4.417 | 2.09E-05 | 0.002 |
| | | 38,119,760 | | *GSDMA* | 3.682 | 3.38E-04 | 0.011 |
| | | 38,128,699 | | *GSDMA* | -5.138 | 9.92E-07 | 0.006 |
| rs11870683 | g.38129841.A>T | 37,913,383 | | | 3.873 | 1.70E-04 | 0.033 |
| | | 38,024,331 | | | -3.401 | 8.92E-04 | 0.017 |
| | | 38,049,272 | | | 3.500 | 6.38E-04 | 0.046 |

| | | | | | | | |
|---|---|---|---|---|---|---|---|
| | | 38,108,710 | | | -4.778 | 4.71E-06 | 0.039 |
| | | 38,128,699 | | *GSDMA* | -3.640 | 3.92E-04 | 0.006 |
| rs60137005 | g.38129996.T>A | 37,910,697 | | | 3.513 | 6.10E-04 | 0.035 |
| | | 38,077,323 | | *ORMDL3* | 3.548 | 5.40E-04 | 0.014 |
| | | 38,077,464 | | *ORMDL3* | 3.885 | 1.62E-04 | 0.014 |
| | | 38,082,022 | | *ORMDL3* | 3.758 | 2.58E-04 | 6.40E-04 |
| | | 38,082,073 | | *ORMDL3* | 4.207 | 4.79E-05 | 6.40E-04 |
| | | 38,082,658 | | *ORMDL3* | 3.720 | 2.95E-04 | 0.011 |
| | | 38,108,710 | | | -5.079 | 1.29E-06 | 0.039 |
| | | 38,119,198 | *GSDMA* | | 3.881 | 1.65E-04 | 0.009 |
| | | 38,119,207 | *GSDMA* | | 4.375 | 2.47E-05 | 0.009 |
| | | 38,119,649 | | *GSDMA* | 4.305 | 3.26E-05 | 0.002 |
| | | 38,119,760 | | *GSDMA* | 3.726 | 2.89E-04 | 0.011 |
| | | 38,128,699 | | *GSDMA* | -5.161 | 8.97E-07 | 0.006 |
| rs56326707 | g.38130139.T>C | 37,910,697 | | | 3.513 | 6.10E-04 | 0.035 |
| | | 38,077,323 | | *ORMDL3* | 3.548 | 5.40E-04 | 0.014 |
| | | 38,077,464 | | *ORMDL3* | 3.885 | 1.62E-04 | 0.014 |
| | | 38,082,022 | | *ORMDL3* | 3.758 | 2.58E-04 | 6.40E-04 |
| | | 38,082,073 | | *ORMDL3* | 4.207 | 4.79E-05 | 6.40E-04 |
| | | 38,082,658 | | *ORMDL3* | 3.720 | 2.95E-04 | 0.011 |
| | | 38,108,710 | | | -5.079 | 1.29E-06 | 0.039 |
| | | 38,119,198 | *GSDMA* | | 3.881 | 1.65E-04 | 0.009 |
| | | 38,119,207 | *GSDMA* | | 4.375 | 2.47E-05 | 0.009 |
| | | 38,119,649 | | *GSDMA* | 4.305 | 3.26E-05 | 0.002 |
| | | 38,119,760 | | *GSDMA* | 3.726 | 2.89E-04 | 0.011 |
| | | 38,128,699 | | *GSDMA* | -5.161 | 8.97E-07 | 0.006 |
| rs62066853 | g.38130492.C>T | 37,828,560 | | *PGAP3* | -3.741 | 2.74E-04 | 0.002 |
| | | 38,081,812 | | *ORMDL3* | -3.662 | 3.63E-04 | 2.01E-04 |
| | | 38,082,022 | | *ORMDL3* | -3.543 | 5.50E-04 | 6.40E-04 |
| | | 38,082,073 | | *ORMDL3* | -4.189 | 5.13E-05 | 6.40E-04 |
| rs56030650 | g.38131187.A>C (Thr.314.Asn) | 37,910,697 | | | 3.513 | 6.10E-04 | 0.035 |
| | | 38,077,323 | | *ORMDL3* | 3.548 | 5.40E-04 | 0.014 |
| | | 38,077,464 | | *ORMDL3* | 3.885 | 1.62E-04 | 0.014 |
| | | 38,082,022 | | *ORMDL3* | 3.758 | 2.58E-04 | 6.40E-04 |
| | | 38,082,073 | | *ORMDL3* | 4.207 | 4.79E-05 | 6.40E-04 |
| | | 38,082,658 | | *ORMDL3* | 3.720 | 2.95E-04 | 0.011 |
| | | 38,108,710 | | | -5.079 | 1.29E-06 | 0.039 |
| | | 38,119,198 | *GSDMA* | | 3.881 | 1.65E-04 | 0.009 |
| | | 38,119,207 | *GSDMA* | | 4.375 | 2.47E-05 | 0.009 |
| | | 38,119,649 | | *GSDMA* | 4.305 | 3.26E-05 | 0.002 |

| | | | | | | | | |
|---|---|---|---|---|---|---|---|---|
| | | | 38,119,760 | | *GSDMA* | 3.726 | 2.89E-04 | 0.011 |
| | | | 38,128,699 | | *GSDMA* | -5.161 | 8.97E-07 | 0.006 |
| | rs60134943 | g.38133792.T>G | 37,910,697 | | | 3.513 | 6.10E-04 | 0.035 |
| | | | 38,077,323 | | *ORMDL3* | 3.548 | 5.40E-04 | 0.014 |
| | | | 38,077,464 | | *ORMDL3* | 3.885 | 1.62E-04 | 0.014 |
| | | | 38,082,022 | | *ORMDL3* | 3.758 | 2.58E-04 | 6.40E-04 |
| | | | 38,082,073 | | *ORMDL3* | 4.207 | 4.79E-05 | 6.40E-04 |
| | | | 38,082,658 | | *ORMDL3* | 3.720 | 2.95E-04 | 0.011 |
| | | | 38,108,710 | | | -5.079 | 1.29E-06 | 0.039 |
| | | | 38,119,198 | *GSDMA* | | 3.881 | 1.65E-04 | 0.009 |
| | | | 38,119,207 | *GSDMA* | | 4.375 | 2.47E-05 | 0.009 |
| | | | 38,119,649 | | *GSDMA* | 4.305 | 3.26E-05 | 0.002 |
| | | | 38,119,760 | | *GSDMA* | 3.726 | 2.89E-04 | 0.011 |
| | | | 38,128,699 | | *GSDMA* | -5.161 | 8.97E-07 | 0.006 |
| | rs139141843 | g.38133914_38133914insA | 38,049,272 | | | 3.618 | 4.24E-04 | 0.046 |
| | | | 38,082,022 | | *ORMDL3* | 3.621 | 4.19E-04 | 6.40E-04 |
| | | | 38,082,073 | | *ORMDL3* | 3.770 | 2.47E-04 | 6.40E-04 |
| | | | 38,108,710 | | | -4.587 | 1.05E-05 | 0.039 |
| | rs921651 | g.38133922.G>A | 37,844,223 | *ERBB2* | *PGAP3* | 3.549 | 5.40E-04 | 7.82E-05 |
| | | | 38,077,464 | | *ORMDL3* | 7.105 | 7.08E-11 | 0.014 |
| | | | 38,084,331 | *ORMDL3* | | 4.005 | 1.04E-04 | 0.005 |
| | | | 38,084,404 | *ORMDL3* | | 3.524 | 5.88E-04 | 0.005 |
| | | | 38,084,450 | *ORMDL3* | | 3.428 | 8.15E-04 | 0.005 |
| NA | rs407307 | g.37827163.G>A | 37,828,195 | | *PGAP3* | -5.022 | 1.65E-06 | |
| | | | 37,828,197 | | *PGAP3* | -4.168 | 5.57E-05 | |
| | | | 37,828,220 | | *PGAP3* | -4.894 | 2.88E-06 | |
| | | | 37,828,273 | | *PGAP3* | -4.635 | 8.55E-06 | 0.010 |
| | | | 37,828,283 | | *PGAP3* | -3.909 | 1.48E-04 | 0.010 |
| | | | 37,828,296 | | *PGAP3* | -5.480 | 2.13E-07 | 0.002 |
| | | | 37,828,338 | | *PGAP3* | -5.026 | 1.63E-06 | 0.002 |
| | | | 37,828,377 | | *PGAP3* | -6.727 | 4.99E-10 | 0.002 |
| | | | 37,828,384 | | *PGAP3* | -5.853 | 3.71E-08 | 0.002 |
| | | | 37,828,560 | | *PGAP3* | -4.434 | 1.95E-05 | 0.002 |
| | | | 37,843,658 | *ERBB2* | *PGAP3* | -4.412 | 2.13E-05 | 7.82E-05 |
| | | | 37,844,480 | *PGAP3* | *ERBB2* | 4.037 | 9.20E-05 | 7.82E-05 |
| | | | 38,072,514 | | *GSDMB* | -3.487 | 6.67E-04 | 0.046 |
| | | | 38,082,022 | | *ORMDL3* | -3.995 | 1.08E-04 | 6.40E-04 |
| | | | 38,082,035 | | *ORMDL3* | -3.688 | 3.31E-04 | 6.40E-04 |
| | rs538149242 | g.37827292.G>T | 37,844,714 | *PGAP3* | *ERBB2* | -3.915 | 1.45E-04 | 7.82E-05 |
| | | | 37,911,152 | | | 5.059 | 1.40E-06 | 0.035 |

| | | | | | | | |
|---|---|---|---|---|---|---|---|
| | | 37,911,186 | | | 3.757 | 2.59E-04 | 0.035 |
| | | 37,914,933 | | | 3.691 | 3.28E-04 | 3.32E-05 |
| | | 38,083,955 | *ORMDL3* | | 3.756 | 2.60E-04 | 0.011 |
| | | 38,115,531 | | | 3.796 | 2.25E-04 | 0.009 |
| rs553962304 | g.37827454.T>C | 37,827,454 | | *PGAP3* | -4.074 | 7.99E-05 | |
| | | 37,910,326 | | | 4.183 | 5.25E-05 | 0.013 |
| | | 37,910,824 | | | 5.551 | 1.53E-07 | 0.035 |
| | | 38,018,269 | | *IKZF3* | -4.192 | 5.08E-05 | 0.046 |
| | | 38,084,404 | *ORMDL3* | | -3.384 | 9.45E-04 | 0.005 |
| | | 38,109,098 | | | 3.636 | 3.98E-04 | 0.002 |
| rs151309334 | g.37831293_37831298del | 37,886,788 | *MIEN1* | *MIEN1* | 4.136 | 6.29E-05 | 0.002 |
| | | 37,974,812 | | *IKZF3* | 3.456 | 7.41E-04 | 0.001 |
| | | 38,084,051 | *ORMDL3* | | 3.893 | 1.58E-04 | 0.005 |
| | | 38,096,571 | | | -3.796 | 2.25E-04 | 0.033 |
| | | 38,096,578 | | | -3.657 | 3.69E-04 | 0.033 |
| rs79524843 | g.37832386.T>G | 37,886,788 | *MIEN1* | *MIEN1* | 4.136 | 6.29E-05 | 0.002 |
| | | 37,974,812 | | *IKZF3* | 3.456 | 7.41E-04 | 0.001 |
| | | 38,084,051 | *ORMDL3* | | 3.893 | 1.58E-04 | 0.005 |
| | | 38,096,571 | | | -3.796 | 2.25E-04 | 0.033 |
| | | 38,096,578 | | | -3.657 | 3.69E-04 | 0.033 |
| rs76096054 | g.37832488.A>C | 37,886,788 | *MIEN1* | *MIEN1* | 4.136 | 6.29E-05 | 0.002 |
| | | 37,974,812 | | *IKZF3* | 3.456 | 7.41E-04 | 0.001 |
| | | 38,084,051 | *ORMDL3* | | 3.893 | 1.58E-04 | 0.005 |
| | | 38,096,571 | | | -3.796 | 2.25E-04 | 0.033 |
| | | 38,096,578 | | | -3.657 | 3.69E-04 | 0.033 |
| rs11656146 | g.37833562.T>C | 37,845,068 | *PGAP3* | *ERBB2* | 4.045 | 8.91E-05 | 7.82E-05 |
| | | 37,886,914 | *MIEN1* | | 3.932 | 1.36E-04 | 0.002 |
| | | 37,893,764 | | | 4.267 | 3.79E-05 | 0.002 |
| | | 38,025,241 | | | -3.977 | 1.15E-04 | 0.046 |
| | | 38,081,995 | | *ORMDL3* | 3.463 | 7.24E-04 | 2.01E-04 |
| | | 38,109,763 | | | -4.134 | 6.36E-05 | 0.002 |
| | | 38,121,965 | | *GSDMA* | 4.322 | 3.05E-05 | 0.011 |
| rs147180766 | g.37835355.A>C | 37,911,173 | | | 5.035 | 1.56E-06 | 0.035 |
| | | 38,051,338 | | | -4.575 | 1.10E-05 | 0.046 |
| | | 38,075,413 | *GSDMB* | | 3.410 | 8.65E-04 | 0.034 |
| | | 38,080,398 | | *ORMDL3* | 3.383 | 9.48E-04 | 0.014 |
| rs113831251 | g.37838200.A>G | 37,886,788 | *MIEN1* | *MIEN1* | 4.136 | 6.29E-05 | 0.002 |
| | | 37,974,812 | | *IKZF3* | 3.456 | 7.41E-04 | 0.001 |
| | | 38,084,051 | *ORMDL3* | | 3.893 | 1.58E-04 | 0.005 |
| | | 38,096,571 | | | -3.796 | 2.25E-04 | 0.033 |

| | | | | | | | |
|---|---|---|---|---|---|---|---|
| | | 38,096,578 | | | -3.657 | 3.69E-04 | 0.033 |
| rs113120593 | g.37840318.T>A | 37,886,788 | *MIEN1* | *MIEN1* | 4.136 | 6.29E-05 | 0.002 |
| | | 37,974,812 | | *IKZF3* | 3.456 | 7.41E-04 | 0.001 |
| | | 38,084,051 | *ORMDL3* | | 3.893 | 1.58E-04 | 0.005 |
| | | 38,096,571 | | | -3.796 | 2.25E-04 | 0.033 |
| | | 38,096,578 | | | -3.657 | 3.69E-04 | 0.033 |
| rs75849983 | g.37843762.G>T | 37,886,788 | *MIEN1* | *MIEN1* | 4.136 | 6.29E-05 | 0.002 |
| | | 37,974,812 | | *IKZF3* | 3.456 | 7.41E-04 | 0.001 |
| | | 38,084,051 | *ORMDL3* | | 3.893 | 1.58E-04 | 0.005 |
| | | 38,096,571 | | | -3.796 | 2.25E-04 | 0.033 |
| | | 38,096,578 | | | -3.657 | 3.69E-04 | 0.033 |
| rs146110680 | g.37847136.A>G | 37,896,010 | | | 3.683 | 3.37E-04 | 0.027 |
| | | 37,910,347 | | | 3.835 | 1.95E-04 | 0.013 |
| | | 37,911,274 | | | 4.371 | 2.50E-05 | 0.035 |
| | | 38,024,237 | | | -3.605 | 4.43E-04 | 0.016 |
| | | 38,024,367 | | | -3.790 | 2.30E-04 | 0.017 |
| | | 38,024,406 | | | -3.391 | 9.24E-04 | 3.75E-04 |
| | | 38,083,590 | | *ORMDL3* | 7.068 | 8.59E-11 | 0.011 |
| | | 38,083,894 | *ORMDL3* | | 4.651 | 8.01E-06 | 0.011 |
| rs201797794 | g.37847811T>A | 37,910,347 | | | 3.546 | 5.45E-04 | 0.013 |
| | | 37,910,684 | | | 3.861 | 1.77E-04 | 0.013 |
| | | 37,910,840 | | | 3.868 | 1.73E-04 | 0.035 |
| | | 38,083,520 | | *ORMDL3* | 3.500 | 6.38E-04 | 0.011 |
| | | 38,119,736 | | *GSDMA* | -3.532 | 5.72E-04 | 0.002 |
| rs116936671 | g.37848072.C>T | 37,828,842 | | *PGAP3* | 3.710 | 3.06E-04 | 0.002 |
| | | 38,083,491 | | *ORMDL3* | 3.709 | 3.07E-04 | 0.011 |
| | | 38,083,864 | | *ORMDL3* | 3.569 | 5.03E-04 | 0.011 |
| | | 38,118,571 | *GSDMA* | | -3.522 | 5.91E-04 | 0.009 |
| rs71383379 | g.37848153.T>G | 37,828,563 | | *PGAP3* | 3.746 | 2.69E-04 | 0.002 |
| | | 37,895,918 | | | -4.856 | 3.38E-06 | 0.027 |
| | | 37,910,171 | | | 5.878 | 3.30E-08 | 0.043 |
| | | 37,910,588 | | | 4.068 | 8.16E-05 | 0.013 |
| | | 37,910,756 | | | 3.747 | 2.68E-04 | 0.035 |
| | | 37,910,781 | | | 3.368 | 9.97E-04 | 0.035 |
| | | 38,074,925 | *GSDMB* | | -4.464 | 1.72E-05 | 0.034 |
| | | 38,077,870 | | *ORMDL3* | -4.042 | 9.04E-05 | 0.014 |
| | | 38,108,710 | | | 3.664 | 3.60E-04 | 0.039 |
| | | 38,125,830 | | *GSDMA* | 3.836 | 1.94E-04 | 0.006 |
| rs572656035 | g.37848683_37848684del | 37,844,476 | *PGAP3* | *ERBB2* | 3.759 | 2.57E-04 | 7.82E-05 |
| | | 37,886,749 | | *MIEN1* | 8.626 | 1.91E-14 | 0.002 |

| | | | | | | | |
|---|---|---|---|---|---|---|---|
| | | 38,077,051 | | | -3.644 | 3.86E-04 | 0.034 |
| | | 38,080,415 | | *ORMDL3* | 3.409 | 8.67E-04 | 0.014 |
| | | 38,080,829 | | *ORMDL3* | 4.279 | 3.61E-05 | 0.014 |
| | | 38,096,571 | | | -4.321 | 3.06E-05 | 0.033 |
| | | 38,096,578 | | | -4.038 | 9.15E-05 | 0.033 |
| | | 38,096,648 | | | -3.914 | 1.46E-04 | 0.033 |
| rs118132312 | g.37851015.G>A | 37,893,894 | | | 4.855 | 3.40E-06 | 0.002 |
| | | 37,896,050 | | | 5.132 | 1.02E-06 | 0.027 |
| | | 37,896,705 | | | 4.375 | 2.47E-05 | 0.008 |
| | | 37,910,542 | | | 3.960 | 1.23E-04 | 0.013 |
| | | 38,023,480 | | | 3.433 | 8.00E-04 | 0.003 |
| | | 38,083,656 | | *ORMDL3* | 5.415 | 2.86E-07 | 0.011 |
| | | 38,084,256 | *ORMDL3* | | -3.795 | 2.25E-04 | 0.005 |
| | | 38,115,535 | | | 3.527 | 5.81E-04 | 0.009 |
| rs551511055 | g.37852066_37852073del | 37,834,869 | | *PGAP3* | 3.725 | 2.90E-04 | 0.020 |
| | | 37,886,474 | | *MIEN1* | 3.817 | 2.08E-04 | 0.002 |
| | | 37,886,749 | | *MIEN1* | 3.615 | 4.28E-04 | 0.002 |
| | | 37,886,894 | *MIEN1* | | 4.276 | 3.66E-05 | 0.002 |
| | | 37,896,827 | | | 3.804 | 2.18E-04 | 0.008 |
| | | 37,910,636 | | | 3.937 | 1.34E-04 | 0.013 |
| | | 38,083,413 | | *ORMDL3* | 4.623 | 9.01E-06 | 0.011 |
| | | 38,083,548 | | *ORMDL3* | 5.058 | 1.41E-06 | 0.011 |
| | | 38,109,660 | | | 3.441 | 7.79E-04 | 0.002 |
| | | 38,133,359 | | *GSDMA* | 4.428 | 2.00E-05 | 0.038 |
| rs553346880 | g.37853810.C>T | 37,827,454 | | *PGAP3* | -4.074 | 7.99E-05 | |
| | | 37,910,326 | | | 4.183 | 5.25E-05 | 0.013 |
| | | 37,910,824 | | | 5.551 | 1.53E-07 | 0.035 |
| | | 38,018,269 | | *IKZF3* | -4.192 | 5.08E-05 | 0.046 |
| | | 38,084,404 | *ORMDL3* | | -3.384 | 9.45E-04 | 0.005 |
| | | 38,109,098 | | | 3.636 | 3.98E-04 | 0.002 |
| rs111700892 | g.37853908.T>C | 37,886,788 | *MIEN1* | *MIEN1* | 4.136 | 6.29E-05 | 0.002 |
| | | 37,974,812 | | *IKZF3* | 3.456 | 7.41E-04 | 0.001 |
| | | 38,084,051 | *ORMDL3* | | 3.893 | 1.58E-04 | 0.005 |
| | | 38,096,571 | | | -3.796 | 2.25E-04 | 0.033 |
| | | 38,096,578 | | | -3.657 | 3.69E-04 | 0.033 |
| rs79747793 | g.37855331.A>C | 37,886,788 | *MIEN1* | *MIEN1* | 4.136 | 6.29E-05 | 0.002 |
| | | 37,974,812 | | *IKZF3* | 3.456 | 7.41E-04 | 0.001 |
| | | 38,084,051 | *ORMDL3* | | 3.893 | 1.58E-04 | 0.005 |
| | | 38,096,571 | | | -3.796 | 2.25E-04 | 0.033 |
| | | 38,096,578 | | | -3.657 | 3.69E-04 | 0.033 |

| | | | | | | | |
|---|---|---|---|---|---|---|---|
| rs12947247 | g.37855541.C>T | 37,886,788 | *MIEN1* | *MIEN1* | 4.136 | 6.29E-05 | 0.002 |
| | | 37,974,812 | | *IKZF3* | 3.456 | 7.41E-04 | 0.001 |
| | | 38,084,051 | *ORMDL3* | | 3.893 | 1.58E-04 | 0.005 |
| | | 38,096,571 | | | -3.796 | 2.25E-04 | 0.033 |
| | | 38,096,578 | | | -3.657 | 3.69E-04 | 0.033 |
| rs34284966 | g.37859083.C>G | 37,886,788 | *MIEN1* | *MIEN1* | 4.136 | 6.29E-05 | 0.002 |
| | | 37,974,812 | | *IKZF3* | 3.456 | 7.41E-04 | 0.001 |
| | | 38,084,051 | *ORMDL3* | | 3.893 | 1.58E-04 | 0.005 |
| | | 38,096,571 | | | -3.796 | 2.25E-04 | 0.033 |
| | | 38,096,578 | | | -3.657 | 3.69E-04 | 0.033 |
| rs138742873 | g.37860716.T>C | 37,896,010 | | | 3.683 | 3.37E-04 | 0.027 |
| | | 37,910,347 | | | 3.835 | 1.95E-04 | 0.013 |
| | | 37,911,274 | | | 4.371 | 2.50E-05 | 0.035 |
| | | 38,024,237 | | | -3.605 | 4.43E-04 | 0.016 |
| | | 38,024,367 | | | -3.790 | 2.30E-04 | 0.017 |
| | | 38,024,406 | | | -3.391 | 9.24E-04 | 3.75E-04 |
| | | 38,083,590 | | *ORMDL3* | 7.068 | 8.59E-11 | 0.011 |
| | | 38,083,894 | *ORMDL3* | | 4.651 | 8.01E-06 | 0.011 |
| rs34006795 | g.37860837.T>C | 37,886,788 | *MIEN1* | *MIEN1* | 4.136 | 6.29E-05 | 0.002 |
| | | 37,974,812 | | *IKZF3* | 3.456 | 7.41E-04 | 0.001 |
| | | 38,084,051 | *ORMDL3* | | 3.893 | 1.58E-04 | 0.005 |
| | | 38,096,571 | | | -3.796 | 2.25E-04 | 0.033 |
| | | 38,096,578 | | | -3.657 | 3.69E-04 | 0.033 |
| rs118036166 | g.37860994.G>A | 37,844,398 | *PGAP3* | *ERBB2* | 3.683 | 3.37E-04 | 7.82E-05 |
| | | 37,911,152 | | | 3.876 | 1.68E-04 | 0.035 |
| | | 37,914,933 | | | 3.429 | 8.12E-04 | 3.32E-05 |
| | | 38,083,537 | | *ORMDL3* | 3.650 | 3.78E-04 | 0.011 |
| rs4252608 | g.37862966.T>C | 37,886,788 | *MIEN1* | *MIEN1* | 4.136 | 6.29E-05 | 0.002 |
| | | 37,974,812 | | *IKZF3* | 3.456 | 7.41E-04 | 0.001 |
| | | 38,084,051 | *ORMDL3* | | 3.893 | 1.58E-04 | 0.005 |
| | | 38,096,571 | | | -3.796 | 2.25E-04 | 0.033 |
| | | 38,096,578 | | | -3.657 | 3.69E-04 | 0.033 |
| rs148040998 | g.37863606_37863606ins TCGGCCA | 37,886,788 | *MIEN1* | *MIEN1* | 4.136 | 6.29E-05 | 0.002 |
| | | 37,974,812 | | *IKZF3* | 3.456 | 7.41E-04 | 0.001 |
| | | 38,084,051 | *ORMDL3* | | 3.893 | 1.58E-04 | 0.005 |
| | | 38,096,571 | | | -3.796 | 2.25E-04 | 0.033 |
| | | 38,096,578 | | | -3.657 | 3.69E-04 | 0.033 |
| rs4252612 | g.37863708.T>C | 37,886,653 | | *MIEN1* | 3.627 | 4.10E-04 | 0.002 |
| | | 37,910,988 | | | 3.518 | 5.99E-04 | 0.035 |
| rs539944790 | g.37867517.C>T | 37,836,263 | | *PGAP3* | -3.933 | 1.36E-04 | 0.019 |

| | | | | | | | |
|---|---|---|---|---|---|---|---|
| | | 37,886,619 | | *MIEN1* | 3.604 | 4.44E-04 | 0.002 |
| | | 37,910,190 | | | 4.487 | 1.57E-05 | 0.043 |
| rs4252623 | g.37867633.G>C | 37,886,827 | *MIEN1* | | 3.395 | 9.09E-04 | 0.002 |
| | | 37,896,063 | | | 3.857 | 1.80E-04 | 0.009 |
| | | 37,910,286 | | | 3.951 | 1.27E-04 | 0.043 |
| | | 37,910,609 | | | 3.427 | 8.18E-04 | 0.013 |
| | | 37,910,707 | | | 3.788 | 2.31E-04 | 0.035 |
| | | 37,913,515 | | | 4.057 | 8.53E-05 | 3.32E-05 |
| | | 38,080,829 | | *ORMDL3* | 5.041 | 1.52E-06 | 0.014 |
| | | 38,083,958 | *ORMDL3* | | 3.426 | 8.20E-04 | 0.011 |
| rs4252624 | g.37867647.T>G | 37,886,788 | *MIEN1* | *MIEN1* | 4.136 | 6.29E-05 | 0.002 |
| | | 37,974,812 | | *IKZF3* | 3.456 | 7.41E-04 | 0.001 |
| | | 38,084,051 | *ORMDL3* | | 3.893 | 1.58E-04 | 0.005 |
| | | 38,096,571 | | | -3.796 | 2.25E-04 | 0.033 |
| | | 38,096,578 | | | -3.657 | 3.69E-04 | 0.033 |
| rs113231716 | g.37869990.G>A | 37,886,788 | *MIEN1* | *MIEN1* | 4.136 | 6.29E-05 | 0.002 |
| | | 37,974,812 | | *IKZF3* | 3.456 | 7.41E-04 | 0.001 |
| | | 38,084,051 | *ORMDL3* | | 3.893 | 1.58E-04 | 0.005 |
| | | 38,096,571 | | | -3.796 | 2.25E-04 | 0.033 |
| | | 38,096,578 | | | -3.657 | 3.69E-04 | 0.033 |
| rs1181234061 | g.37874043_37874049del | 37,844,480 | *PGAP3* | *ERBB2* | 3.923 | 1.41E-04 | 7.82E-05 |
| rs541484690 | g.37874043_37874047del | 37,886,653 | | *MIEN1* | 3.627 | 4.10E-04 | 0.002 |
| | | 37,886,788 | *MIEN1* | *MIEN1* | 4.997 | 1.84E-06 | 0.002 |
| | | 37,910,988 | | | 3.518 | 5.99E-04 | 0.035 |
| | | 38,096,571 | | | -3.604 | 4.44E-04 | 0.033 |
| | | 38,096,578 | | | -3.726 | 2.89E-04 | 0.033 |
| rs4252639 | g.37876179.G>C | 37,886,788 | *MIEN1* | *MIEN1* | 4.136 | 6.29E-05 | 0.002 |
| | | 37,974,812 | | *IKZF3* | 3.456 | 7.41E-04 | 0.001 |
| | | 38,084,051 | *ORMDL3* | | 3.893 | 1.58E-04 | 0.005 |
| | | 38,096,571 | | | -3.796 | 2.25E-04 | 0.033 |
| | | 38,096,578 | | | -3.657 | 3.69E-04 | 0.033 |
| rs111947409 | g.37877716.A>G | 37,886,788 | *MIEN1* | *MIEN1* | 4.136 | 6.29E-05 | 0.002 |
| | | 37,974,812 | | *IKZF3* | 3.456 | 7.41E-04 | 0.001 |
| | | 38,084,051 | *ORMDL3* | | 3.893 | 1.58E-04 | 0.005 |
| | | 38,096,571 | | | -3.796 | 2.25E-04 | 0.033 |
| | | 38,096,578 | | | -3.657 | 3.69E-04 | 0.033 |
| rs115334808 | g.37878113.T>C | 37,886,788 | *MIEN1* | *MIEN1* | 4.136 | 6.29E-05 | 0.002 |
| | | 37,974,812 | | *IKZF3* | 3.456 | 7.41E-04 | 0.001 |
| | | 38,084,051 | *ORMDL3* | | 3.893 | 1.58E-04 | 0.005 |

| | | | | | | | |
|---|---|---|---|---|---|---|---|
| | | 38,096,571 | | | -3.796 | 2.25E-04 | 0.033 |
| | | 38,096,578 | | | -3.657 | 3.69E-04 | 0.033 |
| rs183373485 | g.37878553.T>C | 37,827,454 | | *PGAP3* | -4.074 | 7.99E-05 | |
| | | 37,910,326 | | | 4.183 | 5.25E-05 | 0.013 |
| | | 37,910,824 | | | 5.551 | 1.53E-07 | 0.035 |
| | | 38,018,269 | | *IKZF3* | -4.192 | 5.08E-05 | 0.046 |
| | | 38,084,404 | *ORMDL3* | | -3.384 | 9.45E-04 | 0.005 |
| | | 38,109,098 | | | 3.636 | 3.98E-04 | 0.002 |
| rs4252665 | g.37885383.T>C | 37,856,732 | | *ERBB2* | -4.060 | 8.43E-05 | 0.009 |
| | | 37,886,749 | | *MIEN1* | 4.283 | 3.56E-05 | 0.002 |
| | | 37,895,335 | | | 3.388 | 9.33E-04 | 0.006 |
| | | 37,896,063 | | | 3.530 | 5.75E-04 | 0.009 |
| | | 37,896,203 | | | 3.438 | 7.89E-04 | 0.011 |
| | | 37,976,456 | | *IKZF3* | 3.672 | 3.51E-04 | 0.039 |
| | | 38,076,680 | | | 3.418 | 8.42E-04 | 0.034 |
| | | 38,080,829 | | *ORMDL3* | 3.933 | 1.36E-04 | 0.014 |
| | | 38,096,333 | | | -3.509 | 6.18E-04 | 0.033 |
| | | 38,096,571 | | | -4.910 | 2.68E-06 | 0.033 |
| | | 38,096,578 | | | -5.756 | 5.87E-08 | 0.033 |
| | | 38,096,648 | | | -4.451 | 1.82E-05 | 0.033 |
| | | 38,118,571 | *GSDMA* | | 3.697 | 3.21E-04 | 0.009 |
| rs139233737 | g.37887636_37887636ins T | 37,856,732 | | *ERBB2* | -4.060 | 8.43E-05 | 0.009 |
| | | 37,886,749 | | *MIEN1* | 4.283 | 3.56E-05 | 0.002 |
| | | 37,895,335 | | | 3.388 | 9.33E-04 | 0.006 |
| | | 37,896,063 | | | 3.530 | 5.75E-04 | 0.009 |
| | | 37,896,203 | | | 3.438 | 7.89E-04 | 0.011 |
| | | 37,976,456 | | *IKZF3* | 3.672 | 3.51E-04 | 0.039 |
| | | 38,076,680 | | | 3.418 | 8.42E-04 | 0.034 |
| | | 38,080,829 | | *ORMDL3* | 3.933 | 1.36E-04 | 0.014 |
| | | 38,096,333 | | | -3.509 | 6.18E-04 | 0.033 |
| | | 38,096,571 | | | -4.910 | 2.68E-06 | 0.033 |
| | | 38,096,578 | | | -5.756 | 5.87E-08 | 0.033 |
| | | 38,096,648 | | | -4.451 | 1.82E-05 | 0.033 |
| | | 38,118,571 | *GSDMA* | | 3.697 | 3.21E-04 | 0.009 |
| rs112535478 | g.37888034.G>A | 37,856,732 | | *ERBB2* | -4.060 | 8.43E-05 | 0.009 |
| | | 37,886,749 | | *MIEN1* | 4.283 | 3.56E-05 | 0.002 |
| | | 37,895,335 | | | 3.388 | 9.33E-04 | 0.006 |
| | | 37,896,063 | | | 3.530 | 5.75E-04 | 0.009 |
| | | 37,896,203 | | | 3.438 | 7.89E-04 | 0.011 |
| | | 37,976,456 | | *IKZF3* | 3.672 | 3.51E-04 | 0.039 |

| | | | | | | | |
|---|---|---|---|---|---|---|---|
| | | 38,076,680 | | | 3.418 | 8.42E-04 | 0.034 |
| | | 38,080,829 | | *ORMDL3* | 3.933 | 1.36E-04 | 0.014 |
| | | 38,096,333 | | | -3.509 | 6.18E-04 | 0.033 |
| | | 38,096,571 | | | -4.910 | 2.68E-06 | 0.033 |
| | | 38,096,578 | | | -5.756 | 5.87E-08 | 0.033 |
| | | 38,096,648 | | | -4.451 | 1.82E-05 | 0.033 |
| | | 38,118,571 | *GSDMA* | | 3.697 | 3.21E-04 | 0.009 |
| rs111904390 | g.37888516.T>G | 37,856,732 | | *ERBB2* | -3.668 | 3.56E-04 | 0.009 |
| | | 37,886,749 | | *MIEN1* | 3.945 | 1.30E-04 | 0.002 |
| | | 37,896,063 | | | 3.476 | 6.92E-04 | 0.009 |
| | | 38,020,485 | *IKZF3* | | 3.962 | 1.22E-04 | 0.005 |
| | | 38,080,829 | | *ORMDL3* | 3.995 | 1.08E-04 | 0.014 |
| | | 38,096,333 | | | -3.664 | 3.60E-04 | 0.033 |
| | | 38,096,571 | | | -4.780 | 4.67E-06 | 0.033 |
| | | 38,096,578 | | | -5.642 | 1.00E-07 | 0.033 |
| | | 38,096,648 | | | -3.887 | 1.61E-04 | 0.033 |
| rs144954923 | g.37889025_37889026del | 37,886,788 | *MIEN1* | *MIEN1* | 4.136 | 6.29E-05 | 0.002 |
| | | 37,974,812 | | *IKZF3* | 3.456 | 7.41E-04 | 0.001 |
| | | 38,084,051 | *ORMDL3* | | 3.893 | 1.58E-04 | 0.005 |
| | | 38,096,571 | | | -3.796 | 2.25E-04 | 0.033 |
| | | 38,096,578 | | | -3.657 | 3.69E-04 | 0.033 |
| rs111424867 | g.37889280.T>C | 37,886,788 | *MIEN1* | *MIEN1* | 4.136 | 6.29E-05 | 0.002 |
| | | 37,974,812 | | *IKZF3* | 3.456 | 7.41E-04 | 0.001 |
| | | 38,084,051 | *ORMDL3* | | 3.893 | 1.58E-04 | 0.005 |
| | | 38,096,571 | | | -3.796 | 2.25E-04 | 0.033 |
| | | 38,096,578 | | | -3.657 | 3.69E-04 | 0.033 |
| rs11868060 | g.37891116.G>A | 37,886,619 | | *MIEN1* | 3.647 | 3.82E-04 | 0.002 |
| | | 37,894,916 | | | 4.681 | 7.07E-06 | 0.004 |
| | | 37,896,708 | | | 3.507 | 6.22E-04 | 0.008 |
| | | 37,934,107 | | *IKZF3* | -4.209 | 4.76E-05 | 0.027 |
| | | 37,949,408 | | *IKZF3* | -3.462 | 7.26E-04 | 0.003 |
| | | 38,024,502 | | | -3.690 | 3.29E-04 | 0.005 |
| | | 38,083,275 | | *ORMDL3* | 3.693 | 3.25E-04 | 0.011 |
| | | 38,084,016 | *ORMDL3* | | 3.751 | 2.65E-04 | 0.011 |
| | | 38,109,131 | | | 6.223 | 6.20E-09 | 0.002 |
| rs78841518 | g.37891229.G>A | 37,886,788 | *MIEN1* | *MIEN1* | 4.136 | 6.29E-05 | 0.002 |
| | | 37,974,812 | | *IKZF3* | 3.456 | 7.41E-04 | 0.001 |
| | | 38,084,051 | *ORMDL3* | | 3.893 | 1.58E-04 | 0.005 |
| | | 38,096,571 | | | -3.796 | 2.25E-04 | 0.033 |
| | | 38,096,578 | | | -3.657 | 3.69E-04 | 0.033 |

| | | | | | | | |
|---|---|---|---|---|---|---|---|
| rs77755027 | g.37891235.A>G | 37,886,788 | *MIEN1* | *MIEN1* | 4.136 | 6.29E-05 | 0.002 |
| | | 37,974,812 | | *IKZF3* | 3.456 | 7.41E-04 | 0.001 |
| | | 38,084,051 | *ORMDL3* | | 3.893 | 1.58E-04 | 0.005 |
| | | 38,096,571 | | | -3.796 | 2.25E-04 | 0.033 |
| | | 38,096,578 | | | -3.657 | 3.69E-04 | 0.033 |
| rs112585477 | g.37891654.A>G | 37,856,732 | | *ERBB2* | -3.668 | 3.56E-04 | 0.009 |
| | | 37,886,749 | | *MIEN1* | 3.945 | 1.30E-04 | 0.002 |
| | | 37,896,063 | | | 3.476 | 6.92E-04 | 0.009 |
| | | 38,020,485 | *IKZF3* | | 3.962 | 1.22E-04 | 0.005 |
| | | 38,080,829 | | *ORMDL3* | 3.995 | 1.08E-04 | 0.014 |
| | | 38,096,333 | | | -3.664 | 3.60E-04 | 0.033 |
| | | 38,096,571 | | | -4.780 | 4.67E-06 | 0.033 |
| | | 38,096,578 | | | -5.642 | 1.00E-07 | 0.033 |
| | | 38,096,648 | | | -3.887 | 1.61E-04 | 0.033 |
| rs142447482 | g.37891686.T>C | 37,827,057 | | | 3.449 | 7.60E-04 | |
| | | 37,832,244 | | *PGAP3* | 3.878 | 1.67E-04 | 0.028 |
| | | 37,910,347 | | | 4.224 | 4.48E-05 | 0.013 |
| | | 38,024,014 | | | -4.449 | 1.84E-05 | 0.004 |
| | | 38,024,237 | | | -4.169 | 5.54E-05 | 0.016 |
| | | 38,024,242 | | | -3.883 | 1.63E-04 | 0.016 |
| | | 38,024,244 | | | -3.765 | 2.51E-04 | 0.016 |
| | | 38,083,590 | | *ORMDL3* | 4.237 | 4.25E-05 | 0.011 |
| | | 38,083,894 | *ORMDL3* | | 4.799 | 4.32E-06 | 0.011 |
| rs112327574 | g.37893663.G>C | 37,856,732 | | *ERBB2* | -3.668 | 3.56E-04 | 0.009 |
| | | 37,886,749 | | *MIEN1* | 3.945 | 1.30E-04 | 0.002 |
| | | 37,896,063 | | | 3.476 | 6.92E-04 | 0.009 |
| | | 38,020,485 | *IKZF3* | | 3.962 | 1.22E-04 | 0.005 |
| | | 38,080,829 | | *ORMDL3* | 3.995 | 1.08E-04 | 0.014 |
| | | 38,096,333 | | | -3.664 | 3.60E-04 | 0.033 |
| | | 38,096,571 | | | -4.780 | 4.67E-06 | 0.033 |
| | | 38,096,578 | | | -5.642 | 1.00E-07 | 0.033 |
| | | 38,096,648 | | | -3.887 | 1.61E-04 | 0.033 |
| rs9896940 | g.37895975.G>A | 37,856,732 | | *ERBB2* | -3.668 | 3.56E-04 | 0.009 |
| | | 37,886,749 | | *MIEN1* | 3.945 | 1.30E-04 | 0.002 |
| | | 37,896,063 | | | 3.476 | 6.92E-04 | 0.009 |
| | | 38,020,485 | *IKZF3* | | 3.962 | 1.22E-04 | 0.005 |
| | | 38,080,829 | | *ORMDL3* | 3.995 | 1.08E-04 | 0.014 |
| | | 38,096,333 | | | -3.664 | 3.60E-04 | 0.033 |
| | | 38,096,571 | | | -4.780 | 4.67E-06 | 0.033 |
| | | 38,096,578 | | | -5.642 | 1.00E-07 | 0.033 |
| | | 38,096,648 | | | -3.887 | 1.61E-04 | 0.033 |

| | | | | | | | |
|---|---|---|---|---|---|---|---|
| rs113903339 | g.37898047.T>C | 37,856,732 | | *ERBB2* | -3.668 | 3.56E-04 | 0.009 |
| | | 37,886,749 | | *MIEN1* | 3.945 | 1.30E-04 | 0.002 |
| | | 37,896,063 | | | 3.476 | 6.92E-04 | 0.009 |
| | | 38,020,485 | *IKZF3* | | 3.962 | 1.22E-04 | 0.005 |
| | | 38,080,829 | | *ORMDL3* | 3.995 | 1.08E-04 | 0.014 |
| | | 38,096,333 | | | -3.664 | 3.60E-04 | 0.033 |
| | | 38,096,571 | | | -4.780 | 4.67E-06 | 0.033 |
| | | 38,096,578 | | | -5.642 | 1.00E-07 | 0.033 |
| | | 38,096,648 | | | -3.887 | 1.61E-04 | 0.033 |
| rs76121559 | g.37899092.A>G | 37,844,480 | *PGAP3* | *ERBB2* | 3.929 | 1.38E-04 | 7.82E-05 |
| | | 37,886,788 | *MIEN1* | *MIEN1* | 5.000 | 1.82E-06 | 0.002 |
| | | 38,096,571 | | | -3.587 | 4.72E-04 | 0.033 |
| | | 38,096,578 | | | -3.714 | 3.01E-04 | 0.033 |
| rs796319243 | g.37903880_37903881del | 37,856,732 | | *ERBB2* | -3.668 | 3.56E-04 | 0.009 |
| | | 37,886,749 | | *MIEN1* | 3.945 | 1.30E-04 | 0.002 |
| | | 37,896,063 | | | 3.476 | 6.92E-04 | 0.009 |
| | | 38,020,485 | *IKZF3* | | 3.962 | 1.22E-04 | 0.005 |
| | | 38,080,829 | | *ORMDL3* | 3.995 | 1.08E-04 | 0.014 |
| | | 38,096,333 | | | -3.664 | 3.60E-04 | 0.033 |
| | | 38,096,571 | | | -4.780 | 4.67E-06 | 0.033 |
| | | 38,096,578 | | | -5.642 | 1.00E-07 | 0.033 |
| | | 38,096,648 | | | -3.887 | 1.61E-04 | 0.033 |
| rs9910678 | g.37904211.C>T | 37,856,732 | | *ERBB2* | -3.668 | 3.56E-04 | 0.009 |
| | | 37,886,749 | | *MIEN1* | 3.945 | 1.30E-04 | 0.002 |
| | | 37,896,063 | | | 3.476 | 6.92E-04 | 0.009 |
| | | 38,020,485 | *IKZF3* | | 3.962 | 1.22E-04 | 0.005 |
| | | 38,080,829 | | *ORMDL3* | 3.995 | 1.08E-04 | 0.014 |
| | | 38,096,333 | | | -3.664 | 3.60E-04 | 0.033 |
| | | 38,096,571 | | | -4.780 | 4.67E-06 | 0.033 |
| | | 38,096,578 | | | -5.642 | 1.00E-07 | 0.033 |
| | | 38,096,648 | | | -3.887 | 1.61E-04 | 0.033 |
| rs75933089 | g.37904449.T>G | 37,856,732 | | *ERBB2* | -3.668 | 3.56E-04 | 0.009 |
| | | 37,886,749 | | *MIEN1* | 3.945 | 1.30E-04 | 0.002 |
| | | 37,896,063 | | | 3.476 | 6.92E-04 | 0.009 |
| | | 38,020,485 | *IKZF3* | | 3.962 | 1.22E-04 | 0.005 |
| | | 38,080,829 | | *ORMDL3* | 3.995 | 1.08E-04 | 0.014 |
| | | 38,096,333 | | | -3.664 | 3.60E-04 | 0.033 |
| | | 38,096,571 | | | -4.780 | 4.67E-06 | 0.033 |
| | | 38,096,578 | | | -5.642 | 1.00E-07 | 0.033 |
| | | 38,096,648 | | | -3.887 | 1.61E-04 | 0.033 |

| | | | | | | | |
|---|---|---|---|---|---|---|---|
| rs180809456 | g.37906738.A>G | 37,886,725 | | *MIEN1* | 3.438 | 7.88E-04 | 0.002 |
| | | 37,886,742 | | *MIEN1* | 3.456 | 7.42E-04 | 0.002 |
| | | 37,893,894 | | | 4.915 | 2.63E-06 | 0.002 |
| | | 37,896,050 | | | 5.259 | 5.80E-07 | 0.027 |
| | | 37,896,705 | | | 4.287 | 3.50E-05 | 0.008 |
| | | 37,910,542 | | | 3.913 | 1.46E-04 | 0.013 |
| | | 37,910,569 | | | 3.437 | 7.91E-04 | 0.013 |
| | | 38,083,656 | | *ORMDL3* | 5.466 | 2.26E-07 | 0.011 |
| | | 38,084,256 | *ORMDL3* | | -3.744 | 2.71E-04 | 0.005 |
| | | 38,115,535 | | | 3.580 | 4.84E-04 | 0.009 |
| rs78365632 | g.37906916.A>G | 37,856,732 | | *ERBB2* | -3.668 | 3.56E-04 | 0.009 |
| | | 37,886,749 | | *MIEN1* | 3.945 | 1.30E-04 | 0.002 |
| | | 37,896,063 | | | 3.476 | 6.92E-04 | 0.009 |
| | | 38,020,485 | *IKZF3* | | 3.962 | 1.22E-04 | 0.005 |
| | | 38,080,829 | | *ORMDL3* | 3.995 | 1.08E-04 | 0.014 |
| | | 38,096,333 | | | -3.664 | 3.60E-04 | 0.033 |
| | | 38,096,571 | | | -4.780 | 4.67E-06 | 0.033 |
| | | 38,096,578 | | | -5.642 | 1.00E-07 | 0.033 |
| | | 38,096,648 | | | -3.887 | 1.61E-04 | 0.033 |
| rs9891131 | g.37911521.G>A | 37,856,732 | | *ERBB2* | -3.668 | 3.56E-04 | 0.009 |
| | | 37,886,749 | | *MIEN1* | 3.945 | 1.30E-04 | 0.002 |
| | | 37,896,063 | | | 3.476 | 6.92E-04 | 0.009 |
| | | 38,020,485 | *IKZF3* | | 3.962 | 1.22E-04 | 0.005 |
| | | 38,080,829 | | *ORMDL3* | 3.995 | 1.08E-04 | 0.014 |
| | | 38,096,333 | | | -3.664 | 3.60E-04 | 0.033 |
| | | 38,096,571 | | | -4.780 | 4.67E-06 | 0.033 |
| | | 38,096,578 | | | -5.642 | 1.00E-07 | 0.033 |
| | | 38,096,648 | | | -3.887 | 1.61E-04 | 0.033 |
| rs77013147 | g.37912268.C>T | 37,856,732 | | *ERBB2* | -3.668 | 3.56E-04 | 0.009 |
| | | 37,886,749 | | *MIEN1* | 3.945 | 1.30E-04 | 0.002 |
| | | 37,896,063 | | | 3.476 | 6.92E-04 | 0.009 |
| | | 38,020,485 | *IKZF3* | | 3.962 | 1.22E-04 | 0.005 |
| | | 38,080,829 | | *ORMDL3* | 3.995 | 1.08E-04 | 0.014 |
| | | 38,096,333 | | | -3.664 | 3.60E-04 | 0.033 |
| | | 38,096,571 | | | -4.780 | 4.67E-06 | 0.033 |
| | | 38,096,578 | | | -5.642 | 1.00E-07 | 0.033 |
| | | 38,096,648 | | | -3.887 | 1.61E-04 | 0.033 |
| rs117753409 | g.37912447.A>C | 37,828,773 | | *PGAP3* | 3.405 | 8.80E-04 | 0.002 |
| | | 37,856,857 | | *ERBB2* | -3.409 | 8.68E-04 | 0.009 |
| | | 37,886,765 | | *MIEN1* | 3.518 | 6.00E-04 | 0.002 |
| | | 37,886,921 | *MIEN1* | | 3.552 | 5.33E-04 | 0.002 |

| | | | | | | | |
|---|---|---|---|---|---|---|---|
| | | 37,896,050 | | | 3.683 | 3.37E-04 | 0.027 |
| | | 37,910,406 | | | 3.737 | 2.78E-04 | 0.013 |
| | | 37,910,697 | | | 3.384 | 9.44E-04 | 0.035 |
| | | 37,910,840 | | | 3.970 | 1.18E-04 | 0.035 |
| | | 37,911,360 | | | 3.408 | 8.72E-04 | 0.035 |
| | | 37,975,698 | | *IKZF3* | 5.196 | 7.67E-07 | 0.039 |
| | | 38,020,483 | *IKZF3* | | 4.621 | 9.10E-06 | 0.005 |
| | | 38,024,250 | | | -4.882 | 3.03E-06 | 0.016 |
| | | 38,075,557 | *GSDMB* | | 5.039 | 1.53E-06 | 0.034 |
| | | 38,077,234 | | | -3.420 | 8.36E-04 | 0.014 |
| | | 38,083,520 | | *ORMDL3* | 3.533 | 5.70E-04 | 0.011 |
| | | 38,083,862 | | *ORMDL3* | 3.701 | 3.16E-04 | 0.011 |
| | | 38,108,734 | | | 4.222 | 4.52E-05 | 0.039 |
| | | 38,119,132 | *GSDMA* | | 3.964 | 1.21E-04 | 0.009 |
| rs190852402 | g.37915314.T>A | 37,886,872 | *MIEN1* | | 4.778 | 4.70E-06 | 0.002 |
| | | 37,910,923 | | | 4.844 | 3.55E-06 | 0.035 |
| | | 37,911,173 | | | 6.613 | 8.87E-10 | 0.035 |
| | | 37,947,519 | | *IKZF3* | -3.577 | 4.89E-04 | 0.002 |
| | | 38,020,606 | *IKZF3* | | 3.419 | 8.41E-04 | 0.017 |
| | | 38,109,653 | | | 5.448 | 2.46E-07 | 0.002 |
| rs72832927 | g.37917011.T>C | 37,886,827 | *MIEN1* | | 3.395 | 9.09E-04 | 0.002 |
| | | 37,896,063 | | | 3.857 | 1.80E-04 | 0.009 |
| | | 37,910,286 | | | 3.951 | 1.27E-04 | 0.043 |
| | | 37,910,609 | | | 3.427 | 8.18E-04 | 0.013 |
| | | 37,910,707 | | | 3.788 | 2.31E-04 | 0.035 |
| | | 37,913,515 | | | 4.057 | 8.53E-05 | 3.32E-05 |
| | | 38,080,829 | | *ORMDL3* | 5.041 | 1.52E-06 | 0.014 |
| | | 38,083,958 | *ORMDL3* | | 3.426 | 8.20E-04 | 0.011 |
| rs573534572 | g.37917826.T>C | 37,856,732 | | *ERBB2* | -3.668 | 3.56E-04 | 0.009 |
| | | 37,886,749 | | *MIEN1* | 3.945 | 1.30E-04 | 0.002 |
| | | 37,896,063 | | | 3.476 | 6.92E-04 | 0.009 |
| | | 38,020,485 | *IKZF3* | | 3.962 | 1.22E-04 | 0.005 |
| | | 38,080,829 | | *ORMDL3* | 3.995 | 1.08E-04 | 0.014 |
| | | 38,096,333 | | | -3.664 | 3.60E-04 | 0.033 |
| | | 38,096,571 | | | -4.780 | 4.67E-06 | 0.033 |
| | | 38,096,578 | | | -5.642 | 1.00E-07 | 0.033 |
| | | 38,096,648 | | | -3.887 | 1.61E-04 | 0.033 |
| rs9909432 | g.37918106.C>T | 37,896,063 | | | 3.508 | 6.21E-04 | 0.009 |
| | | 37,976,987 | | *IKZF3* | 3.405 | 8.81E-04 | 0.039 |
| | | 38,077,234 | | | -3.416 | 8.48E-04 | 0.014 |
| | | 38,080,829 | | *ORMDL3* | 4.051 | 8.70E-05 | 0.014 |

| | | | | | | | |
|---|---|---|---|---|---|---|---|
| | | 38,096,333 | | | -3.630 | 4.07E-04 | 0.033 |
| | | 38,096,571 | | | -4.161 | 5.73E-05 | 0.033 |
| | | 38,096,578 | | | -5.082 | 1.27E-06 | 0.033 |
| | | 38,096,648 | | | -3.741 | 2.74E-04 | 0.033 |
| rs67571561 | g.37920847.C>T | 37,856,732 | | *ERBB2* | -3.668 | 3.56E-04 | 0.009 |
| | | 37,886,749 | | *MIEN1* | 3.945 | 1.30E-04 | 0.002 |
| | | 37,896,063 | | | 3.476 | 6.92E-04 | 0.009 |
| | | 38,020,485 | *IKZF3* | | 3.962 | 1.22E-04 | 0.005 |
| | | 38,080,829 | | *ORMDL3* | 3.995 | 1.08E-04 | 0.014 |
| | | 38,096,333 | | | -3.664 | 3.60E-04 | 0.033 |
| | | 38,096,571 | | | -4.780 | 4.67E-06 | 0.033 |
| | | 38,096,578 | | | -5.642 | 1.00E-07 | 0.033 |
| | | 38,096,648 | | | -3.887 | 1.61E-04 | 0.033 |
| rs2941509 | g.37921194.A>G | 37,856,732 | | *ERBB2* | -3.668 | 3.56E-04 | 0.009 |
| | | 37,886,749 | | *MIEN1* | 3.945 | 1.30E-04 | 0.002 |
| | | 37,896,063 | | | 3.476 | 6.92E-04 | 0.009 |
| | | 38,020,485 | *IKZF3* | | 3.962 | 1.22E-04 | 0.005 |
| | | 38,080,829 | | *ORMDL3* | 3.995 | 1.08E-04 | 0.014 |
| | | 38,096,333 | | | -3.664 | 3.60E-04 | 0.033 |
| | | 38,096,571 | | | -4.780 | 4.67E-06 | 0.033 |
| | | 38,096,578 | | | -5.642 | 1.00E-07 | 0.033 |
| | | 38,096,648 | | | -3.887 | 1.61E-04 | 0.033 |
| rs112876941 | g.37922804.T>A | 37,856,732 | | *ERBB2* | -3.668 | 3.56E-04 | 0.009 |
| | | 37,886,749 | | *MIEN1* | 3.945 | 1.30E-04 | 0.002 |
| | | 37,896,063 | | | 3.476 | 6.92E-04 | 0.009 |
| | | 38,020,485 | *IKZF3* | | 3.962 | 1.22E-04 | 0.005 |
| | | 38,080,829 | | *ORMDL3* | 3.995 | 1.08E-04 | 0.014 |
| | | 38,096,333 | | | -3.664 | 3.60E-04 | 0.033 |
| | | 38,096,571 | | | -4.780 | 4.67E-06 | 0.033 |
| | | 38,096,578 | | | -5.642 | 1.00E-07 | 0.033 |
| | | 38,096,648 | | | -3.887 | 1.61E-04 | 0.033 |
| rs112771360 | g.37923771.G>A | 37,856,732 | | *ERBB2* | -3.668 | 3.56E-04 | 0.009 |
| | | 37,886,749 | | *MIEN1* | 3.945 | 1.30E-04 | 0.002 |
| | | 37,896,063 | | | 3.476 | 6.92E-04 | 0.009 |
| | | 38,020,485 | *IKZF3* | | 3.962 | 1.22E-04 | 0.005 |
| | | 38,080,829 | | *ORMDL3* | 3.995 | 1.08E-04 | 0.014 |
| | | 38,096,333 | | | -3.664 | 3.60E-04 | 0.033 |
| | | 38,096,571 | | | -4.780 | 4.67E-06 | 0.033 |
| | | 38,096,578 | | | -5.642 | 1.00E-07 | 0.033 |
| | | 38,096,648 | | | -3.887 | 1.61E-04 | 0.033 |
| rs34758895 | g.37925468.T>C | 37,856,732 | | *ERBB2* | -3.668 | 3.56E-04 | 0.009 |

| | | | | | | | |
|---|---|---|---|---|---|---|---|
| | | 37,886,749 | | *MIEN1* | 3.945 | 1.30E-04 | 0.002 |
| | | 37,896,063 | | | 3.476 | 6.92E-04 | 0.009 |
| | | 38,020,485 | *IKZF3* | | 3.962 | 1.22E-04 | 0.005 |
| | | 38,080,829 | | *ORMDL3* | 3.995 | 1.08E-04 | 0.014 |
| | | 38,096,333 | | | -3.664 | 3.60E-04 | 0.033 |
| | | 38,096,571 | | | -4.780 | 4.67E-06 | 0.033 |
| | | 38,096,578 | | | -5.642 | 1.00E-07 | 0.033 |
| | | 38,096,648 | | | -3.887 | 1.61E-04 | 0.033 |
| rs9894370 | g.37926004.C>G | 37,856,732 | | *ERBB2* | -3.668 | 3.56E-04 | 0.009 |
| | | 37,886,749 | | *MIEN1* | 3.945 | 1.30E-04 | 0.002 |
| | | 37,896,063 | | | 3.476 | 6.92E-04 | 0.009 |
| | | 38,020,485 | *IKZF3* | | 3.962 | 1.22E-04 | 0.005 |
| | | 38,080,829 | | *ORMDL3* | 3.995 | 1.08E-04 | 0.014 |
| | | 38,096,333 | | | -3.664 | 3.60E-04 | 0.033 |
| | | 38,096,571 | | | -4.780 | 4.67E-06 | 0.033 |
| | | 38,096,578 | | | -5.642 | 1.00E-07 | 0.033 |
| | | 38,096,648 | | | -3.887 | 1.61E-04 | 0.033 |
| rs8072612 | g.37927120.G>A | 37,856,732 | | *ERBB2* | -3.668 | 3.56E-04 | 0.009 |
| | | 37,886,749 | | *MIEN1* | 3.945 | 1.30E-04 | 0.002 |
| | | 37,896,063 | | | 3.476 | 6.92E-04 | 0.009 |
| | | 38,020,485 | *IKZF3* | | 3.962 | 1.22E-04 | 0.005 |
| | | 38,080,829 | | *ORMDL3* | 3.995 | 1.08E-04 | 0.014 |
| | | 38,096,333 | | | -3.664 | 3.60E-04 | 0.033 |
| | | 38,096,571 | | | -4.780 | 4.67E-06 | 0.033 |
| | | 38,096,578 | | | -5.642 | 1.00E-07 | 0.033 |
| | | 38,096,648 | | | -3.887 | 1.61E-04 | 0.033 |
| rs71369788 | g.37927145.A>G | 37,856,732 | | *ERBB2* | -3.668 | 3.56E-04 | 0.009 |
| | | 37,886,749 | | *MIEN1* | 3.945 | 1.30E-04 | 0.002 |
| | | 37,896,063 | | | 3.476 | 6.92E-04 | 0.009 |
| | | 38,020,485 | *IKZF3* | | 3.962 | 1.22E-04 | 0.005 |
| | | 38,080,829 | | *ORMDL3* | 3.995 | 1.08E-04 | 0.014 |
| | | 38,096,333 | | | -3.664 | 3.60E-04 | 0.033 |
| | | 38,096,571 | | | -4.780 | 4.67E-06 | 0.033 |
| | | 38,096,578 | | | -5.642 | 1.00E-07 | 0.033 |
| | | 38,096,648 | | | -3.887 | 1.61E-04 | 0.033 |
| rs9652838 | g.37929248.C>T | 37,856,732 | | *ERBB2* | -3.668 | 3.56E-04 | 0.009 |
| | | 37,886,749 | | *MIEN1* | 3.945 | 1.30E-04 | 0.002 |
| | | 37,896,063 | | | 3.476 | 6.92E-04 | 0.009 |
| | | 38,020,485 | *IKZF3* | | 3.962 | 1.22E-04 | 0.005 |
| | | 38,080,829 | | *ORMDL3* | 3.995 | 1.08E-04 | 0.014 |
| | | 38,096,333 | | | -3.664 | 3.60E-04 | 0.033 |

| | | | | | | | |
|---|---|---|---|---|---|---|---|
| | | 38,096,571 | | | -4.780 | 4.67E-06 | 0.033 |
| | | 38,096,578 | | | -5.642 | 1.00E-07 | 0.033 |
| | | 38,096,648 | | | -3.887 | 1.61E-04 | 0.033 |
| rs9652840 | g.37929428.T>A | 37,856,732 | | *ERBB2* | -3.668 | 3.56E-04 | 0.009 |
| | | 37,886,749 | | *MIEN1* | 3.945 | 1.30E-04 | 0.002 |
| | | 37,896,063 | | | 3.476 | 6.92E-04 | 0.009 |
| | | 38,020,485 | *IKZF3* | | 3.962 | 1.22E-04 | 0.005 |
| | | 38,080,829 | | *ORMDL3* | 3.995 | 1.08E-04 | 0.014 |
| | | 38,096,333 | | | -3.664 | 3.60E-04 | 0.033 |
| | | 38,096,571 | | | -4.780 | 4.67E-06 | 0.033 |
| | | 38,096,578 | | | -5.642 | 1.00E-07 | 0.033 |
| | | 38,096,648 | | | -3.887 | 1.61E-04 | 0.033 |
| rs77925972 | g.37930210.C>T | 37,856,732 | | *ERBB2* | -3.668 | 3.56E-04 | 0.009 |
| | | 37,886,749 | | *MIEN1* | 3.945 | 1.30E-04 | 0.002 |
| | | 37,896,063 | | | 3.476 | 6.92E-04 | 0.009 |
| | | 38,020,485 | *IKZF3* | | 3.962 | 1.22E-04 | 0.005 |
| | | 38,080,829 | | *ORMDL3* | 3.995 | 1.08E-04 | 0.014 |
| | | 38,096,333 | | | -3.664 | 3.60E-04 | 0.033 |
| | | 38,096,571 | | | -4.780 | 4.67E-06 | 0.033 |
| | | 38,096,578 | | | -5.642 | 1.00E-07 | 0.033 |
| | | 38,096,648 | | | -3.887 | 1.61E-04 | 0.033 |
| rs7217133 | g.37930262.A>T | 37,856,732 | | *ERBB2* | -3.659 | 3.67E-04 | 0.009 |
| | | 37,886,749 | | *MIEN1* | 3.939 | 1.33E-04 | 0.002 |
| | | 37,896,063 | | | 3.534 | 5.68E-04 | 0.009 |
| | | 38,020,485 | *IKZF3* | | 3.952 | 1.27E-04 | 0.005 |
| | | 38,080,829 | | *ORMDL3* | 4.004 | 1.04E-04 | 0.014 |
| | | 38,096,333 | | | -3.674 | 3.48E-04 | 0.033 |
| | | 38,096,571 | | | -4.755 | 5.19E-06 | 0.033 |
| | | 38,096,578 | | | -5.590 | 1.28E-07 | 0.033 |
| | | 38,096,648 | | | -3.903 | 1.52E-04 | 0.033 |
| rs113730542 | g.37930421.G>C | 37,856,732 | | *ERBB2* | -3.668 | 3.56E-04 | 0.009 |
| | | 37,886,749 | | *MIEN1* | 3.945 | 1.30E-04 | 0.002 |
| | | 37,896,063 | | | 3.476 | 6.92E-04 | 0.009 |
| | | 38,020,485 | *IKZF3* | | 3.962 | 1.22E-04 | 0.005 |
| | | 38,080,829 | | *ORMDL3* | 3.995 | 1.08E-04 | 0.014 |
| | | 38,096,333 | | | -3.664 | 3.60E-04 | 0.033 |
| | | 38,096,571 | | | -4.780 | 4.67E-06 | 0.033 |
| | | 38,096,578 | | | -5.642 | 1.00E-07 | 0.033 |
| | | 38,096,648 | | | -3.887 | 1.61E-04 | 0.033 |
| rs2313429 | g.37930659.C>T | 37,856,732 | | *ERBB2* | -3.668 | 3.56E-04 | 0.009 |
| | | 37,886,749 | | *MIEN1* | 3.945 | 1.30E-04 | 0.002 |

| | | | | | | | |
|---|---|---|---|---|---|---|---|
| | | 37,896,063 | | | 3.476 | 6.92E-04 | 0.009 |
| | | 38,020,485 | *IKZF3* | | 3.962 | 1.22E-04 | 0.005 |
| | | 38,080,829 | | *ORMDL3* | 3.995 | 1.08E-04 | 0.014 |
| | | 38,096,333 | | | -3.664 | 3.60E-04 | 0.033 |
| | | 38,096,571 | | | -4.780 | 4.67E-06 | 0.033 |
| | | 38,096,578 | | | -5.642 | 1.00E-07 | 0.033 |
| | | 38,096,648 | | | -3.887 | 1.61E-04 | 0.033 |
| rs7503018 | g.37931220.G>A | 37,856,732 | | *ERBB2* | -3.668 | 3.56E-04 | 0.009 |
| | | 37,886,749 | | *MIEN1* | 3.945 | 1.30E-04 | 0.002 |
| | | 37,896,063 | | | 3.476 | 6.92E-04 | 0.009 |
| | | 38,020,485 | *IKZF3* | | 3.962 | 1.22E-04 | 0.005 |
| | | 38,080,829 | | *ORMDL3* | 3.995 | 1.08E-04 | 0.014 |
| | | 38,096,333 | | | -3.664 | 3.60E-04 | 0.033 |
| | | 38,096,571 | | | -4.780 | 4.67E-06 | 0.033 |
| | | 38,096,578 | | | -5.642 | 1.00E-07 | 0.033 |
| | | 38,096,648 | | | -3.887 | 1.61E-04 | 0.033 |
| rs181818554 | g.37931904.A>G | 37,828,842 | | *PGAP3* | 3.391 | 9.23E-04 | 0.002 |
| | | 37,844,177 | *ERBB2* | *PGAP3* | 3.475 | 6.94E-04 | 7.82E-05 |
| | | 37,886,619 | | *MIEN1* | 4.005 | 1.04E-04 | 0.002 |
| | | 37,910,190 | | | 3.575 | 4.92E-04 | 0.043 |
| | | 38,083,491 | | *ORMDL3* | 3.461 | 7.28E-04 | 0.011 |
| | | 38,083,864 | | *ORMDL3* | 3.825 | 2.02E-04 | 0.011 |
| rs9913596 | g.37932063.A>G | 37,856,732 | | *ERBB2* | -3.668 | 3.56E-04 | 0.009 |
| | | 37,886,749 | | *MIEN1* | 3.945 | 1.30E-04 | 0.002 |
| | | 37,896,063 | | | 3.476 | 6.92E-04 | 0.009 |
| | | 38,020,485 | *IKZF3* | | 3.962 | 1.22E-04 | 0.005 |
| | | 38,080,829 | | *ORMDL3* | 3.995 | 1.08E-04 | 0.014 |
| | | 38,096,333 | | | -3.664 | 3.60E-04 | 0.033 |
| | | 38,096,571 | | | -4.780 | 4.67E-06 | 0.033 |
| | | 38,096,578 | | | -5.642 | 1.00E-07 | 0.033 |
| | | 38,096,648 | | | -3.887 | 1.61E-04 | 0.033 |
| rs9894898 | g.37932221.C>T | 37,856,732 | | *ERBB2* | -3.668 | 3.56E-04 | 0.009 |
| | | 37,886,749 | | *MIEN1* | 3.945 | 1.30E-04 | 0.002 |
| | | 37,896,063 | | | 3.476 | 6.92E-04 | 0.009 |
| | | 38,020,485 | *IKZF3* | | 3.962 | 1.22E-04 | 0.005 |
| | | 38,080,829 | | *ORMDL3* | 3.995 | 1.08E-04 | 0.014 |
| | | 38,096,333 | | | -3.664 | 3.60E-04 | 0.033 |
| | | 38,096,571 | | | -4.780 | 4.67E-06 | 0.033 |
| | | 38,096,578 | | | -5.642 | 1.00E-07 | 0.033 |
| | | 38,096,648 | | | -3.887 | 1.61E-04 | 0.033 |
| rs9901483 | g.37932774.A>T | 37,856,732 | | *ERBB2* | -3.668 | 3.56E-04 | 0.009 |

| | | | | | | | |
|---|---|---|---|---|---|---|---|
| | | 37,886,749 | | *MIEN1* | 3.945 | 1.30E-04 | 0.002 |
| | | 37,896,063 | | | 3.476 | 6.92E-04 | 0.009 |
| | | 38,020,485 | *IKZF3* | | 3.962 | 1.22E-04 | 0.005 |
| | | 38,080,829 | | *ORMDL3* | 3.995 | 1.08E-04 | 0.014 |
| | | 38,096,333 | | | -3.664 | 3.60E-04 | 0.033 |
| | | 38,096,571 | | | -4.780 | 4.67E-06 | 0.033 |
| | | 38,096,578 | | | -5.642 | 1.00E-07 | 0.033 |
| | | 38,096,648 | | | -3.887 | 1.61E-04 | 0.033 |
| rs113370572 | g.37933468.C>T | 37,856,732 | | *ERBB2* | -3.668 | 3.56E-04 | 0.009 |
| | | 37,886,749 | | *MIEN1* | 3.945 | 1.30E-04 | 0.002 |
| | | 37,896,063 | | | 3.476 | 6.92E-04 | 0.009 |
| | | 38,020,485 | *IKZF3* | | 3.962 | 1.22E-04 | 0.005 |
| | | 38,080,829 | | *ORMDL3* | 3.995 | 1.08E-04 | 0.014 |
| | | 38,096,333 | | | -3.664 | 3.60E-04 | 0.033 |
| | | 38,096,571 | | | -4.780 | 4.67E-06 | 0.033 |
| | | 38,096,578 | | | -5.642 | 1.00E-07 | 0.033 |
| | | 38,096,648 | | | -3.887 | 1.61E-04 | 0.033 |
| rs12709364 | g.37933823.G>A | 37,856,732 | | *ERBB2* | -3.668 | 3.56E-04 | 0.009 |
| | | 37,886,749 | | *MIEN1* | 3.945 | 1.30E-04 | 0.002 |
| | | 37,896,063 | | | 3.476 | 6.92E-04 | 0.009 |
| | | 38,020,485 | *IKZF3* | | 3.962 | 1.22E-04 | 0.005 |
| | | 38,080,829 | | *ORMDL3* | 3.995 | 1.08E-04 | 0.014 |
| | | 38,096,333 | | | -3.664 | 3.60E-04 | 0.033 |
| | | 38,096,571 | | | -4.780 | 4.67E-06 | 0.033 |
| | | 38,096,578 | | | -5.642 | 1.00E-07 | 0.033 |
| | | 38,096,648 | | | -3.887 | 1.61E-04 | 0.033 |
| rs138576332 | g.37934349.C>T | 37,844,365 | *PGAP3ERBB2* | | 4.379 | 2.43E-05 | 7.82E-05 |
| | | 37,893,744 | | | 3.827 | 2.00E-04 | 0.002 |
| | | 37,910,730 | | | 3.640 | 3.92E-04 | 0.035 |
| | | 37,910,939 | | | 3.578 | 4.87E-04 | 0.035 |
| | | 38,077,464 | | *ORMDL3* | 11.387 | 2.90E-21 | 0.014 |
| | | 38,119,760 | | *GSDMA* | 4.349 | 2.74E-05 | 0.011 |
| rs7224641 | g.37934911.C>T | 37,856,732 | | *ERBB2* | -3.668 | 3.56E-04 | 0.009 |
| | | 37,886,749 | | *MIEN1* | 3.945 | 1.30E-04 | 0.002 |
| | | 37,896,063 | | | 3.476 | 6.92E-04 | 0.009 |
| | | 38,020,485 | *IKZF3* | | 3.962 | 1.22E-04 | 0.005 |
| | | 38,080,829 | | *ORMDL3* | 3.995 | 1.08E-04 | 0.014 |
| | | 38,096,333 | | | -3.664 | 3.60E-04 | 0.033 |
| | | 38,096,571 | | | -4.780 | 4.67E-06 | 0.033 |
| | | 38,096,578 | | | -5.642 | 1.00E-07 | 0.033 |
| | | 38,096,648 | | | -3.887 | 1.61E-04 | 0.033 |

| | | | | | | | |
|---|---|---|---|---|---|---|---|
| rs13380871 | g.37936249.C>T | 37,856,732 | | *ERBB2* | -3.668 | 3.56E-04 | 0.009 |
| | | 37,886,749 | | *MIEN1* | 3.945 | 1.30E-04 | 0.002 |
| | | 37,896,063 | | | 3.476 | 6.92E-04 | 0.009 |
| | | 38,020,485 | *IKZF3* | | 3.962 | 1.22E-04 | 0.005 |
| | | 38,080,829 | | *ORMDL3* | 3.995 | 1.08E-04 | 0.014 |
| | | 38,096,333 | | | -3.664 | 3.60E-04 | 0.033 |
| | | 38,096,571 | | | -4.780 | 4.67E-06 | 0.033 |
| | | 38,096,578 | | | -5.642 | 1.00E-07 | 0.033 |
| | | 38,096,648 | | | -3.887 | 1.61E-04 | 0.033 |
| rs35506518 | g.37938094.C>T | 37,856,732 | | *ERBB2* | -3.668 | 3.56E-04 | 0.009 |
| | | 37,886,749 | | *MIEN1* | 3.945 | 1.30E-04 | 0.002 |
| | | 37,896,063 | | | 3.476 | 6.92E-04 | 0.009 |
| | | 38,020,485 | *IKZF3* | | 3.962 | 1.22E-04 | 0.005 |
| | | 38,080,829 | | *ORMDL3* | 3.995 | 1.08E-04 | 0.014 |
| | | 38,096,333 | | | -3.664 | 3.60E-04 | 0.033 |
| | | 38,096,571 | | | -4.780 | 4.67E-06 | 0.033 |
| | | 38,096,578 | | | -5.642 | 1.00E-07 | 0.033 |
| | | 38,096,648 | | | -3.887 | 1.61E-04 | 0.033 |
| rs67605703 | g.37938497.C>T | 37,856,732 | | *ERBB2* | -3.668 | 3.56E-04 | 0.009 |
| | | 37,886,749 | | *MIEN1* | 3.945 | 1.30E-04 | 0.002 |
| | | 37,896,063 | | | 3.476 | 6.92E-04 | 0.009 |
| | | 38,020,485 | *IKZF3* | | 3.962 | 1.22E-04 | 0.005 |
| | | 38,080,829 | | *ORMDL3* | 3.995 | 1.08E-04 | 0.014 |
| | | 38,096,333 | | | -3.664 | 3.60E-04 | 0.033 |
| | | 38,096,571 | | | -4.780 | 4.67E-06 | 0.033 |
| | | 38,096,578 | | | -5.642 | 1.00E-07 | 0.033 |
| | | 38,096,648 | | | -3.887 | 1.61E-04 | 0.033 |
| rs58566623 | g.37938665_37938684del | 37,856,732 | | *ERBB2* | -3.702 | 3.15E-04 | 0.009 |
| | | 37,886,749 | | *MIEN1* | 3.938 | 1.33E-04 | 0.002 |
| | | 37,896,063 | | | 3.471 | 7.05E-04 | 0.009 |
| | | 38,020,485 | *IKZF3* | | 3.965 | 1.20E-04 | 0.005 |
| | | 38,080,829 | | *ORMDL3* | 4.014 | 1.00E-04 | 0.014 |
| | | 38,096,333 | | | -3.752 | 2.63E-04 | 0.033 |
| | | 38,096,571 | | | -4.879 | 3.07E-06 | 0.033 |
| | | 38,096,578 | | | -5.722 | 6.89E-08 | 0.033 |
| | | 38,096,648 | | | -3.912 | 1.47E-04 | 0.033 |
| rs34016964 | g.37938977.T>G | 37,856,732 | | *ERBB2* | -3.668 | 3.56E-04 | 0.009 |
| | | 37,886,749 | | *MIEN1* | 3.945 | 1.30E-04 | 0.002 |
| | | 37,896,063 | | | 3.476 | 6.92E-04 | 0.009 |
| | | 38,020,485 | *IKZF3* | | 3.962 | 1.22E-04 | 0.005 |

| | | | | | | | |
|---|---|---|---|---|---|---|---|
| | | 38,080,829 | | *ORMDL3* | 3.995 | 1.08E-04 | 0.014 |
| | | 38,096,333 | | | -3.664 | 3.60E-04 | 0.033 |
| | | 38,096,571 | | | -4.780 | 4.67E-06 | 0.033 |
| | | 38,096,578 | | | -5.642 | 1.00E-07 | 0.033 |
| | | 38,096,648 | | | -3.887 | 1.61E-04 | 0.033 |
| rs72832928 | g.37939398.T>G | 37,886,827 | *MIEN1* | | 3.395 | 9.09E-04 | 0.002 |
| | | 37,896,063 | | | 3.857 | 1.80E-04 | 0.009 |
| | | 37,910,286 | | | 3.951 | 1.27E-04 | 0.043 |
| | | 37,910,609 | | | 3.427 | 8.18E-04 | 0.013 |
| | | 37,910,707 | | | 3.788 | 2.31E-04 | 0.035 |
| | | 37,913,515 | | | 4.057 | 8.53E-05 | 3.32E-05 |
| | | 38,080,829 | | *ORMDL3* | 5.041 | 1.52E-06 | 0.014 |
| | | 38,083,958 | *ORMDL3* | | 3.426 | 8.20E-04 | 0.011 |
| rs9909365 | g.37939959.G>A | 37,856,732 | | *ERBB2* | -3.668 | 3.56E-04 | 0.009 |
| | | 37,886,749 | | *MIEN1* | 3.945 | 1.30E-04 | 0.002 |
| | | 37,896,063 | | | 3.476 | 6.92E-04 | 0.009 |
| | | 38,020,485 | *IKZF3* | | 3.962 | 1.22E-04 | 0.005 |
| | | 38,080,829 | | *ORMDL3* | 3.995 | 1.08E-04 | 0.014 |
| | | 38,096,333 | | | -3.664 | 3.60E-04 | 0.033 |
| | | 38,096,571 | | | -4.780 | 4.67E-06 | 0.033 |
| | | 38,096,578 | | | -5.642 | 1.00E-07 | 0.033 |
| | | 38,096,648 | | | -3.887 | 1.61E-04 | 0.033 |
| rs113812449 | g.37940168.C>T | 37,856,732 | | *ERBB2* | -3.668 | 3.56E-04 | 0.009 |
| | | 37,886,749 | | *MIEN1* | 3.945 | 1.30E-04 | 0.002 |
| | | 37,896,063 | | | 3.476 | 6.92E-04 | 0.009 |
| | | 38,020,485 | *IKZF3* | | 3.962 | 1.22E-04 | 0.005 |
| | | 38,080,829 | | *ORMDL3* | 3.995 | 1.08E-04 | 0.014 |
| | | 38,096,333 | | | -3.664 | 3.60E-04 | 0.033 |
| | | 38,096,571 | | | -4.780 | 4.67E-06 | 0.033 |
| | | 38,096,578 | | | -5.642 | 1.00E-07 | 0.033 |
| | | 38,096,648 | | | -3.887 | 1.61E-04 | 0.033 |
| rs1510475 | g.37941380.G>T | 37,856,732 | | *ERBB2* | -3.668 | 3.56E-04 | 0.009 |
| | | 37,886,749 | | *MIEN1* | 3.945 | 1.30E-04 | 0.002 |
| | | 37,896,063 | | | 3.476 | 6.92E-04 | 0.009 |
| | | 38,020,485 | *IKZF3* | | 3.962 | 1.22E-04 | 0.005 |
| | | 38,080,829 | | *ORMDL3* | 3.995 | 1.08E-04 | 0.014 |
| | | 38,096,333 | | | -3.664 | 3.60E-04 | 0.033 |
| | | 38,096,571 | | | -4.780 | 4.67E-06 | 0.033 |
| | | 38,096,578 | | | -5.642 | 1.00E-07 | 0.033 |
| | | 38,096,648 | | | -3.887 | 1.61E-04 | 0.033 |
| rs112345383 | g.37942018.T>C | 37,856,732 | | *ERBB2* | -3.668 | 3.56E-04 | 0.009 |

| | | | | | | | |
|---|---|---|---|---|---|---|---|
| | | 37,886,749 | | *MIEN1* | 3.945 | 1.30E-04 | 0.002 |
| | | 37,896,063 | | | 3.476 | 6.92E-04 | 0.009 |
| | | 38,020,485 | *IKZF3* | | 3.962 | 1.22E-04 | 0.005 |
| | | 38,080,829 | | *ORMDL3* | 3.995 | 1.08E-04 | 0.014 |
| | | 38,096,333 | | | -3.664 | 3.60E-04 | 0.033 |
| | | 38,096,571 | | | -4.780 | 4.67E-06 | 0.033 |
| | | 38,096,578 | | | -5.642 | 1.00E-07 | 0.033 |
| | | 38,096,648 | | | -3.887 | 1.61E-04 | 0.033 |
| rs34599546 | g.37942972.T>C | 37,856,732 | | *ERBB2* | -3.668 | 3.56E-04 | 0.009 |
| | | 37,886,749 | | *MIEN1* | 3.945 | 1.30E-04 | 0.002 |
| | | 37,896,063 | | | 3.476 | 6.92E-04 | 0.009 |
| | | 38,020,485 | *IKZF3* | | 3.962 | 1.22E-04 | 0.005 |
| | | 38,080,829 | | *ORMDL3* | 3.995 | 1.08E-04 | 0.014 |
| | | 38,096,333 | | | -3.664 | 3.60E-04 | 0.033 |
| | | 38,096,571 | | | -4.780 | 4.67E-06 | 0.033 |
| | | 38,096,578 | | | -5.642 | 1.00E-07 | 0.033 |
| | | 38,096,648 | | | -3.887 | 1.61E-04 | 0.033 |
| rs111862642 | g.37942984.G>C | 37,856,732 | | *ERBB2* | -3.668 | 3.56E-04 | 0.009 |
| | | 37,886,749 | | *MIEN1* | 3.945 | 1.30E-04 | 0.002 |
| | | 37,896,063 | | | 3.476 | 6.92E-04 | 0.009 |
| | | 38,020,485 | *IKZF3* | | 3.962 | 1.22E-04 | 0.005 |
| | | 38,080,829 | | *ORMDL3* | 3.995 | 1.08E-04 | 0.014 |
| | | 38,096,333 | | | -3.664 | 3.60E-04 | 0.033 |
| | | 38,096,571 | | | -4.780 | 4.67E-06 | 0.033 |
| | | 38,096,578 | | | -5.642 | 1.00E-07 | 0.033 |
| | | 38,096,648 | | | -3.887 | 1.61E-04 | 0.033 |
| rs9911634 | g.37943731.A>C | 37,856,732 | | *ERBB2* | -3.668 | 3.56E-04 | 0.009 |
| | | 37,886,749 | | *MIEN1* | 3.945 | 1.30E-04 | 0.002 |
| | | 37,896,063 | | | 3.476 | 6.92E-04 | 0.009 |
| | | 38,020,485 | *IKZF3* | | 3.962 | 1.22E-04 | 0.005 |
| | | 38,080,829 | | *ORMDL3* | 3.995 | 1.08E-04 | 0.014 |
| | | 38,096,333 | | | -3.664 | 3.60E-04 | 0.033 |
| | | 38,096,571 | | | -4.780 | 4.67E-06 | 0.033 |
| | | 38,096,578 | | | -5.642 | 1.00E-07 | 0.033 |
| | | 38,096,648 | | | -3.887 | 1.61E-04 | 0.033 |
| rs9911669 | g.37943767.G>C | 37,856,732 | | *ERBB2* | -3.668 | 3.56E-04 | 0.009 |
| | | 37,886,749 | | *MIEN1* | 3.945 | 1.30E-04 | 0.002 |
| | | 37,896,063 | | | 3.476 | 6.92E-04 | 0.009 |
| | | 38,020,485 | *IKZF3* | | 3.962 | 1.22E-04 | 0.005 |
| | | 38,080,829 | | *ORMDL3* | 3.995 | 1.08E-04 | 0.014 |
| | | 38,096,333 | | | -3.664 | 3.60E-04 | 0.033 |

| | | | | | | | |
|---|---|---|---|---|---|---|---|
| | | 38,096,571 | | | -4.780 | 4.67E-06 | 0.033 |
| | | 38,096,578 | | | -5.642 | 1.00E-07 | 0.033 |
| | | 38,096,648 | | | -3.887 | 1.61E-04 | 0.033 |
| rs9911688 | g.37943801.T>C | 37,856,732 | | *ERBB2* | -3.668 | 3.56E-04 | 0.009 |
| | | 37,886,749 | | *MIEN1* | 3.945 | 1.30E-04 | 0.002 |
| | | 37,896,063 | | | 3.476 | 6.92E-04 | 0.009 |
| | | 38,020,485 | *IKZF3* | | 3.962 | 1.22E-04 | 0.005 |
| | | 38,080,829 | | *ORMDL3* | 3.995 | 1.08E-04 | 0.014 |
| | | 38,096,333 | | | -3.664 | 3.60E-04 | 0.033 |
| | | 38,096,571 | | | -4.780 | 4.67E-06 | 0.033 |
| | | 38,096,578 | | | -5.642 | 1.00E-07 | 0.033 |
| | | 38,096,648 | | | -3.887 | 1.61E-04 | 0.033 |
| rs111536507 | g.37944276.T>A | 37,886,788 | *MIEN1* | *MIEN1* | 4.149 | 6.00E-05 | 0.002 |
| | | 38,024,592 | | | 4.400 | 2.23E-05 | 0.046 |
| | | 38,024,606 | | | 4.304 | 3.27E-05 | 0.046 |
| | | 38,083,737 | | *ORMDL3* | 3.512 | 6.13E-04 | 0.011 |
| | | 38,096,333 | | | -3.527 | 5.80E-04 | 0.033 |
| | | 38,096,571 | | | -4.910 | 2.68E-06 | 0.033 |
| | | 38,096,578 | | | -4.517 | 1.39E-05 | 0.033 |
| rs34291217 | g.37944411.A>C | 37,856,732 | | *ERBB2* | -3.668 | 3.56E-04 | 0.009 |
| | | 37,886,749 | | *MIEN1* | 3.945 | 1.30E-04 | 0.002 |
| | | 37,896,063 | | | 3.476 | 6.92E-04 | 0.009 |
| | | 38,020,485 | *IKZF3* | | 3.962 | 1.22E-04 | 0.005 |
| | | 38,080,829 | | *ORMDL3* | 3.995 | 1.08E-04 | 0.014 |
| | | 38,096,333 | | | -3.664 | 3.60E-04 | 0.033 |
| | | 38,096,571 | | | -4.780 | 4.67E-06 | 0.033 |
| | | 38,096,578 | | | -5.642 | 1.00E-07 | 0.033 |
| | | 38,096,648 | | | -3.887 | 1.61E-04 | 0.033 |
| rs35088469 | g.37944482.T>C | 37,856,732 | | *ERBB2* | -3.668 | 3.56E-04 | 0.009 |
| | | 37,886,749 | | *MIEN1* | 3.945 | 1.30E-04 | 0.002 |
| | | 37,896,063 | | | 3.476 | 6.92E-04 | 0.009 |
| | | 38,020,485 | *IKZF3* | | 3.962 | 1.22E-04 | 0.005 |
| | | 38,080,829 | | *ORMDL3* | 3.995 | 1.08E-04 | 0.014 |
| | | 38,096,333 | | | -3.664 | 3.60E-04 | 0.033 |
| | | 38,096,571 | | | -4.780 | 4.67E-06 | 0.033 |
| | | 38,096,578 | | | -5.642 | 1.00E-07 | 0.033 |
| | | 38,096,648 | | | -3.887 | 1.61E-04 | 0.033 |
| rs112301322 | g.37944519.G>C (Gly.234.Ala) | 37,856,732 | | *ERBB2* | -3.668 | 3.56E-04 | 0.009 |
| | | 37,886,749 | | *MIEN1* | 3.945 | 1.30E-04 | 0.002 |
| | | 37,896,063 | | | 3.476 | 6.92E-04 | 0.009 |

| | | | | | | | |
|---|---|---|---|---|---|---|---|
| | | 38,020,485 | *IKZF3* | | 3.962 | 1.22E-04 | 0.005 |
| | | 38,080,829 | | *ORMDL3* | 3.995 | 1.08E-04 | 0.014 |
| | | 38,096,333 | | | -3.664 | 3.60E-04 | 0.033 |
| | | 38,096,571 | | | -4.780 | 4.67E-06 | 0.033 |
| | | 38,096,578 | | | -5.642 | 1.00E-07 | 0.033 |
| | | 38,096,648 | | | -3.887 | 1.61E-04 | 0.033 |
| rs112771646 | g.37945709.C>A | 37,856,732 | | *ERBB2* | -3.668 | 3.56E-04 | 0.009 |
| | | 37,886,749 | | *MIEN1* | 3.945 | 1.30E-04 | 0.002 |
| | | 37,896,063 | | | 3.476 | 6.92E-04 | 0.009 |
| | | 38,020,485 | *IKZF3* | | 3.962 | 1.22E-04 | 0.005 |
| | | 38,080,829 | | *ORMDL3* | 3.995 | 1.08E-04 | 0.014 |
| | | 38,096,333 | | | -3.664 | 3.60E-04 | 0.033 |
| | | 38,096,571 | | | -4.780 | 4.67E-06 | 0.033 |
| | | 38,096,578 | | | -5.642 | 1.00E-07 | 0.033 |
| | | 38,096,648 | | | -3.887 | 1.61E-04 | 0.033 |
| rs9913795 | g.37945841.T>C | 37,856,732 | | *ERBB2* | -3.668 | 3.56E-04 | 0.009 |
| | | 37,886,749 | | *MIEN1* | 3.945 | 1.30E-04 | 0.002 |
| | | 37,896,063 | | | 3.476 | 6.92E-04 | 0.009 |
| | | 38,020,485 | *IKZF3* | | 3.962 | 1.22E-04 | 0.005 |
| | | 38,080,829 | | *ORMDL3* | 3.995 | 1.08E-04 | 0.014 |
| | | 38,096,333 | | | -3.664 | 3.60E-04 | 0.033 |
| | | 38,096,571 | | | -4.780 | 4.67E-06 | 0.033 |
| | | 38,096,578 | | | -5.642 | 1.00E-07 | 0.033 |
| | | 38,096,648 | | | -3.887 | 1.61E-04 | 0.033 |
| rs9914516 | g.37946125.T>C | 37,856,732 | | *ERBB2* | -3.668 | 3.56E-04 | 0.009 |
| | | 37,886,749 | | *MIEN1* | 3.945 | 1.30E-04 | 0.002 |
| | | 37,896,063 | | | 3.476 | 6.92E-04 | 0.009 |
| | | 38,020,485 | *IKZF3* | | 3.962 | 1.22E-04 | 0.005 |
| | | 38,080,829 | | *ORMDL3* | 3.995 | 1.08E-04 | 0.014 |
| | | 38,096,333 | | | -3.664 | 3.60E-04 | 0.033 |
| | | 38,096,571 | | | -4.780 | 4.67E-06 | 0.033 |
| | | 38,096,578 | | | -5.642 | 1.00E-07 | 0.033 |
| | | 38,096,648 | | | -3.887 | 1.61E-04 | 0.033 |
| rs9913044 | g.37946228.A>G | 37,856,732 | | *ERBB2* | -3.668 | 3.56E-04 | 0.009 |
| | | 37,886,749 | | *MIEN1* | 3.945 | 1.30E-04 | 0.002 |
| | | 37,896,063 | | | 3.476 | 6.92E-04 | 0.009 |
| | | 38,020,485 | *IKZF3* | | 3.962 | 1.22E-04 | 0.005 |
| | | 38,080,829 | | *ORMDL3* | 3.995 | 1.08E-04 | 0.014 |
| | | 38,096,333 | | | -3.664 | 3.60E-04 | 0.033 |
| | | 38,096,571 | | | -4.780 | 4.67E-06 | 0.033 |
| | | 38,096,578 | | | -5.642 | 1.00E-07 | 0.033 |

| | | | | | | | |
|---|---|---|---|---|---|---|---|
| | | 38,096,648 | | | -3.887 | 1.61E-04 | 0.033 |
| rs71355416 | g.37946601_37946602del | 37,856,732 | | *ERBB2* | -3.668 | 3.56E-04 | 0.009 |
| | | 37,886,749 | | *MIEN1* | 3.945 | 1.30E-04 | 0.002 |
| | | 37,896,063 | | | 3.476 | 6.92E-04 | 0.009 |
| | | 38,020,485 | *IKZF3* | | 3.962 | 1.22E-04 | 0.005 |
| | | 38,080,829 | | *ORMDL3* | 3.995 | 1.08E-04 | 0.014 |
| | | 38,096,333 | | | -3.664 | 3.60E-04 | 0.033 |
| | | 38,096,571 | | | -4.780 | 4.67E-06 | 0.033 |
| | | 38,096,578 | | | -5.642 | 1.00E-07 | 0.033 |
| | | 38,096,648 | | | -3.887 | 1.61E-04 | 0.033 |
| rs35030093 | g.37946972_37946972insT | 37,896,063 | | | 3.508 | 6.21E-04 | 0.009 |
| | | 37,976,987 | | *IKZF3* | 3.405 | 8.81E-04 | 0.039 |
| | | 38,077,234 | | | -3.416 | 8.48E-04 | 0.014 |
| | | 38,080,829 | | *ORMDL3* | 4.051 | 8.70E-05 | 0.014 |
| | | 38,096,333 | | | -3.630 | 4.07E-04 | 0.033 |
| | | 38,096,571 | | | -4.161 | 5.73E-05 | 0.033 |
| | | 38,096,578 | | | -5.082 | 1.27E-06 | 0.033 |
| | | 38,096,648 | | | -3.741 | 2.74E-04 | 0.033 |
| rs117574970 | g.37947427.G>A | 37,910,190 | | | 3.537 | 5.61E-04 | 0.043 |
| | | 38,024,426 | | | -3.792 | 2.28E-04 | 3.75E-04 |
| rs35352075 | g.37949791.C>T | 37,856,732 | | *ERBB2* | -3.668 | 3.56E-04 | 0.009 |
| | | 37,886,749 | | *MIEN1* | 3.945 | 1.30E-04 | 0.002 |
| | | 37,896,063 | | | 3.476 | 6.92E-04 | 0.009 |
| | | 38,020,485 | *IKZF3* | | 3.962 | 1.22E-04 | 0.005 |
| | | 38,080,829 | | *ORMDL3* | 3.995 | 1.08E-04 | 0.014 |
| | | 38,096,333 | | | -3.664 | 3.60E-04 | 0.033 |
| | | 38,096,571 | | | -4.780 | 4.67E-06 | 0.033 |
| | | 38,096,578 | | | -5.642 | 1.00E-07 | 0.033 |
| | | 38,096,648 | | | -3.887 | 1.61E-04 | 0.033 |
| rs35105110 | g.37950422.A>G | 37,856,732 | | *ERBB2* | -3.668 | 3.56E-04 | 0.009 |
| | | 37,886,749 | | *MIEN1* | 3.945 | 1.30E-04 | 0.002 |
| | | 37,896,063 | | | 3.476 | 6.92E-04 | 0.009 |
| | | 38,020,485 | *IKZF3* | | 3.962 | 1.22E-04 | 0.005 |
| | | 38,080,829 | | *ORMDL3* | 3.995 | 1.08E-04 | 0.014 |
| | | 38,096,333 | | | -3.664 | 3.60E-04 | 0.033 |
| | | 38,096,571 | | | -4.780 | 4.67E-06 | 0.033 |
| | | 38,096,578 | | | -5.642 | 1.00E-07 | 0.033 |
| | | 38,096,648 | | | -3.887 | 1.61E-04 | 0.033 |
| rs35938199 | g.37950813.T>C | 37,856,732 | | *ERBB2* | -3.668 | 3.56E-04 | 0.009 |

| | | | | | | | |
|---|---|---|---|---|---|---|---|
| | | 37,886,749 | | *MIEN1* | 3.945 | 1.30E-04 | 0.002 |
| | | 37,896,063 | | | 3.476 | 6.92E-04 | 0.009 |
| | | 38,020,485 | *IKZF3* | | 3.962 | 1.22E-04 | 0.005 |
| | | 38,080,829 | | *ORMDL3* | 3.995 | 1.08E-04 | 0.014 |
| | | 38,096,333 | | | -3.664 | 3.60E-04 | 0.033 |
| | | 38,096,571 | | | -4.780 | 4.67E-06 | 0.033 |
| | | 38,096,578 | | | -5.642 | 1.00E-07 | 0.033 |
| | | 38,096,648 | | | -3.887 | 1.61E-04 | 0.033 |
| rs12938749 | g.37951848.T>C | 37,856,732 | | *ERBB2* | -3.668 | 3.56E-04 | 0.009 |
| | | 37,886,749 | | *MIEN1* | 3.945 | 1.30E-04 | 0.002 |
| | | 37,896,063 | | | 3.476 | 6.92E-04 | 0.009 |
| | | 38,020,485 | *IKZF3* | | 3.962 | 1.22E-04 | 0.005 |
| | | 38,080,829 | | *ORMDL3* | 3.995 | 1.08E-04 | 0.014 |
| | | 38,096,333 | | | -3.664 | 3.60E-04 | 0.033 |
| | | 38,096,571 | | | -4.780 | 4.67E-06 | 0.033 |
| | | 38,096,578 | | | -5.642 | 1.00E-07 | 0.033 |
| | | 38,096,648 | | | -3.887 | 1.61E-04 | 0.033 |
| rs56928975 | g.37952032.G>A | 37,856,732 | | *ERBB2* | -3.668 | 3.56E-04 | 0.009 |
| | | 37,886,749 | | *MIEN1* | 3.945 | 1.30E-04 | 0.002 |
| | | 37,896,063 | | | 3.476 | 6.92E-04 | 0.009 |
| | | 38,020,485 | *IKZF3* | | 3.962 | 1.22E-04 | 0.005 |
| | | 38,080,829 | | *ORMDL3* | 3.995 | 1.08E-04 | 0.014 |
| | | 38,096,333 | | | -3.664 | 3.60E-04 | 0.033 |
| | | 38,096,571 | | | -4.780 | 4.67E-06 | 0.033 |
| | | 38,096,578 | | | -5.642 | 1.00E-07 | 0.033 |
| | | 38,096,648 | | | -3.887 | 1.61E-04 | 0.033 |
| rs113159227 | g.37952092.A>G | 37,856,732 | | *ERBB2* | -3.668 | 3.56E-04 | 0.009 |
| | | 37,886,749 | | *MIEN1* | 3.945 | 1.30E-04 | 0.002 |
| | | 37,896,063 | | | 3.476 | 6.92E-04 | 0.009 |
| | | 38,020,485 | *IKZF3* | | 3.962 | 1.22E-04 | 0.005 |
| | | 38,080,829 | | *ORMDL3* | 3.995 | 1.08E-04 | 0.014 |
| | | 38,096,333 | | | -3.664 | 3.60E-04 | 0.033 |
| | | 38,096,571 | | | -4.780 | 4.67E-06 | 0.033 |
| | | 38,096,578 | | | -5.642 | 1.00E-07 | 0.033 |
| | | 38,096,648 | | | -3.887 | 1.61E-04 | 0.033 |
| rs73302152 | g.37952351.C>G | 37,856,732 | | *ERBB2* | -3.668 | 3.56E-04 | 0.009 |
| | | 37,886,749 | | *MIEN1* | 3.945 | 1.30E-04 | 0.002 |
| | | 37,896,063 | | | 3.476 | 6.92E-04 | 0.009 |
| | | 38,020,485 | *IKZF3* | | 3.962 | 1.22E-04 | 0.005 |
| | | 38,080,829 | | *ORMDL3* | 3.995 | 1.08E-04 | 0.014 |
| | | 38,096,333 | | | -3.664 | 3.60E-04 | 0.033 |

| | | | | | | | |
|---|---|---|---|---|---|---|---|
| | | 38,096,571 | | | -4.780 | 4.67E-06 | 0.033 |
| | | 38,096,578 | | | -5.642 | 1.00E-07 | 0.033 |
| | | 38,096,648 | | | -3.887 | 1.61E-04 | 0.033 |
| rs75148376 | g.37952509.T>C | 37,856,873 | | *ERBB2* | -4.032 | 9.35E-05 | 0.009 |
| | | 37,886,749 | | *MIEN1* | 5.501 | 1.93E-07 | 0.002 |
| | | 37,893,638 | | | -3.510 | 6.16E-04 | 0.002 |
| | | 37,894,352 | | | -4.012 | 1.01E-04 | 0.013 |
| | | 37,896,203 | | | 3.371 | 9.87E-04 | 0.011 |
| | | 38,080,829 | | *ORMDL3* | 4.552 | 1.21E-05 | 0.014 |
| | | 38,096,571 | | | -3.892 | 1.58E-04 | 0.033 |
| | | 38,096,578 | | | -3.664 | 3.60E-04 | 0.033 |
| rs113369293 | g.37952655.T>C | 37,856,732 | | *ERBB2* | -3.668 | 3.56E-04 | 0.009 |
| | | 37,886,749 | | *MIEN1* | 3.945 | 1.30E-04 | 0.002 |
| | | 37,896,063 | | | 3.476 | 6.92E-04 | 0.009 |
| | | 38,020,485 | *IKZF3* | | 3.962 | 1.22E-04 | 0.005 |
| | | 38,080,829 | | *ORMDL3* | 3.995 | 1.08E-04 | 0.014 |
| | | 38,096,333 | | | -3.664 | 3.60E-04 | 0.033 |
| | | 38,096,571 | | | -4.780 | 4.67E-06 | 0.033 |
| | | 38,096,578 | | | -5.642 | 1.00E-07 | 0.033 |
| | | 38,096,648 | | | -3.887 | 1.61E-04 | 0.033 |
| rs201873500 | g.37953666_37953670del | 37,886,788 | *MIEN1* | *MIEN1* | 4.083 | 7.71E-05 | 0.002 |
| | | 37,974,812 | | *IKZF3* | 3.466 | 7.16E-04 | 0.001 |
| | | 38,084,051 | *ORMDL3* | | 3.937 | 1.34E-04 | 0.005 |
| | | 38,096,571 | | | -3.877 | 1.67E-04 | 0.033 |
| | | 38,096,578 | | | -3.665 | 3.59E-04 | 0.033 |
| rs13313561 | g.37954451.C>G | 37,856,732 | | *ERBB2* | -3.668 | 3.56E-04 | 0.009 |
| | | 37,886,749 | | *MIEN1* | 3.945 | 1.30E-04 | 0.002 |
| | | 37,896,063 | | | 3.476 | 6.92E-04 | 0.009 |
| | | 38,020,485 | *IKZF3* | | 3.962 | 1.22E-04 | 0.005 |
| | | 38,080,829 | | *ORMDL3* | 3.995 | 1.08E-04 | 0.014 |
| | | 38,096,333 | | | -3.664 | 3.60E-04 | 0.033 |
| | | 38,096,571 | | | -4.780 | 4.67E-06 | 0.033 |
| | | 38,096,578 | | | -5.642 | 1.00E-07 | 0.033 |
| | | 38,096,648 | | | -3.887 | 1.61E-04 | 0.033 |
| rs34137841 | g.37954533_37954534del | 37,856,732 | | *ERBB2* | -3.668 | 3.56E-04 | 0.009 |
| | | 37,886,749 | | *MIEN1* | 3.945 | 1.30E-04 | 0.002 |
| | | 37,896,063 | | | 3.476 | 6.92E-04 | 0.009 |
| | | 38,020,485 | *IKZF3* | | 3.962 | 1.22E-04 | 0.005 |
| | | 38,080,829 | | *ORMDL3* | 3.995 | 1.08E-04 | 0.014 |

| | | | | | | | |
|---|---|---|---|---|---|---|---|
| | | 38,096,333 | | | -3.664 | 3.60E-04 | 0.033 |
| | | 38,096,571 | | | -4.780 | 4.67E-06 | 0.033 |
| | | 38,096,578 | | | -5.642 | 1.00E-07 | 0.033 |
| | | 38,096,648 | | | -3.887 | 1.61E-04 | 0.033 |
| rs9899345 | g.37954758.A>G | 37,856,732 | | *ERBB2* | -3.668 | 3.56E-04 | 0.009 |
| | | 37,886,749 | | *MIEN1* | 3.945 | 1.30E-04 | 0.002 |
| | | 37,896,063 | | | 3.476 | 6.92E-04 | 0.009 |
| | | 38,020,485 | *IKZF3* | | 3.962 | 1.22E-04 | 0.005 |
| | | 38,080,829 | | *ORMDL3* | 3.995 | 1.08E-04 | 0.014 |
| | | 38,096,333 | | | -3.664 | 3.60E-04 | 0.033 |
| | | 38,096,571 | | | -4.780 | 4.67E-06 | 0.033 |
| | | 38,096,578 | | | -5.642 | 1.00E-07 | 0.033 |
| | | 38,096,648 | | | -3.887 | 1.61E-04 | 0.033 |
| rs34516015 | g.37954798.A>T | 37,856,732 | | *ERBB2* | -3.668 | 3.56E-04 | 0.009 |
| | | 37,886,749 | | *MIEN1* | 3.945 | 1.30E-04 | 0.002 |
| | | 37,896,063 | | | 3.476 | 6.92E-04 | 0.009 |
| | | 38,020,485 | *IKZF3* | | 3.962 | 1.22E-04 | 0.005 |
| | | 38,080,829 | | *ORMDL3* | 3.995 | 1.08E-04 | 0.014 |
| | | 38,096,333 | | | -3.664 | 3.60E-04 | 0.033 |
| | | 38,096,571 | | | -4.780 | 4.67E-06 | 0.033 |
| | | 38,096,578 | | | -5.642 | 1.00E-07 | 0.033 |
| | | 38,096,648 | | | -3.887 | 1.61E-04 | 0.033 |
| rs34876670 | g.37954800.C>T | 37,856,732 | | *ERBB2* | -3.668 | 3.56E-04 | 0.009 |
| | | 37,886,749 | | *MIEN1* | 3.945 | 1.30E-04 | 0.002 |
| | | 37,896,063 | | | 3.476 | 6.92E-04 | 0.009 |
| | | 38,020,485 | *IKZF3* | | 3.962 | 1.22E-04 | 0.005 |
| | | 38,080,829 | | *ORMDL3* | 3.995 | 1.08E-04 | 0.014 |
| | | 38,096,333 | | | -3.664 | 3.60E-04 | 0.033 |
| | | 38,096,571 | | | -4.780 | 4.67E-06 | 0.033 |
| | | 38,096,578 | | | -5.642 | 1.00E-07 | 0.033 |
| | | 38,096,648 | | | -3.887 | 1.61E-04 | 0.033 |
| rs9907988 | g.37954883.C>T | 37,856,732 | | *ERBB2* | -3.668 | 3.56E-04 | 0.009 |
| | | 37,886,749 | | *MIEN1* | 3.945 | 1.30E-04 | 0.002 |
| | | 37,896,063 | | | 3.476 | 6.92E-04 | 0.009 |
| | | 38,020,485 | *IKZF3* | | 3.962 | 1.22E-04 | 0.005 |
| | | 38,080,829 | | *ORMDL3* | 3.995 | 1.08E-04 | 0.014 |
| | | 38,096,333 | | | -3.664 | 3.60E-04 | 0.033 |
| | | 38,096,571 | | | -4.780 | 4.67E-06 | 0.033 |
| | | 38,096,578 | | | -5.642 | 1.00E-07 | 0.033 |
| | | 38,096,648 | | | -3.887 | 1.61E-04 | 0.033 |
| rs149214511 | g.37955019.G>A | 37,894,397 | | | 3.473 | 6.99E-04 | 0.013 |

| | | | | | | | |
|---|---|---|---|---|---|---|---|
| | | 38,024,063 | | | -3.617 | 4.25E-04 | 0.004 |
| | | 38,024,146 | | | -3.985 | 1.12E-04 | 0.004 |
| | | 38,024,189 | | | -4.393 | 2.30E-05 | 0.016 |
| | | 38,024,317 | | | -3.506 | 6.26E-04 | 0.023 |
| | | 38,024,353 | | | -3.614 | 4.29E-04 | 0.017 |
| | | 38,075,656 | *GSDMB* | | -3.703 | 3.13E-04 | 0.034 |
| rs34344462 | g.37955194.G>A | 37,879,682 | | *ERBB2* | 3.639 | 3.93E-04 | 0.002 |
| | | 37,896,063 | | | 3.755 | 2.60E-04 | 0.009 |
| | | 38,083,269 | | *ORMDL3* | 3.399 | 8.99E-04 | 0.011 |
| | | 38,096,333 | | | -3.520 | 5.95E-04 | 0.033 |
| | | 38,096,578 | | | -4.078 | 7.88E-05 | 0.033 |
| rs13313564 | g.37955555.A>G | 37,856,732 | | *ERBB2* | -3.668 | 3.56E-04 | 0.009 |
| | | 37,886,749 | | *MIEN1* | 3.945 | 1.30E-04 | 0.002 |
| | | 37,896,063 | | | 3.476 | 6.92E-04 | 0.009 |
| | | 38,020,485 | *IKZF3* | | 3.962 | 1.22E-04 | 0.005 |
| | | 38,080,829 | | *ORMDL3* | 3.995 | 1.08E-04 | 0.014 |
| | | 38,096,333 | | | -3.664 | 3.60E-04 | 0.033 |
| | | 38,096,571 | | | -4.780 | 4.67E-06 | 0.033 |
| | | 38,096,578 | | | -5.642 | 1.00E-07 | 0.033 |
| | | 38,096,648 | | | -3.887 | 1.61E-04 | 0.033 |
| rs13313573 | g.37955818.T>G | 37,856,732 | | *ERBB2* | -3.668 | 3.56E-04 | 0.009 |
| | | 37,886,749 | | *MIEN1* | 3.945 | 1.30E-04 | 0.002 |
| | | 37,896,063 | | | 3.476 | 6.92E-04 | 0.009 |
| | | 38,020,485 | *IKZF3* | | 3.962 | 1.22E-04 | 0.005 |
| | | 38,080,829 | | *ORMDL3* | 3.995 | 1.08E-04 | 0.014 |
| | | 38,096,333 | | | -3.664 | 3.60E-04 | 0.033 |
| | | 38,096,571 | | | -4.780 | 4.67E-06 | 0.033 |
| | | 38,096,578 | | | -5.642 | 1.00E-07 | 0.033 |
| | | 38,096,648 | | | -3.887 | 1.61E-04 | 0.033 |
| rs1563104 | | 37,894,916 | | | 3.861 | 1.77E-04 | 0.004 |
| | | 37,896,708 | | | 4.821 | 3.93E-06 | 0.008 |
| | | 38,020,618 | *IKZF3* | | 3.400 | 8.95E-04 | 0.017 |
| | | 38,024,502 | | | -4.416 | 2.10E-05 | 0.005 |
| | | 38,024,505 | | | -3.441 | 7.79E-04 | 0.005 |
| | | 38,024,606 | | | 3.913 | 1.46E-04 | 0.046 |
| | | 38,024,614 | | | 3.658 | 3.68E-04 | 0.046 |
| | | 38,109,131 | | | 6.420 | 2.34E-09 | 0.002 |
| | | 38,115,470 | | | 4.754 | 5.21E-06 | 0.009 |
| rs17607816 | g.37957235.C>T | 37,828,773 | | *PGAP3* | 3.405 | 8.80E-04 | 0.002 |
| | | 37,856,857 | | *ERBB2* | -3.409 | 8.68E-04 | 0.009 |
| | | 37,886,765 | | *MIEN1* | 3.518 | 6.00E-04 | 0.002 |

| | | | | | | | |
|---|---|---|---|---|---|---|---|
| | | 37,886,921 | *MIEN1* | | 3.552 | 5.33E-04 | 0.002 |
| | | 37,896,050 | | | 3.683 | 3.37E-04 | 0.027 |
| | | 37,910,406 | | | 3.737 | 2.78E-04 | 0.013 |
| | | 37,910,697 | | | 3.384 | 9.44E-04 | 0.035 |
| | | 37,910,840 | | | 3.970 | 1.18E-04 | 0.035 |
| | | 37,911,360 | | | 3.408 | 8.72E-04 | 0.035 |
| | | 37,975,698 | | *IKZF3* | 5.196 | 7.67E-07 | 0.039 |
| | | 38,020,483 | *IKZF3* | | 4.621 | 9.10E-06 | 0.005 |
| | | 38,024,250 | | | -4.882 | 3.03E-06 | 0.016 |
| | | 38,075,557 | *GSDMB* | | 5.039 | 1.53E-06 | 0.034 |
| | | 38,077,234 | | | -3.420 | 8.36E-04 | 0.014 |
| | | 38,083,520 | | *ORMDL3* | 3.533 | 5.70E-04 | 0.011 |
| | | 38,083,862 | | *ORMDL3* | 3.701 | 3.16E-04 | 0.011 |
| | | 38,108,734 | | | 4.222 | 4.52E-05 | 0.039 |
| | | 38,119,132 | *GSDMA* | | 3.964 | 1.21E-04 | 0.009 |
| rs12937330 | g.37957317.A>C | 37,856,732 | | *ERBB2* | -3.668 | 3.56E-04 | 0.009 |
| | | 37,886,749 | | *MIEN1* | 3.945 | 1.30E-04 | 0.002 |
| | | 37,896,063 | | | 3.476 | 6.92E-04 | 0.009 |
| | | 38,020,485 | *IKZF3* | | 3.962 | 1.22E-04 | 0.005 |
| | | 38,080,829 | | *ORMDL3* | 3.995 | 1.08E-04 | 0.014 |
| | | 38,096,333 | | | -3.664 | 3.60E-04 | 0.033 |
| | | 38,096,571 | | | -4.780 | 4.67E-06 | 0.033 |
| | | 38,096,578 | | | -5.642 | 1.00E-07 | 0.033 |
| | | 38,096,648 | | | -3.887 | 1.61E-04 | 0.033 |
| rs34988504 | g.37957632.T>C | 37,856,732 | | *ERBB2* | -3.668 | 3.56E-04 | 0.009 |
| | | 37,886,749 | | *MIEN1* | 3.945 | 1.30E-04 | 0.002 |
| | | 37,896,063 | | | 3.476 | 6.92E-04 | 0.009 |
| | | 38,020,485 | *IKZF3* | | 3.962 | 1.22E-04 | 0.005 |
| | | 38,080,829 | | *ORMDL3* | 3.995 | 1.08E-04 | 0.014 |
| | | 38,096,333 | | | -3.664 | 3.60E-04 | 0.033 |
| | | 38,096,571 | | | -4.780 | 4.67E-06 | 0.033 |
| | | 38,096,578 | | | -5.642 | 1.00E-07 | 0.033 |
| | | 38,096,648 | | | -3.887 | 1.61E-04 | 0.033 |
| rs36097841 | g.37958113.A>G | 37,856,732 | | *ERBB2* | -3.668 | 3.56E-04 | 0.009 |
| | | 37,886,749 | | *MIEN1* | 3.945 | 1.30E-04 | 0.002 |
| | | 37,896,063 | | | 3.476 | 6.92E-04 | 0.009 |
| | | 38,020,485 | *IKZF3* | | 3.962 | 1.22E-04 | 0.005 |
| | | 38,080,829 | | *ORMDL3* | 3.995 | 1.08E-04 | 0.014 |
| | | 38,096,333 | | | -3.664 | 3.60E-04 | 0.033 |
| | | 38,096,571 | | | -4.780 | 4.67E-06 | 0.033 |
| | | 38,096,578 | | | -5.642 | 1.00E-07 | 0.033 |

| | | | | | | | |
|---|---|---|---|---|---|---|---|
| | | 38,096,648 | | | -3.887 | 1.61E-04 | 0.033 |
| rs7211998 | g.37959789.G>A | 37,856,732 | | *ERBB2* | -3.668 | 3.56E-04 | 0.009 |
| | | 37,886,749 | | *MIEN1* | 3.945 | 1.30E-04 | 0.002 |
| | | 37,896,063 | | | 3.476 | 6.92E-04 | 0.009 |
| | | 38,020,485 | *IKZF3* | | 3.962 | 1.22E-04 | 0.005 |
| | | 38,080,829 | | *ORMDL3* | 3.995 | 1.08E-04 | 0.014 |
| | | 38,096,333 | | | -3.664 | 3.60E-04 | 0.033 |
| | | 38,096,571 | | | -4.780 | 4.67E-06 | 0.033 |
| | | 38,096,578 | | | -5.642 | 1.00E-07 | 0.033 |
| | | 38,096,648 | | | -3.887 | 1.61E-04 | 0.033 |
| rs113064843 | g.37960422.C>T | 37,856,732 | | *ERBB2* | -3.668 | 3.56E-04 | 0.009 |
| | | 37,886,749 | | *MIEN1* | 3.945 | 1.30E-04 | 0.002 |
| | | 37,896,063 | | | 3.476 | 6.92E-04 | 0.009 |
| | | 38,020,485 | *IKZF3* | | 3.962 | 1.22E-04 | 0.005 |
| | | 38,080,829 | | *ORMDL3* | 3.995 | 1.08E-04 | 0.014 |
| | | 38,096,333 | | | -3.664 | 3.60E-04 | 0.033 |
| | | 38,096,571 | | | -4.780 | 4.67E-06 | 0.033 |
| | | 38,096,578 | | | -5.642 | 1.00E-07 | 0.033 |
| | | 38,096,648 | | | -3.887 | 1.61E-04 | 0.033 |
| rs9894680 | g.37962237.C>T | 37,856,732 | | *ERBB2* | -3.668 | 3.56E-04 | 0.009 |
| | | 37,886,749 | | *MIEN1* | 3.945 | 1.30E-04 | 0.002 |
| | | 37,896,063 | | | 3.476 | 6.92E-04 | 0.009 |
| | | 38,020,485 | *IKZF3* | | 3.962 | 1.22E-04 | 0.005 |
| | | 38,080,829 | | *ORMDL3* | 3.995 | 1.08E-04 | 0.014 |
| | | 38,096,333 | | | -3.664 | 3.60E-04 | 0.033 |
| | | 38,096,571 | | | -4.780 | 4.67E-06 | 0.033 |
| | | 38,096,578 | | | -5.642 | 1.00E-07 | 0.033 |
| | | 38,096,648 | | | -3.887 | 1.61E-04 | 0.033 |
| rs9914302 | g.37962408.A>G | 37,856,732 | | *ERBB2* | -3.668 | 3.56E-04 | 0.009 |
| | | 37,886,749 | | *MIEN1* | 3.945 | 1.30E-04 | 0.002 |
| | | 37,896,063 | | | 3.476 | 6.92E-04 | 0.009 |
| | | 38,020,485 | *IKZF3* | | 3.962 | 1.22E-04 | 0.005 |
| | | 38,080,829 | | *ORMDL3* | 3.995 | 1.08E-04 | 0.014 |
| | | 38,096,333 | | | -3.664 | 3.60E-04 | 0.033 |
| | | 38,096,571 | | | -4.780 | 4.67E-06 | 0.033 |
| | | 38,096,578 | | | -5.642 | 1.00E-07 | 0.033 |
| | | 38,096,648 | | | -3.887 | 1.61E-04 | 0.033 |
| rs79989390 | g.37962845.A>G | 37,856,732 | | *ERBB2* | -3.668 | 3.56E-04 | 0.009 |
| | | 37,886,749 | | *MIEN1* | 3.945 | 1.30E-04 | 0.002 |
| | | 37,896,063 | | | 3.476 | 6.92E-04 | 0.009 |
| | | 38,020,485 | *IKZF3* | | 3.962 | 1.22E-04 | 0.005 |

| | | | | | | | |
|---|---|---|---|---|---|---|---|
| | | 38,080,829 | | *ORMDL3* | 3.995 | 1.08E-04 | 0.014 |
| | | 38,096,333 | | | -3.664 | 3.60E-04 | 0.033 |
| | | 38,096,571 | | | -4.780 | 4.67E-06 | 0.033 |
| | | 38,096,578 | | | -5.642 | 1.00E-07 | 0.033 |
| | | 38,096,648 | | | -3.887 | 1.61E-04 | 0.033 |
| rs77232433 | g.37962846.T>C | 37,856,732 | | *ERBB2* | -3.668 | 3.56E-04 | 0.009 |
| | | 37,886,749 | | *MIEN1* | 3.945 | 1.30E-04 | 0.002 |
| | | 37,896,063 | | | 3.476 | 6.92E-04 | 0.009 |
| | | 38,020,485 | *IKZF3* | | 3.962 | 1.22E-04 | 0.005 |
| | | 38,080,829 | | *ORMDL3* | 3.995 | 1.08E-04 | 0.014 |
| | | 38,096,333 | | | -3.664 | 3.60E-04 | 0.033 |
| | | 38,096,571 | | | -4.780 | 4.67E-06 | 0.033 |
| | | 38,096,578 | | | -5.642 | 1.00E-07 | 0.033 |
| | | 38,096,648 | | | -3.887 | 1.61E-04 | 0.033 |
| rs34860387 | g.37962889_37962889ins G | 37,894,916 | | | 3.388 | 9.33E-04 | 0.004 |
| | | 37,934,107 | | *IKZF3* | -4.486 | 1.58E-05 | 0.027 |
| | | 38,084,016 | *ORMDL3* | | 4.474 | 1.66E-05 | 0.011 |
| | | 38,109,131 | | | 4.746 | 5.39E-06 | 0.002 |
| rs151330975 | g.37963506.T>C | 37,886,755 | | *MIEN1* | 3.707 | 3.09E-04 | 0.002 |
| | | 37,886,872 | *MIEN1* | | 5.079 | 1.28E-06 | 0.002 |
| | | 37,886,883 | *MIEN1* | | 3.425 | 8.23E-04 | 0.002 |
| | | 37,886,921 | *MIEN1* | | 5.093 | 1.21E-06 | 0.002 |
| | | 37,894,469 | | | -5.578 | 1.35E-07 | 0.013 |
| | | 37,894,530 | | | -4.833 | 3.73E-06 | 0.013 |
| | | 37,910,294 | | | 3.526 | 5.83E-04 | 0.043 |
| | | 37,910,882 | | | 4.551 | 1.21E-05 | 0.035 |
| | | 37,911,096 | | | 4.159 | 5.77E-05 | 0.035 |
| | | 38,022,345 | | | 3.942 | 1.31E-04 | 0.003 |
| | | 38,022,353 | | | 7.270 | 2.99E-11 | 0.003 |
| | | 38,083,257 | | *ORMDL3* | 3.784 | 2.35E-04 | 0.011 |
| | | 38,083,556 | | *ORMDL3* | 4.599 | 9.93E-06 | 0.011 |
| | | 38,083,891 | *ORMDL3* | | 4.022 | 9.73E-05 | 0.011 |
| | | 38,083,958 | *ORMDL3* | | 3.799 | 2.22E-04 | 0.011 |
| | | 38,109,541 | | | 5.770 | 5.52E-08 | 0.002 |
| | | 38,109,595 | | | 4.038 | 9.17E-05 | 0.002 |
| rs9901617 | g.37964176.C>G | 37,856,732 | | *ERBB2* | -3.668 | 3.56E-04 | 0.009 |
| | | 37,886,749 | | *MIEN1* | 3.945 | 1.30E-04 | 0.002 |
| | | 37,896,063 | | | 3.476 | 6.92E-04 | 0.009 |
| | | 38,020,485 | *IKZF3* | | 3.962 | 1.22E-04 | 0.005 |
| | | 38,080,829 | | *ORMDL3* | 3.995 | 1.08E-04 | 0.014 |

| | | | | | | | |
|---|---|---|---|---|---|---|---|
| | | 38,096,333 | | | -3.664 | 3.60E-04 | 0.033 |
| | | 38,096,571 | | | -4.780 | 4.67E-06 | 0.033 |
| | | 38,096,578 | | | -5.642 | 1.00E-07 | 0.033 |
| | | 38,096,648 | | | -3.887 | 1.61E-04 | 0.033 |
| rs4337325 | g.37964436.T>C | 37,856,732 | | *ERBB2* | -3.668 | 3.56E-04 | 0.009 |
| | | 37,886,749 | | *MIEN1* | 3.945 | 1.30E-04 | 0.002 |
| | | 37,896,063 | | | 3.476 | 6.92E-04 | 0.009 |
| | | 38,020,485 | *IKZF3* | | 3.962 | 1.22E-04 | 0.005 |
| | | 38,080,829 | | *ORMDL3* | 3.995 | 1.08E-04 | 0.014 |
| | | 38,096,333 | | | -3.664 | 3.60E-04 | 0.033 |
| | | 38,096,571 | | | -4.780 | 4.67E-06 | 0.033 |
| | | 38,096,578 | | | -5.642 | 1.00E-07 | 0.033 |
| | | 38,096,648 | | | -3.887 | 1.61E-04 | 0.033 |
| rs28580691 | g.37965567.C>T | 37,856,732 | | *ERBB2* | -3.668 | 3.56E-04 | 0.009 |
| | | 37,886,749 | | *MIEN1* | 3.945 | 1.30E-04 | 0.002 |
| | | 37,896,063 | | | 3.476 | 6.92E-04 | 0.009 |
| | | 38,020,485 | *IKZF3* | | 3.962 | 1.22E-04 | 0.005 |
| | | 38,080,829 | | *ORMDL3* | 3.995 | 1.08E-04 | 0.014 |
| | | 38,096,333 | | | -3.664 | 3.60E-04 | 0.033 |
| | | 38,096,571 | | | -4.780 | 4.67E-06 | 0.033 |
| | | 38,096,578 | | | -5.642 | 1.00E-07 | 0.033 |
| | | 38,096,648 | | | -3.887 | 1.61E-04 | 0.033 |
| rs79026872 | g.37965932.C>T | 37,910,260 | | | 3.665 | 3.60E-04 | 0.043 |
| | | 37,911,152 | | | 3.607 | 4.41E-04 | 0.035 |
| rs9892781 | g.37967438.C>T | 37,856,732 | | *ERBB2* | -3.668 | 3.56E-04 | 0.009 |
| | | 37,886,749 | | *MIEN1* | 3.945 | 1.30E-04 | 0.002 |
| | | 37,896,063 | | | 3.476 | 6.92E-04 | 0.009 |
| | | 38,020,485 | *IKZF3* | | 3.962 | 1.22E-04 | 0.005 |
| | | 38,080,829 | | *ORMDL3* | 3.995 | 1.08E-04 | 0.014 |
| | | 38,096,333 | | | -3.664 | 3.60E-04 | 0.033 |
| | | 38,096,571 | | | -4.780 | 4.67E-06 | 0.033 |
| | | 38,096,578 | | | -5.642 | 1.00E-07 | 0.033 |
| | | 38,096,648 | | | -3.887 | 1.61E-04 | 0.033 |
| rs9914757 | g.37967562.G>A | 37,856,732 | | *ERBB2* | -3.668 | 3.56E-04 | 0.009 |
| | | 37,886,749 | | *MIEN1* | 3.945 | 1.30E-04 | 0.002 |
| | | 37,896,063 | | | 3.476 | 6.92E-04 | 0.009 |
| | | 38,020,485 | *IKZF3* | | 3.962 | 1.22E-04 | 0.005 |
| | | 38,080,829 | | *ORMDL3* | 3.995 | 1.08E-04 | 0.014 |
| | | 38,096,333 | | | -3.664 | 3.60E-04 | 0.033 |
| | | 38,096,571 | | | -4.780 | 4.67E-06 | 0.033 |
| | | 38,096,578 | | | -5.642 | 1.00E-07 | 0.033 |

| | | | | | | | |
|---|---|---|---|---|---|---|---|
| | | 38,096,648 | | | -3.887 | 1.61E-04 | 0.033 |
| rs558811024 | g.37967634_37967650del | 37,856,732 | | *ERBB2* | -3.668 | 3.56E-04 | 0.009 |
| | | 37,886,749 | | *MIEN1* | 3.945 | 1.30E-04 | 0.002 |
| | | 37,896,063 | | | 3.476 | 6.92E-04 | 0.009 |
| | | 38,020,485 | *IKZF3* | | 3.962 | 1.22E-04 | 0.005 |
| | | 38,080,829 | | *ORMDL3* | 3.995 | 1.08E-04 | 0.014 |
| | | 38,096,333 | | | -3.664 | 3.60E-04 | 0.033 |
| | | 38,096,571 | | | -4.780 | 4.67E-06 | 0.033 |
| | | 38,096,578 | | | -5.642 | 1.00E-07 | 0.033 |
| | | 38,096,648 | | | -3.887 | 1.61E-04 | 0.033 |
| rs536870805 | g.37967644.G>A | 37,834,864 | | *PGAP3* | 3.530 | 5.74E-04 | 0.020 |
| | | 37,886,749 | | *MIEN1* | 4.574 | 1.10E-05 | 0.002 |
| | | 37,896,006 | | | 3.419 | 8.39E-04 | 0.027 |
| | | 37,910,177 | | | 3.413 | 8.56E-04 | 0.043 |
| | | 38,020,485 | *IKZF3* | | 4.732 | 5.72E-06 | 0.005 |
| | | 38,080,829 | | *ORMDL3* | 3.568 | 5.05E-04 | 0.014 |
| | | 38,096,333 | | | -3.536 | 5.64E-04 | 0.033 |
| | | 38,096,571 | | | -4.852 | 3.44E-06 | 0.033 |
| | | 38,096,578 | | | -5.932 | 2.54E-08 | 0.033 |
| | | 38,096,648 | | | -4.480 | 1.62E-05 | 0.033 |
| rs114777282 | g.37967650.A>C | 37,856,732 | | *ERBB2* | -3.706 | 3.10E-04 | 0.009 |
| | | 37,879,682 | | *ERBB2* | 3.563 | 5.14E-04 | 0.002 |
| | | 37,896,063 | | | 3.800 | 2.21E-04 | 0.009 |
| | | 37,903,448 | | | 3.534 | 5.67E-04 | 0.008 |
| | | 38,024,589 | | | 3.628 | 4.10E-04 | 0.046 |
| | | 38,076,680 | | | 3.575 | 4.92E-04 | 0.034 |
| | | 38,083,269 | | *ORMDL3* | 3.753 | 2.63E-04 | 0.011 |
| | | 38,096,333 | | | -3.375 | 9.73E-04 | 0.033 |
| | | 38,096,578 | | | -4.025 | 9.62E-05 | 0.033 |
| rs67135646 | g.37967872.G>C | 37,856,732 | | *ERBB2* | -3.668 | 3.56E-04 | 0.009 |
| | | 37,886,749 | | *MIEN1* | 3.945 | 1.30E-04 | 0.002 |
| | | 37,896,063 | | | 3.476 | 6.92E-04 | 0.009 |
| | | 38,020,485 | *IKZF3* | | 3.962 | 1.22E-04 | 0.005 |
| | | 38,080,829 | | *ORMDL3* | 3.995 | 1.08E-04 | 0.014 |
| | | 38,096,333 | | | -3.664 | 3.60E-04 | 0.033 |
| | | 38,096,571 | | | -4.780 | 4.67E-06 | 0.033 |
| | | 38,096,578 | | | -5.642 | 1.00E-07 | 0.033 |
| | | 38,096,648 | | | -3.887 | 1.61E-04 | 0.033 |
| rs112238900 | g.37968495.T>C | 37,856,732 | | *ERBB2* | -3.668 | 3.56E-04 | 0.009 |
| | | 37,886,749 | | *MIEN1* | 3.945 | 1.30E-04 | 0.002 |

| | | | | | | | |
|---|---|---|---|---|---|---|---|
| | | 37,896,063 | | | 3.476 | 6.92E-04 | 0.009 |
| | | 38,020,485 | *IKZF3* | | 3.962 | 1.22E-04 | 0.005 |
| | | 38,080,829 | | *ORMDL3* | 3.995 | 1.08E-04 | 0.014 |
| | | 38,096,333 | | | -3.664 | 3.60E-04 | 0.033 |
| | | 38,096,571 | | | -4.780 | 4.67E-06 | 0.033 |
| | | 38,096,578 | | | -5.642 | 1.00E-07 | 0.033 |
| | | 38,096,648 | | | -3.887 | 1.61E-04 | 0.033 |
| rs111307979 | g.37969363.G>A | 37,879,682 | | *ERBB2* | 3.904 | 1.51E-04 | 0.002 |
| | | 37,896,063 | | | 4.195 | 5.02E-05 | 0.009 |
| | | 38,083,269 | | *ORMDL3* | 4.154 | 5.87E-05 | 0.011 |
| | | 38,108,720 | | | 4.104 | 7.12E-05 | 0.039 |
| | | 38,119,834 | | *GSDMA* | 3.403 | 8.85E-04 | 0.011 |
| rs534820301 | g.37969371.G>A | 37,856,732 | | *ERBB2* | -3.668 | 3.56E-04 | 0.009 |
| | | 37,886,749 | | *MIEN1* | 3.945 | 1.30E-04 | 0.002 |
| | | 37,896,063 | | | 3.476 | 6.92E-04 | 0.009 |
| | | 38,020,485 | *IKZF3* | | 3.962 | 1.22E-04 | 0.005 |
| | | 38,080,829 | | *ORMDL3* | 3.995 | 1.08E-04 | 0.014 |
| | | 38,096,333 | | | -3.664 | 3.60E-04 | 0.033 |
| | | 38,096,571 | | | -4.780 | 4.67E-06 | 0.033 |
| | | 38,096,578 | | | -5.642 | 1.00E-07 | 0.033 |
| | | 38,096,648 | | | -3.887 | 1.61E-04 | 0.033 |
| rs12943633 | g.37970365.T>C | 37,828,560 | | *PGAP3* | -3.584 | 4.78E-04 | 0.002 |
| | | 37,856,732 | | *ERBB2* | -3.530 | 5.75E-04 | 0.009 |
| | | 37,886,749 | | *MIEN1* | 4.365 | 2.57E-05 | 0.002 |
| | | 37,894,413 | | | -3.750 | 2.65E-04 | 0.013 |
| | | 38,020,485 | *IKZF3* | | 3.653 | 3.75E-04 | 0.005 |
| | | 38,024,589 | | | 3.797 | 2.24E-04 | 0.046 |
| | | 38,076,680 | | | 3.932 | 1.36E-04 | 0.034 |
| | | 38,096,578 | | | -4.064 | 8.29E-05 | 0.033 |
| rs113115305 | g.37970687.C>A | 37,856,732 | | *ERBB2* | -3.668 | 3.56E-04 | 0.009 |
| | | 37,886,749 | | *MIEN1* | 3.945 | 1.30E-04 | 0.002 |
| | | 37,896,063 | | | 3.476 | 6.92E-04 | 0.009 |
| | | 38,020,485 | *IKZF3* | | 3.962 | 1.22E-04 | 0.005 |
| | | 38,080,829 | | *ORMDL3* | 3.995 | 1.08E-04 | 0.014 |
| | | 38,096,333 | | | -3.664 | 3.60E-04 | 0.033 |
| | | 38,096,571 | | | -4.780 | 4.67E-06 | 0.033 |
| | | 38,096,578 | | | -5.642 | 1.00E-07 | 0.033 |
| | | 38,096,648 | | | -3.887 | 1.61E-04 | 0.033 |
| rs12952146 | g.37970903.T>C | 37,856,732 | | *ERBB2* | -3.668 | 3.56E-04 | 0.009 |
| | | 37,886,749 | | *MIEN1* | 3.945 | 1.30E-04 | 0.002 |
| | | 37,896,063 | | | 3.476 | 6.92E-04 | 0.009 |

| | | | | | | | |
|---|---|---|---|---|---|---|---|
| | | 38,020,485 | *IKZF3* | | 3.962 | 1.22E-04 | 0.005 |
| | | 38,080,829 | | *ORMDL3* | 3.995 | 1.08E-04 | 0.014 |
| | | 38,096,333 | | | -3.664 | 3.60E-04 | 0.033 |
| | | 38,096,571 | | | -4.780 | 4.67E-06 | 0.033 |
| | | 38,096,578 | | | -5.642 | 1.00E-07 | 0.033 |
| | | 38,096,648 | | | -3.887 | 1.61E-04 | 0.033 |
| rs112412105 | g.37971636.G>A | 37,856,732 | | *ERBB2* | -3.668 | 3.56E-04 | 0.009 |
| | | 37,886,749 | | *MIEN1* | 3.945 | 1.30E-04 | 0.002 |
| | | 37,896,063 | | | 3.476 | 6.92E-04 | 0.009 |
| | | 38,020,485 | *IKZF3* | | 3.962 | 1.22E-04 | 0.005 |
| | | 38,080,829 | | *ORMDL3* | 3.995 | 1.08E-04 | 0.014 |
| | | 38,096,333 | | | -3.664 | 3.60E-04 | 0.033 |
| | | 38,096,571 | | | -4.780 | 4.67E-06 | 0.033 |
| | | 38,096,578 | | | -5.642 | 1.00E-07 | 0.033 |
| | | 38,096,648 | | | -3.887 | 1.61E-04 | 0.033 |
| rs9898031 | g.37972648.G>C | 37,856,732 | | *ERBB2* | -3.668 | 3.56E-04 | 0.009 |
| | | 37,886,749 | | *MIEN1* | 3.945 | 1.30E-04 | 0.002 |
| | | 37,896,063 | | | 3.476 | 6.92E-04 | 0.009 |
| | | 38,020,485 | *IKZF3* | | 3.962 | 1.22E-04 | 0.005 |
| | | 38,080,829 | | *ORMDL3* | 3.995 | 1.08E-04 | 0.014 |
| | | 38,096,333 | | | -3.664 | 3.60E-04 | 0.033 |
| | | 38,096,571 | | | -4.780 | 4.67E-06 | 0.033 |
| | | 38,096,578 | | | -5.642 | 1.00E-07 | 0.033 |
| | | 38,096,648 | | | -3.887 | 1.61E-04 | 0.033 |
| rs9902621 | g.37973011.A>G | 37,856,732 | | *ERBB2* | -3.668 | 3.56E-04 | 0.009 |
| | | 37,886,749 | | *MIEN1* | 3.945 | 1.30E-04 | 0.002 |
| | | 37,896,063 | | | 3.476 | 6.92E-04 | 0.009 |
| | | 38,020,485 | *IKZF3* | | 3.962 | 1.22E-04 | 0.005 |
| | | 38,080,829 | | *ORMDL3* | 3.995 | 1.08E-04 | 0.014 |
| | | 38,096,333 | | | -3.664 | 3.60E-04 | 0.033 |
| | | 38,096,571 | | | -4.780 | 4.67E-06 | 0.033 |
| | | 38,096,578 | | | -5.642 | 1.00E-07 | 0.033 |
| | | 38,096,648 | | | -3.887 | 1.61E-04 | 0.033 |
| rs9913957 | g.37974499.G>A | 37,856,732 | | *ERBB2* | -3.668 | 3.56E-04 | 0.009 |
| | | 37,886,749 | | *MIEN1* | 3.945 | 1.30E-04 | 0.002 |
| | | 37,896,063 | | | 3.476 | 6.92E-04 | 0.009 |
| | | 38,020,485 | *IKZF3* | | 3.962 | 1.22E-04 | 0.005 |
| | | 38,080,829 | | *ORMDL3* | 3.995 | 1.08E-04 | 0.014 |
| | | 38,096,333 | | | -3.664 | 3.60E-04 | 0.033 |
| | | 38,096,571 | | | -4.780 | 4.67E-06 | 0.033 |
| | | 38,096,578 | | | -5.642 | 1.00E-07 | 0.033 |

| | | | | | | | |
|---|---|---|---|---|---|---|---|
| | | 38,096,648 | | | -3.887 | 1.61E-04 | 0.033 |
| rs58075375 | g.37975593.T>C | 37,856,732 | | *ERBB2* | -3.668 | 3.56E-04 | 0.009 |
| | | 37,886,749 | | *MIEN1* | 3.945 | 1.30E-04 | 0.002 |
| | | 37,896,063 | | | 3.476 | 6.92E-04 | 0.009 |
| | | 38,020,485 | *IKZF3* | | 3.962 | 1.22E-04 | 0.005 |
| | | 38,080,829 | | *ORMDL3* | 3.995 | 1.08E-04 | 0.014 |
| | | 38,096,333 | | | -3.664 | 3.60E-04 | 0.033 |
| | | 38,096,571 | | | -4.780 | 4.67E-06 | 0.033 |
| | | 38,096,578 | | | -5.642 | 1.00E-07 | 0.033 |
| | | 38,096,648 | | | -3.887 | 1.61E-04 | 0.033 |
| rs34053394 | g.37975661.G>A | 37,856,732 | | *ERBB2* | -3.668 | 3.56E-04 | 0.009 |
| | | 37,886,749 | | *MIEN1* | 3.945 | 1.30E-04 | 0.002 |
| | | 37,896,063 | | | 3.476 | 6.92E-04 | 0.009 |
| | | 38,020,485 | *IKZF3* | | 3.962 | 1.22E-04 | 0.005 |
| | | 38,080,829 | | *ORMDL3* | 3.995 | 1.08E-04 | 0.014 |
| | | 38,096,333 | | | -3.664 | 3.60E-04 | 0.033 |
| | | 38,096,571 | | | -4.780 | 4.67E-06 | 0.033 |
| | | 38,096,578 | | | -5.642 | 1.00E-07 | 0.033 |
| | | 38,096,648 | | | -3.887 | 1.61E-04 | 0.033 |
| rs112743130 | g.37975856.C>G | 37,856,732 | | *ERBB2* | -3.668 | 3.56E-04 | 0.009 |
| | | 37,886,749 | | *MIEN1* | 3.945 | 1.30E-04 | 0.002 |
| | | 37,896,063 | | | 3.476 | 6.92E-04 | 0.009 |
| | | 38,020,485 | *IKZF3* | | 3.962 | 1.22E-04 | 0.005 |
| | | 38,080,829 | | *ORMDL3* | 3.995 | 1.08E-04 | 0.014 |
| | | 38,096,333 | | | -3.664 | 3.60E-04 | 0.033 |
| | | 38,096,571 | | | -4.780 | 4.67E-06 | 0.033 |
| | | 38,096,578 | | | -5.642 | 1.00E-07 | 0.033 |
| | | 38,096,648 | | | -3.887 | 1.61E-04 | 0.033 |
| rs9901917 | g.37976206.C>G | 37,856,732 | | *ERBB2* | -3.668 | 3.56E-04 | 0.009 |
| | | 37,886,749 | | *MIEN1* | 3.945 | 1.30E-04 | 0.002 |
| | | 37,896,063 | | | 3.476 | 6.92E-04 | 0.009 |
| | | 38,020,485 | *IKZF3* | | 3.962 | 1.22E-04 | 0.005 |
| | | 38,080,829 | | *ORMDL3* | 3.995 | 1.08E-04 | 0.014 |
| | | 38,096,333 | | | -3.664 | 3.60E-04 | 0.033 |
| | | 38,096,571 | | | -4.780 | 4.67E-06 | 0.033 |
| | | 38,096,578 | | | -5.642 | 1.00E-07 | 0.033 |
| | | 38,096,648 | | | -3.887 | 1.61E-04 | 0.033 |
| rs9904834 | g.37976501.G>A | 37,856,732 | | *ERBB2* | -3.668 | 3.56E-04 | 0.009 |
| | | 37,886,749 | | *MIEN1* | 3.945 | 1.30E-04 | 0.002 |
| | | 37,896,063 | | | 3.476 | 6.92E-04 | 0.009 |
| | | 38,020,485 | *IKZF3* | | 3.962 | 1.22E-04 | 0.005 |

| | | | | | | |
|---|---|---|---|---|---|---|
| | | 38,080,829 | *ORMDL3* | 3.995 | 1.08E-04 | 0.014 |
| | | 38,096,333 | | -3.664 | 3.60E-04 | 0.033 |
| | | 38,096,571 | | -4.780 | 4.67E-06 | 0.033 |
| | | 38,096,578 | | -5.642 | 1.00E-07 | 0.033 |
| | | 38,096,648 | | -3.887 | 1.61E-04 | 0.033 |
| rs9904277 | g.37976599.T>C | 37,856,732 | *ERBB2* | -3.668 | 3.56E-04 | 0.009 |
| | | 37,886,749 | *MIEN1* | 3.945 | 1.30E-04 | 0.002 |
| | | 37,896,063 | | 3.476 | 6.92E-04 | 0.009 |
| | | 38,020,485 | *IKZF3* | 3.962 | 1.22E-04 | 0.005 |
| | | 38,080,829 | *ORMDL3* | 3.995 | 1.08E-04 | 0.014 |
| | | 38,096,333 | | -3.664 | 3.60E-04 | 0.033 |
| | | 38,096,571 | | -4.780 | 4.67E-06 | 0.033 |
| | | 38,096,578 | | -5.642 | 1.00E-07 | 0.033 |
| | | 38,096,648 | | -3.887 | 1.61E-04 | 0.033 |
| rs9911069 | g.37976602.C>T | 37,856,732 | *ERBB2* | -3.668 | 3.56E-04 | 0.009 |
| | | 37,886,749 | *MIEN1* | 3.945 | 1.30E-04 | 0.002 |
| | | 37,896,063 | | 3.476 | 6.92E-04 | 0.009 |
| | | 38,020,485 | *IKZF3* | 3.962 | 1.22E-04 | 0.005 |
| | | 38,080,829 | *ORMDL3* | 3.995 | 1.08E-04 | 0.014 |
| | | 38,096,333 | | -3.664 | 3.60E-04 | 0.033 |
| | | 38,096,571 | | -4.780 | 4.67E-06 | 0.033 |
| | | 38,096,578 | | -5.642 | 1.00E-07 | 0.033 |
| | | 38,096,648 | | -3.887 | 1.61E-04 | 0.033 |
| rs9908983 | g.37976927.A>G | 37,856,732 | *ERBB2* | -3.668 | 3.56E-04 | 0.009 |
| | | 37,886,749 | *MIEN1* | 3.945 | 1.30E-04 | 0.002 |
| | | 37,896,063 | | 3.476 | 6.92E-04 | 0.009 |
| | | 38,020,485 | *IKZF3* | 3.962 | 1.22E-04 | 0.005 |
| | | 38,080,829 | *ORMDL3* | 3.995 | 1.08E-04 | 0.014 |
| | | 38,096,333 | | -3.664 | 3.60E-04 | 0.033 |
| | | 38,096,571 | | -4.780 | 4.67E-06 | 0.033 |
| | | 38,096,578 | | -5.642 | 1.00E-07 | 0.033 |
| | | 38,096,648 | | -3.887 | 1.61E-04 | 0.033 |
| rs8076347 | g.37977541.T>G | 37,856,732 | *ERBB2* | -3.668 | 3.56E-04 | 0.009 |
| | | 37,886,749 | *MIEN1* | 3.945 | 1.30E-04 | 0.002 |
| | | 37,896,063 | | 3.476 | 6.92E-04 | 0.009 |
| | | 38,020,485 | *IKZF3* | 3.962 | 1.22E-04 | 0.005 |
| | | 38,080,829 | *ORMDL3* | 3.995 | 1.08E-04 | 0.014 |
| | | 38,096,333 | | -3.664 | 3.60E-04 | 0.033 |
| | | 38,096,571 | | -4.780 | 4.67E-06 | 0.033 |
| | | 38,096,578 | | -5.642 | 1.00E-07 | 0.033 |
| | | 38,096,648 | | -3.887 | 1.61E-04 | 0.033 |

| | | | | | | | |
|---|---|---|---|---|---|---|---|
| rs8067523 | g.37977814.C>T | 37,856,732 | | *ERBB2* | -3.668 | 3.56E-04 | 0.009 |
| | | 37,886,749 | | *MIEN1* | 3.945 | 1.30E-04 | 0.002 |
| | | 37,896,063 | | | 3.476 | 6.92E-04 | 0.009 |
| | | 38,020,485 | *IKZF3* | | 3.962 | 1.22E-04 | 0.005 |
| | | 38,080,829 | | *ORMDL3* | 3.995 | 1.08E-04 | 0.014 |
| | | 38,096,333 | | | -3.664 | 3.60E-04 | 0.033 |
| | | 38,096,571 | | | -4.780 | 4.67E-06 | 0.033 |
| | | 38,096,578 | | | -5.642 | 1.00E-07 | 0.033 |
| | | 38,096,648 | | | -3.887 | 1.61E-04 | 0.033 |
| rs111907649 | g.37979125.A>G | 37,856,873 | | *ERBB2* | -4.032 | 9.35E-05 | 0.009 |
| | | 37,886,749 | | *MIEN1* | 5.501 | 1.93E-07 | 0.002 |
| | | 37,893,638 | | | -3.510 | 6.16E-04 | 0.002 |
| | | 37,894,352 | | | -4.012 | 1.01E-04 | 0.013 |
| | | 37,896,203 | | | 3.371 | 9.87E-04 | 0.011 |
| | | 38,080,829 | | *ORMDL3* | 4.552 | 1.21E-05 | 0.014 |
| | | 38,096,571 | | | -3.892 | 1.58E-04 | 0.033 |
| | | 38,096,578 | | | -3.664 | 3.60E-04 | 0.033 |
| rs12940616 | g.37979780.G>A | 37,856,732 | | *ERBB2* | -3.668 | 3.56E-04 | 0.009 |
| | | 37,886,749 | | *MIEN1* | 3.945 | 1.30E-04 | 0.002 |
| | | 37,896,063 | | | 3.476 | 6.92E-04 | 0.009 |
| | | 38,020,485 | *IKZF3* | | 3.962 | 1.22E-04 | 0.005 |
| | | 38,080,829 | | *ORMDL3* | 3.995 | 1.08E-04 | 0.014 |
| | | 38,096,333 | | | -3.664 | 3.60E-04 | 0.033 |
| | | 38,096,571 | | | -4.780 | 4.67E-06 | 0.033 |
| | | 38,096,578 | | | -5.642 | 1.00E-07 | 0.033 |
| | | 38,096,648 | | | -3.887 | 1.61E-04 | 0.033 |
| rs72832930 | g.37979950.A>G | 37,886,653 | | *MIEN1* | 3.383 | 9.48E-04 | 0.002 |
| | | 37,886,971 | *MIEN1* | | 3.465 | 7.18E-04 | 0.002 |
| rs62066986 | g.37979986.T>C | 37,845,068 | *PGAP3* | *ERBB2* | 3.467 | 7.13E-04 | 7.82E-05 |
| | | 37,886,914 | *MIEN1* | | 3.877 | 1.67E-04 | 0.002 |
| | | 37,910,225 | | | 3.381 | 9.54E-04 | 0.043 |
| | | 38,019,871 | | *IKZF3* | -3.550 | 5.36E-04 | 0.046 |
| | | 38,085,031 | *ORMDL3* | | 4.637 | 8.49E-06 | 0.005 |
| | | 38,109,763 | | | -3.693 | 3.25E-04 | 0.002 |
| rs28880017 | g.37980023.T>C | 37,856,732 | | *ERBB2* | -3.668 | 3.56E-04 | 0.009 |
| | | 37,886,749 | | *MIEN1* | 3.945 | 1.30E-04 | 0.002 |
| | | 37,896,063 | | | 3.476 | 6.92E-04 | 0.009 |
| | | 38,020,485 | *IKZF3* | | 3.962 | 1.22E-04 | 0.005 |
| | | 38,080,829 | | *ORMDL3* | 3.995 | 1.08E-04 | 0.014 |
| | | 38,096,333 | | | -3.664 | 3.60E-04 | 0.033 |
| | | 38,096,571 | | | -4.780 | 4.67E-06 | 0.033 |

| | | | | | | | |
|---|---|---|---|---|---|---|---|
| | | 38,096,578 | | | -5.642 | 1.00E-07 | 0.033 |
| | | 38,096,648 | | | -3.887 | 1.61E-04 | 0.033 |
| rs9914665 | g.37980567.C>T | 37,856,732 | | *ERBB2* | -3.668 | 3.56E-04 | 0.009 |
| | | 37,886,749 | | *MIEN1* | 3.945 | 1.30E-04 | 0.002 |
| | | 37,896,063 | | | 3.476 | 6.92E-04 | 0.009 |
| | | 38,020,485 | *IKZF3* | | 3.962 | 1.22E-04 | 0.005 |
| | | 38,080,829 | | *ORMDL3* | 3.995 | 1.08E-04 | 0.014 |
| | | 38,096,333 | | | -3.664 | 3.60E-04 | 0.033 |
| | | 38,096,571 | | | -4.780 | 4.67E-06 | 0.033 |
| | | 38,096,578 | | | -5.642 | 1.00E-07 | 0.033 |
| | | 38,096,648 | | | -3.887 | 1.61E-04 | 0.033 |
| rs34925681 | g.37981129.T>C | 37,856,732 | | *ERBB2* | -3.668 | 3.56E-04 | 0.009 |
| | | 37,886,749 | | *MIEN1* | 3.945 | 1.30E-04 | 0.002 |
| | | 37,896,063 | | | 3.476 | 6.92E-04 | 0.009 |
| | | 38,020,485 | *IKZF3* | | 3.962 | 1.22E-04 | 0.005 |
| | | 38,080,829 | | *ORMDL3* | 3.995 | 1.08E-04 | 0.014 |
| | | 38,096,333 | | | -3.664 | 3.60E-04 | 0.033 |
| | | 38,096,571 | | | -4.780 | 4.67E-06 | 0.033 |
| | | 38,096,578 | | | -5.642 | 1.00E-07 | 0.033 |
| | | 38,096,648 | | | -3.887 | 1.61E-04 | 0.033 |
| rs34864615 | g.37981224.A>G | 37,856,732 | | *ERBB2* | -3.668 | 3.56E-04 | 0.009 |
| | | 37,886,749 | | *MIEN1* | 3.945 | 1.30E-04 | 0.002 |
| | | 37,896,063 | | | 3.476 | 6.92E-04 | 0.009 |
| | | 38,020,485 | *IKZF3* | | 3.962 | 1.22E-04 | 0.005 |
| | | 38,080,829 | | *ORMDL3* | 3.995 | 1.08E-04 | 0.014 |
| | | 38,096,333 | | | -3.664 | 3.60E-04 | 0.033 |
| | | 38,096,571 | | | -4.780 | 4.67E-06 | 0.033 |
| | | 38,096,578 | | | -5.642 | 1.00E-07 | 0.033 |
| | | 38,096,648 | | | -3.887 | 1.61E-04 | 0.033 |
| rs34640000 | g.37981238.A>G | 37,856,732 | | *ERBB2* | -3.668 | 3.56E-04 | 0.009 |
| | | 37,886,749 | | *MIEN1* | 3.945 | 1.30E-04 | 0.002 |
| | | 37,896,063 | | | 3.476 | 6.92E-04 | 0.009 |
| | | 38,020,485 | *IKZF3* | | 3.962 | 1.22E-04 | 0.005 |
| | | 38,080,829 | | *ORMDL3* | 3.995 | 1.08E-04 | 0.014 |
| | | 38,096,333 | | | -3.664 | 3.60E-04 | 0.033 |
| | | 38,096,571 | | | -4.780 | 4.67E-06 | 0.033 |
| | | 38,096,578 | | | -5.642 | 1.00E-07 | 0.033 |
| | | 38,096,648 | | | -3.887 | 1.61E-04 | 0.033 |
| rs12942660 | g.37982038.T>C | 37,856,732 | | *ERBB2* | -3.668 | 3.56E-04 | 0.009 |
| | | 37,886,749 | | *MIEN1* | 3.945 | 1.30E-04 | 0.002 |
| | | 37,896,063 | | | 3.476 | 6.92E-04 | 0.009 |

| | | | | | | | |
|---|---|---|---|---|---|---|---|
| | | 38,020,485 | *IKZF3* | | 3.962 | 1.22E-04 | 0.005 |
| | | 38,080,829 | | *ORMDL3* | 3.995 | 1.08E-04 | 0.014 |
| | | 38,096,333 | | | -3.664 | 3.60E-04 | 0.033 |
| | | 38,096,571 | | | -4.780 | 4.67E-06 | 0.033 |
| | | 38,096,578 | | | -5.642 | 1.00E-07 | 0.033 |
| | | 38,096,648 | | | -3.887 | 1.61E-04 | 0.033 |
| rs111469562 | g.37982697.C>T | 37,856,732 | | *ERBB2* | -3.668 | 3.56E-04 | 0.009 |
| | | 37,886,749 | | *MIEN1* | 3.945 | 1.30E-04 | 0.002 |
| | | 37,896,063 | | | 3.476 | 6.92E-04 | 0.009 |
| | | 38,020,485 | *IKZF3* | | 3.962 | 1.22E-04 | 0.005 |
| | | 38,080,829 | | *ORMDL3* | 3.995 | 1.08E-04 | 0.014 |
| | | 38,096,333 | | | -3.664 | 3.60E-04 | 0.033 |
| | | 38,096,571 | | | -4.780 | 4.67E-06 | 0.033 |
| | | 38,096,578 | | | -5.642 | 1.00E-07 | 0.033 |
| | | 38,096,648 | | | -3.887 | 1.61E-04 | 0.033 |
| rs35130019 | g.37983142.G>A | 37,856,732 | | *ERBB2* | -3.668 | 3.56E-04 | 0.009 |
| | | 37,886,749 | | *MIEN1* | 3.945 | 1.30E-04 | 0.002 |
| | | 37,896,063 | | | 3.476 | 6.92E-04 | 0.009 |
| | | 38,020,485 | *IKZF3* | | 3.962 | 1.22E-04 | 0.005 |
| | | 38,080,829 | | *ORMDL3* | 3.995 | 1.08E-04 | 0.014 |
| | | 38,096,333 | | | -3.664 | 3.60E-04 | 0.033 |
| | | 38,096,571 | | | -4.780 | 4.67E-06 | 0.033 |
| | | 38,096,578 | | | -5.642 | 1.00E-07 | 0.033 |
| | | 38,096,648 | | | -3.887 | 1.61E-04 | 0.033 |
| rs112437508 | g.37983513.A>G | 37,856,732 | | *ERBB2* | -3.668 | 3.56E-04 | 0.009 |
| | | 37,886,749 | | *MIEN1* | 3.945 | 1.30E-04 | 0.002 |
| | | 37,896,063 | | | 3.476 | 6.92E-04 | 0.009 |
| | | 38,020,485 | *IKZF3* | | 3.962 | 1.22E-04 | 0.005 |
| | | 38,080,829 | | *ORMDL3* | 3.995 | 1.08E-04 | 0.014 |
| | | 38,096,333 | | | -3.664 | 3.60E-04 | 0.033 |
| | | 38,096,571 | | | -4.780 | 4.67E-06 | 0.033 |
| | | 38,096,578 | | | -5.642 | 1.00E-07 | 0.033 |
| | | 38,096,648 | | | -3.887 | 1.61E-04 | 0.033 |
| rs112797570 | g.37983752.A>G | 37,856,732 | | *ERBB2* | -3.668 | 3.56E-04 | 0.009 |
| | | 37,886,749 | | *MIEN1* | 3.945 | 1.30E-04 | 0.002 |
| | | 37,896,063 | | | 3.476 | 6.92E-04 | 0.009 |
| | | 38,020,485 | *IKZF3* | | 3.962 | 1.22E-04 | 0.005 |
| | | 38,080,829 | | *ORMDL3* | 3.995 | 1.08E-04 | 0.014 |
| | | 38,096,333 | | | -3.664 | 3.60E-04 | 0.033 |
| | | 38,096,571 | | | -4.780 | 4.67E-06 | 0.033 |
| | | 38,096,578 | | | -5.642 | 1.00E-07 | 0.033 |

| | | | | | | | |
|---|---|---|---|---|---|---|---|
| | | 38,096,648 | | | -3.887 | 1.61E-04 | 0.033 |
| rs66763967 | g.37984078.C>T | 37,856,732 | | *ERBB2* | -3.668 | 3.56E-04 | 0.009 |
| | | 37,886,749 | | *MIEN1* | 3.945 | 1.30E-04 | 0.002 |
| | | 37,896,063 | | | 3.476 | 6.92E-04 | 0.009 |
| | | 38,020,485 | *IKZF3* | | 3.962 | 1.22E-04 | 0.005 |
| | | 38,080,829 | | *ORMDL3* | 3.995 | 1.08E-04 | 0.014 |
| | | 38,096,333 | | | -3.664 | 3.60E-04 | 0.033 |
| | | 38,096,571 | | | -4.780 | 4.67E-06 | 0.033 |
| | | 38,096,578 | | | -5.642 | 1.00E-07 | 0.033 |
| | | 38,096,648 | | | -3.887 | 1.61E-04 | 0.033 |
| rs71369791 | g.37984086.G>A | 37,856,732 | | *ERBB2* | -3.668 | 3.56E-04 | 0.009 |
| | | 37,886,749 | | *MIEN1* | 3.945 | 1.30E-04 | 0.002 |
| | | 37,896,063 | | | 3.476 | 6.92E-04 | 0.009 |
| | | 38,020,485 | *IKZF3* | | 3.962 | 1.22E-04 | 0.005 |
| | | 38,080,829 | | *ORMDL3* | 3.995 | 1.08E-04 | 0.014 |
| | | 38,096,333 | | | -3.664 | 3.60E-04 | 0.033 |
| | | 38,096,571 | | | -4.780 | 4.67E-06 | 0.033 |
| | | 38,096,578 | | | -5.642 | 1.00E-07 | 0.033 |
| | | 38,096,648 | | | -3.887 | 1.61E-04 | 0.033 |
| rs12946188 | g.37984430.A>C | 37,856,732 | | *ERBB2* | -3.668 | 3.56E-04 | 0.009 |
| | | 37,886,749 | | *MIEN1* | 3.945 | 1.30E-04 | 0.002 |
| | | 37,896,063 | | | 3.476 | 6.92E-04 | 0.009 |
| | | 38,020,485 | *IKZF3* | | 3.962 | 1.22E-04 | 0.005 |
| | | 38,080,829 | | *ORMDL3* | 3.995 | 1.08E-04 | 0.014 |
| | | 38,096,333 | | | -3.664 | 3.60E-04 | 0.033 |
| | | 38,096,571 | | | -4.780 | 4.67E-06 | 0.033 |
| | | 38,096,578 | | | -5.642 | 1.00E-07 | 0.033 |
| | | 38,096,648 | | | -3.887 | 1.61E-04 | 0.033 |
| rs111247722 | g.37984480.A>C | 37,886,788 | *MIEN1* | *MIEN1* | 3.557 | 5.25E-04 | 0.002 |
| | | 37,910,144 | | | 3.620 | 4.20E-04 | 0.043 |
| | | 38,024,592 | | | 4.175 | 5.43E-05 | 0.046 |
| | | 38,024,606 | | | 3.541 | 5.55E-04 | 0.046 |
| | | 38,083,447 | | *ORMDL3* | 3.530 | 5.75E-04 | 0.011 |
| | | 38,096,333 | | | -3.727 | 2.88E-04 | 0.033 |
| | | 38,096,571 | | | -5.028 | 1.61E-06 | 0.033 |
| | | 38,096,578 | | | -4.429 | 1.99E-05 | 0.033 |
| | | 38,109,575 | | | 3.440 | 7.82E-04 | 0.002 |
| rs12946792 | g.37985322.C>T | 37,856,732 | | *ERBB2* | -3.668 | 3.56E-04 | 0.009 |
| | | 37,886,749 | | *MIEN1* | 3.945 | 1.30E-04 | 0.002 |
| | | 37,896,063 | | | 3.476 | 6.92E-04 | 0.009 |
| | | 38,020,485 | *IKZF3* | | 3.962 | 1.22E-04 | 0.005 |

| | | | | | | | |
|---|---|---|---|---|---|---|---|
| | | 38,080,829 | | *ORMDL3* | 3.995 | 1.08E-04 | 0.014 |
| | | 38,096,333 | | | -3.664 | 3.60E-04 | 0.033 |
| | | 38,096,571 | | | -4.780 | 4.67E-06 | 0.033 |
| | | 38,096,578 | | | -5.642 | 1.00E-07 | 0.033 |
| | | 38,096,648 | | | -3.887 | 1.61E-04 | 0.033 |
| rs138214281 | g.37986106.A>G | 37,832,244 | | *PGAP3* | 3.917 | 1.44E-04 | 0.028 |
| | | 37,886,847 | *MIEN1* | | 4.412 | 2.12E-05 | 0.002 |
| | | 37,886,894 | *MIEN1* | | 6.956 | 1.53E-10 | 0.002 |
| | | 37,895,018 | | | 4.670 | 7.42E-06 | 0.004 |
| | | 37,895,429 | | | -3.385 | 9.40E-04 | 0.006 |
| | | 37,910,190 | | | 3.612 | 4.33E-04 | 0.043 |
| | | 37,910,193 | | | 3.915 | 1.45E-04 | 0.043 |
| | | 37,910,326 | | | 3.383 | 9.48E-04 | 0.013 |
| | | 37,910,588 | | | 3.558 | 5.22E-04 | 0.013 |
| | | 37,910,631 | | | 4.220 | 4.56E-05 | 0.013 |
| | | 37,910,784 | | | 3.560 | 5.19E-04 | 0.035 |
| | | 37,911,103 | | | 4.381 | 2.41E-05 | 0.035 |
| | | 38,020,858 | *IKZF3* | | 3.797 | 2.23E-04 | 0.017 |
| | | 38,081,530 | | *ORMDL3* | 3.815 | 2.09E-04 | 2.01E-04 |
| | | 38,083,134 | | *ORMDL3* | 3.404 | 8.83E-04 | 0.011 |
| | | 38,083,548 | | *ORMDL3* | 3.632 | 4.04E-04 | 0.011 |
| | | 38,109,084 | | | 4.389 | 2.33E-05 | 0.002 |
| | | 38,109,949 | | | 3.768 | 2.48E-04 | 0.038 |
| | | 38,133,359 | | *GSDMA* | 6.407 | 2.49E-09 | 0.038 |
| rs9913769 | g.37986337.G>A | 37,856,732 | | *ERBB2* | -3.668 | 3.56E-04 | 0.009 |
| | | 37,886,749 | | *MIEN1* | 3.945 | 1.30E-04 | 0.002 |
| | | 37,896,063 | | | 3.476 | 6.92E-04 | 0.009 |
| | | 38,020,485 | *IKZF3* | | 3.962 | 1.22E-04 | 0.005 |
| | | 38,080,829 | | *ORMDL3* | 3.995 | 1.08E-04 | 0.014 |
| | | 38,096,333 | | | -3.664 | 3.60E-04 | 0.033 |
| | | 38,096,571 | | | -4.780 | 4.67E-06 | 0.033 |
| | | 38,096,578 | | | -5.642 | 1.00E-07 | 0.033 |
| | | 38,096,648 | | | -3.887 | 1.61E-04 | 0.033 |
| rs534965497 | g.37986744.C>A | 37,894,027 | | | 5.163 | 8.88E-07 | 0.013 |
| | | 37,895,775 | | | -3.735 | 2.80E-04 | 0.027 |
| | | 38,024,290 | | | 3.731 | 2.84E-04 | 0.019 |
| rs71369792 | g.37986873.A>G | 37,856,732 | | *ERBB2* | -3.668 | 3.56E-04 | 0.009 |
| | | 37,886,749 | | *MIEN1* | 3.945 | 1.30E-04 | 0.002 |
| | | 37,896,063 | | | 3.476 | 6.92E-04 | 0.009 |
| | | 38,020,485 | *IKZF3* | | 3.962 | 1.22E-04 | 0.005 |
| | | 38,080,829 | | *ORMDL3* | 3.995 | 1.08E-04 | 0.014 |

| | | | | | | | |
|---|---|---|---|---|---|---|---|
| | | 38,096,333 | | | -3.664 | 3.60E-04 | 0.033 |
| | | 38,096,571 | | | -4.780 | 4.67E-06 | 0.033 |
| | | 38,096,578 | | | -5.642 | 1.00E-07 | 0.033 |
| | | 38,096,648 | | | -3.887 | 1.61E-04 | 0.033 |
| rs113479772 | g.37987043.A>G | 37,856,732 | | *ERBB2* | -3.668 | 3.56E-04 | 0.009 |
| | | 37,886,749 | | *MIEN1* | 3.945 | 1.30E-04 | 0.002 |
| | | 37,896,063 | | | 3.476 | 6.92E-04 | 0.009 |
| | | 38,020,485 | *IKZF3* | | 3.962 | 1.22E-04 | 0.005 |
| | | 38,080,829 | | *ORMDL3* | 3.995 | 1.08E-04 | 0.014 |
| | | 38,096,333 | | | -3.664 | 3.60E-04 | 0.033 |
| | | 38,096,571 | | | -4.780 | 4.67E-06 | 0.033 |
| | | 38,096,578 | | | -5.642 | 1.00E-07 | 0.033 |
| | | 38,096,648 | | | -3.887 | 1.61E-04 | 0.033 |
| rs111734595 | g.37987400.T>C | 37,856,732 | | *ERBB2* | -3.668 | 3.56E-04 | 0.009 |
| | | 37,886,749 | | *MIEN1* | 3.945 | 1.30E-04 | 0.002 |
| | | 37,896,063 | | | 3.476 | 6.92E-04 | 0.009 |
| | | 38,020,485 | *IKZF3* | | 3.962 | 1.22E-04 | 0.005 |
| | | 38,080,829 | | *ORMDL3* | 3.995 | 1.08E-04 | 0.014 |
| | | 38,096,333 | | | -3.664 | 3.60E-04 | 0.033 |
| | | 38,096,571 | | | -4.780 | 4.67E-06 | 0.033 |
| | | 38,096,578 | | | -5.642 | 1.00E-07 | 0.033 |
| | | 38,096,648 | | | -3.887 | 1.61E-04 | 0.033 |
| rs112141468 | g.37987465.T>C | 37,856,732 | | *ERBB2* | -3.668 | 3.56E-04 | 0.009 |
| | | 37,886,749 | | *MIEN1* | 3.945 | 1.30E-04 | 0.002 |
| | | 37,896,063 | | | 3.476 | 6.92E-04 | 0.009 |
| | | 38,020,485 | *IKZF3* | | 3.962 | 1.22E-04 | 0.005 |
| | | 38,080,829 | | *ORMDL3* | 3.995 | 1.08E-04 | 0.014 |
| | | 38,096,333 | | | -3.664 | 3.60E-04 | 0.033 |
| | | 38,096,571 | | | -4.780 | 4.67E-06 | 0.033 |
| | | 38,096,578 | | | -5.642 | 1.00E-07 | 0.033 |
| | | 38,096,648 | | | -3.887 | 1.61E-04 | 0.033 |
| rs73304123 | g.37987589.T>C | 37,856,732 | | *ERBB2* | -3.668 | 3.56E-04 | 0.009 |
| | | 37,886,749 | | *MIEN1* | 3.945 | 1.30E-04 | 0.002 |
| | | 37,896,063 | | | 3.476 | 6.92E-04 | 0.009 |
| | | 38,020,485 | *IKZF3* | | 3.962 | 1.22E-04 | 0.005 |
| | | 38,080,829 | | *ORMDL3* | 3.995 | 1.08E-04 | 0.014 |
| | | 38,096,333 | | | -3.664 | 3.60E-04 | 0.033 |
| | | 38,096,571 | | | -4.780 | 4.67E-06 | 0.033 |
| | | 38,096,578 | | | -5.642 | 1.00E-07 | 0.033 |
| | | 38,096,648 | | | -3.887 | 1.61E-04 | 0.033 |
| rs8069893 | g.37988293.T>C | 37,856,732 | | *ERBB2* | -3.668 | 3.56E-04 | 0.009 |

| | | | | | | | |
|---|---|---|---|---|---|---|---|
| | | 37,886,749 | | *MIEN1* | 3.945 | 1.30E-04 | 0.002 |
| | | 37,896,063 | | | 3.476 | 6.92E-04 | 0.009 |
| | | 38,020,485 | *IKZF3* | | 3.962 | 1.22E-04 | 0.005 |
| | | 38,080,829 | | *ORMDL3* | 3.995 | 1.08E-04 | 0.014 |
| | | 38,096,333 | | | -3.664 | 3.60E-04 | 0.033 |
| | | 38,096,571 | | | -4.780 | 4.67E-06 | 0.033 |
| | | 38,096,578 | | | -5.642 | 1.00E-07 | 0.033 |
| | | 38,096,648 | | | -3.887 | 1.61E-04 | 0.033 |
| rs111944912 | g.37988477.C>T | 37,856,732 | | *ERBB2* | -3.668 | 3.56E-04 | 0.009 |
| | | 37,886,749 | | *MIEN1* | 3.945 | 1.30E-04 | 0.002 |
| | | 37,896,063 | | | 3.476 | 6.92E-04 | 0.009 |
| | | 38,020,485 | *IKZF3* | | 3.962 | 1.22E-04 | 0.005 |
| | | 38,080,829 | | *ORMDL3* | 3.995 | 1.08E-04 | 0.014 |
| | | 38,096,333 | | | -3.664 | 3.60E-04 | 0.033 |
| | | 38,096,571 | | | -4.780 | 4.67E-06 | 0.033 |
| | | 38,096,578 | | | -5.642 | 1.00E-07 | 0.033 |
| | | 38,096,648 | | | -3.887 | 1.61E-04 | 0.033 |
| rs34521860 | g.37988775_37988775insAT | 37,856,732 | | *ERBB2* | -3.659 | 3.67E-04 | 0.009 |
| | | 37,886,749 | | *MIEN1* | 3.939 | 1.33E-04 | 0.002 |
| | | 37,896,063 | | | 3.534 | 5.68E-04 | 0.009 |
| | | 38,020,485 | *IKZF3* | | 3.952 | 1.27E-04 | 0.005 |
| | | 38,080,829 | | *ORMDL3* | 4.004 | 1.04E-04 | 0.014 |
| | | 38,096,333 | | | -3.674 | 3.48E-04 | 0.033 |
| | | 38,096,571 | | | -4.755 | 5.19E-06 | 0.033 |
| | | 38,096,578 | | | -5.590 | 1.28E-07 | 0.033 |
| | | 38,096,648 | | | -3.903 | 1.52E-04 | 0.033 |
| rs12945253 | g.37991328.G>C | 37,856,732 | | *ERBB2* | -3.668 | 3.56E-04 | 0.009 |
| | | 37,886,749 | | *MIEN1* | 3.945 | 1.30E-04 | 0.002 |
| | | 37,896,063 | | | 3.476 | 6.92E-04 | 0.009 |
| | | 38,020,485 | *IKZF3* | | 3.962 | 1.22E-04 | 0.005 |
| | | 38,080,829 | | *ORMDL3* | 3.995 | 1.08E-04 | 0.014 |
| | | 38,096,333 | | | -3.664 | 3.60E-04 | 0.033 |
| | | 38,096,571 | | | -4.780 | 4.67E-06 | 0.033 |
| | | 38,096,578 | | | -5.642 | 1.00E-07 | 0.033 |
| | | 38,096,648 | | | -3.887 | 1.61E-04 | 0.033 |
| rs28449671 | g.37991631.C>T | 37,856,732 | | *ERBB2* | -3.668 | 3.56E-04 | 0.009 |
| | | 37,886,749 | | *MIEN1* | 3.945 | 1.30E-04 | 0.002 |
| | | 37,896,063 | | | 3.476 | 6.92E-04 | 0.009 |
| | | 38,020,485 | *IKZF3* | | 3.962 | 1.22E-04 | 0.005 |
| | | 38,080,829 | | *ORMDL3* | 3.995 | 1.08E-04 | 0.014 |

| | | | | | | | |
|---|---|---|---|---|---|---|---|
| | | 38,096,333 | | | -3.664 | 3.60E-04 | 0.033 |
| | | 38,096,571 | | | -4.780 | 4.67E-06 | 0.033 |
| | | 38,096,578 | | | -5.642 | 1.00E-07 | 0.033 |
| | | 38,096,648 | | | -3.887 | 1.61E-04 | 0.033 |
| rs34518282 | g.37992001.C>G | 37,856,732 | | *ERBB2* | -3.668 | 3.56E-04 | 0.009 |
| | | 37,886,749 | | *MIEN1* | 3.945 | 1.30E-04 | 0.002 |
| | | 37,896,063 | | | 3.476 | 6.92E-04 | 0.009 |
| | | 38,020,485 | *IKZF3* | | 3.962 | 1.22E-04 | 0.005 |
| | | 38,080,829 | | *ORMDL3* | 3.995 | 1.08E-04 | 0.014 |
| | | 38,096,333 | | | -3.664 | 3.60E-04 | 0.033 |
| | | 38,096,571 | | | -4.780 | 4.67E-06 | 0.033 |
| | | 38,096,578 | | | -5.642 | 1.00E-07 | 0.033 |
| | | 38,096,648 | | | -3.887 | 1.61E-04 | 0.033 |
| rs28392589 | g.37992885.G>A | 37,856,732 | | *ERBB2* | -3.668 | 3.56E-04 | 0.009 |
| | | 37,886,749 | | *MIEN1* | 3.945 | 1.30E-04 | 0.002 |
| | | 37,896,063 | | | 3.476 | 6.92E-04 | 0.009 |
| | | 38,020,485 | *IKZF3* | | 3.962 | 1.22E-04 | 0.005 |
| | | 38,080,829 | | *ORMDL3* | 3.995 | 1.08E-04 | 0.014 |
| | | 38,096,333 | | | -3.664 | 3.60E-04 | 0.033 |
| | | 38,096,571 | | | -4.780 | 4.67E-06 | 0.033 |
| | | 38,096,578 | | | -5.642 | 1.00E-07 | 0.033 |
| | | 38,096,648 | | | -3.887 | 1.61E-04 | 0.033 |
| rs111691913 | g.37993239.T>C | 37,856,732 | | *ERBB2* | -3.668 | 3.56E-04 | 0.009 |
| | | 37,886,749 | | *MIEN1* | 3.945 | 1.30E-04 | 0.002 |
| | | 37,896,063 | | | 3.476 | 6.92E-04 | 0.009 |
| | | 38,020,485 | *IKZF3* | | 3.962 | 1.22E-04 | 0.005 |
| | | 38,080,829 | | *ORMDL3* | 3.995 | 1.08E-04 | 0.014 |
| | | 38,096,333 | | | -3.664 | 3.60E-04 | 0.033 |
| | | 38,096,571 | | | -4.780 | 4.67E-06 | 0.033 |
| | | 38,096,578 | | | -5.642 | 1.00E-07 | 0.033 |
| | | 38,096,648 | | | -3.887 | 1.61E-04 | 0.033 |
| rs12938617 | g.37993352.A>T | 37,856,732 | | *ERBB2* | -3.668 | 3.56E-04 | 0.009 |
| | | 37,886,749 | | *MIEN1* | 3.945 | 1.30E-04 | 0.002 |
| | | 37,896,063 | | | 3.476 | 6.92E-04 | 0.009 |
| | | 38,020,485 | *IKZF3* | | 3.962 | 1.22E-04 | 0.005 |
| | | 38,080,829 | | *ORMDL3* | 3.995 | 1.08E-04 | 0.014 |
| | | 38,096,333 | | | -3.664 | 3.60E-04 | 0.033 |
| | | 38,096,571 | | | -4.780 | 4.67E-06 | 0.033 |
| | | 38,096,578 | | | -5.642 | 1.00E-07 | 0.033 |
| | | 38,096,648 | | | -3.887 | 1.61E-04 | 0.033 |
| rs9898464 | g.37993992.C>T | 37,856,732 | | *ERBB2* | -3.668 | 3.56E-04 | 0.009 |

| | | | | | | | |
|---|---|---|---|---|---|---|---|
| | | 37,886,749 | | *MIEN1* | 3.945 | 1.30E-04 | 0.002 |
| | | 37,896,063 | | | 3.476 | 6.92E-04 | 0.009 |
| | | 38,020,485 | *IKZF3* | | 3.962 | 1.22E-04 | 0.005 |
| | | 38,080,829 | | *ORMDL3* | 3.995 | 1.08E-04 | 0.014 |
| | | 38,096,333 | | | -3.664 | 3.60E-04 | 0.033 |
| | | 38,096,571 | | | -4.780 | 4.67E-06 | 0.033 |
| | | 38,096,578 | | | -5.642 | 1.00E-07 | 0.033 |
| | | 38,096,648 | | | -3.887 | 1.61E-04 | 0.033 |
| rs9900019 | g.37994625.C>T | 37,856,732 | | *ERBB2* | -3.668 | 3.56E-04 | 0.009 |
| | | 37,886,749 | | *MIEN1* | 3.945 | 1.30E-04 | 0.002 |
| | | 37,896,063 | | | 3.476 | 6.92E-04 | 0.009 |
| | | 38,020,485 | *IKZF3* | | 3.962 | 1.22E-04 | 0.005 |
| | | 38,080,829 | | *ORMDL3* | 3.995 | 1.08E-04 | 0.014 |
| | | 38,096,333 | | | -3.664 | 3.60E-04 | 0.033 |
| | | 38,096,571 | | | -4.780 | 4.67E-06 | 0.033 |
| | | 38,096,578 | | | -5.642 | 1.00E-07 | 0.033 |
| | | 38,096,648 | | | -3.887 | 1.61E-04 | 0.033 |
| rs34485135 | g.37994722.C>T | 37,856,732 | | *ERBB2* | -3.668 | 3.56E-04 | 0.009 |
| | | 37,856,732 | | *ERBB2* | -3.668 | 3.56E-04 | 0.009 |
| | | 37,886,749 | | *MIEN1* | 3.945 | 1.30E-04 | 0.002 |
| | | 37,886,749 | | *MIEN1* | 3.945 | 1.30E-04 | 0.002 |
| | | 37,896,063 | | | 3.476 | 6.92E-04 | 0.009 |
| | | 37,896,063 | | | 3.476 | 6.92E-04 | 0.009 |
| | | 38,020,485 | *IKZF3* | | 3.962 | 1.22E-04 | 0.005 |
| | | 38,020,485 | *IKZF3* | | 3.962 | 1.22E-04 | 0.005 |
| | | 38,080,829 | | *ORMDL3* | 3.995 | 1.08E-04 | 0.014 |
| | | 38,080,829 | | *ORMDL3* | 3.995 | 1.08E-04 | 0.014 |
| | | 38,096,333 | | | -3.664 | 3.60E-04 | 0.033 |
| | | 38,096,333 | | | -3.664 | 3.60E-04 | 0.033 |
| | | 38,096,571 | | | -4.780 | 4.67E-06 | 0.033 |
| | | 38,096,571 | | | -4.780 | 4.67E-06 | 0.033 |
| | | 38,096,578 | | | -5.642 | 1.00E-07 | 0.033 |
| | | 38,096,578 | | | -5.642 | 1.00E-07 | 0.033 |
| | | 38,096,648 | | | -3.887 | 1.61E-04 | 0.033 |
| | | 38,096,648 | | | -3.887 | 1.61E-04 | 0.033 |
| rs183751548 | g.37995365.C>T | 37,828,283 | | *PGAP3* | -5.371 | 3.50E-07 | 0.010 |
| | | 37,828,296 | | *PGAP3* | -3.944 | 1.30E-04 | 0.002 |
| | | 37,835,902 | | *PGAP3* | 3.381 | 9.55E-04 | 0.020 |
| | | 37,856,596 | | *ERBB2* | 4.242 | 4.17E-05 | 0.006 |
| | | 37,896,708 | | | 3.811 | 2.13E-04 | 0.008 |
| | | 37,896,750 | | | 4.751 | 5.27E-06 | 0.008 |

| | | | | | | | |
|---|---|---|---|---|---|---|---|
| | | 37,910,179 | | | 3.839 | 1.92E-04 | 0.043 |
| | | 37,929,495 | | *IKZF3* | 3.889 | 1.60E-04 | 0.001 |
| | | 38,077,176 | | | -3.402 | 8.89E-04 | 0.034 |
| | | 38,108,868 | | | -3.825 | 2.02E-04 | 0.014 |
| rs113930796 | g.37995897.C>T | 37,886,788 | *MIEN1* | *MIEN1* | 4.145 | 6.09E-05 | 0.002 |
| | | 38,024,592 | | | 4.411 | 2.14E-05 | 0.046 |
| | | 38,024,606 | | | 4.294 | 3.40E-05 | 0.046 |
| | | 38,083,737 | | *ORMDL3* | 3.503 | 6.32E-04 | 0.011 |
| | | 38,096,333 | | | -3.549 | 5.39E-04 | 0.033 |
| | | 38,096,571 | | | -4.936 | 2.40E-06 | 0.033 |
| | | 38,096,578 | | | -4.534 | 1.30E-05 | 0.033 |
| rs9900541 | g.37996071.C>T | 37,856,732 | | *ERBB2* | -3.668 | 3.56E-04 | 0.009 |
| | | 37,886,749 | | *MIEN1* | 3.945 | 1.30E-04 | 0.002 |
| | | 37,896,063 | | | 3.476 | 6.92E-04 | 0.009 |
| | | 38,020,485 | *IKZF3* | | 3.962 | 1.22E-04 | 0.005 |
| | | 38,080,829 | | *ORMDL3* | 3.995 | 1.08E-04 | 0.014 |
| | | 38,096,333 | | | -3.664 | 3.60E-04 | 0.033 |
| | | 38,096,571 | | | -4.780 | 4.67E-06 | 0.033 |
| | | 38,096,578 | | | -5.642 | 1.00E-07 | 0.033 |
| | | 38,096,648 | | | -3.887 | 1.61E-04 | 0.033 |
| rs9908694 | g.37997772.T>C | 37,856,732 | | *ERBB2* | -3.668 | 3.56E-04 | 0.009 |
| | | 37,886,749 | | *MIEN1* | 3.945 | 1.30E-04 | 0.002 |
| | | 37,896,063 | | | 3.476 | 6.92E-04 | 0.009 |
| | | 38,020,485 | *IKZF3* | | 3.962 | 1.22E-04 | 0.005 |
| | | 38,080,829 | | *ORMDL3* | 3.995 | 1.08E-04 | 0.014 |
| | | 38,096,333 | | | -3.664 | 3.60E-04 | 0.033 |
| | | 38,096,571 | | | -4.780 | 4.67E-06 | 0.033 |
| | | 38,096,578 | | | -5.642 | 1.00E-07 | 0.033 |
| | | 38,096,648 | | | -3.887 | 1.61E-04 | 0.033 |
| rs35405174 | g.37998705.G>A | 37,856,732 | | *ERBB2* | -3.668 | 3.56E-04 | 0.009 |
| | | 37,886,749 | | *MIEN1* | 3.945 | 1.30E-04 | 0.002 |
| | | 37,896,063 | | | 3.476 | 6.92E-04 | 0.009 |
| | | 38,020,485 | *IKZF3* | | 3.962 | 1.22E-04 | 0.005 |
| | | 38,080,829 | | *ORMDL3* | 3.995 | 1.08E-04 | 0.014 |
| | | 38,096,333 | | | -3.664 | 3.60E-04 | 0.033 |
| | | 38,096,571 | | | -4.780 | 4.67E-06 | 0.033 |
| | | 38,096,578 | | | -5.642 | 1.00E-07 | 0.033 |
| | | 38,096,648 | | | -3.887 | 1.61E-04 | 0.033 |
| rs28403951 | g.37999534.G>C | 37,856,732 | | *ERBB2* | -3.668 | 3.56E-04 | 0.009 |
| | | 37,886,749 | | *MIEN1* | 3.945 | 1.30E-04 | 0.002 |
| | | 37,896,063 | | | 3.476 | 6.92E-04 | 0.009 |

| | | | | | | | |
|---|---|---|---|---|---|---|---|
| | | 38,020,485 | *IKZF3* | | 3.962 | 1.22E-04 | 0.005 |
| | | 38,080,829 | | *ORMDL3* | 3.995 | 1.08E-04 | 0.014 |
| | | 38,096,333 | | | -3.664 | 3.60E-04 | 0.033 |
| | | 38,096,571 | | | -4.780 | 4.67E-06 | 0.033 |
| | | 38,096,578 | | | -5.642 | 1.00E-07 | 0.033 |
| | | 38,096,648 | | | -3.887 | 1.61E-04 | 0.033 |
| rs34135674 | g.37999691_37999702del | 37,856,732 | | *ERBB2* | -4.052 | 8.70E-05 | 0.009 |
| | | 37,886,749 | | *MIEN1* | 5.111 | 1.12E-06 | 0.002 |
| | | 37,894,413 | | | -4.249 | 4.07E-05 | 0.013 |
| | | 37,896,063 | | | 3.439 | 7.85E-04 | 0.009 |
| | | 37,903,456 | | | -3.594 | 4.60E-04 | 0.008 |
| | | 38,020,485 | *IKZF3* | | 3.401 | 8.92E-04 | 0.005 |
| | | 38,024,589 | | | 3.715 | 3.00E-04 | 0.046 |
| | | 38,080,829 | | *ORMDL3* | 3.806 | 2.17E-04 | 0.014 |
| | | 38,096,571 | | | -4.430 | 1.98E-05 | 0.033 |
| | | 38,096,578 | | | -4.627 | 8.87E-06 | 0.033 |
| | | 38,118,571 | *GSDMA* | | 3.463 | 7.25E-04 | 0.009 |
| rs12946040 | g.38000018.A>C | 37,856,732 | | *ERBB2* | -3.668 | 3.56E-04 | 0.009 |
| | | 37,886,749 | | *MIEN1* | 3.945 | 1.30E-04 | 0.002 |
| | | 37,896,063 | | | 3.476 | 6.92E-04 | 0.009 |
| | | 38,020,485 | *IKZF3* | | 3.962 | 1.22E-04 | 0.005 |
| | | 38,080,829 | | *ORMDL3* | 3.995 | 1.08E-04 | 0.014 |
| | | 38,096,333 | | | -3.664 | 3.60E-04 | 0.033 |
| | | 38,096,571 | | | -4.780 | 4.67E-06 | 0.033 |
| | | 38,096,578 | | | -5.642 | 1.00E-07 | 0.033 |
| | | 38,096,648 | | | -3.887 | 1.61E-04 | 0.033 |
| rs36041042 | g.38000057_38000057insT | 37,856,732 | | *ERBB2* | -3.673 | 3.49E-04 | 0.009 |
| | | 37,886,749 | | *MIEN1* | 3.935 | 1.35E-04 | 0.002 |
| | | 37,895,335 | | | 3.386 | 9.40E-04 | 0.006 |
| | | 37,896,006 | | | 3.369 | 9.92E-04 | 0.027 |
| | | 37,896,063 | | | 3.428 | 8.14E-04 | 0.009 |
| | | 38,020,485 | *IKZF3* | | 3.941 | 1.32E-04 | 0.005 |
| | | 38,080,829 | | *ORMDL3* | 4.031 | 9.41E-05 | 0.014 |
| | | 38,096,333 | | | -3.665 | 3.59E-04 | 0.033 |
| | | 38,096,571 | | | -4.755 | 5.18E-06 | 0.033 |
| | | 38,096,578 | | | -5.642 | 1.00E-07 | 0.033 |
| | | 38,096,648 | | | -3.844 | 1.88E-04 | 0.033 |
| rs67600807 | g.38001559.G>A | 37,856,732 | | *ERBB2* | -3.668 | 3.56E-04 | 0.009 |
| | | 37,886,749 | | *MIEN1* | 3.945 | 1.30E-04 | 0.002 |

| | | | | | | | |
|---|---|---|---|---|---|---|---|
| | | 37,896,063 | | | 3.476 | 6.92E-04 | 0.009 |
| | | 38,020,485 | *IKZF3* | | 3.962 | 1.22E-04 | 0.005 |
| | | 38,080,829 | | *ORMDL3* | 3.995 | 1.08E-04 | 0.014 |
| | | 38,096,333 | | | -3.664 | 3.60E-04 | 0.033 |
| | | 38,096,571 | | | -4.780 | 4.67E-06 | 0.033 |
| | | 38,096,578 | | | -5.642 | 1.00E-07 | 0.033 |
| | | 38,096,648 | | | -3.887 | 1.61E-04 | 0.033 |
| rs34039260 | g.38001659_38001659ins C | 37,856,732 | | *ERBB2* | -3.668 | 3.56E-04 | 0.009 |
| | | 37,886,749 | | *MIEN1* | 3.945 | 1.30E-04 | 0.002 |
| | | 37,896,063 | | | 3.476 | 6.92E-04 | 0.009 |
| | | 38,020,485 | *IKZF3* | | 3.962 | 1.22E-04 | 0.005 |
| | | 38,080,829 | | *ORMDL3* | 3.995 | 1.08E-04 | 0.014 |
| | | 38,096,333 | | | -3.664 | 3.60E-04 | 0.033 |
| | | 38,096,571 | | | -4.780 | 4.67E-06 | 0.033 |
| | | 38,096,578 | | | -5.642 | 1.00E-07 | 0.033 |
| | | 38,096,648 | | | -3.887 | 1.61E-04 | 0.033 |
| rs112677036 | g.38002153.A>G | 37,856,732 | | *ERBB2* | -3.668 | 3.56E-04 | 0.009 |
| | | 37,886,749 | | *MIEN1* | 3.945 | 1.30E-04 | 0.002 |
| | | 37,896,063 | | | 3.476 | 6.92E-04 | 0.009 |
| | | 38,020,485 | *IKZF3* | | 3.962 | 1.22E-04 | 0.005 |
| | | 38,080,829 | | *ORMDL3* | 3.995 | 1.08E-04 | 0.014 |
| | | 38,096,333 | | | -3.664 | 3.60E-04 | 0.033 |
| | | 38,096,571 | | | -4.780 | 4.67E-06 | 0.033 |
| | | 38,096,578 | | | -5.642 | 1.00E-07 | 0.033 |
| | | 38,096,648 | | | -3.887 | 1.61E-04 | 0.033 |
| rs117576441 | g.38002455.G>A | 37,844,261 | *ERBB2* | *PGAP3* | 3.593 | 4.62E-04 | 7.82E-05 |
| | | 37,863,585 | | *ERBB2* | 3.448 | 7.62E-04 | 0.002 |
| | | 37,894,397 | | | 3.615 | 4.29E-04 | 0.013 |
| | | 38,024,146 | | | -4.008 | 1.03E-04 | 0.004 |
| | | 38,024,353 | | | -3.936 | 1.34E-04 | 0.017 |
| | | 38,024,483 | | | -3.505 | 6.26E-04 | 0.002 |
| rs573641447 | g.38002747.G>T | 37,856,732 | | *ERBB2* | -4.062 | 8.36E-05 | 0.009 |
| | | 37,886,749 | | *MIEN1* | 4.283 | 3.55E-05 | 0.002 |
| | | 37,895,335 | | | 3.383 | 9.47E-04 | 0.006 |
| | | 37,896,063 | | | 3.534 | 5.67E-04 | 0.009 |
| | | 37,896,203 | | | 3.432 | 8.04E-04 | 0.011 |
| | | 37,976,456 | | *IKZF3* | 3.676 | 3.45E-04 | 0.039 |
| | | 38,076,680 | | | 3.416 | 8.49E-04 | 0.034 |
| | | 38,080,829 | | *ORMDL3* | 3.936 | 1.34E-04 | 0.014 |
| | | 38,096,333 | | | -3.509 | 6.19E-04 | 0.033 |

| | | | | | | | |
|---|---|---|---|---|---|---|---|
| | | 38,096,571 | | | -4.908 | 2.70E-06 | 0.033 |
| | | 38,096,578 | | | -5.754 | 5.94E-08 | 0.033 |
| | | 38,096,648 | | | -4.445 | 1.87E-05 | 0.033 |
| | | 38,118,571 | *GSDMA* | | 3.693 | 3.25E-04 | 0.009 |
| rs28535471 | g.38002916.C>G | 37,856,732 | | *ERBB2* | -3.668 | 3.56E-04 | 0.009 |
| | | 37,886,749 | | *MIEN1* | 3.945 | 1.30E-04 | 0.002 |
| | | 37,896,063 | | | 3.476 | 6.92E-04 | 0.009 |
| | | 38,020,485 | *IKZF3* | | 3.962 | 1.22E-04 | 0.005 |
| | | 38,080,829 | | *ORMDL3* | 3.995 | 1.08E-04 | 0.014 |
| | | 38,096,333 | | | -3.664 | 3.60E-04 | 0.033 |
| | | 38,096,571 | | | -4.780 | 4.67E-06 | 0.033 |
| | | 38,096,578 | | | -5.642 | 1.00E-07 | 0.033 |
| | | 38,096,648 | | | -3.887 | 1.61E-04 | 0.033 |
| rs28648398 | g.38003025.C>G | 37,856,732 | | *ERBB2* | -3.668 | 3.56E-04 | 0.009 |
| | | 37,886,749 | | *MIEN1* | 3.945 | 1.30E-04 | 0.002 |
| | | 37,896,063 | | | 3.476 | 6.92E-04 | 0.009 |
| | | 38,020,485 | *IKZF3* | | 3.962 | 1.22E-04 | 0.005 |
| | | 38,080,829 | | *ORMDL3* | 3.995 | 1.08E-04 | 0.014 |
| | | 38,096,333 | | | -3.664 | 3.60E-04 | 0.033 |
| | | 38,096,571 | | | -4.780 | 4.67E-06 | 0.033 |
| | | 38,096,578 | | | -5.642 | 1.00E-07 | 0.033 |
| | | 38,096,648 | | | -3.887 | 1.61E-04 | 0.033 |
| rs28695941 | g.38003330.T>A | 37,856,732 | | *ERBB2* | -3.668 | 3.56E-04 | 0.009 |
| | | 37,886,749 | | *MIEN1* | 3.945 | 1.30E-04 | 0.002 |
| | | 37,896,063 | | | 3.476 | 6.92E-04 | 0.009 |
| | | 38,020,485 | *IKZF3* | | 3.962 | 1.22E-04 | 0.005 |
| | | 38,080,829 | | *ORMDL3* | 3.995 | 1.08E-04 | 0.014 |
| | | 38,096,333 | | | -3.664 | 3.60E-04 | 0.033 |
| | | 38,096,571 | | | -4.780 | 4.67E-06 | 0.033 |
| | | 38,096,578 | | | -5.642 | 1.00E-07 | 0.033 |
| | | 38,096,648 | | | -3.887 | 1.61E-04 | 0.033 |
| rs34654193 | g.38003859.C>G | 37,856,732 | | *ERBB2* | -3.668 | 3.56E-04 | 0.009 |
| | | 37,886,749 | | *MIEN1* | 3.945 | 1.30E-04 | 0.002 |
| | | 37,896,063 | | | 3.476 | 6.92E-04 | 0.009 |
| | | 38,020,485 | *IKZF3* | | 3.962 | 1.22E-04 | 0.005 |
| | | 38,080,829 | | *ORMDL3* | 3.995 | 1.08E-04 | 0.014 |
| | | 38,096,333 | | | -3.664 | 3.60E-04 | 0.033 |
| | | 38,096,571 | | | -4.780 | 4.67E-06 | 0.033 |
| | | 38,096,578 | | | -5.642 | 1.00E-07 | 0.033 |
| | | 38,096,648 | | | -3.887 | 1.61E-04 | 0.033 |

| | | | | | | | |
|---|---|---|---|---|---|---|---|
| rs555695098 | g.38004454_38004454ins | 37,856,732 | | *ERBB2* | -3.668 | 3.56E-04 | 0.009 |
| | | 37,886,749 | | *MIEN1* | 3.945 | 1.30E-04 | 0.002 |
| | | 37,896,063 | | | 3.476 | 6.92E-04 | 0.009 |
| | | 38,020,485 | *IKZF3* | | 3.962 | 1.22E-04 | 0.005 |
| | | 38,080,829 | | *ORMDL3* | 3.995 | 1.08E-04 | 0.014 |
| | | 38,096,333 | | | -3.664 | 3.60E-04 | 0.033 |
| | | 38,096,571 | | | -4.780 | 4.67E-06 | 0.033 |
| | | 38,096,578 | | | -5.642 | 1.00E-07 | 0.033 |
| | | 38,096,648 | | | -3.887 | 1.61E-04 | 0.033 |
| rs28537796 | g.38004906.T>C | 37,856,732 | | *ERBB2* | -3.668 | 3.56E-04 | 0.009 |
| | | 37,886,749 | | *MIEN1* | 3.945 | 1.30E-04 | 0.002 |
| | | 37,896,063 | | | 3.476 | 6.92E-04 | 0.009 |
| | | 38,020,485 | *IKZF3* | | 3.962 | 1.22E-04 | 0.005 |
| | | 38,080,829 | | *ORMDL3* | 3.995 | 1.08E-04 | 0.014 |
| | | 38,096,333 | | | -3.664 | 3.60E-04 | 0.033 |
| | | 38,096,571 | | | -4.780 | 4.67E-06 | 0.033 |
| | | 38,096,578 | | | -5.642 | 1.00E-07 | 0.033 |
| | | 38,096,648 | | | -3.887 | 1.61E-04 | 0.033 |
| rs28896477 | g.38005534.T>C | 37,856,732 | | *ERBB2* | -3.668 | 3.56E-04 | 0.009 |
| | | 37,886,749 | | *MIEN1* | 3.945 | 1.30E-04 | 0.002 |
| | | 37,896,063 | | | 3.476 | 6.92E-04 | 0.009 |
| | | 38,020,485 | *IKZF3* | | 3.962 | 1.22E-04 | 0.005 |
| | | 38,080,829 | | *ORMDL3* | 3.995 | 1.08E-04 | 0.014 |
| | | 38,096,333 | | | -3.664 | 3.60E-04 | 0.033 |
| | | 38,096,571 | | | -4.780 | 4.67E-06 | 0.033 |
| | | 38,096,578 | | | -5.642 | 1.00E-07 | 0.033 |
| | | 38,096,648 | | | -3.887 | 1.61E-04 | 0.033 |
| rs28842324 | g.38005742.T>C | 37,856,732 | | *ERBB2* | -3.668 | 3.56E-04 | 0.009 |
| | | 37,886,749 | | *MIEN1* | 3.945 | 1.30E-04 | 0.002 |
| | | 37,896,063 | | | 3.476 | 6.92E-04 | 0.009 |
| | | 38,020,485 | *IKZF3* | | 3.962 | 1.22E-04 | 0.005 |
| | | 38,080,829 | | *ORMDL3* | 3.995 | 1.08E-04 | 0.014 |
| | | 38,096,333 | | | -3.664 | 3.60E-04 | 0.033 |
| | | 38,096,571 | | | -4.780 | 4.67E-06 | 0.033 |
| | | 38,096,578 | | | -5.642 | 1.00E-07 | 0.033 |
| | | 38,096,648 | | | -3.887 | 1.61E-04 | 0.033 |
| rs528792070 | g.38006700_38006700ins TAC | 37,856,732 | | *ERBB2* | -3.668 | 3.56E-04 | 0.009 |
| | | 37,886,749 | | *MIEN1* | 3.945 | 1.30E-04 | 0.002 |
| | | 37,896,063 | | | 3.476 | 6.92E-04 | 0.009 |

| | | | | | | | |
|---|---|---|---|---|---|---|---|
| | | 38,020,485 | *IKZF3* | | 3.962 | 1.22E-04 | 0.005 |
| | | 38,080,829 | | *ORMDL3* | 3.995 | 1.08E-04 | 0.014 |
| | | 38,096,333 | | | -3.664 | 3.60E-04 | 0.033 |
| | | 38,096,571 | | | -4.780 | 4.67E-06 | 0.033 |
| | | 38,096,578 | | | -5.642 | 1.00E-07 | 0.033 |
| | | 38,096,648 | | | -3.887 | 1.61E-04 | 0.033 |
| rs553196378 | g.38006967_38006969del | 37,856,732 | | *ERBB2* | -3.668 | 3.56E-04 | 0.009 |
| | | 37,886,749 | | *MIEN1* | 3.945 | 1.30E-04 | 0.002 |
| | | 37,896,063 | | | 3.476 | 6.92E-04 | 0.009 |
| | | 38,020,485 | *IKZF3* | | 3.962 | 1.22E-04 | 0.005 |
| | | 38,080,829 | | *ORMDL3* | 3.995 | 1.08E-04 | 0.014 |
| | | 38,096,333 | | | -3.664 | 3.60E-04 | 0.033 |
| | | 38,096,571 | | | -4.780 | 4.67E-06 | 0.033 |
| | | 38,096,578 | | | -5.642 | 1.00E-07 | 0.033 |
| | | 38,096,648 | | | -3.887 | 1.61E-04 | 0.033 |
| rs143123127 | g.38007190.A>G | 37,856,732 | | *ERBB2* | -3.668 | 3.56E-04 | 0.009 |
| | | 37,886,749 | | *MIEN1* | 3.945 | 1.30E-04 | 0.002 |
| | | 37,896,063 | | | 3.476 | 6.92E-04 | 0.009 |
| | | 38,020,485 | *IKZF3* | | 3.962 | 1.22E-04 | 0.005 |
| | | 38,080,829 | | *ORMDL3* | 3.995 | 1.08E-04 | 0.014 |
| | | 38,096,333 | | | -3.664 | 3.60E-04 | 0.033 |
| | | 38,096,571 | | | -4.780 | 4.67E-06 | 0.033 |
| | | 38,096,578 | | | -5.642 | 1.00E-07 | 0.033 |
| | | 38,096,648 | | | -3.887 | 1.61E-04 | 0.033 |
| rs202041850 | g.38007197_38007201del | 37,856,732 | | *ERBB2* | -3.668 | 3.56E-04 | 0.009 |
| | | 37,886,749 | | *MIEN1* | 3.945 | 1.30E-04 | 0.002 |
| | | 37,896,063 | | | 3.476 | 6.92E-04 | 0.009 |
| | | 38,020,485 | *IKZF3* | | 3.962 | 1.22E-04 | 0.005 |
| | | 38,080,829 | | *ORMDL3* | 3.995 | 1.08E-04 | 0.014 |
| | | 38,096,333 | | | -3.664 | 3.60E-04 | 0.033 |
| | | 38,096,571 | | | -4.780 | 4.67E-06 | 0.033 |
| | | 38,096,578 | | | -5.642 | 1.00E-07 | 0.033 |
| | | 38,096,648 | | | -3.887 | 1.61E-04 | 0.033 |
| rs114400736 | g.38007319.G>T | 37,856,732 | | *ERBB2* | -3.668 | 3.56E-04 | 0.009 |
| | | 37,886,749 | | *MIEN1* | 3.945 | 1.30E-04 | 0.002 |
| | | 37,896,063 | | | 3.476 | 6.92E-04 | 0.009 |
| | | 38,020,485 | *IKZF3* | | 3.962 | 1.22E-04 | 0.005 |
| | | 38,080,829 | | *ORMDL3* | 3.995 | 1.08E-04 | 0.014 |
| | | 38,096,333 | | | -3.664 | 3.60E-04 | 0.033 |

| | | | | | | | |
|---|---|---|---|---|---|---|---|
| | | 38,096,571 | | | -4.780 | 4.67E-06 | 0.033 |
| | | 38,096,578 | | | -5.642 | 1.00E-07 | 0.033 |
| | | 38,096,648 | | | -3.887 | 1.61E-04 | 0.033 |
| rs12938117 | g.38008165.T>C | 37,856,732 | | *ERBB2* | -3.668 | 3.56E-04 | 0.009 |
| | | 37,886,749 | | *MIEN1* | 3.945 | 1.30E-04 | 0.002 |
| | | 37,896,063 | | | 3.476 | 6.92E-04 | 0.009 |
| | | 38,020,485 | *IKZF3* | | 3.962 | 1.22E-04 | 0.005 |
| | | 38,080,829 | | *ORMDL3* | 3.995 | 1.08E-04 | 0.014 |
| | | 38,096,333 | | | -3.664 | 3.60E-04 | 0.033 |
| | | 38,096,571 | | | -4.780 | 4.67E-06 | 0.033 |
| | | 38,096,578 | | | -5.642 | 1.00E-07 | 0.033 |
| | | 38,096,648 | | | -3.887 | 1.61E-04 | 0.033 |
| rs113233720 | g.38008191.T>C | 37,856,732 | | *ERBB2* | -3.668 | 3.56E-04 | 0.009 |
| | | 37,886,749 | | *MIEN1* | 3.945 | 1.30E-04 | 0.002 |
| | | 37,896,063 | | | 3.476 | 6.92E-04 | 0.009 |
| | | 38,020,485 | *IKZF3* | | 3.962 | 1.22E-04 | 0.005 |
| | | 38,080,829 | | *ORMDL3* | 3.995 | 1.08E-04 | 0.014 |
| | | 38,096,333 | | | -3.664 | 3.60E-04 | 0.033 |
| | | 38,096,571 | | | -4.780 | 4.67E-06 | 0.033 |
| | | 38,096,578 | | | -5.642 | 1.00E-07 | 0.033 |
| | | 38,096,648 | | | -3.887 | 1.61E-04 | 0.033 |
| rs8068894 | g.38009000.G>A | 37,856,732 | | *ERBB2* | -3.668 | 3.56E-04 | 0.009 |
| | | 37,886,749 | | *MIEN1* | 3.945 | 1.30E-04 | 0.002 |
| | | 37,896,063 | | | 3.476 | 6.92E-04 | 0.009 |
| | | 38,020,485 | *IKZF3* | | 3.962 | 1.22E-04 | 0.005 |
| | | 38,080,829 | | *ORMDL3* | 3.995 | 1.08E-04 | 0.014 |
| | | 38,096,333 | | | -3.664 | 3.60E-04 | 0.033 |
| | | 38,096,571 | | | -4.780 | 4.67E-06 | 0.033 |
| | | 38,096,578 | | | -5.642 | 1.00E-07 | 0.033 |
| | | 38,096,648 | | | -3.887 | 1.61E-04 | 0.033 |
| rs202130396 | g.38009223_38009223ins A | 37,856,732 | | *ERBB2* | -3.667 | 3.56E-04 | 0.009 |
| | | 37,886,749 | | *MIEN1* | 3.944 | 1.30E-04 | 0.002 |
| | | 37,896,063 | | | 3.481 | 6.80E-04 | 0.009 |
| | | 38,020,485 | *IKZF3* | | 3.957 | 1.24E-04 | 0.005 |
| | | 38,080,829 | | *ORMDL3* | 3.999 | 1.06E-04 | 0.014 |
| | | 38,096,333 | | | -3.662 | 3.63E-04 | 0.033 |
| | | 38,096,571 | | | -4.784 | 4.59E-06 | 0.033 |
| | | 38,096,578 | | | -5.649 | 9.73E-08 | 0.033 |
| | | 38,096,648 | | | -3.894 | 1.57E-04 | 0.033 |
| rs8069531 | g.38009344.T>A | 37,856,732 | | *ERBB2* | -3.668 | 3.56E-04 | 0.009 |

| | | | | | | | |
|---|---|---|---|---|---|---|---|
| | | 37,886,749 | | *MIEN1* | 3.945 | 1.30E-04 | 0.002 |
| | | 37,896,063 | | | 3.476 | 6.92E-04 | 0.009 |
| | | 38,020,485 | *IKZF3* | | 3.962 | 1.22E-04 | 0.005 |
| | | 38,080,829 | | *ORMDL3* | 3.995 | 1.08E-04 | 0.014 |
| | | 38,096,333 | | | -3.664 | 3.60E-04 | 0.033 |
| | | 38,096,571 | | | -4.780 | 4.67E-06 | 0.033 |
| | | 38,096,578 | | | -5.642 | 1.00E-07 | 0.033 |
| | | 38,096,648 | | | -3.887 | 1.61E-04 | 0.033 |
| rs79113275 | g.38009350.T>C | 37,886,872 | *MIEN1* | | 4.778 | 4.70E-06 | 0.002 |
| | | 37,910,923 | | | 4.844 | 3.55E-06 | 0.035 |
| | | 37,911,173 | | | 6.613 | 8.87E-10 | 0.035 |
| | | 37,947,519 | | *IKZF3* | -3.577 | 4.89E-04 | 0.002 |
| | | 38,020,606 | *IKZF3* | | 3.419 | 8.41E-04 | 0.017 |
| | | 38,109,653 | | | 5.448 | 2.46E-07 | 0.002 |
| rs12947480 | g.38009676.A>G | 37,856,732 | | *ERBB2* | -3.668 | 3.56E-04 | 0.009 |
| | | 37,886,749 | | *MIEN1* | 3.945 | 1.30E-04 | 0.002 |
| | | 37,896,063 | | | 3.476 | 6.92E-04 | 0.009 |
| | | 38,020,485 | *IKZF3* | | 3.962 | 1.22E-04 | 0.005 |
| | | 38,080,829 | | *ORMDL3* | 3.995 | 1.08E-04 | 0.014 |
| | | 38,096,333 | | | -3.664 | 3.60E-04 | 0.033 |
| | | 38,096,571 | | | -4.780 | 4.67E-06 | 0.033 |
| | | 38,096,578 | | | -5.642 | 1.00E-07 | 0.033 |
| | | 38,096,648 | | | -3.887 | 1.61E-04 | 0.033 |
| rs9907291 | g.38010037.G>A | 37,856,732 | | *ERBB2* | -3.668 | 3.56E-04 | 0.009 |
| | | 37,886,749 | | *MIEN1* | 3.945 | 1.30E-04 | 0.002 |
| | | 37,896,063 | | | 3.476 | 6.92E-04 | 0.009 |
| | | 38,020,485 | *IKZF3* | | 3.962 | 1.22E-04 | 0.005 |
| | | 38,080,829 | | *ORMDL3* | 3.995 | 1.08E-04 | 0.014 |
| | | 38,096,333 | | | -3.664 | 3.60E-04 | 0.033 |
| | | 38,096,571 | | | -4.780 | 4.67E-06 | 0.033 |
| | | 38,096,578 | | | -5.642 | 1.00E-07 | 0.033 |
| | | 38,096,648 | | | -3.887 | 1.61E-04 | 0.033 |
| rs9907339 | g.38010103.C>A | 37,856,732 | | *ERBB2* | -3.668 | 3.56E-04 | 0.009 |
| | | 37,886,749 | | *MIEN1* | 3.945 | 1.30E-04 | 0.002 |
| | | 37,896,063 | | | 3.476 | 6.92E-04 | 0.009 |
| | | 38,020,485 | *IKZF3* | | 3.962 | 1.22E-04 | 0.005 |
| | | 38,080,829 | | *ORMDL3* | 3.995 | 1.08E-04 | 0.014 |
| | | 38,096,333 | | | -3.664 | 3.60E-04 | 0.033 |
| | | 38,096,571 | | | -4.780 | 4.67E-06 | 0.033 |
| | | 38,096,578 | | | -5.642 | 1.00E-07 | 0.033 |
| | | 38,096,648 | | | -3.887 | 1.61E-04 | 0.033 |

| | | | | | | | |
|---|---|---|---|---|---|---|---|
| rs9907498 | g.38010104.G>A | 37,856,732 | | *ERBB2* | -3.668 | 3.56E-04 | 0.009 |
| | | 37,886,749 | | *MIEN1* | 3.945 | 1.30E-04 | 0.002 |
| | | 37,896,063 | | | 3.476 | 6.92E-04 | 0.009 |
| | | 38,020,485 | *IKZF3* | | 3.962 | 1.22E-04 | 0.005 |
| | | 38,080,829 | | *ORMDL3* | 3.995 | 1.08E-04 | 0.014 |
| | | 38,096,333 | | | -3.664 | 3.60E-04 | 0.033 |
| | | 38,096,571 | | | -4.780 | 4.67E-06 | 0.033 |
| | | 38,096,578 | | | -5.642 | 1.00E-07 | 0.033 |
| | | 38,096,648 | | | -3.887 | 1.61E-04 | 0.033 |
| rs8077365 | g.38010758.T>G | 37,856,732 | | *ERBB2* | -3.694 | 3.23E-04 | 0.009 |
| | | 37,879,682 | | *ERBB2* | 3.380 | 9.58E-04 | 0.002 |
| | | 37,896,063 | | | 3.678 | 3.42E-04 | 0.009 |
| | | 38,020,485 | *IKZF3* | | 3.611 | 4.35E-04 | 0.005 |
| | | 38,096,333 | | | -3.536 | 5.63E-04 | 0.033 |
| | | 38,096,578 | | | -4.659 | 7.76E-06 | 0.033 |
| rs8078409 | g.38010759.T>C | 37,856,732 | | *ERBB2* | -3.694 | 3.23E-04 | 0.009 |
| | | 37,879,682 | | *ERBB2* | 3.380 | 9.58E-04 | 0.002 |
| | | 37,896,063 | | | 3.678 | 3.42E-04 | 0.009 |
| | | 38,020,485 | *IKZF3* | | 3.611 | 4.35E-04 | 0.005 |
| | | 38,096,333 | | | -3.536 | 5.63E-04 | 0.033 |
| | | 38,096,578 | | | -4.659 | 7.76E-06 | 0.033 |
| rs8079075 | g.38010815.G>A | 37,856,732 | | *ERBB2* | -3.668 | 3.56E-04 | 0.009 |
| | | 37,886,749 | | *MIEN1* | 3.945 | 1.30E-04 | 0.002 |
| | | 37,896,063 | | | 3.476 | 6.92E-04 | 0.009 |
| | | 38,020,485 | *IKZF3* | | 3.962 | 1.22E-04 | 0.005 |
| | | 38,080,829 | | *ORMDL3* | 3.995 | 1.08E-04 | 0.014 |
| | | 38,096,333 | | | -3.664 | 3.60E-04 | 0.033 |
| | | 38,096,571 | | | -4.780 | 4.67E-06 | 0.033 |
| | | 38,096,578 | | | -5.642 | 1.00E-07 | 0.033 |
| | | 38,096,648 | | | -3.887 | 1.61E-04 | 0.033 |
| rs28621669 | g.38011266.C>T | 37,856,732 | | *ERBB2* | -3.668 | 3.56E-04 | 0.009 |
| | | 37,886,749 | | *MIEN1* | 3.945 | 1.30E-04 | 0.002 |
| | | 37,896,063 | | | 3.476 | 6.92E-04 | 0.009 |
| | | 38,020,485 | *IKZF3* | | 3.962 | 1.22E-04 | 0.005 |
| | | 38,080,829 | | *ORMDL3* | 3.995 | 1.08E-04 | 0.014 |
| | | 38,096,333 | | | -3.664 | 3.60E-04 | 0.033 |
| | | 38,096,571 | | | -4.780 | 4.67E-06 | 0.033 |
| | | 38,096,578 | | | -5.642 | 1.00E-07 | 0.033 |
| | | 38,096,648 | | | -3.887 | 1.61E-04 | 0.033 |
| rs28723017 | g.38011451.T>C | 37,856,732 | | *ERBB2* | -3.668 | 3.56E-04 | 0.009 |
| | | 37,886,749 | | *MIEN1* | 3.945 | 1.30E-04 | 0.002 |

| | | | | | | | |
|---|---|---|---|---|---|---|---|
| | | 37,896,063 | | | 3.476 | 6.92E-04 | 0.009 |
| | | 38,020,485 | *IKZF3* | | 3.962 | 1.22E-04 | 0.005 |
| | | 38,080,829 | | *ORMDL3* | 3.995 | 1.08E-04 | 0.014 |
| | | 38,096,333 | | | -3.664 | 3.60E-04 | 0.033 |
| | | 38,096,571 | | | -4.780 | 4.67E-06 | 0.033 |
| | | 38,096,578 | | | -5.642 | 1.00E-07 | 0.033 |
| | | 38,096,648 | | | -3.887 | 1.61E-04 | 0.033 |
| rs9900538 | g.38011709.C>T | 37,856,732 | | *ERBB2* | -3.668 | 3.56E-04 | 0.009 |
| | | 37,886,749 | | *MIEN1* | 3.945 | 1.30E-04 | 0.002 |
| | | 37,896,063 | | | 3.476 | 6.92E-04 | 0.009 |
| | | 38,020,485 | *IKZF3* | | 3.962 | 1.22E-04 | 0.005 |
| | | 38,080,829 | | *ORMDL3* | 3.995 | 1.08E-04 | 0.014 |
| | | 38,096,333 | | | -3.664 | 3.60E-04 | 0.033 |
| | | 38,096,571 | | | -4.780 | 4.67E-06 | 0.033 |
| | | 38,096,578 | | | -5.642 | 1.00E-07 | 0.033 |
| | | 38,096,648 | | | -3.887 | 1.61E-04 | 0.033 |
| rs546918444 | g.38012317.C>A | 37,879,682 | | *ERBB2* | 3.886 | 1.61E-04 | 0.002 |
| | | 37,896,063 | | | 4.183 | 5.25E-05 | 0.009 |
| | | 38,083,269 | | *ORMDL3* | 4.158 | 5.79E-05 | 0.011 |
| | | 38,108,720 | | | 4.114 | 6.86E-05 | 0.039 |
| | | 38,119,834 | | *GSDMA* | 3.383 | 9.47E-04 | 0.011 |
| rs559395711 | g.38012318.A>C | 37,879,682 | | *ERBB2* | 3.886 | 1.61E-04 | 0.002 |
| | | 37,896,063 | | | 4.183 | 5.25E-05 | 0.009 |
| | | 38,083,269 | | *ORMDL3* | 4.158 | 5.79E-05 | 0.011 |
| | | 38,108,720 | | | 4.114 | 6.86E-05 | 0.039 |
| | | 38,119,834 | | *GSDMA* | 3.383 | 9.47E-04 | 0.011 |
| rs113466546 | g.38012587.A>G | 37,856,732 | | *ERBB2* | -3.668 | 3.56E-04 | 0.009 |
| | | 37,886,749 | | *MIEN1* | 3.945 | 1.30E-04 | 0.002 |
| | | 37,896,063 | | | 3.476 | 6.92E-04 | 0.009 |
| | | 38,020,485 | *IKZF3* | | 3.962 | 1.22E-04 | 0.005 |
| | | 38,080,829 | | *ORMDL3* | 3.995 | 1.08E-04 | 0.014 |
| | | 38,096,333 | | | -3.664 | 3.60E-04 | 0.033 |
| | | 38,096,571 | | | -4.780 | 4.67E-06 | 0.033 |
| | | 38,096,578 | | | -5.642 | 1.00E-07 | 0.033 |
| | | 38,096,648 | | | -3.887 | 1.61E-04 | 0.033 |
| rs35770042 | g.38012991.G>A | 37,856,732 | | *ERBB2* | -3.668 | 3.56E-04 | 0.009 |
| | | 37,886,749 | | *MIEN1* | 3.945 | 1.30E-04 | 0.002 |
| | | 37,896,063 | | | 3.476 | 6.92E-04 | 0.009 |
| | | 38,020,485 | *IKZF3* | | 3.962 | 1.22E-04 | 0.005 |
| | | 38,080,829 | | *ORMDL3* | 3.995 | 1.08E-04 | 0.014 |
| | | 38,096,333 | | | -3.664 | 3.60E-04 | 0.033 |

| | | | | | | | |
|---|---|---|---|---|---|---|---|
| | | 38,096,571 | | | -4.780 | 4.67E-06 | 0.033 |
| | | 38,096,578 | | | -5.642 | 1.00E-07 | 0.033 |
| | | 38,096,648 | | | -3.887 | 1.61E-04 | 0.033 |
| rs118017471 | g.38013360.A>G | 37,832,244 | | *PGAP3* | 3.805 | 2.18E-04 | 0.028 |
| | | 37,844,412 | *PGAP3* | *ERBB2* | 3.384 | 9.44E-04 | 7.82E-05 |
| | | 37,895,409 | | | -4.216 | 4.63E-05 | 0.006 |
| | | 37,910,347 | | | 4.937 | 2.39E-06 | 0.013 |
| | | 37,911,853 | | | -3.692 | 3.27E-04 | 0.035 |
| | | 37,933,052 | | *IKZF3* | 3.654 | 3.73E-04 | 0.027 |
| | | 38,020,220 | | *IKZF3* | 3.399 | 8.97E-04 | 0.046 |
| | | 38,022,345 | | | 3.422 | 8.30E-04 | 0.003 |
| | | 38,024,014 | | | -6.059 | 1.38E-08 | 0.004 |
| | | 38,024,237 | | | -5.276 | 5.36E-07 | 0.016 |
| | | 38,024,242 | | | -4.433 | 1.95E-05 | 0.016 |
| | | 38,024,244 | | | -4.214 | 4.67E-05 | 0.016 |
| | | 38,083,894 | *ORMDL3* | | 4.544 | 1.25E-05 | 0.011 |
| rs28661421 | g.38013807.G>A | 37,856,732 | | *ERBB2* | -3.668 | 3.56E-04 | 0.009 |
| | | 37,886,749 | | *MIEN1* | 3.945 | 1.30E-04 | 0.002 |
| | | 37,896,063 | | | 3.476 | 6.92E-04 | 0.009 |
| | | 38,020,485 | *IKZF3* | | 3.962 | 1.22E-04 | 0.005 |
| | | 38,080,829 | | *ORMDL3* | 3.995 | 1.08E-04 | 0.014 |
| | | 38,096,333 | | | -3.664 | 3.60E-04 | 0.033 |
| | | 38,096,571 | | | -4.780 | 4.67E-06 | 0.033 |
| | | 38,096,578 | | | -5.642 | 1.00E-07 | 0.033 |
| | | 38,096,648 | | | -3.887 | 1.61E-04 | 0.033 |
| rs4795396 | g.38013982.T>C | 37,832,464 | | *PGAP3* | 3.560 | 5.18E-04 | 0.028 |
| | | 37,844,223 | *ERBB2* | *PGAP3* | 3.400 | 8.95E-04 | 7.82E-05 |
| | | 37,844,365 | *PGAP3ERBB2* | | 3.793 | 2.27E-04 | 7.82E-05 |
| | | 38,077,464 | | *ORMDL3* | 6.661 | 6.98E-10 | 0.014 |
| rs16965367 | g.38014316.C>T | 37,856,732 | | *ERBB2* | -3.668 | 3.56E-04 | 0.009 |
| | | 37,886,749 | | *MIEN1* | 3.945 | 1.30E-04 | 0.002 |
| | | 37,896,063 | | | 3.476 | 6.92E-04 | 0.009 |
| | | 38,020,485 | *IKZF3* | | 3.962 | 1.22E-04 | 0.005 |
| | | 38,080,829 | | *ORMDL3* | 3.995 | 1.08E-04 | 0.014 |
| | | 38,096,333 | | | -3.664 | 3.60E-04 | 0.033 |
| | | 38,096,571 | | | -4.780 | 4.67E-06 | 0.033 |
| | | 38,096,578 | | | -5.642 | 1.00E-07 | 0.033 |
| | | 38,096,648 | | | -3.887 | 1.61E-04 | 0.033 |
| rs9912095 | g.38014533.G>A | 37,856,732 | | *ERBB2* | -3.668 | 3.56E-04 | 0.009 |
| | | 37,886,749 | | *MIEN1* | 3.945 | 1.30E-04 | 0.002 |
| | | 37,896,063 | | | 3.476 | 6.92E-04 | 0.009 |

| | | | | | | | |
|---|---|---|---|---|---|---|---|
| | | 38,020,485 | *IKZF3* | | 3.962 | 1.22E-04 | 0.005 |
| | | 38,080,829 | | *ORMDL3* | 3.995 | 1.08E-04 | 0.014 |
| | | 38,096,333 | | | -3.664 | 3.60E-04 | 0.033 |
| | | 38,096,571 | | | -4.780 | 4.67E-06 | 0.033 |
| | | 38,096,578 | | | -5.642 | 1.00E-07 | 0.033 |
| | | 38,096,648 | | | -3.887 | 1.61E-04 | 0.033 |
| rs9915509 | g.38014705.A>G | 37,856,732 | | *ERBB2* | -3.668 | 3.56E-04 | 0.009 |
| | | 37,886,749 | | *MIEN1* | 3.945 | 1.30E-04 | 0.002 |
| | | 37,896,063 | | | 3.476 | 6.92E-04 | 0.009 |
| | | 38,020,485 | *IKZF3* | | 3.962 | 1.22E-04 | 0.005 |
| | | 38,080,829 | | *ORMDL3* | 3.995 | 1.08E-04 | 0.014 |
| | | 38,096,333 | | | -3.664 | 3.60E-04 | 0.033 |
| | | 38,096,571 | | | -4.780 | 4.67E-06 | 0.033 |
| | | 38,096,578 | | | -5.642 | 1.00E-07 | 0.033 |
| | | 38,096,648 | | | -3.887 | 1.61E-04 | 0.033 |
| rs201152777 | g.38014758_38014758insT | 37,856,732 | | *ERBB2* | -3.668 | 3.56E-04 | 0.009 |
| | | 37,886,749 | | *MIEN1* | 3.945 | 1.30E-04 | 0.002 |
| | | 37,896,063 | | | 3.476 | 6.92E-04 | 0.009 |
| | | 38,020,485 | *IKZF3* | | 3.962 | 1.22E-04 | 0.005 |
| | | 38,080,829 | | *ORMDL3* | 3.995 | 1.08E-04 | 0.014 |
| | | 38,096,333 | | | -3.664 | 3.60E-04 | 0.033 |
| | | 38,096,571 | | | -4.780 | 4.67E-06 | 0.033 |
| | | 38,096,578 | | | -5.642 | 1.00E-07 | 0.033 |
| | | 38,096,648 | | | -3.887 | 1.61E-04 | 0.033 |
| rs9915797 | g.38014868.A>G | 37,856,732 | | *ERBB2* | -3.668 | 3.56E-04 | 0.009 |
| | | 37,886,749 | | *MIEN1* | 3.945 | 1.30E-04 | 0.002 |
| | | 37,896,063 | | | 3.476 | 6.92E-04 | 0.009 |
| | | 38,020,485 | *IKZF3* | | 3.962 | 1.22E-04 | 0.005 |
| | | 38,080,829 | | *ORMDL3* | 3.995 | 1.08E-04 | 0.014 |
| | | 38,096,333 | | | -3.664 | 3.60E-04 | 0.033 |
| | | 38,096,571 | | | -4.780 | 4.67E-06 | 0.033 |
| | | 38,096,578 | | | -5.642 | 1.00E-07 | 0.033 |
| | | 38,096,648 | | | -3.887 | 1.61E-04 | 0.033 |
| rs9916259 | g.38015060.T>G | 37,856,732 | | *ERBB2* | -3.668 | 3.56E-04 | 0.009 |
| | | 37,886,749 | | *MIEN1* | 3.945 | 1.30E-04 | 0.002 |
| | | 37,896,063 | | | 3.476 | 6.92E-04 | 0.009 |
| | | 38,020,485 | *IKZF3* | | 3.962 | 1.22E-04 | 0.005 |
| | | 38,080,829 | | *ORMDL3* | 3.995 | 1.08E-04 | 0.014 |
| | | 38,096,333 | | | -3.664 | 3.60E-04 | 0.033 |
| | | 38,096,571 | | | -4.780 | 4.67E-06 | 0.033 |

| | | | | | | | |
|---|---|---|---|---|---|---|---|
| | | 38,096,578 | | | -5.642 | 1.00E-07 | 0.033 |
| | | 38,096,648 | | | -3.887 | 1.61E-04 | 0.033 |
| rs142025263 | g.38015188_38015188ins ACT | 37,856,732 | | *ERBB2* | -3.668 | 3.56E-04 | 0.009 |
| | | 37,886,749 | | *MIEN1* | 3.945 | 1.30E-04 | 0.002 |
| | | 37,896,063 | | | 3.476 | 6.92E-04 | 0.009 |
| | | 38,020,485 | *IKZF3* | | 3.962 | 1.22E-04 | 0.005 |
| | | 38,080,829 | | *ORMDL3* | 3.995 | 1.08E-04 | 0.014 |
| | | 38,096,333 | | | -3.664 | 3.60E-04 | 0.033 |
| | | 38,096,571 | | | -4.780 | 4.67E-06 | 0.033 |
| | | 38,096,578 | | | -5.642 | 1.00E-07 | 0.033 |
| | | 38,096,648 | | | -3.887 | 1.61E-04 | 0.033 |
| rs8068125 | g.38015517.C>T | 37,856,732 | | *ERBB2* | -3.668 | 3.56E-04 | 0.009 |
| | | 37,886,749 | | *MIEN1* | 3.945 | 1.30E-04 | 0.002 |
| | | 37,896,063 | | | 3.476 | 6.92E-04 | 0.009 |
| | | 38,020,485 | *IKZF3* | | 3.962 | 1.22E-04 | 0.005 |
| | | 38,080,829 | | *ORMDL3* | 3.995 | 1.08E-04 | 0.014 |
| | | 38,096,333 | | | -3.664 | 3.60E-04 | 0.033 |
| | | 38,096,571 | | | -4.780 | 4.67E-06 | 0.033 |
| | | 38,096,578 | | | -5.642 | 1.00E-07 | 0.033 |
| | | 38,096,648 | | | -3.887 | 1.61E-04 | 0.033 |
| rs77924338 | g.38016357.T>C | 37,856,732 | | *ERBB2* | -3.668 | 3.56E-04 | 0.009 |
| | | 37,886,749 | | *MIEN1* | 3.945 | 1.30E-04 | 0.002 |
| | | 37,896,063 | | | 3.476 | 6.92E-04 | 0.009 |
| | | 38,020,485 | *IKZF3* | | 3.962 | 1.22E-04 | 0.005 |
| | | 38,080,829 | | *ORMDL3* | 3.995 | 1.08E-04 | 0.014 |
| | | 38,096,333 | | | -3.664 | 3.60E-04 | 0.033 |
| | | 38,096,571 | | | -4.780 | 4.67E-06 | 0.033 |
| | | 38,096,578 | | | -5.642 | 1.00E-07 | 0.033 |
| | | 38,096,648 | | | -3.887 | 1.61E-04 | 0.033 |
| rs9899006 | g.38017065.A>T | 37,856,732 | | *ERBB2* | -3.668 | 3.56E-04 | 0.009 |
| | | 37,886,749 | | *MIEN1* | 3.945 | 1.30E-04 | 0.002 |
| | | 37,896,063 | | | 3.476 | 6.92E-04 | 0.009 |
| | | 38,020,485 | *IKZF3* | | 3.962 | 1.22E-04 | 0.005 |
| | | 38,080,829 | | *ORMDL3* | 3.995 | 1.08E-04 | 0.014 |
| | | 38,096,333 | | | -3.664 | 3.60E-04 | 0.033 |
| | | 38,096,571 | | | -4.780 | 4.67E-06 | 0.033 |
| | | 38,096,578 | | | -5.642 | 1.00E-07 | 0.033 |
| | | 38,096,648 | | | -3.887 | 1.61E-04 | 0.033 |
| rs67837020 | g.38017313_38017320del | 37,856,732 | | *ERBB2* | -3.668 | 3.56E-04 | 0.009 |

| | | | | | | | |
|---|---|---|---|---|---|---|---|
| | | 37,886,749 | | *MIEN1* | 3.945 | 1.30E-04 | 0.002 |
| | | 37,896,063 | | | 3.476 | 6.92E-04 | 0.009 |
| | | 38,020,485 | *IKZF3* | | 3.962 | 1.22E-04 | 0.005 |
| | | 38,080,829 | | *ORMDL3* | 3.995 | 1.08E-04 | 0.014 |
| | | 38,096,333 | | | -3.664 | 3.60E-04 | 0.033 |
| | | 38,096,571 | | | -4.780 | 4.67E-06 | 0.033 |
| | | 38,096,578 | | | -5.642 | 1.00E-07 | 0.033 |
| | | 38,096,648 | | | -3.887 | 1.61E-04 | 0.033 |
| rs112773712 | g.38017704.A>G | 37,886,788 | *MIEN1* | *MIEN1* | 4.149 | 6.00E-05 | 0.002 |
| | | 38,024,592 | | | 4.400 | 2.23E-05 | 0.046 |
| | | 38,024,606 | | | 4.304 | 3.27E-05 | 0.046 |
| | | 38,083,737 | | *ORMDL3* | 3.512 | 6.13E-04 | 0.011 |
| | | 38,096,333 | | | -3.527 | 5.80E-04 | 0.033 |
| | | 38,096,571 | | | -4.910 | 2.68E-06 | 0.033 |
| | | 38,096,578 | | | -4.517 | 1.39E-05 | 0.033 |
| rs9899336 | g.38017780.T>C | 37,856,732 | | *ERBB2* | -3.668 | 3.56E-04 | 0.009 |
| | | 37,886,749 | | *MIEN1* | 3.945 | 1.30E-04 | 0.002 |
| | | 37,896,063 | | | 3.476 | 6.92E-04 | 0.009 |
| | | 38,020,485 | *IKZF3* | | 3.962 | 1.22E-04 | 0.005 |
| | | 38,080,829 | | *ORMDL3* | 3.995 | 1.08E-04 | 0.014 |
| | | 38,096,333 | | | -3.664 | 3.60E-04 | 0.033 |
| | | 38,096,571 | | | -4.780 | 4.67E-06 | 0.033 |
| | | 38,096,578 | | | -5.642 | 1.00E-07 | 0.033 |
| | | 38,096,648 | | | -3.887 | 1.61E-04 | 0.033 |
| rs77137867 | g.38017969.T>C | 37,836,263 | | *PGAP3* | -3.933 | 1.36E-04 | 0.019 |
| | | 37,886,619 | | *MIEN1* | 3.604 | 4.44E-04 | 0.002 |
| | | 37,910,190 | | | 4.487 | 1.57E-05 | 0.043 |
| rs202227901 | g.38018498_38018500del | 37,894,916 | | | 3.440 | 7.83E-04 | 0.004 |
| | | 37,934,107 | | *IKZF3* | -4.490 | 1.55E-05 | 0.027 |
| | | 38,084,016 | *ORMDL3* | | 4.466 | 1.72E-05 | 0.011 |
| | | 38,109,131 | | | 4.741 | 5.50E-06 | 0.002 |
| rs9907096 | g.38018774.T>C | 37,856,732 | | *ERBB2* | -3.668 | 3.56E-04 | 0.009 |
| | | 37,886,749 | | *MIEN1* | 3.945 | 1.30E-04 | 0.002 |
| | | 37,896,063 | | | 3.476 | 6.92E-04 | 0.009 |
| | | 38,020,485 | *IKZF3* | | 3.962 | 1.22E-04 | 0.005 |
| | | 38,080,829 | | *ORMDL3* | 3.995 | 1.08E-04 | 0.014 |
| | | 38,096,333 | | | -3.664 | 3.60E-04 | 0.033 |
| | | 38,096,571 | | | -4.780 | 4.67E-06 | 0.033 |
| | | 38,096,578 | | | -5.642 | 1.00E-07 | 0.033 |
| | | 38,096,648 | | | -3.887 | 1.61E-04 | 0.033 |

| | | | | | | |
|---|---|---|---|---|---|---|
| rs9907966 | g.38018803.C>A | 37,856,732 | *ERBB2* | -3.694 | 3.23E-04 | 0.009 |
| | | 37,879,682 | *ERBB2* | 3.380 | 9.58E-04 | 0.002 |
| | | 37,896,063 | | 3.678 | 3.42E-04 | 0.009 |
| | | 38,020,485 | *IKZF3* | 3.611 | 4.35E-04 | 0.005 |
| | | 38,096,333 | | -3.536 | 5.63E-04 | 0.033 |
| | | 38,096,578 | | -4.659 | 7.76E-06 | 0.033 |
| rs9905881 | g.38018955.A>G | 37,856,732 | *ERBB2* | -3.668 | 3.56E-04 | 0.009 |
| | | 37,886,749 | *MIEN1* | 3.945 | 1.30E-04 | 0.002 |
| | | 37,896,063 | | 3.476 | 6.92E-04 | 0.009 |
| | | 38,020,485 | *IKZF3* | 3.962 | 1.22E-04 | 0.005 |
| | | 38,080,829 | *ORMDL3* | 3.995 | 1.08E-04 | 0.014 |
| | | 38,096,333 | | -3.664 | 3.60E-04 | 0.033 |
| | | 38,096,571 | | -4.780 | 4.67E-06 | 0.033 |
| | | 38,096,578 | | -5.642 | 1.00E-07 | 0.033 |
| | | 38,096,648 | | -3.887 | 1.61E-04 | 0.033 |
| rs9907564 | g.38018967.T>C | 37,856,732 | *ERBB2* | -3.668 | 3.56E-04 | 0.009 |
| | | 37,886,749 | *MIEN1* | 3.945 | 1.30E-04 | 0.002 |
| | | 37,896,063 | | 3.476 | 6.92E-04 | 0.009 |
| | | 38,020,485 | *IKZF3* | 3.962 | 1.22E-04 | 0.005 |
| | | 38,080,829 | *ORMDL3* | 3.995 | 1.08E-04 | 0.014 |
| | | 38,096,333 | | -3.664 | 3.60E-04 | 0.033 |
| | | 38,096,571 | | -4.780 | 4.67E-06 | 0.033 |
| | | 38,096,578 | | -5.642 | 1.00E-07 | 0.033 |
| | | 38,096,648 | | -3.887 | 1.61E-04 | 0.033 |
| rs9907794 | g.38019026.G>C | 37,856,732 | *ERBB2* | -3.668 | 3.56E-04 | 0.009 |
| | | 37,886,749 | *MIEN1* | 3.945 | 1.30E-04 | 0.002 |
| | | 37,896,063 | | 3.476 | 6.92E-04 | 0.009 |
| | | 38,020,485 | *IKZF3* | 3.962 | 1.22E-04 | 0.005 |
| | | 38,080,829 | *ORMDL3* | 3.995 | 1.08E-04 | 0.014 |
| | | 38,096,333 | | -3.664 | 3.60E-04 | 0.033 |
| | | 38,096,571 | | -4.780 | 4.67E-06 | 0.033 |
| | | 38,096,578 | | -5.642 | 1.00E-07 | 0.033 |
| | | 38,096,648 | | -3.887 | 1.61E-04 | 0.033 |
| rs188353899 | g.38019856.T>G | 37,886,725 | *MIEN1* | 3.428 | 8.16E-04 | 0.002 |
| | | 37,886,742 | *MIEN1* | 3.445 | 7.70E-04 | 0.002 |
| | | 37,893,894 | | 4.943 | 2.33E-06 | 0.002 |
| | | 37,896,050 | | 5.253 | 5.94E-07 | 0.027 |
| | | 37,896,705 | | 4.320 | 3.07E-05 | 0.008 |
| | | 37,910,542 | | 3.878 | 1.66E-04 | 0.013 |
| | | 37,910,569 | | 3.428 | 8.16E-04 | 0.013 |
| | | 38,083,656 | *ORMDL3* | 5.520 | 1.77E-07 | 0.011 |

| | | | | | | | |
|---|---|---|---|---|---|---|---|
| | | 38,084,256 | *ORMDL3* | | -3.681 | 3.39E-04 | 0.005 |
| | | 38,115,535 | | | 3.527 | 5.82E-04 | 0.009 |
| rs111678394 | g.38021117.C>G | 37,856,732 | | *ERBB2* | -3.668 | 3.56E-04 | 0.009 |
| | | 37,886,749 | | *MIEN1* | 3.945 | 1.30E-04 | 0.002 |
| | | 37,896,063 | | | 3.476 | 6.92E-04 | 0.009 |
| | | 38,020,485 | *IKZF3* | | 3.962 | 1.22E-04 | 0.005 |
| | | 38,080,829 | | *ORMDL3* | 3.995 | 1.08E-04 | 0.014 |
| | | 38,096,333 | | | -3.664 | 3.60E-04 | 0.033 |
| | | 38,096,571 | | | -4.780 | 4.67E-06 | 0.033 |
| | | 38,096,578 | | | -5.642 | 1.00E-07 | 0.033 |
| | | 38,096,648 | | | -3.887 | 1.61E-04 | 0.033 |
| rs11284476 | g.38022534_38022535del | 37,856,732 | | *ERBB2* | -3.668 | 3.56E-04 | 0.009 |
| | | 37,886,749 | | *MIEN1* | 3.945 | 1.30E-04 | 0.002 |
| | | 37,896,063 | | | 3.476 | 6.92E-04 | 0.009 |
| | | 38,020,485 | *IKZF3* | | 3.962 | 1.22E-04 | 0.005 |
| | | 38,080,829 | | *ORMDL3* | 3.995 | 1.08E-04 | 0.014 |
| | | 38,096,333 | | | -3.664 | 3.60E-04 | 0.033 |
| | | 38,096,571 | | | -4.780 | 4.67E-06 | 0.033 |
| | | 38,096,578 | | | -5.642 | 1.00E-07 | 0.033 |
| | | 38,096,648 | | | -3.887 | 1.61E-04 | 0.033 |
| rs4622539 | g.38023031.A>T | 37,856,732 | | *ERBB2* | -3.668 | 3.56E-04 | 0.009 |
| | | 37,886,749 | | *MIEN1* | 3.945 | 1.30E-04 | 0.002 |
| | | 37,896,063 | | | 3.476 | 6.92E-04 | 0.009 |
| | | 38,020,485 | *IKZF3* | | 3.962 | 1.22E-04 | 0.005 |
| | | 38,080,829 | | *ORMDL3* | 3.995 | 1.08E-04 | 0.014 |
| | | 38,096,333 | | | -3.664 | 3.60E-04 | 0.033 |
| | | 38,096,571 | | | -4.780 | 4.67E-06 | 0.033 |
| | | 38,096,578 | | | -5.642 | 1.00E-07 | 0.033 |
| | | 38,096,648 | | | -3.887 | 1.61E-04 | 0.033 |
| rs1453560 | g.38023441.C>A | 37,856,732 | | *ERBB2* | -3.668 | 3.56E-04 | 0.009 |
| | | 37,886,749 | | *MIEN1* | 3.945 | 1.30E-04 | 0.002 |
| | | 37,896,063 | | | 3.476 | 6.92E-04 | 0.009 |
| | | 38,020,485 | *IKZF3* | | 3.962 | 1.22E-04 | 0.005 |
| | | 38,080,829 | | *ORMDL3* | 3.995 | 1.08E-04 | 0.014 |
| | | 38,096,333 | | | -3.664 | 3.60E-04 | 0.033 |
| | | 38,096,571 | | | -4.780 | 4.67E-06 | 0.033 |
| | | 38,096,578 | | | -5.642 | 1.00E-07 | 0.033 |
| | | 38,096,648 | | | -3.887 | 1.61E-04 | 0.033 |
| rs112880843 | g.38024298.A>G | 37,856,732 | | *ERBB2* | -3.668 | 3.56E-04 | 0.009 |
| | | 37,886,749 | | *MIEN1* | 3.945 | 1.30E-04 | 0.002 |

| | | | | | | | |
|---|---|---|---|---|---|---|---|
| | | 37,896,063 | | | 3.476 | 6.92E-04 | 0.009 |
| | | 38,020,485 | *IKZF3* | | 3.962 | 1.22E-04 | 0.005 |
| | | 38,080,829 | | *ORMDL3* | 3.995 | 1.08E-04 | 0.014 |
| | | 38,096,333 | | | -3.664 | 3.60E-04 | 0.033 |
| | | 38,096,571 | | | -4.780 | 4.67E-06 | 0.033 |
| | | 38,096,578 | | | -5.642 | 1.00E-07 | 0.033 |
| | | 38,096,648 | | | -3.887 | 1.61E-04 | 0.033 |
| rs35591738 | g.38027030.G>C (Pro.68.Ala) | 37,886,795 | *MIEN1* | | 3.828 | 2.00E-04 | 0.002 |
| | | 37,894,936 | | | 3.666 | 3.58E-04 | 0.004 |
| | | 37,896,708 | | | 3.573 | 4.96E-04 | 0.008 |
| | | 38,083,360 | | *ORMDL3* | 3.576 | 4.90E-04 | 0.011 |
| | | 38,083,475 | | *ORMDL3* | 3.777 | 2.40E-04 | 0.011 |
| | | 38,108,868 | | | -5.205 | 7.38E-07 | 0.014 |
| | | 38,109,131 | | | 4.062 | 8.35E-05 | 0.002 |
| rs76599245 | g.38027963.G>C | 37,894,916 | | | 3.440 | 7.83E-04 | 0.004 |
| | | 37,934,107 | | *IKZF3* | -4.490 | 1.55E-05 | 0.027 |
| | | 38,084,016 | *ORMDL3* | | 4.466 | 1.72E-05 | 0.011 |
| | | 38,109,131 | | | 4.741 | 5.50E-06 | 0.002 |
| rs150803188 | g.38032505.A>G | 37,856,732 | | *ERBB2* | -3.668 | 3.56E-04 | 0.009 |
| | | 37,886,749 | | *MIEN1* | 3.945 | 1.30E-04 | 0.002 |
| | | 37,896,063 | | | 3.476 | 6.92E-04 | 0.009 |
| | | 38,020,485 | *IKZF3* | | 3.962 | 1.22E-04 | 0.005 |
| | | 38,080,829 | | *ORMDL3* | 3.995 | 1.08E-04 | 0.014 |
| | | 38,096,333 | | | -3.664 | 3.60E-04 | 0.033 |
| | | 38,096,571 | | | -4.780 | 4.67E-06 | 0.033 |
| | | 38,096,578 | | | -5.642 | 1.00E-07 | 0.033 |
| | | 38,096,648 | | | -3.887 | 1.61E-04 | 0.033 |
| rs372063139 | g.38040332.T>C | 37,844,950 | *PGAP3* | *ERBB2* | -3.642 | 3.89E-04 | 7.82E-05 |
| | | 37,856,133 | | *ERBB2* | -3.562 | 5.15E-04 | 0.027 |
| | | 37,886,765 | | *MIEN1* | 4.206 | 4.81E-05 | 0.002 |
| | | 37,886,813 | *MIEN1* | | 5.914 | 2.78E-08 | 0.002 |
| | | 37,910,840 | | | 5.449 | 2.45E-07 | 0.035 |
| | | 37,910,911 | | | 4.735 | 5.65E-06 | 0.035 |
| | | 37,911,360 | | | 4.000 | 1.06E-04 | 0.035 |
| | | 38,024,244 | | | -4.970 | 2.07E-06 | 0.016 |
| | | 38,082,018 | | *ORMDL3* | -3.370 | 9.91E-04 | 6.40E-04 |
| | | 38,083,520 | | *ORMDL3* | 4.454 | 1.80E-05 | 0.011 |
| | | 38,108,734 | | | 3.680 | 3.41E-04 | 0.039 |
| rs79062097 | g.38043467.T>C | 37,886,390 | | *MIEN1* | -4.017 | 9.91E-05 | 0.002 |
| | | 37,911,186 | | | 3.578 | 4.87E-04 | 0.035 |

| | | | | | | | |
|---|---|---|---|---|---|---|---|
| | | 38,083,134 | | *ORMDL3* | 3.416 | 8.49E-04 | 0.011 |
| | | 38,085,031 | *ORMDL3* | | 4.013 | 1.01E-04 | 0.005 |
| rs146767035 | g.38045084.A>G | 37,913,515 | | | 4.384 | 2.38E-05 | 3.32E-05 |
| | | 38,020,616 | *IKZF3* | | 4.017 | 9.93E-05 | 0.017 |
| | | 38,080,829 | | *ORMDL3* | 3.639 | 3.94E-04 | 0.014 |
| | | 38,083,884 | *ORMDL3* | *ORMDL3* | 4.028 | 9.51E-05 | 0.011 |
| | | 38,108,509 | | | 3.792 | 2.27E-04 | 0.007 |
| | | 38,108,576 | | | 3.631 | 4.05E-04 | 0.007 |
| | | 38,109,541 | | | 3.944 | 1.30E-04 | 0.002 |
| rs138038828 | g.38047171.A>G | 37,894,014 | | | 3.503 | 6.30E-04 | 0.013 |
| | | 37,896,010 | | | 4.546 | 1.23E-05 | 0.027 |
| | | 37,911,274 | | | 6.369 | 3.01E-09 | 0.035 |
| | | 38,003,906 | | *IKZF3* | -3.846 | 1.87E-04 | 0.012 |
| | | 38,020,485 | *IKZF3* | | 3.664 | 3.60E-04 | 0.005 |
| | | 38,024,367 | | | -4.155 | 5.85E-05 | 0.017 |
| | | 38,083,590 | | *ORMDL3* | 10.853 | 6.22E-20 | 0.011 |
| | | 38,083,757 | | *ORMDL3* | 4.645 | 8.22E-06 | 0.011 |
| | | 38,083,894 | *ORMDL3* | | 3.498 | 6.42E-04 | 0.011 |
| rs143199092 | g.38050906.A>G | 37,886,795 | *MIEN1* | | 3.416 | 8.50E-04 | 0.002 |
| | | 38,020,460 | *IKZF3* | | 3.592 | 4.63E-04 | 0.005 |
| | | 38,024,409 | | | -4.146 | 6.06E-05 | 3.75E-04 |
| | | 38,077,826 | | *ORMDL3* | -3.402 | 8.89E-04 | 0.014 |
| | | 38,108,868 | | | -5.106 | 1.14E-06 | 0.014 |
| | | 38,109,131 | | | 4.071 | 8.08E-05 | 0.002 |
| rs141561785 | g.38058007.T>A | 37,910,819 | | | 5.067 | 1.36E-06 | 0.035 |
| | | 37,911,127 | | | 6.609 | 9.04E-10 | 0.035 |
| | | 37,911,390 | | | 4.749 | 5.32E-06 | 0.035 |
| | | 37,914,766 | | | -3.925 | 1.40E-04 | 3.32E-05 |
| | | 37,947,493 | | *IKZF3* | 6.569 | 1.11E-09 | 0.002 |
| | | 37,948,989 | | *IKZF3* | 3.427 | 8.18E-04 | 0.002 |
| | | 38,020,407 | | *IKZF3* | 3.992 | 1.09E-04 | 0.010 |
| | | 38,080,315 | | *ORMDL3* | 3.371 | 9.88E-04 | 0.014 |
| | | 38,083,227 | | *ORMDL3* | 3.651 | 3.78E-04 | 0.011 |
| | | 38,084,256 | *ORMDL3* | | -3.951 | 1.27E-04 | 0.005 |
| rs764747088 | g.38062863_38062864del | 37,856,228 | | *ERBB2* | -5.068 | 1.35E-06 | 0.015 |
| | | 37,857,668 | | *ERBB2* | 3.602 | 4.48E-04 | 0.003 |
| | | 37,910,382 | | | 3.981 | 1.14E-04 | 0.013 |
| | | 37,931,456 | | *IKZF3* | 3.588 | 4.71E-04 | 0.027 |
| | | 38,024,146 | | | 4.490 | 1.55E-05 | 0.004 |
| | | 38,077,098 | | | -4.936 | 2.40E-06 | 0.034 |

| | | | | | | | |
|---|---|---|---|---|---|---|---|
| | | 38,077,176 | | | -3.604 | 4.46E-04 | 0.034 |
| | | 38,077,277 | | | -3.792 | 2.28E-04 | 0.014 |
| | | 38,081,368 | | *ORMDL3* | -3.858 | 1.79E-04 | 2.01E-04 |
| | | 38,084,037 | *ORMDL3* | | 3.754 | 2.61E-04 | 0.005 |
| | | 38,084,450 | *ORMDL3* | | 3.869 | 1.72E-04 | 0.005 |
| | | 38,104,907 | | | -3.732 | 2.83E-04 | 0.007 |
| | | 38,108,622 | | | -3.621 | 4.19E-04 | 0.007 |
| rs1475291182 | g.38063809_38063811del | 37,856,732 | | *ERBB2* | -3.668 | 3.56E-04 | 0.009 |
| | | 37,886,749 | | *MIEN1* | 3.945 | 1.30E-04 | 0.002 |
| | | 37,896,063 | | | 3.476 | 6.92E-04 | 0.009 |
| | | 38,020,485 | *IKZF3* | | 3.962 | 1.22E-04 | 0.005 |
| | | 38,080,829 | | *ORMDL3* | 3.995 | 1.08E-04 | 0.014 |
| | | 38,096,333 | | | -3.664 | 3.60E-04 | 0.033 |
| | | 38,096,571 | | | -4.780 | 4.67E-06 | 0.033 |
| | | 38,096,578 | | | -5.642 | 1.00E-07 | 0.033 |
| | | 38,096,648 | | | -3.887 | 1.61E-04 | 0.033 |
| rs538657578 | g.38071007.G>T | 37,913,515 | | | 4.384 | 2.38E-05 | 3.32E-05 |
| | | 38,020,616 | *IKZF3* | | 4.017 | 9.93E-05 | 0.017 |
| | | 38,080,829 | | *ORMDL3* | 3.639 | 3.94E-04 | 0.014 |
| | | 38,083,884 | *ORMDL3* | *ORMDL3* | 4.028 | 9.51E-05 | 0.011 |
| | | 38,108,509 | | | 3.792 | 2.27E-04 | 0.007 |
| | | 38,108,576 | | | 3.631 | 4.05E-04 | 0.007 |
| | | 38,109,541 | | | 3.944 | 1.30E-04 | 0.002 |
| rs534518916 | g.38074323.T>C | 37,844,031 | *ERBB2* | *PGAP3* | 3.407 | 8.74E-04 | 7.82E-05 |
| | | 37,856,031 | | *ERBB2* | -3.684 | 3.35E-04 | 0.027 |
| | | 37,894,027 | | | 3.718 | 2.97E-04 | 0.013 |
| | | 37,895,026 | | | 3.829 | 1.99E-04 | 0.004 |
| | | 38,024,290 | | | 3.799 | 2.23E-04 | 0.019 |
| rs143706978 | g.38075501.C>T | 37,894,530 | | | -3.418 | 8.44E-04 | 0.013 |
| | | 37,895,757 | | | 3.875 | 1.68E-04 | 0.027 |
| | | 37,907,916 | | | -3.415 | 8.50E-04 | 0.008 |
| | | 38,081,995 | | *ORMDL3* | 3.707 | 3.09E-04 | 2.01E-04 |
| | | 38,082,022 | | *ORMDL3* | 3.542 | 5.52E-04 | 6.40E-04 |
| | | 38,108,523 | | | -3.382 | 9.50E-04 | 0.007 |
| rs145809834 | g.38077298.G>C | 37,844,452 | *PGAP3* | *ERBB2* | 4.836 | 3.67E-06 | 7.82E-05 |
| | | 37,886,979 | *MIEN1* | | 6.347 | 3.36E-09 | 0.002 |
| | | 38,083,590 | | *ORMDL3* | 3.815 | 2.09E-04 | 0.011 |
| rs3169572 | g.38077412.T>C | 37,856,732 | | *ERBB2* | -3.694 | 3.23E-04 | 0.009 |
| | | 37,879,682 | | *ERBB2* | 3.380 | 9.58E-04 | 0.002 |
| | | 37,896,063 | | | 3.678 | 3.42E-04 | 0.009 |

| | | | | | | | |
|---|---|---|---|---|---|---|---|
| | | 38,020,485 | *IKZF3* | | 3.611 | 4.35E-04 | 0.005 |
| | | 38,096,333 | | | -3.536 | 5.63E-04 | 0.033 |
| | | 38,096,578 | | | -4.659 | 7.76E-06 | 0.033 |
| rs7942 | g.38077464.C>T | 37,844,223 | *ERBB2* | *PGAP3* | 4.079 | 7.85E-05 | 7.82E-05 |
| | | 37,844,398 | *PGAP3* | *ERBB2* | 4.056 | 8.56E-05 | 7.82E-05 |
| | | 37,856,636 | | *ERBB2* | 3.382 | 9.52E-04 | 0.006 |
| | | 37,910,990 | | | 3.422 | 8.32E-04 | 0.035 |
| | | 38,077,464 | | *ORMDL3* | 14.617 | 3.19E-29 | 0.014 |
| | | 38,084,331 | *ORMDL3* | | 3.910 | 1.48E-04 | 0.005 |
| | | 38,084,450 | *ORMDL3* | | 3.855 | 1.81E-04 | 0.005 |
| | | 38,084,474 | *ORMDL3* | | 3.943 | 1.31E-04 | 0.005 |
| | | 38,100,110 | | | 3.877 | 1.67E-04 | 0.033 |
| | | 38,119,760 | | *GSDMA* | 4.044 | 8.94E-05 | 0.011 |
| rs111935829 | g.38081394.G>C | 37,886,390 | | *MIEN1* | -3.843 | 1.89E-04 | 0.002 |
| | | 37,911,186 | | | 3.707 | 3.09E-04 | 0.035 |
| | | 38,020,758 | *IKZF3* | | 3.577 | 4.90E-04 | 0.017 |
| | | 38,025,241 | | | -3.423 | 8.29E-04 | 0.046 |
| | | 38,083,134 | | *ORMDL3* | 3.911 | 1.47E-04 | 0.011 |
| rs539216949 | g.38081519.G>C | 37,827,454 | | *PGAP3* | -4.074 | 7.99E-05 | |
| | | 37,910,326 | | | 4.183 | 5.25E-05 | 0.013 |
| | | 37,910,824 | | | 5.551 | 1.53E-07 | 0.035 |
| | | 38,018,269 | | *IKZF3* | -4.192 | 5.08E-05 | 0.046 |
| | | 38,084,404 | *ORMDL3* | | -3.384 | 9.45E-04 | 0.005 |
| | | 38,109,098 | | | 3.636 | 3.98E-04 | 0.002 |
| rs534063293 | g.38082034.C>G | 37,856,228 | | *ERBB2* | -5.068 | 1.35E-06 | 0.015 |
| | | 37,857,668 | | *ERBB2* | 3.602 | 4.48E-04 | 0.003 |
| | | 37,910,382 | | | 3.981 | 1.14E-04 | 0.013 |
| | | 37,931,456 | | *IKZF3* | 3.588 | 4.71E-04 | 0.027 |
| | | 38,024,146 | | | 4.490 | 1.55E-05 | 0.004 |
| | | 38,077,098 | | | -4.936 | 2.40E-06 | 0.034 |
| | | 38,077,176 | | | -3.604 | 4.46E-04 | 0.034 |
| | | 38,077,277 | | | -3.792 | 2.28E-04 | 0.014 |
| | | 38,081,368 | | *ORMDL3* | -3.858 | 1.79E-04 | 2.01E-04 |
| | | 38,084,037 | *ORMDL3* | | 3.754 | 2.61E-04 | 0.005 |
| | | 38,084,450 | *ORMDL3* | | 3.869 | 1.72E-04 | 0.005 |
| | | 38,104,907 | | | -3.732 | 2.83E-04 | 0.007 |
| | | 38,108,622 | | | -3.621 | 4.19E-04 | 0.007 |
| rs17608925 | g.38082831.C>T | 38,020,208 | | *IKZF3* | 3.931 | 1.37E-04 | 0.046 |
| | | 38,077,098 | | | 4.998 | 1.83E-06 | 0.034 |
| | | 38,077,104 | | | 4.344 | 2.80E-05 | 0.034 |
| | | 38,077,107 | | | 4.705 | 6.40E-06 | 0.034 |

| | | | | | | |
|---|---|---|---|---|---|---|
| | | 38,077,191 | | 3.710 | 3.06E-04 | 0.034 |
| | | 38,077,703 | *ORMDL3* | 4.613 | 9.37E-06 | 0.014 |
| | | 38,081,368 | *ORMDL3* | 3.714 | 3.02E-04 | 2.01E-04 |
| | | 38,081,492 | *ORMDL3* | 4.502 | 1.48E-05 | 2.01E-04 |
| | | 38,081,812 | *ORMDL3* | 5.412 | 2.90E-07 | 2.01E-04 |
| | | 38,081,829 | *ORMDL3* | 5.895 | 3.04E-08 | 2.01E-04 |
| | | 38,081,995 | *ORMDL3* | 8.677 | 1.44E-14 | 2.01E-04 |
| | | 38,082,018 | *ORMDL3* | 6.324 | 3.78E-09 | 6.40E-04 |
| | | 38,082,022 | *ORMDL3* | 6.403 | 2.55E-09 | 6.40E-04 |
| | | 38,082,035 | *ORMDL3* | 5.423 | 2.76E-07 | 6.40E-04 |
| | | 38,082,073 | *ORMDL3* | 4.814 | 4.04E-06 | 6.40E-04 |
| | | 38,082,206 | *ORMDL3* | 5.231 | 6.55E-07 | 0.004 |
| | | 38,082,235 | *ORMDL3* | 5.631 | 1.06E-07 | 0.004 |
| | | 38,082,346 | *ORMDL3* | 6.028 | 1.60E-08 | 0.011 |
| | | 38,082,448 | *ORMDL3* | 7.671 | 3.54E-12 | 0.011 |
| | | 38,082,658 | *ORMDL3* | 5.217 | 7.00E-07 | 0.011 |
| | | 38,082,778 | *ORMDL3* | 4.642 | 8.34E-06 | 0.011 |
| | | 38,082,921 | *ORMDL3* | 6.140 | 9.32E-09 | 0.011 |
| rs538034067 | g.38089312_38089316del | 37,910,539 | | 3.747 | 2.68E-04 | 0.013 |
| | | 37,910,678 | | 3.680 | 3.41E-04 | 0.013 |
| | | 38,104,883 | | -3.610 | 4.36E-04 | 0.007 |
| rs73985231 | g.38090888.C>T | 37,886,461 | *MIEN1* | 3.753 | 2.62E-04 | 0.002 |
| | | 38,083,375 | *ORMDL3* | 4.737 | 5.60E-06 | 0.011 |
| rs79057213 | g.38091558.G>C | 37,886,461 | *MIEN1* | 3.753 | 2.62E-04 | 0.002 |
| | | 38,083,375 | *ORMDL3* | 4.737 | 5.60E-06 | 0.011 |
| rs1395015138 | g.38093219_38093220del | 37,886,461 | *MIEN1* | 3.753 | 2.62E-04 | 0.002 |
| | | 38,083,375 | *ORMDL3* | 4.737 | 5.60E-06 | 0.011 |
| rs112218945 | g.38093534.T>C | 37,886,390 | *MIEN1* | -3.759 | 2.57E-04 | 0.002 |
| | | 38,020,758 | *IKZF3* | 4.232 | 4.35E-05 | 0.017 |
| | | 38,077,104 | | -3.748 | 2.67E-04 | 0.034 |
| | | 38,083,308 | *ORMDL3* | 3.811 | 2.13E-04 | 0.011 |
| rs139807160 | g.38093764_38093765del | 37,886,461 | *MIEN1* | 3.812 | 2.12E-04 | 0.002 |
| | | 37,886,755 | *MIEN1* | 3.402 | 8.89E-04 | 0.002 |
| | | 37,895,953 | | -3.419 | 8.39E-04 | 0.027 |
| | | 37,910,423 | | 3.543 | 5.50E-04 | 0.013 |
| | | 37,911,096 | | 3.458 | 7.36E-04 | 0.035 |
| | | 37,912,185 | | -3.832 | 1.97E-04 | 0.033 |

| | | | | | | | |
|---|---|---|---|---|---|---|---|
| rs201516952 | g.38094390_38094390ins A | 37,886,461 | | *MIEN1* | 3.753 | 2.62E-04 | 0.002 |
| | | 38,083,375 | | *ORMDL3* | 4.737 | 5.60E-06 | 0.011 |
| rs117079275 | g.38097078.C>T | 37,886,461 | | *MIEN1* | 3.753 | 2.62E-04 | 0.002 |
| | | 38,083,375 | | *ORMDL3* | 4.737 | 5.60E-06 | 0.011 |
| rs150985855 | g.38097193.T>C | 37,886,461 | | *MIEN1* | 3.753 | 2.62E-04 | 0.002 |
| | | 38,083,375 | | *ORMDL3* | 4.737 | 5.60E-06 | 0.011 |
| rs75270719 | g.38097674.A>C | 37,828,560 | | *PGAP3* | -3.779 | 2.38E-04 | 0.002 |
| | | 37,910,347 | | | 3.805 | 2.17E-04 | 0.013 |
| | | 38,109,601 | | | 3.765 | 2.52E-04 | 0.002 |
| | | 38,125,830 | | *GSDMA* | 3.373 | 9.81E-04 | 0.006 |
| rs80320567 | g.38098042.T>C | 37,886,461 | | *MIEN1* | 3.753 | 2.62E-04 | 0.002 |
| | | 38,083,375 | | *ORMDL3* | 4.737 | 5.60E-06 | 0.011 |
| rs72832968 | g.38100673.G>C (Leu.172.Val) | 37,886,653 | | *MIEN1* | 3.452 | 7.51E-04 | 0.002 |
| | | 37,886,971 | *MIEN1* | | 3.710 | 3.06E-04 | 0.002 |
| | | 37,910,988 | | | 3.591 | 4.65E-04 | 0.035 |
| | | 38,083,560 | | *ORMDL3* | 3.372 | 9.82E-04 | 0.011 |
| rs117381273 | g.38101719.A>G | 37,886,461 | | *MIEN1* | 3.719 | 2.96E-04 | 0.002 |
| | | 37,886,838 | *MIEN1* | | 4.205 | 4.83E-05 | 0.002 |
| rs72832969 | g.38103440.A>G | 37,856,596 | | *ERBB2* | 3.504 | 6.29E-04 | 0.006 |
| | | 37,896,708 | | | 3.587 | 4.72E-04 | 0.008 |
| | | 38,077,176 | | | -3.962 | 1.22E-04 | 0.034 |
| rs112779146 | g.38103953.T>C | 37,886,390 | | *MIEN1* | -3.752 | 2.64E-04 | 0.002 |
| | | 38,020,758 | *IKZF3* | | 4.239 | 4.23E-05 | 0.017 |
| | | 38,077,104 | | | -3.756 | 2.59E-04 | 0.034 |
| | | 38,083,308 | | *ORMDL3* | 3.819 | 2.06E-04 | 0.011 |
| rs147041950 | g.38104802.A>G | 37,886,653 | | *MIEN1* | 3.452 | 7.51E-04 | 0.002 |
| | | 37,886,971 | *MIEN1* | | 3.710 | 3.06E-04 | 0.002 |
| | | 37,910,988 | | | 3.591 | 4.65E-04 | 0.035 |
| | | 38,083,560 | | *ORMDL3* | 3.372 | 9.82E-04 | 0.011 |
| rs73985233 | g.38105151.G>T | 37,886,461 | | *MIEN1* | 4.084 | 7.70E-05 | 0.002 |
| | | 37,886,838 | *MIEN1* | | 3.774 | 2.43E-04 | 0.002 |
| | | 38,077,621 | | *ORMDL3* | 3.383 | 9.48E-04 | 0.014 |
| | | 38,083,375 | | *ORMDL3* | 4.358 | 2.65E-05 | 0.011 |
| rs16965455 | g.38105754.C>T | 37,886,461 | | *MIEN1* | 4.084 | 7.70E-05 | 0.002 |
| | | 37,886,838 | *MIEN1* | | 3.774 | 2.43E-04 | 0.002 |
| | | 38,077,621 | | *ORMDL3* | 3.383 | 9.48E-04 | 0.014 |
| | | 38,083,375 | | *ORMDL3* | 4.358 | 2.65E-05 | 0.011 |
| rs147647857 | g.38106100_38106101del | 37,886,461 | | *MIEN1* | 4.084 | 7.70E-05 | 0.002 |

| | | | | | | | |
|---|---|---|---|---|---|---|---|
| | | 37,886,838 | *MIEN1* | | 3.774 | 2.43E-04 | 0.002 |
| | | 38,077,621 | | *ORMDL3* | 3.383 | 9.48E-04 | 0.014 |
| | | 38,083,375 | | *ORMDL3* | 4.358 | 2.65E-05 | 0.011 |
| rs73985234 | g.38106544.C>A | 37,886,461 | | *MIEN1* | 4.084 | 7.70E-05 | 0.002 |
| | | 37,886,838 | *MIEN1* | | 3.774 | 2.43E-04 | 0.002 |
| | | 38,077,621 | | *ORMDL3* | 3.383 | 9.48E-04 | 0.014 |
| | | 38,083,375 | | *ORMDL3* | 4.358 | 2.65E-05 | 0.011 |
| rs16965456 | g.38107095.G>T | 37,886,461 | | *MIEN1* | 4.084 | 7.70E-05 | 0.002 |
| | | 37,886,838 | *MIEN1* | | 3.774 | 2.43E-04 | 0.002 |
| | | 38,077,621 | | *ORMDL3* | 3.383 | 9.48E-04 | 0.014 |
| | | 38,083,375 | | *ORMDL3* | 4.358 | 2.65E-05 | 0.011 |
| rs138971063 | g.38107934.G>C | 37,886,461 | | *MIEN1* | 4.084 | 7.70E-05 | 0.002 |
| | | 37,886,838 | *MIEN1* | | 3.774 | 2.43E-04 | 0.002 |
| | | 38,077,621 | | *ORMDL3* | 3.383 | 9.48E-04 | 0.014 |
| | | 38,083,375 | | *ORMDL3* | 4.358 | 2.65E-05 | 0.011 |
| rs75481677 | g.38108230.T>A | 37,886,461 | | *MIEN1* | 4.084 | 7.70E-05 | 0.002 |
| | | 37,886,838 | *MIEN1* | | 3.774 | 2.43E-04 | 0.002 |
| | | 38,077,621 | | *ORMDL3* | 3.383 | 9.48E-04 | 0.014 |
| | | 38,083,375 | | *ORMDL3* | 4.358 | 2.65E-05 | 0.011 |
| rs142598968 | g.38109073.A>G | 37,844,253 | *ERBB2* | *PGAP3* | 4.095 | 7.38E-05 | 7.82E-05 |
| | | 37,844,398 | *PGAP3* | *ERBB2* | 3.787 | 2.32E-04 | 7.82E-05 |
| | | 37,844,714 | *PGAP3* | *ERBB2* | -5.587 | 1.30E-07 | 7.82E-05 |
| | | 37,886,692 | | *MIEN1* | 3.571 | 4.99E-04 | 0.002 |
| | | 37,910,269 | | | 4.677 | 7.20E-06 | 0.043 |
| | | 37,910,627 | | | 4.327 | 2.99E-05 | 0.013 |
| | | 37,910,811 | | | 3.646 | 3.84E-04 | 0.035 |
| | | 37,911,152 | | | 5.359 | 3.69E-07 | 0.035 |
| | | 38,024,702 | | | 3.674 | 3.48E-04 | 0.046 |
| | | 38,082,206 | | *ORMDL3* | -3.780 | 2.38E-04 | 0.004 |
| | | 38,083,257 | | *ORMDL3* | 4.093 | 7.42E-05 | 0.011 |
| | | 38,083,418 | | *ORMDL3* | 3.940 | 1.32E-04 | 0.011 |
| | | 38,083,560 | | *ORMDL3* | 5.249 | 6.06E-07 | 0.011 |
| | | 38,108,576 | | | 3.496 | 6.48E-04 | 0.007 |
| | | 38,108,703 | | | 3.399 | 8.99E-04 | 0.007 |
| rs116705564 | g.38111089.C>G | 37,886,461 | | *MIEN1* | 4.084 | 7.70E-05 | 0.002 |
| | | 37,886,838 | *MIEN1* | | 3.774 | 2.43E-04 | 0.002 |
| | | 38,077,621 | | *ORMDL3* | 3.383 | 9.48E-04 | 0.014 |
| | | 38,083,375 | | *ORMDL3* | 4.358 | 2.65E-05 | 0.011 |
| rs146082680 | g.38112684.A>G | 37,831,371 | | *PGAP3* | -3.533 | 5.70E-04 | 0.028 |
| | | 37,894,027 | | | 4.501 | 1.48E-05 | 0.013 |
| | | 37,895,775 | | | -3.386 | 9.39E-04 | 0.027 |

| | | | | | | | |
|---|---|---|---|---|---|---|---|
| rs146886475 | g.38112819.A>G | 37,886,461 | | *MIEN1* | 4.159 | 5.78E-05 | 0.002 |
| | | 37,886,838 | *MIEN1* | | 4.384 | 2.38E-05 | 0.002 |
| | | 37,912,185 | | | -3.918 | 1.43E-04 | 0.033 |
| rs79445771 | g.38116760.G>A | 37,886,461 | | *MIEN1* | 4.210 | 4.74E-05 | 0.002 |
| | | 37,886,838 | *MIEN1* | | 4.498 | 1.51E-05 | 0.002 |
| | | 37,912,185 | | | -3.859 | 1.79E-04 | 0.033 |
| rs77500917 | g.38116780.A>C | 37,886,461 | | *MIEN1* | 4.210 | 4.74E-05 | 0.002 |
| | | 37,886,838 | *MIEN1* | | 4.498 | 1.51E-05 | 0.002 |
| | | 37,912,185 | | | -3.859 | 1.79E-04 | 0.033 |
| rs74879852 | g.38116842.A>G | 37,886,461 | | *MIEN1* | 4.210 | 4.74E-05 | 0.002 |
| | | 37,886,838 | *MIEN1* | | 4.498 | 1.51E-05 | 0.002 |
| | | 37,912,185 | | | -3.859 | 1.79E-04 | 0.033 |
| rs75042551 | g.38118705.A>G | 37,886,921 | *MIEN1* | | 3.410 | 8.65E-04 | 0.002 |
| | | 37,894,014 | | | -3.458 | 7.36E-04 | 0.013 |
| | | 37,910,837 | | | 3.396 | 9.08E-04 | 0.035 |
| | | 38,083,354 | | *ORMDL3* | 3.496 | 6.47E-04 | 0.011 |
| rs373861118 | g.38118929_38118930ins CACACACACACA | 37,910,697 | | | 3.953 | 1.26E-04 | 0.035 |
| | | 38,077,051 | | | 3.453 | 7.49E-04 | 0.034 |
| | | 38,077,277 | | | 3.831 | 1.98E-04 | 0.014 |
| | | 38,077,464 | | *ORMDL3* | 3.902 | 1.53E-04 | 0.014 |
| | | 38,081,530 | | *ORMDL3* | 3.467 | 7.15E-04 | 2.01E-04 |
| | | 38,081,812 | | *ORMDL3* | 3.860 | 1.78E-04 | 2.01E-04 |
| | | 38,082,022 | | *ORMDL3* | 4.551 | 1.21E-05 | 6.40E-04 |
| | | 38,082,035 | | *ORMDL3* | 3.739 | 2.75E-04 | 6.40E-04 |
| | | 38,082,073 | | *ORMDL3* | 4.664 | 7.59E-06 | 6.40E-04 |
| | | 38,082,346 | | *ORMDL3* | 3.451 | 7.53E-04 | 0.011 |
| | | 38,082,658 | | *ORMDL3* | 4.367 | 2.55E-05 | 0.011 |
| | | 38,108,710 | | | -6.588 | 1.01E-09 | 0.039 |
| | | 38,119,071 | *GSDMA* | | 4.797 | 4.34E-06 | 0.009 |
| | | 38,119,198 | *GSDMA* | | 6.080 | 1.25E-08 | 0.009 |
| | | 38,119,207 | *GSDMA* | | 7.436 | 1.24E-11 | 0.009 |
| | | 38,119,649 | | *GSDMA* | 6.672 | 6.60E-10 | 0.002 |
| | | 38,119,760 | | *GSDMA* | 4.107 | 7.03E-05 | 0.011 |
| | | 38,120,234 | | *GSDMA* | 3.564 | 5.11E-04 | 0.011 |
| | | 38,128,699 | | *GSDMA* | -4.427 | 2.01E-05 | 0.006 |
| rs118073438 | g.38120999.T>G | 37,844,714 | *PGAP3* | *ERBB2* | -3.886 | 1.62E-04 | 7.82E-05 |
| | | 37,911,152 | | | 5.058 | 1.41E-06 | 0.035 |
| | | 37,911,186 | | | 3.745 | 2.70E-04 | 0.035 |
| | | 37,914,933 | | | 3.738 | 2.77E-04 | 3.32E-05 |
| | | 38,083,955 | *ORMDL3* | | 3.793 | 2.27E-04 | 0.011 |

| | | | | | | | |
|---|---|---|---|---|---|---|---|
| | | 38,115,531 | | | 3.775 | 2.43E-04 | 0.009 |
| rs80204597 | g.38121038.A>G | 37,886,461 | | *MIEN1* | 4.210 | 4.74E-05 | 0.002 |
| | | 37,886,838 | *MIEN1* | | 4.498 | 1.51E-05 | 0.002 |
| | | 37,912,185 | | | -3.859 | 1.79E-04 | 0.033 |
| rs141558041 | g.38121479_38121482del | 37,886,838 | *MIEN1* | | 5.673 | 8.70E-08 | 0.002 |
| | | 37,910,230 | | | 3.570 | 5.00E-04 | 0.043 |
| | | 38,024,634 | | | 3.375 | 9.74E-04 | 0.046 |
| | | 38,108,703 | | | 4.229 | 4.39E-05 | 0.007 |
| rs149553751 | g.38122099.T>C (Tyr.53.Tyr) | 37,858,063 | | *ERBB2* | 4.102 | 7.19E-05 | 0.040 |
| | | 37,893,994 | | | -3.747 | 2.68E-04 | 0.013 |
| | | 37,894,027 | | | -3.543 | 5.50E-04 | 0.013 |
| | | 37,910,184 | | | 3.652 | 3.75E-04 | 0.043 |
| | | 37,910,418 | | | 4.225 | 4.47E-05 | 0.013 |
| | | 37,910,840 | | | 3.669 | 3.54E-04 | 0.035 |
| | | 37,910,939 | | | 3.903 | 1.52E-04 | 0.035 |
| | | 37,911,189 | | | 5.398 | 3.09E-07 | 0.035 |
| | | 38,024,237 | | | 3.653 | 3.74E-04 | 0.016 |
| rs34076738 | g.38126222_38126222ins CAC | 38,077,464 | | *ORMDL3* | 3.910 | 1.48E-04 | 0.014 |
| | | 38,096,571 | | | -4.991 | 1.89E-06 | 0.033 |
| | | 38,096,578 | | | -4.278 | 3.63E-05 | 0.033 |
| rs55757004 | g.38126395.C>G | 38,119,198 | *GSDMA* | | -3.390 | 9.26E-04 | 0.009 |
| | | 38,119,207 | *GSDMA* | | -3.954 | 1.26E-04 | 0.009 |
| rs145291298 | g.38127058.C>T | 37,910,539 | | | 3.747 | 2.68E-04 | 0.013 |
| | | 37,910,678 | | | 3.680 | 3.41E-04 | 0.013 |
| | | 38,104,883 | | | -3.610 | 4.36E-04 | 0.007 |
| rs375455727 | g.38128786.A>G | 37,836,263 | | *PGAP3* | -3.933 | 1.36E-04 | 0.019 |
| | | 37,886,619 | | *MIEN1* | 3.604 | 4.44E-04 | 0.002 |
| | | 37,910,190 | | | 4.487 | 1.57E-05 | 0.043 |
| rs116955962 | g.38129201.C>T | 37,886,894 | *MIEN1* | | 5.588 | 1.29E-07 | 0.002 |
| | | 37,886,971 | *MIEN1* | | 4.584 | 1.06E-05 | 0.002 |
| | | 37,910,177 | | | 6.407 | 2.50E-09 | 0.043 |
| | | 37,910,283 | | | 3.823 | 2.03E-04 | 0.043 |
| | | 37,911,127 | | | 4.608 | 9.58E-06 | 0.035 |
| | | 37,930,190 | | *IKZF3* | -3.752 | 2.63E-04 | 0.027 |
| | | 38,083,250 | | *ORMDL3* | 3.733 | 2.82E-04 | 0.011 |
| rs149274005 | g.38129546.G>A | 37,844,040 | *ERBB2* | *PGAP3* | 4.951 | 2.25E-06 | 7.82E-05 |
| | | 37,844,950 | *PGAP3* | *ERBB2* | -3.560 | 5.18E-04 | 7.82E-05 |
| | | 37,857,767 | | *ERBB2* | 4.473 | 1.67E-05 | 0.003 |

| | | | | | | | |
|---|---|---|---|---|---|---|---|
| | | 37,886,204 | | *MIEN1* | 4.411 | 2.14E-05 | 0.002 |
| | | 37,886,610 | | *MIEN1* | 4.597 | 1.00E-05 | 0.002 |
| | | 37,893,689 | | | 3.690 | 3.28E-04 | 0.002 |
| | | 37,910,109 | | | 4.153 | 5.91E-05 | 0.043 |
| | | 38,081,368 | | *ORMDL3* | 4.138 | 6.26E-05 | 2.01E-04 |
| | | 38,081,995 | | *ORMDL3* | 3.875 | 1.68E-04 | 2.01E-04 |
| | | 38,082,022 | | *ORMDL3* | 3.541 | 5.54E-04 | 6.40E-04 |
| | | 38,082,035 | | *ORMDL3* | 3.478 | 6.87E-04 | 6.40E-04 |
| | | 38,082,346 | | *ORMDL3* | 4.591 | 1.03E-05 | 0.011 |
| | | 38,082,448 | | *ORMDL3* | 3.696 | 3.21E-04 | 0.011 |
| | | 38,109,639 | | | 3.747 | 2.68E-04 | 0.002 |
| rs184771266 | g.38130783.A>G | 37,835,700 | | *PGAP3* | 3.898 | 1.55E-04 | 0.020 |
| | | 37,853,032 | | *ERBB2* | 4.302 | 3.30E-05 | 0.027 |
| | | 37,856,873 | | *ERBB2* | -5.487 | 2.06E-07 | 0.009 |
| | | 37,863,630 | | *ERBB2* | 3.424 | 8.25E-04 | 0.002 |
| | | 37,894,352 | | | -3.692 | 3.26E-04 | 0.013 |
| | | 37,895,460 | | | 3.745 | 2.70E-04 | 0.006 |
| | | 37,907,982 | | | 3.436 | 7.94E-04 | 0.043 |
| | | 37,910,279 | | | 3.994 | 1.08E-04 | 0.043 |
| | | 37,910,523 | | | 4.314 | 3.14E-05 | 0.013 |
| | | 37,910,756 | | | 3.578 | 4.88E-04 | 0.035 |
| | | 37,911,394 | | | -3.447 | 7.64E-04 | 0.035 |
| | | 38,083,590 | | *ORMDL3* | 3.618 | 4.24E-04 | 0.011 |
| rs12946335 | g.38133764.T>C | 37,844,040 | *ERBB2* | *PGAP3* | 4.951 | 2.25E-06 | 7.82E-05 |
| | | 37,844,950 | *PGAP3* | *ERBB2* | -3.560 | 5.18E-04 | 7.82E-05 |
| | | 37,857,767 | | *ERBB2* | 4.473 | 1.67E-05 | 0.003 |
| | | 37,886,204 | | *MIEN1* | 4.411 | 2.14E-05 | 0.002 |
| | | 37,886,610 | | *MIEN1* | 4.597 | 1.00E-05 | 0.002 |
| | | 37,893,689 | | | 3.690 | 3.28E-04 | 0.002 |
| | | 37,910,109 | | | 4.153 | 5.91E-05 | 0.043 |
| | | 38,081,368 | | *ORMDL3* | 4.138 | 6.26E-05 | 2.01E-04 |
| | | 38,081,995 | | *ORMDL3* | 3.875 | 1.68E-04 | 2.01E-04 |
| | | 38,082,022 | | *ORMDL3* | 3.541 | 5.54E-04 | 6.40E-04 |
| | | 38,082,035 | | *ORMDL3* | 3.478 | 6.87E-04 | 6.40E-04 |
| | | 38,082,346 | | *ORMDL3* | 4.591 | 1.03E-05 | 0.011 |
| | | 38,082,448 | | *ORMDL3* | 3.696 | 3.21E-04 | 0.011 |
| | | 38,109,639 | | | 3.747 | 2.68E-04 | 0.002 |
| rs59132767 | g.38134300_38134301del | 37,910,697 | | | 3.513 | 6.10E-04 | 0.035 |
| | | 38,077,323 | | *ORMDL3* | 3.548 | 5.40E-04 | 0.014 |
| | | 38,077,464 | | *ORMDL3* | 3.885 | 1.62E-04 | 0.014 |

| | | | | | |
|---|---|---|---|---|---|
| 38,082,022 | | *ORMDL3* | 3.758 | 2.58E-04 | 6.40E-04 |
| 38,082,073 | | *ORMDL3* | 4.207 | 4.79E-05 | 6.40E-04 |
| 38,082,658 | | *ORMDL3* | 3.720 | 2.95E-04 | 0.011 |
| 38,108,710 | | | -5.079 | 1.29E-06 | 0.039 |
| 38,119,198 | *GSDMA* | | 3.881 | 1.65E-04 | 0.009 |
| 38,119,207 | *GSDMA* | | 4.375 | 2.47E-05 | 0.009 |
| 38,119,649 | | *GSDMA* | 4.305 | 3.26E-05 | 0.002 |
| 38,119,760 | | *GSDMA* | 3.726 | 2.89E-04 | 0.011 |
| 38,128,699 | | *GSDMA* | -5.161 | 8.97E-07 | 0.006 |

[a]Genes are indicated in the fifth and/or sixth columns if the CpG position is within 1,500 bp from the transcription start site (Gene is methylated [1.5 kb from TSS]) or in the gene body (Gene is methylated [gene body]). [b]P values for the association between the SNPs included in the mQTL and asthma. Ps are only shown if significant (p <0.001). [c]FDR values for the association between the methylation levels of CpG included in the mQTL and asthma.
Abbreviation: TSS = transcription start site.
