## Supplemental Table 11 for "*GSDMA* drives the most replicated association with asthma in naïve CD4^+^ T cells"

**Table S11:** Significant eQTLs in eosinophils

| Haplotype block | SNP id | HGVS names (hg19) | Gene | Statistic | P eQTL | P asthma[a] |
|---|---|---|---|---|---|---|
| First block | rs34503675 | g.37885640_37885640insG | *GSDMB* | -3.579 | 4.90E-04 | |
| | | | *MIEN1* | -3.782 | 2.39E-04 | |
| | rs66459548 | g.37892697.A>T | *GSDMB* | -3.755 | 2.63E-04 | |
| | | | *MIEN1* | -3.898 | 1.57E-04 | |
| | rs67597968 | g.37893458.G>A | *GSDMB* | -3.755 | 2.63E-04 | |
| | | | *MIEN1* | -3.898 | 1.57E-04 | |
| Second block | rs9896218 | g.37894463.C>A | *GSDMB* | -3.755 | 2.63E-04 | |
| | | | *MIEN1* | -3.898 | 1.57E-04 | |
| | rs71355409 | g.37895367_37895371del | *GSDMB* | -3.464 | 7.27E-04 | |
| | | | *MIEN1* | -4.001 | 1.07E-04 | |
| | rs36079893 | g.37898909.C>A (Pro.105.Pro) | *GSDMB* | -3.755 | 2.63E-04 | |
| | | | *MIEN1* | -3.898 | 1.57E-04 | |
| | rs34093201 | g.37898249.A>G | *GSDMB* | -4.120 | 6.80E-05 | |
| | | | *MIEN1* | -3.386 | 9.46E-04 | |
| | rs35585925 | g.37898210.C>T | *GSDMB* | -3.755 | 2.63E-04 | |
| | | | *MIEN1* | -3.898 | 1.57E-04 | |
| | rs59667301 | g.37895379.C>T | *GSDMB* | -3.755 | 2.63E-04 | |
| | | | *MIEN1* | -3.898 | 1.57E-04 | |
| Third block | rs9747973 | g.37905107.C>T | *GSDMB* | -3.498 | 6.47E-04 | |
| | | | *ORMDL3* | -5.042 | 1.57E-06 | |
| | rs11078921 | g.37908867.A>C | *ORMDL3* | -4.417 | 2.13E-05 | |
| | rs2941522 | g.37910368.G>A | *GSDMB* | -3.815 | 2.12E-04 | |
| | | | *ORMDL3* | -5.319 | 4.60E-07 | |
| | rs12946510 | g.37912377.T>C | *ORMDL3* | -4.679 | 7.32E-06 | |
| | rs35833706 | g.37916390_37916390insT | *GSDMB* | -4.279 | 3.68E-05 | |
| | | | *ORMDL3* | -6.116 | 1.12E-08 | |
| | rs907091 | g.37921742.C>T | *GSDMB* | -4.350 | 2.78E-05 | |
| | | | *ORMDL3* | -6.167 | 8.71E-09 | |
| | rs2952140 | g.37928059.G>A | *GSDMB* | -4.515 | 1.44E-05 | |
| | | | *ORMDL3* | -6.294 | 4.70E-09 | |
| | rs2313430 | g.37929816.A>G | *GSDMB* | -4.577 | 1.11E-05 | |
| | | | *ORMDL3* | -6.362 | 3.36E-09 | |
| | rs10445308 | g.37938047.T>C | *ORMDL3* | -4.891 | 2.99E-06 | |
| | rs12942330 | g.37939839.T>C | *ORMDL3* | -4.891 | 2.99E-06 | |
| | rs11658993 | g.37940808.T>C | *ORMDL3* | -4.891 | 2.99E-06 | |
| | rs2952144 | g.37960017.C>T | *GSDMB* | -4.253 | 4.08E-05 | |
| | | | *ORMDL3* | -6.075 | 1.36E-08 | |
| | rs4795395 | g.37962987.A>T | *ORMDL3* | -4.891 | 2.99E-06 | |
| | rs9909593 | g.37970149.G>A | *ORMDL3* | -4.891 | 2.99E-06 | |
| | rs907092 | g.37922259.A>G (Ser.438.Ser) | *ORMDL3* | -4.729 | 5.95E-06 | |
| Fourth block | rs113897057 | g.37975214_37975214insTTCTA | *GSDMB* | -4.587 | 1.07E-05 | |
| | | | *ORMDL3* | -6.438 | 2.31E-09 | |
| | rs12944882 | g.37983492.C>T | *GSDMB* | -5.121 | 1.11E-06 | |
| | | | *ORMDL3* | -7.078 | 9.04E-11 | |
| | rs3816470 | g.37985801.C>T | *GSDMB* | -5.121 | 1.11E-06 | |
| | | | *ORMDL3* | -7.078 | 9.04E-11 | |
| | rs62066988 | g.37992281.T>C | *ORMDL3* | -5.077 | 1.34E-06 | |

| | | | | |
|---|---|---|---|---|
| rs34233420 | g.38004929_38004933del | *GSDMB* | -3.546 | 5.49E-04 |
| | | *ORMDL3* | -5.219 | 7.20E-07 |
| rs9916765 | g.38005595.C>T | *GSDMB* | -4.481 | 1.65E-05 |
| | | *ORMDL3* | -6.225 | 6.59E-09 |
| rs35564481 | g.38020058_38020058insC | *GSDMB* | -4.380 | 2.47E-05 |
| | | *ORMDL3* | -5.891 | 3.27E-08 |
| rs1453559 | g.38020419.G>A | *GSDMB* | -4.481 | 1.65E-05 |
| | | *ORMDL3* | -6.225 | 6.59E-09 |
| rs4795397 | g.38023745.G>A | *ORMDL3* | -4.754 | 5.35E-06 |
| rs11557466 | g.38024626.T>C (Leu.7.Leu) | *ORMDL3* | -4.757 | 5.29E-06 |
| rs11078925 | g.38025208.C>T | *ORMDL3* | -4.757 | 5.29E-06 |
| rs111940435 | g.38025904.T>C | *GSDMB* | -3.382 | 9.59E-04 |
| | | *ORMDL3* | -3.485 | 6.77E-04 |
| rs34120102 | g.38026035.A>G | *ORMDL3* | -4.757 | 5.29E-06 |
| rs11655198 | g.38026169.T>C | *GSDMB* | -4.474 | 1.69E-05 |
| | | *ORMDL3* | -6.205 | 7.26E-09 |
| rs11650661 | g.38026286.T>A | *GSDMB* | -4.474 | 1.69E-05 |
| | | *ORMDL3* | -6.205 | 7.26E-09 |
| rs11655292 | g.38026361.G>C | *GSDMB* | -4.474 | 1.69E-05 |
| | | *ORMDL3* | -6.205 | 7.26E-09 |
| rs12709365 | g.38027400.G>A | *ORMDL3* | -4.757 | 5.29E-06 |
| rs13380815 | g.38027583.G>A | *ORMDL3* | -4.757 | 5.29E-06 |
| rs113606921 | g.38028254.C>T | *GSDMB* | -3.382 | 9.59E-04 |
| | | *ORMDL3* | -3.485 | 6.77E-04 |
| rs111845187 | g.38028497.G>C | *GSDMB* | -3.382 | 9.59E-04 |
| | | *ORMDL3* | -3.485 | 6.77E-04 |
| rs112561568 | g.38028857.A>G | *GSDMB* | -3.382 | 9.59E-04 |
| | | *ORMDL3* | -3.485 | 6.77E-04 |
| rs12936231 | g.38029120.G>C | *GSDMB* | -4.474 | 1.69E-05 |
| | | *ORMDL3* | -6.205 | 7.26E-09 |
| rs147134135 | g.38029867.C>A | *GSDMB* | -3.382 | 9.59E-04 |
| | | *ORMDL3* | -3.485 | 6.77E-04 |
| rs11870965 | g.38030205.A>T | *ORMDL3* | -4.794 | 4.53E-06 |
| rs9903250 | g.38031030.A>G | *GSDMB* | -4.474 | 1.69E-05 |
| | | *ORMDL3* | -6.205 | 7.26E-09 |
| rs112389508 | g.38031067.A>G | *GSDMB* | -3.382 | 9.59E-04 |
| | | *ORMDL3* | -3.485 | 6.77E-04 |
| rs111396994 | g.38031325.G>A | *GSDMB* | -3.382 | 9.59E-04 |
| | | *ORMDL3* | -3.485 | 6.77E-04 |
| rs10852935 | g.38031674.T>C (Cys.292.Cys) | *ORMDL3* | -4.757 | 5.29E-06 |
| rs10852936 | g.38031714.T>C | *ORMDL3* | -4.708 | 6.48E-06 |
| rs9891174 | g.38031802.A>T | *ORMDL3* | -4.757 | 5.29E-06 |
| rs781123329 | g.38031846_38031848del | *GSDMB* | -3.382 | 9.59E-04 |
| | | *ORMDL3* | -3.485 | 6.77E-04 |
| rs59716545 | g.38031857.G>T | *GSDMB* | -3.408 | 8.78E-04 |
| | | *ORMDL3* | -4.811 | 4.21E-06 |
| rs34189114 | g.38032460.T>C | *ORMDL3* | -4.757 | 5.29E-06 |
| rs35736272 | g.38032680.C>T | *ORMDL3* | -4.757 | 5.29E-06 |
| rs1054609 | g.38033277.C>A | *ORMDL3* | -4.757 | 5.29E-06 |
| rs117846478 | g.38033447.G>A | *GSDMB* | -3.382 | 9.59E-04 |
| | | *ORMDL3* | -3.485 | 6.77E-04 |

| | | | | |
|---|---|---|---|---|
| rs75972983 | g.38035050.T>C | *GSDMB* | -3.382 | 9.59E-04 |
| | | *ORMDL3* | -3.485 | 6.77E-04 |
| rs9907088 | g.38035116.A>G | *ORMDL3* | -4.757 | 5.29E-06 |
| rs36038753 | g.38035370.T>G | *ORMDL3* | -4.757 | 5.29E-06 |
| rs112750506 | g.38035410.G>A | *GSDMB* | -3.382 | 9.59E-04 |
| | | *ORMDL3* | -3.485 | 6.77E-04 |
| rs35569035 | g.38035624.T>C | *ORMDL3* | -4.445 | 1.90E-05 |
| rs9910826 | g.38035648.G>A | *ORMDL3* | -4.445 | 1.90E-05 |
| rs199882592 | g.38035706_38035707del | *GSDMB* | -3.382 | 9.59E-04 |
| | | *ORMDL3* | -3.485 | 6.77E-04 |
| rs34074973 | g.38035766_38035769del | *ORMDL3* | -4.757 | 5.29E-06 |
| rs9904624 | g.38036586.G>A | *ORMDL3* | -4.757 | 5.29E-06 |
| rs112750995 | g.38037559.T>A | *GSDMB* | -3.382 | 9.59E-04 |
| | | *ORMDL3* | -3.485 | 6.77E-04 |
| rs4795398 | g.38038179.T>C | *ORMDL3* | -4.757 | 5.29E-06 |
| rs148094956 | g.38039561_38039565del | *GSDMB* | -4.474 | 1.69E-05 |
| | | *ORMDL3* | -6.205 | 7.26E-09 |
| rs112569955 | g.38039806.G>A | *GSDMB* | -3.382 | 9.59E-04 |
| | | *ORMDL3* | -3.485 | 6.77E-04 |
| rs12232497 | g.38040119.C>T | *ORMDL3* | -4.757 | 5.29E-06 |
| rs112928243 | g.38040193.A>G | *GSDMB* | -3.382 | 9.59E-04 |
| | | *ORMDL3* | -3.485 | 6.77E-04 |
| rs12232498 | g.38040363.C>T | *ORMDL3* | -4.803 | 4.36E-06 |
| rs12941333 | g.38040534.T>C | *ORMDL3* | -4.757 | 5.29E-06 |
| rs113731681 | g.38040895.T>G | *GSDMB* | -3.382 | 9.59E-04 |
| | | *ORMDL3* | -3.485 | 6.77E-04 |
| rs112931462 | g.38041246.A>G | *GSDMB* | -3.382 | 9.59E-04 |
| | | *ORMDL3* | -3.485 | 6.77E-04 |
| rs12948927 | g.38041484.T>G | *GSDMB* | -3.382 | 9.59E-04 |
| | | *ORMDL3* | -3.485 | 6.77E-04 |
| rs9908132 | g.38042777.A>T | *GSDMB* | -4.474 | 1.69E-05 |
| | | *ORMDL3* | -6.205 | 7.26E-09 |
| rs9901146 | g.38043343.A>G | *GSDMB* | -4.474 | 1.69E-05 |
| | | *ORMDL3* | -6.205 | 7.26E-09 |
| rs12936409 | g.38043649.T>C | *ORMDL3* | -4.757 | 5.29E-06 |
| rs111317272 | g.38044525.G>C | *GSDMB* | -3.382 | 9.59E-04 |
| | | *ORMDL3* | -3.485 | 6.77E-04 |
| rs113486187 | g.38045043.A>G | *GSDMB* | -3.382 | 9.59E-04 |
| | | *ORMDL3* | -3.485 | 6.77E-04 |
| rs12103884 | g.38045725.T>C | *GSDMB* | -4.474 | 1.69E-05 |
| | | *ORMDL3* | -6.205 | 7.26E-09 |
| rs112049571 | g.38047658.A>G | *GSDMB* | -3.382 | 9.59E-04 |
| | | *ORMDL3* | -3.485 | 6.77E-04 |
| rs9906951 | g.38048244.C>T | *GSDMB* | -4.474 | 1.69E-05 |
| | | *ORMDL3* | -6.205 | 7.26E-09 |
| rs75760678 | g.38048721.T>C | *GSDMB* | -3.382 | 9.59E-04 |
| | | *ORMDL3* | -3.485 | 6.77E-04 |
| rs12950209 | g.38049102.C>T | *GSDMB* | -4.474 | 1.69E-05 |
| | | *ORMDL3* | -6.205 | 7.26E-09 |
| rs12950743 | g.38049233.C>T | *GSDMB* | -4.474 | 1.69E-05 |
| | | *ORMDL3* | -6.205 | 7.26E-09 |

| | | | | |
|---|---|---|---|---|
| rs7359623 | g.38049589.T>C | *GSDMB* | -4.474 | 1.69E-05 |
| | | *ORMDL3* | -6.205 | 7.26E-09 |
| rs17676923 | g.38050125.T>C | *GSDMB* | -3.387 | 9.43E-04 |
| | | *ORMDL3* | -3.504 | 6.36E-04 |
| rs35336365 | g.38050715.G>A | *GSDMB* | -3.382 | 9.59E-04 |
| | | *ORMDL3* | -3.485 | 6.77E-04 |
| rs12453507 | g.38053207.G>C | *GSDMB* | -3.968 | 1.21E-04 |
| | | *ORMDL3* | -5.597 | 1.30E-07 |
| rs2123685 | g.38053889.C>T | *GSDMB* | -3.382 | 9.59E-04 |
| | | *ORMDL3* | -3.485 | 6.77E-04 |
| rs11651596 | g.38056116.C>T | *ORMDL3* | -4.347 | 2.82E-05 |
| rs79457595 | g.38056247.T>C | *GSDMB* | -3.382 | 9.59E-04 |
| | | *ORMDL3* | -3.485 | 6.77E-04 |
| rs12949100 | g.38057189.A>G | *ORMDL3* | -4.348 | 2.81E-05 |
| rs8069176 | g.38057197.A>G | *ORMDL3* | -4.348 | 2.81E-05 |
| rs151192757 | g.38057566.T>C | *GSDMB* | -3.382 | 9.59E-04 |
| | | *ORMDL3* | -3.485 | 6.77E-04 |
| rs113348879 | g.38057605.T>A | *GSDMB* | -3.382 | 9.59E-04 |
| | | *ORMDL3* | -3.485 | 6.77E-04 |
| rs11657449 | g.38057841.C>G | *ORMDL3* | -4.506 | 1.49E-05 |
| rs4795399 | g.38061439.C>T | *ORMDL3* | -4.347 | 2.82E-05 |
| rs2305479 | g.38062217.A>G (Gly.304.Arg) | *GSDMB* | -3.918 | 1.46E-04 |
| | | *ORMDL3* | -5.729 | 7.04E-08 |
| rs35104165 | g.38062503.C>T (Asp.250.Gly) | *GSDMB* | -3.382 | 9.59E-04 |
| | | *ORMDL3* | -3.485 | 6.77E-04 |
| rs111655187 | g.38062738.A>C | *GSDMB* | -3.382 | 9.59E-04 |
| | | *ORMDL3* | -3.485 | 6.77E-04 |
| rs883770 | g.38063381.T>C | *GSDMB* | -3.918 | 1.46E-04 |
| | | *ORMDL3* | -5.729 | 7.04E-08 |
| rs62067034 | g.38063738.T>C | *GSDMB* | -3.918 | 1.46E-04 |
| | | *ORMDL3* | -5.729 | 7.04E-08 |
| rs12949547 | g.38063826.A>C | *GSDMB* | -3.382 | 9.59E-04 |
| | | *ORMDL3* | -3.485 | 6.77E-04 |
| rs113263629 | g.38063849.T>A | *GSDMB* | -3.382 | 9.59E-04 |
| | | *ORMDL3* | -3.485 | 6.77E-04 |
| rs36000226 | g.38063929.C>T | *GSDMB* | -3.918 | 1.46E-04 |
| | | *ORMDL3* | -5.729 | 7.04E-08 |
| rs36084703 | g.38063980_38063981del | *GSDMB* | -3.918 | 1.46E-04 |
| | | *ORMDL3* | -5.729 | 7.04E-08 |
| rs11078927 | g.38064405.T>C | *ORMDL3* | -4.347 | 2.82E-05 |
| rs11078928 | g.38064469.C>T (Leu.224.Leu) | *ORMDL3* | -4.347 | 2.82E-05 |
| rs111826178 | g.38064924.T>C | *GSDMB* | -3.382 | 9.59E-04 |
| | | *ORMDL3* | -3.485 | 6.77E-04 |
| rs113657930 | g.38065485.G>C | *GSDMB* | -3.382 | 9.59E-04 |
| | | *ORMDL3* | -3.485 | 6.77E-04 |
| rs1008723 | g.38066267.T>G | *GSDMB* | -3.918 | 1.46E-04 |
| | | *ORMDL3* | -5.729 | 7.04E-08 |
| rs56380902 | g.38066372.C>T | *GSDMB* | -3.918 | 1.46E-04 |
| | | *ORMDL3* | -5.729 | 7.04E-08 |
| rs4795400 | g.38067020.T>C | *ORMDL3* | -4.347 | 2.82E-05 |
| rs183631218 | g.38067532.C>T | *GSDMB* | -3.382 | 9.59E-04 |

| | | | | | | |
|---|---|---|---|---|---|---|
| | | | *ORMDL3* | -3.485 | 6.77E-04 | |
| | rs4795401 | g.38067533.G>A | *GSDMB* | -3.918 | 1.46E-04 | |
| | | | *ORMDL3* | -5.729 | 7.04E-08 | |
| | rs869402 | g.38068043.T>C | *ORMDL3* | -5.665 | 9.48E-08 | 5.35E-04 |
| | rs11078926 | g.38062976.A>G | *ORMDL3* | -4.347 | 2.82E-05 | |
| | rs112557679 | g.38044164.G>A | *GSDMB* | -3.382 | 9.59E-04 | |
| | | | *ORMDL3* | -3.485 | 6.77E-04 | |
| | rs11557467 | g.38028634.T>G (Ser.173.Ile) | *GSDMB* | -4.474 | 1.69E-05 | |
| | | | *ORMDL3* | -6.205 | 7.26E-09 | |
| | rs11658278 | g.38031164.C>T | *GSDMB* | -4.474 | 1.69E-05 | |
| | | | *ORMDL3* | -6.205 | 7.26E-09 | |
| | rs12150079 | g.38025417.A>G | *ORMDL3* | -4.858 | 3.46E-06 | |
| | rs12939457 | g.38032188.C>T | *ORMDL3* | -4.744 | 5.57E-06 | |
| | rs12939565 | g.38038389.T>A | *GSDMB* | -4.474 | 1.69E-05 | |
| | | | *ORMDL3* | -6.205 | 7.26E-09 | |
| | rs12939566 | g.38038390.T>A | *ORMDL3* | -4.757 | 5.29E-06 | |
| | rs189027840 | g.38052195.A>T | *GSDMB* | -3.405 | 8.89E-04 | |
| | | | *ORMDL3* | -3.497 | 6.51E-04 | |
| | rs200216139 | g.38032132_38032132insC | *GSDMB* | -4.253 | 4.08E-05 | |
| | | | *ORMDL3* | -6.010 | 1.86E-08 | |
| | rs201674348 | g.38062297_38062297insCTGCA CTCATAGCA | *GSDMB* | -3.382 | 9.59E-04 | |
| | | | *ORMDL3* | -3.485 | 6.77E-04 | |
| | rs202126107 | g.38006767_38006776del | *ORMDL3* | -4.754 | 5.35E-06 | |
| | rs2290400 | g.38066240.G>A | *GSDMB* | -3.918 | 1.46E-04 | |
| | | | *ORMDL3* | -5.729 | 7.04E-08 | |
| | rs2305480 | g.38062196.T>C (Pro.311.Ser) | *ORMDL3* | -4.347 | 2.82E-05 | |
| | rs2872507 | g.38040763.A>G | *ORMDL3* | -4.757 | 5.29E-06 | |
| | rs33938760 | | *GSDMB* | -4.474 | 1.69E-05 | |
| | | | *ORMDL3* | -6.205 | 7.26E-09 | |
| | rs34170568 | g.38055921_38055922ins | *ORMDL3* | -4.478 | 1.67E-05 | |
| | rs35196450 | g.38062942_38062942insC | *ORMDL3* | -4.347 | 2.82E-05 | |
| | rs35503505 | g.38057780_38057783del | *ORMDL3* | -4.703 | 6.62E-06 | |
| | rs36095411 | g.38031865.G>T | *GSDMB* | -3.831 | 2.01E-04 | 7.00E-04 |
| | | | *ORMDL3* | -5.707 | 7.79E-08 | 7.00E-04 |
| | rs367998020 | g.38032200_38032217del | *ORMDL3* | -4.744 | 5.57E-06 | |
| | rs56750287 | g.38062944.C>A | *ORMDL3* | -4.347 | 2.82E-05 | |
| | rs68122720 | | *ORMDL3* | -4.757 | 5.29E-06 | |
| | rs74978817 | g.38037989.G>A | *GSDMB* | -3.382 | 9.59E-04 | |
| | | | *ORMDL3* | -3.485 | 6.77E-04 | |
| | rs8067378 | g.38051348.G>A | *GSDMB* | -4.474 | 1.69E-05 | |
| | | | *ORMDL3* | -6.205 | 7.26E-09 | |
| | rs9303277 | g.37976469.T>C | *GSDMB* | -4.577 | 1.11E-05 | |
| | | | *ORMDL3* | -6.362 | 3.36E-09 | |
| | rs9905959 | g.38031138.G>A | *ORMDL3* | -4.757 | 5.29E-06 | |
| Fifth block | rs870829 | g.38068382.T>G | *ORMDL3* | -5.546 | 1.64E-07 | 7.23E-04 |
| | rs1011082 | g.38068514.A>G | *ORMDL3* | -5.672 | 9.17E-08 | 7.66E-04 |
| | rs921650 | g.38069076.C>T | *ORMDL3* | -5.665 | 9.48E-08 | 5.34E-04 |
| | rs921649 | g.38069274.G>A | *ORMDL3* | -5.665 | 9.48E-08 | 5.34E-04 |
| | rs5820308 | g.38069364_38069369del | *ORMDL3* | -4.303 | 3.35E-05 | 8.13E-04 |
| | rs6503524 | g.38069809.C>T | *ORMDL3* | -5.665 | 9.48E-08 | 5.34E-04 |

| | | | | | | |
|---|---|---|---|---|---|---|
| | rs7216558 | g.38070071.C>T | *ORMDL3* | -5.665 | 9.48E-08 | 5.34E-04 |
| | rs7221605 | g.38070789.C>T | *ORMDL3* | -5.681 | 8.78E-08 | |
| | rs1031458 | g.38072173.G>T | *ORMDL3* | -5.681 | 8.78E-08 | |
| | rs1031459 | g.38072245.C>G | *ORMDL3* | -4.278 | 3.70E-05 | |
| | rs1031460 | g.38072247.T>G | *ORMDL3* | -5.681 | 8.78E-08 | |
| | rs8065777 | g.38072402.C>T | *ORMDL3* | -5.681 | 8.78E-08 | |
| | rs112599791 | g.38072475.A>G | *GSDMB* | -3.382 | 9.59E-04 | |
| | | | *ORMDL3* | -3.485 | 6.77E-04 | |
| | rs2872516 | g.38072727.C>T | *ORMDL3* | -4.334 | 2.97E-05 | |
| | rs113894104 | g.38072955.T>C | *GSDMB* | -3.382 | 9.59E-04 | |
| | | | *ORMDL3* | -3.485 | 6.77E-04 | |
| | rs9303279 | g.38073968.G>C | *ORMDL3* | -4.303 | 3.35E-05 | 6.13E-04 |
| | rs9303280 | g.38074031.T>C | *ORMDL3* | -5.681 | 8.78E-08 | 8.49E-04 |
| | rs9303281 | g.38074046.G>A | *ORMDL3* | -5.665 | 9.48E-08 | 2.60E-04 |
| | rs7219923 | g.38074518.C>T | *ORMDL3* | -5.665 | 9.48E-08 | 2.75E-04 |
| | rs111379006 | g.38075100.A>G | *GSDMB* | -3.382 | 9.59E-04 | |
| | | | *ORMDL3* | -3.485 | 6.77E-04 | |
| | rs7224129 | g.38075426.G>A | *ORMDL3* | -5.789 | 5.32E-08 | 3.47E-04 |
| | rs8074437 | g.38076137.T>G | *ORMDL3* | -5.789 | 5.32E-08 | 3.47E-04 |
| | rs71971950 | g.38076198_38076201del | *ORMDL3* | -5.767 | 5.88E-08 | 3.77E-04 |
| | rs3859186 | g.38076329.T>C | *GSDMB* | -3.382 | 9.59E-04 | |
| | | | *ORMDL3* | -3.485 | 6.77E-04 | |
| | rs112260932 | g.38077185.C>T | *GSDMB* | -3.382 | 9.59E-04 | |
| | | | *ORMDL3* | -3.485 | 6.77E-04 | |
| | rs3169574 | g.38077485.T>C | *GSDMB* | -3.382 | 9.59E-04 | |
| | | | *ORMDL3* | -3.485 | 6.77E-04 | |
| | rs78199107 | g.38077750.C>T | *GSDMB* | -3.382 | 9.59E-04 | |
| | | | *ORMDL3* | -3.485 | 6.77E-04 | |
| | rs4065275 | g.38080865.A>G | *ORMDL3* | -5.525 | 1.80E-07 | 8.19E-04 |
| | rs8076131 | g.38080912.G>A | *ORMDL3* | -4.287 | 3.57E-05 | 5.13E-04 |
| | rs12603332 | g.38082807.T>C | *ORMDL3* | -5.525 | 1.80E-07 | 4.23E-04 |
| | rs145604770 | g.38086736.A>G | *GSDMB* | -3.382 | 9.59E-04 | |
| | | | *ORMDL3* | -3.485 | 6.77E-04 | |
| | rs143385463 | g.38071855_38071858del | *ORMDL3* | -5.681 | 8.78E-08 | |
| | rs150597688 | g.38071086_38071086insTT | *ORMDL3* | -5.681 | 8.78E-08 | |
| | rs4795402 | g.38085385.A>C | *ORMDL3* | -3.532 | 5.77E-04 | |
| | rs7216389 | g.38069949.C>T | *ORMDL3* | -5.665 | 9.48E-08 | 5.34E-04 |
| Sixth block | rs112191651 | g.38088150.T>C | *GSDMB* | -3.382 | 9.59E-04 | |
| | | | *ORMDL3* | -3.485 | 6.77E-04 | |
| | rs56199421 | g.38090808.C>T | *ORMDL3* | -3.880 | 1.68E-04 | |
| | rs76285844 | g.38092183.C>T | *GSDMB* | 4.228 | 4.50E-05 | |
| | | | *ORMDL3* | 4.661 | 7.89E-06 | |
| | rs12601749 | g.38093315.A>G | *ORMDL3* | -3.711 | 3.09E-04 | |
| | rs12603481 | g.38093339.A>G | *ORMDL3* | -4.630 | 8.96E-06 | |
| | rs6503525 | g.38095174.C>G | *ORMDL3* | -4.630 | 8.96E-06 | |
| | rs7216564 | g.38097172.A>T | *ORMDL3* | -4.630 | 8.96E-06 | |
| | rs144007425 | g.38098781_38098781insG | *ORMDL3* | -4.630 | 8.96E-06 | |
| | rs8065126 | g.38099035.T>C | *ORMDL3* | -3.596 | 4.63E-04 | |
| | rs4795406 | g.38100134.C>G | *ORMDL3* | -4.630 | 8.96E-06 | |
| | rs3848395 | g.38101468.T>C | *ORMDL3* | -3.596 | 4.63E-04 | |
| | rs4065985 | g.38101932.G>C | *ORMDL3* | -3.711 | 3.09E-04 | |

| | | | | |
|---|---|---|---|---|
| rs3893044 | g.38103016.A>G | *ORMDL3* | -3.711 | 3.09E-04 |
| rs62068170 | g.38103210.G>A | *ORMDL3* | -3.711 | 3.09E-04 |
| rs62068171 | g.38103242.G>A | *ORMDL3* | -3.711 | 3.09E-04 |
| rs8080734 | g.38103285.G>A | *ORMDL3* | -3.555 | 5.33E-04 |
| rs8071050 | g.38106599.A>G | *ORMDL3* | -4.372 | 2.55E-05 |
| rs4795408 | g.38107627.A>G | *ORMDL3* | -4.630 | 8.96E-06 |
| rs10589831 | g.38108023_38108025del | *ORMDL3* | -3.441 | 7.86E-04 |
| rs9889716 | g.38108298.A>G | *ORMDL3* | -3.501 | 6.42E-04 |
| rs9895948 | g.38108363.T>C | *ORMDL3* | -3.441 | 7.86E-04 |
| rs7223318 | g.38108553.G>C | *ORMDL3* | -3.441 | 7.86E-04 |
| rs7209742 | g.38108708.A>G | *ORMDL3* | -3.441 | 7.86E-04 |
| rs28618095 | g.38109075.T>C | *ORMDL3* | -3.402 | 8.96E-04 |
| rs56301252 | g.38109155.T>G | *ORMDL3* | -3.441 | 7.86E-04 |
| rs60667221 | g.38110390.T>A | *ORMDL3* | -3.405 | 8.89E-04 |
| rs17609240 | g.38110689.T>G | *ORMDL3* | -3.405 | 8.89E-04 |
| rs8076474 | g.38111234.G>C | *ORMDL3* | -3.405 | 8.89E-04 |
| rs1007655 | g.38111419.G>A | *ORMDL3* | -3.405 | 8.89E-04 |
| rs1563103 | g.38111740.G>A | *ORMDL3* | -3.405 | 8.89E-04 |
| rs2313640 | g.38111845.G>A | *ORMDL3* | -3.405 | 8.89E-04 |
| rs8068522 | g.38112076.T>C | *ORMDL3* | -3.405 | 8.89E-04 |
| rs8081437 | g.38112114.G>A | *ORMDL3* | -3.422 | 8.39E-04 |
| rs8081462 | g.38112190.C>G | *ORMDL3* | -4.630 | 8.96E-06 |
| rs62068174 | g.38112255.G>C | *ORMDL3* | -3.405 | 8.89E-04 |
| rs62068175 | g.38112300.A>C | *ORMDL3* | -3.405 | 8.89E-04 |
| rs62068176 | g.38112438.A>G | *ORMDL3* | -3.484 | 6.80E-04 |
| rs62068177 | g.38112601.G>A | *ORMDL3* | -3.564 | 5.16E-04 |
| rs62068178 | g.38112608.C>T | *ORMDL3* | -3.564 | 5.16E-04 |
| rs62068179 | g.38112617.G>A | *ORMDL3* | -3.564 | 5.16E-04 |
| rs67480438 | g.38112774.A>G | *ORMDL3* | -3.394 | 9.21E-04 |
| rs62068182 | g.38112864.G>A | *ORMDL3* | -3.394 | 9.21E-04 |
| rs62068184 | g.38113054.C>T | *ORMDL3* | -3.405 | 8.89E-04 |
| rs201351591 | g.38113956_38113956insG | *GSDMB* | 3.941 | 1.34E-04 |
| | | *ORMDL3* | 4.108 | 7.14E-05 |
| rs7218742 | g.38114361.A>G | *ORMDL3* | -3.405 | 8.89E-04 |
| rs7218321 | g.38114469.C>T | *ORMDL3* | -3.405 | 8.89E-04 |
| rs6503526 | g.38114598.T>C | *ORMDL3* | -4.586 | 1.07E-05 |
| rs6503527 | g.38114719.G>A | *ORMDL3* | -3.405 | 8.89E-04 |
| rs3931960 | g.38114977.G>C | *ORMDL3* | -4.372 | 2.55E-05 |
| rs7223717 | g.38115333.C>A | *ORMDL3* | -4.372 | 2.55E-05 |
| rs10693935 | g.38115429_38115429insTG | *ORMDL3* | -4.372 | 2.55E-05 |
| rs564455927 | g.38118235_38118235insA | *ORMDL3* | -4.521 | 1.40E-05 |
| rs55739615 | g.38119638.C>T | *ORMDL3* | -3.614 | 4.34E-04 |
| rs12451084 | g.38119757.T>G | *ORMDL3* | -4.630 | 8.96E-06 |
| rs12451100 | g.38119831.T>G | *ORMDL3* | -4.630 | 8.96E-06 |
| rs4795409 | g.38120736.T>C | *ORMDL3* | -4.630 | 8.96E-06 |
| rs8069202 | g.38122200.A>G | *ORMDL3* | -4.578 | 1.11E-05 |
| rs1007654 | g.38111354.A>G | *ORMDL3* | -3.405 | 8.89E-04 |
| rs35123741 | g.38092930.G>A | *ORMDL3* | -4.630 | 8.96E-06 |
| rs3894194 | g.38121993.T>C (Arg.18.Gln) | *ORMDL3* | -4.578 | 1.11E-05 |
| rs3902025 | g.38119254.C>A | *ORMDL3* | -3.614 | 4.34E-04 |
| rs4065986 | g.38102641.A>G | *ORMDL3* | -4.630 | 8.96E-06 |

| | | | | | |
|---|---|---|---|---|---|
| | rs4134417 | g.38113278.G>A | *ORMDL3* | -3.382 | 9.60E-04 |
| | rs4134498 | g.38113282.G>A | *ORMDL3* | -3.382 | 9.60E-04 |
| | rs4134499 | g.38113274.G>A | *ORMDL3* | -3.382 | 9.60E-04 |
| | rs4134500 | g.38113270.G>A | *ORMDL3* | -3.382 | 9.60E-04 |
| | rs4134501 | g.38113290.G>A | *ORMDL3* | -3.382 | 9.60E-04 |
| | rs4134502 | g.38113286.G>A | *ORMDL3* | -3.382 | 9.60E-04 |
| | rs4795405 | g.38088417.T>C | *ORMDL3* | -4.117 | 6.90E-05 |
| | rs55927420 | g.38109251.G>T | *ORMDL3* | -3.441 | 7.86E-04 |
| | rs56340811 | g.38115430.A>C | *ORMDL3* | -4.372 | 2.55E-05 |
| | rs56410675 | g.38109254.A>G | *ORMDL3* | -3.441 | 7.86E-04 |
| | rs62068180 | g.38112825.T>C | *ORMDL3* | -3.394 | 9.21E-04 |
| | rs62068181 | g.38112832.T>C | *ORMDL3* | -3.394 | 9.21E-04 |
| | rs62068183 | g.38112918.C>T | *ORMDL3* | -3.405 | 8.89E-04 |
| | rs7219080 | g.38114516.A>C | *ORMDL3* | -3.405 | 8.89E-04 |
| | rs8079416 | g.38092713.C>T | *ORMDL3* | -4.630 | 8.96E-06 |
| Seventh block | rs150060808 | g.38126911_38126919del | *ORMDL3* | -4.079 | 7.95E-05 |
| NA | rs79524843 | g.37832386.T>G | *MIEN1* | -4.005 | 1.05E-04 |
| | rs76096054 | g.37832488.A>C | *MIEN1* | -4.005 | 1.05E-04 |
| | rs113831251 | g.37838200.A>G | *MIEN1* | -4.005 | 1.05E-04 |
| | rs75849983 | g.37843762.G>T | *MIEN1* | -4.005 | 1.05E-04 |
| | rs572656035 | g.37848683_37848684del | *ORMDL3* | -3.618 | 4.29E-04 |
| | rs551511055 | g.37852066_37852073del | *ORMDL3* | -3.847 | 1.89E-04 |
| | rs111700892 | g.37853908.T>C | *MIEN1* | -4.005 | 1.05E-04 |
| | rs79747793 | g.37855331.A>C | *MIEN1* | -4.005 | 1.05E-04 |
| | rs12947247 | g.37855541.C>T | *MIEN1* | -4.005 | 1.05E-04 |
| | rs34284966 | g.37859083.C>G | *MIEN1* | -4.005 | 1.05E-04 |
| | rs34006795 | g.37860837.T>C | *MIEN1* | -4.005 | 1.05E-04 |
| | rs148040998 | g.37863606_37863606insTCGGCCA | *MIEN1* | -4.005 | 1.05E-04 |
| | rs4252624 | g.37867647.T>G | *MIEN1* | -4.005 | 1.05E-04 |
| | rs1181234061 | g.37874043_37874049del | *MIEN1* | -4.678 | 7.35E-06 |
| | rs4252639 | g.37876179.G>C | *MIEN1* | -4.005 | 1.05E-04 |
| | rs111947409 | g.37877716.A>G | *MIEN1* | -4.005 | 1.05E-04 |
| | rs115334808 | g.37878113.T>C | *MIEN1* | -4.005 | 1.05E-04 |
| | rs4252665 | g.37885383.T>C | *ORMDL3* | -3.667 | 3.60E-04 |
| | rs139233737 | g.37887636_37887636insT | *ORMDL3* | -3.667 | 3.60E-04 |
| | rs112535478 | g.37888034.G>A | *ORMDL3* | -3.667 | 3.60E-04 |
| | rs144954923 | g.37889025_37889026del | *MIEN1* | -4.005 | 1.05E-04 |
| | rs111424867 | g.37889280.T>C | *MIEN1* | -4.005 | 1.05E-04 |
| | rs78841518 | g.37891229.G>A | *MIEN1* | -4.005 | 1.05E-04 |
| | rs77755027 | g.37891235.A>G | *MIEN1* | -4.005 | 1.05E-04 |
| | rs112585477 | g.37891654.A>G | *GSDMB* | -3.382 | 9.59E-04 |
| | | | *ORMDL3* | -3.485 | 6.77E-04 |
| | rs112327574 | g.37893663.G>C | *GSDMB* | -3.382 | 9.59E-04 |
| | | | *ORMDL3* | -3.485 | 6.77E-04 |
| | rs9896940 | g.37895975.G>A | *GSDMB* | -3.382 | 9.59E-04 |
| | | | *ORMDL3* | -3.485 | 6.77E-04 |
| | rs113903339 | g.37898047.T>C | *GSDMB* | -3.382 | 9.59E-04 |
| | | | *ORMDL3* | -3.485 | 6.77E-04 |
| | rs76121559 | g.37899092.A>G | *MIEN1* | -4.672 | 7.53E-06 |
| | rs796319243 | g.37903880_37903881del | *GSDMB* | -3.382 | 9.59E-04 |

| | | | | |
|---|---|---|---|---|
| | | *ORMDL3* | -3.485 | 6.77E-04 |
| rs9910678 | g.37904211.C>T | *GSDMB* | -3.382 | 9.59E-04 |
| | | *ORMDL3* | -3.485 | 6.77E-04 |
| rs75933089 | g.37904449.T>G | *GSDMB* | -3.382 | 9.59E-04 |
| | | *ORMDL3* | -3.485 | 6.77E-04 |
| rs78365632 | g.37906916.A>G | *GSDMB* | -3.382 | 9.59E-04 |
| | | *ORMDL3* | -3.485 | 6.77E-04 |
| rs9891131 | g.37911521.G>A | *GSDMB* | -3.382 | 9.59E-04 |
| | | *ORMDL3* | -3.485 | 6.77E-04 |
| rs77013147 | g.37912268.C>T | *GSDMB* | -3.382 | 9.59E-04 |
| | | *ORMDL3* | -3.485 | 6.77E-04 |
| rs573534572 | g.37917826.T>C | *GSDMB* | -3.382 | 9.59E-04 |
| | | *ORMDL3* | -3.485 | 6.77E-04 |
| rs67571561 | g.37920847.C>T | *GSDMB* | -3.382 | 9.59E-04 |
| | | *ORMDL3* | -3.485 | 6.77E-04 |
| rs2941509 | g.37921194.A>G | *GSDMB* | -3.382 | 9.59E-04 |
| | | *ORMDL3* | -3.485 | 6.77E-04 |
| rs112876941 | g.37922804.T>A | *GSDMB* | -3.382 | 9.59E-04 |
| | | *ORMDL3* | -3.485 | 6.77E-04 |
| rs112771360 | g.37923771.G>A | *GSDMB* | -3.382 | 9.59E-04 |
| | | *ORMDL3* | -3.485 | 6.77E-04 |
| rs34758895 | g.37925468.T>C | *GSDMB* | -3.382 | 9.59E-04 |
| | | *ORMDL3* | -3.485 | 6.77E-04 |
| rs9894370 | g.37926004.C>G | *GSDMB* | -3.382 | 9.59E-04 |
| | | *ORMDL3* | -3.485 | 6.77E-04 |
| rs8072612 | g.37927120.G>A | *GSDMB* | -3.382 | 9.59E-04 |
| | | *ORMDL3* | -3.485 | 6.77E-04 |
| rs71369788 | g.37927145.A>G | *GSDMB* | -3.382 | 9.59E-04 |
| | | *ORMDL3* | -3.485 | 6.77E-04 |
| rs9652838 | g.37929248.C>T | *GSDMB* | -3.382 | 9.59E-04 |
| | | *ORMDL3* | -3.485 | 6.77E-04 |
| rs9652840 | g.37929428.T>A | *GSDMB* | -3.382 | 9.59E-04 |
| | | *ORMDL3* | -3.485 | 6.77E-04 |
| rs7217133 | g.37930262.A>T | *GSDMB* | -3.382 | 9.59E-04 |
| | | *ORMDL3* | -3.485 | 6.77E-04 |
| rs113730542 | g.37930421.G>C | *GSDMB* | -3.382 | 9.59E-04 |
| | | *ORMDL3* | -3.485 | 6.77E-04 |
| rs2313429 | g.37930659.C>T | *GSDMB* | -3.382 | 9.59E-04 |
| | | *ORMDL3* | -3.485 | 6.77E-04 |
| rs7503018 | g.37931220.G>A | *GSDMB* | -3.382 | 9.59E-04 |
| | | *ORMDL3* | -3.485 | 6.77E-04 |
| rs9913596 | g.37932063.A>G | *GSDMB* | -3.382 | 9.59E-04 |
| | | *ORMDL3* | -3.485 | 6.77E-04 |
| rs9894898 | g.37932221.C>T | *GSDMB* | -3.382 | 9.59E-04 |
| | | *ORMDL3* | -3.485 | 6.77E-04 |
| rs9901483 | g.37932774.A>T | *GSDMB* | -3.382 | 9.59E-04 |
| | | *ORMDL3* | -3.485 | 6.77E-04 |
| rs113370572 | g.37933468.C>T | *GSDMB* | -3.382 | 9.59E-04 |
| | | *ORMDL3* | -3.485 | 6.77E-04 |
| rs12709364 | g.37933823.G>A | *GSDMB* | -3.382 | 9.59E-04 |
| | | *ORMDL3* | -3.485 | 6.77E-04 |

| | | | | |
|---|---|---|---|---|
| rs7224641 | g.37934911.C>T | *GSDMB* | -3.382 | 9.59E-04 |
| | | *ORMDL3* | -3.485 | 6.77E-04 |
| rs13380871 | g.37936249.C>T | *GSDMB* | -3.382 | 9.59E-04 |
| | | *ORMDL3* | -3.485 | 6.77E-04 |
| rs35506518 | g.37938094.C>T | *GSDMB* | -3.382 | 9.59E-04 |
| | | *ORMDL3* | -3.485 | 6.77E-04 |
| rs67605703 | g.37938497.C>T | *GSDMB* | -3.382 | 9.59E-04 |
| | | *ORMDL3* | -3.485 | 6.77E-04 |
| rs34016964 | g.37938977.T>G | *GSDMB* | -3.382 | 9.59E-04 |
| | | *ORMDL3* | -3.485 | 6.77E-04 |
| rs9909365 | g.37939959.G>A | *GSDMB* | -3.382 | 9.59E-04 |
| | | *ORMDL3* | -3.485 | 6.77E-04 |
| rs113812449 | g.37940168.C>T | *GSDMB* | -3.382 | 9.59E-04 |
| | | *ORMDL3* | -3.485 | 6.77E-04 |
| rs1510475 | g.37941380.G>T | *GSDMB* | -3.382 | 9.59E-04 |
| | | *ORMDL3* | -3.485 | 6.77E-04 |
| rs112345383 | g.37942018.T>C | *GSDMB* | -3.382 | 9.59E-04 |
| | | *ORMDL3* | -3.485 | 6.77E-04 |
| rs34599546 | g.37942972.T>C | *GSDMB* | -3.382 | 9.59E-04 |
| | | *ORMDL3* | -3.485 | 6.77E-04 |
| rs111862642 | g.37942984.G>C | *GSDMB* | -3.382 | 9.59E-04 |
| | | *ORMDL3* | -3.485 | 6.77E-04 |
| rs9911634 | g.37943731.A>C | *GSDMB* | -3.382 | 9.59E-04 |
| | | *ORMDL3* | -3.485 | 6.77E-04 |
| rs9911669 | g.37943767.G>C | *GSDMB* | -3.382 | 9.59E-04 |
| | | *ORMDL3* | -3.485 | 6.77E-04 |
| rs34291217 | g.37944411.A>C | *GSDMB* | -3.382 | 9.59E-04 |
| | | *ORMDL3* | -3.485 | 6.77E-04 |
| rs35088469 | g.37944482.T>C | *GSDMB* | -3.382 | 9.59E-04 |
| | | *ORMDL3* | -3.485 | 6.77E-04 |
| rs112301322 | g.37944519.G>C (Gly.234.Ala) | *GSDMB* | -3.382 | 9.59E-04 |
| | | *ORMDL3* | -3.485 | 6.77E-04 |
| rs112771646 | g.37945709.C>A | *GSDMB* | -3.382 | 9.59E-04 |
| | | *ORMDL3* | -3.485 | 6.77E-04 |
| rs9913795 | g.37945841.T>C | *GSDMB* | -3.382 | 9.59E-04 |
| | | *ORMDL3* | -3.485 | 6.77E-04 |
| rs9914516 | g.37946125.T>C | *GSDMB* | -3.382 | 9.59E-04 |
| | | *ORMDL3* | -3.485 | 6.77E-04 |
| rs9913044 | g.37946228.A>G | *GSDMB* | -3.382 | 9.59E-04 |
| | | *ORMDL3* | -3.485 | 6.77E-04 |
| rs71355416 | g.37946601_37946602del | *GSDMB* | -3.382 | 9.59E-04 |
| | | *ORMDL3* | -3.485 | 6.77E-04 |
| rs35352075 | g.37949791.C>T | *GSDMB* | -3.382 | 9.59E-04 |
| | | *ORMDL3* | -3.485 | 6.77E-04 |
| rs35105110 | g.37950422.A>G | *GSDMB* | -3.382 | 9.59E-04 |
| | | *ORMDL3* | -3.485 | 6.77E-04 |
| rs35938199 | g.37950813.T>C | *GSDMB* | -3.382 | 9.59E-04 |
| | | *ORMDL3* | -3.485 | 6.77E-04 |
| rs12938749 | g.37951848.T>C | *GSDMB* | -3.382 | 9.59E-04 |
| | | *ORMDL3* | -3.485 | 6.77E-04 |
| rs56928975 | g.37952032.G>A | *GSDMB* | -3.382 | 9.59E-04 |

| | | | | |
|---|---|---|---|---|
| | | *ORMDL3* | -3.485 | 6.77E-04 |
| rs113159227 | g.37952092.A>G | *GSDMB* | -3.382 | 9.59E-04 |
| | | *ORMDL3* | -3.485 | 6.77E-04 |
| rs73302152 | g.37952351.C>G | *GSDMB* | -3.382 | 9.59E-04 |
| | | *ORMDL3* | -3.485 | 6.77E-04 |
| rs75148376 | g.37952509.T>C | *ORMDL3* | -3.452 | 7.58E-04 |
| rs113369293 | g.37952655.T>C | *GSDMB* | -3.382 | 9.59E-04 |
| | | *ORMDL3* | -3.485 | 6.77E-04 |
| rs13313561 | g.37954451.C>G | *GSDMB* | -3.382 | 9.59E-04 |
| | | *ORMDL3* | -3.485 | 6.77E-04 |
| rs34137841 | g.37954533_37954534del | *GSDMB* | -3.382 | 9.59E-04 |
| | | *ORMDL3* | -3.485 | 6.77E-04 |
| rs34516015 | g.37954798.A>T | *GSDMB* | -3.382 | 9.59E-04 |
| | | *ORMDL3* | -3.485 | 6.77E-04 |
| rs34876670 | g.37954800.C>T | *GSDMB* | -3.382 | 9.59E-04 |
| | | *ORMDL3* | -3.485 | 6.77E-04 |
| rs9907988 | g.37954883.C>T | *GSDMB* | -3.382 | 9.59E-04 |
| | | *ORMDL3* | -3.485 | 6.77E-04 |
| rs13313564 | g.37955555.A>G | *GSDMB* | -3.382 | 9.59E-04 |
| | | *ORMDL3* | -3.485 | 6.77E-04 |
| rs13313573 | g.37955818.T>G | *GSDMB* | -3.382 | 9.59E-04 |
| | | *ORMDL3* | -3.485 | 6.77E-04 |
| rs12937330 | g.37957317.A>C | *GSDMB* | -3.382 | 9.59E-04 |
| | | *ORMDL3* | -3.485 | 6.77E-04 |
| rs34988504 | g.37957632.T>C | *GSDMB* | -3.382 | 9.59E-04 |
| | | *ORMDL3* | -3.485 | 6.77E-04 |
| rs36097841 | g.37958113.A>G | *GSDMB* | -3.382 | 9.59E-04 |
| | | *ORMDL3* | -3.485 | 6.77E-04 |
| rs113064843 | g.37960422.C>T | *GSDMB* | -3.382 | 9.59E-04 |
| | | *ORMDL3* | -3.485 | 6.77E-04 |
| rs9894680 | g.37962237.C>T | *GSDMB* | -3.382 | 9.59E-04 |
| | | *ORMDL3* | -3.485 | 6.77E-04 |
| rs9914302 | g.37962408.A>G | *GSDMB* | -3.382 | 9.59E-04 |
| | | *ORMDL3* | -3.485 | 6.77E-04 |
| rs79989390 | g.37962845.A>G | *GSDMB* | -3.382 | 9.59E-04 |
| | | *ORMDL3* | -3.485 | 6.77E-04 |
| rs77232433 | g.37962846.T>C | *GSDMB* | -3.382 | 9.59E-04 |
| | | *ORMDL3* | -3.485 | 6.77E-04 |
| rs9901617 | g.37964176.C>G | *GSDMB* | -3.382 | 9.59E-04 |
| | | *ORMDL3* | -3.485 | 6.77E-04 |
| rs4337325 | g.37964436.T>C | *GSDMB* | -3.382 | 9.59E-04 |
| | | *ORMDL3* | -3.485 | 6.77E-04 |
| rs28580691 | g.37965567.C>T | *GSDMB* | -3.382 | 9.59E-04 |
| | | *ORMDL3* | -3.485 | 6.77E-04 |
| rs9892781 | g.37967438.C>T | *GSDMB* | -3.382 | 9.59E-04 |
| | | *ORMDL3* | -3.485 | 6.77E-04 |
| rs9914757 | g.37967562.G>A | *GSDMB* | -3.382 | 9.59E-04 |
| | | *ORMDL3* | -3.485 | 6.77E-04 |
| rs558811024 | g.37967634_37967650del | *GSDMB* | -3.382 | 9.59E-04 |
| | | *ORMDL3* | -3.485 | 6.77E-04 |
| rs67135646 | g.37967872.G>C | *GSDMB* | -3.382 | 9.59E-04 |

| | | | | |
|---|---|---|---|---|
| | | *ORMDL3* | -3.485 | 6.77E-04 |
| rs112238900 | g.37968495.T>C | *GSDMB* | -3.382 | 9.59E-04 |
| | | *ORMDL3* | -3.485 | 6.77E-04 |
| rs534820301 | g.37969371.G>A | *GSDMB* | -3.382 | 9.59E-04 |
| | | *ORMDL3* | -3.485 | 6.77E-04 |
| rs113115305 | g.37970687.C>A | *GSDMB* | -3.382 | 9.59E-04 |
| | | *ORMDL3* | -3.485 | 6.77E-04 |
| rs12952146 | g.37970903.T>C | *GSDMB* | -3.382 | 9.59E-04 |
| | | *ORMDL3* | -3.485 | 6.77E-04 |
| rs112412105 | g.37971636.G>A | *GSDMB* | -3.382 | 9.59E-04 |
| | | *ORMDL3* | -3.485 | 6.77E-04 |
| rs9898031 | g.37972648.G>C | *GSDMB* | -3.382 | 9.59E-04 |
| | | *ORMDL3* | -3.485 | 6.77E-04 |
| rs9902621 | g.37973011.A>G | *GSDMB* | -3.382 | 9.59E-04 |
| | | *ORMDL3* | -3.485 | 6.77E-04 |
| rs9913957 | g.37974499.G>A | *GSDMB* | -3.382 | 9.59E-04 |
| | | *ORMDL3* | -3.485 | 6.77E-04 |
| rs58075375 | g.37975593.T>C | *GSDMB* | -3.382 | 9.59E-04 |
| | | *ORMDL3* | -3.485 | 6.77E-04 |
| rs34053394 | g.37975661.G>A | *GSDMB* | -3.382 | 9.59E-04 |
| | | *ORMDL3* | -3.485 | 6.77E-04 |
| rs112743130 | g.37975856.C>G | *GSDMB* | -3.382 | 9.59E-04 |
| | | *ORMDL3* | -3.485 | 6.77E-04 |
| rs9901917 | g.37976206.C>G | *GSDMB* | -3.382 | 9.59E-04 |
| | | *ORMDL3* | -3.485 | 6.77E-04 |
| rs9904834 | g.37976501.G>A | *GSDMB* | -3.382 | 9.59E-04 |
| | | *ORMDL3* | -3.485 | 6.77E-04 |
| rs9904277 | g.37976599.T>C | *GSDMB* | -3.382 | 9.59E-04 |
| | | *ORMDL3* | -3.485 | 6.77E-04 |
| rs9911069 | g.37976602.C>T | *GSDMB* | -3.382 | 9.59E-04 |
| | | *ORMDL3* | -3.485 | 6.77E-04 |
| rs9908983 | g.37976927.A>G | *GSDMB* | -3.382 | 9.59E-04 |
| | | *ORMDL3* | -3.485 | 6.77E-04 |
| rs8076347 | g.37977541.T>G | *GSDMB* | -3.382 | 9.59E-04 |
| | | *ORMDL3* | -3.485 | 6.77E-04 |
| rs8067523 | g.37977814.C>T | *GSDMB* | -3.382 | 9.59E-04 |
| | | *ORMDL3* | -3.485 | 6.77E-04 |
| rs111907649 | g.37979125.A>G | *ORMDL3* | -3.452 | 7.58E-04 |
| rs12940616 | g.37979780.G>A | *GSDMB* | -3.382 | 9.59E-04 |
| | | *ORMDL3* | -3.485 | 6.77E-04 |
| rs28880017 | g.37980023.T>C | *GSDMB* | -3.382 | 9.59E-04 |
| | | *ORMDL3* | -3.485 | 6.77E-04 |
| rs9914665 | g.37980567.C>T | *GSDMB* | -3.382 | 9.59E-04 |
| | | *ORMDL3* | -3.485 | 6.77E-04 |
| rs34925681 | g.37981129.T>C | *GSDMB* | -3.382 | 9.59E-04 |
| | | *ORMDL3* | -3.485 | 6.77E-04 |
| rs34864615 | g.37981224.A>G | *GSDMB* | -3.382 | 9.59E-04 |
| | | *ORMDL3* | -3.485 | 6.77E-04 |
| rs34640000 | g.37981238.A>G | *GSDMB* | -3.382 | 9.59E-04 |
| | | *ORMDL3* | -3.485 | 6.77E-04 |
| rs12942660 | g.37982038.T>C | *GSDMB* | -3.382 | 9.59E-04 |

| | | | | |
|---|---|---|---|---|
| | | *ORMDL3* | -3.485 | 6.77E-04 |
| rs111469562 | g.37982697.C>T | *GSDMB* | -3.382 | 9.59E-04 |
| | | *ORMDL3* | -3.485 | 6.77E-04 |
| rs35130019 | g.37983142.G>A | *GSDMB* | -3.382 | 9.59E-04 |
| | | *ORMDL3* | -3.485 | 6.77E-04 |
| rs112437508 | g.37983513.A>G | *GSDMB* | -3.382 | 9.59E-04 |
| | | *ORMDL3* | -3.485 | 6.77E-04 |
| rs112797570 | g.37983752.A>G | *GSDMB* | -3.382 | 9.59E-04 |
| | | *ORMDL3* | -3.485 | 6.77E-04 |
| rs66763967 | g.37984078.C>T | *GSDMB* | -3.382 | 9.59E-04 |
| | | *ORMDL3* | -3.485 | 6.77E-04 |
| rs71369791 | g.37984086.G>A | *GSDMB* | -3.382 | 9.59E-04 |
| | | *ORMDL3* | -3.485 | 6.77E-04 |
| rs12946188 | g.37984430.A>C | *GSDMB* | -3.382 | 9.59E-04 |
| | | *ORMDL3* | -3.485 | 6.77E-04 |
| rs12946792 | g.37985322.C>T | *GSDMB* | -3.382 | 9.59E-04 |
| | | *ORMDL3* | -3.485 | 6.77E-04 |
| rs9913769 | g.37986337.G>A | *GSDMB* | -3.382 | 9.59E-04 |
| | | *ORMDL3* | -3.485 | 6.77E-04 |
| rs71369792 | g.37986873.A>G | *GSDMB* | -3.382 | 9.59E-04 |
| | | *ORMDL3* | -3.485 | 6.77E-04 |
| rs113479772 | g.37987043.A>G | *GSDMB* | -3.382 | 9.59E-04 |
| | | *ORMDL3* | -3.485 | 6.77E-04 |
| rs111734595 | g.37987400.T>C | *GSDMB* | -3.382 | 9.59E-04 |
| | | *ORMDL3* | -3.485 | 6.77E-04 |
| rs112141468 | g.37987465.T>C | *GSDMB* | -3.382 | 9.59E-04 |
| | | *ORMDL3* | -3.485 | 6.77E-04 |
| rs73304123 | g.37987589.T>C | *GSDMB* | -3.382 | 9.59E-04 |
| | | *ORMDL3* | -3.485 | 6.77E-04 |
| rs8069893 | g.37988293.T>C | *GSDMB* | -3.382 | 9.59E-04 |
| | | *ORMDL3* | -3.485 | 6.77E-04 |
| rs111944912 | g.37988477.C>T | *GSDMB* | -3.382 | 9.59E-04 |
| | | *ORMDL3* | -3.485 | 6.77E-04 |
| rs34521860 | g.37988775_37988775insAT | *GSDMB* | -3.382 | 9.59E-04 |
| | | *ORMDL3* | -3.485 | 6.77E-04 |
| rs12945253 | g.37991328.G>C | *GSDMB* | -3.382 | 9.59E-04 |
| | | *ORMDL3* | -3.485 | 6.77E-04 |
| rs28449671 | g.37991631.C>T | *GSDMB* | -3.382 | 9.59E-04 |
| | | *ORMDL3* | -3.485 | 6.77E-04 |
| rs34518282 | g.37992001.C>G | *GSDMB* | -3.382 | 9.59E-04 |
| | | *ORMDL3* | -3.485 | 6.77E-04 |
| rs28392589 | g.37992885.G>A | *GSDMB* | -3.382 | 9.59E-04 |
| | | *ORMDL3* | -3.485 | 6.77E-04 |
| rs111691913 | g.37993239.T>C | *GSDMB* | -3.382 | 9.59E-04 |
| | | *ORMDL3* | -3.485 | 6.77E-04 |
| rs12938617 | g.37993352.A>T | *GSDMB* | -3.382 | 9.59E-04 |
| | | *ORMDL3* | -3.485 | 6.77E-04 |
| rs9898464 | g.37993992.C>T | *GSDMB* | -3.382 | 9.59E-04 |
| | | *ORMDL3* | -3.485 | 6.77E-04 |
| rs9900019 | g.37994625.C>T | *GSDMB* | -3.382 | 9.59E-04 |
| | | *ORMDL3* | -3.485 | 6.77E-04 |

| | | | | |
|---|---|---|---|---|
| rs34485135 | g.37994722.C>T | *GSDMB* | -3.382 | 9.59E-04 |
| | | *ORMDL3* | -3.485 | 6.77E-04 |
| | | *GSDMB* | -3.382 | 9.59E-04 |
| | | *ORMDL3* | -3.485 | 6.77E-04 |
| rs9900541 | g.37996071.C>T | *GSDMB* | -3.382 | 9.59E-04 |
| | | *ORMDL3* | -3.485 | 6.77E-04 |
| rs35405174 | g.37998705.G>A | *GSDMB* | -3.382 | 9.59E-04 |
| | | *ORMDL3* | -3.485 | 6.77E-04 |
| rs28403951 | g.37999534.G>C | *GSDMB* | -3.382 | 9.59E-04 |
| | | *ORMDL3* | -3.485 | 6.77E-04 |
| rs12946040 | g.38000018.A>C | *GSDMB* | -3.382 | 9.59E-04 |
| | | *ORMDL3* | -3.485 | 6.77E-04 |
| rs36041042 | g.38000057_38000057insT | *ORMDL3* | -3.477 | 6.96E-04 |
| rs67600807 | g.38001559.G>A | *GSDMB* | -3.382 | 9.59E-04 |
| | | *ORMDL3* | -3.485 | 6.77E-04 |
| rs34039260 | g.38001659_38001659insC | *GSDMB* | -3.382 | 9.59E-04 |
| | | *ORMDL3* | -3.485 | 6.77E-04 |
| rs112677036 | g.38002153.A>G | *GSDMB* | -3.382 | 9.59E-04 |
| | | *ORMDL3* | -3.485 | 6.77E-04 |
| rs573641447 | g.38002747.G>T | *ORMDL3* | -3.654 | 3.78E-04 |
| rs28535471 | g.38002916.C>G | *GSDMB* | -3.382 | 9.59E-04 |
| | | *ORMDL3* | -3.485 | 6.77E-04 |
| rs28648398 | g.38003025.C>G | *GSDMB* | -3.382 | 9.59E-04 |
| | | *ORMDL3* | -3.485 | 6.77E-04 |
| rs28695941 | g.38003330.T>A | *GSDMB* | -3.382 | 9.59E-04 |
| | | *ORMDL3* | -3.485 | 6.77E-04 |
| rs34654193 | g.38003859.C>G | *GSDMB* | -3.382 | 9.59E-04 |
| | | *ORMDL3* | -3.485 | 6.77E-04 |
| rs555695098 | g.38004454_38004454ins | *GSDMB* | -3.382 | 9.59E-04 |
| | | *ORMDL3* | -3.485 | 6.77E-04 |
| rs28537796 | g.38004906.T>C | *GSDMB* | -3.382 | 9.59E-04 |
| | | *ORMDL3* | -3.485 | 6.77E-04 |
| rs28896477 | g.38005534.T>C | *GSDMB* | -3.382 | 9.59E-04 |
| | | *ORMDL3* | -3.485 | 6.77E-04 |
| rs28842324 | g.38005742.T>C | *GSDMB* | -3.382 | 9.59E-04 |
| | | *ORMDL3* | -3.485 | 6.77E-04 |
| rs528792070 | g.38006700_38006700insTAC | *GSDMB* | -3.382 | 9.59E-04 |
| | | *ORMDL3* | -3.485 | 6.77E-04 |
| rs553196378 | g.38006967_38006969del | *GSDMB* | -3.382 | 9.59E-04 |
| | | *ORMDL3* | -3.485 | 6.77E-04 |
| rs114400736 | g.38007319.G>T | *GSDMB* | -3.382 | 9.59E-04 |
| | | *ORMDL3* | -3.485 | 6.77E-04 |
| rs12938117 | g.38008165.T>C | *GSDMB* | -3.382 | 9.59E-04 |
| | | *ORMDL3* | -3.485 | 6.77E-04 |
| rs113233720 | g.38008191.T>C | *GSDMB* | -3.382 | 9.59E-04 |
| | | *ORMDL3* | -3.485 | 6.77E-04 |
| rs8068894 | g.38009000.G>A | *GSDMB* | -3.382 | 9.59E-04 |
| | | *ORMDL3* | -3.485 | 6.77E-04 |
| rs202130396 | g.38009223_38009223insA | *GSDMB* | -3.382 | 9.59E-04 |
| | | *ORMDL3* | -3.485 | 6.77E-04 |
| rs8069531 | g.38009344.T>A | *GSDMB* | -3.382 | 9.59E-04 |

| | | | | |
|---|---|---|---|---|
| | | *ORMDL3* | -3.485 | 6.77E-04 |
| rs12947480 | g.38009676.A>G | *GSDMB* | -3.382 | 9.59E-04 |
| | | *ORMDL3* | -3.485 | 6.77E-04 |
| rs9907291 | g.38010037.G>A | *GSDMB* | -3.382 | 9.59E-04 |
| | | *ORMDL3* | -3.485 | 6.77E-04 |
| rs9907339 | g.38010103.C>A | *GSDMB* | -3.382 | 0.001 |
| | | *ORMDL3* | -3.485 | 0.001 |
| rs9907498 | g.38010104.G>A | *GSDMB* | -3.382 | 9.59E-04 |
| | | *ORMDL3* | -3.485 | 6.77E-04 |
| rs8077365 | g.38010758.T>G | *ORMDL3* | -3.364 | 1.02E-03 |
| rs8078409 | g.38010759.T>C | *ORMDL3* | -3.364 | 1.02E-03 |
| rs8079075 | g.38010815.G>A | *GSDMB* | -3.382 | 9.59E-04 |
| | | *ORMDL3* | -3.485 | 6.77E-04 |
| rs28621669 | g.38011266.C>T | *GSDMB* | -3.382 | 9.59E-04 |
| | | *ORMDL3* | -3.485 | 6.77E-04 |
| rs28723017 | g.38011451.T>C | *GSDMB* | -3.382 | 9.59E-04 |
| | | *ORMDL3* | -3.485 | 6.77E-04 |
| rs9900538 | g.38011709.C>T | *GSDMB* | -3.382 | 9.59E-04 |
| | | *ORMDL3* | -3.485 | 6.77E-04 |
| rs113466546 | g.38012587.A>G | *GSDMB* | -3.382 | 9.59E-04 |
| | | *ORMDL3* | -3.485 | 6.77E-04 |
| rs35770042 | g.38012991.G>A | *GSDMB* | -3.382 | 9.59E-04 |
| | | *ORMDL3* | -3.485 | 6.77E-04 |
| rs28661421 | g.38013807.G>A | *GSDMB* | -3.382 | 9.59E-04 |
| | | *ORMDL3* | -3.485 | 6.77E-04 |
| rs16965367 | g.38014316.C>T | *GSDMB* | -3.382 | 9.59E-04 |
| | | *ORMDL3* | -3.485 | 6.77E-04 |
| rs9912095 | g.38014533.G>A | *GSDMB* | -3.382 | 9.59E-04 |
| | | *ORMDL3* | -3.485 | 6.77E-04 |
| rs9915509 | g.38014705.A>G | *GSDMB* | -3.382 | 9.59E-04 |
| | | *ORMDL3* | -3.485 | 6.77E-04 |
| rs201152777 | g.38014758_38014758insT | *GSDMB* | -3.382 | 9.59E-04 |
| | | *ORMDL3* | -3.485 | 6.77E-04 |
| rs9915797 | g.38014868.A>G | *GSDMB* | -3.382 | 9.59E-04 |
| | | *ORMDL3* | -3.485 | 6.77E-04 |
| rs9916259 | g.38015060.T>G | *GSDMB* | -3.382 | 9.59E-04 |
| | | *ORMDL3* | -3.485 | 6.77E-04 |
| rs142025263 | g.38015188_38015188insACT | *GSDMB* | -3.382 | 9.59E-04 |
| | | *ORMDL3* | -3.485 | 6.77E-04 |
| rs8068125 | g.38015517.C>T | *GSDMB* | -3.382 | 9.59E-04 |
| | | *ORMDL3* | -3.485 | 6.77E-04 |
| rs77924338 | g.38016357.T>C | *GSDMB* | -3.382 | 9.59E-04 |
| | | *ORMDL3* | -3.485 | 6.77E-04 |
| rs9899006 | g.38017065.A>T | *GSDMB* | -3.382 | 9.59E-04 |
| | | *ORMDL3* | -3.485 | 6.77E-04 |
| rs67837020 | g.38017313_38017320del | *GSDMB* | -3.382 | 9.59E-04 |
| | | *ORMDL3* | -3.485 | 6.77E-04 |
| rs9899336 | g.38017780.T>C | *GSDMB* | -3.382 | 0.001 |
| | | *ORMDL3* | -3.485 | 6.77E-04 |
| rs9907096 | g.38018774.T>C | *GSDMB* | -3.382 | 9.59E-04 |
| | | *ORMDL3* | -3.485 | 6.77E-04 |

| | | | | |
|---|---|---|---|---|
| rs9907966 | g.38018803.C>A | *ORMDL3* | -3.364 | 1.02E-03 |
| rs9905881 | g.38018955.A>G | *GSDMB* | -3.382 | 9.59E-04 |
| | | *ORMDL3* | -3.485 | 6.77E-04 |
| rs9907564 | g.38018967.T>C | *GSDMB* | -3.382 | 9.59E-04 |
| | | *ORMDL3* | -3.485 | 6.77E-04 |
| rs9907794 | g.38019026.G>C | *GSDMB* | -3.382 | 9.59E-04 |
| | | *ORMDL3* | -3.485 | 6.77E-04 |
| rs111678394 | g.38021117.C>G | *GSDMB* | -3.382 | 9.59E-04 |
| | | *ORMDL3* | -3.485 | 6.77E-04 |
| rs11284476 | g.38022534_38022535del | *GSDMB* | -3.382 | 9.59E-04 |
| | | *ORMDL3* | -3.485 | 6.77E-04 |
| rs4622539 | g.38023031.A>T | *GSDMB* | -3.382 | 9.59E-04 |
| | | *ORMDL3* | -3.485 | 6.77E-04 |
| rs112880843 | g.38024298.A>G | *GSDMB* | -3.382 | 9.59E-04 |
| | | *ORMDL3* | -3.485 | 6.77E-04 |
| rs150803188 | g.38032505.A>G | *GSDMB* | -3.382 | 0.001 |
| | | *ORMDL3* | -3.485 | 6.77E-04 |
| rs1475291182 | g.38063809_38063811del | *GSDMB* | -3.382 | 9.59E-04 |
| | | *ORMDL3* | -3.485 | 6.77E-04 |
| rs3169572 | g.38077412.T>C | *ORMDL3* | -3.364 | 1.02E-03 |
| rs373861118 | g.38118929_38118930insCACACACACACA | *ORMDL3* | -4.069 | 8.27E-05 |
| rs111904390 | g.37888516.T>G | *GSDMB* | -3.382 | 9.59E-04 |
| | | *ORMDL3* | -3.485 | 6.77E-04 |
| rs113120593 | g.37840318.T>A | *MIEN1* | -4.005 | 1.05E-04 |
| rs113231716 | g.37869990.G>A | *MIEN1* | -4.005 | 1.05E-04 |
| rs143123127 | g.38007190.A>G | *GSDMB* | -3.382 | 9.59E-04 |
| | | *ORMDL3* | -3.485 | 6.77E-04 |
| rs1453560 | g.38023441.C>A | *GSDMB* | -3.382 | 9.59E-04 |
| | | *ORMDL3* | -3.485 | 6.77E-04 |
| rs151309334 | g.37831293_37831298del | *MIEN1* | -4.005 | 1.05E-04 |
| rs202041850 | g.38007197_38007201del | *GSDMB* | -3.382 | 9.59E-04 |
| | | *ORMDL3* | -3.485 | 6.77E-04 |
| rs4252608 | g.37862966.T>C | *MIEN1* | -4.005 | 1.05E-04 |
| rs55757004 | g.38126395.C>G | *GSDMB* | 3.710 | 3.09E-04 |
| | | *ORMDL3* | 3.900 | 1.55E-04 |
| rs58566623 | g.37938665_37938684del | *GSDMB* | -3.373 | 9.89E-04 |
| | | *ORMDL3* | -3.461 | 7.35E-04 |
| rs7211998 | g.37959789.G>A | *GSDMB* | -3.382 | 9.59E-04 |
| | | *ORMDL3* | -3.485 | 6.77E-04 |
| rs77925972 | g.37930210.C>T | *GSDMB* | -3.382 | 9.59E-04 |
| | | *ORMDL3* | -3.485 | 6.77E-04 |
| rs9899345 | g.37954758.A>G | *GSDMB* | -3.382 | 0.001 |
| | | *ORMDL3* | -3.485 | 6.77E-04 |
| rs9908694 | g.37997772.T>C | *GSDMB* | -3.382 | 9.59E-04 |
| | | *ORMDL3* | -3.485 | 6.77E-04 |
| rs9911688 | g.37943801.T>C | *GSDMB* | -3.382 | 9.59E-04 |
| | | *ORMDL3* | -3.485 | 6.77E-04 |

[a] P values for the association between the SNPs included in the eQTL and asthma. Ps are only shown if significant (p <0.001).
