## Supplemental Table 13 for "*GSDMA* drives the most replicated association with asthma in naïve CD4^+^ T cells"

**Table S13:** Significant eQTLs with *GSDMA* expression in naïve CD4$^+$ T cells for individuals with asthma and controls only

| Haplotype block | SNP id | HGVS names (hg19) | Gene | Statistic | P eQTL | P eQTL, permutations | P asthma[a] |
|---|---|---|---|---|---|---|---|
| Controls only | | | | | | | |
| Fifth block | rs870829 | g.38068382.T>G | GSDMA | -4.8744595 | 5.43E-06 | 0.019 | 7.23E-04 |
| | rs1011082 | g.38068514.A>G | GSDMA | -4.2004886 | 6.87E-05 | 0.022 | 7.66E-04 |
| | rs921650 | g.38069076.C>T | GSDMA | -4.3718177 | 3.67E-05 | 0.020 | 5.34E-04 |
| | rs921649 | g.38069274.G>A | GSDMA | -4.3718177 | 3.67E-05 | 0.020 | 5.34E-04 |
| | rs5820308 | g.38069364_38069369del | GSDMA | -5.3361787 | 8.63E-07 | 0.012 | 8.13E-04 |
| | rs6503524 | g.38069809.C>T | GSDMA | -4.3718177 | 3.67E-05 | 0.020 | 5.34E-04 |
| | rs7216389 | g.38069949.C>T | GSDMA | -4.3718177 | 3.67E-05 | 0.020 | 5.34E-04 |
| | rs7216558 | g.38070071.C>T | GSDMA | -4.3718177 | 3.67E-05 | 0.020 | 5.34E-04 |
| | rs7221605 | g.38070789.C>T | GSDMA | -4.3077336 | 4.65E-05 | 0.023 | |
| | rs150597688 | g.38071086_38071086insTT | GSDMA | -4.3077336 | 4.65E-05 | 0.023 | |
| | rs143385463 | g.38071855_38071858del | GSDMA | -4.3077336 | 4.65E-05 | 0.023 | |
| | rs1031458 | g.38072173.G>T | GSDMA | -4.3077336 | 4.65E-05 | 0.023 | |
| | rs1031459 | g.38072245.C>G | GSDMA | -5.9519307 | 6.70E-08 | 0.012 | |
| | rs1031460 | g.38072247.T>G | GSDMA | -4.3077336 | 4.65E-05 | 0.023 | |
| | rs8065777 | g.38072402.C>T | GSDMA | -4.3077336 | 4.65E-05 | 0.023 | |
| | rs2872516 | g.38072727.C>T | GSDMA | -5.2419919 | 1.26E-06 | 0.017 | |
| | rs9303279 | g.38073968.G>C | GSDMA | -5.3361787 | 8.63E-07 | 0.012 | 6.13E-04 |
| | rs9303280 | g.38074031.T>C | GSDMA | -4.3077336 | 4.65E-05 | 0.023 | 8.49E-04 |
| | rs9303281 | g.38074046.G>A | GSDMA | -4.3718177 | 3.67E-05 | 0.020 | 2.60E-04 |
| | rs7219923 | g.38074518.C>T | GSDMA | -4.3718177 | 3.67E-05 | 0.020 | 2.75E-04 |
| | rs7224129 | g.38075426.G>A | GSDMA | -4.3718177 | 3.67E-05 | 0.020 | 3.47E-04 |
| | rs8074437 | g.38076137.T>G | GSDMA | -4.3718177 | 3.67E-05 | 0.020 | 3.47E-04 |
| | rs71971950 | g.38076198_38076201del | GSDMA | -4.6997018 | 1.07E-05 | 0.016 | 3.77E-04 |
| | rs4065275 | g.38080865.A>G | GSDMA | -4.5594211 | 1.82E-05 | 0.025 | 8.19E-04 |
| | rs8076131 | g.38080912.G>A | GSDMA | -5.587804 | 3.08E-07 | 0.008 | 5.13E-04 |
| | rs12603332 | g.38082807.T>C | GSDMA | -4.516065 | 2.15E-05 | 0.025 | 4.23E-04 |
| | rs35557848 | g.38087439.T>C | GSDMA | -3.6997813 | 3.94E-04 | 0.030 | |

| | | | | | | | |
|---|---|---|---|---|---|---|---|
| Individuals with asthma only | | | | | | | |
| Fifth block | rs870829 | g.38068382.T>G | *GSDMA* | -4.874 | 5.43E-06 | 0.032 | 7.23E-04 |
| | rs921650 | g.38069076.C>T | *GSDMA* | -4.372 | 3.67E-05 | 0.044 | 5.34E-04 |
| | rs921649 | g.38069274.G>A | *GSDMA* | -4.372 | 3.67E-05 | 0.044 | 5.34E-04 |
| | rs5820308 | g.38069364_38069369del | *GSDMA* | -5.336 | 8.63E-07 | 0.035 | 8.13E-04 |
| | rs6503524 | g.38069809.C>T | *GSDMA* | -4.372 | 3.67E-05 | 0.044 | 5.34E-04 |
| | rs7216389 | g.38069949.C>T | *GSDMA* | -4.372 | 3.67E-05 | 0.044 | 5.34E-04 |
| | rs7216558 | g.38070071.C>T | *GSDMA* | -4.372 | 3.67E-05 | 0.044 | 5.34E-04 |
| | rs1031459 | g.38072245.C>G | *GSDMA* | -5.952 | 6.70E-08 | 0.047 | |
| | rs2872516 | g.38072727.C>T | *GSDMA* | -5.242 | 1.26E-06 | 0.053 | |
| | rs9303279 | g.38073968.G>C | *GSDMA* | -5.336 | 8.63E-07 | 0.035 | 6.13E-04 |
| | rs9303281 | g.38074046.G>A | *GSDMA* | -4.372 | 3.67E-05 | 0.044 | 2.60E-04 |
| | rs7219923 | g.38074518.C>T | *GSDMA* | -4.372 | 3.67E-05 | 0.044 | 2.75E-04 |
| | rs7224129 | g.38075426.G>A | *GSDMA* | -4.372 | 3.67E-05 | 0.039 | 3.47E-04 |
| | rs8074437 | g.38076137.T>G | *GSDMA* | -4.372 | 3.67E-05 | 0.039 | 3.47E-04 |
| | rs71971950 | g.38076198_38076201del | *GSDMA* | -4.700 | 1.07E-05 | 0.025 | 3.77E-04 |
| | rs4065275 | g.38080865.A>G | *GSDMA* | -4.559 | 1.82E-05 | 0.055 | 8.19E-04 |
| | rs8076131 | g.38080912.G>A | *GSDMA* | -5.588 | 3.08E-07 | 0.041 | 5.13E-04 |
| | rs12603332 | g.38082807.T>C | *GSDMA* | -4.516 | 2.15E-05 | 0.056 | 4.23E-04 |

[a] P values for the association between the SNPs from the eQTL and asthma. Ps are only shown if significant (p <0.001 and p for permutations <0.05).
